## Supplementary File 1 for "A spatial and temporal atlas of tubulin isotype gene expression during vertebrate embryonic development"

### Single Cell RNA-Seq Analysis: Tubulins in Neural Crest EMT

Rogers Lab, UC Davis School of Veterinary Medicine

### 1. Set-Up and Configuration

```
# Threshold configurations as per original manuscripts
get_qc_params <- function(sample_name) {
  # Williams et al. 2022 Parameters
  if (grepl("HH[45]", sample_name) && grepl("Williams", sample_name)) {
    return(list(min_feat=500, max_feat=3500, max_mt=0.5))
  } else if (grepl("HH[67]", sample_name) && grepl("Williams", sample_name)) {
    return(list(min_feat=500, max_feat=5000, max_mt=0.5))
  }
  # Pajanoja 2023/2025 Parameters
  else {
    return(list(min_feat=200, max_feat=4000, max_mt=5.0))
  }
}

# Global list to track cell counts through filtering steps
qc_tracker <- list()

# Quality Control Plotter
qc_tester <- function(obj) {
  p <- VlnPlot(obj, features = c("nFeature_RNA", "nCount_RNA", "percent.mt"),
               group.by = "seurat_clusters", pt.size = 0, ncol = 1)
  print(p)
}
```

### 2. Processing Pajanoja et al Datasets (GSE221190)

```
pajanoja_samples <- c("HH5_1", "HH5_2", "HH7_1", "HH7_2", "HH8_1", "HH8_2", "HH9_1", "HH9_2",
"HH11")
pajanoja_list <- list()

for (sample in pajanoja_samples) {

  # Ensure correct sample_id and reference for HH11 (2025 vs 2023)
  if (sample == "HH11") {
    sample_full_name <- "Pajanoja2025_HH11"
    ref_year <- "Pajanoja_2025"
  } else {
    sample_full_name <- paste0("Pajanoja2023_", sample)
```

```
    ref_year <- "Pajanoja_2023"
  }

  params <- get_qc_params(sample_full_name)

  # Load Data
  print(paste0("Loading ",sample))
  raw_mat <- Read10X(paste0("E:\\Rogers Lab\\Pajanoja 2023\\INPUT raw\\", sample))
  filt_mat <- Read10X(paste0("E:\\Rogers Lab\\Pajanoja 2023\\INPUT\\", sample))

  # SoupX: Auto-estimate contamination
  print(paste0("Running SoupX for ",sample))
  sc = SoupChannel(toc=filt_mat, tod=raw_mat, calcSoupProfile = FALSE,  keepDroplets = TRUE )
  sc = estimateSoup(sc)

  tmp_seu <- CreateSeuratObject(filt_mat) %>%
    NormalizeData(verbose=T) %>%
    FindVariableFeatures(verbose=T) %>%
    ScaleData(verbose=T) %>%
    RunPCA(verbose=T) %>%
    FindNeighbors(dims=1:20, verbose=T) %>%
    FindClusters(resolution=0.5, verbose=T)
  sc = setClusters(sc, tmp_seu$seurat_clusters)

  adj_matrix <- tryCatch({
    print(plotMarkerDistribution(sc)) # Attempt to plot
    sc <- autoEstCont(sc)             # Attempt to estimate
    adjustCounts(sc)                  # Output adjusted matrix
  }, error = function(e) {
    # fallback to filt_mat if the object does need
    message(paste0("\n[!] SoupX failed for ", sample_full_name, " with error: ", e$message))
    message("[!] Proceeding with standard unadjusted filtered counts instead.\n")
    return(filt_mat)
  })

  # Create Clean Seurat Object & Record Pre-QC Count
  seu <- CreateSeuratObject(counts = adj_matrix, project = sample_full_name, min.cells = 3, mi
n.features = params$min_feat)
  seu[["percent.mt"]] <- PercentageFeatureSet(seu, pattern = "^MT-")
  count_pre_qc <- ncol(seu)

  # Apply Strict QC & Record Post-QC Count
  print(paste0("Running QC for ",sample))
  seu <- subset(seu, subset = nFeature_RNA <= params$max_feat & percent.mt < params$max_mt)
  count_post_qc <- ncol(seu)
  rm(raw_mat, filt_mat, adj_matrix)
  gc()

  # DoubletFinder
  print(paste0("Running DoubletFinder for ",sample))
  seu <- NormalizeData(seu, verbose=F) %>%
    FindVariableFeatures(verbose=F) %>%
    ScaleData(verbose=F) %>%
    RunPCA(verbose=F) %>%
```

```
    RunUMAP(dims=1:20, verbose=F)
  sweep.res <- paramSweep(seu, PCs = 1:20, sct = FALSE)
  sweep.stats <- summarizeSweep(sweep.res, GT = FALSE)
  bcmvn <- find.pK(sweep.stats)
  pK_val <- as.numeric(as.character(bcmvn$pK[which.max(bcmvn$BCmetric)]))
  nExp_poi <- round(ncol(seu) * 8e-6 * ncol(seu)) # Dynamic 10X rate

  seu <- doubletFinder(seu, PCs = 1:20, pN = 0.25, pK = pK_val, nExp = nExp_poi, reuse.pANN = 
NULL, sct = FALSE)
  df_col <- grep("DF.classifications", colnames(seu@meta.data), value = TRUE)
  seu$Doublet_Classification <- seu@meta.data[[df_col]]

  # Generate Pre-Removal Doublet UMAP
  p1 <- DimPlot(seu, group.by = "Doublet_Classification")+ggtitle(label = paste0(sample_full_n
ame, ": Pre-Removal"))

  # Subset Singlets & Add Metadata
  seu <- subset(seu, subset = Doublet_Classification == "Singlet")
  seu$sample_id <- sample_full_name
  seu$stage <- strsplit(sample, "_")[[1]][1]
  seu$ref <- ref_year
  count_post_df <- ncol(seu)

  # Generate Post-Removal Singlet UMAP and plot side-by-side
  p2 <- DimPlot(seu, group.by = "Doublet_Classification")+ggtitle(label = "Post-Removal (Singl
ets)")
  print(p1 | p2)

  # Append to Tracker
  qc_tracker[[sample_full_name]] <- data.frame(Sample = sample_full_name, Pre_QC = count_pre_q
c, Post_QC = count_post_qc, Post_DoubletFinder = count_post_df)

  pajanoja_list[[sample]] <- seu

  # save copy of corrected object
  saveRDS(seu,paste0("E:\\Rogers Lab\\Rogers NC EMT\\Pajanoja_", sample,"_v1.rds"))
}
```

```
## [1] "Loading HH5_1"
## [1] "Running SoupX for HH5_1"
```

```
## Modularity Optimizer version 1.3.0 by Ludo Waltman and Nees Jan van Eck
##
## Number of nodes: 12944
## Number of edges: 426989
##
## Running Louvain algorithm...
## Maximum modularity in 10 random starts: 0.8908
## Number of communities: 13
## Elapsed time: 1 seconds
```

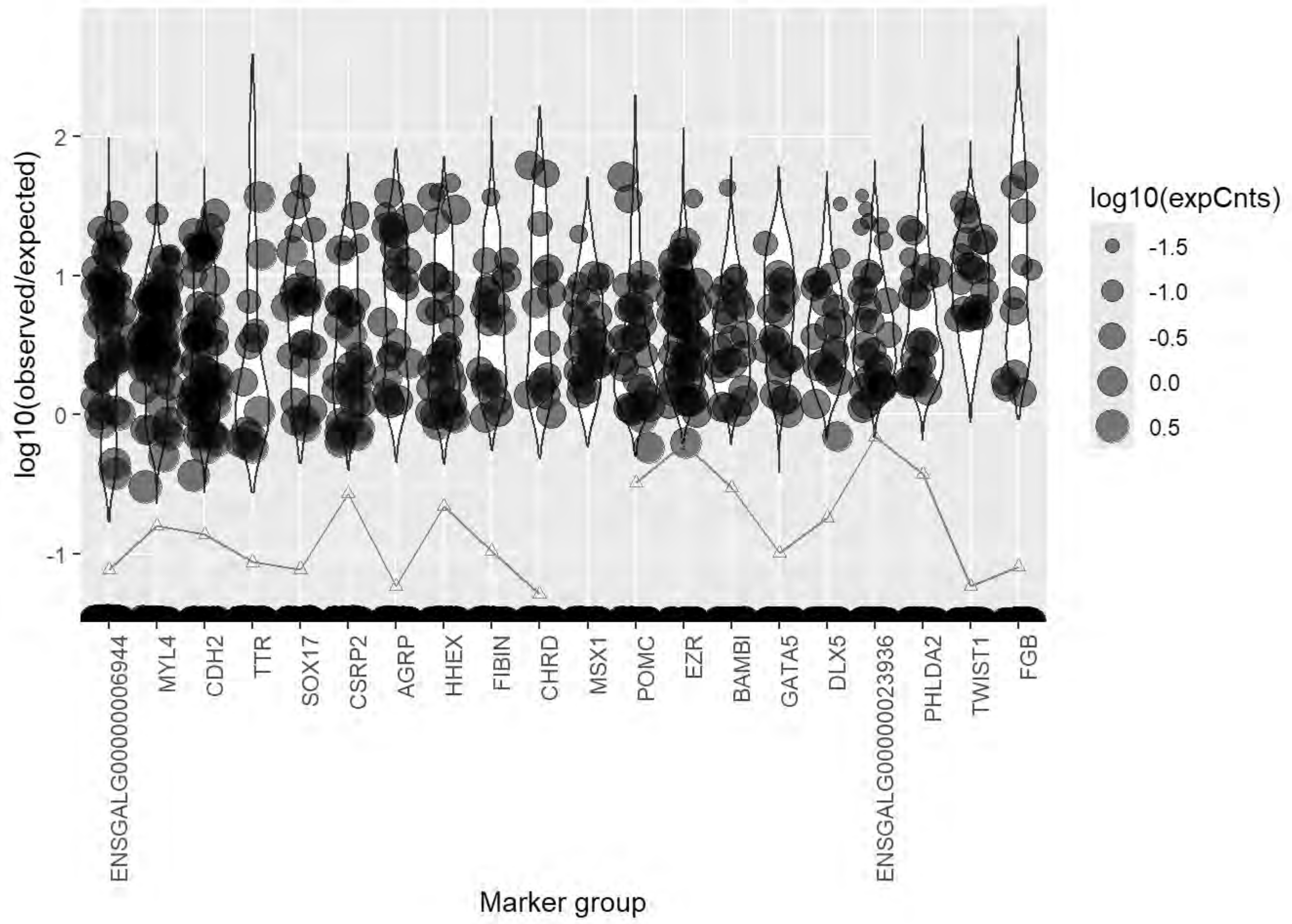
log10(observed/expected)
2
1
0
-1
log10(expCnts)
-1.5
-1.0
-0.5
0.0
0.5
ENSGALG00000006944
MYL4
CDH2
TTR
SOX17
CSRP2
AGRP
HHEX
FIBIN
CHRD
MSX1
POMC
EZR
BAMBI
GATA5
DLX5
ENSGALG00000023936
PHLDA2
TWIST1
FGB
Marker group

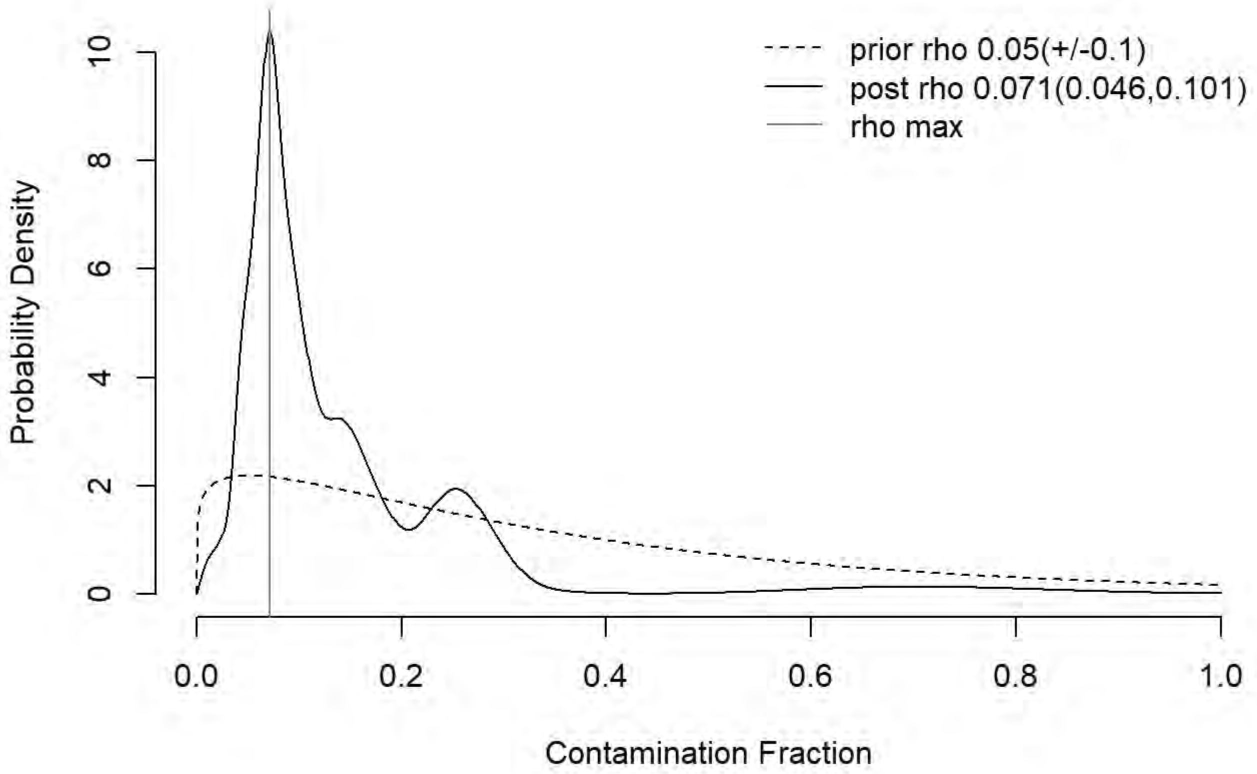
prior rho 0.05(+/-0.1)
post rho 0.071(0.046,0.101)
rho max
Probability Density
0
2
4
6
8
10
0.0
0.2
0.4
0.6
0.8
1.0
Contamination Fraction

```
## [1] "Running QC for HH5_1"
## [1] "Running DoubletFinder for HH5_1"
```

```
## [1] "Creating artificial doublets for pN = 5%"
## [1] "Creating Seurat object..."
## [1] "Normalizing Seurat object..."
```

```
## [1] "Finding variable genes..."
```

```
## [1] "Scaling data..."
```

```
## [1] "Running PCA..."
## [1] "Calculating PC distance matrix..."
## [1] "Defining neighborhoods..."
## [1] "Computing pANN across all pK..."
## [1] "pK = 0.001..."
## [1] "pK = 0.005..."
## [1] "pK = 0.01..."
## [1] "pK = 0.02..."
## [1] "pK = 0.03..."
## [1] "pK = 0.04..."
## [1] "pK = 0.05..."
## [1] "pK = 0.06..."
## [1] "pK = 0.07..."
## [1] "pK = 0.08..."
## [1] "pK = 0.09..."
## [1] "pK = 0.1..."
## [1] "pK = 0.11..."
## [1] "pK = 0.12..."
## [1] "pK = 0.13..."
## [1] "pK = 0.14..."
## [1] "pK = 0.15..."
## [1] "pK = 0.16..."
## [1] "pK = 0.17..."
## [1] "pK = 0.18..."
## [1] "pK = 0.19..."
## [1] "pK = 0.2..."
## [1] "pK = 0.21..."
## [1] "pK = 0.22..."
## [1] "pK = 0.23..."
## [1] "pK = 0.24..."
## [1] "pK = 0.25..."
## [1] "pK = 0.26..."
## [1] "pK = 0.27..."
## [1] "pK = 0.28..."
## [1] "pK = 0.29..."
## [1] "pK = 0.3..."
## [1] "Creating artificial doublets for pN = 10%"
## [1] "Creating Seurat object..."
## [1] "Normalizing Seurat object..."
```

```
## [1] "Finding variable genes..."
```

```
## [1] "Scaling data..."
```

```
## [1] "Running PCA..."
## [1] "Calculating PC distance matrix..."
## [1] "Defining neighborhoods..."
## [1] "Computing pANN across all pK..."
## [1] "pK = 0.001..."
## [1] "pK = 0.005..."
## [1] "pK = 0.01..."
## [1] "pK = 0.02..."
## [1] "pK = 0.03..."
## [1] "pK = 0.04..."
## [1] "pK = 0.05..."
## [1] "pK = 0.06..."
## [1] "pK = 0.07..."
## [1] "pK = 0.08..."
## [1] "pK = 0.09..."
## [1] "pK = 0.1..."
## [1] "pK = 0.11..."
## [1] "pK = 0.12..."
## [1] "pK = 0.13..."
## [1] "pK = 0.14..."
## [1] "pK = 0.15..."
## [1] "pK = 0.16..."
## [1] "pK = 0.17..."
## [1] "pK = 0.18..."
## [1] "pK = 0.19..."
## [1] "pK = 0.2..."
## [1] "pK = 0.21..."
## [1] "pK = 0.22..."
## [1] "pK = 0.23..."
## [1] "pK = 0.24..."
## [1] "pK = 0.25..."
## [1] "pK = 0.26..."
## [1] "pK = 0.27..."
## [1] "pK = 0.28..."
## [1] "pK = 0.29..."
## [1] "pK = 0.3..."
## [1] "Creating artificial doublets for pN = 15%"
## [1] "Creating Seurat object..."
## [1] "Normalizing Seurat object..."
```

```
## [1] "Finding variable genes..."
```

```
## [1] "Scaling data..."
```

```
## [1] "Running PCA..."
## [1] "Calculating PC distance matrix..."
```

```
## [1] "Defining neighborhoods..."
## [1] "Computing pANN across all pK..."
## [1] "pK = 0.001..."
## [1] "pK = 0.005..."
## [1] "pK = 0.01..."
## [1] "pK = 0.02..."
## [1] "pK = 0.03..."
## [1] "pK = 0.04..."
## [1] "pK = 0.05..."
## [1] "pK = 0.06..."
## [1] "pK = 0.07..."
## [1] "pK = 0.08..."
## [1] "pK = 0.09..."
## [1] "pK = 0.1..."
## [1] "pK = 0.11..."
## [1] "pK = 0.12..."
## [1] "pK = 0.13..."
## [1] "pK = 0.14..."
## [1] "pK = 0.15..."
## [1] "pK = 0.16..."
## [1] "pK = 0.17..."
## [1] "pK = 0.18..."
## [1] "pK = 0.19..."
## [1] "pK = 0.2..."
## [1] "pK = 0.21..."
## [1] "pK = 0.22..."
## [1] "pK = 0.23..."
## [1] "pK = 0.24..."
## [1] "pK = 0.25..."
## [1] "pK = 0.26..."
## [1] "pK = 0.27..."
## [1] "pK = 0.28..."
## [1] "pK = 0.29..."
## [1] "pK = 0.3..."
## [1] "Creating artificial doublets for pN = 20%"
## [1] "Creating Seurat object..."
## [1] "Normalizing Seurat object..."
```

```
## [1] "Finding variable genes..."
```

```
## [1] "Scaling data..."
```

```
## [1] "Running PCA..."
## [1] "Calculating PC distance matrix..."
## [1] "Defining neighborhoods..."
## [1] "Computing pANN across all pK..."
## [1] "pK = 0.001..."
## [1] "pK = 0.005..."
## [1] "pK = 0.01..."
## [1] "pK = 0.02..."
## [1] "pK = 0.03..."
## [1] "pK = 0.04..."
```

```
## [1] "pK = 0.05..."
## [1] "pK = 0.06..."
## [1] "pK = 0.07..."
## [1] "pK = 0.08..."
## [1] "pK = 0.09..."
## [1] "pK = 0.1..."
## [1] "pK = 0.11..."
## [1] "pK = 0.12..."
## [1] "pK = 0.13..."
## [1] "pK = 0.14..."
## [1] "pK = 0.15..."
## [1] "pK = 0.16..."
## [1] "pK = 0.17..."
## [1] "pK = 0.18..."
## [1] "pK = 0.19..."
## [1] "pK = 0.2..."
## [1] "pK = 0.21..."
## [1] "pK = 0.22..."
## [1] "pK = 0.23..."
## [1] "pK = 0.24..."
## [1] "pK = 0.25..."
## [1] "pK = 0.26..."
## [1] "pK = 0.27..."
## [1] "pK = 0.28..."
## [1] "pK = 0.29..."
## [1] "pK = 0.3..."
## [1] "Creating artificial doublets for pN = 25%"
## [1] "Creating Seurat object..."
## [1] "Normalizing Seurat object..."
```

```
## [1] "Finding variable genes..."
```

```
## [1] "Scaling data..."
```

```
## [1] "Running PCA..."
## [1] "Calculating PC distance matrix..."
## [1] "Defining neighborhoods..."
## [1] "Computing pANN across all pK..."
## [1] "pK = 0.001..."
## [1] "pK = 0.005..."
## [1] "pK = 0.01..."
## [1] "pK = 0.02..."
## [1] "pK = 0.03..."
## [1] "pK = 0.04..."
## [1] "pK = 0.05..."
## [1] "pK = 0.06..."
## [1] "pK = 0.07..."
## [1] "pK = 0.08..."
## [1] "pK = 0.09..."
## [1] "pK = 0.1..."
## [1] "pK = 0.11..."
## [1] "pK = 0.12..."
```

```
## [1] "pK = 0.13..."
## [1] "pK = 0.14..."
## [1] "pK = 0.15..."
## [1] "pK = 0.16..."
## [1] "pK = 0.17..."
## [1] "pK = 0.18..."
## [1] "pK = 0.19..."
## [1] "pK = 0.2..."
## [1] "pK = 0.21..."
## [1] "pK = 0.22..."
## [1] "pK = 0.23..."
## [1] "pK = 0.24..."
## [1] "pK = 0.25..."
## [1] "pK = 0.26..."
## [1] "pK = 0.27..."
## [1] "pK = 0.28..."
## [1] "pK = 0.29..."
## [1] "pK = 0.3..."
## [1] "Creating artificial doublets for pN = 30%"
## [1] "Creating Seurat object..."
## [1] "Normalizing Seurat object..."
```

```
## [1] "Finding variable genes..."
```

```
## [1] "Scaling data..."
```

```
## [1] "Running PCA..."
## [1] "Calculating PC distance matrix..."
## [1] "Defining neighborhoods..."
## [1] "Computing pANN across all pK..."
## [1] "pK = 0.001..."
## [1] "pK = 0.005..."
## [1] "pK = 0.01..."
## [1] "pK = 0.02..."
## [1] "pK = 0.03..."
## [1] "pK = 0.04..."
## [1] "pK = 0.05..."
## [1] "pK = 0.06..."
## [1] "pK = 0.07..."
## [1] "pK = 0.08..."
## [1] "pK = 0.09..."
## [1] "pK = 0.1..."
## [1] "pK = 0.11..."
## [1] "pK = 0.12..."
## [1] "pK = 0.13..."
## [1] "pK = 0.14..."
## [1] "pK = 0.15..."
## [1] "pK = 0.16..."
## [1] "pK = 0.17..."
## [1] "pK = 0.18..."
## [1] "pK = 0.19..."
## [1] "pK = 0.2..."
```

```
## [1] "pK = 0.21..."
## [1] "pK = 0.22..."
## [1] "pK = 0.23..."
## [1] "pK = 0.24..."
## [1] "pK = 0.25..."
## [1] "pK = 0.26..."
## [1] "pK = 0.27..."
## [1] "pK = 0.28..."
## [1] "pK = 0.29..."
## [1] "pK = 0.3..."
```

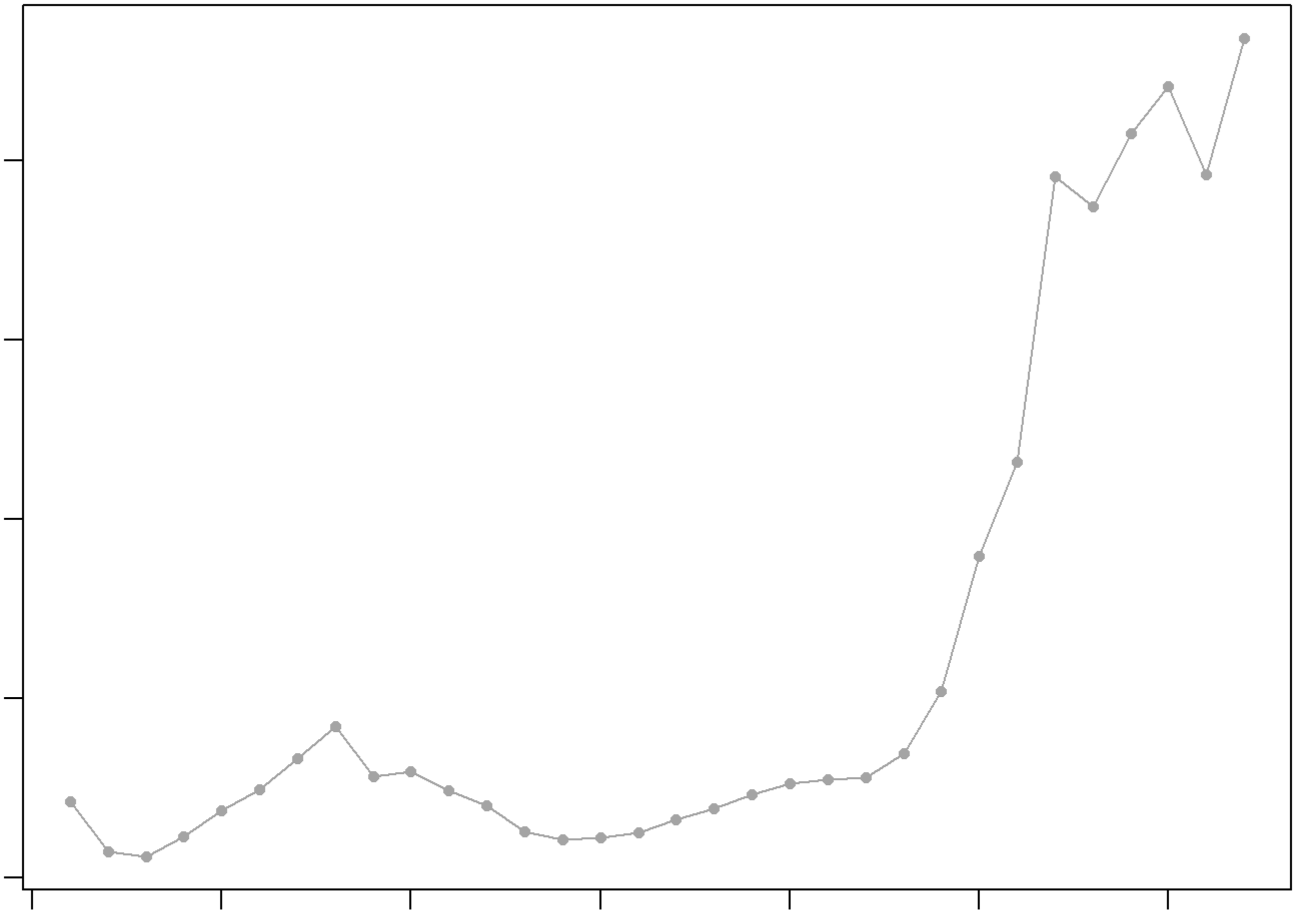

```
## NULL
## [1] "Creating 4283 artificial doublets..."
## [1] "Creating Seurat object..."
## [1] "Normalizing Seurat object..."
```

```
## [1] "Finding variable genes..."
```

```
## [1] "Scaling data..."
```

```
## [1] "Running PCA..."
## [1] "Calculating PC distance matrix..."
## [1] "Computing pANN..."
## [1] "Classifying doublets.."
```

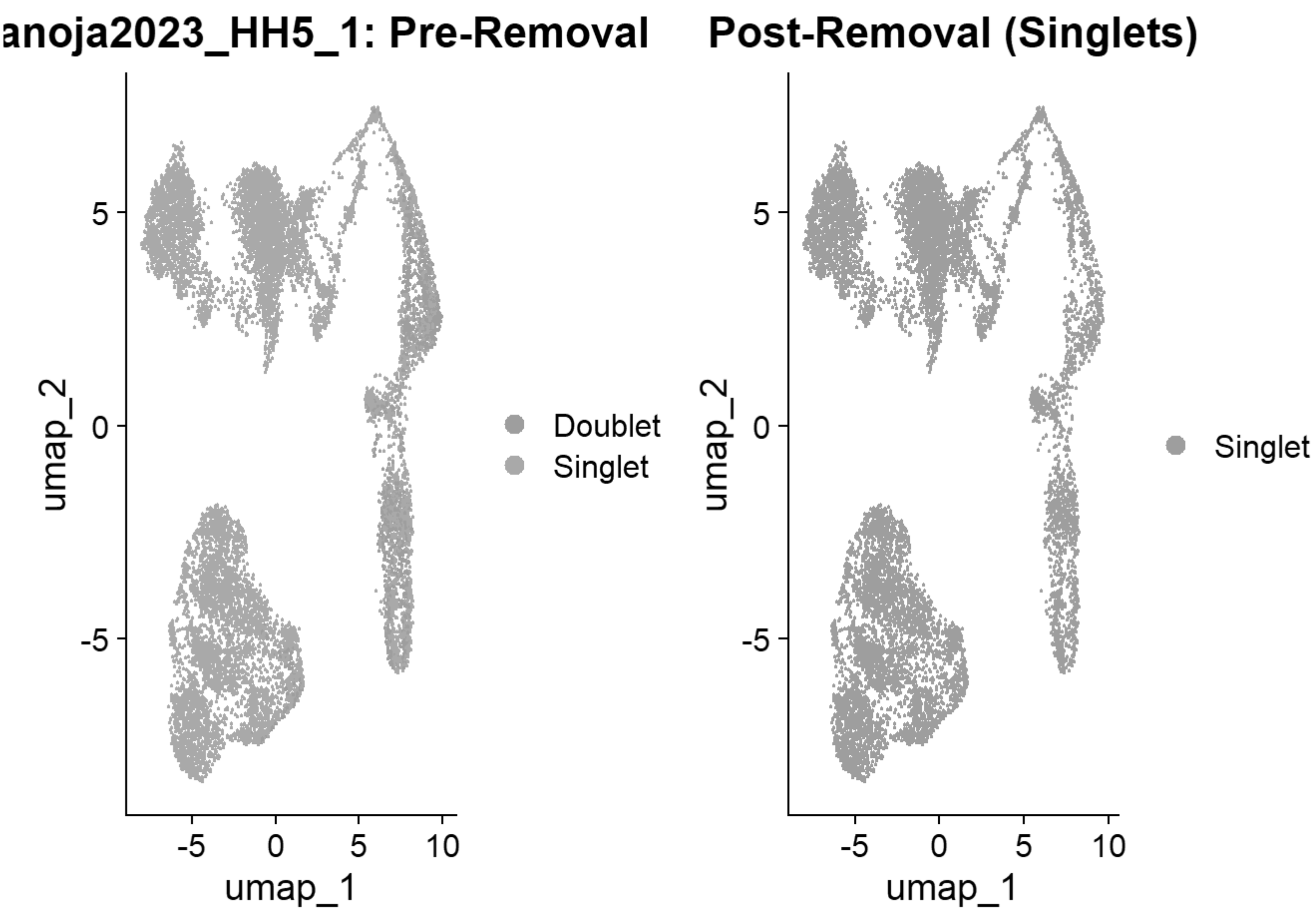


```
## [1] "Loading HH5_2"
## [1] "Running SoupX for HH5_2"
```

```
## Modularity Optimizer version 1.3.0 by Ludo Waltman and Nees Jan van Eck
##
## Number of nodes: 5535
## Number of edges: 207887
##
## Running Louvain algorithm...
## Maximum modularity in 10 random starts: 0.8551
## Number of communities: 8
## Elapsed time: 0 seconds
```

```
## [1] "Running QC for HH5_2"
## [1] "Running DoubletFinder for HH5_2"
## [1] "Creating artificial doublets for pN = 5%"
## [1] "Creating Seurat object..."
## [1] "Normalizing Seurat object..."
```

```
## [1] "Finding variable genes..."
```

```
## [1] "Scaling data..."
```

```
## [1] "Running PCA..."
## [1] "Calculating PC distance matrix..."
## [1] "Defining neighborhoods..."
## [1] "Computing pANN across all pK..."
## [1] "pK = 0.005..."
## [1] "pK = 0.01..."
## [1] "pK = 0.02..."
## [1] "pK = 0.03..."
## [1] "pK = 0.04..."
## [1] "pK = 0.05..."
## [1] "pK = 0.06..."
## [1] "pK = 0.07..."
## [1] "pK = 0.08..."
## [1] "pK = 0.09..."
## [1] "pK = 0.1..."
## [1] "pK = 0.11..."
## [1] "pK = 0.12..."
## [1] "pK = 0.13..."
## [1] "pK = 0.14..."
## [1] "pK = 0.15..."
## [1] "pK = 0.16..."
## [1] "pK = 0.17..."
## [1] "pK = 0.18..."
## [1] "pK = 0.19..."
## [1] "pK = 0.2..."
## [1] "pK = 0.21..."
## [1] "pK = 0.22..."
## [1] "pK = 0.23..."
## [1] "pK = 0.24..."
## [1] "pK = 0.25..."
## [1] "pK = 0.26..."
## [1] "pK = 0.27..."
## [1] "pK = 0.28..."
## [1] "pK = 0.29..."
## [1] "pK = 0.3..."
## [1] "Creating artificial doublets for pN = 10%"
## [1] "Creating Seurat object..."
## [1] "Normalizing Seurat object..."
```

```
## [1] "Finding variable genes..."
```

```
## [1] "Scaling data..."
```

```
## [1] "Running PCA..."
## [1] "Calculating PC distance matrix..."
## [1] "Defining neighborhoods..."
## [1] "Computing pANN across all pK..."
## [1] "pK = 0.005..."
## [1] "pK = 0.01..."
## [1] "pK = 0.02..."
## [1] "pK = 0.03..."
```

```
## [1] "pK = 0.04..."
## [1] "pK = 0.05..."
## [1] "pK = 0.06..."
## [1] "pK = 0.07..."
## [1] "pK = 0.08..."
## [1] "pK = 0.09..."
## [1] "pK = 0.1..."
## [1] "pK = 0.11..."
## [1] "pK = 0.12..."
## [1] "pK = 0.13..."
## [1] "pK = 0.14..."
## [1] "pK = 0.15..."
## [1] "pK = 0.16..."
## [1] "pK = 0.17..."
## [1] "pK = 0.18..."
## [1] "pK = 0.19..."
## [1] "pK = 0.2..."
## [1] "pK = 0.21..."
## [1] "pK = 0.22..."
## [1] "pK = 0.23..."
## [1] "pK = 0.24..."
## [1] "pK = 0.25..."
## [1] "pK = 0.26..."
## [1] "pK = 0.27..."
## [1] "pK = 0.28..."
## [1] "pK = 0.29..."
## [1] "pK = 0.3..."
## [1] "Creating artificial doublets for pN = 15%"
## [1] "Creating Seurat object..."
## [1] "Normalizing Seurat object..."
```

```
## [1] "Finding variable genes..."
```

```
## [1] "Scaling data..."
```

```
## [1] "Running PCA..."
## [1] "Calculating PC distance matrix..."
## [1] "Defining neighborhoods..."
## [1] "Computing pANN across all pK..."
## [1] "pK = 0.005..."
## [1] "pK = 0.01..."
## [1] "pK = 0.02..."
## [1] "pK = 0.03..."
## [1] "pK = 0.04..."
## [1] "pK = 0.05..."
## [1] "pK = 0.06..."
## [1] "pK = 0.07..."
## [1] "pK = 0.08..."
## [1] "pK = 0.09..."
## [1] "pK = 0.1..."
## [1] "pK = 0.11..."
## [1] "pK = 0.12..."
```

```
## [1] "pK = 0.13..."
## [1] "pK = 0.14..."
## [1] "pK = 0.15..."
## [1] "pK = 0.16..."
## [1] "pK = 0.17..."
## [1] "pK = 0.18..."
## [1] "pK = 0.19..."
## [1] "pK = 0.2..."
## [1] "pK = 0.21..."
## [1] "pK = 0.22..."
## [1] "pK = 0.23..."
## [1] "pK = 0.24..."
## [1] "pK = 0.25..."
## [1] "pK = 0.26..."
## [1] "pK = 0.27..."
## [1] "pK = 0.28..."
## [1] "pK = 0.29..."
## [1] "pK = 0.3..."
## [1] "Creating artificial doublets for pN = 20%"
## [1] "Creating Seurat object..."
## [1] "Normalizing Seurat object..."
```

```
## [1] "Finding variable genes..."
```

```
## [1] "Scaling data..."
```

```
## [1] "Running PCA..."
## [1] "Calculating PC distance matrix..."
## [1] "Defining neighborhoods..."
## [1] "Computing pANN across all pK..."
## [1] "pK = 0.005..."
## [1] "pK = 0.01..."
## [1] "pK = 0.02..."
## [1] "pK = 0.03..."
## [1] "pK = 0.04..."
## [1] "pK = 0.05..."
## [1] "pK = 0.06..."
## [1] "pK = 0.07..."
## [1] "pK = 0.08..."
## [1] "pK = 0.09..."
## [1] "pK = 0.1..."
## [1] "pK = 0.11..."
## [1] "pK = 0.12..."
## [1] "pK = 0.13..."
## [1] "pK = 0.14..."
## [1] "pK = 0.15..."
## [1] "pK = 0.16..."
## [1] "pK = 0.17..."
## [1] "pK = 0.18..."
## [1] "pK = 0.19..."
## [1] "pK = 0.2..."
## [1] "pK = 0.21..."
```

```
## [1] "pK = 0.22..."
## [1] "pK = 0.23..."
## [1] "pK = 0.24..."
## [1] "pK = 0.25..."
## [1] "pK = 0.26..."
## [1] "pK = 0.27..."
## [1] "pK = 0.28..."
## [1] "pK = 0.29..."
## [1] "pK = 0.3..."
## [1] "Creating artificial doublets for pN = 25%"
## [1] "Creating Seurat object..."
## [1] "Normalizing Seurat object..."
```

```
## [1] "Finding variable genes..."
```

```
## [1] "Scaling data..."
```

```
## [1] "Running PCA..."
## [1] "Calculating PC distance matrix..."
## [1] "Defining neighborhoods..."
## [1] "Computing pANN across all pK..."
## [1] "pK = 0.005..."
## [1] "pK = 0.01..."
## [1] "pK = 0.02..."
## [1] "pK = 0.03..."
## [1] "pK = 0.04..."
## [1] "pK = 0.05..."
## [1] "pK = 0.06..."
## [1] "pK = 0.07..."
## [1] "pK = 0.08..."
## [1] "pK = 0.09..."
## [1] "pK = 0.1..."
## [1] "pK = 0.11..."
## [1] "pK = 0.12..."
## [1] "pK = 0.13..."
## [1] "pK = 0.14..."
## [1] "pK = 0.15..."
## [1] "pK = 0.16..."
## [1] "pK = 0.17..."
## [1] "pK = 0.18..."
## [1] "pK = 0.19..."
## [1] "pK = 0.2..."
## [1] "pK = 0.21..."
## [1] "pK = 0.22..."
## [1] "pK = 0.23..."
## [1] "pK = 0.24..."
## [1] "pK = 0.25..."
## [1] "pK = 0.26..."
## [1] "pK = 0.27..."
## [1] "pK = 0.28..."
## [1] "pK = 0.29..."
## [1] "pK = 0.3..."
```

```
## [1] "Creating artificial doublets for pN = 30%"
## [1] "Creating Seurat object..."
## [1] "Normalizing Seurat object..."
```

```
## [1] "Finding variable genes..."
```

```
## [1] "Scaling data..."
```

```
## [1] "Running PCA..."
## [1] "Calculating PC distance matrix..."
## [1] "Defining neighborhoods..."
## [1] "Computing pANN across all pK..."
## [1] "pK = 0.005..."
## [1] "pK = 0.01..."
## [1] "pK = 0.02..."
## [1] "pK = 0.03..."
## [1] "pK = 0.04..."
## [1] "pK = 0.05..."
## [1] "pK = 0.06..."
## [1] "pK = 0.07..."
## [1] "pK = 0.08..."
## [1] "pK = 0.09..."
## [1] "pK = 0.1..."
## [1] "pK = 0.11..."
## [1] "pK = 0.12..."
## [1] "pK = 0.13..."
## [1] "pK = 0.14..."
## [1] "pK = 0.15..."
## [1] "pK = 0.16..."
## [1] "pK = 0.17..."
## [1] "pK = 0.18..."
## [1] "pK = 0.19..."
## [1] "pK = 0.2..."
## [1] "pK = 0.21..."
## [1] "pK = 0.22..."
## [1] "pK = 0.23..."
## [1] "pK = 0.24..."
## [1] "pK = 0.25..."
## [1] "pK = 0.26..."
## [1] "pK = 0.27..."
## [1] "pK = 0.28..."
## [1] "pK = 0.29..."
## [1] "pK = 0.3..."
```

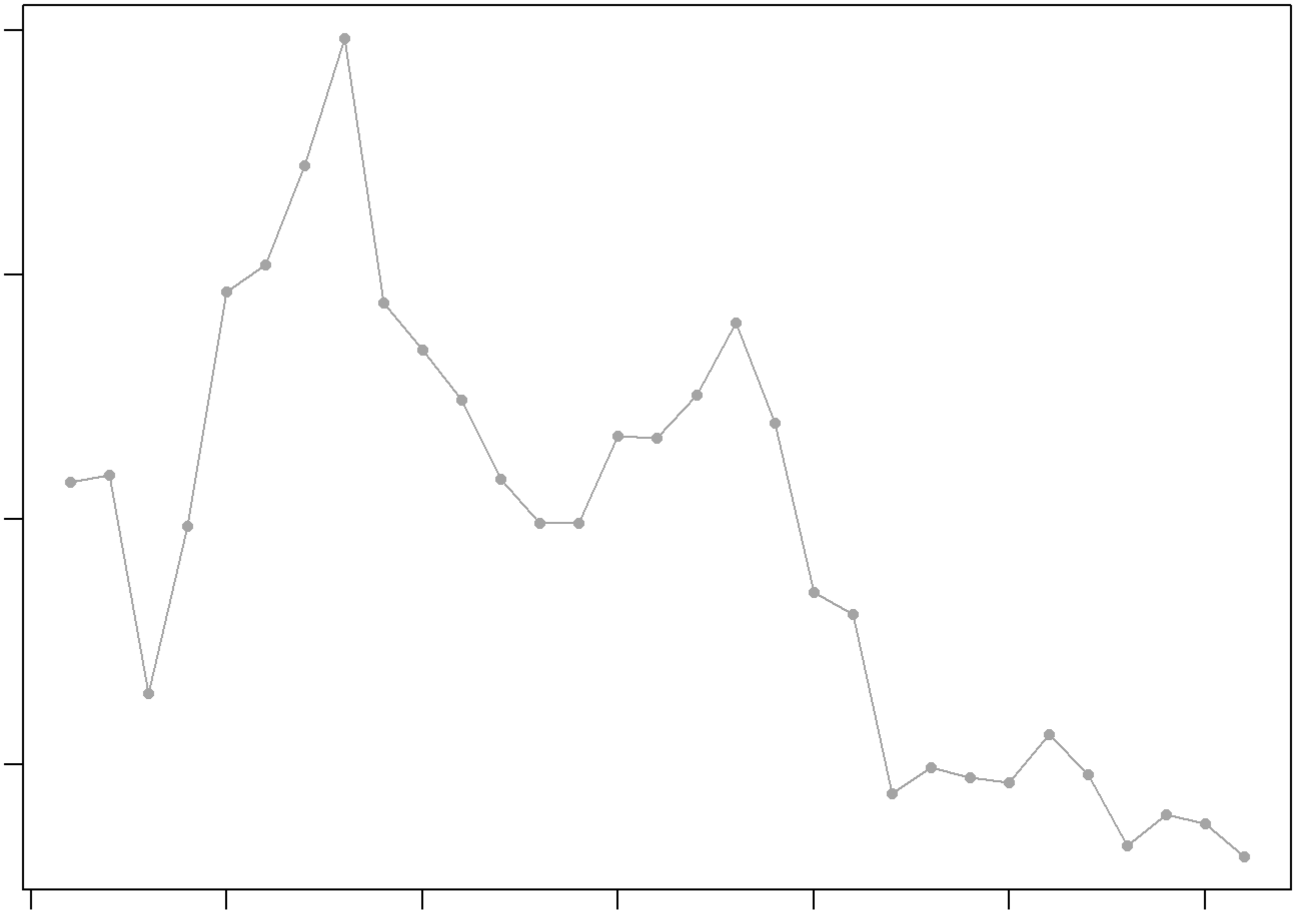

```
## NULL
## [1] "Creating 1843 artificial doublets..."
## [1] "Creating Seurat object..."
## [1] "Normalizing Seurat object..."
```

```
## [1] "Finding variable genes..."
```

```
## [1] "Scaling data..."
```

```
## [1] "Running PCA..."
## [1] "Calculating PC distance matrix..."
## [1] "Computing pANN..."
## [1] "Classifying doublets.."
```

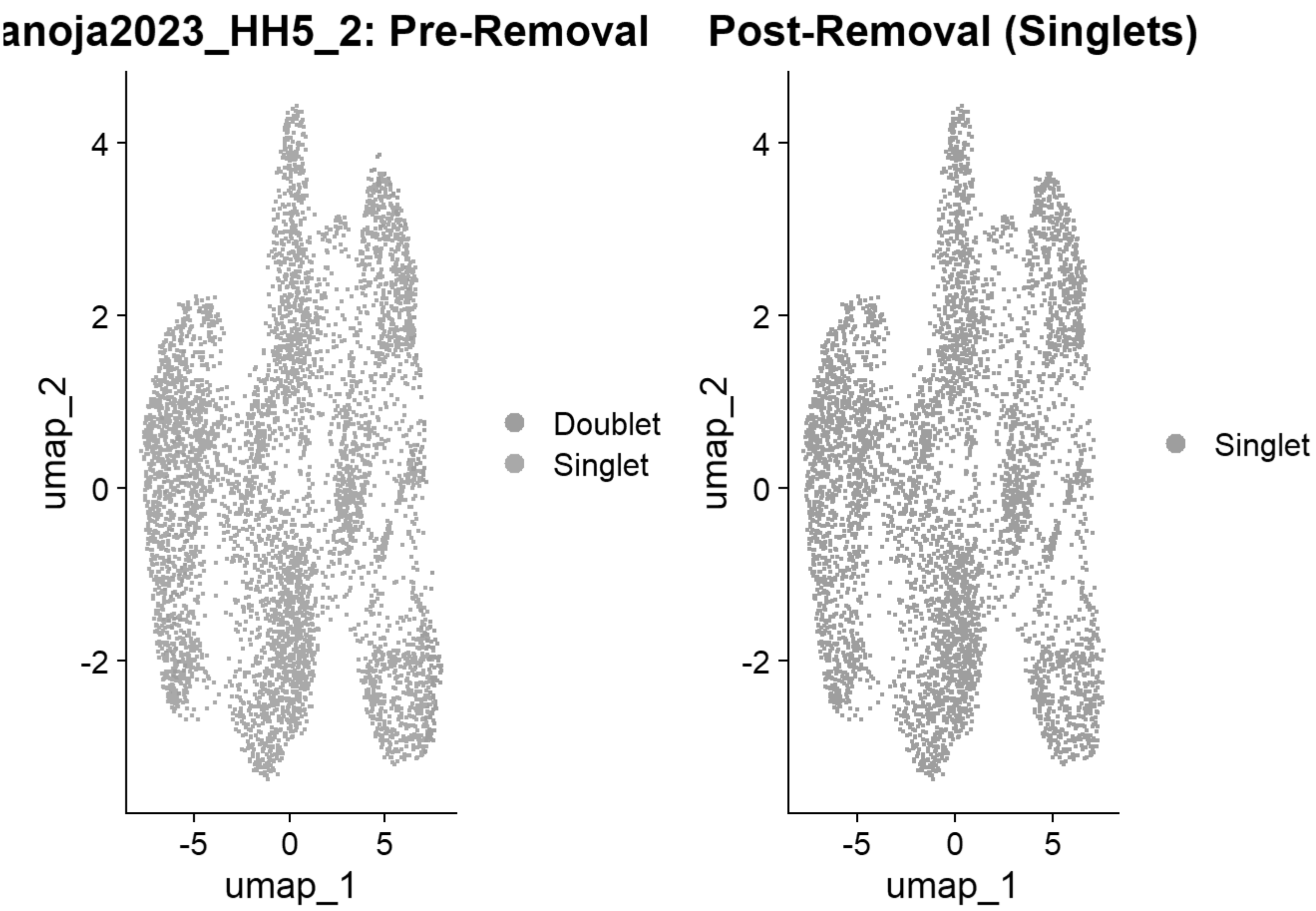


```
## [1] "Loading HH7_1"
## [1] "Running SoupX for HH7_1"
```

```
## Modularity Optimizer version 1.3.0 by Ludo Waltman and Nees Jan van Eck
##
## Number of nodes: 6141
## Number of edges: 209749
##
## Running Louvain algorithm...
## Maximum modularity in 10 random starts: 0.8609
## Number of communities: 9
## Elapsed time: 0 seconds
```

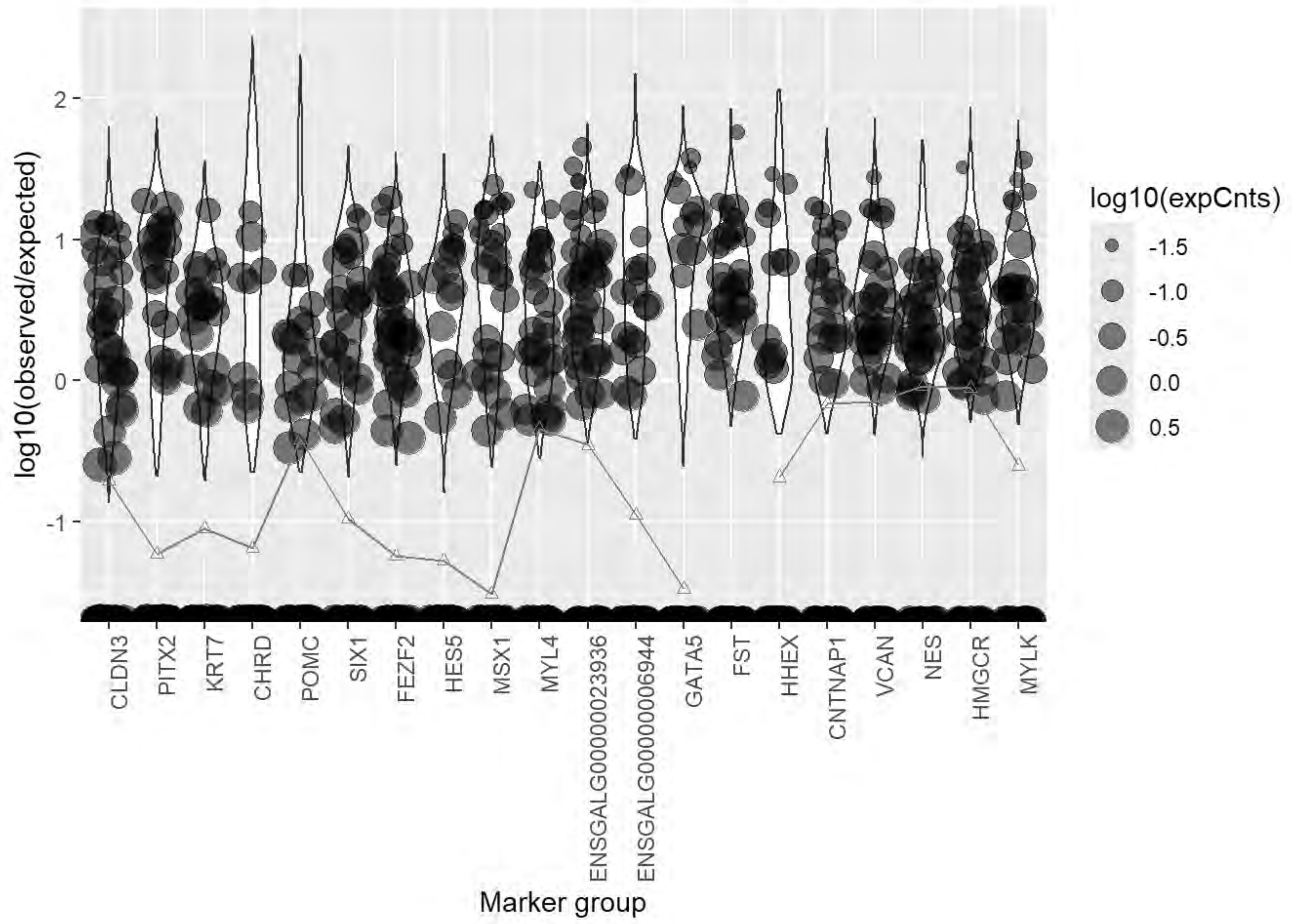
log10(observed/expected)
2
1
0
-1
CLDN3
PITX2
KRT7
CHRD
POMC
SIX1
FEZF2
HES5
MSX1
MYL4
ENSGALG00000023936
ENSGALG00000006944
GATA5
FST
HHEX
CNTNAP1
VCAN
NES
HMGCR
MYLK
Marker group
log10(expCnts)
-1.5
-1.0
-0.5
0.0
0.5

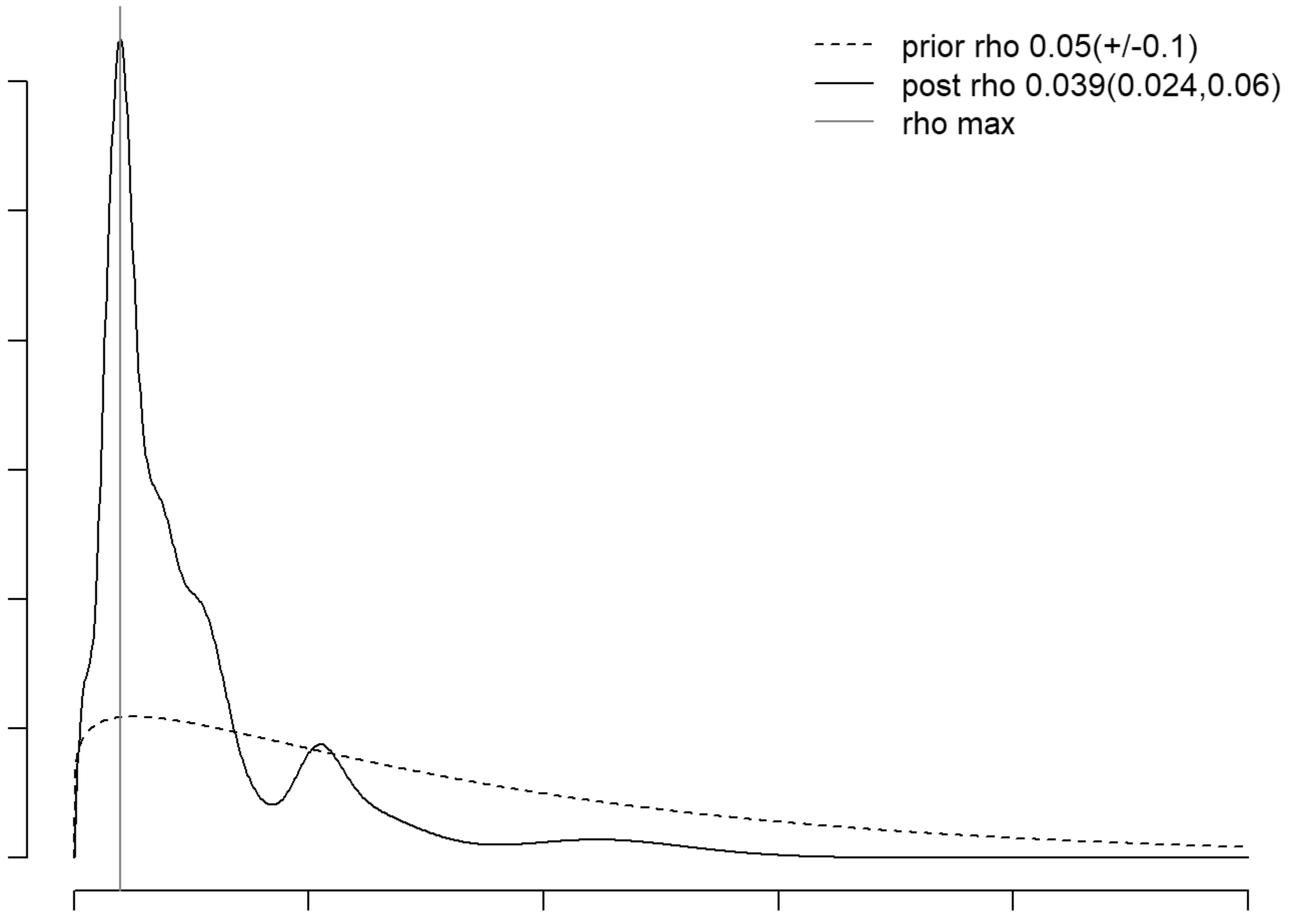
prior rho 0.05(+/-0.1)
post rho 0.039(0.024,0.06)
rho max

```
## [1] "Running QC for HH7_1"
## [1] "Running DoubletFinder for HH7_1"
## [1] "Creating artificial doublets for pN = 5%"
## [1] "Creating Seurat object..."
## [1] "Normalizing Seurat object..."
```

```
## [1] "Finding variable genes..."
```

```
## [1] "Scaling data..."
```

```
## [1] "Running PCA..."
## [1] "Calculating PC distance matrix..."
## [1] "Defining neighborhoods..."
## [1] "Computing pANN across all pK..."
## [1] "pK = 0.005..."
## [1] "pK = 0.01..."
## [1] "pK = 0.02..."
## [1] "pK = 0.03..."
## [1] "pK = 0.04..."
## [1] "pK = 0.05..."
## [1] "pK = 0.06..."
## [1] "pK = 0.07..."
## [1] "pK = 0.08..."
## [1] "pK = 0.09..."
## [1] "pK = 0.1..."
## [1] "pK = 0.11..."
## [1] "pK = 0.12..."
## [1] "pK = 0.13..."
## [1] "pK = 0.14..."
## [1] "pK = 0.15..."
## [1] "pK = 0.16..."
## [1] "pK = 0.17..."
## [1] "pK = 0.18..."
## [1] "pK = 0.19..."
## [1] "pK = 0.2..."
## [1] "pK = 0.21..."
## [1] "pK = 0.22..."
## [1] "pK = 0.23..."
## [1] "pK = 0.24..."
## [1] "pK = 0.25..."
## [1] "pK = 0.26..."
## [1] "pK = 0.27..."
## [1] "pK = 0.28..."
## [1] "pK = 0.29..."
## [1] "pK = 0.3..."
## [1] "Creating artificial doublets for pN = 10%"
## [1] "Creating Seurat object..."
## [1] "Normalizing Seurat object..."
```

```
## [1] "Finding variable genes..."
```

```
## [1] "Scaling data..."
```

```
## [1] "Running PCA..."
## [1] "Calculating PC distance matrix..."
## [1] "Defining neighborhoods..."
## [1] "Computing pANN across all pK..."
## [1] "pK = 0.005..."
## [1] "pK = 0.01..."
## [1] "pK = 0.02..."
## [1] "pK = 0.03..."
## [1] "pK = 0.04..."
## [1] "pK = 0.05..."
## [1] "pK = 0.06..."
## [1] "pK = 0.07..."
## [1] "pK = 0.08..."
## [1] "pK = 0.09..."
## [1] "pK = 0.1..."
## [1] "pK = 0.11..."
## [1] "pK = 0.12..."
## [1] "pK = 0.13..."
## [1] "pK = 0.14..."
## [1] "pK = 0.15..."
## [1] "pK = 0.16..."
## [1] "pK = 0.17..."
## [1] "pK = 0.18..."
## [1] "pK = 0.19..."
## [1] "pK = 0.2..."
## [1] "pK = 0.21..."
## [1] "pK = 0.22..."
## [1] "pK = 0.23..."
## [1] "pK = 0.24..."
## [1] "pK = 0.25..."
## [1] "pK = 0.26..."
## [1] "pK = 0.27..."
## [1] "pK = 0.28..."
## [1] "pK = 0.29..."
## [1] "pK = 0.3..."
## [1] "Creating artificial doublets for pN = 15%"
## [1] "Creating Seurat object..."
## [1] "Normalizing Seurat object..."
```

```
## [1] "Finding variable genes..."
```

```
## [1] "Scaling data..."
```

```
## [1] "Running PCA..."
## [1] "Calculating PC distance matrix..."
## [1] "Defining neighborhoods..."
## [1] "Computing pANN across all pK..."
## [1] "pK = 0.005..."
```

```
## [1] "pK = 0.01..."
## [1] "pK = 0.02..."
## [1] "pK = 0.03..."
## [1] "pK = 0.04..."
## [1] "pK = 0.05..."
## [1] "pK = 0.06..."
## [1] "pK = 0.07..."
## [1] "pK = 0.08..."
## [1] "pK = 0.09..."
## [1] "pK = 0.1..."
## [1] "pK = 0.11..."
## [1] "pK = 0.12..."
## [1] "pK = 0.13..."
## [1] "pK = 0.14..."
## [1] "pK = 0.15..."
## [1] "pK = 0.16..."
## [1] "pK = 0.17..."
## [1] "pK = 0.18..."
## [1] "pK = 0.19..."
## [1] "pK = 0.2..."
## [1] "pK = 0.21..."
## [1] "pK = 0.22..."
## [1] "pK = 0.23..."
## [1] "pK = 0.24..."
## [1] "pK = 0.25..."
## [1] "pK = 0.26..."
## [1] "pK = 0.27..."
## [1] "pK = 0.28..."
## [1] "pK = 0.29..."
## [1] "pK = 0.3..."
## [1] "Creating artificial doublets for pN = 20%"
## [1] "Creating Seurat object..."
## [1] "Normalizing Seurat object..."
```

```
## [1] "Finding variable genes..."
```

```
## [1] "Scaling data..."
```

```
## [1] "Running PCA..."
## [1] "Calculating PC distance matrix..."
## [1] "Defining neighborhoods..."
## [1] "Computing pANN across all pK..."
## [1] "pK = 0.005..."
## [1] "pK = 0.01..."
## [1] "pK = 0.02..."
## [1] "pK = 0.03..."
## [1] "pK = 0.04..."
## [1] "pK = 0.05..."
## [1] "pK = 0.06..."
## [1] "pK = 0.07..."
## [1] "pK = 0.08..."
## [1] "pK = 0.09..."
```

```
## [1] "pK = 0.1..."
## [1] "pK = 0.11..."
## [1] "pK = 0.12..."
## [1] "pK = 0.13..."
## [1] "pK = 0.14..."
## [1] "pK = 0.15..."
## [1] "pK = 0.16..."
## [1] "pK = 0.17..."
## [1] "pK = 0.18..."
## [1] "pK = 0.19..."
## [1] "pK = 0.2..."
## [1] "pK = 0.21..."
## [1] "pK = 0.22..."
## [1] "pK = 0.23..."
## [1] "pK = 0.24..."
## [1] "pK = 0.25..."
## [1] "pK = 0.26..."
## [1] "pK = 0.27..."
## [1] "pK = 0.28..."
## [1] "pK = 0.29..."
## [1] "pK = 0.3..."
## [1] "Creating artificial doublets for pN = 25%"
## [1] "Creating Seurat object..."
## [1] "Normalizing Seurat object..."
```

```
## [1] "Finding variable genes..."
```

```
## [1] "Scaling data..."
```

```
## [1] "Running PCA..."
## [1] "Calculating PC distance matrix..."
## [1] "Defining neighborhoods..."
## [1] "Computing pANN across all pK..."
## [1] "pK = 0.005..."
## [1] "pK = 0.01..."
## [1] "pK = 0.02..."
## [1] "pK = 0.03..."
## [1] "pK = 0.04..."
## [1] "pK = 0.05..."
## [1] "pK = 0.06..."
## [1] "pK = 0.07..."
## [1] "pK = 0.08..."
## [1] "pK = 0.09..."
## [1] "pK = 0.1..."
## [1] "pK = 0.11..."
## [1] "pK = 0.12..."
## [1] "pK = 0.13..."
## [1] "pK = 0.14..."
## [1] "pK = 0.15..."
## [1] "pK = 0.16..."
## [1] "pK = 0.17..."
## [1] "pK = 0.18..."
```

```
## [1] "pK = 0.19..."
## [1] "pK = 0.2..."
## [1] "pK = 0.21..."
## [1] "pK = 0.22..."
## [1] "pK = 0.23..."
## [1] "pK = 0.24..."
## [1] "pK = 0.25..."
## [1] "pK = 0.26..."
## [1] "pK = 0.27..."
## [1] "pK = 0.28..."
## [1] "pK = 0.29..."
## [1] "pK = 0.3..."
## [1] "Creating artificial doublets for pN = 30%"
## [1] "Creating Seurat object..."
## [1] "Normalizing Seurat object..."
```

```
## [1] "Finding variable genes..."
```

```
## [1] "Scaling data..."
```

```
## [1] "Running PCA..."
## [1] "Calculating PC distance matrix..."
## [1] "Defining neighborhoods..."
## [1] "Computing pANN across all pK..."
## [1] "pK = 0.005..."
## [1] "pK = 0.01..."
## [1] "pK = 0.02..."
## [1] "pK = 0.03..."
## [1] "pK = 0.04..."
## [1] "pK = 0.05..."
## [1] "pK = 0.06..."
## [1] "pK = 0.07..."
## [1] "pK = 0.08..."
## [1] "pK = 0.09..."
## [1] "pK = 0.1..."
## [1] "pK = 0.11..."
## [1] "pK = 0.12..."
## [1] "pK = 0.13..."
## [1] "pK = 0.14..."
## [1] "pK = 0.15..."
## [1] "pK = 0.16..."
## [1] "pK = 0.17..."
## [1] "pK = 0.18..."
## [1] "pK = 0.19..."
## [1] "pK = 0.2..."
## [1] "pK = 0.21..."
## [1] "pK = 0.22..."
## [1] "pK = 0.23..."
## [1] "pK = 0.24..."
## [1] "pK = 0.25..."
## [1] "pK = 0.26..."
## [1] "pK = 0.27..."
```

```
## [1] "pK = 0.28..."
## [1] "pK = 0.29..."
## [1] "pK = 0.3..."
```

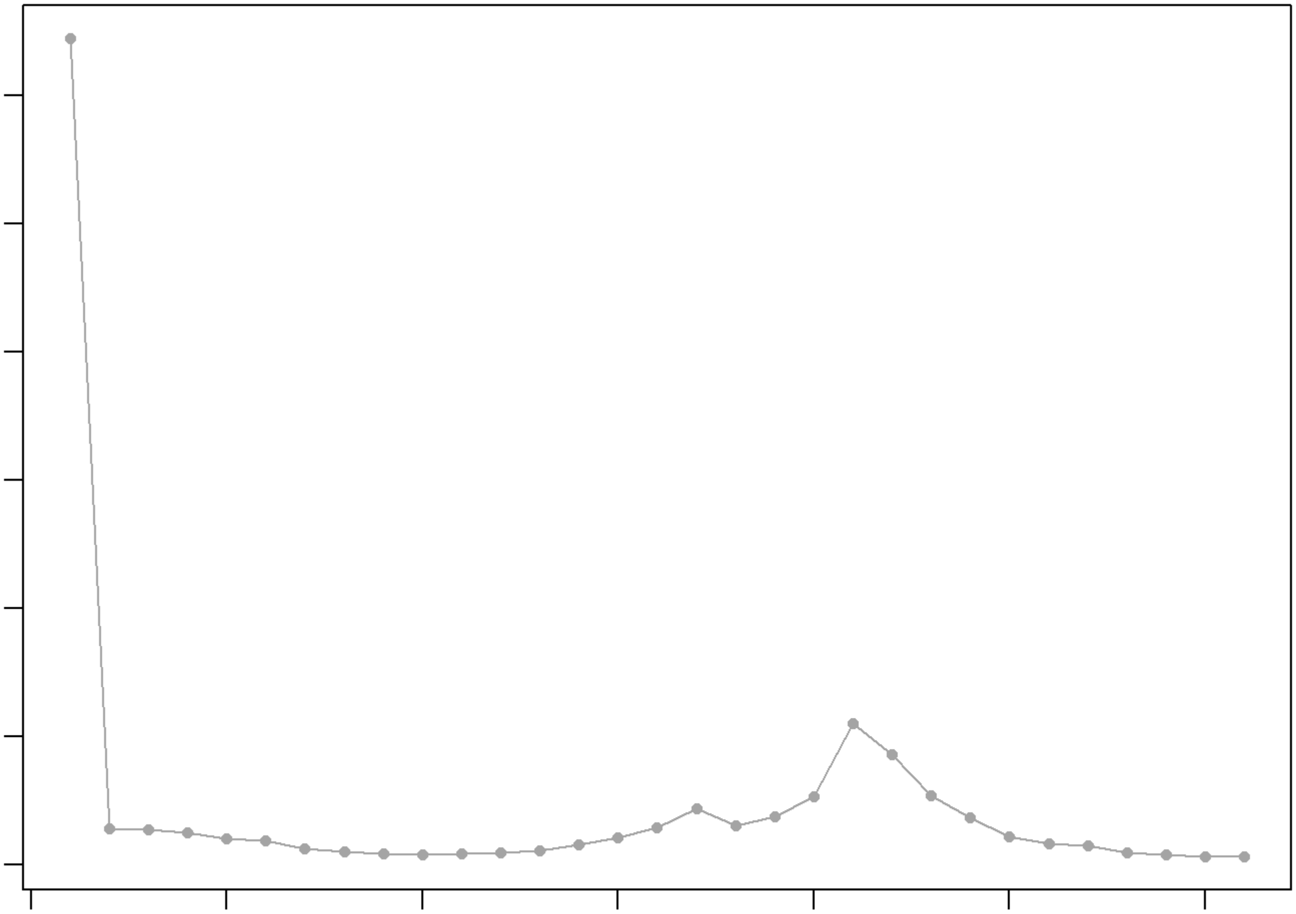

```
## NULL
## [1] "Creating 1929 artificial doublets..."
## [1] "Creating Seurat object..."
## [1] "Normalizing Seurat object..."
```

```
## [1] "Finding variable genes..."
```

```
## [1] "Scaling data..."
```

```
## [1] "Running PCA..."
## [1] "Calculating PC distance matrix..."
## [1] "Computing pANN..."
## [1] "Classifying doublets.."
```

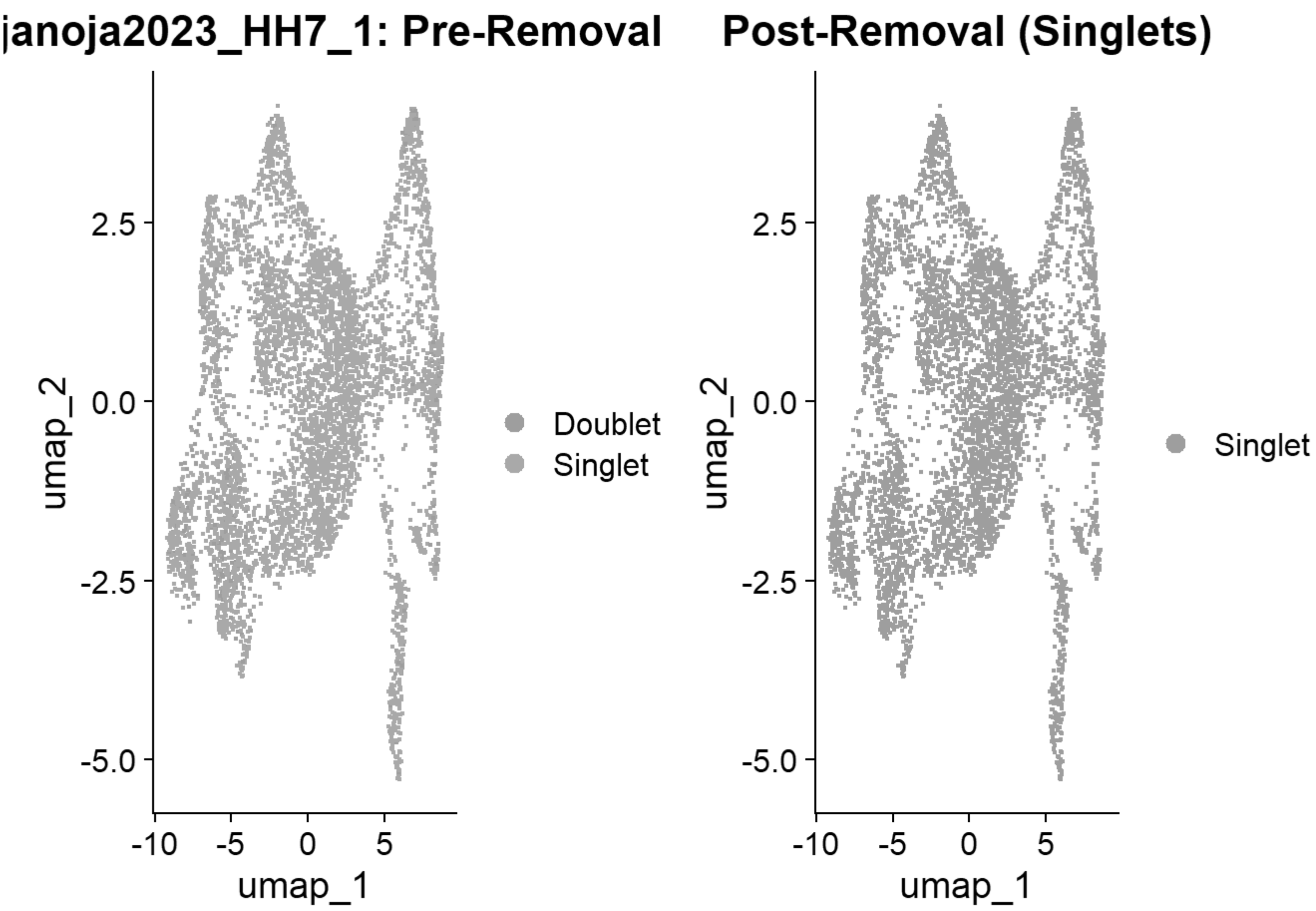


```
## [1] "Loading HH7_2"
## [1] "Running SoupX for HH7_2"
```

```
## Modularity Optimizer version 1.3.0 by Ludo Waltman and Nees Jan van Eck
##
## Number of nodes: 6225
## Number of edges: 212635
##
## Running Louvain algorithm...
## Maximum modularity in 10 random starts: 0.8589
## Number of communities: 10
## Elapsed time: 0 seconds
```

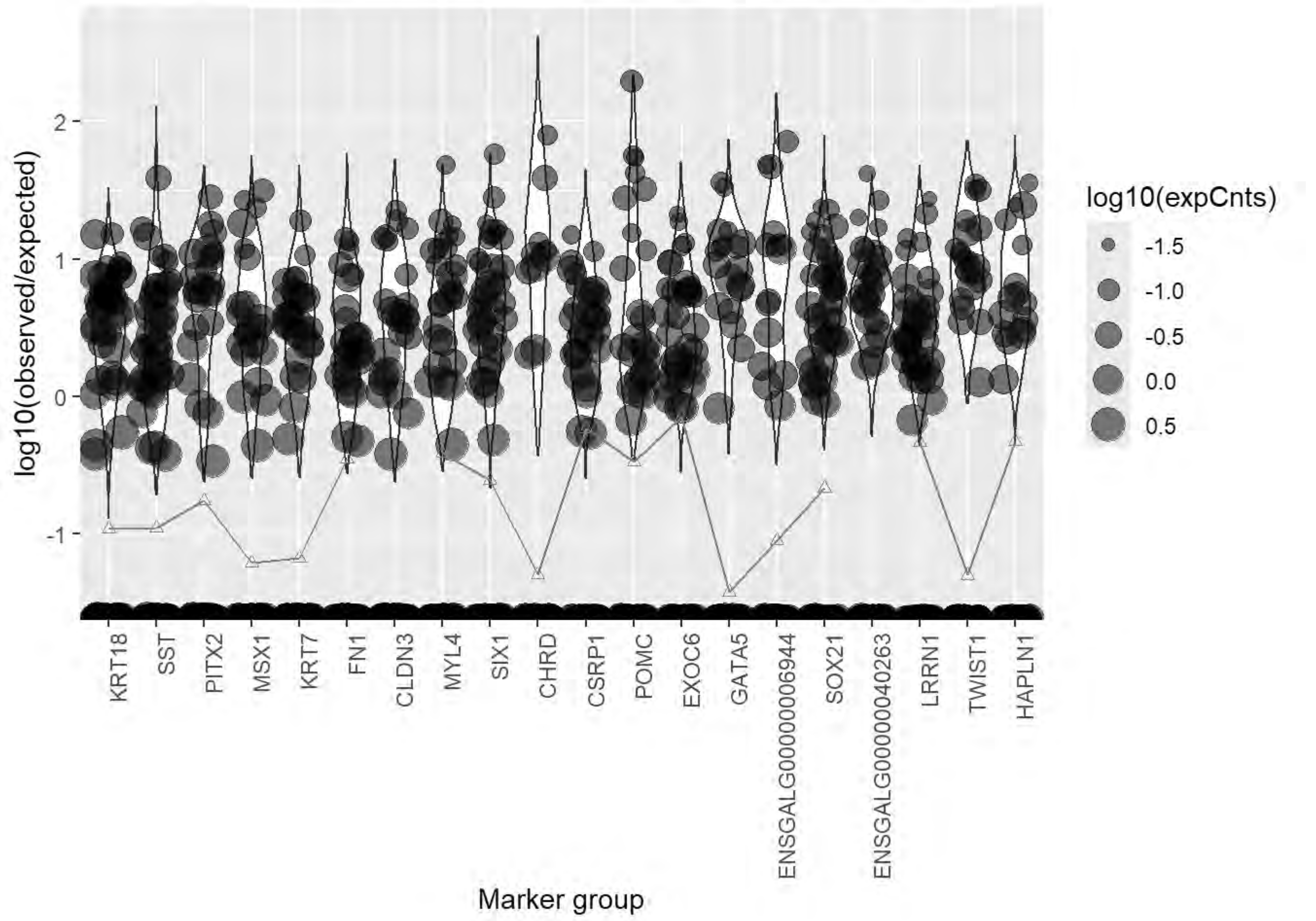
log10(observed/expected)
2
1
0
-1
KRT18
SST
PITX2
MSX1
KRT7
FN1
CLDN3
MYL4
SIX1
CHRD
CSRP1
POMC
EXOC6
GATA5
ENSGALG00000006944
SOX21
ENSGALG00000040263
LRRN1
TWIST1
HAPLN1
Marker group
log10(expCnts)
-1.5
-1.0
-0.5
0.0
0.5

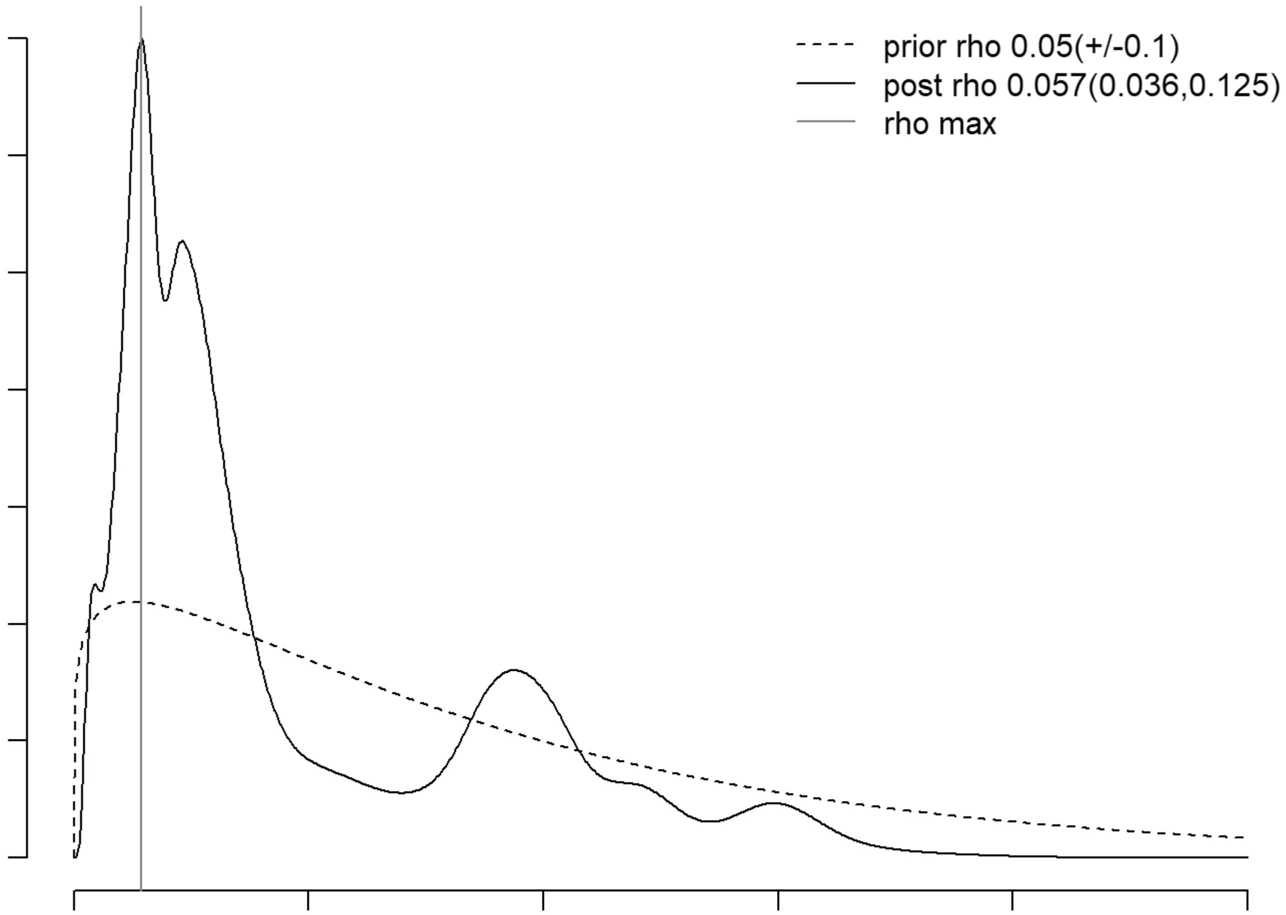
prior rho 0.05(+/-0.1)
post rho 0.057(0.036,0.125)
rho max

```
## [1] "Running QC for HH7_2"
## [1] "Running DoubletFinder for HH7_2"
## [1] "Creating artificial doublets for pN = 5%"
## [1] "Creating Seurat object..."
## [1] "Normalizing Seurat object..."
```

```
## [1] "Finding variable genes..."
```

```
## [1] "Scaling data..."
```

```
## [1] "Running PCA..."
## [1] "Calculating PC distance matrix..."
## [1] "Defining neighborhoods..."
## [1] "Computing pANN across all pK..."
## [1] "pK = 0.005..."
## [1] "pK = 0.01..."
## [1] "pK = 0.02..."
## [1] "pK = 0.03..."
## [1] "pK = 0.04..."
## [1] "pK = 0.05..."
## [1] "pK = 0.06..."
## [1] "pK = 0.07..."
## [1] "pK = 0.08..."
## [1] "pK = 0.09..."
## [1] "pK = 0.1..."
## [1] "pK = 0.11..."
## [1] "pK = 0.12..."
## [1] "pK = 0.13..."
## [1] "pK = 0.14..."
## [1] "pK = 0.15..."
## [1] "pK = 0.16..."
## [1] "pK = 0.17..."
## [1] "pK = 0.18..."
## [1] "pK = 0.19..."
## [1] "pK = 0.2..."
## [1] "pK = 0.21..."
## [1] "pK = 0.22..."
## [1] "pK = 0.23..."
## [1] "pK = 0.24..."
## [1] "pK = 0.25..."
## [1] "pK = 0.26..."
## [1] "pK = 0.27..."
## [1] "pK = 0.28..."
## [1] "pK = 0.29..."
## [1] "pK = 0.3..."
## [1] "Creating artificial doublets for pN = 10%"
## [1] "Creating Seurat object..."
## [1] "Normalizing Seurat object..."
```

```
## [1] "Finding variable genes..."
```

```
## [1] "Scaling data..."
```

```
## [1] "Running PCA..."
## [1] "Calculating PC distance matrix..."
## [1] "Defining neighborhoods..."
## [1] "Computing pANN across all pK..."
## [1] "pK = 0.005..."
## [1] "pK = 0.01..."
## [1] "pK = 0.02..."
## [1] "pK = 0.03..."
## [1] "pK = 0.04..."
## [1] "pK = 0.05..."
## [1] "pK = 0.06..."
## [1] "pK = 0.07..."
## [1] "pK = 0.08..."
## [1] "pK = 0.09..."
## [1] "pK = 0.1..."
## [1] "pK = 0.11..."
## [1] "pK = 0.12..."
## [1] "pK = 0.13..."
## [1] "pK = 0.14..."
## [1] "pK = 0.15..."
## [1] "pK = 0.16..."
## [1] "pK = 0.17..."
## [1] "pK = 0.18..."
## [1] "pK = 0.19..."
## [1] "pK = 0.2..."
## [1] "pK = 0.21..."
## [1] "pK = 0.22..."
## [1] "pK = 0.23..."
## [1] "pK = 0.24..."
## [1] "pK = 0.25..."
## [1] "pK = 0.26..."
## [1] "pK = 0.27..."
## [1] "pK = 0.28..."
## [1] "pK = 0.29..."
## [1] "pK = 0.3..."
## [1] "Creating artificial doublets for pN = 15%"
## [1] "Creating Seurat object..."
## [1] "Normalizing Seurat object..."
```

```
## [1] "Finding variable genes..."
```

```
## [1] "Scaling data..."
```

```
## [1] "Running PCA..."
## [1] "Calculating PC distance matrix..."
## [1] "Defining neighborhoods..."
## [1] "Computing pANN across all pK..."
## [1] "pK = 0.005..."
```

```
## [1] "pK = 0.01..."
## [1] "pK = 0.02..."
## [1] "pK = 0.03..."
## [1] "pK = 0.04..."
## [1] "pK = 0.05..."
## [1] "pK = 0.06..."
## [1] "pK = 0.07..."
## [1] "pK = 0.08..."
## [1] "pK = 0.09..."
## [1] "pK = 0.1..."
## [1] "pK = 0.11..."
## [1] "pK = 0.12..."
## [1] "pK = 0.13..."
## [1] "pK = 0.14..."
## [1] "pK = 0.15..."
## [1] "pK = 0.16..."
## [1] "pK = 0.17..."
## [1] "pK = 0.18..."
## [1] "pK = 0.19..."
## [1] "pK = 0.2..."
## [1] "pK = 0.21..."
## [1] "pK = 0.22..."
## [1] "pK = 0.23..."
## [1] "pK = 0.24..."
## [1] "pK = 0.25..."
## [1] "pK = 0.26..."
## [1] "pK = 0.27..."
## [1] "pK = 0.28..."
## [1] "pK = 0.29..."
## [1] "pK = 0.3..."
## [1] "Creating artificial doublets for pN = 20%"
## [1] "Creating Seurat object..."
## [1] "Normalizing Seurat object..."
```

```
## [1] "Finding variable genes..."
```

```
## [1] "Scaling data..."
```

```
## [1] "Running PCA..."
## [1] "Calculating PC distance matrix..."
## [1] "Defining neighborhoods..."
## [1] "Computing pANN across all pK..."
## [1] "pK = 0.005..."
## [1] "pK = 0.01..."
## [1] "pK = 0.02..."
## [1] "pK = 0.03..."
## [1] "pK = 0.04..."
## [1] "pK = 0.05..."
## [1] "pK = 0.06..."
## [1] "pK = 0.07..."
## [1] "pK = 0.08..."
## [1] "pK = 0.09..."
```

```
## [1] "pK = 0.1..."
## [1] "pK = 0.11..."
## [1] "pK = 0.12..."
## [1] "pK = 0.13..."
## [1] "pK = 0.14..."
## [1] "pK = 0.15..."
## [1] "pK = 0.16..."
## [1] "pK = 0.17..."
## [1] "pK = 0.18..."
## [1] "pK = 0.19..."
## [1] "pK = 0.2..."
## [1] "pK = 0.21..."
## [1] "pK = 0.22..."
## [1] "pK = 0.23..."
## [1] "pK = 0.24..."
## [1] "pK = 0.25..."
## [1] "pK = 0.26..."
## [1] "pK = 0.27..."
## [1] "pK = 0.28..."
## [1] "pK = 0.29..."
## [1] "pK = 0.3..."
## [1] "Creating artificial doublets for pN = 25%"
## [1] "Creating Seurat object..."
## [1] "Normalizing Seurat object..."
```

```
## [1] "Finding variable genes..."
```

```
## [1] "Scaling data..."
```

```
## [1] "Running PCA..."
## [1] "Calculating PC distance matrix..."
## [1] "Defining neighborhoods..."
## [1] "Computing pANN across all pK..."
## [1] "pK = 0.005..."
## [1] "pK = 0.01..."
## [1] "pK = 0.02..."
## [1] "pK = 0.03..."
## [1] "pK = 0.04..."
## [1] "pK = 0.05..."
## [1] "pK = 0.06..."
## [1] "pK = 0.07..."
## [1] "pK = 0.08..."
## [1] "pK = 0.09..."
## [1] "pK = 0.1..."
## [1] "pK = 0.11..."
## [1] "pK = 0.12..."
## [1] "pK = 0.13..."
## [1] "pK = 0.14..."
## [1] "pK = 0.15..."
## [1] "pK = 0.16..."
## [1] "pK = 0.17..."
## [1] "pK = 0.18..."
```

```
## [1] "pK = 0.19..."
## [1] "pK = 0.2..."
## [1] "pK = 0.21..."
## [1] "pK = 0.22..."
## [1] "pK = 0.23..."
## [1] "pK = 0.24..."
## [1] "pK = 0.25..."
## [1] "pK = 0.26..."
## [1] "pK = 0.27..."
## [1] "pK = 0.28..."
## [1] "pK = 0.29..."
## [1] "pK = 0.3..."
## [1] "Creating artificial doublets for pN = 30%"
## [1] "Creating Seurat object..."
## [1] "Normalizing Seurat object..."
```

```
## [1] "Finding variable genes..."
```

```
## [1] "Scaling data..."
```

```
## [1] "Running PCA..."
## [1] "Calculating PC distance matrix..."
## [1] "Defining neighborhoods..."
## [1] "Computing pANN across all pK..."
## [1] "pK = 0.005..."
## [1] "pK = 0.01..."
## [1] "pK = 0.02..."
## [1] "pK = 0.03..."
## [1] "pK = 0.04..."
## [1] "pK = 0.05..."
## [1] "pK = 0.06..."
## [1] "pK = 0.07..."
## [1] "pK = 0.08..."
## [1] "pK = 0.09..."
## [1] "pK = 0.1..."
## [1] "pK = 0.11..."
## [1] "pK = 0.12..."
## [1] "pK = 0.13..."
## [1] "pK = 0.14..."
## [1] "pK = 0.15..."
## [1] "pK = 0.16..."
## [1] "pK = 0.17..."
## [1] "pK = 0.18..."
## [1] "pK = 0.19..."
## [1] "pK = 0.2..."
## [1] "pK = 0.21..."
## [1] "pK = 0.22..."
## [1] "pK = 0.23..."
## [1] "pK = 0.24..."
## [1] "pK = 0.25..."
## [1] "pK = 0.26..."
## [1] "pK = 0.27..."
```

```
## [1] "pK = 0.28..."
## [1] "pK = 0.29..."
## [1] "pK = 0.3..."
```

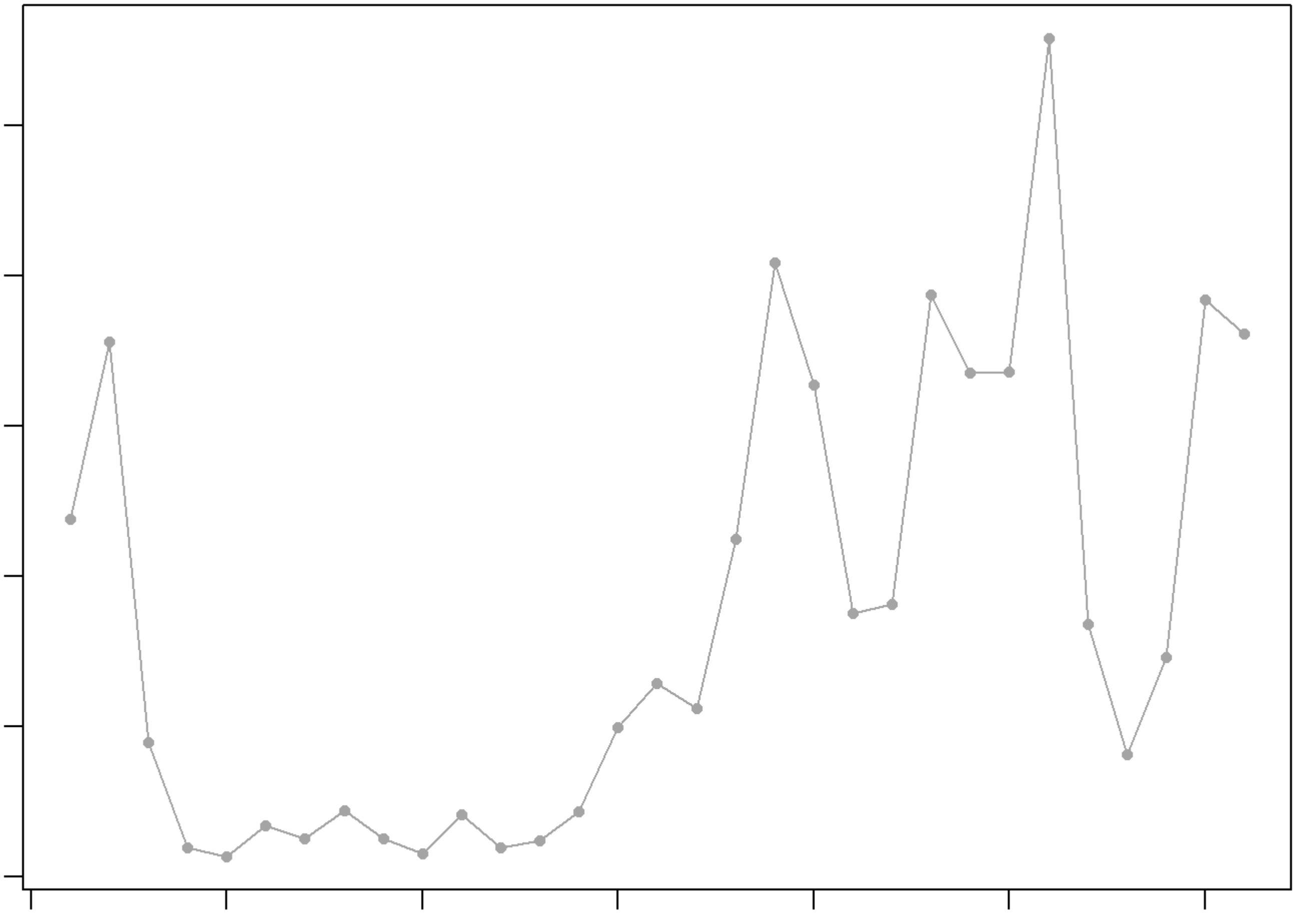

```
## NULL
## [1] "Creating 2051 artificial doublets..."
## [1] "Creating Seurat object..."
## [1] "Normalizing Seurat object..."
```

```
## [1] "Finding variable genes..."
```

```
## [1] "Scaling data..."
```

```
## [1] "Running PCA..."
## [1] "Calculating PC distance matrix..."
## [1] "Computing pANN..."
## [1] "Classifying doublets.."
```

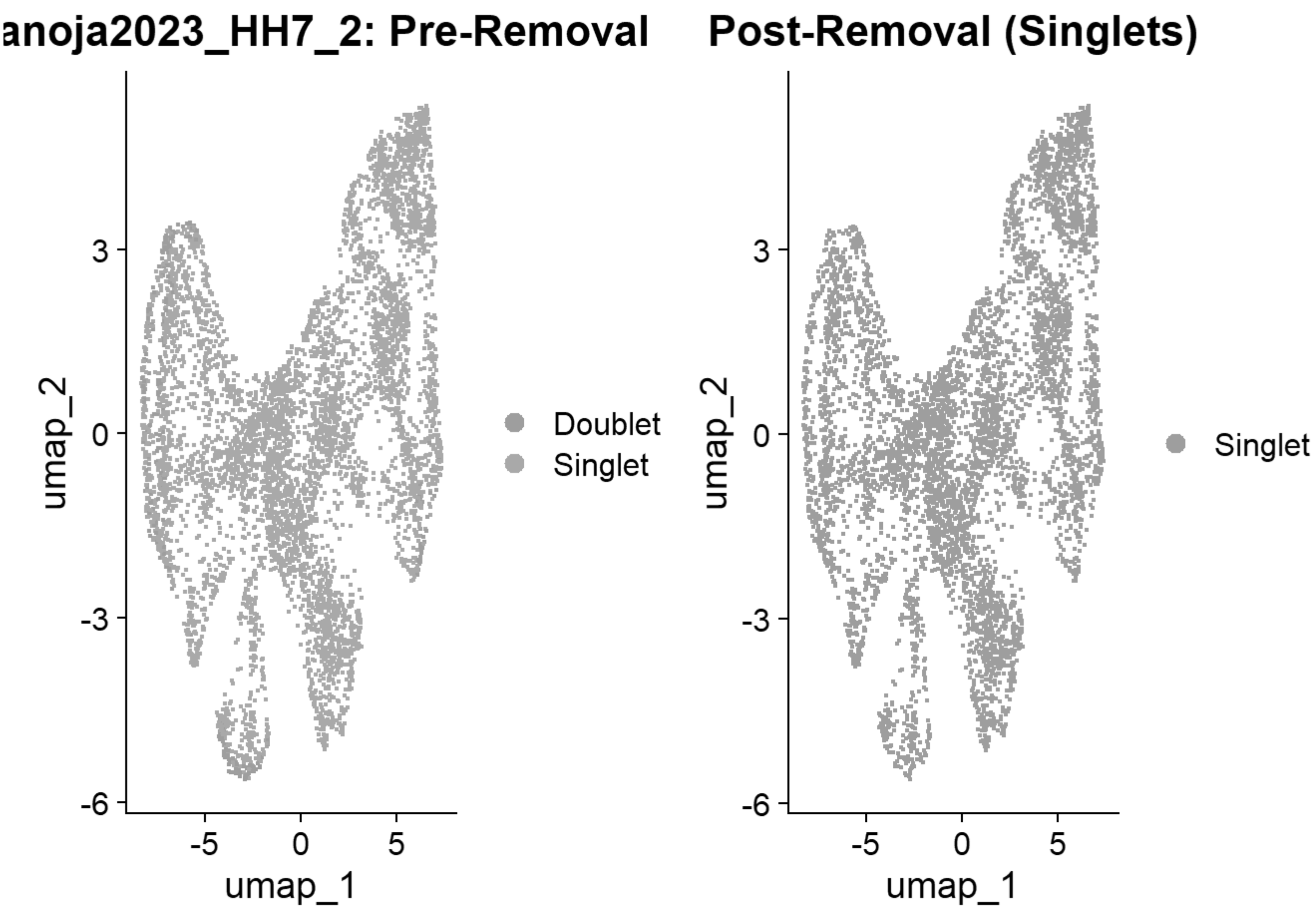


```
## [1] "Loading HH8_1"
## [1] "Running SoupX for HH8_1"
```

```
## Modularity Optimizer version 1.3.0 by Ludo Waltman and Nees Jan van Eck
##
## Number of nodes: 10012
## Number of edges: 344818
##
## Running Louvain algorithm...
## Maximum modularity in 10 random starts: 0.8910
## Number of communities: 12
## Elapsed time: 0 seconds
```

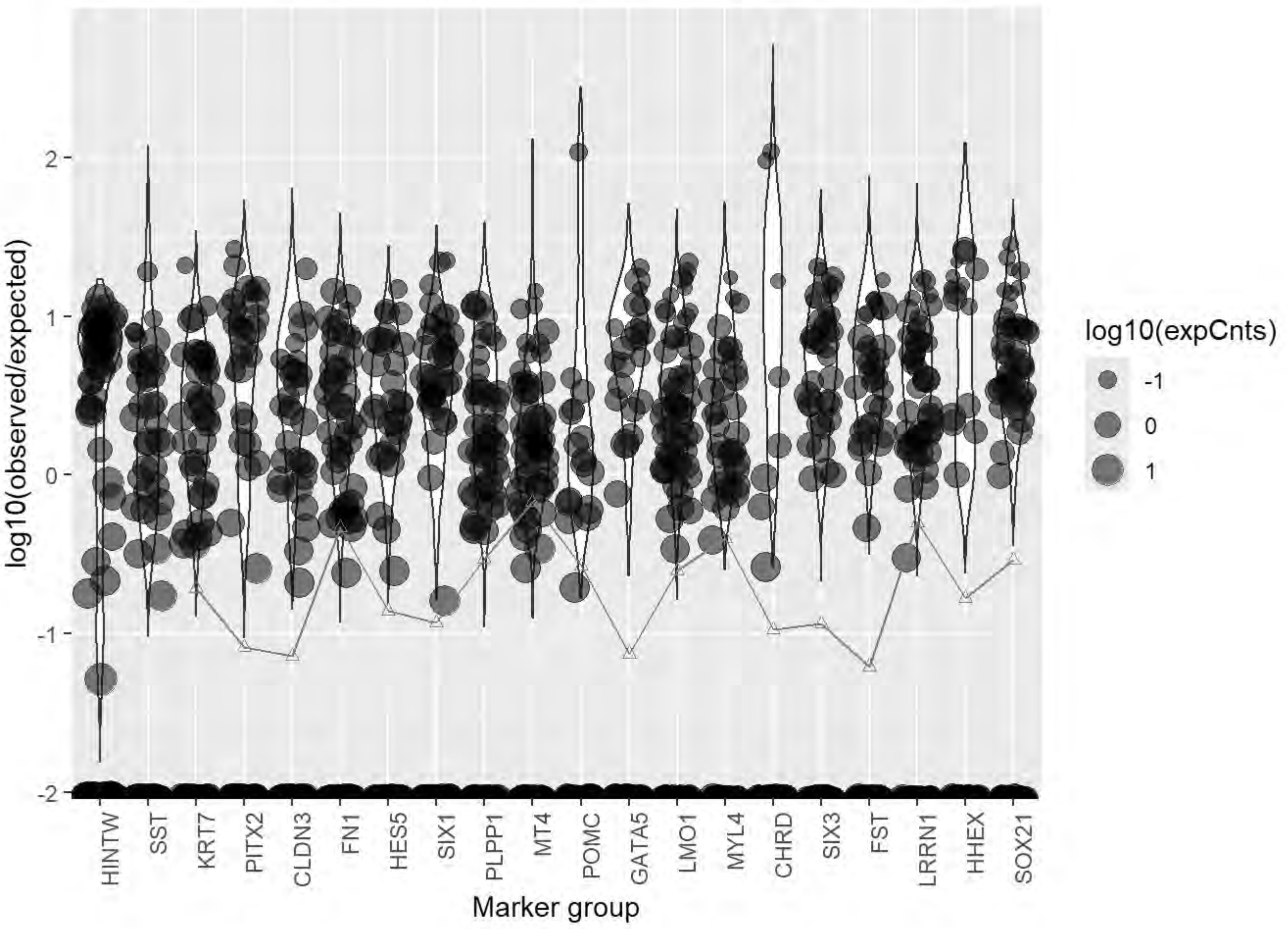
log10(observed/expected)
2
1
0
-1
-2
HINTW
SST
KRT7
PITX2
CLDN3
FN1
HES5
SIX1
PLPP1
MT4
POMC
GATA5
LMO1
MYL4
CHRD
SIX3
FST
LRRN1
HHEX
SOX21
Marker group
log10(expCnts)
-1
0
1

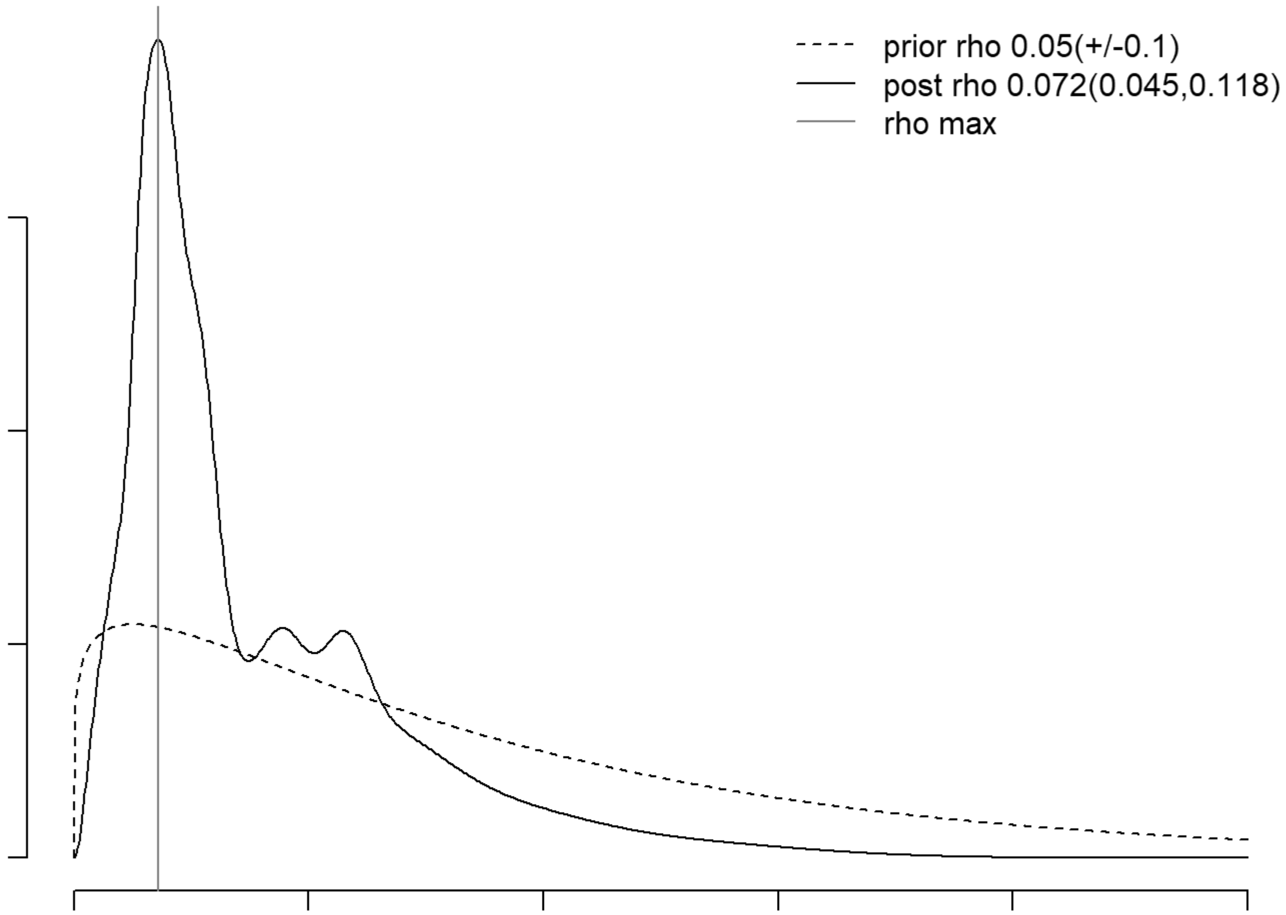
prior rho 0.05(+/-0.1)
post rho 0.072(0.045,0.118)
rho max

```
## [1] "Running QC for HH8_1"
## [1] "Running DoubletFinder for HH8_1"
## [1] "Creating artificial doublets for pN = 5%"
## [1] "Creating Seurat object..."
## [1] "Normalizing Seurat object..."
```

```
## [1] "Finding variable genes..."
```

```
## [1] "Scaling data..."
```

```
## [1] "Running PCA..."
## [1] "Calculating PC distance matrix..."
## [1] "Defining neighborhoods..."
## [1] "Computing pANN across all pK..."
## [1] "pK = 0.001..."
## [1] "pK = 0.005..."
## [1] "pK = 0.01..."
## [1] "pK = 0.02..."
## [1] "pK = 0.03..."
## [1] "pK = 0.04..."
## [1] "pK = 0.05..."
## [1] "pK = 0.06..."
## [1] "pK = 0.07..."
## [1] "pK = 0.08..."
## [1] "pK = 0.09..."
## [1] "pK = 0.1..."
## [1] "pK = 0.11..."
## [1] "pK = 0.12..."
## [1] "pK = 0.13..."
## [1] "pK = 0.14..."
## [1] "pK = 0.15..."
## [1] "pK = 0.16..."
## [1] "pK = 0.17..."
## [1] "pK = 0.18..."
## [1] "pK = 0.19..."
## [1] "pK = 0.2..."
## [1] "pK = 0.21..."
## [1] "pK = 0.22..."
## [1] "pK = 0.23..."
## [1] "pK = 0.24..."
## [1] "pK = 0.25..."
## [1] "pK = 0.26..."
## [1] "pK = 0.27..."
## [1] "pK = 0.28..."
## [1] "pK = 0.29..."
## [1] "pK = 0.3..."
## [1] "Creating artificial doublets for pN = 10%"
## [1] "Creating Seurat object..."
## [1] "Normalizing Seurat object..."
```

```
## [1] "Finding variable genes..."
```

```
## [1] "Scaling data..."
```

```
## [1] "Running PCA..."
## [1] "Calculating PC distance matrix..."
## [1] "Defining neighborhoods..."
## [1] "Computing pANN across all pK..."
## [1] "pK = 0.001..."
## [1] "pK = 0.005..."
## [1] "pK = 0.01..."
## [1] "pK = 0.02..."
## [1] "pK = 0.03..."
## [1] "pK = 0.04..."
## [1] "pK = 0.05..."
## [1] "pK = 0.06..."
## [1] "pK = 0.07..."
## [1] "pK = 0.08..."
## [1] "pK = 0.09..."
## [1] "pK = 0.1..."
## [1] "pK = 0.11..."
## [1] "pK = 0.12..."
## [1] "pK = 0.13..."
## [1] "pK = 0.14..."
## [1] "pK = 0.15..."
## [1] "pK = 0.16..."
## [1] "pK = 0.17..."
## [1] "pK = 0.18..."
## [1] "pK = 0.19..."
## [1] "pK = 0.2..."
## [1] "pK = 0.21..."
## [1] "pK = 0.22..."
## [1] "pK = 0.23..."
## [1] "pK = 0.24..."
## [1] "pK = 0.25..."
## [1] "pK = 0.26..."
## [1] "pK = 0.27..."
## [1] "pK = 0.28..."
## [1] "pK = 0.29..."
## [1] "pK = 0.3..."
## [1] "Creating artificial doublets for pN = 15%"
## [1] "Creating Seurat object..."
## [1] "Normalizing Seurat object..."
```

```
## [1] "Finding variable genes..."
```

```
## [1] "Scaling data..."
```

```
## [1] "Running PCA..."
## [1] "Calculating PC distance matrix..."
## [1] "Defining neighborhoods..."
```

```
## [1] "Computing pANN across all pK..."
## [1] "pK = 0.001..."
## [1] "pK = 0.005..."
## [1] "pK = 0.01..."
## [1] "pK = 0.02..."
## [1] "pK = 0.03..."
## [1] "pK = 0.04..."
## [1] "pK = 0.05..."
## [1] "pK = 0.06..."
## [1] "pK = 0.07..."
## [1] "pK = 0.08..."
## [1] "pK = 0.09..."
## [1] "pK = 0.1..."
## [1] "pK = 0.11..."
## [1] "pK = 0.12..."
## [1] "pK = 0.13..."
## [1] "pK = 0.14..."
## [1] "pK = 0.15..."
## [1] "pK = 0.16..."
## [1] "pK = 0.17..."
## [1] "pK = 0.18..."
## [1] "pK = 0.19..."
## [1] "pK = 0.2..."
## [1] "pK = 0.21..."
## [1] "pK = 0.22..."
## [1] "pK = 0.23..."
## [1] "pK = 0.24..."
## [1] "pK = 0.25..."
## [1] "pK = 0.26..."
## [1] "pK = 0.27..."
## [1] "pK = 0.28..."
## [1] "pK = 0.29..."
## [1] "pK = 0.3..."
## [1] "Creating artificial doublets for pN = 20%"
## [1] "Creating Seurat object..."
## [1] "Normalizing Seurat object..."
```

```
## [1] "Finding variable genes..."
```

```
## [1] "Scaling data..."
```

```
## [1] "Running PCA..."
## [1] "Calculating PC distance matrix..."
## [1] "Defining neighborhoods..."
## [1] "Computing pANN across all pK..."
## [1] "pK = 0.001..."
## [1] "pK = 0.005..."
## [1] "pK = 0.01..."
## [1] "pK = 0.02..."
## [1] "pK = 0.03..."
## [1] "pK = 0.04..."
## [1] "pK = 0.05..."
```

```
## [1] "pK = 0.06..."
## [1] "pK = 0.07..."
## [1] "pK = 0.08..."
## [1] "pK = 0.09..."
## [1] "pK = 0.1..."
## [1] "pK = 0.11..."
## [1] "pK = 0.12..."
## [1] "pK = 0.13..."
## [1] "pK = 0.14..."
## [1] "pK = 0.15..."
## [1] "pK = 0.16..."
## [1] "pK = 0.17..."
## [1] "pK = 0.18..."
## [1] "pK = 0.19..."
## [1] "pK = 0.2..."
## [1] "pK = 0.21..."
## [1] "pK = 0.22..."
## [1] "pK = 0.23..."
## [1] "pK = 0.24..."
## [1] "pK = 0.25..."
## [1] "pK = 0.26..."
## [1] "pK = 0.27..."
## [1] "pK = 0.28..."
## [1] "pK = 0.29..."
## [1] "pK = 0.3..."
## [1] "Creating artificial doublets for pN = 25%"
## [1] "Creating Seurat object..."
## [1] "Normalizing Seurat object..."
```

```
## [1] "Finding variable genes..."
```

```
## [1] "Scaling data..."
```

```
## [1] "Running PCA..."
## [1] "Calculating PC distance matrix..."
## [1] "Defining neighborhoods..."
## [1] "Computing pANN across all pK..."
## [1] "pK = 0.001..."
## [1] "pK = 0.005..."
## [1] "pK = 0.01..."
## [1] "pK = 0.02..."
## [1] "pK = 0.03..."
## [1] "pK = 0.04..."
## [1] "pK = 0.05..."
## [1] "pK = 0.06..."
## [1] "pK = 0.07..."
## [1] "pK = 0.08..."
## [1] "pK = 0.09..."
## [1] "pK = 0.1..."
## [1] "pK = 0.11..."
## [1] "pK = 0.12..."
## [1] "pK = 0.13..."
```

```
## [1] "pK = 0.14..."
## [1] "pK = 0.15..."
## [1] "pK = 0.16..."
## [1] "pK = 0.17..."
## [1] "pK = 0.18..."
## [1] "pK = 0.19..."
## [1] "pK = 0.2..."
## [1] "pK = 0.21..."
## [1] "pK = 0.22..."
## [1] "pK = 0.23..."
## [1] "pK = 0.24..."
## [1] "pK = 0.25..."
## [1] "pK = 0.26..."
## [1] "pK = 0.27..."
## [1] "pK = 0.28..."
## [1] "pK = 0.29..."
## [1] "pK = 0.3..."
## [1] "Creating artificial doublets for pN = 30%"
## [1] "Creating Seurat object..."
## [1] "Normalizing Seurat object..."
```

```
## [1] "Finding variable genes..."
```

```
## [1] "Scaling data..."
```

```
## [1] "Running PCA..."
## [1] "Calculating PC distance matrix..."
## [1] "Defining neighborhoods..."
## [1] "Computing pANN across all pK..."
## [1] "pK = 0.001..."
## [1] "pK = 0.005..."
## [1] "pK = 0.01..."
## [1] "pK = 0.02..."
## [1] "pK = 0.03..."
## [1] "pK = 0.04..."
## [1] "pK = 0.05..."
## [1] "pK = 0.06..."
## [1] "pK = 0.07..."
## [1] "pK = 0.08..."
## [1] "pK = 0.09..."
## [1] "pK = 0.1..."
## [1] "pK = 0.11..."
## [1] "pK = 0.12..."
## [1] "pK = 0.13..."
## [1] "pK = 0.14..."
## [1] "pK = 0.15..."
## [1] "pK = 0.16..."
## [1] "pK = 0.17..."
## [1] "pK = 0.18..."
## [1] "pK = 0.19..."
## [1] "pK = 0.2..."
## [1] "pK = 0.21..."
```

```
## [1] "pK = 0.22..."
## [1] "pK = 0.23..."
## [1] "pK = 0.24..."
## [1] "pK = 0.25..."
## [1] "pK = 0.26..."
## [1] "pK = 0.27..."
## [1] "pK = 0.28..."
## [1] "pK = 0.29..."
## [1] "pK = 0.3..."
```

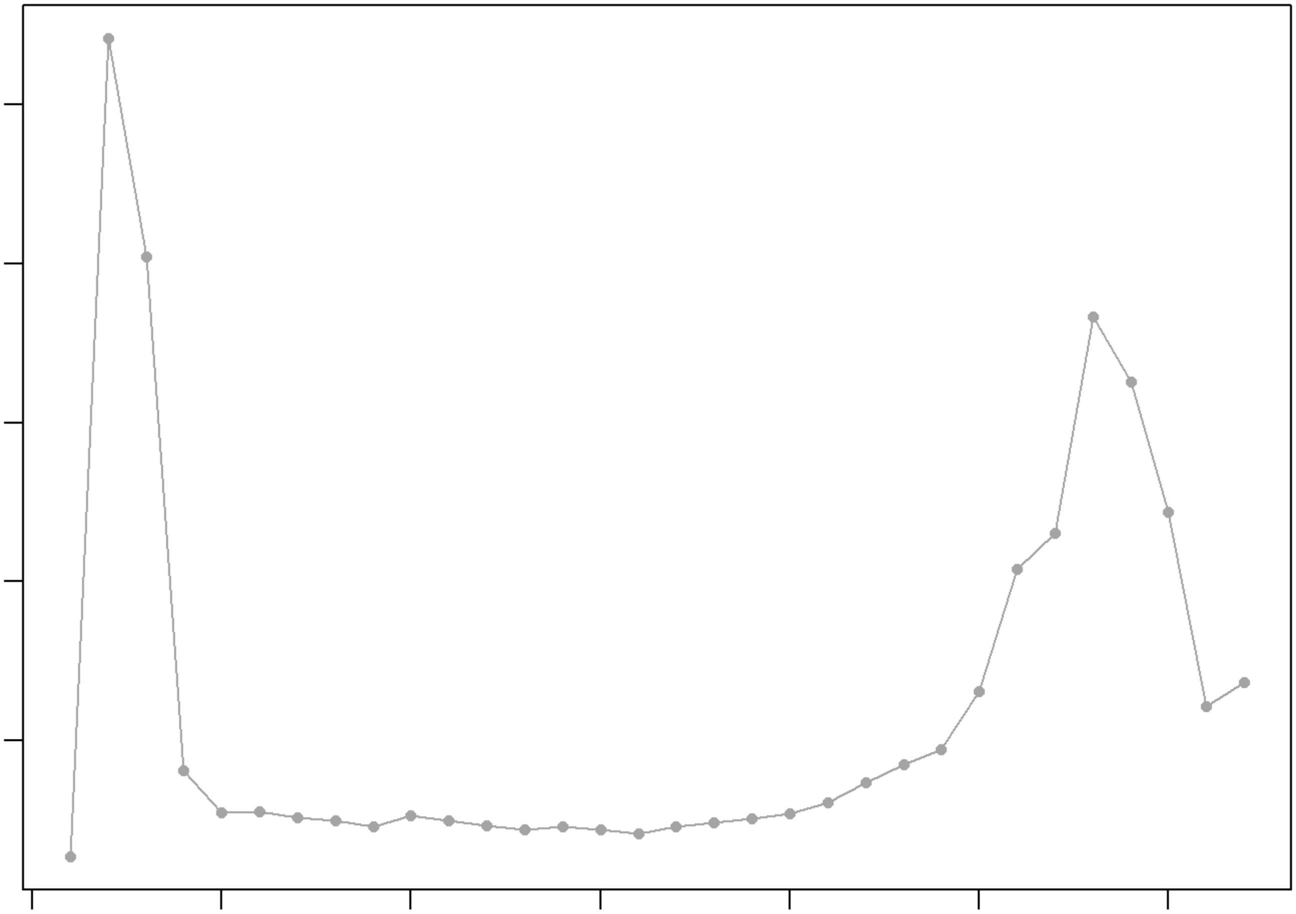

```
## NULL
## [1] "Creating 3261 artificial doublets..."
## [1] "Creating Seurat object..."
## [1] "Normalizing Seurat object..."
```

```
## [1] "Finding variable genes..."
```

```
## [1] "Scaling data..."
```

```
## [1] "Running PCA..."
## [1] "Calculating PC distance matrix..."
## [1] "Computing pANN..."
## [1] "Classifying doublets.."
```

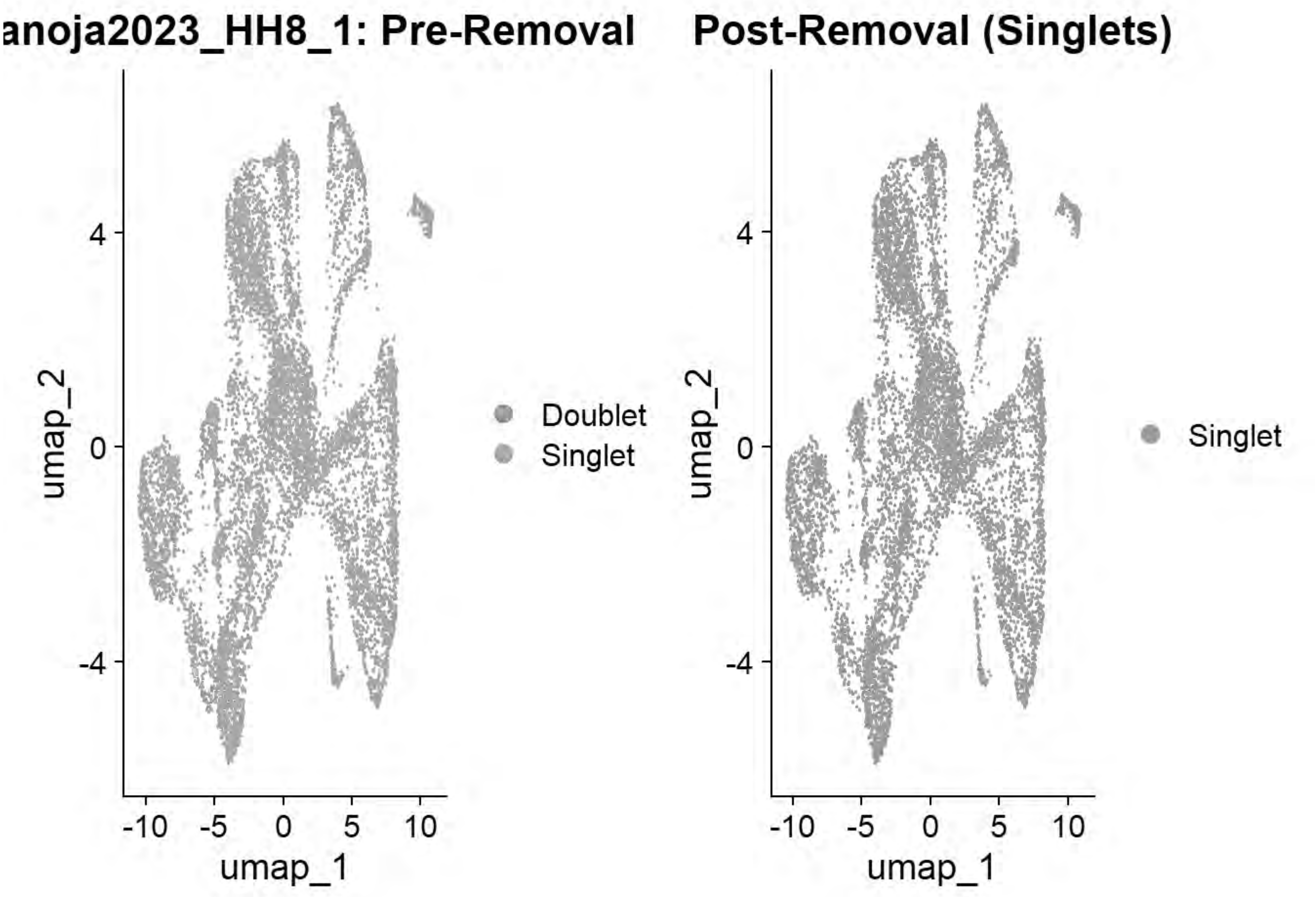


```
## [1] "Loading HH8_2"
## [1] "Running SoupX for HH8_2"
```

```
## Modularity Optimizer version 1.3.0 by Ludo Waltman and Nees Jan van Eck
##
## Number of nodes: 23489
## Number of edges: 681856
##
## Running Louvain algorithm...
## Maximum modularity in 10 random starts: 0.8741
## Number of communities: 12
## Elapsed time: 2 seconds
```

```
## [1] "Running QC for HH8_2"
## [1] "Running DoubletFinder for HH8_2"
## [1] "Creating artificial doublets for pN = 5%"
## [1] "Creating Seurat object..."
## [1] "Normalizing Seurat object..."
```

```
## [1] "Finding variable genes..."
```

```
## [1] "Scaling data..."
```

```
## [1] "Running PCA..."
## [1] "Calculating PC distance matrix..."
## [1] "Defining neighborhoods..."
## [1] "Computing pANN across all pK..."
## [1] "pK = 5e-04..."
## [1] "pK = 0.001..."
## [1] "pK = 0.005..."
## [1] "pK = 0.01..."
## [1] "pK = 0.02..."
## [1] "pK = 0.03..."
## [1] "pK = 0.04..."
## [1] "pK = 0.05..."
## [1] "pK = 0.06..."
## [1] "pK = 0.07..."
## [1] "pK = 0.08..."
## [1] "pK = 0.09..."
## [1] "pK = 0.1..."
## [1] "pK = 0.11..."
## [1] "pK = 0.12..."
## [1] "pK = 0.13..."
## [1] "pK = 0.14..."
## [1] "pK = 0.15..."
## [1] "pK = 0.16..."
## [1] "pK = 0.17..."
## [1] "pK = 0.18..."
## [1] "pK = 0.19..."
## [1] "pK = 0.2..."
## [1] "pK = 0.21..."
## [1] "pK = 0.22..."
## [1] "pK = 0.23..."
## [1] "pK = 0.24..."
## [1] "pK = 0.25..."
## [1] "pK = 0.26..."
## [1] "pK = 0.27..."
## [1] "pK = 0.28..."
## [1] "pK = 0.29..."
## [1] "pK = 0.3..."
## [1] "Creating artificial doublets for pN = 10%"
## [1] "Creating Seurat object..."
## [1] "Normalizing Seurat object..."
```

```
## [1] "Finding variable genes..."
```

```
## [1] "Scaling data..."
```

```
## [1] "Running PCA..."
## [1] "Calculating PC distance matrix..."
## [1] "Defining neighborhoods..."
## [1] "Computing pANN across all pK..."
## [1] "pK = 5e-04..."
## [1] "pK = 0.001..."
## [1] "pK = 0.005..."
```

```
## [1] "pK = 0.01..."
## [1] "pK = 0.02..."
## [1] "pK = 0.03..."
## [1] "pK = 0.04..."
## [1] "pK = 0.05..."
## [1] "pK = 0.06..."
## [1] "pK = 0.07..."
## [1] "pK = 0.08..."
## [1] "pK = 0.09..."
## [1] "pK = 0.1..."
## [1] "pK = 0.11..."
## [1] "pK = 0.12..."
## [1] "pK = 0.13..."
## [1] "pK = 0.14..."
## [1] "pK = 0.15..."
## [1] "pK = 0.16..."
## [1] "pK = 0.17..."
## [1] "pK = 0.18..."
## [1] "pK = 0.19..."
## [1] "pK = 0.2..."
## [1] "pK = 0.21..."
## [1] "pK = 0.22..."
## [1] "pK = 0.23..."
## [1] "pK = 0.24..."
## [1] "pK = 0.25..."
## [1] "pK = 0.26..."
## [1] "pK = 0.27..."
## [1] "pK = 0.28..."
## [1] "pK = 0.29..."
## [1] "pK = 0.3..."
## [1] "Creating artificial doublets for pN = 15%"
## [1] "Creating Seurat object..."
## [1] "Normalizing Seurat object..."
```

```
## [1] "Finding variable genes..."
```

```
## [1] "Scaling data..."
```

```
## [1] "Running PCA..."
## [1] "Calculating PC distance matrix..."
## [1] "Defining neighborhoods..."
## [1] "Computing pANN across all pK..."
## [1] "pK = 5e-04..."
## [1] "pK = 0.001..."
## [1] "pK = 0.005..."
## [1] "pK = 0.01..."
## [1] "pK = 0.02..."
## [1] "pK = 0.03..."
## [1] "pK = 0.04..."
## [1] "pK = 0.05..."
## [1] "pK = 0.06..."
## [1] "pK = 0.07..."
```

```
## [1] "pK = 0.08..."
## [1] "pK = 0.09..."
## [1] "pK = 0.1..."
## [1] "pK = 0.11..."
## [1] "pK = 0.12..."
## [1] "pK = 0.13..."
## [1] "pK = 0.14..."
## [1] "pK = 0.15..."
## [1] "pK = 0.16..."
## [1] "pK = 0.17..."
## [1] "pK = 0.18..."
## [1] "pK = 0.19..."
## [1] "pK = 0.2..."
## [1] "pK = 0.21..."
## [1] "pK = 0.22..."
## [1] "pK = 0.23..."
## [1] "pK = 0.24..."
## [1] "pK = 0.25..."
## [1] "pK = 0.26..."
## [1] "pK = 0.27..."
## [1] "pK = 0.28..."
## [1] "pK = 0.29..."
## [1] "pK = 0.3..."
## [1] "Creating artificial doublets for pN = 20%"
## [1] "Creating Seurat object..."
## [1] "Normalizing Seurat object..."
```

```
## [1] "Finding variable genes..."
```

```
## [1] "Scaling data..."
```

```
## [1] "Running PCA..."
## [1] "Calculating PC distance matrix..."
## [1] "Defining neighborhoods..."
## [1] "Computing pANN across all pK..."
## [1] "pK = 5e-04..."
## [1] "pK = 0.001..."
## [1] "pK = 0.005..."
## [1] "pK = 0.01..."
## [1] "pK = 0.02..."
## [1] "pK = 0.03..."
## [1] "pK = 0.04..."
## [1] "pK = 0.05..."
## [1] "pK = 0.06..."
## [1] "pK = 0.07..."
## [1] "pK = 0.08..."
## [1] "pK = 0.09..."
## [1] "pK = 0.1..."
## [1] "pK = 0.11..."
## [1] "pK = 0.12..."
## [1] "pK = 0.13..."
## [1] "pK = 0.14..."
```

```
## [1] "pK = 0.15..."
## [1] "pK = 0.16..."
## [1] "pK = 0.17..."
## [1] "pK = 0.18..."
## [1] "pK = 0.19..."
## [1] "pK = 0.2..."
## [1] "pK = 0.21..."
## [1] "pK = 0.22..."
## [1] "pK = 0.23..."
## [1] "pK = 0.24..."
## [1] "pK = 0.25..."
## [1] "pK = 0.26..."
## [1] "pK = 0.27..."
## [1] "pK = 0.28..."
## [1] "pK = 0.29..."
## [1] "pK = 0.3..."
## [1] "Creating artificial doublets for pN = 25%"
## [1] "Creating Seurat object..."
## [1] "Normalizing Seurat object..."
```

```
## [1] "Finding variable genes..."
```

```
## [1] "Scaling data..."
```

```
## [1] "Running PCA..."
## [1] "Calculating PC distance matrix..."
## [1] "Defining neighborhoods..."
## [1] "Computing pANN across all pK..."
## [1] "pK = 5e-04..."
## [1] "pK = 0.001..."
## [1] "pK = 0.005..."
## [1] "pK = 0.01..."
## [1] "pK = 0.02..."
## [1] "pK = 0.03..."
## [1] "pK = 0.04..."
## [1] "pK = 0.05..."
## [1] "pK = 0.06..."
## [1] "pK = 0.07..."
## [1] "pK = 0.08..."
## [1] "pK = 0.09..."
## [1] "pK = 0.1..."
## [1] "pK = 0.11..."
## [1] "pK = 0.12..."
## [1] "pK = 0.13..."
## [1] "pK = 0.14..."
## [1] "pK = 0.15..."
## [1] "pK = 0.16..."
## [1] "pK = 0.17..."
## [1] "pK = 0.18..."
## [1] "pK = 0.19..."
## [1] "pK = 0.2..."
## [1] "pK = 0.21..."
```

```
## [1] "pK = 0.22..."
## [1] "pK = 0.23..."
## [1] "pK = 0.24..."
## [1] "pK = 0.25..."
## [1] "pK = 0.26..."
## [1] "pK = 0.27..."
## [1] "pK = 0.28..."
## [1] "pK = 0.29..."
## [1] "pK = 0.3..."
## [1] "Creating artificial doublets for pN = 30%"
## [1] "Creating Seurat object..."
## [1] "Normalizing Seurat object..."
```

```
## [1] "Finding variable genes..."
```

```
## [1] "Scaling data..."
```

```
## [1] "Running PCA..."
## [1] "Calculating PC distance matrix..."
## [1] "Defining neighborhoods..."
## [1] "Computing pANN across all pK..."
## [1] "pK = 5e-04..."
## [1] "pK = 0.001..."
## [1] "pK = 0.005..."
## [1] "pK = 0.01..."
## [1] "pK = 0.02..."
## [1] "pK = 0.03..."
## [1] "pK = 0.04..."
## [1] "pK = 0.05..."
## [1] "pK = 0.06..."
## [1] "pK = 0.07..."
## [1] "pK = 0.08..."
## [1] "pK = 0.09..."
## [1] "pK = 0.1..."
## [1] "pK = 0.11..."
## [1] "pK = 0.12..."
## [1] "pK = 0.13..."
## [1] "pK = 0.14..."
## [1] "pK = 0.15..."
## [1] "pK = 0.16..."
## [1] "pK = 0.17..."
## [1] "pK = 0.18..."
## [1] "pK = 0.19..."
## [1] "pK = 0.2..."
## [1] "pK = 0.21..."
## [1] "pK = 0.22..."
## [1] "pK = 0.23..."
## [1] "pK = 0.24..."
## [1] "pK = 0.25..."
## [1] "pK = 0.26..."
## [1] "pK = 0.27..."
## [1] "pK = 0.28..."
```

```
## [1] "pK = 0.29..."
## [1] "pK = 0.3..."
```

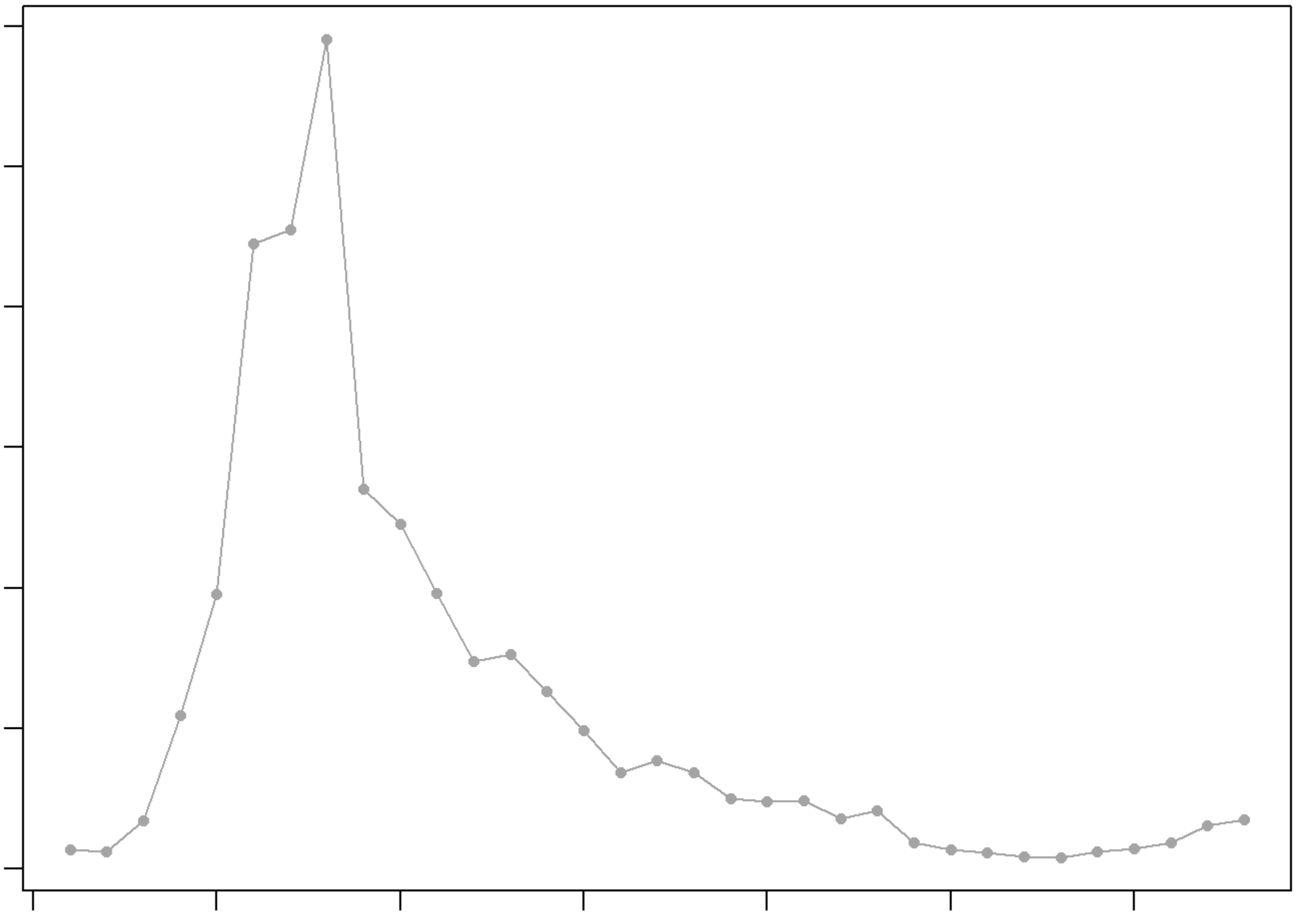

```
## NULL
## [1] "Creating 7563 artificial doublets..."
## [1] "Creating Seurat object..."
## [1] "Normalizing Seurat object..."
```

```
## [1] "Finding variable genes..."
```

```
## [1] "Scaling data..."
```

```
## [1] "Running PCA..."
## [1] "Calculating PC distance matrix..."
## [1] "Computing pANN..."
## [1] "Classifying doublets.."
```

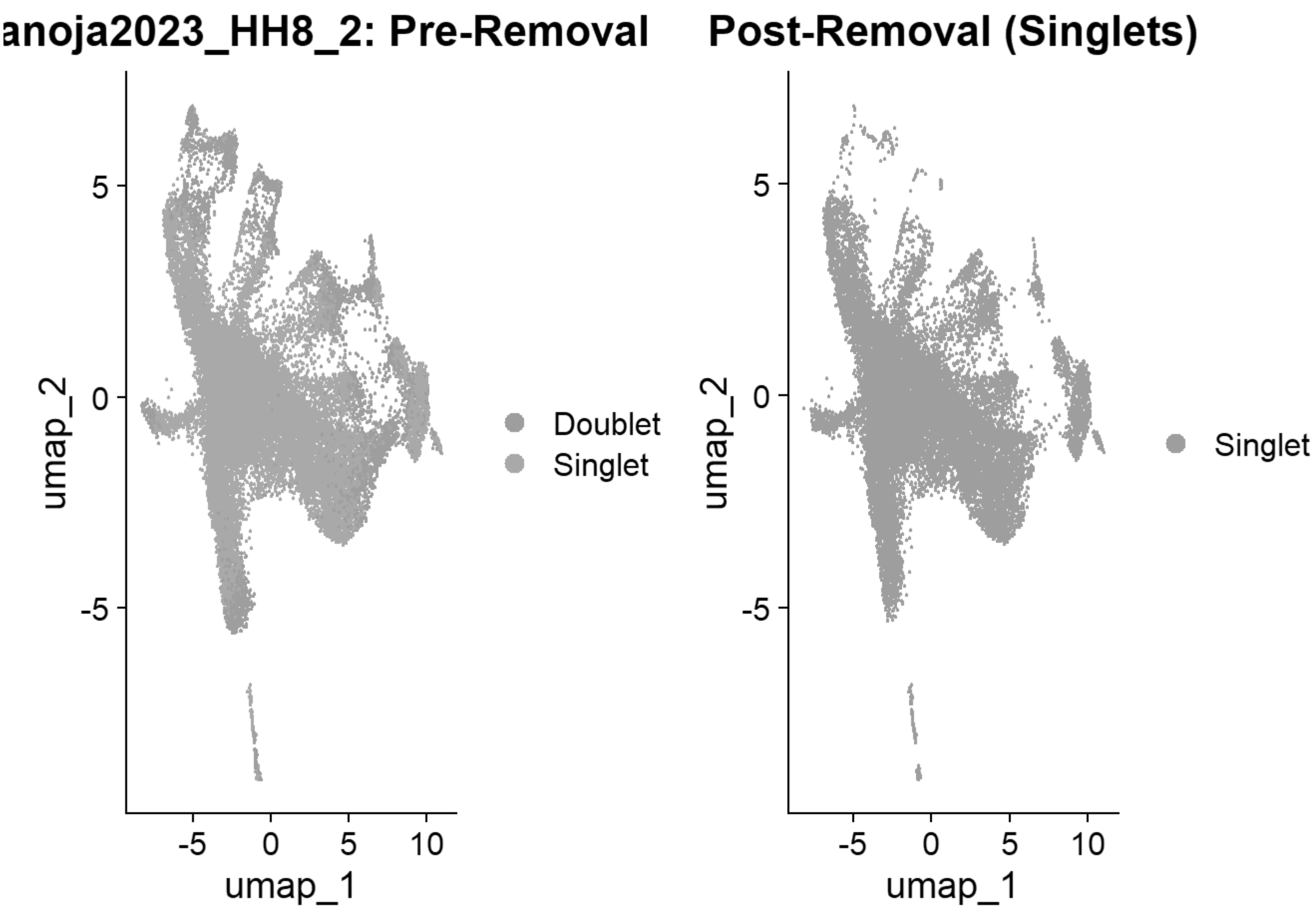


```
## [1] "Loading HH9_1"
## [1] "Running SoupX for HH9_1"
```

```
## Modularity Optimizer version 1.3.0 by Ludo Waltman and Nees Jan van Eck
##
## Number of nodes: 16839
## Number of edges: 531221
##
## Running Louvain algorithm...
## Maximum modularity in 10 random starts: 0.8824
## Number of communities: 18
## Elapsed time: 2 seconds
```

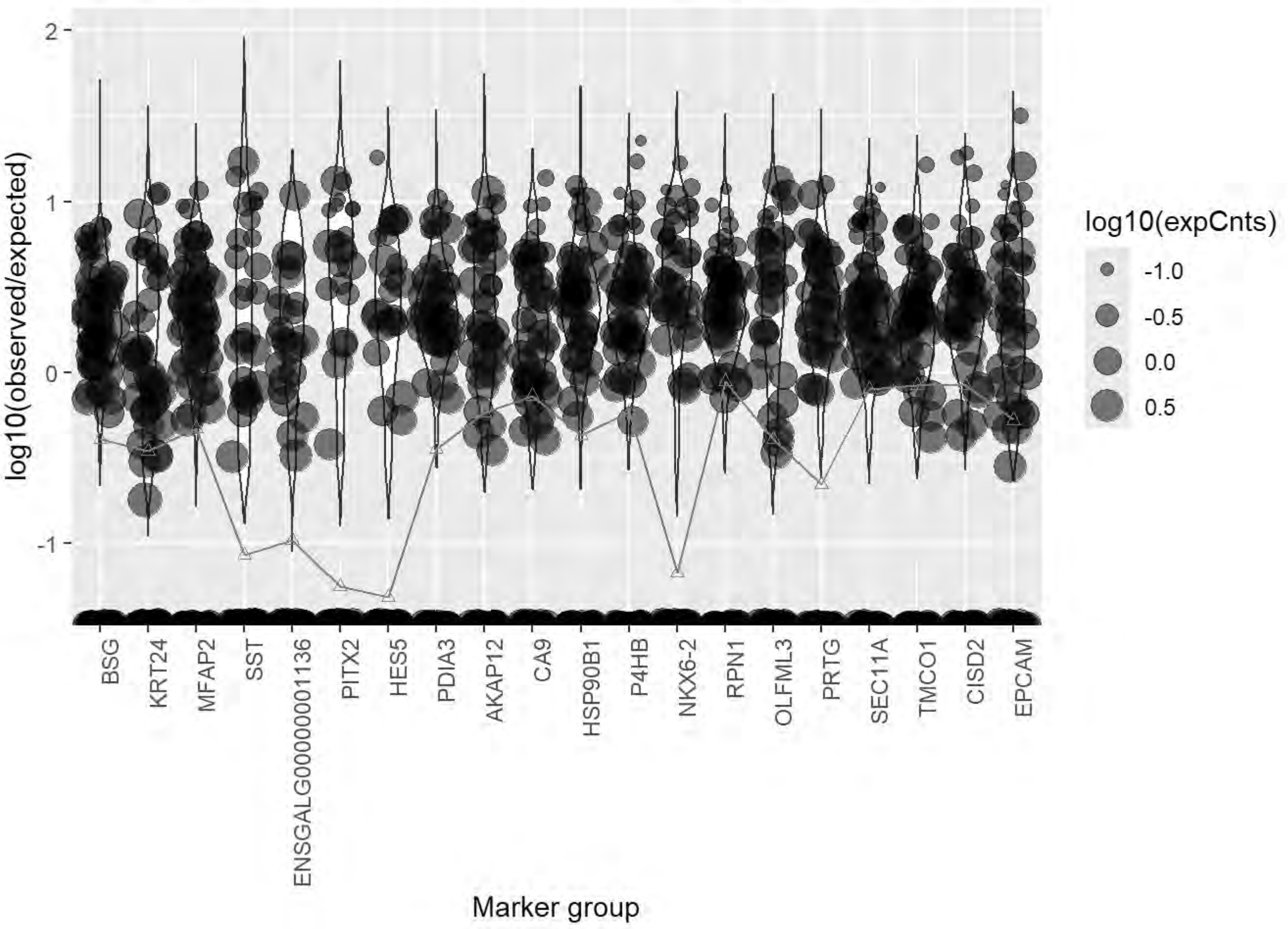
log10(observed/expected)
2
1
0
-1
BSG
KRT24
MFAP2
SST
ENSGALG00000001136
PITX2
HES5
PDIA3
AKAP12
CA9
HSP90B1
P4HB
NKX6-2
RPN1
OLFML3
PRTG
SEC11A
TMCO1
CISD2
EPCAM
Marker group
log10(expCnts)
-1.0
-0.5
0.0
0.5

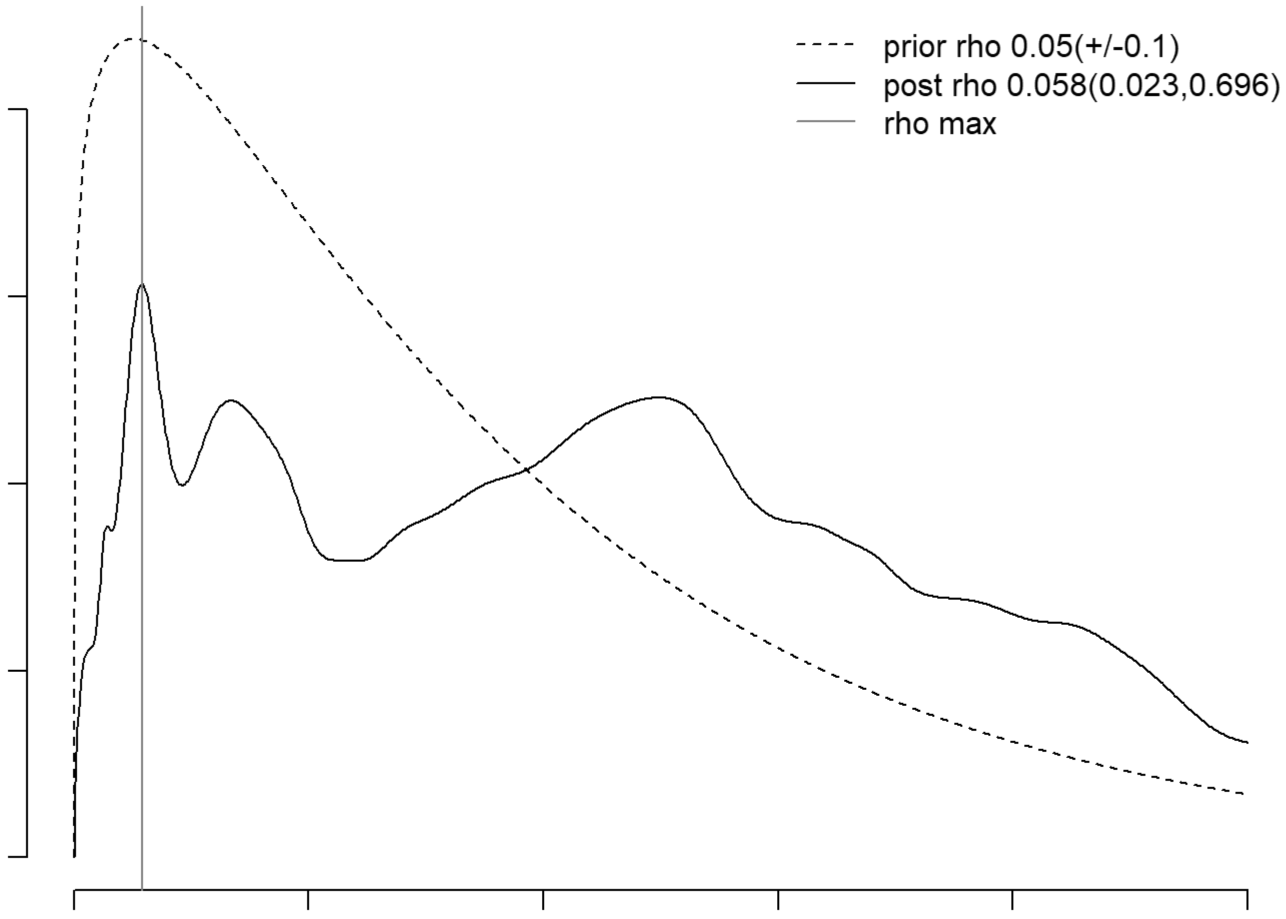
prior rho 0.05(+/-0.1)
post rho 0.058(0.023,0.696)
rho max

```
## [1] "Running QC for HH9_1"
## [1] "Running DoubletFinder for HH9_1"
## [1] "Creating artificial doublets for pN = 5%"
## [1] "Creating Seurat object..."
## [1] "Normalizing Seurat object..."
```

```
## [1] "Finding variable genes..."
```

```
## [1] "Scaling data..."
```

```
## [1] "Running PCA..."
## [1] "Calculating PC distance matrix..."
## [1] "Defining neighborhoods..."
## [1] "Computing pANN across all pK..."
## [1] "pK = 0.001..."
## [1] "pK = 0.005..."
## [1] "pK = 0.01..."
## [1] "pK = 0.02..."
## [1] "pK = 0.03..."
## [1] "pK = 0.04..."
## [1] "pK = 0.05..."
## [1] "pK = 0.06..."
## [1] "pK = 0.07..."
## [1] "pK = 0.08..."
## [1] "pK = 0.09..."
## [1] "pK = 0.1..."
## [1] "pK = 0.11..."
## [1] "pK = 0.12..."
## [1] "pK = 0.13..."
## [1] "pK = 0.14..."
## [1] "pK = 0.15..."
## [1] "pK = 0.16..."
## [1] "pK = 0.17..."
## [1] "pK = 0.18..."
## [1] "pK = 0.19..."
## [1] "pK = 0.2..."
## [1] "pK = 0.21..."
## [1] "pK = 0.22..."
## [1] "pK = 0.23..."
## [1] "pK = 0.24..."
## [1] "pK = 0.25..."
## [1] "pK = 0.26..."
## [1] "pK = 0.27..."
## [1] "pK = 0.28..."
## [1] "pK = 0.29..."
## [1] "pK = 0.3..."
## [1] "Creating artificial doublets for pN = 10%"
## [1] "Creating Seurat object..."
## [1] "Normalizing Seurat object..."
```

```
## [1] "Finding variable genes..."
```

```
## [1] "Scaling data..."
```

```
## [1] "Running PCA..."
## [1] "Calculating PC distance matrix..."
## [1] "Defining neighborhoods..."
## [1] "Computing pANN across all pK..."
## [1] "pK = 0.001..."
## [1] "pK = 0.005..."
## [1] "pK = 0.01..."
## [1] "pK = 0.02..."
## [1] "pK = 0.03..."
## [1] "pK = 0.04..."
## [1] "pK = 0.05..."
## [1] "pK = 0.06..."
## [1] "pK = 0.07..."
## [1] "pK = 0.08..."
## [1] "pK = 0.09..."
## [1] "pK = 0.1..."
## [1] "pK = 0.11..."
## [1] "pK = 0.12..."
## [1] "pK = 0.13..."
## [1] "pK = 0.14..."
## [1] "pK = 0.15..."
## [1] "pK = 0.16..."
## [1] "pK = 0.17..."
## [1] "pK = 0.18..."
## [1] "pK = 0.19..."
## [1] "pK = 0.2..."
## [1] "pK = 0.21..."
## [1] "pK = 0.22..."
## [1] "pK = 0.23..."
## [1] "pK = 0.24..."
## [1] "pK = 0.25..."
## [1] "pK = 0.26..."
## [1] "pK = 0.27..."
## [1] "pK = 0.28..."
## [1] "pK = 0.29..."
## [1] "pK = 0.3..."
## [1] "Creating artificial doublets for pN = 15%"
## [1] "Creating Seurat object..."
## [1] "Normalizing Seurat object..."
```

```
## [1] "Finding variable genes..."
```

```
## [1] "Scaling data..."
```

```
## [1] "Running PCA..."
## [1] "Calculating PC distance matrix..."
## [1] "Defining neighborhoods..."
```

```
## [1] "Computing pANN across all pK..."
## [1] "pK = 0.001..."
## [1] "pK = 0.005..."
## [1] "pK = 0.01..."
## [1] "pK = 0.02..."
## [1] "pK = 0.03..."
## [1] "pK = 0.04..."
## [1] "pK = 0.05..."
## [1] "pK = 0.06..."
## [1] "pK = 0.07..."
## [1] "pK = 0.08..."
## [1] "pK = 0.09..."
## [1] "pK = 0.1..."
## [1] "pK = 0.11..."
## [1] "pK = 0.12..."
## [1] "pK = 0.13..."
## [1] "pK = 0.14..."
## [1] "pK = 0.15..."
## [1] "pK = 0.16..."
## [1] "pK = 0.17..."
## [1] "pK = 0.18..."
## [1] "pK = 0.19..."
## [1] "pK = 0.2..."
## [1] "pK = 0.21..."
## [1] "pK = 0.22..."
## [1] "pK = 0.23..."
## [1] "pK = 0.24..."
## [1] "pK = 0.25..."
## [1] "pK = 0.26..."
## [1] "pK = 0.27..."
## [1] "pK = 0.28..."
## [1] "pK = 0.29..."
## [1] "pK = 0.3..."
## [1] "Creating artificial doublets for pN = 20%"
## [1] "Creating Seurat object..."
## [1] "Normalizing Seurat object..."
```

```
## [1] "Finding variable genes..."
```

```
## [1] "Scaling data..."
```

```
## [1] "Running PCA..."
## [1] "Calculating PC distance matrix..."
## [1] "Defining neighborhoods..."
## [1] "Computing pANN across all pK..."
## [1] "pK = 0.001..."
## [1] "pK = 0.005..."
## [1] "pK = 0.01..."
## [1] "pK = 0.02..."
## [1] "pK = 0.03..."
## [1] "pK = 0.04..."
## [1] "pK = 0.05..."
```

```
## [1] "pK = 0.06..."
## [1] "pK = 0.07..."
## [1] "pK = 0.08..."
## [1] "pK = 0.09..."
## [1] "pK = 0.1..."
## [1] "pK = 0.11..."
## [1] "pK = 0.12..."
## [1] "pK = 0.13..."
## [1] "pK = 0.14..."
## [1] "pK = 0.15..."
## [1] "pK = 0.16..."
## [1] "pK = 0.17..."
## [1] "pK = 0.18..."
## [1] "pK = 0.19..."
## [1] "pK = 0.2..."
## [1] "pK = 0.21..."
## [1] "pK = 0.22..."
## [1] "pK = 0.23..."
## [1] "pK = 0.24..."
## [1] "pK = 0.25..."
## [1] "pK = 0.26..."
## [1] "pK = 0.27..."
## [1] "pK = 0.28..."
## [1] "pK = 0.29..."
## [1] "pK = 0.3..."
## [1] "Creating artificial doublets for pN = 25%"
## [1] "Creating Seurat object..."
## [1] "Normalizing Seurat object..."
```

```
## [1] "Finding variable genes..."
```

```
## [1] "Scaling data..."
```

```
## [1] "Running PCA..."
## [1] "Calculating PC distance matrix..."
## [1] "Defining neighborhoods..."
## [1] "Computing pANN across all pK..."
## [1] "pK = 0.001..."
## [1] "pK = 0.005..."
## [1] "pK = 0.01..."
## [1] "pK = 0.02..."
## [1] "pK = 0.03..."
## [1] "pK = 0.04..."
## [1] "pK = 0.05..."
## [1] "pK = 0.06..."
## [1] "pK = 0.07..."
## [1] "pK = 0.08..."
## [1] "pK = 0.09..."
## [1] "pK = 0.1..."
## [1] "pK = 0.11..."
## [1] "pK = 0.12..."
## [1] "pK = 0.13..."
```

```
## [1] "pK = 0.14..."
## [1] "pK = 0.15..."
## [1] "pK = 0.16..."
## [1] "pK = 0.17..."
## [1] "pK = 0.18..."
## [1] "pK = 0.19..."
## [1] "pK = 0.2..."
## [1] "pK = 0.21..."
## [1] "pK = 0.22..."
## [1] "pK = 0.23..."
## [1] "pK = 0.24..."
## [1] "pK = 0.25..."
## [1] "pK = 0.26..."
## [1] "pK = 0.27..."
## [1] "pK = 0.28..."
## [1] "pK = 0.29..."
## [1] "pK = 0.3..."
## [1] "Creating artificial doublets for pN = 30%"
## [1] "Creating Seurat object..."
## [1] "Normalizing Seurat object..."
```

```
## [1] "Finding variable genes..."
```

```
## [1] "Scaling data..."
```

```
## [1] "Running PCA..."
## [1] "Calculating PC distance matrix..."
## [1] "Defining neighborhoods..."
## [1] "Computing pANN across all pK..."
## [1] "pK = 0.001..."
## [1] "pK = 0.005..."
## [1] "pK = 0.01..."
## [1] "pK = 0.02..."
## [1] "pK = 0.03..."
## [1] "pK = 0.04..."
## [1] "pK = 0.05..."
## [1] "pK = 0.06..."
## [1] "pK = 0.07..."
## [1] "pK = 0.08..."
## [1] "pK = 0.09..."
## [1] "pK = 0.1..."
## [1] "pK = 0.11..."
## [1] "pK = 0.12..."
## [1] "pK = 0.13..."
## [1] "pK = 0.14..."
## [1] "pK = 0.15..."
## [1] "pK = 0.16..."
## [1] "pK = 0.17..."
## [1] "pK = 0.18..."
## [1] "pK = 0.19..."
## [1] "pK = 0.2..."
## [1] "pK = 0.21..."
```

```
## [1] "pK = 0.22..."
## [1] "pK = 0.23..."
## [1] "pK = 0.24..."
## [1] "pK = 0.25..."
## [1] "pK = 0.26..."
## [1] "pK = 0.27..."
## [1] "pK = 0.28..."
## [1] "pK = 0.29..."
## [1] "pK = 0.3..."
```

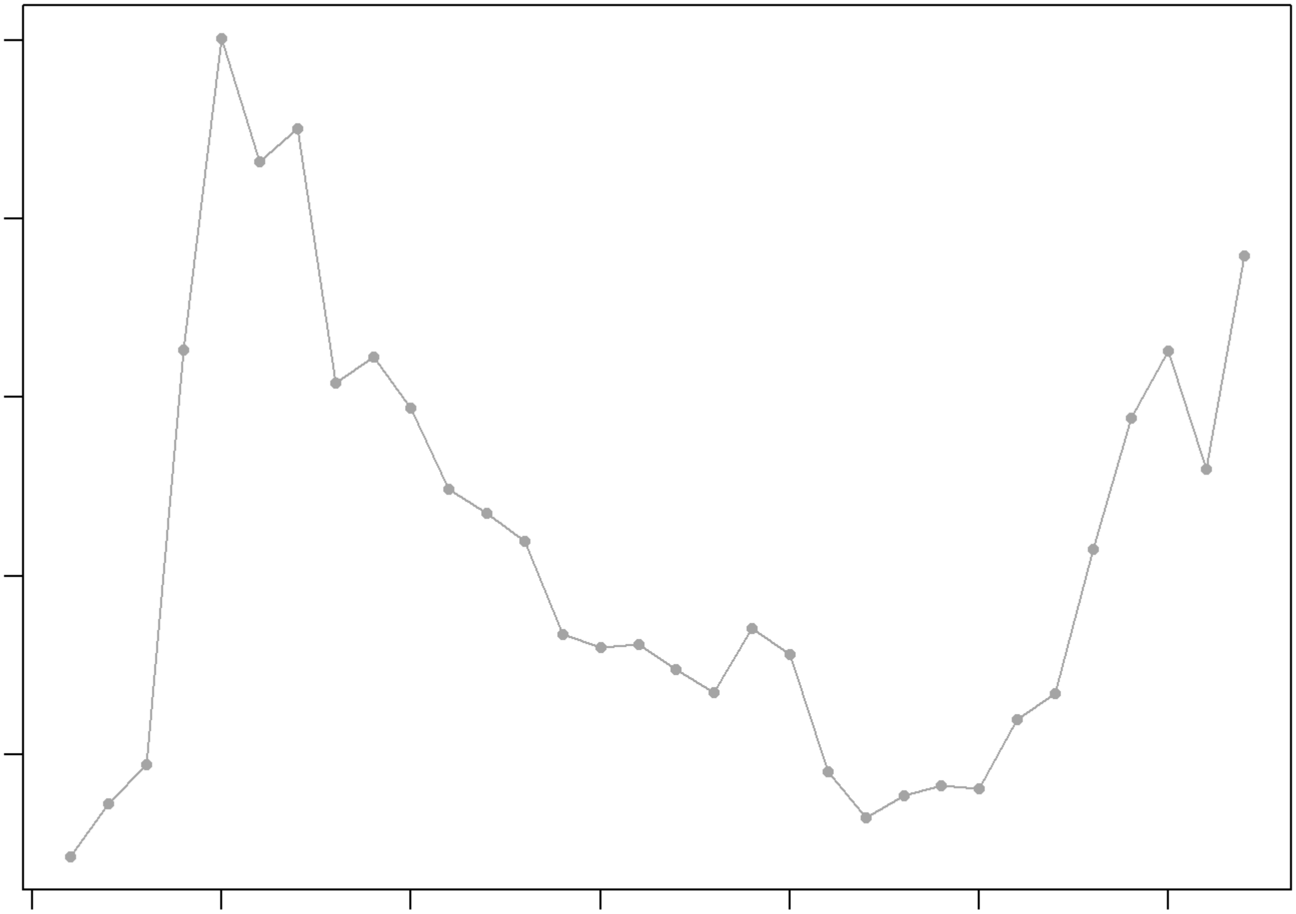

```
## NULL
## [1] "Creating 5472 artificial doublets..."
## [1] "Creating Seurat object..."
## [1] "Normalizing Seurat object..."
```

```
## [1] "Finding variable genes..."
```

```
## [1] "Scaling data..."
```

```
## [1] "Running PCA..."
## [1] "Calculating PC distance matrix..."
## [1] "Computing pANN..."
## [1] "Classifying doublets.."
```

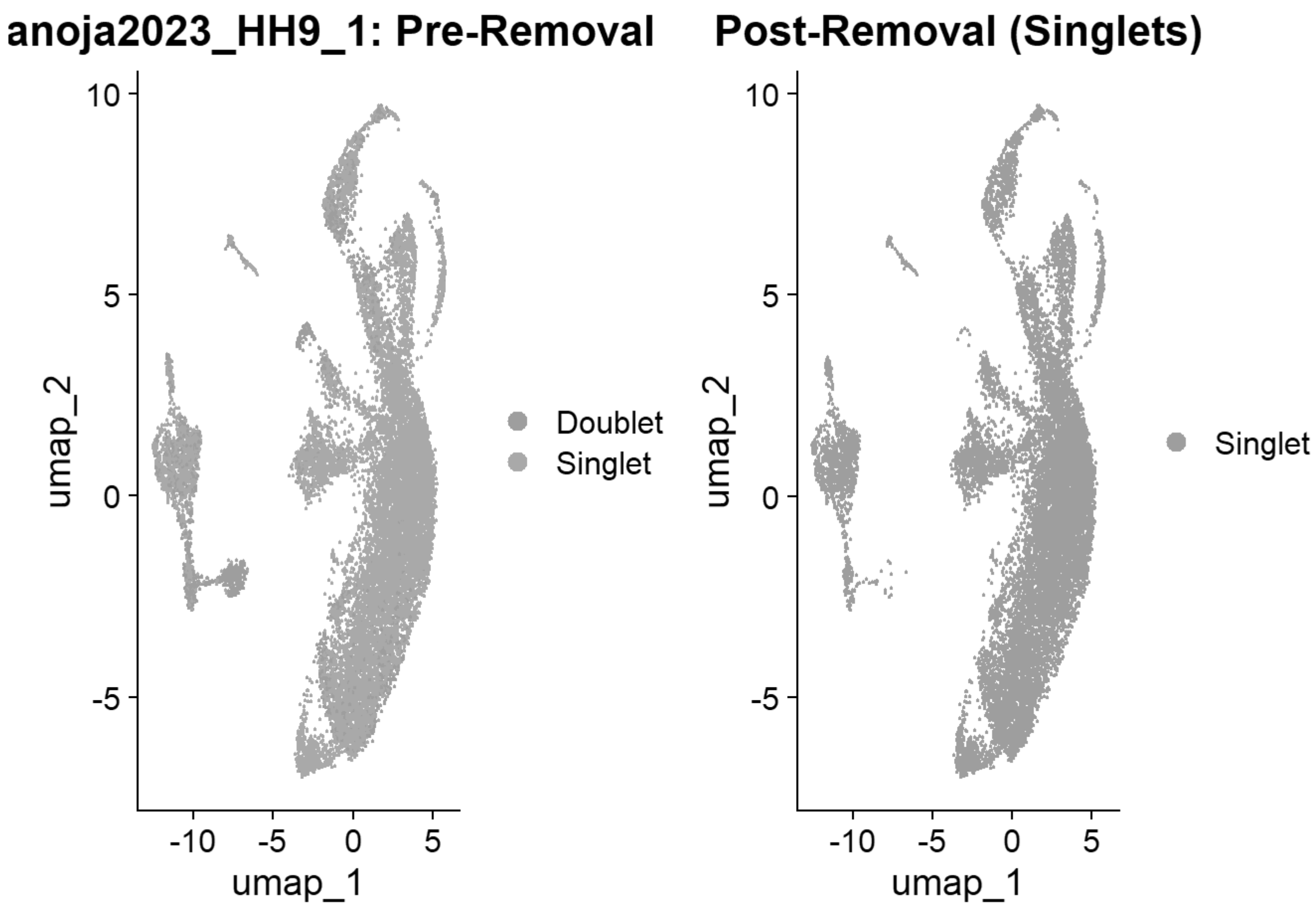


```
## [1] "Loading HH9_2"
## [1] "Running SoupX for HH9_2"
```

```
## Modularity Optimizer version 1.3.0 by Ludo Waltman and Nees Jan van Eck
##
## Number of nodes: 16758
## Number of edges: 537534
##
## Running Louvain algorithm...
## Maximum modularity in 10 random starts: 0.8820
## Number of communities: 19
## Elapsed time: 2 seconds
```

```
## [1] "Running QC for HH9_2"
## [1] "Running DoubletFinder for HH9_2"
## [1] "Creating artificial doublets for pN = 5%"
## [1] "Creating Seurat object..."
## [1] "Normalizing Seurat object..."
```

```
## [1] "Finding variable genes..."
```

```
## [1] "Scaling data..."
```

```
## [1] "Running PCA..."
## [1] "Calculating PC distance matrix..."
## [1] "Defining neighborhoods..."
## [1] "Computing pANN across all pK..."
## [1] "pK = 0.001..."
## [1] "pK = 0.005..."
## [1] "pK = 0.01..."
## [1] "pK = 0.02..."
## [1] "pK = 0.03..."
## [1] "pK = 0.04..."
## [1] "pK = 0.05..."
## [1] "pK = 0.06..."
## [1] "pK = 0.07..."
## [1] "pK = 0.08..."
## [1] "pK = 0.09..."
## [1] "pK = 0.1..."
## [1] "pK = 0.11..."
## [1] "pK = 0.12..."
## [1] "pK = 0.13..."
## [1] "pK = 0.14..."
## [1] "pK = 0.15..."
## [1] "pK = 0.16..."
## [1] "pK = 0.17..."
## [1] "pK = 0.18..."
## [1] "pK = 0.19..."
## [1] "pK = 0.2..."
## [1] "pK = 0.21..."
## [1] "pK = 0.22..."
## [1] "pK = 0.23..."
## [1] "pK = 0.24..."
## [1] "pK = 0.25..."
## [1] "pK = 0.26..."
## [1] "pK = 0.27..."
## [1] "pK = 0.28..."
## [1] "pK = 0.29..."
## [1] "pK = 0.3..."
## [1] "Creating artificial doublets for pN = 10%"
## [1] "Creating Seurat object..."
## [1] "Normalizing Seurat object..."
```

```
## [1] "Finding variable genes..."
```

```
## [1] "Scaling data..."
```

```
## [1] "Running PCA..."
## [1] "Calculating PC distance matrix..."
## [1] "Defining neighborhoods..."
## [1] "Computing pANN across all pK..."
## [1] "pK = 0.001..."
## [1] "pK = 0.005..."
## [1] "pK = 0.01..."
## [1] "pK = 0.02..."
```

```
## [1] "pK = 0.03..."
## [1] "pK = 0.04..."
## [1] "pK = 0.05..."
## [1] "pK = 0.06..."
## [1] "pK = 0.07..."
## [1] "pK = 0.08..."
## [1] "pK = 0.09..."
## [1] "pK = 0.1..."
## [1] "pK = 0.11..."
## [1] "pK = 0.12..."
## [1] "pK = 0.13..."
## [1] "pK = 0.14..."
## [1] "pK = 0.15..."
## [1] "pK = 0.16..."
## [1] "pK = 0.17..."
## [1] "pK = 0.18..."
## [1] "pK = 0.19..."
## [1] "pK = 0.2..."
## [1] "pK = 0.21..."
## [1] "pK = 0.22..."
## [1] "pK = 0.23..."
## [1] "pK = 0.24..."
## [1] "pK = 0.25..."
## [1] "pK = 0.26..."
## [1] "pK = 0.27..."
## [1] "pK = 0.28..."
## [1] "pK = 0.29..."
## [1] "pK = 0.3..."
## [1] "Creating artificial doublets for pN = 15%"
## [1] "Creating Seurat object..."
## [1] "Normalizing Seurat object..."
```

```
## [1] "Finding variable genes..."
```

```
## [1] "Scaling data..."
```

```
## [1] "Running PCA..."
## [1] "Calculating PC distance matrix..."
## [1] "Defining neighborhoods..."
## [1] "Computing pANN across all pK..."
## [1] "pK = 0.001..."
## [1] "pK = 0.005..."
## [1] "pK = 0.01..."
## [1] "pK = 0.02..."
## [1] "pK = 0.03..."
## [1] "pK = 0.04..."
## [1] "pK = 0.05..."
## [1] "pK = 0.06..."
## [1] "pK = 0.07..."
## [1] "pK = 0.08..."
## [1] "pK = 0.09..."
## [1] "pK = 0.1..."
```

```
## [1] "pK = 0.11..."
## [1] "pK = 0.12..."
## [1] "pK = 0.13..."
## [1] "pK = 0.14..."
## [1] "pK = 0.15..."
## [1] "pK = 0.16..."
## [1] "pK = 0.17..."
## [1] "pK = 0.18..."
## [1] "pK = 0.19..."
## [1] "pK = 0.2..."
## [1] "pK = 0.21..."
## [1] "pK = 0.22..."
## [1] "pK = 0.23..."
## [1] "pK = 0.24..."
## [1] "pK = 0.25..."
## [1] "pK = 0.26..."
## [1] "pK = 0.27..."
## [1] "pK = 0.28..."
## [1] "pK = 0.29..."
## [1] "pK = 0.3..."
## [1] "Creating artificial doublets for pN = 20%"
## [1] "Creating Seurat object..."
## [1] "Normalizing Seurat object..."
```

```
## [1] "Finding variable genes..."
```

```
## [1] "Scaling data..."
```

```
## [1] "Running PCA..."
## [1] "Calculating PC distance matrix..."
## [1] "Defining neighborhoods..."
## [1] "Computing pANN across all pK..."
## [1] "pK = 0.001..."
## [1] "pK = 0.005..."
## [1] "pK = 0.01..."
## [1] "pK = 0.02..."
## [1] "pK = 0.03..."
## [1] "pK = 0.04..."
## [1] "pK = 0.05..."
## [1] "pK = 0.06..."
## [1] "pK = 0.07..."
## [1] "pK = 0.08..."
## [1] "pK = 0.09..."
## [1] "pK = 0.1..."
## [1] "pK = 0.11..."
## [1] "pK = 0.12..."
## [1] "pK = 0.13..."
## [1] "pK = 0.14..."
## [1] "pK = 0.15..."
## [1] "pK = 0.16..."
## [1] "pK = 0.17..."
## [1] "pK = 0.18..."
```

```
## [1] "pK = 0.19..."
## [1] "pK = 0.2..."
## [1] "pK = 0.21..."
## [1] "pK = 0.22..."
## [1] "pK = 0.23..."
## [1] "pK = 0.24..."
## [1] "pK = 0.25..."
## [1] "pK = 0.26..."
## [1] "pK = 0.27..."
## [1] "pK = 0.28..."
## [1] "pK = 0.29..."
## [1] "pK = 0.3..."
## [1] "Creating artificial doublets for pN = 25%"
## [1] "Creating Seurat object..."
## [1] "Normalizing Seurat object..."
```

```
## [1] "Finding variable genes..."
```

```
## [1] "Scaling data..."
```

```
## [1] "Running PCA..."
## [1] "Calculating PC distance matrix..."
## [1] "Defining neighborhoods..."
## [1] "Computing pANN across all pK..."
## [1] "pK = 0.001..."
## [1] "pK = 0.005..."
## [1] "pK = 0.01..."
## [1] "pK = 0.02..."
## [1] "pK = 0.03..."
## [1] "pK = 0.04..."
## [1] "pK = 0.05..."
## [1] "pK = 0.06..."
## [1] "pK = 0.07..."
## [1] "pK = 0.08..."
## [1] "pK = 0.09..."
## [1] "pK = 0.1..."
## [1] "pK = 0.11..."
## [1] "pK = 0.12..."
## [1] "pK = 0.13..."
## [1] "pK = 0.14..."
## [1] "pK = 0.15..."
## [1] "pK = 0.16..."
## [1] "pK = 0.17..."
## [1] "pK = 0.18..."
## [1] "pK = 0.19..."
## [1] "pK = 0.2..."
## [1] "pK = 0.21..."
## [1] "pK = 0.22..."
## [1] "pK = 0.23..."
## [1] "pK = 0.24..."
## [1] "pK = 0.25..."
## [1] "pK = 0.26..."
```

```
## [1] "pK = 0.27..."
## [1] "pK = 0.28..."
## [1] "pK = 0.29..."
## [1] "pK = 0.3..."
## [1] "Creating artificial doublets for pN = 30%"
## [1] "Creating Seurat object..."
## [1] "Normalizing Seurat object..."
```

```
## [1] "Finding variable genes..."
```

```
## [1] "Scaling data..."
```

```
## [1] "Running PCA..."
## [1] "Calculating PC distance matrix..."
## [1] "Defining neighborhoods..."
## [1] "Computing pANN across all pK..."
## [1] "pK = 0.001..."
## [1] "pK = 0.005..."
## [1] "pK = 0.01..."
## [1] "pK = 0.02..."
## [1] "pK = 0.03..."
## [1] "pK = 0.04..."
## [1] "pK = 0.05..."
## [1] "pK = 0.06..."
## [1] "pK = 0.07..."
## [1] "pK = 0.08..."
## [1] "pK = 0.09..."
## [1] "pK = 0.1..."
## [1] "pK = 0.11..."
## [1] "pK = 0.12..."
## [1] "pK = 0.13..."
## [1] "pK = 0.14..."
## [1] "pK = 0.15..."
## [1] "pK = 0.16..."
## [1] "pK = 0.17..."
## [1] "pK = 0.18..."
## [1] "pK = 0.19..."
## [1] "pK = 0.2..."
## [1] "pK = 0.21..."
## [1] "pK = 0.22..."
## [1] "pK = 0.23..."
## [1] "pK = 0.24..."
## [1] "pK = 0.25..."
## [1] "pK = 0.26..."
## [1] "pK = 0.27..."
## [1] "pK = 0.28..."
## [1] "pK = 0.29..."
## [1] "pK = 0.3..."
```

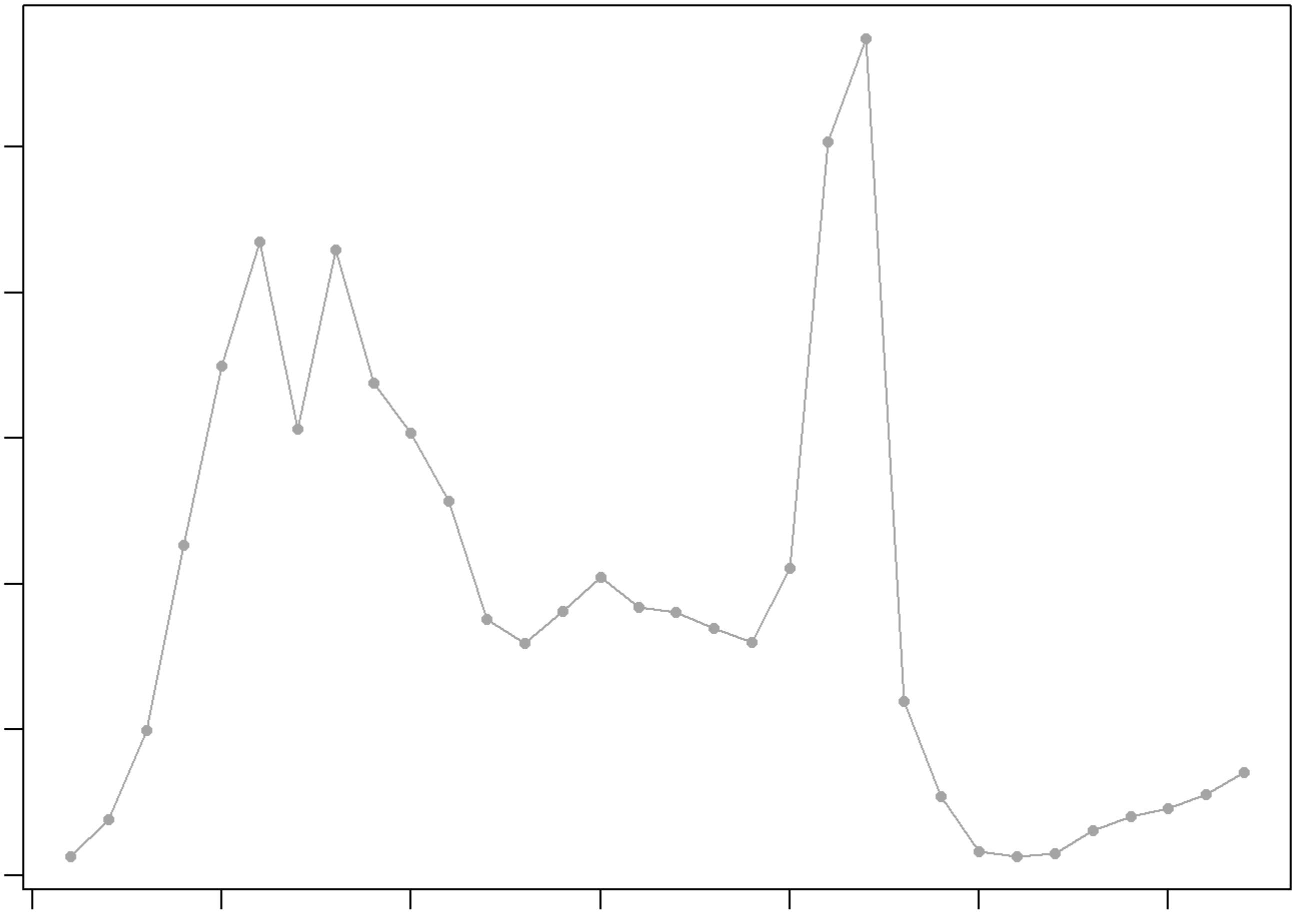

```
## NULL
## [1] "Creating 5356 artificial doublets..."
## [1] "Creating Seurat object..."
## [1] "Normalizing Seurat object..."
```

```
## [1] "Finding variable genes..."
```

```
## [1] "Scaling data..."
```

```
## [1] "Running PCA..."
## [1] "Calculating PC distance matrix..."
## [1] "Computing pANN..."
## [1] "Classifying doublets.."
```

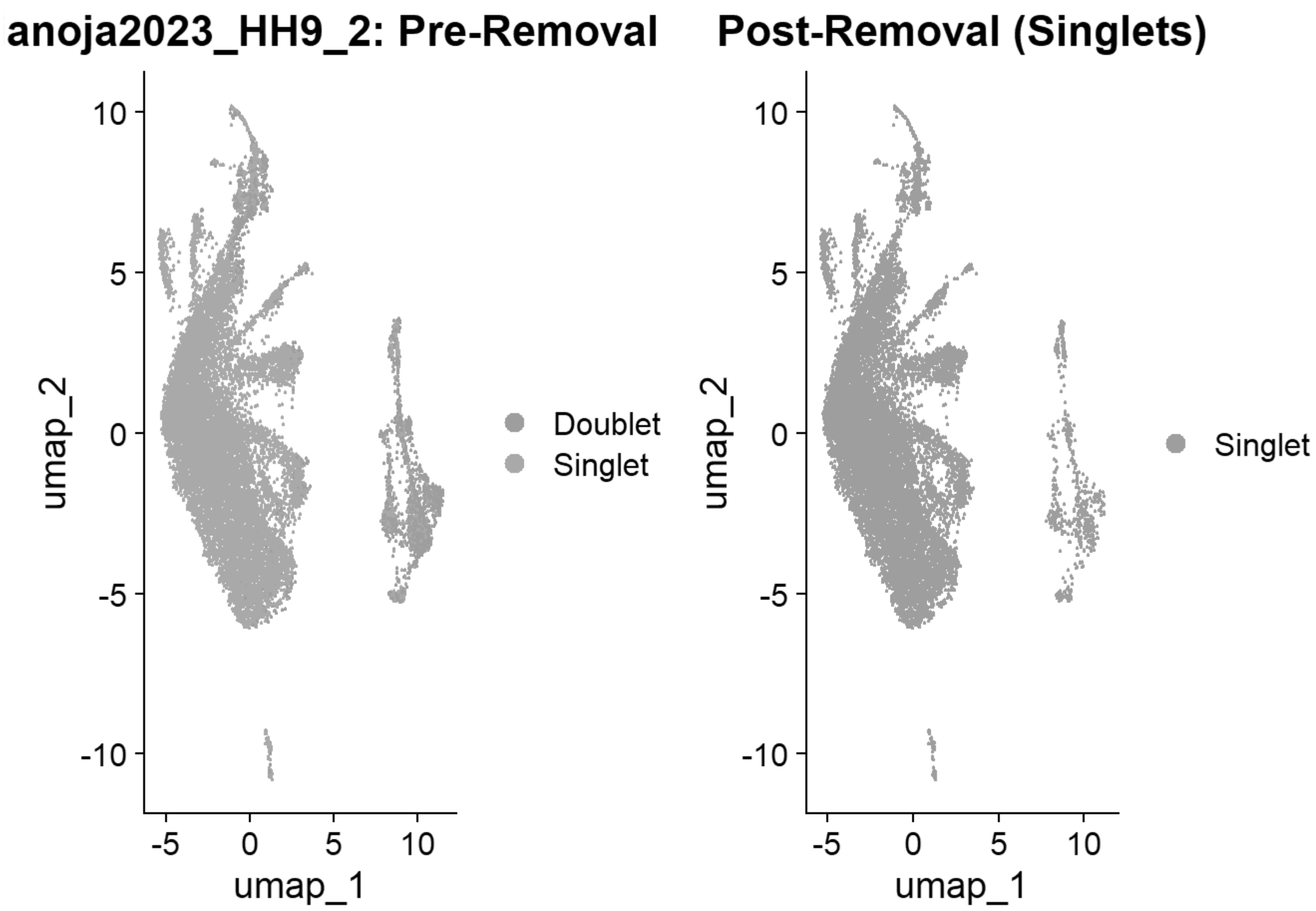


```
## [1] "Loading HH11"
## [1] "Running SoupX for HH11"
```

```
## Modularity Optimizer version 1.3.0 by Ludo Waltman and Nees Jan van Eck
##
## Number of nodes: 9806
## Number of edges: 337624
##
## Running Louvain algorithm...
## Maximum modularity in 10 random starts: 0.9112
## Number of communities: 15
## Elapsed time: 0 seconds
```

```
## [1] "Running QC for HH11"
## [1] "Running DoubletFinder for HH11"
## [1] "Creating artificial doublets for pN = 5%"
## [1] "Creating Seurat object..."
## [1] "Normalizing Seurat object..."
```

```
## [1] "Finding variable genes..."
```

```
## [1] "Scaling data..."
```

```
## [1] "Running PCA..."
## [1] "Calculating PC distance matrix..."
## [1] "Defining neighborhoods..."
## [1] "Computing pANN across all pK..."
## [1] "pK = 0.001..."
## [1] "pK = 0.005..."
## [1] "pK = 0.01..."
## [1] "pK = 0.02..."
## [1] "pK = 0.03..."
## [1] "pK = 0.04..."
## [1] "pK = 0.05..."
## [1] "pK = 0.06..."
## [1] "pK = 0.07..."
## [1] "pK = 0.08..."
## [1] "pK = 0.09..."
## [1] "pK = 0.1..."
## [1] "pK = 0.11..."
## [1] "pK = 0.12..."
## [1] "pK = 0.13..."
## [1] "pK = 0.14..."
## [1] "pK = 0.15..."
## [1] "pK = 0.16..."
## [1] "pK = 0.17..."
## [1] "pK = 0.18..."
## [1] "pK = 0.19..."
## [1] "pK = 0.2..."
## [1] "pK = 0.21..."
## [1] "pK = 0.22..."
## [1] "pK = 0.23..."
## [1] "pK = 0.24..."
## [1] "pK = 0.25..."
## [1] "pK = 0.26..."
## [1] "pK = 0.27..."
## [1] "pK = 0.28..."
## [1] "pK = 0.29..."
## [1] "pK = 0.3..."
## [1] "Creating artificial doublets for pN = 10%"
## [1] "Creating Seurat object..."
## [1] "Normalizing Seurat object..."
```

```
## [1] "Finding variable genes..."
```

```
## [1] "Scaling data..."
```

```
## [1] "Running PCA..."
## [1] "Calculating PC distance matrix..."
## [1] "Defining neighborhoods..."
## [1] "Computing pANN across all pK..."
## [1] "pK = 0.001..."
## [1] "pK = 0.005..."
## [1] "pK = 0.01..."
## [1] "pK = 0.02..."
```

```
## [1] "pK = 0.03..."
## [1] "pK = 0.04..."
## [1] "pK = 0.05..."
## [1] "pK = 0.06..."
## [1] "pK = 0.07..."
## [1] "pK = 0.08..."
## [1] "pK = 0.09..."
## [1] "pK = 0.1..."
## [1] "pK = 0.11..."
## [1] "pK = 0.12..."
## [1] "pK = 0.13..."
## [1] "pK = 0.14..."
## [1] "pK = 0.15..."
## [1] "pK = 0.16..."
## [1] "pK = 0.17..."
## [1] "pK = 0.18..."
## [1] "pK = 0.19..."
## [1] "pK = 0.2..."
## [1] "pK = 0.21..."
## [1] "pK = 0.22..."
## [1] "pK = 0.23..."
## [1] "pK = 0.24..."
## [1] "pK = 0.25..."
## [1] "pK = 0.26..."
## [1] "pK = 0.27..."
## [1] "pK = 0.28..."
## [1] "pK = 0.29..."
## [1] "pK = 0.3..."
## [1] "Creating artificial doublets for pN = 15%"
## [1] "Creating Seurat object..."
## [1] "Normalizing Seurat object..."
```

```
## [1] "Finding variable genes..."
```

```
## [1] "Scaling data..."
```

```
## [1] "Running PCA..."
## [1] "Calculating PC distance matrix..."
## [1] "Defining neighborhoods..."
## [1] "Computing pANN across all pK..."
## [1] "pK = 0.001..."
## [1] "pK = 0.005..."
## [1] "pK = 0.01..."
## [1] "pK = 0.02..."
## [1] "pK = 0.03..."
## [1] "pK = 0.04..."
## [1] "pK = 0.05..."
## [1] "pK = 0.06..."
## [1] "pK = 0.07..."
## [1] "pK = 0.08..."
## [1] "pK = 0.09..."
## [1] "pK = 0.1..."
```

```
## [1] "pK = 0.11..."
## [1] "pK = 0.12..."
## [1] "pK = 0.13..."
## [1] "pK = 0.14..."
## [1] "pK = 0.15..."
## [1] "pK = 0.16..."
## [1] "pK = 0.17..."
## [1] "pK = 0.18..."
## [1] "pK = 0.19..."
## [1] "pK = 0.2..."
## [1] "pK = 0.21..."
## [1] "pK = 0.22..."
## [1] "pK = 0.23..."
## [1] "pK = 0.24..."
## [1] "pK = 0.25..."
## [1] "pK = 0.26..."
## [1] "pK = 0.27..."
## [1] "pK = 0.28..."
## [1] "pK = 0.29..."
## [1] "pK = 0.3..."
## [1] "Creating artificial doublets for pN = 20%"
## [1] "Creating Seurat object..."
## [1] "Normalizing Seurat object..."
```

```
## [1] "Finding variable genes..."
```

```
## [1] "Scaling data..."
```

```
## [1] "Running PCA..."
## [1] "Calculating PC distance matrix..."
## [1] "Defining neighborhoods..."
## [1] "Computing pANN across all pK..."
## [1] "pK = 0.001..."
## [1] "pK = 0.005..."
## [1] "pK = 0.01..."
## [1] "pK = 0.02..."
## [1] "pK = 0.03..."
## [1] "pK = 0.04..."
## [1] "pK = 0.05..."
## [1] "pK = 0.06..."
## [1] "pK = 0.07..."
## [1] "pK = 0.08..."
## [1] "pK = 0.09..."
## [1] "pK = 0.1..."
## [1] "pK = 0.11..."
## [1] "pK = 0.12..."
## [1] "pK = 0.13..."
## [1] "pK = 0.14..."
## [1] "pK = 0.15..."
## [1] "pK = 0.16..."
## [1] "pK = 0.17..."
## [1] "pK = 0.18..."
```

```
## [1] "pK = 0.19..."
## [1] "pK = 0.2..."
## [1] "pK = 0.21..."
## [1] "pK = 0.22..."
## [1] "pK = 0.23..."
## [1] "pK = 0.24..."
## [1] "pK = 0.25..."
## [1] "pK = 0.26..."
## [1] "pK = 0.27..."
## [1] "pK = 0.28..."
## [1] "pK = 0.29..."
## [1] "pK = 0.3..."
## [1] "Creating artificial doublets for pN = 25%"
## [1] "Creating Seurat object..."
## [1] "Normalizing Seurat object..."
```

```
## [1] "Finding variable genes..."
```

```
## [1] "Scaling data..."
```

```
## [1] "Running PCA..."
## [1] "Calculating PC distance matrix..."
## [1] "Defining neighborhoods..."
## [1] "Computing pANN across all pK..."
## [1] "pK = 0.001..."
## [1] "pK = 0.005..."
## [1] "pK = 0.01..."
## [1] "pK = 0.02..."
## [1] "pK = 0.03..."
## [1] "pK = 0.04..."
## [1] "pK = 0.05..."
## [1] "pK = 0.06..."
## [1] "pK = 0.07..."
## [1] "pK = 0.08..."
## [1] "pK = 0.09..."
## [1] "pK = 0.1..."
## [1] "pK = 0.11..."
## [1] "pK = 0.12..."
## [1] "pK = 0.13..."
## [1] "pK = 0.14..."
## [1] "pK = 0.15..."
## [1] "pK = 0.16..."
## [1] "pK = 0.17..."
## [1] "pK = 0.18..."
## [1] "pK = 0.19..."
## [1] "pK = 0.2..."
## [1] "pK = 0.21..."
## [1] "pK = 0.22..."
## [1] "pK = 0.23..."
## [1] "pK = 0.24..."
## [1] "pK = 0.25..."
## [1] "pK = 0.26..."
```

```
## [1] "pK = 0.27..."
## [1] "pK = 0.28..."
## [1] "pK = 0.29..."
## [1] "pK = 0.3..."
## [1] "Creating artificial doublets for pN = 30%"
## [1] "Creating Seurat object..."
## [1] "Normalizing Seurat object..."
```

```
## [1] "Finding variable genes..."
```

```
## [1] "Scaling data..."
```

```
## [1] "Running PCA..."
## [1] "Calculating PC distance matrix..."
## [1] "Defining neighborhoods..."
## [1] "Computing pANN across all pK..."
## [1] "pK = 0.001..."
## [1] "pK = 0.005..."
## [1] "pK = 0.01..."
## [1] "pK = 0.02..."
## [1] "pK = 0.03..."
## [1] "pK = 0.04..."
## [1] "pK = 0.05..."
## [1] "pK = 0.06..."
## [1] "pK = 0.07..."
## [1] "pK = 0.08..."
## [1] "pK = 0.09..."
## [1] "pK = 0.1..."
## [1] "pK = 0.11..."
## [1] "pK = 0.12..."
## [1] "pK = 0.13..."
## [1] "pK = 0.14..."
## [1] "pK = 0.15..."
## [1] "pK = 0.16..."
## [1] "pK = 0.17..."
## [1] "pK = 0.18..."
## [1] "pK = 0.19..."
## [1] "pK = 0.2..."
## [1] "pK = 0.21..."
## [1] "pK = 0.22..."
## [1] "pK = 0.23..."
## [1] "pK = 0.24..."
## [1] "pK = 0.25..."
## [1] "pK = 0.26..."
## [1] "pK = 0.27..."
## [1] "pK = 0.28..."
## [1] "pK = 0.29..."
## [1] "pK = 0.3..."
```

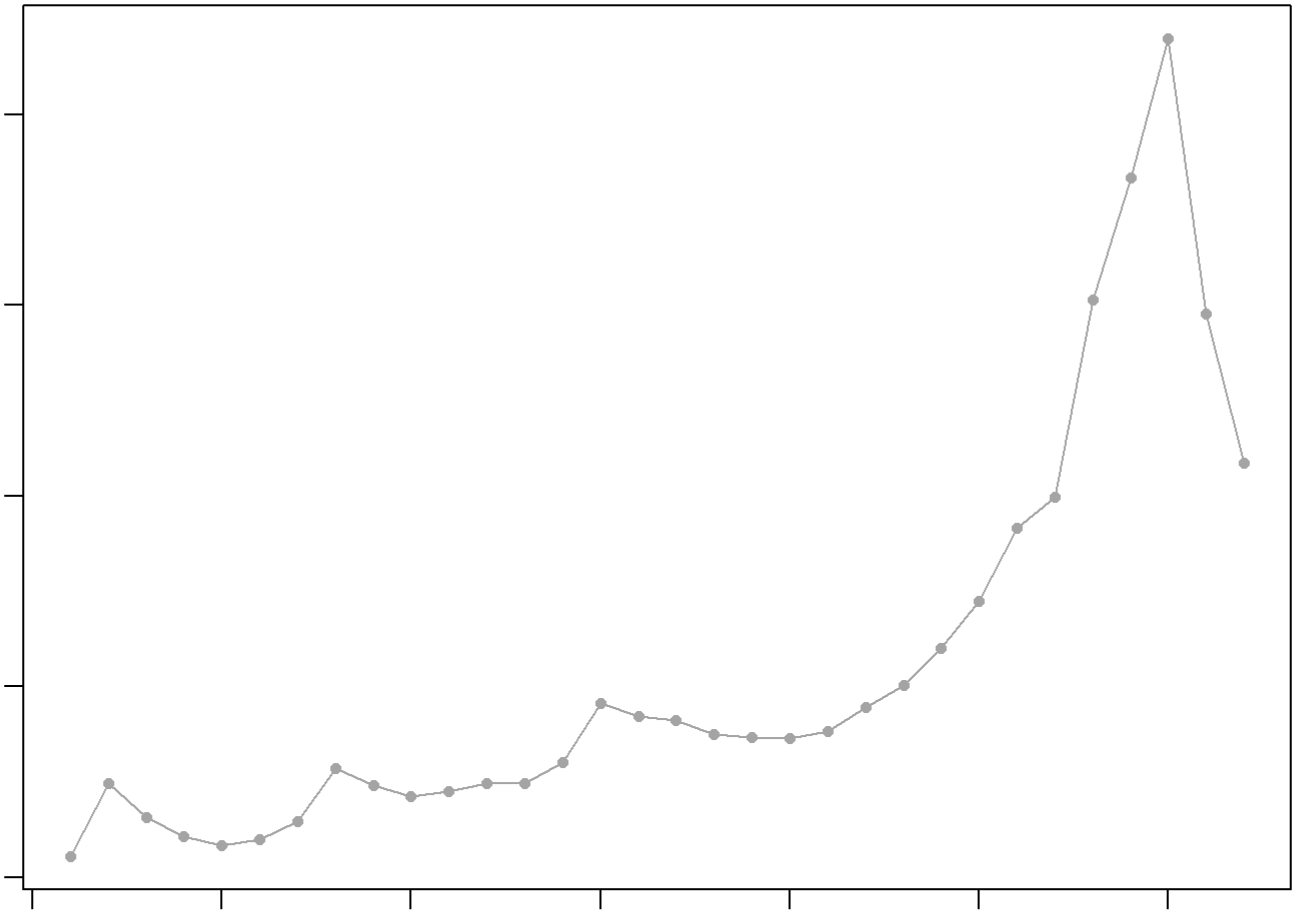

```
## NULL
## [1] "Creating 3203 artificial doublets..."
## [1] "Creating Seurat object..."
## [1] "Normalizing Seurat object..."
```

```
## [1] "Finding variable genes..."
```

```
## [1] "Scaling data..."
```

```
## [1] "Running PCA..."
## [1] "Calculating PC distance matrix..."
## [1] "Computing pANN..."
## [1] "Classifying doublets.."
```

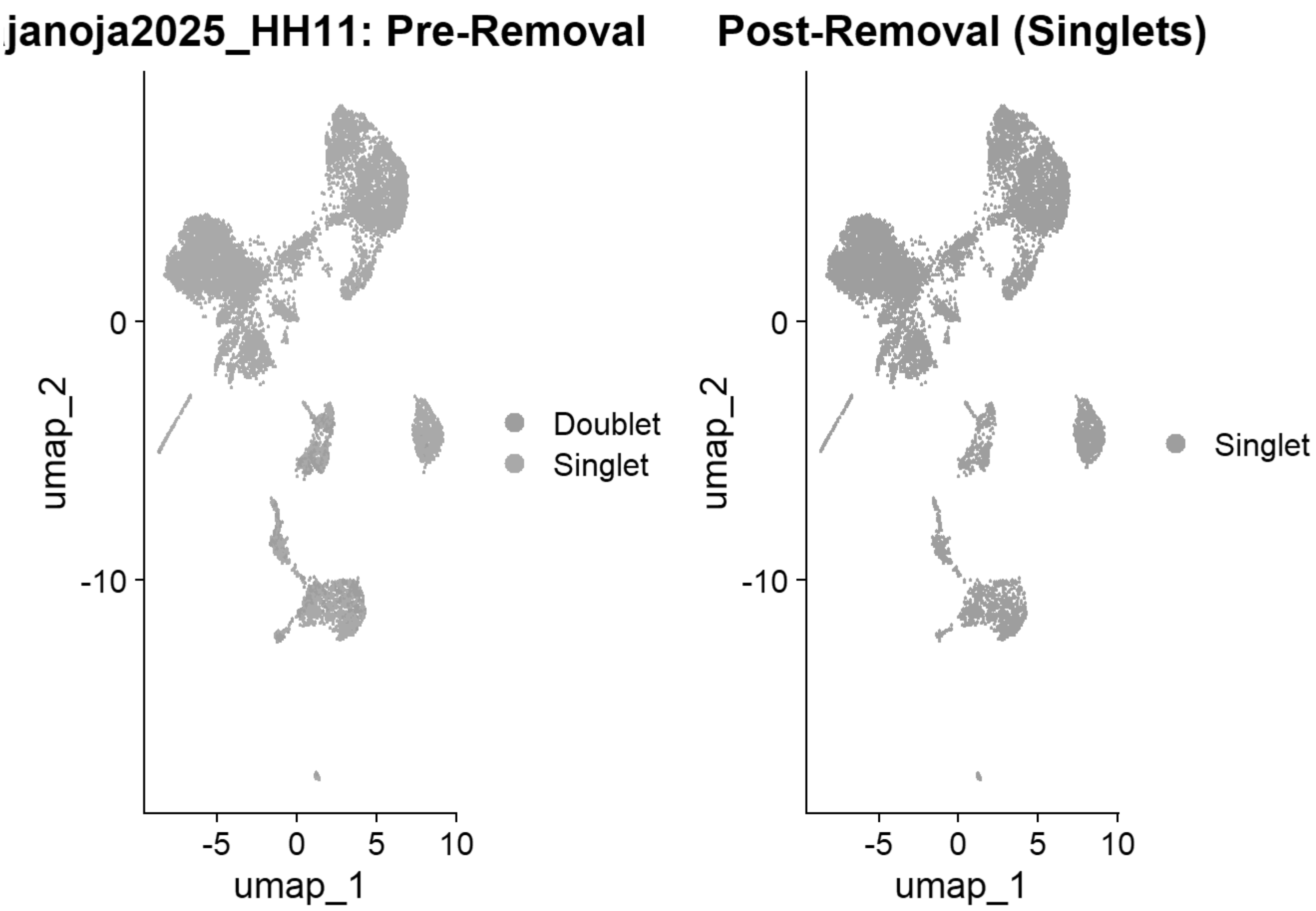


### 3. Processing Williams et al Dataset (GSE181577)

```
williams_samples <- c("HH4", "HH5", "HH6", "HH7")
williams_list <- list()

for (sample in williams_samples) {
  sample_full_name <- paste0("Williams_", sample)
  params <- get_qc_params(sample_full_name)

  filt_mat <- Read10X(paste0("E:\\Rogers Lab\\Williams 2021\\INPUT\\", sample))

  # Create Seurat Object & Record Pre-QC Count
  seu <- CreateSeuratObject(counts = filt_mat, project = sample_full_name, min.cells = 3, min.
features = params$min_feat)
  seu[["percent.mt"]] <- PercentageFeatureSet(seu, pattern = "^MT-")
  count_pre_qc <- ncol(seu)

  # Apply Strict QC & Record Post-QC Count
  seu <- subset(seu, subset = nFeature_RNA <= params$max_feat & percent.mt < params$max_mt)
  count_post_qc <- ncol(seu)
  rm(filt_mat)
  gc()
```

```
  # DoubletFinder
  seu <- NormalizeData(seu, verbose=F) %>% FindVariableFeatures(verbose=F) %>% ScaleData(verbo
se=F) %>% RunPCA(verbose=F) %>% RunUMAP(dims=1:20, verbose=F)
  sweep.res <- paramSweep(seu, PCs = 1:20, sct = FALSE)
  sweep.stats <- summarizeSweep(sweep.res, GT = FALSE)
  bcmvn <- find.pK(sweep.stats)
  pK_val <- as.numeric(as.character(bcmvn$pK[which.max(bcmvn$BCmetric)]))
  nExp_poi <- round(ncol(seu) * 8e-6 * ncol(seu))

  seu <- doubletFinder(seu, PCs = 1:20, pN = 0.25, pK = pK_val, nExp = nExp_poi, reuse.pANN =
NULL, sct = FALSE)
  df_col <- grep("DF.classifications", colnames(seu@meta.data), value = TRUE)
  seu$Doublet_Classification <- seu@meta.data[[df_col]]

  # Generate Pre-Removal Doublet UMAP
  p1 <- DimPlot(seu, group.by = "Doublet_Classification")+ggtitle(label = paste0(sample_full_n
ame, ": Pre-Removal"))

  # Subset Singlets & Add Metadata
  seu <- subset(seu, subset = Doublet_Classification == "Singlet")
  seu$sample_id <- sample_full_name
  seu$stage <- sample
  seu$ref <- "Williams_2022"
  count_post_df <- ncol(seu)

  # Generate Post-Removal Singlet UMAP and plot side-by-side
  p2 <- DimPlot(seu, group.by = "Doublet_Classification") +ggtitle(label = "Post-Removal (Sing
lets)")
  print(p1 | p2)

  # Append to Tracker
  qc_tracker[[sample_full_name]] <- data.frame(Sample = sample_full_name, Pre_QC = count_pre_q
c, Post_QC = count_post_qc, Post_DoubletFinder = count_post_df)

  williams_list[[sample]] <- seu

  # save copy of corrected object
  saveRDS(seu,paste0("E:\\Rogers Lab\\Rogers NC EMT\\Williams_", sample,"_v1.rds"))
}
```

```
## [1] "Creating artificial doublets for pN = 5%"
## [1] "Creating Seurat object..."
## [1] "Normalizing Seurat object..."
```

```
## [1] "Finding variable genes..."
```

```
## [1] "Scaling data..."
```

```
## [1] "Running PCA..."
## [1] "Calculating PC distance matrix..."
## [1] "Defining neighborhoods..."
```

```
## [1] "Computing pANN across all pK..."
## [1] "pK = 0.005..."
## [1] "pK = 0.01..."
## [1] "pK = 0.02..."
## [1] "pK = 0.03..."
## [1] "pK = 0.04..."
## [1] "pK = 0.05..."
## [1] "pK = 0.06..."
## [1] "pK = 0.07..."
## [1] "pK = 0.08..."
## [1] "pK = 0.09..."
## [1] "pK = 0.1..."
## [1] "pK = 0.11..."
## [1] "pK = 0.12..."
## [1] "pK = 0.13..."
## [1] "pK = 0.14..."
## [1] "pK = 0.15..."
## [1] "pK = 0.16..."
## [1] "pK = 0.17..."
## [1] "pK = 0.18..."
## [1] "pK = 0.19..."
## [1] "pK = 0.2..."
## [1] "pK = 0.21..."
## [1] "pK = 0.22..."
## [1] "pK = 0.23..."
## [1] "pK = 0.24..."
## [1] "pK = 0.25..."
## [1] "pK = 0.26..."
## [1] "pK = 0.27..."
## [1] "pK = 0.28..."
## [1] "pK = 0.29..."
## [1] "pK = 0.3..."
## [1] "Creating artificial doublets for pN = 10%"
## [1] "Creating Seurat object..."
## [1] "Normalizing Seurat object..."
```

```
## [1] "Finding variable genes..."
```

```
## [1] "Scaling data..."
```

```
## [1] "Running PCA..."
## [1] "Calculating PC distance matrix..."
## [1] "Defining neighborhoods..."
## [1] "Computing pANN across all pK..."
## [1] "pK = 0.005..."
## [1] "pK = 0.01..."
## [1] "pK = 0.02..."
## [1] "pK = 0.03..."
## [1] "pK = 0.04..."
## [1] "pK = 0.05..."
## [1] "pK = 0.06..."
## [1] "pK = 0.07..."
```

```
## [1] "pK = 0.08..."
## [1] "pK = 0.09..."
## [1] "pK = 0.1..."
## [1] "pK = 0.11..."
## [1] "pK = 0.12..."
## [1] "pK = 0.13..."
## [1] "pK = 0.14..."
## [1] "pK = 0.15..."
## [1] "pK = 0.16..."
## [1] "pK = 0.17..."
## [1] "pK = 0.18..."
## [1] "pK = 0.19..."
## [1] "pK = 0.2..."
## [1] "pK = 0.21..."
## [1] "pK = 0.22..."
## [1] "pK = 0.23..."
## [1] "pK = 0.24..."
## [1] "pK = 0.25..."
## [1] "pK = 0.26..."
## [1] "pK = 0.27..."
## [1] "pK = 0.28..."
## [1] "pK = 0.29..."
## [1] "pK = 0.3..."
## [1] "Creating artificial doublets for pN = 15%"
## [1] "Creating Seurat object..."
## [1] "Normalizing Seurat object..."
```

```
## [1] "Finding variable genes..."
```

```
## [1] "Scaling data..."
```

```
## [1] "Running PCA..."
## [1] "Calculating PC distance matrix..."
## [1] "Defining neighborhoods..."
## [1] "Computing pANN across all pK..."
## [1] "pK = 0.005..."
## [1] "pK = 0.01..."
## [1] "pK = 0.02..."
## [1] "pK = 0.03..."
## [1] "pK = 0.04..."
## [1] "pK = 0.05..."
## [1] "pK = 0.06..."
## [1] "pK = 0.07..."
## [1] "pK = 0.08..."
## [1] "pK = 0.09..."
## [1] "pK = 0.1..."
## [1] "pK = 0.11..."
## [1] "pK = 0.12..."
## [1] "pK = 0.13..."
## [1] "pK = 0.14..."
## [1] "pK = 0.15..."
## [1] "pK = 0.16..."
```

```
## [1] "pK = 0.17..."
## [1] "pK = 0.18..."
## [1] "pK = 0.19..."
## [1] "pK = 0.2..."
## [1] "pK = 0.21..."
## [1] "pK = 0.22..."
## [1] "pK = 0.23..."
## [1] "pK = 0.24..."
## [1] "pK = 0.25..."
## [1] "pK = 0.26..."
## [1] "pK = 0.27..."
## [1] "pK = 0.28..."
## [1] "pK = 0.29..."
## [1] "pK = 0.3..."
## [1] "Creating artificial doublets for pN = 20%"
## [1] "Creating Seurat object..."
## [1] "Normalizing Seurat object..."
```

```
## [1] "Finding variable genes..."
```

```
## [1] "Scaling data..."
```

```
## [1] "Running PCA..."
## [1] "Calculating PC distance matrix..."
## [1] "Defining neighborhoods..."
## [1] "Computing pANN across all pK..."
## [1] "pK = 0.005..."
## [1] "pK = 0.01..."
## [1] "pK = 0.02..."
## [1] "pK = 0.03..."
## [1] "pK = 0.04..."
## [1] "pK = 0.05..."
## [1] "pK = 0.06..."
## [1] "pK = 0.07..."
## [1] "pK = 0.08..."
## [1] "pK = 0.09..."
## [1] "pK = 0.1..."
## [1] "pK = 0.11..."
## [1] "pK = 0.12..."
## [1] "pK = 0.13..."
## [1] "pK = 0.14..."
## [1] "pK = 0.15..."
## [1] "pK = 0.16..."
## [1] "pK = 0.17..."
## [1] "pK = 0.18..."
## [1] "pK = 0.19..."
## [1] "pK = 0.2..."
## [1] "pK = 0.21..."
## [1] "pK = 0.22..."
## [1] "pK = 0.23..."
## [1] "pK = 0.24..."
## [1] "pK = 0.25..."
```

```
## [1] "pK = 0.26..."
## [1] "pK = 0.27..."
## [1] "pK = 0.28..."
## [1] "pK = 0.29..."
## [1] "pK = 0.3..."
## [1] "Creating artificial doublets for pN = 25%"
## [1] "Creating Seurat object..."
## [1] "Normalizing Seurat object..."
```

```
## [1] "Finding variable genes..."
```

```
## [1] "Scaling data..."
```

```
## [1] "Running PCA..."
## [1] "Calculating PC distance matrix..."
## [1] "Defining neighborhoods..."
## [1] "Computing pANN across all pK..."
## [1] "pK = 0.005..."
## [1] "pK = 0.01..."
## [1] "pK = 0.02..."
## [1] "pK = 0.03..."
## [1] "pK = 0.04..."
## [1] "pK = 0.05..."
## [1] "pK = 0.06..."
## [1] "pK = 0.07..."
## [1] "pK = 0.08..."
## [1] "pK = 0.09..."
## [1] "pK = 0.1..."
## [1] "pK = 0.11..."
## [1] "pK = 0.12..."
## [1] "pK = 0.13..."
## [1] "pK = 0.14..."
## [1] "pK = 0.15..."
## [1] "pK = 0.16..."
## [1] "pK = 0.17..."
## [1] "pK = 0.18..."
## [1] "pK = 0.19..."
## [1] "pK = 0.2..."
## [1] "pK = 0.21..."
## [1] "pK = 0.22..."
## [1] "pK = 0.23..."
## [1] "pK = 0.24..."
## [1] "pK = 0.25..."
## [1] "pK = 0.26..."
## [1] "pK = 0.27..."
## [1] "pK = 0.28..."
## [1] "pK = 0.29..."
## [1] "pK = 0.3..."
## [1] "Creating artificial doublets for pN = 30%"
## [1] "Creating Seurat object..."
## [1] "Normalizing Seurat object..."
```

```
## [1] "Finding variable genes..."
```

```
## [1] "Scaling data..."
```

```
## [1] "Running PCA..."
## [1] "Calculating PC distance matrix..."
## [1] "Defining neighborhoods..."
## [1] "Computing pANN across all pK..."
## [1] "pK = 0.005..."
## [1] "pK = 0.01..."
## [1] "pK = 0.02..."
## [1] "pK = 0.03..."
## [1] "pK = 0.04..."
## [1] "pK = 0.05..."
## [1] "pK = 0.06..."
## [1] "pK = 0.07..."
## [1] "pK = 0.08..."
## [1] "pK = 0.09..."
## [1] "pK = 0.1..."
## [1] "pK = 0.11..."
## [1] "pK = 0.12..."
## [1] "pK = 0.13..."
## [1] "pK = 0.14..."
## [1] "pK = 0.15..."
## [1] "pK = 0.16..."
## [1] "pK = 0.17..."
## [1] "pK = 0.18..."
## [1] "pK = 0.19..."
## [1] "pK = 0.2..."
## [1] "pK = 0.21..."
## [1] "pK = 0.22..."
## [1] "pK = 0.23..."
## [1] "pK = 0.24..."
## [1] "pK = 0.25..."
## [1] "pK = 0.26..."
## [1] "pK = 0.27..."
## [1] "pK = 0.28..."
## [1] "pK = 0.29..."
## [1] "pK = 0.3..."
```

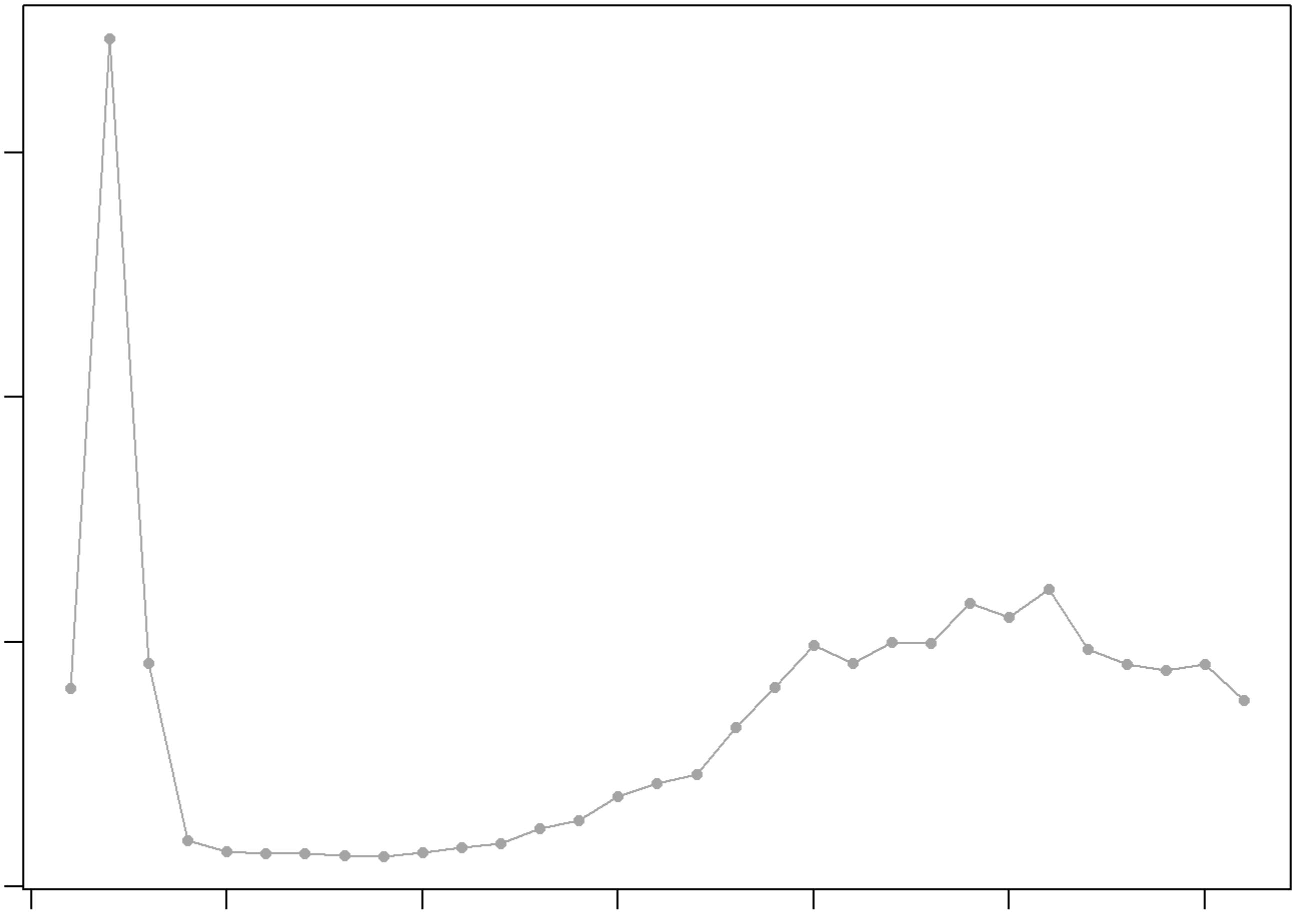

```
## NULL
## [1] "Creating 728 artificial doublets..."
## [1] "Creating Seurat object..."
## [1] "Normalizing Seurat object..."
```

```
## [1] "Finding variable genes..."
```

```
## [1] "Scaling data..."
```

```
## [1] "Running PCA..."
## [1] "Calculating PC distance matrix..."
## [1] "Computing pANN..."
## [1] "Classifying doublets.."
```

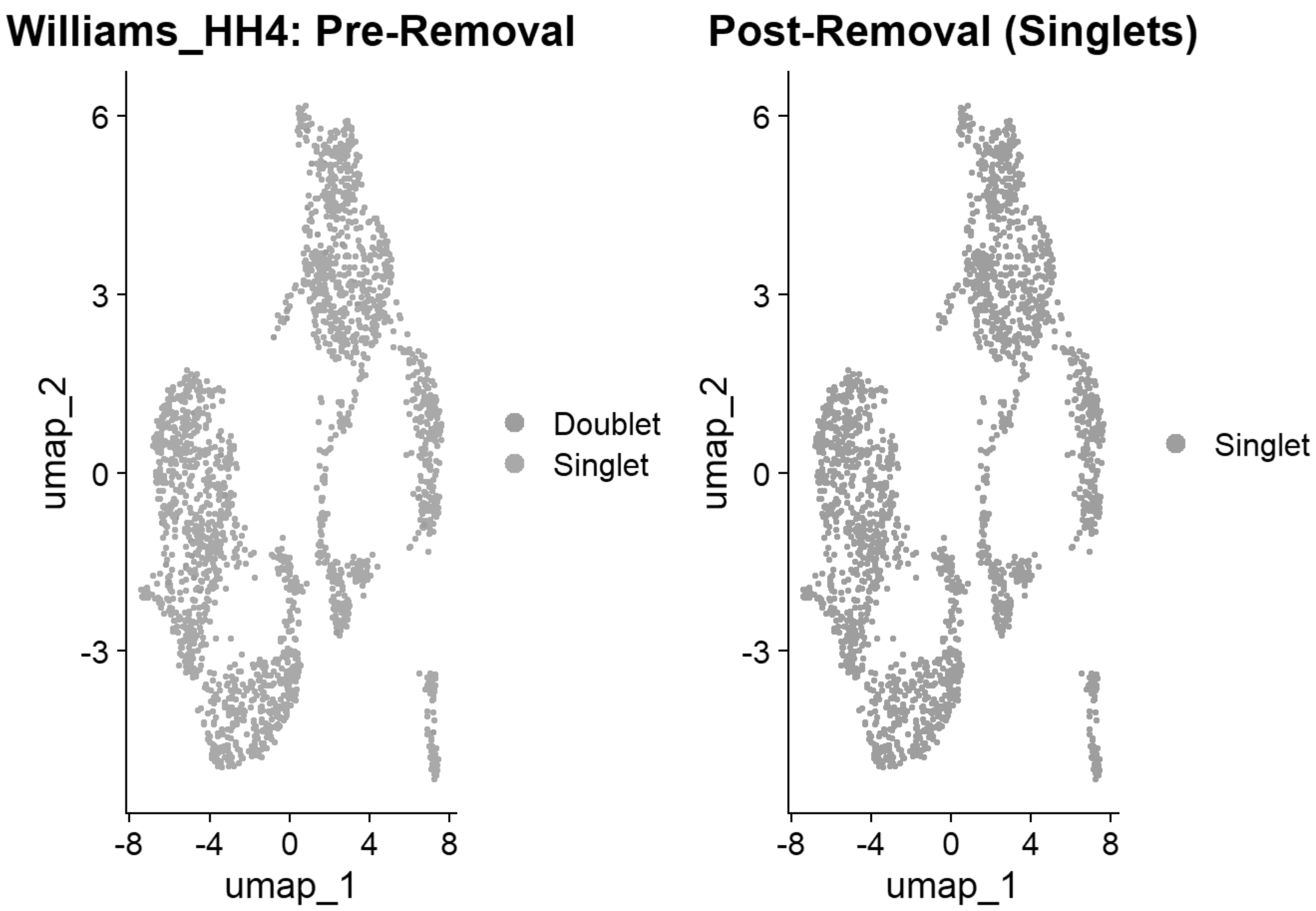


```
## [1] "Creating artificial doublets for pN = 5%"
## [1] "Creating Seurat object..."
## [1] "Normalizing Seurat object..."
```

```
## [1] "Finding variable genes..."
```

```
## [1] "Scaling data..."
```

```
## [1] "Running PCA..."
## [1] "Calculating PC distance matrix..."
## [1] "Defining neighborhoods..."
## [1] "Computing pANN across all pK..."
## [1] "pK = 0.005..."
## [1] "pK = 0.01..."
## [1] "pK = 0.02..."
## [1] "pK = 0.03..."
## [1] "pK = 0.04..."
## [1] "pK = 0.05..."
## [1] "pK = 0.06..."
## [1] "pK = 0.07..."
## [1] "pK = 0.08..."
## [1] "pK = 0.09..."
## [1] "pK = 0.1..."
## [1] "pK = 0.11..."
```

```
## [1] "pK = 0.12..."
## [1] "pK = 0.13..."
## [1] "pK = 0.14..."
## [1] "pK = 0.15..."
## [1] "pK = 0.16..."
## [1] "pK = 0.17..."
## [1] "pK = 0.18..."
## [1] "pK = 0.19..."
## [1] "pK = 0.2..."
## [1] "pK = 0.21..."
## [1] "pK = 0.22..."
## [1] "pK = 0.23..."
## [1] "pK = 0.24..."
## [1] "pK = 0.25..."
## [1] "pK = 0.26..."
## [1] "pK = 0.27..."
## [1] "pK = 0.28..."
## [1] "pK = 0.29..."
## [1] "pK = 0.3..."
## [1] "Creating artificial doublets for pN = 10%"
## [1] "Creating Seurat object..."
## [1] "Normalizing Seurat object..."
```

```
## [1] "Finding variable genes..."
```

```
## [1] "Scaling data..."
```

```
## [1] "Running PCA..."
## [1] "Calculating PC distance matrix..."
## [1] "Defining neighborhoods..."
## [1] "Computing pANN across all pK..."
## [1] "pK = 0.005..."
## [1] "pK = 0.01..."
## [1] "pK = 0.02..."
## [1] "pK = 0.03..."
## [1] "pK = 0.04..."
## [1] "pK = 0.05..."
## [1] "pK = 0.06..."
## [1] "pK = 0.07..."
## [1] "pK = 0.08..."
## [1] "pK = 0.09..."
## [1] "pK = 0.1..."
## [1] "pK = 0.11..."
## [1] "pK = 0.12..."
## [1] "pK = 0.13..."
## [1] "pK = 0.14..."
## [1] "pK = 0.15..."
## [1] "pK = 0.16..."
## [1] "pK = 0.17..."
## [1] "pK = 0.18..."
## [1] "pK = 0.19..."
## [1] "pK = 0.2..."
```

```
## [1] "pK = 0.21..."
## [1] "pK = 0.22..."
## [1] "pK = 0.23..."
## [1] "pK = 0.24..."
## [1] "pK = 0.25..."
## [1] "pK = 0.26..."
## [1] "pK = 0.27..."
## [1] "pK = 0.28..."
## [1] "pK = 0.29..."
## [1] "pK = 0.3..."
## [1] "Creating artificial doublets for pN = 15%"
## [1] "Creating Seurat object..."
## [1] "Normalizing Seurat object..."
```

```
## [1] "Finding variable genes..."
```

```
## [1] "Scaling data..."
```

```
## [1] "Running PCA..."
## [1] "Calculating PC distance matrix..."
## [1] "Defining neighborhoods..."
## [1] "Computing pANN across all pK..."
## [1] "pK = 0.005..."
## [1] "pK = 0.01..."
## [1] "pK = 0.02..."
## [1] "pK = 0.03..."
## [1] "pK = 0.04..."
## [1] "pK = 0.05..."
## [1] "pK = 0.06..."
## [1] "pK = 0.07..."
## [1] "pK = 0.08..."
## [1] "pK = 0.09..."
## [1] "pK = 0.1..."
## [1] "pK = 0.11..."
## [1] "pK = 0.12..."
## [1] "pK = 0.13..."
## [1] "pK = 0.14..."
## [1] "pK = 0.15..."
## [1] "pK = 0.16..."
## [1] "pK = 0.17..."
## [1] "pK = 0.18..."
## [1] "pK = 0.19..."
## [1] "pK = 0.2..."
## [1] "pK = 0.21..."
## [1] "pK = 0.22..."
## [1] "pK = 0.23..."
## [1] "pK = 0.24..."
## [1] "pK = 0.25..."
## [1] "pK = 0.26..."
## [1] "pK = 0.27..."
## [1] "pK = 0.28..."
## [1] "pK = 0.29..."
```

```
## [1] "pK = 0.3..."
## [1] "Creating artificial doublets for pN = 20%"
## [1] "Creating Seurat object..."
## [1] "Normalizing Seurat object..."
```

```
## [1] "Finding variable genes..."
```

```
## [1] "Scaling data..."
```

```
## [1] "Running PCA..."
## [1] "Calculating PC distance matrix..."
## [1] "Defining neighborhoods..."
## [1] "Computing pANN across all pK..."
## [1] "pK = 0.005..."
## [1] "pK = 0.01..."
## [1] "pK = 0.02..."
## [1] "pK = 0.03..."
## [1] "pK = 0.04..."
## [1] "pK = 0.05..."
## [1] "pK = 0.06..."
## [1] "pK = 0.07..."
## [1] "pK = 0.08..."
## [1] "pK = 0.09..."
## [1] "pK = 0.1..."
## [1] "pK = 0.11..."
## [1] "pK = 0.12..."
## [1] "pK = 0.13..."
## [1] "pK = 0.14..."
## [1] "pK = 0.15..."
## [1] "pK = 0.16..."
## [1] "pK = 0.17..."
## [1] "pK = 0.18..."
## [1] "pK = 0.19..."
## [1] "pK = 0.2..."
## [1] "pK = 0.21..."
## [1] "pK = 0.22..."
## [1] "pK = 0.23..."
## [1] "pK = 0.24..."
## [1] "pK = 0.25..."
## [1] "pK = 0.26..."
## [1] "pK = 0.27..."
## [1] "pK = 0.28..."
## [1] "pK = 0.29..."
## [1] "pK = 0.3..."
## [1] "Creating artificial doublets for pN = 25%"
## [1] "Creating Seurat object..."
## [1] "Normalizing Seurat object..."
```

```
## [1] "Finding variable genes..."
```

```
## [1] "Scaling data..."
```

```
## [1] "Running PCA..."
## [1] "Calculating PC distance matrix..."
## [1] "Defining neighborhoods..."
## [1] "Computing pANN across all pK..."
## [1] "pK = 0.005..."
## [1] "pK = 0.01..."
## [1] "pK = 0.02..."
## [1] "pK = 0.03..."
## [1] "pK = 0.04..."
## [1] "pK = 0.05..."
## [1] "pK = 0.06..."
## [1] "pK = 0.07..."
## [1] "pK = 0.08..."
## [1] "pK = 0.09..."
## [1] "pK = 0.1..."
## [1] "pK = 0.11..."
## [1] "pK = 0.12..."
## [1] "pK = 0.13..."
## [1] "pK = 0.14..."
## [1] "pK = 0.15..."
## [1] "pK = 0.16..."
## [1] "pK = 0.17..."
## [1] "pK = 0.18..."
## [1] "pK = 0.19..."
## [1] "pK = 0.2..."
## [1] "pK = 0.21..."
## [1] "pK = 0.22..."
## [1] "pK = 0.23..."
## [1] "pK = 0.24..."
## [1] "pK = 0.25..."
## [1] "pK = 0.26..."
## [1] "pK = 0.27..."
## [1] "pK = 0.28..."
## [1] "pK = 0.29..."
## [1] "pK = 0.3..."
## [1] "Creating artificial doublets for pN = 30%"
## [1] "Creating Seurat object..."
## [1] "Normalizing Seurat object..."
```

```
## [1] "Finding variable genes..."
```

```
## [1] "Scaling data..."
```

```
## [1] "Running PCA..."
## [1] "Calculating PC distance matrix..."
## [1] "Defining neighborhoods..."
## [1] "Computing pANN across all pK..."
## [1] "pK = 0.005..."
## [1] "pK = 0.01..."
## [1] "pK = 0.02..."
```

```
## [1] "pK = 0.03..."
## [1] "pK = 0.04..."
## [1] "pK = 0.05..."
## [1] "pK = 0.06..."
## [1] "pK = 0.07..."
## [1] "pK = 0.08..."
## [1] "pK = 0.09..."
## [1] "pK = 0.1..."
## [1] "pK = 0.11..."
## [1] "pK = 0.12..."
## [1] "pK = 0.13..."
## [1] "pK = 0.14..."
## [1] "pK = 0.15..."
## [1] "pK = 0.16..."
## [1] "pK = 0.17..."
## [1] "pK = 0.18..."
## [1] "pK = 0.19..."
## [1] "pK = 0.2..."
## [1] "pK = 0.21..."
## [1] "pK = 0.22..."
## [1] "pK = 0.23..."
## [1] "pK = 0.24..."
## [1] "pK = 0.25..."
## [1] "pK = 0.26..."
## [1] "pK = 0.27..."
## [1] "pK = 0.28..."
## [1] "pK = 0.29..."
## [1] "pK = 0.3..."
```

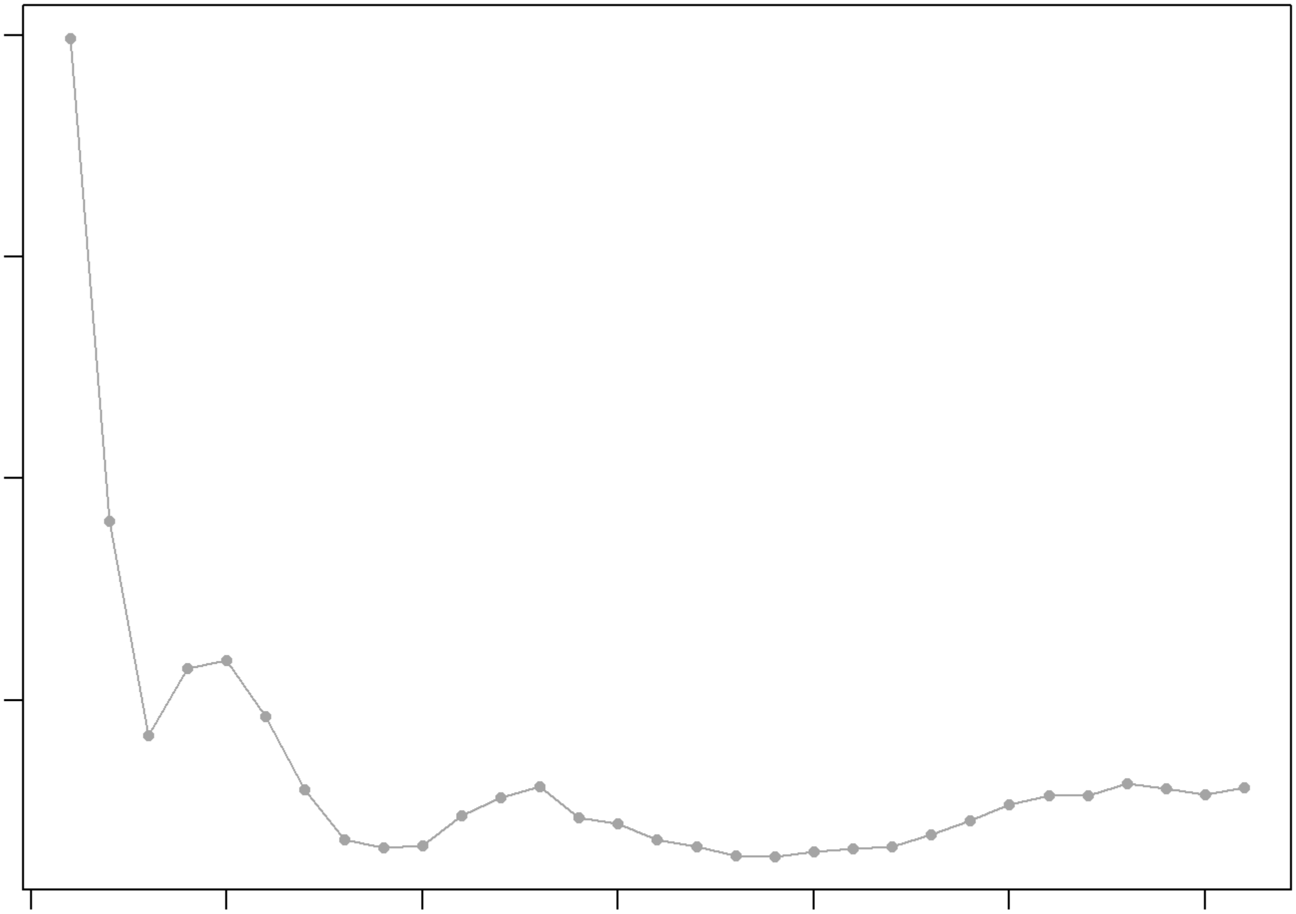

```
## NULL
## [1] "Creating 841 artificial doublets..."
## [1] "Creating Seurat object..."
## [1] "Normalizing Seurat object..."
```

```
## [1] "Finding variable genes..."
```

```
## [1] "Scaling data..."
```

```
## [1] "Running PCA..."
## [1] "Calculating PC distance matrix..."
## [1] "Computing pANN..."
## [1] "Classifying doublets.."
```

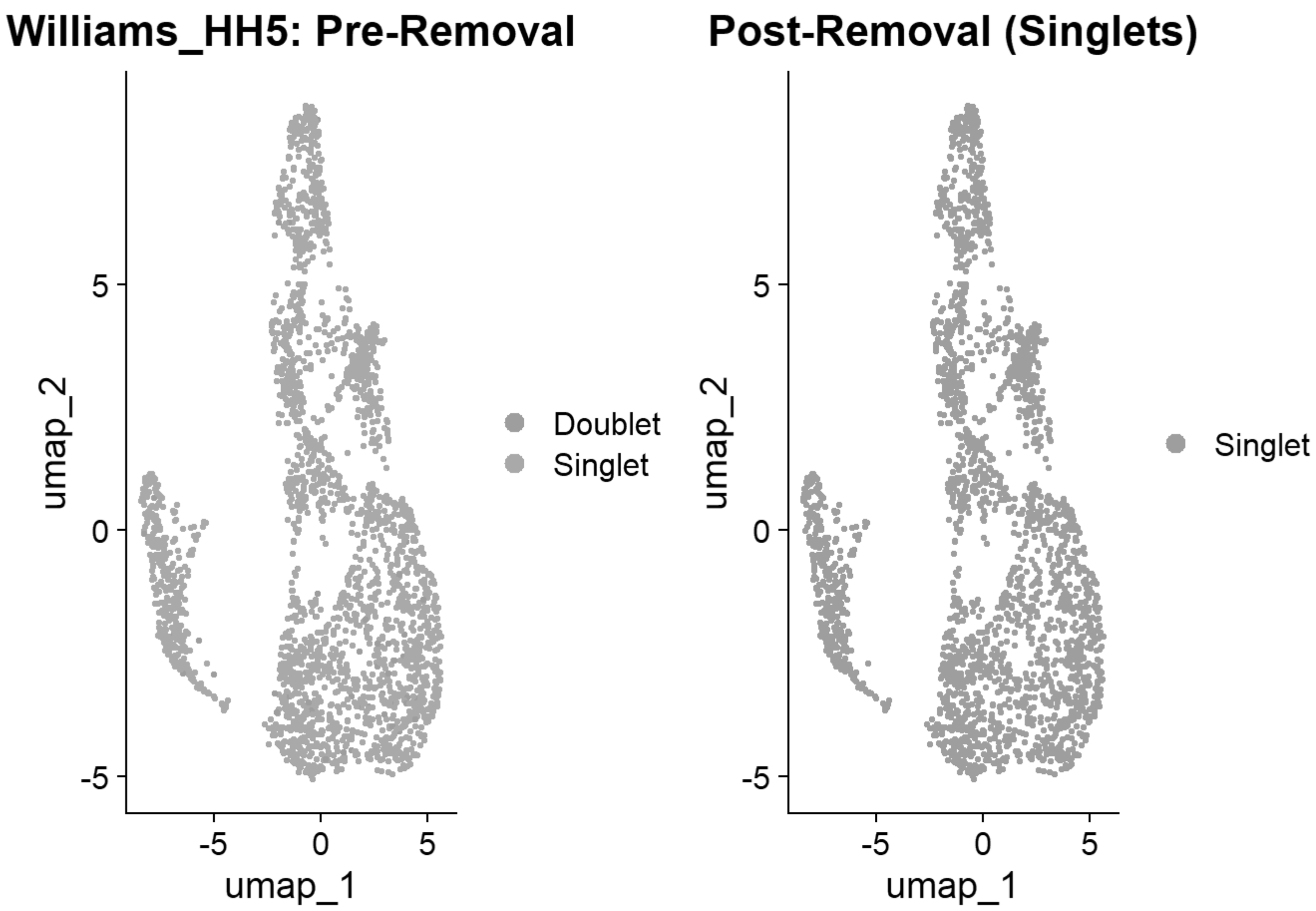


```
## [1] "Creating artificial doublets for pN = 5%"
## [1] "Creating Seurat object..."
## [1] "Normalizing Seurat object..."
```

```
## [1] "Finding variable genes..."
```

```
## [1] "Scaling data..."
```

```
## [1] "Running PCA..."
## [1] "Calculating PC distance matrix..."
## [1] "Defining neighborhoods..."
## [1] "Computing pANN across all pK..."
## [1] "pK = 0.005..."
## [1] "pK = 0.01..."
## [1] "pK = 0.02..."
## [1] "pK = 0.03..."
## [1] "pK = 0.04..."
## [1] "pK = 0.05..."
## [1] "pK = 0.06..."
## [1] "pK = 0.07..."
## [1] "pK = 0.08..."
## [1] "pK = 0.09..."
## [1] "pK = 0.1..."
## [1] "pK = 0.11..."
```

```
## [1] "pK = 0.12..."
## [1] "pK = 0.13..."
## [1] "pK = 0.14..."
## [1] "pK = 0.15..."
## [1] "pK = 0.16..."
## [1] "pK = 0.17..."
## [1] "pK = 0.18..."
## [1] "pK = 0.19..."
## [1] "pK = 0.2..."
## [1] "pK = 0.21..."
## [1] "pK = 0.22..."
## [1] "pK = 0.23..."
## [1] "pK = 0.24..."
## [1] "pK = 0.25..."
## [1] "pK = 0.26..."
## [1] "pK = 0.27..."
## [1] "pK = 0.28..."
## [1] "pK = 0.29..."
## [1] "pK = 0.3..."
## [1] "Creating artificial doublets for pN = 10%"
## [1] "Creating Seurat object..."
## [1] "Normalizing Seurat object..."
```

```
## [1] "Finding variable genes..."
```

```
## [1] "Scaling data..."
```

```
## [1] "Running PCA..."
## [1] "Calculating PC distance matrix..."
## [1] "Defining neighborhoods..."
## [1] "Computing pANN across all pK..."
## [1] "pK = 0.005..."
## [1] "pK = 0.01..."
## [1] "pK = 0.02..."
## [1] "pK = 0.03..."
## [1] "pK = 0.04..."
## [1] "pK = 0.05..."
## [1] "pK = 0.06..."
## [1] "pK = 0.07..."
## [1] "pK = 0.08..."
## [1] "pK = 0.09..."
## [1] "pK = 0.1..."
## [1] "pK = 0.11..."
## [1] "pK = 0.12..."
## [1] "pK = 0.13..."
## [1] "pK = 0.14..."
## [1] "pK = 0.15..."
## [1] "pK = 0.16..."
## [1] "pK = 0.17..."
## [1] "pK = 0.18..."
## [1] "pK = 0.19..."
## [1] "pK = 0.2..."
```

```
## [1] "pK = 0.21..."
## [1] "pK = 0.22..."
## [1] "pK = 0.23..."
## [1] "pK = 0.24..."
## [1] "pK = 0.25..."
## [1] "pK = 0.26..."
## [1] "pK = 0.27..."
## [1] "pK = 0.28..."
## [1] "pK = 0.29..."
## [1] "pK = 0.3..."
## [1] "Creating artificial doublets for pN = 15%"
## [1] "Creating Seurat object..."
## [1] "Normalizing Seurat object..."
```

```
## [1] "Finding variable genes..."
```

```
## [1] "Scaling data..."
```

```
## [1] "Running PCA..."
## [1] "Calculating PC distance matrix..."
## [1] "Defining neighborhoods..."
## [1] "Computing pANN across all pK..."
## [1] "pK = 0.005..."
## [1] "pK = 0.01..."
## [1] "pK = 0.02..."
## [1] "pK = 0.03..."
## [1] "pK = 0.04..."
## [1] "pK = 0.05..."
## [1] "pK = 0.06..."
## [1] "pK = 0.07..."
## [1] "pK = 0.08..."
## [1] "pK = 0.09..."
## [1] "pK = 0.1..."
## [1] "pK = 0.11..."
## [1] "pK = 0.12..."
## [1] "pK = 0.13..."
## [1] "pK = 0.14..."
## [1] "pK = 0.15..."
## [1] "pK = 0.16..."
## [1] "pK = 0.17..."
## [1] "pK = 0.18..."
## [1] "pK = 0.19..."
## [1] "pK = 0.2..."
## [1] "pK = 0.21..."
## [1] "pK = 0.22..."
## [1] "pK = 0.23..."
## [1] "pK = 0.24..."
## [1] "pK = 0.25..."
## [1] "pK = 0.26..."
## [1] "pK = 0.27..."
## [1] "pK = 0.28..."
## [1] "pK = 0.29..."
```

```
## [1] "pK = 0.3..."
## [1] "Creating artificial doublets for pN = 20%"
## [1] "Creating Seurat object..."
## [1] "Normalizing Seurat object..."
```

```
## [1] "Finding variable genes..."
```

```
## [1] "Scaling data..."
```

```
## [1] "Running PCA..."
## [1] "Calculating PC distance matrix..."
## [1] "Defining neighborhoods..."
## [1] "Computing pANN across all pK..."
## [1] "pK = 0.005..."
## [1] "pK = 0.01..."
## [1] "pK = 0.02..."
## [1] "pK = 0.03..."
## [1] "pK = 0.04..."
## [1] "pK = 0.05..."
## [1] "pK = 0.06..."
## [1] "pK = 0.07..."
## [1] "pK = 0.08..."
## [1] "pK = 0.09..."
## [1] "pK = 0.1..."
## [1] "pK = 0.11..."
## [1] "pK = 0.12..."
## [1] "pK = 0.13..."
## [1] "pK = 0.14..."
## [1] "pK = 0.15..."
## [1] "pK = 0.16..."
## [1] "pK = 0.17..."
## [1] "pK = 0.18..."
## [1] "pK = 0.19..."
## [1] "pK = 0.2..."
## [1] "pK = 0.21..."
## [1] "pK = 0.22..."
## [1] "pK = 0.23..."
## [1] "pK = 0.24..."
## [1] "pK = 0.25..."
## [1] "pK = 0.26..."
## [1] "pK = 0.27..."
## [1] "pK = 0.28..."
## [1] "pK = 0.29..."
## [1] "pK = 0.3..."
## [1] "Creating artificial doublets for pN = 25%"
## [1] "Creating Seurat object..."
## [1] "Normalizing Seurat object..."
```

```
## [1] "Finding variable genes..."
```

```
## [1] "Scaling data..."
```

```
## [1] "Running PCA..."
## [1] "Calculating PC distance matrix..."
## [1] "Defining neighborhoods..."
## [1] "Computing pANN across all pK..."
## [1] "pK = 0.005..."
## [1] "pK = 0.01..."
## [1] "pK = 0.02..."
## [1] "pK = 0.03..."
## [1] "pK = 0.04..."
## [1] "pK = 0.05..."
## [1] "pK = 0.06..."
## [1] "pK = 0.07..."
## [1] "pK = 0.08..."
## [1] "pK = 0.09..."
## [1] "pK = 0.1..."
## [1] "pK = 0.11..."
## [1] "pK = 0.12..."
## [1] "pK = 0.13..."
## [1] "pK = 0.14..."
## [1] "pK = 0.15..."
## [1] "pK = 0.16..."
## [1] "pK = 0.17..."
## [1] "pK = 0.18..."
## [1] "pK = 0.19..."
## [1] "pK = 0.2..."
## [1] "pK = 0.21..."
## [1] "pK = 0.22..."
## [1] "pK = 0.23..."
## [1] "pK = 0.24..."
## [1] "pK = 0.25..."
## [1] "pK = 0.26..."
## [1] "pK = 0.27..."
## [1] "pK = 0.28..."
## [1] "pK = 0.29..."
## [1] "pK = 0.3..."
## [1] "Creating artificial doublets for pN = 30%"
## [1] "Creating Seurat object..."
## [1] "Normalizing Seurat object..."
```

```
## [1] "Finding variable genes..."
```

```
## [1] "Scaling data..."
```

```
## [1] "Running PCA..."
## [1] "Calculating PC distance matrix..."
## [1] "Defining neighborhoods..."
## [1] "Computing pANN across all pK..."
## [1] "pK = 0.005..."
## [1] "pK = 0.01..."
## [1] "pK = 0.02..."
```

```
## [1] "pK = 0.03..."
## [1] "pK = 0.04..."
## [1] "pK = 0.05..."
## [1] "pK = 0.06..."
## [1] "pK = 0.07..."
## [1] "pK = 0.08..."
## [1] "pK = 0.09..."
## [1] "pK = 0.1..."
## [1] "pK = 0.11..."
## [1] "pK = 0.12..."
## [1] "pK = 0.13..."
## [1] "pK = 0.14..."
## [1] "pK = 0.15..."
## [1] "pK = 0.16..."
## [1] "pK = 0.17..."
## [1] "pK = 0.18..."
## [1] "pK = 0.19..."
## [1] "pK = 0.2..."
## [1] "pK = 0.21..."
## [1] "pK = 0.22..."
## [1] "pK = 0.23..."
## [1] "pK = 0.24..."
## [1] "pK = 0.25..."
## [1] "pK = 0.26..."
## [1] "pK = 0.27..."
## [1] "pK = 0.28..."
## [1] "pK = 0.29..."
## [1] "pK = 0.3..."
```

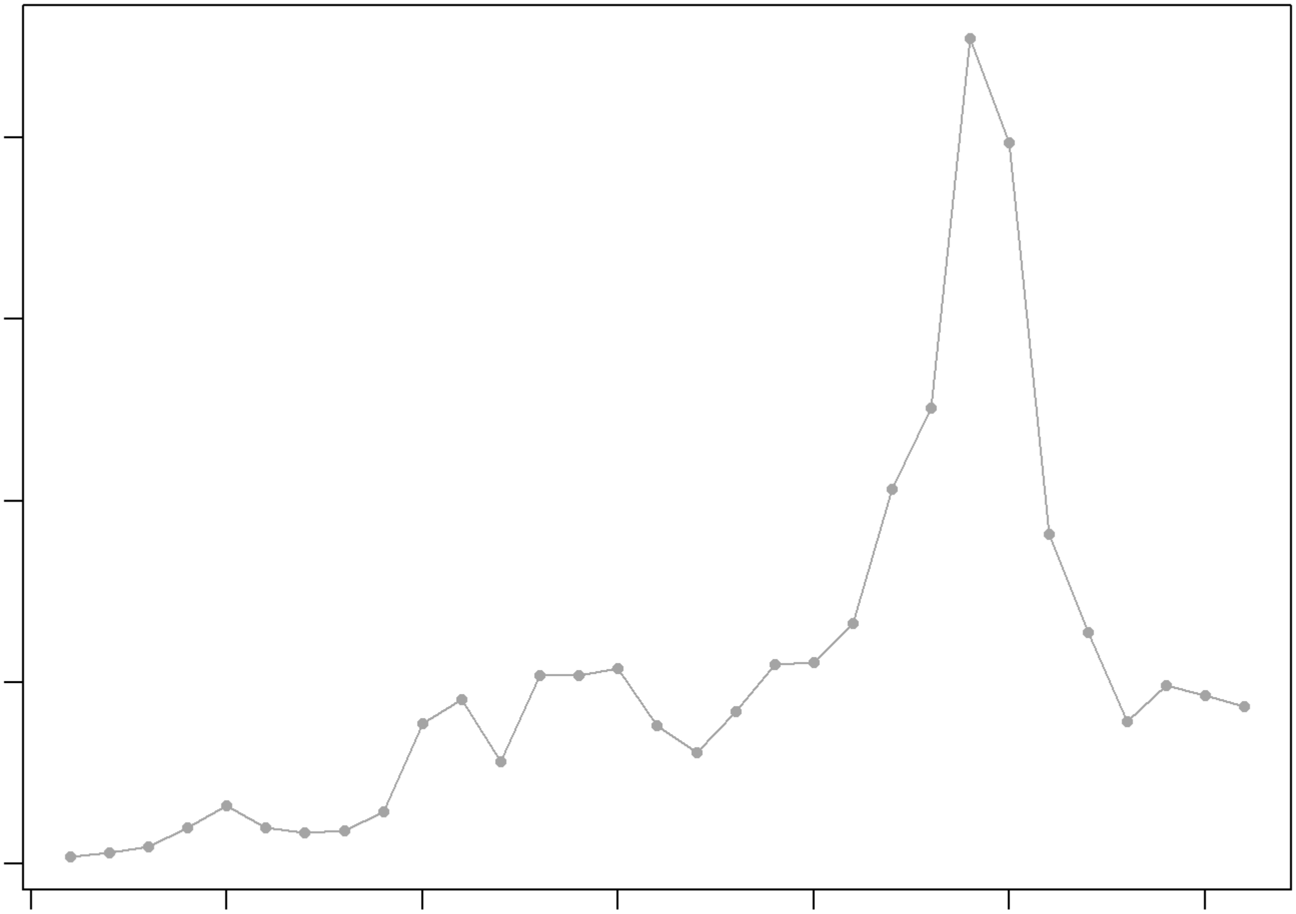

```
## NULL
## [1] "Creating 1442 artificial doublets..."
## [1] "Creating Seurat object..."
## [1] "Normalizing Seurat object..."
```

```
## [1] "Finding variable genes..."
```

```
## [1] "Scaling data..."
```

```
## [1] "Running PCA..."
## [1] "Calculating PC distance matrix..."
## [1] "Computing pANN..."
## [1] "Classifying doublets.."
```

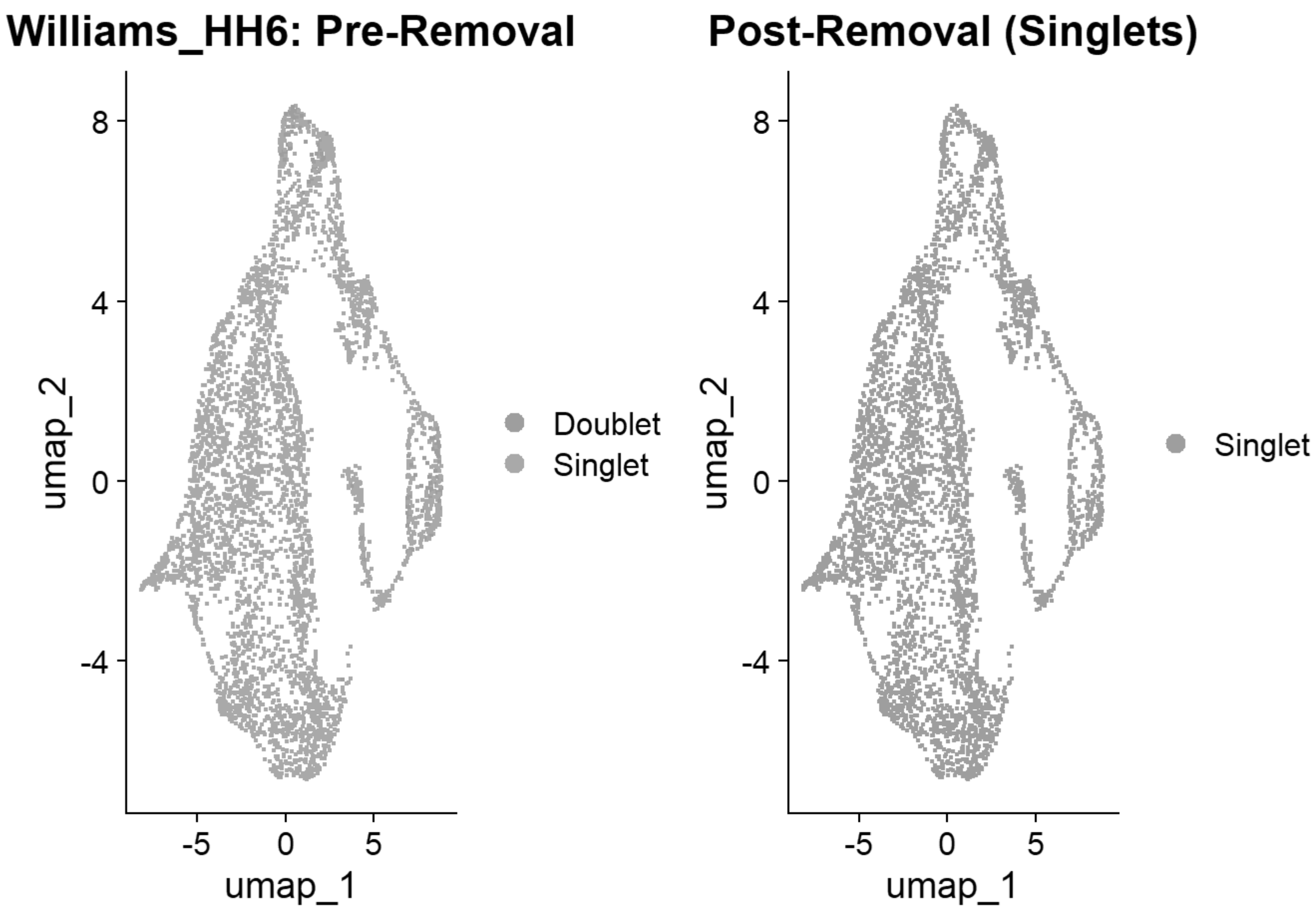


```
## [1] "Creating artificial doublets for pN = 5%"
## [1] "Creating Seurat object..."
## [1] "Normalizing Seurat object..."
```

```
## [1] "Finding variable genes..."
```

```
## [1] "Scaling data..."
```

```
## [1] "Running PCA..."
## [1] "Calculating PC distance matrix..."
## [1] "Defining neighborhoods..."
## [1] "Computing pANN across all pK..."
## [1] "pK = 0.005..."
## [1] "pK = 0.01..."
## [1] "pK = 0.02..."
## [1] "pK = 0.03..."
## [1] "pK = 0.04..."
## [1] "pK = 0.05..."
## [1] "pK = 0.06..."
## [1] "pK = 0.07..."
## [1] "pK = 0.08..."
## [1] "pK = 0.09..."
## [1] "pK = 0.1..."
## [1] "pK = 0.11..."
```

```
## [1] "pK = 0.12..."
## [1] "pK = 0.13..."
## [1] "pK = 0.14..."
## [1] "pK = 0.15..."
## [1] "pK = 0.16..."
## [1] "pK = 0.17..."
## [1] "pK = 0.18..."
## [1] "pK = 0.19..."
## [1] "pK = 0.2..."
## [1] "pK = 0.21..."
## [1] "pK = 0.22..."
## [1] "pK = 0.23..."
## [1] "pK = 0.24..."
## [1] "pK = 0.25..."
## [1] "pK = 0.26..."
## [1] "pK = 0.27..."
## [1] "pK = 0.28..."
## [1] "pK = 0.29..."
## [1] "pK = 0.3..."
## [1] "Creating artificial doublets for pN = 10%"
## [1] "Creating Seurat object..."
## [1] "Normalizing Seurat object..."
```

```
## [1] "Finding variable genes..."
```

```
## [1] "Scaling data..."
```

```
## [1] "Running PCA..."
## [1] "Calculating PC distance matrix..."
## [1] "Defining neighborhoods..."
## [1] "Computing pANN across all pK..."
## [1] "pK = 0.005..."
## [1] "pK = 0.01..."
## [1] "pK = 0.02..."
## [1] "pK = 0.03..."
## [1] "pK = 0.04..."
## [1] "pK = 0.05..."
## [1] "pK = 0.06..."
## [1] "pK = 0.07..."
## [1] "pK = 0.08..."
## [1] "pK = 0.09..."
## [1] "pK = 0.1..."
## [1] "pK = 0.11..."
## [1] "pK = 0.12..."
## [1] "pK = 0.13..."
## [1] "pK = 0.14..."
## [1] "pK = 0.15..."
## [1] "pK = 0.16..."
## [1] "pK = 0.17..."
## [1] "pK = 0.18..."
## [1] "pK = 0.19..."
## [1] "pK = 0.2..."
```

```
## [1] "pK = 0.21..."
## [1] "pK = 0.22..."
## [1] "pK = 0.23..."
## [1] "pK = 0.24..."
## [1] "pK = 0.25..."
## [1] "pK = 0.26..."
## [1] "pK = 0.27..."
## [1] "pK = 0.28..."
## [1] "pK = 0.29..."
## [1] "pK = 0.3..."
## [1] "Creating artificial doublets for pN = 15%"
## [1] "Creating Seurat object..."
## [1] "Normalizing Seurat object..."
```

```
## [1] "Finding variable genes..."
```

```
## [1] "Scaling data..."
```

```
## [1] "Running PCA..."
## [1] "Calculating PC distance matrix..."
## [1] "Defining neighborhoods..."
## [1] "Computing pANN across all pK..."
## [1] "pK = 0.005..."
## [1] "pK = 0.01..."
## [1] "pK = 0.02..."
## [1] "pK = 0.03..."
## [1] "pK = 0.04..."
## [1] "pK = 0.05..."
## [1] "pK = 0.06..."
## [1] "pK = 0.07..."
## [1] "pK = 0.08..."
## [1] "pK = 0.09..."
## [1] "pK = 0.1..."
## [1] "pK = 0.11..."
## [1] "pK = 0.12..."
## [1] "pK = 0.13..."
## [1] "pK = 0.14..."
## [1] "pK = 0.15..."
## [1] "pK = 0.16..."
## [1] "pK = 0.17..."
## [1] "pK = 0.18..."
## [1] "pK = 0.19..."
## [1] "pK = 0.2..."
## [1] "pK = 0.21..."
## [1] "pK = 0.22..."
## [1] "pK = 0.23..."
## [1] "pK = 0.24..."
## [1] "pK = 0.25..."
## [1] "pK = 0.26..."
## [1] "pK = 0.27..."
## [1] "pK = 0.28..."
## [1] "pK = 0.29..."
```

```
## [1] "pK = 0.3..."
## [1] "Creating artificial doublets for pN = 20%"
## [1] "Creating Seurat object..."
## [1] "Normalizing Seurat object..."
```

```
## [1] "Finding variable genes..."
```

```
## [1] "Scaling data..."
```

```
## [1] "Running PCA..."
## [1] "Calculating PC distance matrix..."
## [1] "Defining neighborhoods..."
## [1] "Computing pANN across all pK..."
## [1] "pK = 0.005..."
## [1] "pK = 0.01..."
## [1] "pK = 0.02..."
## [1] "pK = 0.03..."
## [1] "pK = 0.04..."
## [1] "pK = 0.05..."
## [1] "pK = 0.06..."
## [1] "pK = 0.07..."
## [1] "pK = 0.08..."
## [1] "pK = 0.09..."
## [1] "pK = 0.1..."
## [1] "pK = 0.11..."
## [1] "pK = 0.12..."
## [1] "pK = 0.13..."
## [1] "pK = 0.14..."
## [1] "pK = 0.15..."
## [1] "pK = 0.16..."
## [1] "pK = 0.17..."
## [1] "pK = 0.18..."
## [1] "pK = 0.19..."
## [1] "pK = 0.2..."
## [1] "pK = 0.21..."
## [1] "pK = 0.22..."
## [1] "pK = 0.23..."
## [1] "pK = 0.24..."
## [1] "pK = 0.25..."
## [1] "pK = 0.26..."
## [1] "pK = 0.27..."
## [1] "pK = 0.28..."
## [1] "pK = 0.29..."
## [1] "pK = 0.3..."
## [1] "Creating artificial doublets for pN = 25%"
## [1] "Creating Seurat object..."
## [1] "Normalizing Seurat object..."
```

```
## [1] "Finding variable genes..."
```

```
## [1] "Scaling data..."
```

```
## [1] "Running PCA..."
## [1] "Calculating PC distance matrix..."
## [1] "Defining neighborhoods..."
## [1] "Computing pANN across all pK..."
## [1] "pK = 0.005..."
## [1] "pK = 0.01..."
## [1] "pK = 0.02..."
## [1] "pK = 0.03..."
## [1] "pK = 0.04..."
## [1] "pK = 0.05..."
## [1] "pK = 0.06..."
## [1] "pK = 0.07..."
## [1] "pK = 0.08..."
## [1] "pK = 0.09..."
## [1] "pK = 0.1..."
## [1] "pK = 0.11..."
## [1] "pK = 0.12..."
## [1] "pK = 0.13..."
## [1] "pK = 0.14..."
## [1] "pK = 0.15..."
## [1] "pK = 0.16..."
## [1] "pK = 0.17..."
## [1] "pK = 0.18..."
## [1] "pK = 0.19..."
## [1] "pK = 0.2..."
## [1] "pK = 0.21..."
## [1] "pK = 0.22..."
## [1] "pK = 0.23..."
## [1] "pK = 0.24..."
## [1] "pK = 0.25..."
## [1] "pK = 0.26..."
## [1] "pK = 0.27..."
## [1] "pK = 0.28..."
## [1] "pK = 0.29..."
## [1] "pK = 0.3..."
## [1] "Creating artificial doublets for pN = 30%"
## [1] "Creating Seurat object..."
## [1] "Normalizing Seurat object..."
```

```
## [1] "Finding variable genes..."
```

```
## [1] "Scaling data..."
```

```
## [1] "Running PCA..."
## [1] "Calculating PC distance matrix..."
## [1] "Defining neighborhoods..."
## [1] "Computing pANN across all pK..."
## [1] "pK = 0.005..."
## [1] "pK = 0.01..."
## [1] "pK = 0.02..."
```

```
## [1] "pK = 0.03..."
## [1] "pK = 0.04..."
## [1] "pK = 0.05..."
## [1] "pK = 0.06..."
## [1] "pK = 0.07..."
## [1] "pK = 0.08..."
## [1] "pK = 0.09..."
## [1] "pK = 0.1..."
## [1] "pK = 0.11..."
## [1] "pK = 0.12..."
## [1] "pK = 0.13..."
## [1] "pK = 0.14..."
## [1] "pK = 0.15..."
## [1] "pK = 0.16..."
## [1] "pK = 0.17..."
## [1] "pK = 0.18..."
## [1] "pK = 0.19..."
## [1] "pK = 0.2..."
## [1] "pK = 0.21..."
## [1] "pK = 0.22..."
## [1] "pK = 0.23..."
## [1] "pK = 0.24..."
## [1] "pK = 0.25..."
## [1] "pK = 0.26..."
## [1] "pK = 0.27..."
## [1] "pK = 0.28..."
## [1] "pK = 0.29..."
## [1] "pK = 0.3..."
```

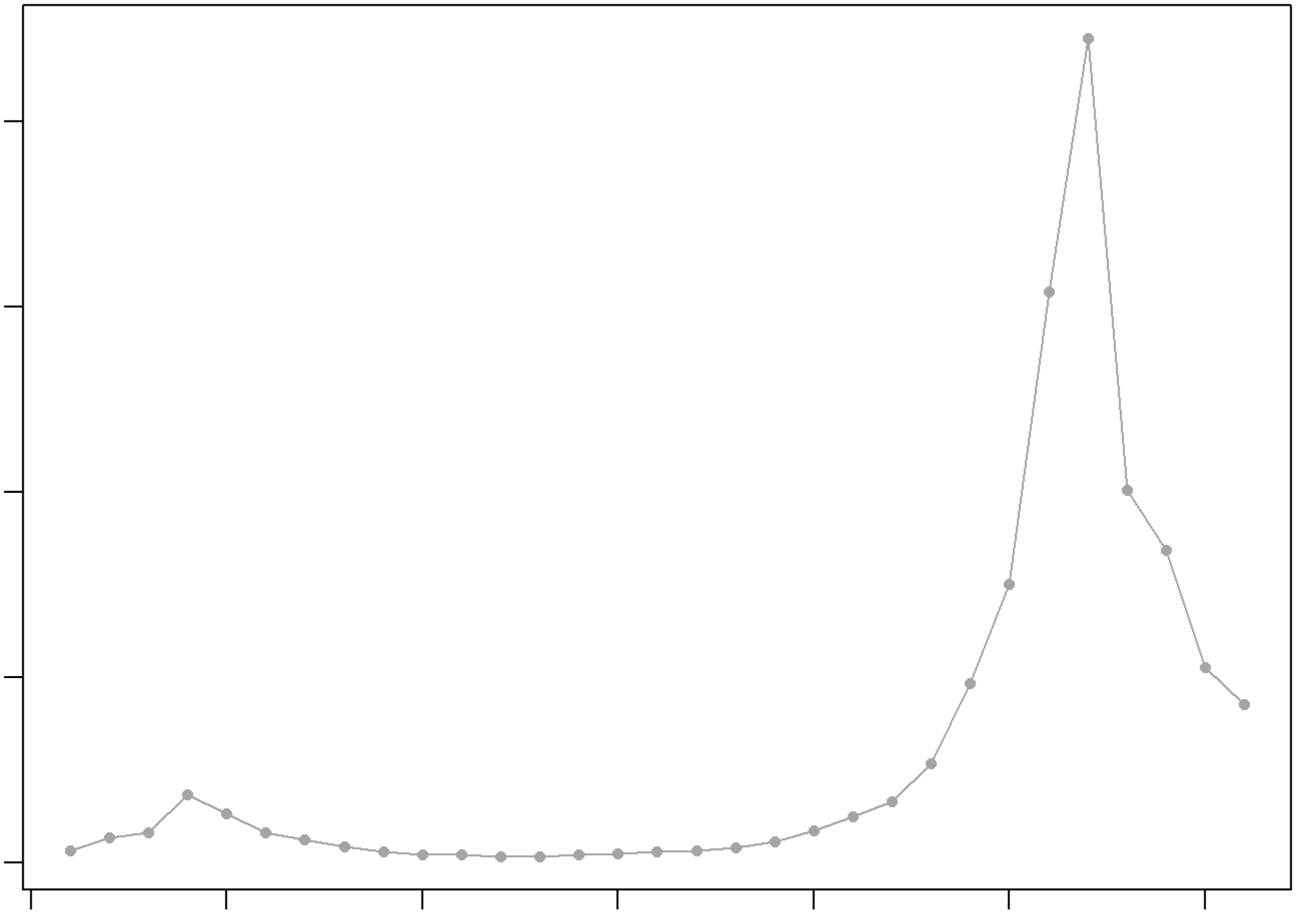

```
## NULL
## [1] "Creating 2049 artificial doublets..."
## [1] "Creating Seurat object..."
## [1] "Normalizing Seurat object..."
```

```
## [1] "Finding variable genes..."
```

```
## [1] "Scaling data..."
```

```
## [1] "Running PCA..."
## [1] "Calculating PC distance matrix..."
## [1] "Computing pANN..."
## [1] "Classifying doublets.."
```

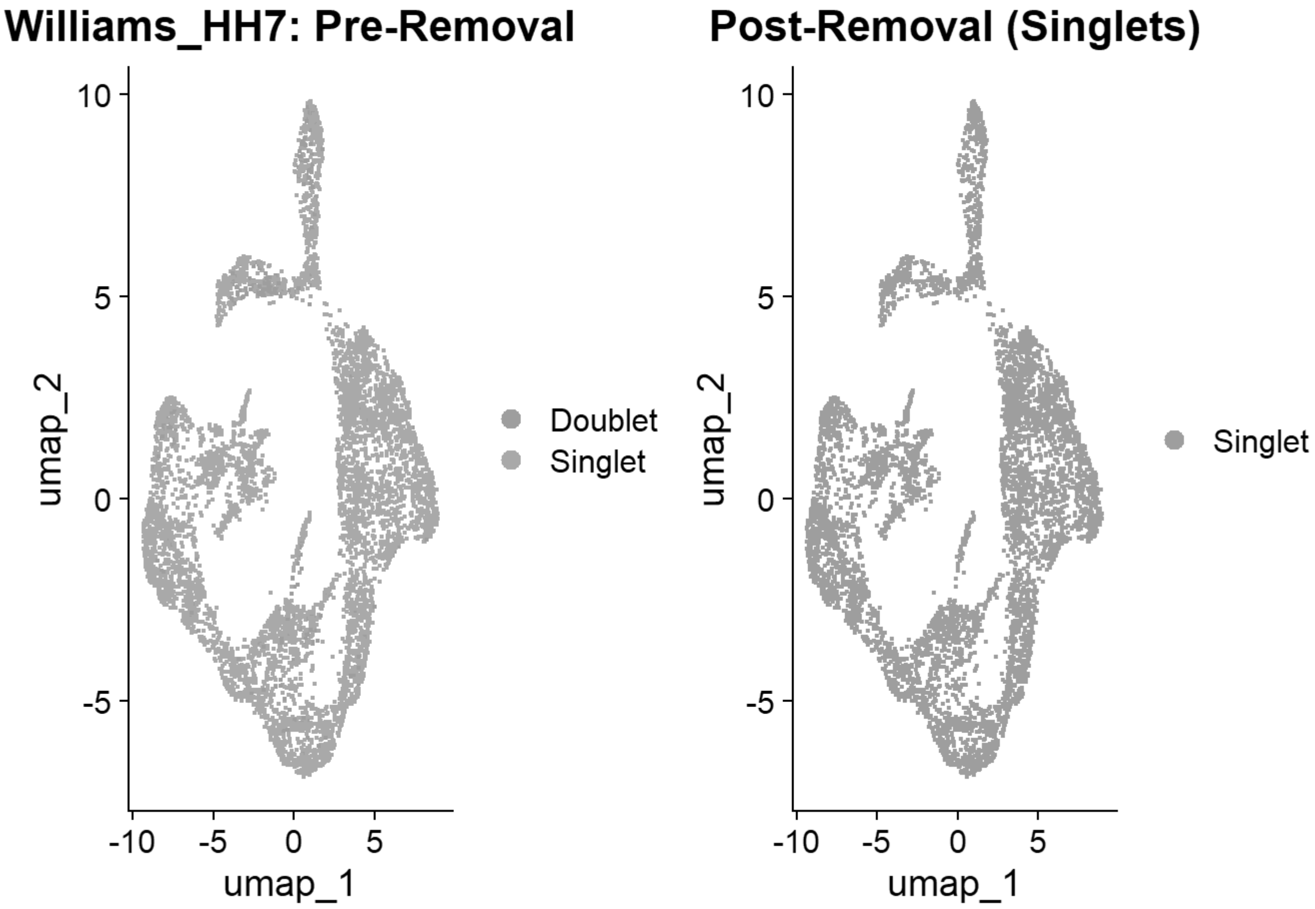


### 4. Global Integration & Annotation

```
# Compile and render the Quality Control Tracking Table
qc_summary_df <- do.call(rbind, qc_tracker)
rownames(qc_summary_df) <- NULL
kable(qc_summary_df, caption = "Cell Counts Recovered After Sequential Filtering Steps")
```

Cell Counts Recovered After Sequential Filtering Steps

| Sample | Pre_QC | Post_QC | Post_DoubletFinder |
|---|---:|---:|---:|
| Pajanoja2023_HH5_1 | 12943 | 12849 | 11528 |
| Pajanoja2023_HH5_2 | 5535 | 5530 | 5285 |
| Pajanoja2023_HH7_1 | 6141 | 5786 | 5518 |
| Pajanoja2023_HH7_2 | 6225 | 6153 | 5850 |
| Pajanoja2023_HH8_1 | 10012 | 9782 | 9016 |
| Pajanoja2023_HH8_2 | 23489 | 22688 | 18570 |
| Pajanoja2023_HH9_1 | 16839 | 16416 | 14260 |
| Pajanoja2023_HH9_2 | 16755 | 16069 | 14003 |

| | | | |
|---|---|---|---|
| Pajanoja2025_HH11 | 9801 | 9610 | 8871 |
| Williams_HH4 | 2187 | 2184 | 2146 |
| Williams_HH5 | 6411 | 2523 | 2472 |
| Williams_HH6 | 4333 | 4327 | 4177 |
| Williams_HH7 | 6149 | 6148 | 5846 |

```
# Merge all clean singlets
all_samples <- c(pajanoja_list, williams_list)
chick <- merge(all_samples[[1]], y = all_samples[2:length(all_samples)])
saveRDS(chick, "E:\\Rogers Lab\\Rogers NC EMT\\Global_chick_v2.rds")
rm(pajanoja_list, williams_list, all_samples, tmp_seu, seu, sc, sweep.res,bmcvn); gc()
```

```
##             used   (Mb) gc trigger    (Mb)   max used    (Mb)
## Ncells   4399027  235.0   17064884   911.4   21331105  1139.3
## Vcells 795694595 6070.7 2326593630 17750.6 2423451037 18489.5
```

```
# Standard Pre-processing
chick <- JoinLayers(chick)
chick[["RNA"]] <- split(chick[["RNA"]], f = chick$sample_id)
```

```
## Splitting 'counts', 'data' layers. Not splitting 'scale.data'. If you would like to split o
ther layers, set in `layers` argument.
```

```
chick
```

```
## An object of class Seurat
## 18523 features across 107542 samples within 1 assay
## Active assay: RNA (18523 features, 2000 variable features)
##  27 layers present: counts.Pajanoja2023_HH5_1, counts.Pajanoja2023_HH5_2, counts.Pajanoja20
23_HH7_1, counts.Pajanoja2023_HH7_2, counts.Pajanoja2023_HH8_1, counts.Pajanoja2023_HH8_2, cou
nts.Pajanoja2023_HH9_1, counts.Pajanoja2023_HH9_2, counts.Pajanoja2025_HH11, counts.Williams_H
H4, counts.Williams_HH5, counts.Williams_HH6, counts.Williams_HH7, scale.data, data.Pajanoja20
23_HH5_1, data.Pajanoja2023_HH5_2, data.Pajanoja2023_HH7_1, data.Pajanoja2023_HH7_2, data.Paja
noja2023_HH8_1, data.Pajanoja2023_HH8_2, data.Pajanoja2023_HH9_1, data.Pajanoja2023_HH9_2, dat
a.Pajanoja2025_HH11, data.Williams_HH4, data.Williams_HH5, data.Williams_HH6, data.Williams_HH
7
```

```
chick <- NormalizeData(chick)
```

```
## Normalizing layer: counts.Pajanoja2023_HH5_1
```

```
## Normalizing layer: counts.Pajanoja2023_HH5_2
```

```
## Normalizing layer: counts.Pajanoja2023_HH7_1
```

```
## Normalizing layer: counts.Pajanoja2023_HH7_2
```

```
## Normalizing layer: counts.Pajanoja2023_HH8_1
```

```
## Normalizing layer: counts.Pajanoja2023_HH8_2
```

```
## Normalizing layer: counts.Pajanoja2023_HH9_1
```

```
## Normalizing layer: counts.Pajanoja2023_HH9_2
```

```
## Normalizing layer: counts.Pajanoja2025_HH11
```

```
## Normalizing layer: counts.Williams_HH4
```

```
## Normalizing layer: counts.Williams_HH5
```

```
## Normalizing layer: counts.Williams_HH6
```

```
## Normalizing layer: counts.Williams_HH7
```

```
chick <- CellCycleScoring(chick, s.features = cc.genes$s.genes, g2m.features = cc.genes$g2m.ge
nes)
chick <- FindVariableFeatures(chick)
```

```
## Finding variable features for layer counts.Pajanoja2023_HH5_1
```

```
## Finding variable features for layer counts.Pajanoja2023_HH5_2
```

```
## Finding variable features for layer counts.Pajanoja2023_HH7_1
```

```
## Finding variable features for layer counts.Pajanoja2023_HH7_2
```

```
## Finding variable features for layer counts.Pajanoja2023_HH8_1
```

```
## Finding variable features for layer counts.Pajanoja2023_HH8_2
```

```
## Finding variable features for layer counts.Pajanoja2023_HH9_1
```

```
## Finding variable features for layer counts.Pajanoja2023_HH9_2
```

```
## Finding variable features for layer counts.Pajanoja2025_HH11
```

```
## Finding variable features for layer counts.Williams_HH4
```

```
## Finding variable features for layer counts.Williams_HH5
```

```
## Finding variable features for layer counts.Williams_HH6
```

```
## Finding variable features for layer counts.Williams_HH7
```

```
chick <- ScaleData(chick, vars.to.regress = c("percent.mt", "S.Score", "G2M.Score"))
```

```
## Regressing out percent.mt, S.Score, G2M.Score
```

```
## Centering and scaling data matrix
```

```
chick <- RunPCA(chick, npcs = 50, verbose = FALSE)

# Harmony Integration
chick <- IntegrateLayers(object = chick, method = HarmonyIntegration, orig.reduction = "pca",
new.reduction = "harmony", assay = "RNA", group.by = "sample_id", verbose = TRUE)
```

```
## The `features` argument is ignored by `HarmonyIntegration`.
## This message is displayed once per session.
```

```
## Transposing data matrix
```

```
## Using automatic lambda estimation
```

```
## Initializing state using k-means centroids initialization
```

```
## Harmony 1/10
```

```
## Harmony 2/10
```

```
## Harmony 3/10
```

```
## Harmony 4/10
```

```
## Harmony 5/10
```

```
## Harmony converged after 5 iterations
```

```
chick <- FindNeighbors(chick, reduction = "harmony", dims = 1:30)
```

```
## Computing nearest neighbor graph
```

```
## Computing SNN
```

```
chick <- RunUMAP(chick, reduction = "harmony", dims = 1:30)
```

```
## 22:13:50 UMAP embedding parameters a = 0.9922 b = 1.112
```

```
## 22:13:50 Read 107542 rows and found 30 numeric columns
```

```
## 22:13:50 Using Annoy for neighbor search, n_neighbors = 30
```

```
## 22:13:50 Building Annoy index with metric = cosine, n_trees = 50
```

```
## 0%   10   20   30   40   50   60   70   80   90   100%
```

```
## [----|----|----|----|----|----|----|----|----|----|
```

```
## **************************************************|
## 22:13:59 Writing NN index file to temp file C:\Users\RANEES~1\AppData\Local\Temp\RtmpqcWgVb
\file6a2c6a4d5741
## 22:13:59 Searching Annoy index using 1 thread, search_k = 3000
## 22:14:35 Annoy recall = 100%
## 22:14:36 Commencing smooth kNN distance calibration using 1 thread with target n_neighbors
= 30
## 22:14:41 Initializing from normalized Laplacian + noise (using RSpectra)
## 22:14:44 Commencing optimization for 200 epochs, with 5173630 positive edges
## 22:14:44 Using rng type: pcg
## 22:16:23 Optimization finished
```

```
chick <- FindClusters(chick, resolution = 0.4)
```

```
## Modularity Optimizer version 1.3.0 by Ludo Waltman and Nees Jan van Eck
##
## Number of nodes: 107542
## Number of edges: 3064558
##
## Running Louvain algorithm...
## Maximum modularity in 10 random starts: 0.9250
## Number of communities: 18
## Elapsed time: 29 seconds
```

```
# Run QC Sanity Tester
qc_tester(chick)
```

```
## Rasterizing points since number of points exceeds 100,000.
## To disable this behavior set `raster=FALSE`
## Rasterizing points since number of points exceeds 100,000.
## To disable this behavior set `raster=FALSE`
## Rasterizing points since number of points exceeds 100,000.
## To disable this behavior set `raster=FALSE`
```

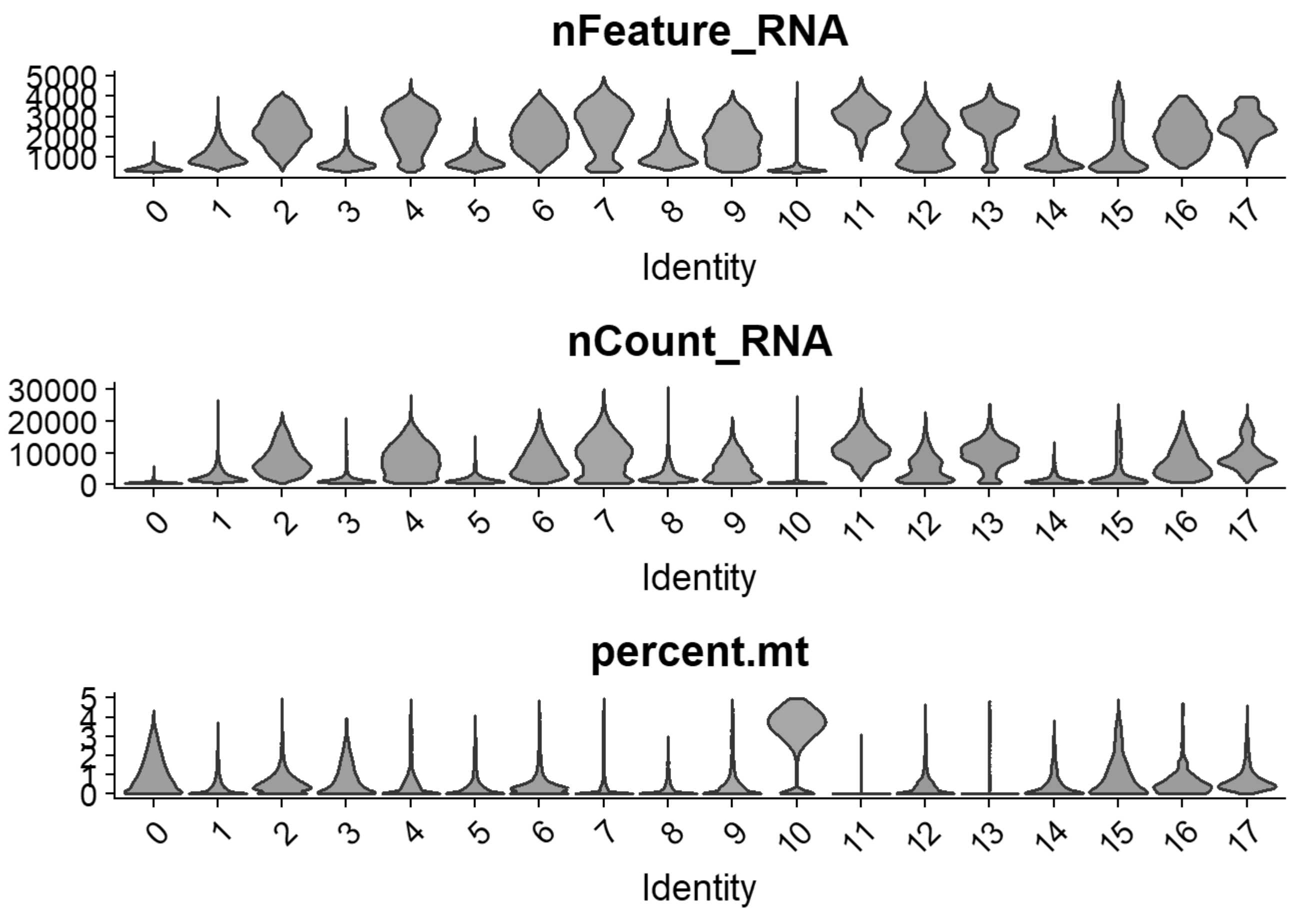


```
cols= grep("^RNA_snn_res.",names(chick@meta.data))
chick@meta.data[cols] <- NULL
#Picking higher resolution to remove low quality clusters (likely 0, 10)
chick <- FindClusters(chick, resolution = 1.0, verbose = TRUE)
```

```
## Modularity Optimizer version 1.3.0 by Ludo Waltman and Nees Jan van Eck
##
## Number of nodes: 107542
## Number of edges: 3064558
##
## Running Louvain algorithm...
## Maximum modularity in 10 random starts: 0.8822
## Number of communities: 27
## Elapsed time: 27 seconds
```

```
p6=DimPlot(chick, label = TRUE)+NoLegend()
```

```
## Rasterizing points since number of points exceeds 100,000.
## To disable this behavior set `raster=FALSE`
```

```
print(p6)
```

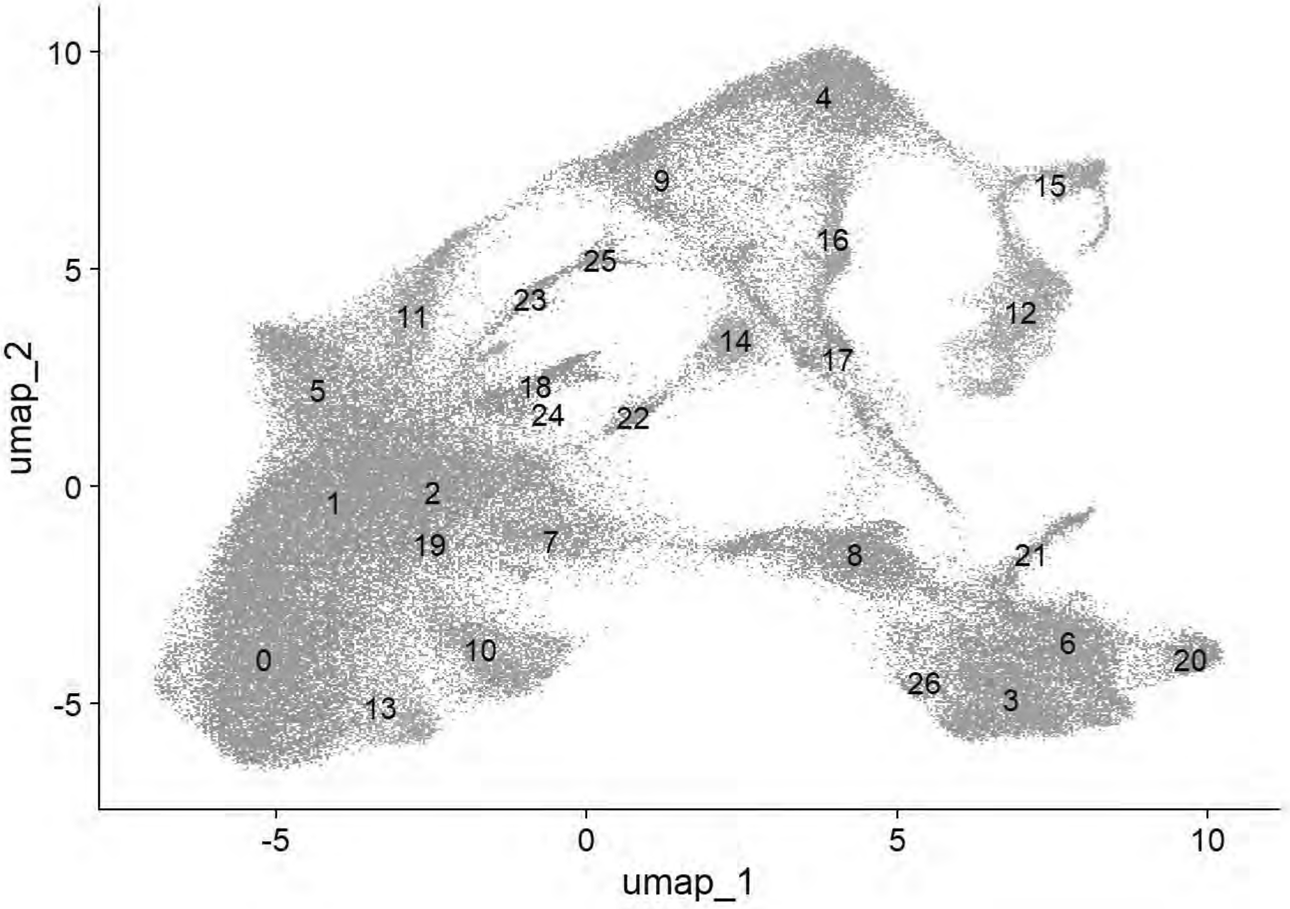


```
p6=DimPlot(chick, group.by = "ref")
```

```
## Rasterizing points since number of points exceeds 100,000.
## To disable this behavior set `raster=FALSE`
```

```
print(p6)
```

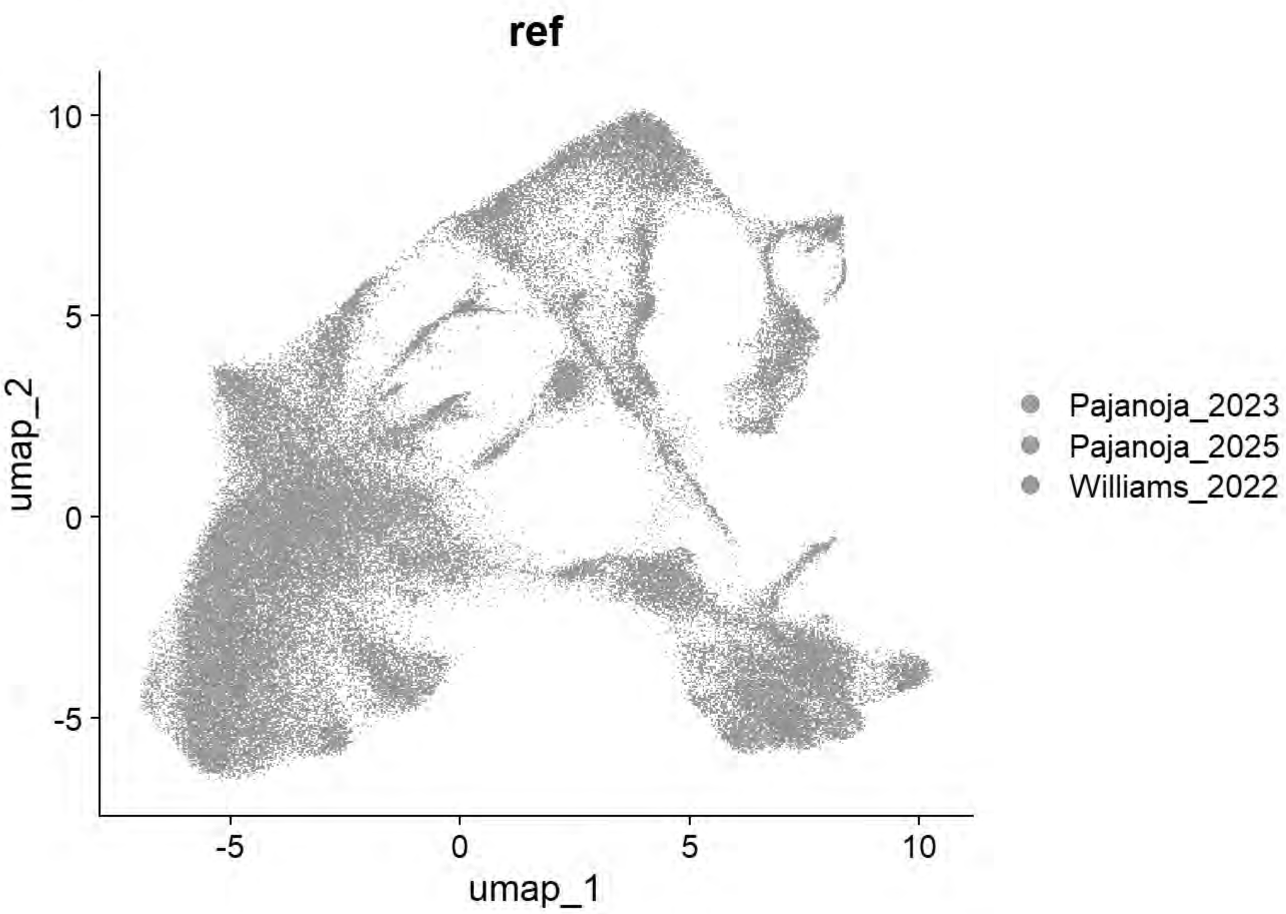


```
p6=DimPlot(chick, group.by = "Phase")
```

```
## Rasterizing points since number of points exceeds 100,000.
## To disable this behavior set `raster=FALSE`
```

```
print(p6)
```

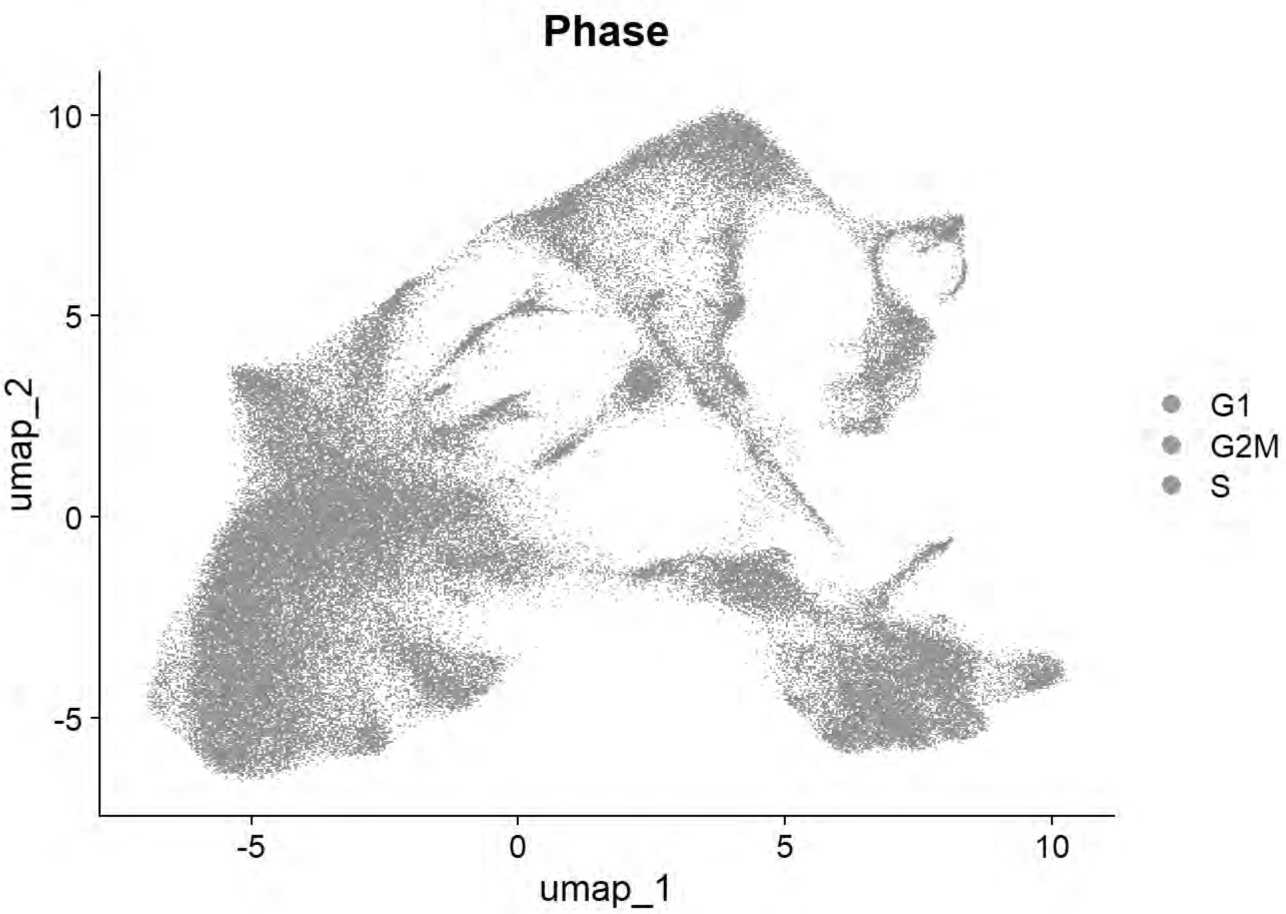


```
table(chick$seurat_clusters,chick$sample_id)
```

```
##
##      Pajanoja2023_HH5_1 Pajanoja2023_HH5_2 Pajanoja2023_HH7_1
##   0                1369                251                896
##   1                1295                469                735
##   2                 526                268                182
##   3                1706                676                429
##   4                 579                351                319
##   5                 285                355                235
##   6                1026                318                578
##   7                 589                799                490
##   8                1373                489                307
##   9                 208                 97                132
##   10                445                 76                163
##   11                156                151                173
##   12                845                305                197
##   13                 23                  3                 46
##   14                 92                 43                  3
##   15                351                185                108
##   16                 16                 45                  0
##   17                  0                  4                  0
##   18                356                191                 71
##   19                 28                123                327
##   20                178                 66                 86
```

```
##   21                 15                  2                 21
##   22                 51                  8                  2
##   23                  2                  4                  5
##   24                 11                  5                  5
##   25                  2                  1                  7
##   26                  1                  0                  1
##    
##      Pajanoja2023_HH7_2 Pajanoja2023_HH8_1 Pajanoja2023_HH8_2
##   0                 516               1089               3485
##   1                 567                928               2655
##   2                 199                187               3148
##   3                 799                907                273
##   4                 423                534                634
##   5                 371                458               1482
##   6                 428                230                525
##   7                 624                912                182
##   8                 370                575                156
##   9                 229                374                626
##   10                133                347                670
##   11                264                562               1274
##   12                164                401                300
##   13                 30                196                466
##   14                  8                  6               1412
##   15                175                156                129
##   16                  1                  0                126
##   17                  1                  0                  0
##   18                112                204                196
##   19                213                460                  2
##   20                162                181                 53
##   21                 39                157                106
##   22                  5                  2                373
##   23                  8                 74                169
##   24                  3                 23                  2
##   25                  6                 53                126
##   26                  0                  0                  0
##    
##      Pajanoja2023_HH9_1 Pajanoja2023_HH9_2 Pajanoja2025_HH11 Williams_HH4
##   0                3453               3877              1525            9
##   1                3485               3190              1037           62
##   2                1439               1535               213           21
##   3                 513                272              1572          204
##   4                 531                381               667          105
##   5                1022               1341               217            4
##   6                 379                 49               783          142
##   7                 114                121               147          244
##   8                  15                120               319          275
##   9                  83                 57                42          307
##   10                846                662               513           32
##   11                556                314                48           37
##   12                 11                 53               166          222
##   13                697                838               268            0
##   14                 67                148               125           96
##   15                 76                 96                58           40
##   16                  1                  5                 3           58
```

```
##   17                  0                   0                  0             74
##   18                238                 206                116             13
##   19                 65                  57                 38              1
##   20                 90                  73                290             10
##   21                146                 112                560              1
##   22                166                 203                 26             31
##   23                227                 239                 61              6
##   24                  7                   4                  9            113
##   25                 33                  50                 68             38
##   26                  0                   0                  0              1
##
##      Williams_HH5 Williams_HH6 Williams_HH7
##   0            11           13           13
##   1            32           12           15
##   2           106           13           43
##   3            74           13          163
##   4           411          611         1294
##   5            51           11           43
##   6            20           13          152
##   7            44           46          105
##   8            56          103          187
##   9           305         1017          709
##   10            3            4            4
##   11           12           26           42
##   12          297          281          336
##   13            0            0            0
##   14           62           77           39
##   15           38          174          390
##   16          360          530          830
##   17          258          775          863
##   18           24           30           30
##   19            2            0            1
##   20            3           17           86
##   21            0            1           14
##   22           18            3           22
##   23            2           22            3
##   24          177          178          159
##   25           21          112           55
##   26           85           95          248
```

```
qc_tester(chick)
```

```
## Rasterizing points since number of points exceeds 100,000.
## To disable this behavior set `raster=FALSE`
## Rasterizing points since number of points exceeds 100,000.
## To disable this behavior set `raster=FALSE`
## Rasterizing points since number of points exceeds 100,000.
## To disable this behavior set `raster=FALSE`
```

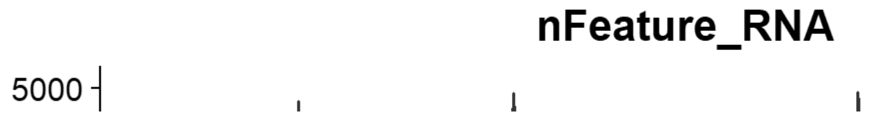

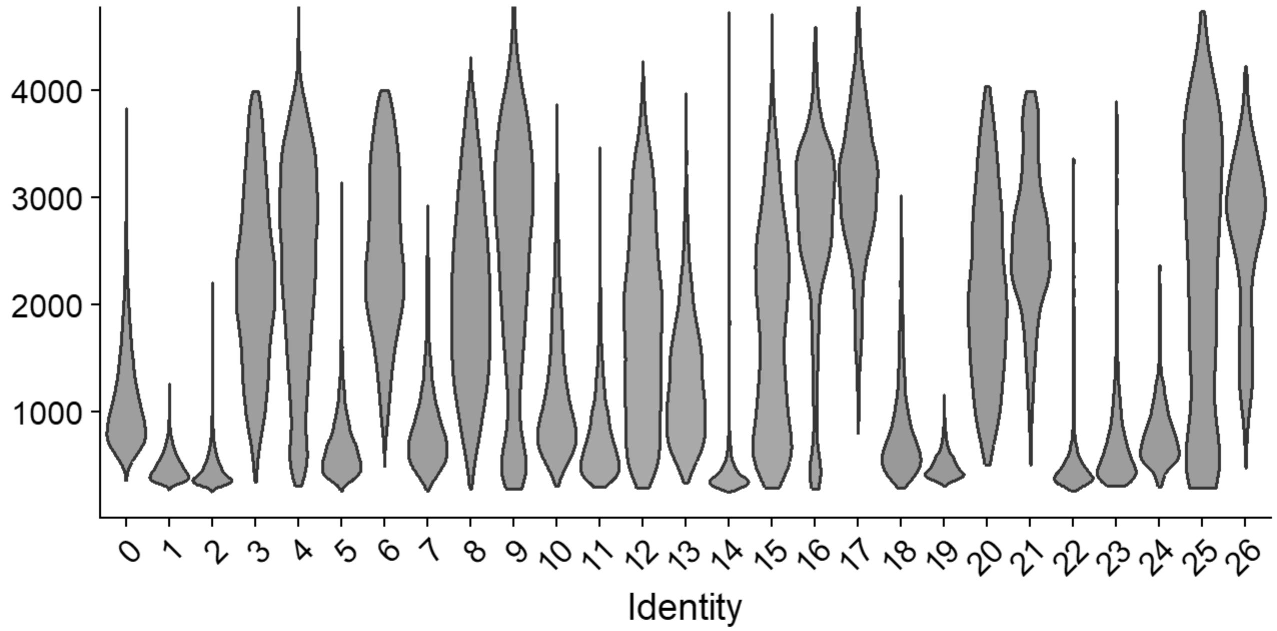


**nCount_RNA**

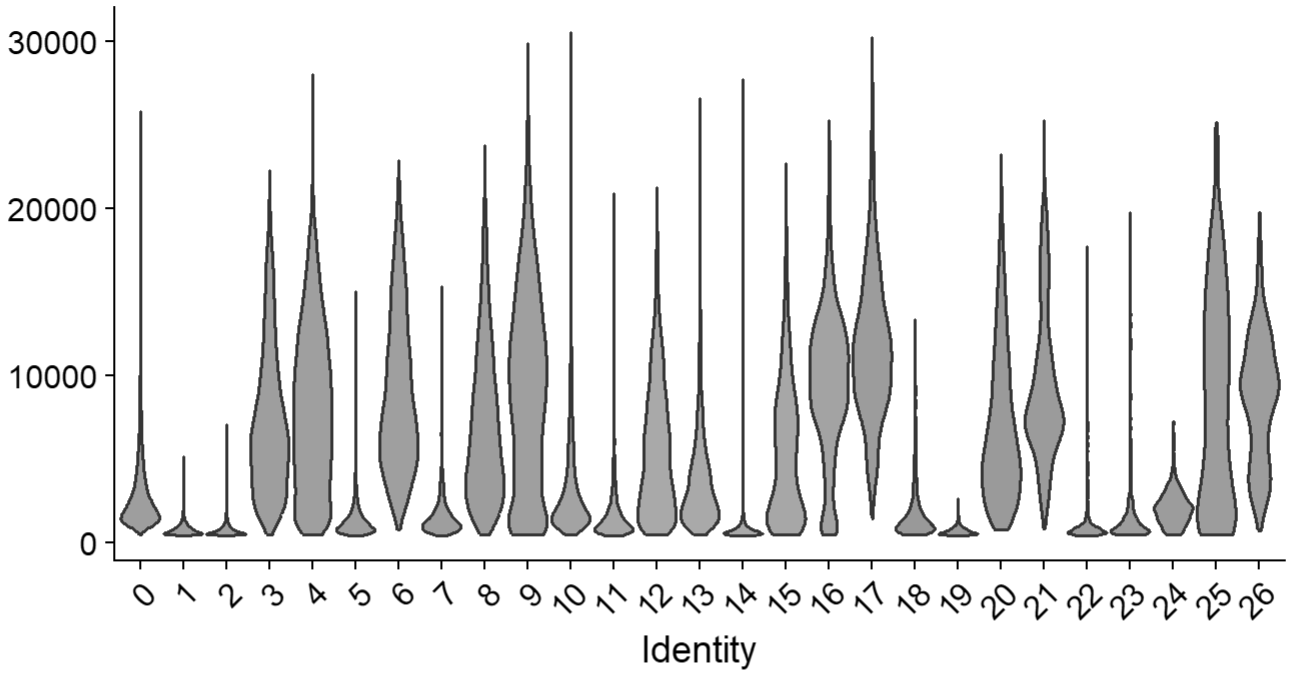


**percent.mt**

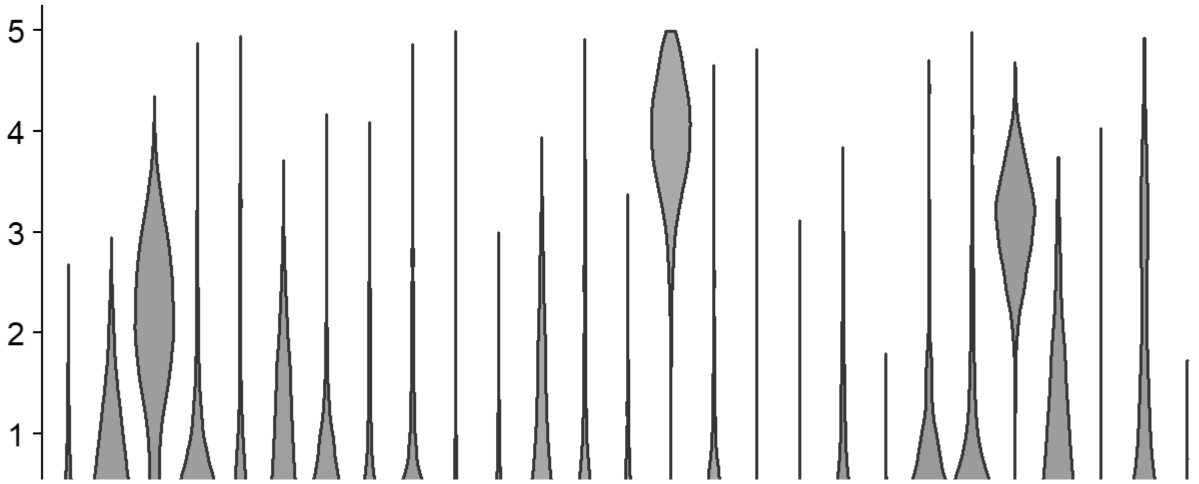

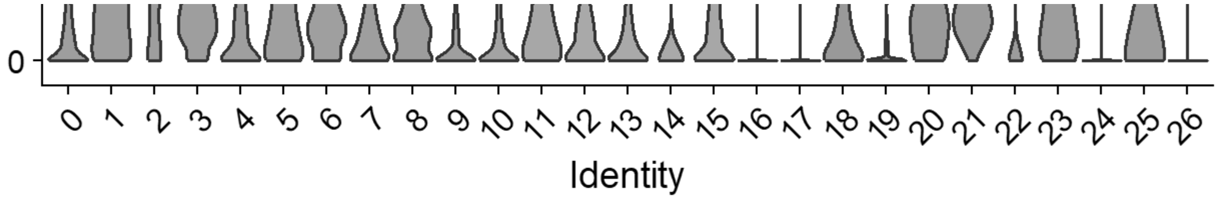


```r
# Williams: Neural plate/tube
FeaturePlot(chick, features = c("CLDN1","FRZB","DNMT3A"))
```

```
## Rasterizing points since number of points exceeds 100,000.
## To disable this behavior set `raster=FALSE`
## Rasterizing points since number of points exceeds 100,000.
## To disable this behavior set `raster=FALSE`
## Rasterizing points since number of points exceeds 100,000.
## To disable this behavior set `raster=FALSE`
```

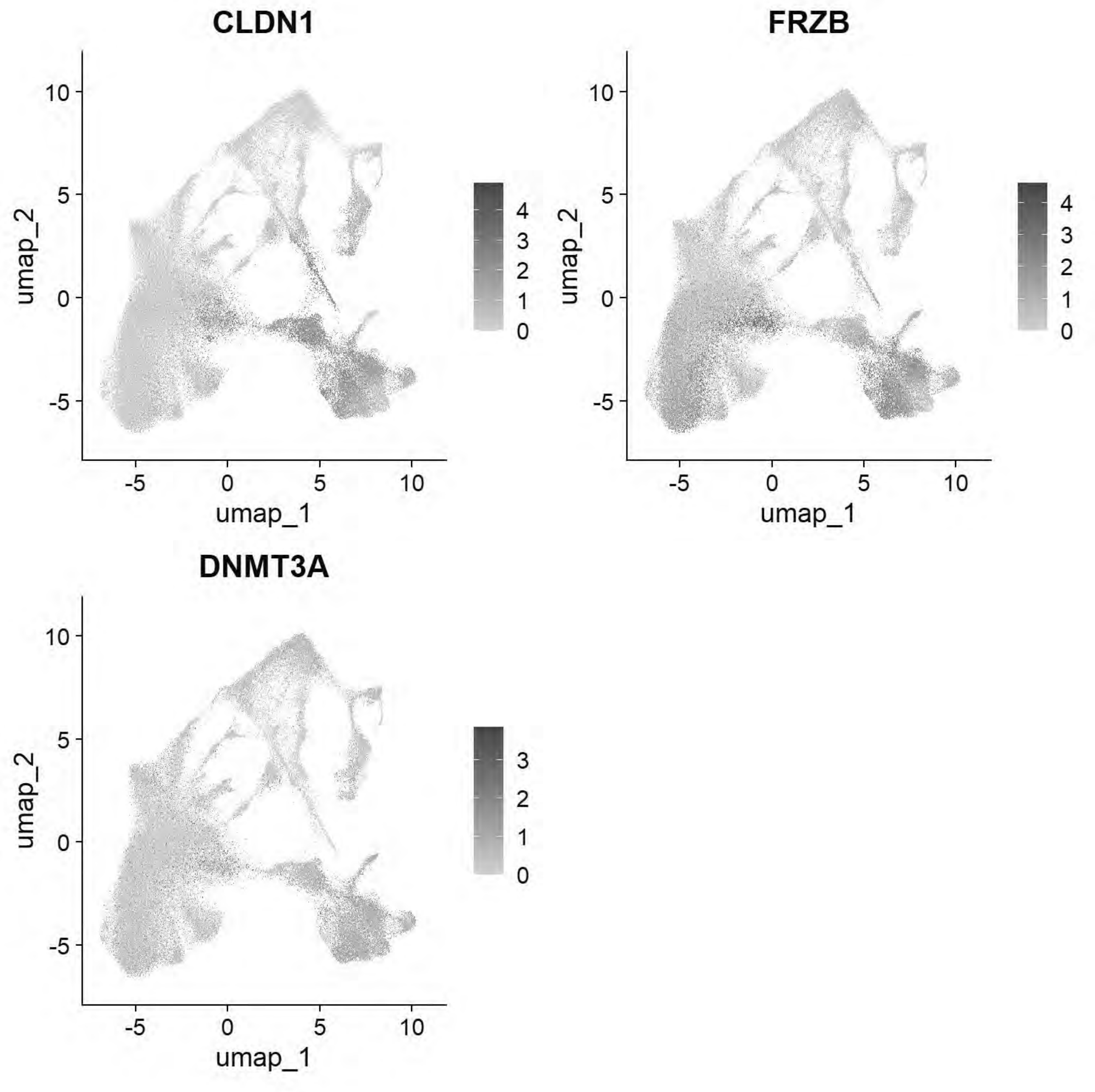


```
# Williams: Non-Neural Ectoderm
FeaturePlot(chick, features = c("DLX5","TFAP2A","ASTL","PAX6"))
```

```
## Rasterizing points since number of points exceeds 100,000.
## To disable this behavior set `raster=FALSE`
## Rasterizing points since number of points exceeds 100,000.
## To disable this behavior set `raster=FALSE`
## Rasterizing points since number of points exceeds 100,000.
## To disable this behavior set `raster=FALSE`
## Rasterizing points since number of points exceeds 100,000.
```

```
## To disable this behavior set `raster=FALSE`
```

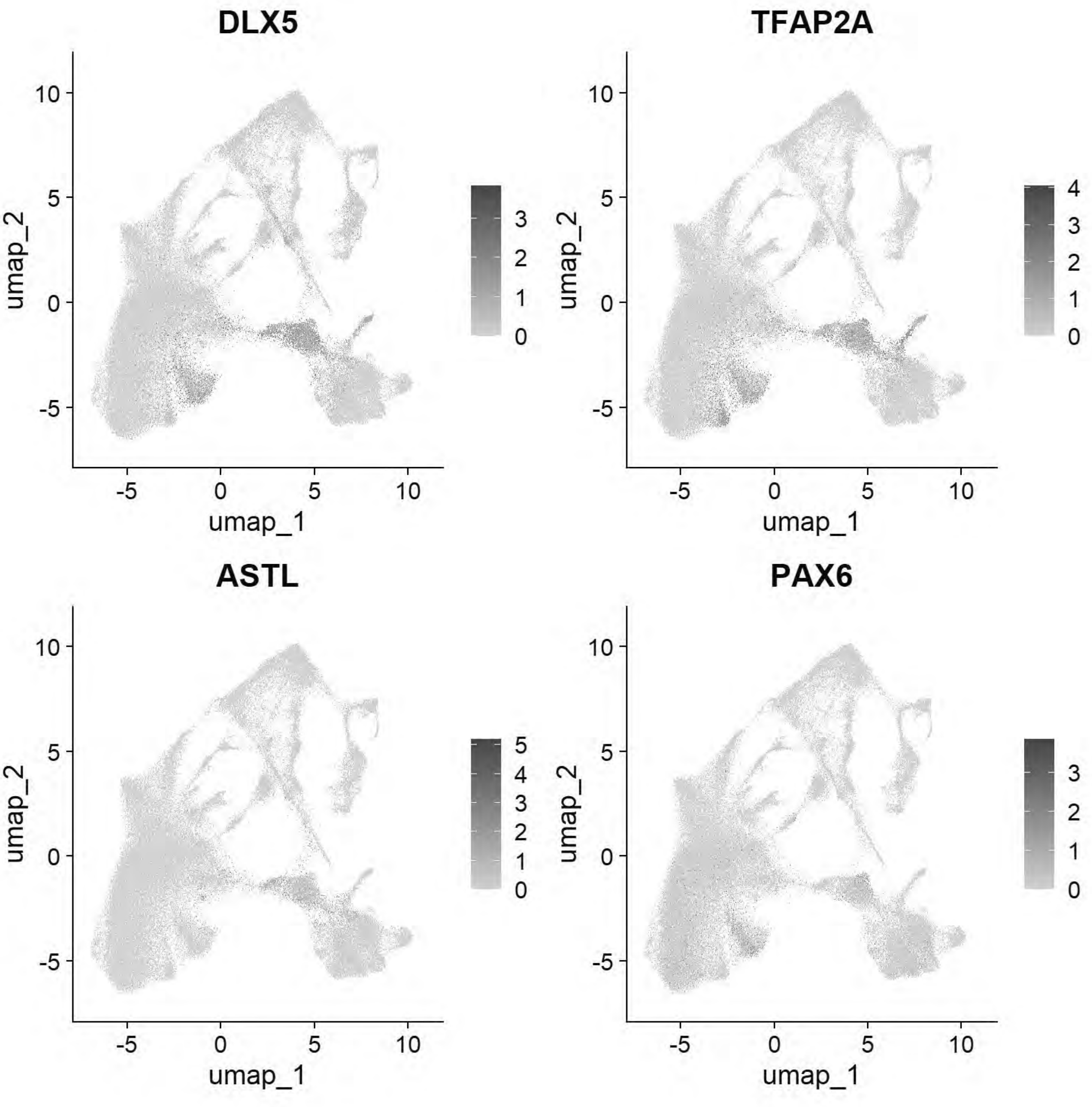


```
# Williams: Ectoderm (Neural Plate)
FeaturePlot(chick, features = c("SOX2","SOX3","SOX21","FRZB","SFRP2"))
```

```
## Rasterizing points since number of points exceeds 100,000.
## To disable this behavior set `raster=FALSE`
## Rasterizing points since number of points exceeds 100,000.
## To disable this behavior set `raster=FALSE`
## Rasterizing points since number of points exceeds 100,000.
```

```
## To disable this behavior set `raster=FALSE`
## Rasterizing points since number of points exceeds 100,000.
## To disable this behavior set `raster=FALSE`
## Rasterizing points since number of points exceeds 100,000.
## To disable this behavior set `raster=FALSE`
```

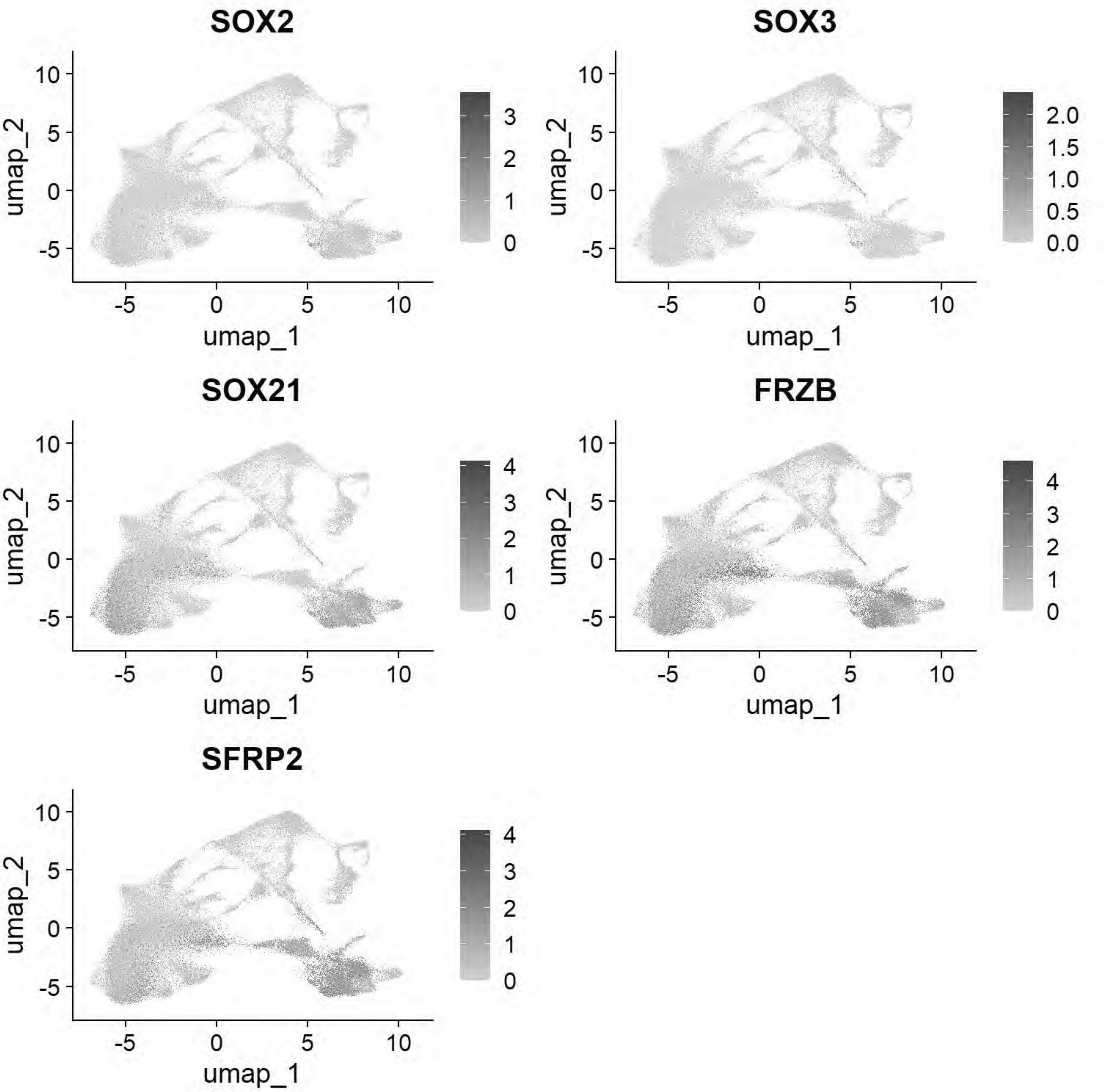


```
# Williams: Ectoderm (Neural Plate Border)
FeaturePlot(chick, features = c("PAX7","TFAP2A","DLX5","BMP4","MSX1","DRAXIN","TFAP2B"))
```

```
## Rasterizing points since number of points exceeds 100,000.
```

```
## To disable this behavior set `raster=FALSE`
## Rasterizing points since number of points exceeds 100,000.
## To disable this behavior set `raster=FALSE`
## Rasterizing points since number of points exceeds 100,000.
## To disable this behavior set `raster=FALSE`
## Rasterizing points since number of points exceeds 100,000.
## To disable this behavior set `raster=FALSE`
## Rasterizing points since number of points exceeds 100,000.
## To disable this behavior set `raster=FALSE`
## Rasterizing points since number of points exceeds 100,000.
## To disable this behavior set `raster=FALSE`
## Rasterizing points since number of points exceeds 100,000.
## To disable this behavior set `raster=FALSE`
```

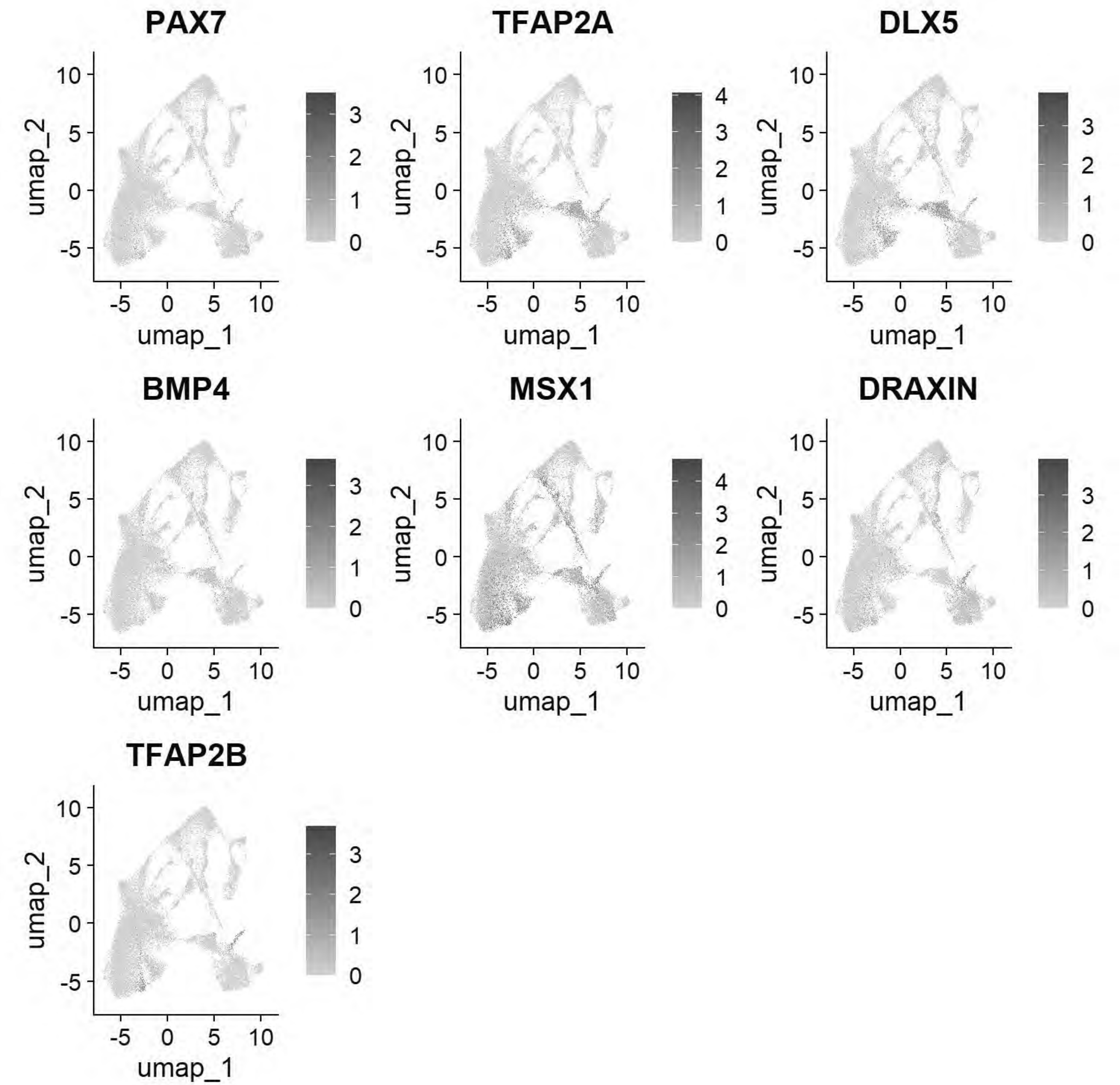


```
# Williams:Epiblast stem cell
FeaturePlot(chick, features = c("ID3","SALL4","TGIF1","ELAVL1"))
```

```
## Rasterizing points since number of points exceeds 100,000.
## To disable this behavior set `raster=FALSE`
## Rasterizing points since number of points exceeds 100,000.
## To disable this behavior set `raster=FALSE`
## Rasterizing points since number of points exceeds 100,000.
## To disable this behavior set `raster=FALSE`
## Rasterizing points since number of points exceeds 100,000.
```

```
## To disable this behavior set `raster=FALSE`
```

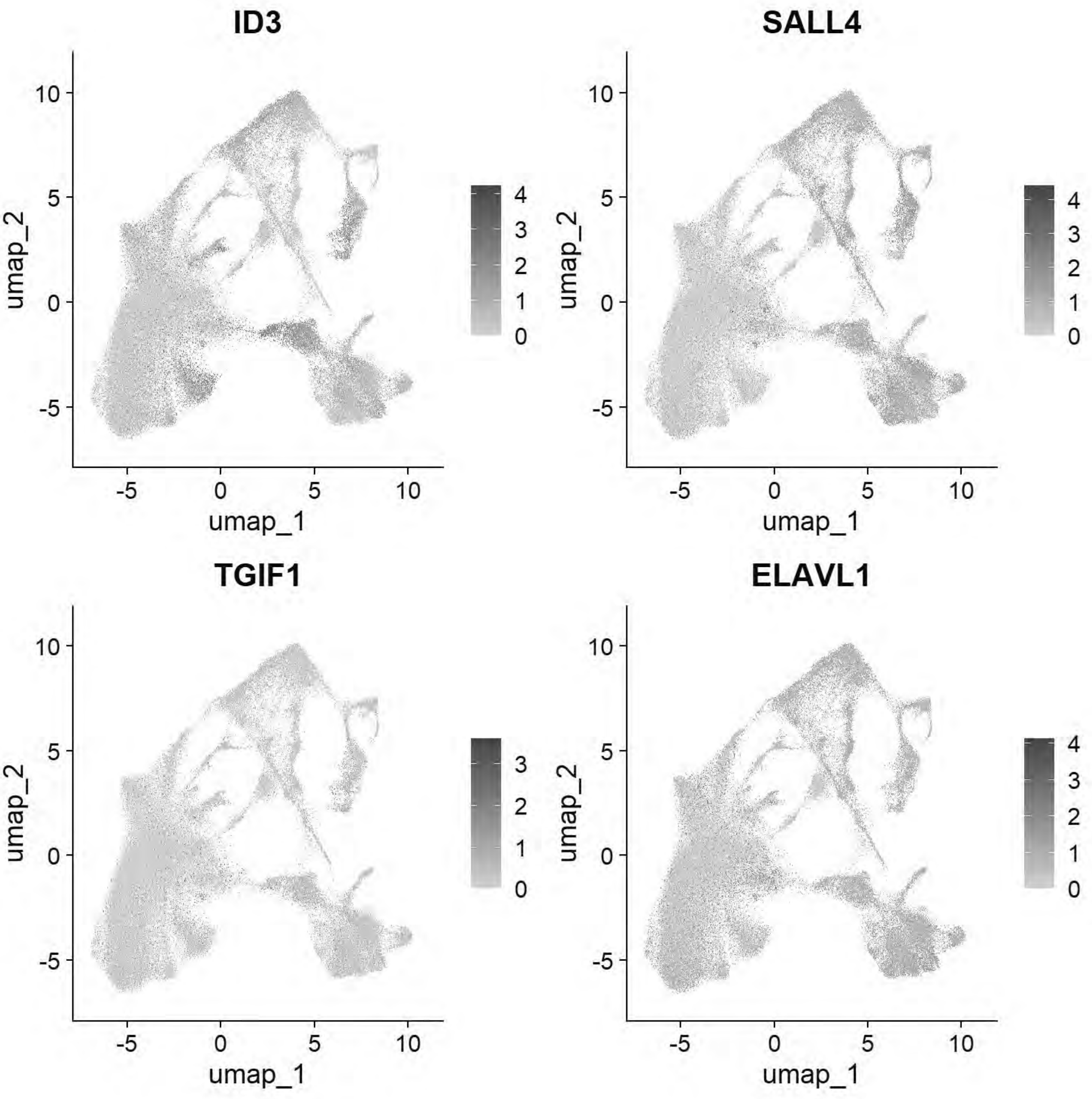


```
# Williams: Posterior Lateral Plate Mesoderm
FeaturePlot(chick, features = c("GATA2","HOXB5","CDX4"))
```

```
## Rasterizing points since number of points exceeds 100,000.
## To disable this behavior set `raster=FALSE`
## Rasterizing points since number of points exceeds 100,000.
## To disable this behavior set `raster=FALSE`
## Rasterizing points since number of points exceeds 100,000.
```

```
## To disable this behavior set `raster=FALSE`
```

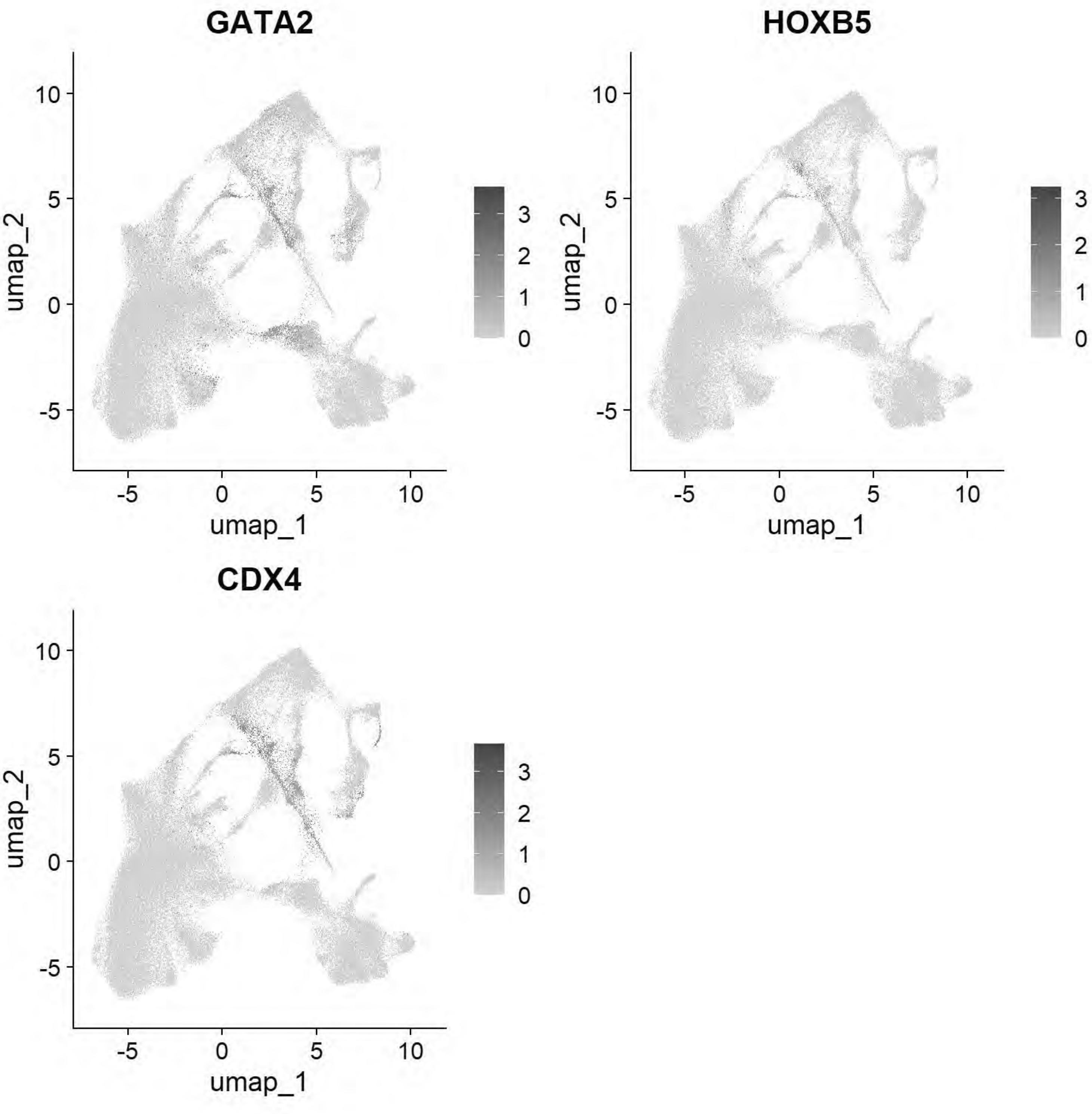


```
# Williams: Paraxial Mesoderm
FeaturePlot(chick, features = c("MSGN1","MESP1","MEOX1"))
```

```
## Rasterizing points since number of points exceeds 100,000.
## To disable this behavior set `raster=FALSE`
## Rasterizing points since number of points exceeds 100,000.
## To disable this behavior set `raster=FALSE`
## Rasterizing points since number of points exceeds 100,000.
```

```
## To disable this behavior set `raster=FALSE`
```

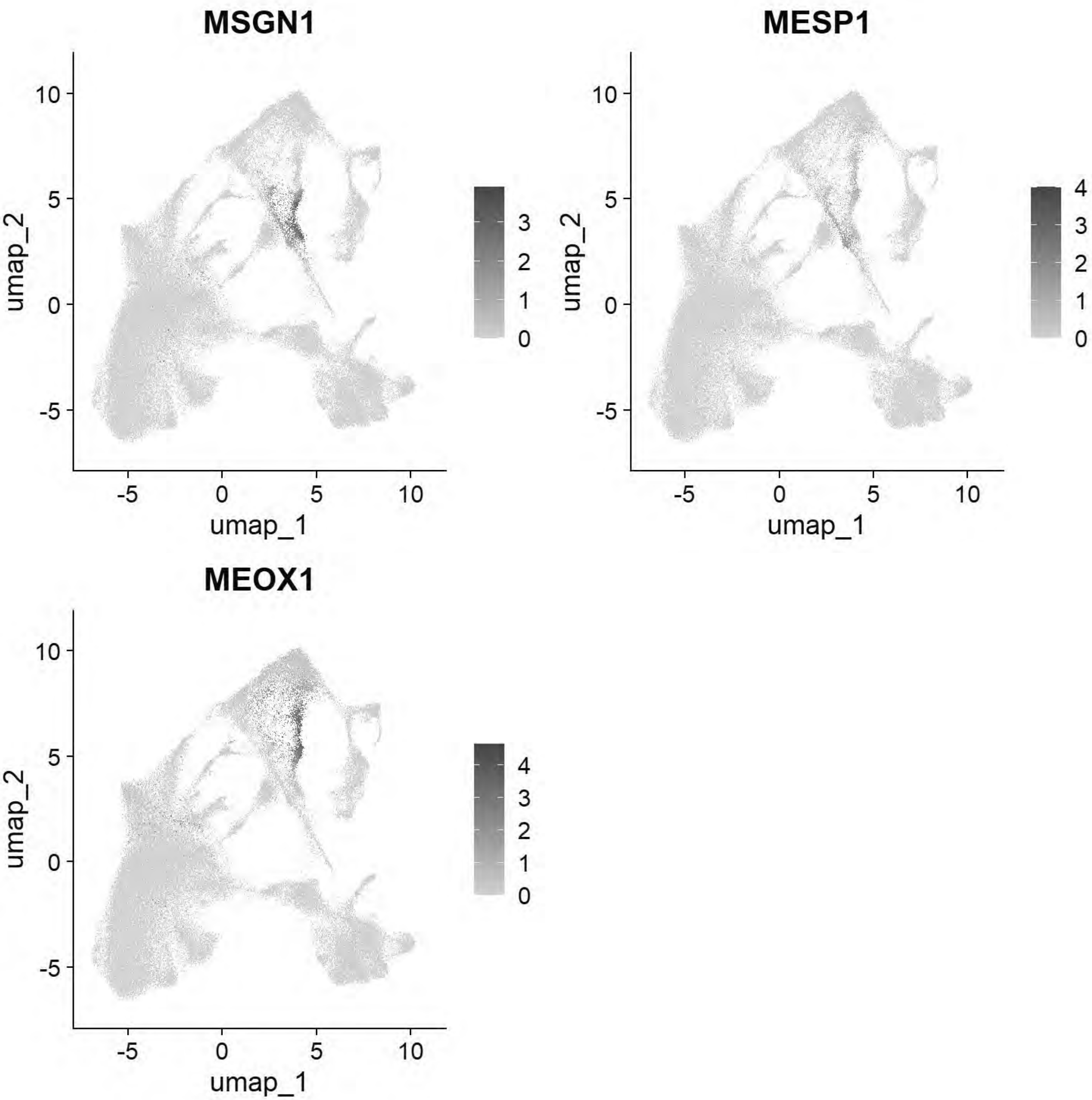


```
# Williams: Lateral Plate Mesoderm
FeaturePlot(chick, features = c("PITX2","ALX1","OLFML3","SIX1","TWIST1"))
```

```
## Rasterizing points since number of points exceeds 100,000.
## To disable this behavior set `raster=FALSE`
## Rasterizing points since number of points exceeds 100,000.
## To disable this behavior set `raster=FALSE`
## Rasterizing points since number of points exceeds 100,000.
```

```
## To disable this behavior set `raster=FALSE`
## Rasterizing points since number of points exceeds 100,000.
## To disable this behavior set `raster=FALSE`
## Rasterizing points since number of points exceeds 100,000.
## To disable this behavior set `raster=FALSE`
```

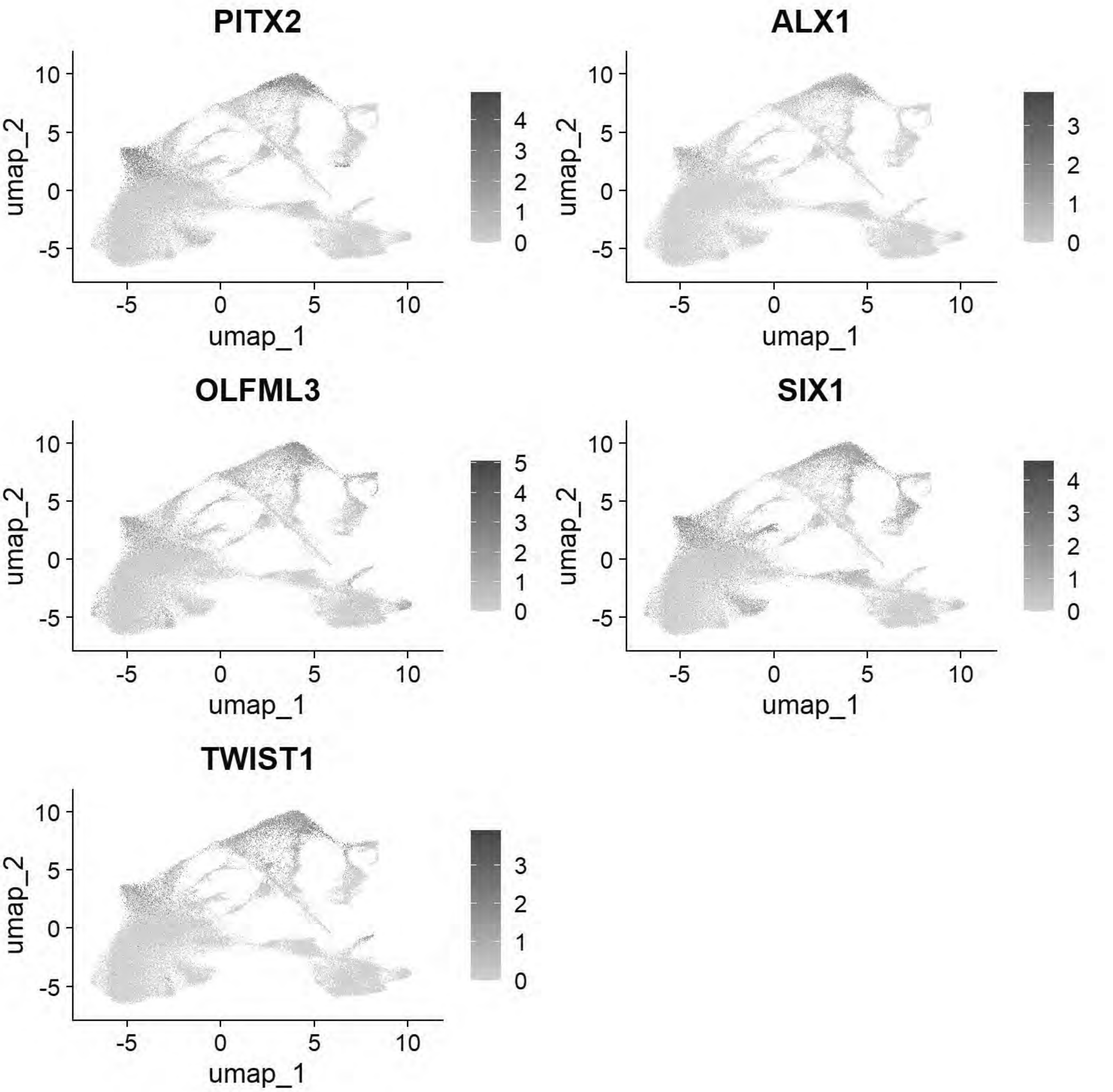


```
# Williams: Cardiac Mesoderm
FeaturePlot(chick, features = c("TCF21","GATA5","LMO2","ETS1","KDR"))
```

```
## Rasterizing points since number of points exceeds 100,000.
```

```
## To disable this behavior set `raster=FALSE`
## Rasterizing points since number of points exceeds 100,000.
## To disable this behavior set `raster=FALSE`
## Rasterizing points since number of points exceeds 100,000.
## To disable this behavior set `raster=FALSE`
## Rasterizing points since number of points exceeds 100,000.
## To disable this behavior set `raster=FALSE`
## Rasterizing points since number of points exceeds 100,000.
## To disable this behavior set `raster=FALSE`
```

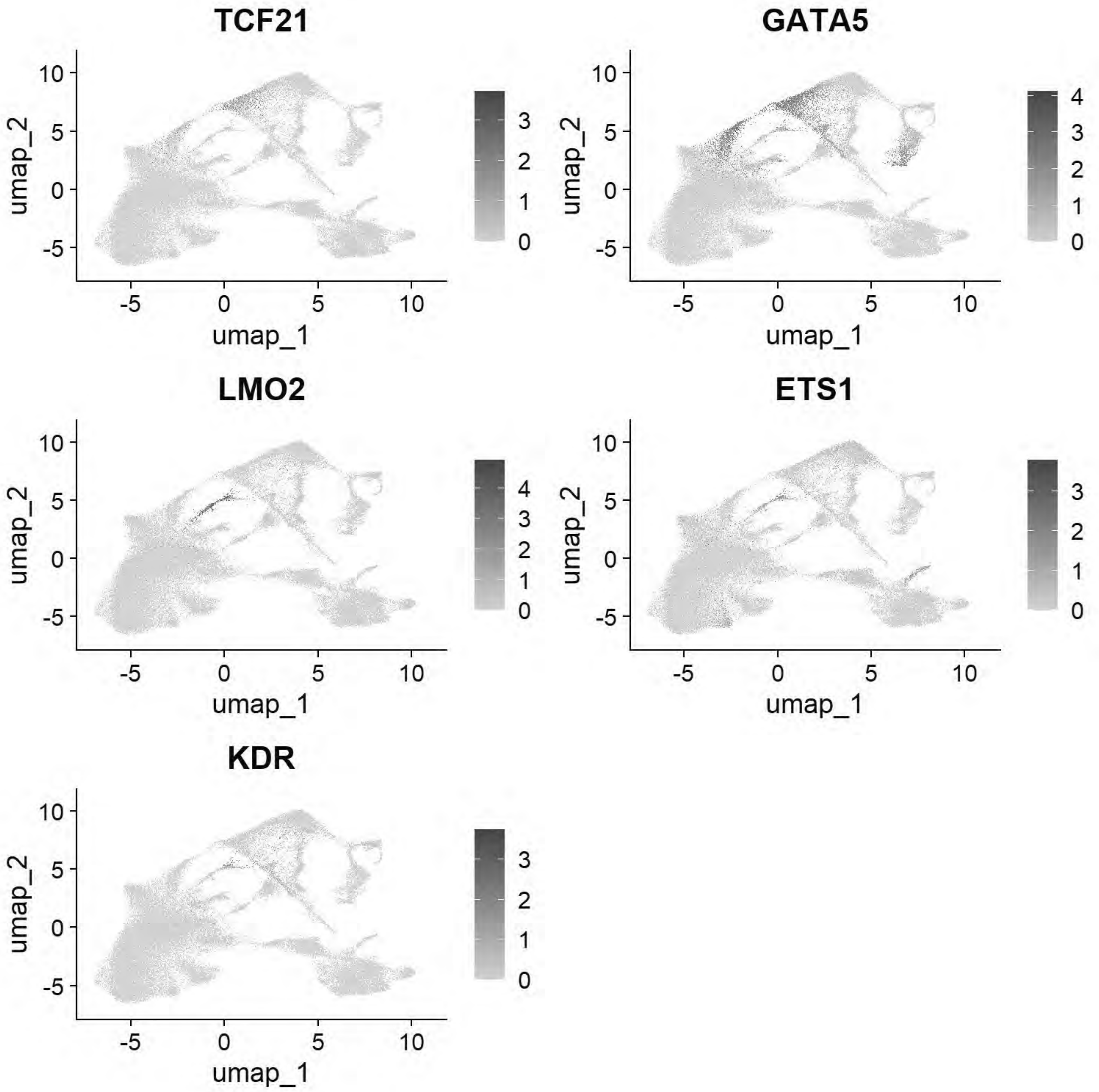

```
# Williams: Head Mesenchyme
FeaturePlot(chick, features = c("TCF21","GATA5","LMO2","ETS1","KDR"))
```

```
## Rasterizing points since number of points exceeds 100,000.
## To disable this behavior set `raster=FALSE`
## Rasterizing points since number of points exceeds 100,000.
## To disable this behavior set `raster=FALSE`
## Rasterizing points since number of points exceeds 100,000.
## To disable this behavior set `raster=FALSE`
## Rasterizing points since number of points exceeds 100,000.
## To disable this behavior set `raster=FALSE`
## Rasterizing points since number of points exceeds 100,000.
## To disable this behavior set `raster=FALSE`
```

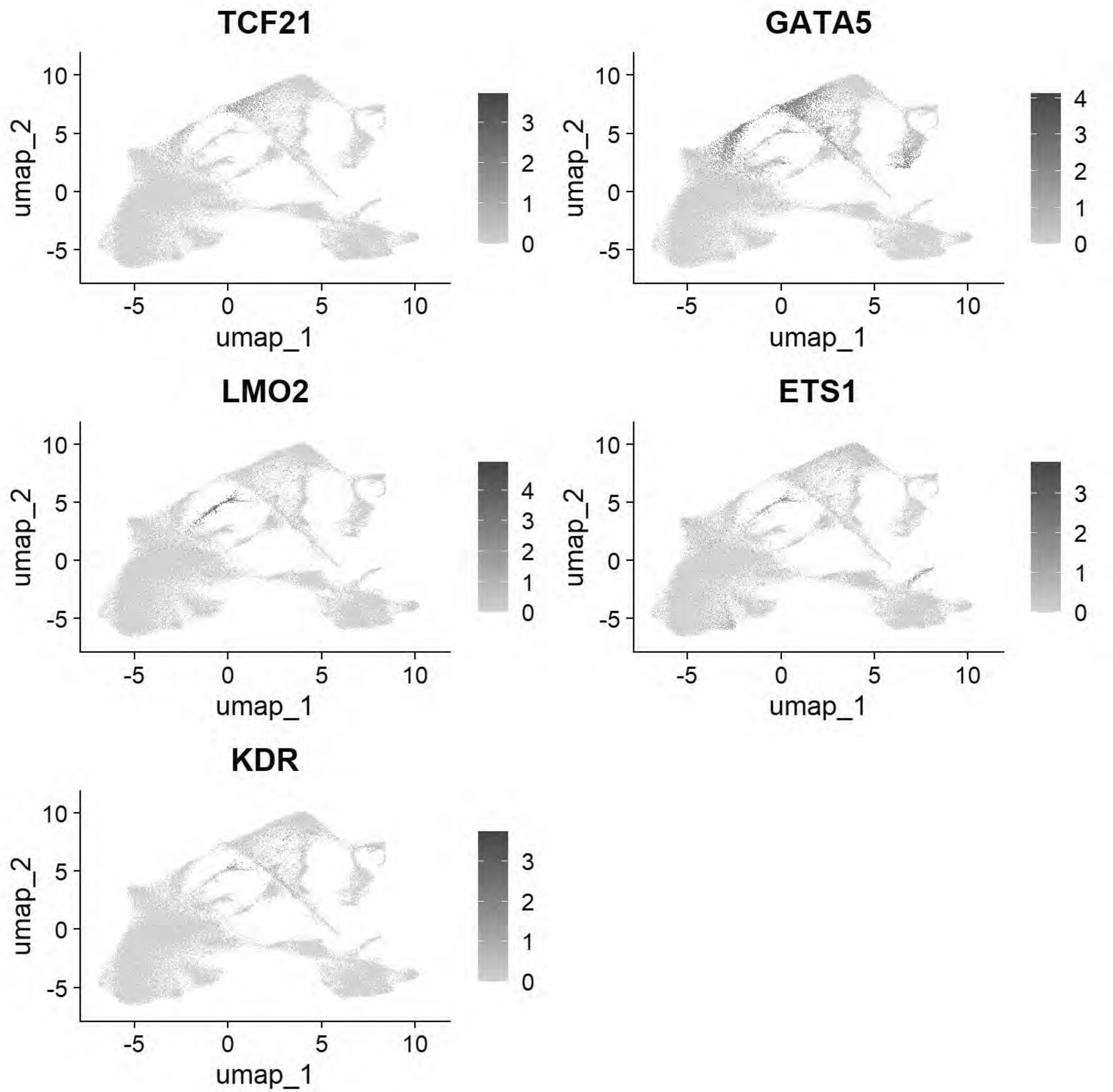


```
# Williams: Hensen's Node & Primitive Streak
FeaturePlot(chick, features = c("DLL1","FGF8","NOTO","CHRD"))
```

```
## Rasterizing points since number of points exceeds 100,000.
## To disable this behavior set `raster=FALSE`
## Rasterizing points since number of points exceeds 100,000.
## To disable this behavior set `raster=FALSE`
## Rasterizing points since number of points exceeds 100,000.
## To disable this behavior set `raster=FALSE`
## Rasterizing points since number of points exceeds 100,000.
```

```
## To disable this behavior set `raster=FALSE`
```

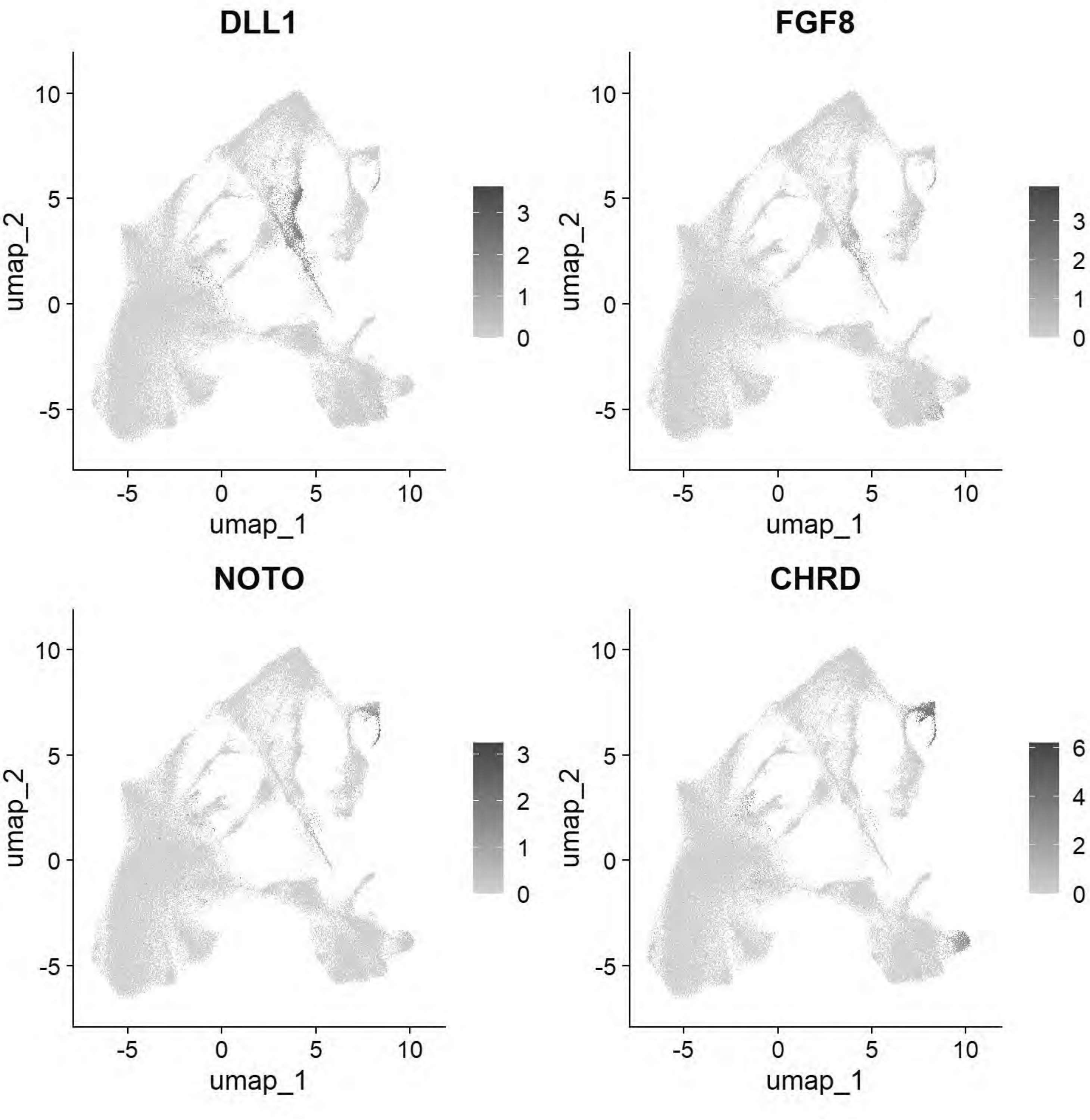


```
# Williams: Endoderm
FeaturePlot(chick, features = c("SOX17","FOXA2","CXCR4"))
```

```
## Rasterizing points since number of points exceeds 100,000.
## To disable this behavior set `raster=FALSE`
## Rasterizing points since number of points exceeds 100,000.
## To disable this behavior set `raster=FALSE`
## Rasterizing points since number of points exceeds 100,000.
```

```
## To disable this behavior set `raster=FALSE`
```

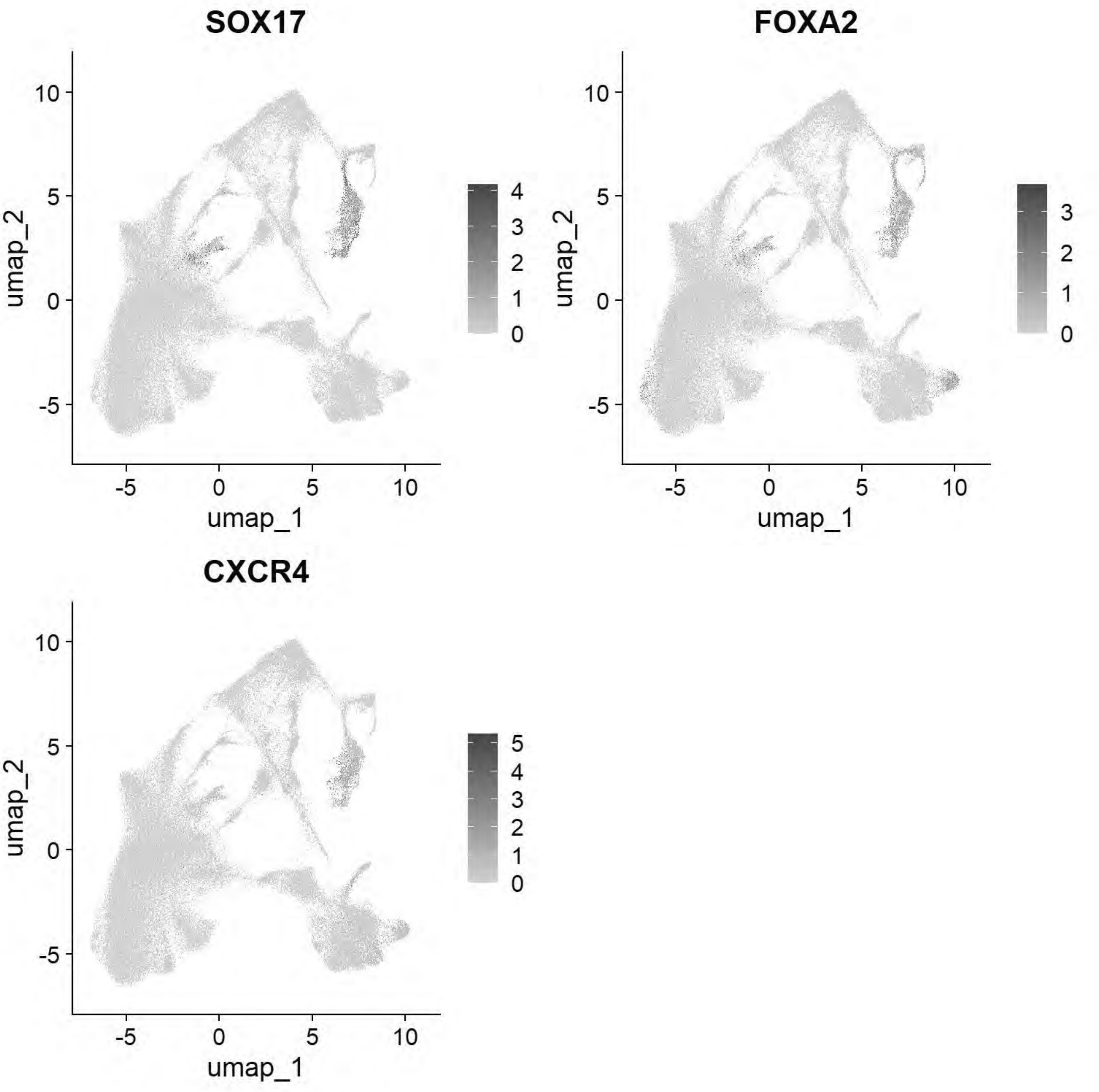


```
# Pajanoja: Ectoderm
FeaturePlot(chick, features = c("TFAP2A","DLX5","CLDN1","SOX2","NESTIN","MYCN"))
```

```
## Warning: The following requested variables were not found: NESTIN
```

```
## Rasterizing points since number of points exceeds 100,000.
## To disable this behavior set `raster=FALSE`
```

```
## Rasterizing points since number of points exceeds 100,000.
## To disable this behavior set `raster=FALSE`
## Rasterizing points since number of points exceeds 100,000.
## To disable this behavior set `raster=FALSE`
## Rasterizing points since number of points exceeds 100,000.
## To disable this behavior set `raster=FALSE`
## Rasterizing points since number of points exceeds 100,000.
## To disable this behavior set `raster=FALSE`
```

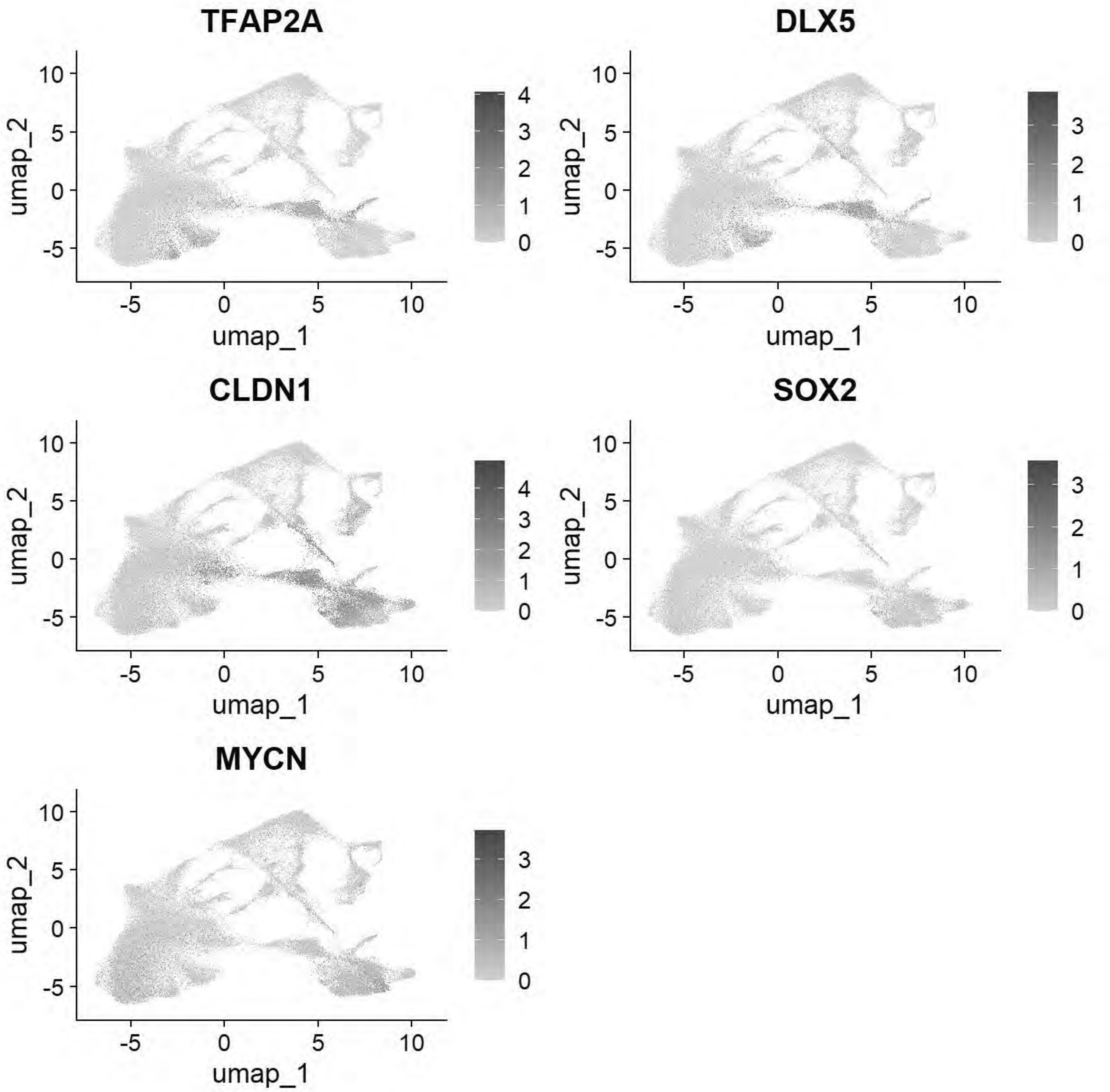


```
# Pajanoja: Endoderm
```

```
FeaturePlot(chick, features = c("SOX17","KRT17","HHEX"))
```

```
## Rasterizing points since number of points exceeds 100,000.
## To disable this behavior set `raster=FALSE`
## Rasterizing points since number of points exceeds 100,000.
## To disable this behavior set `raster=FALSE`
## Rasterizing points since number of points exceeds 100,000.
## To disable this behavior set `raster=FALSE`
```

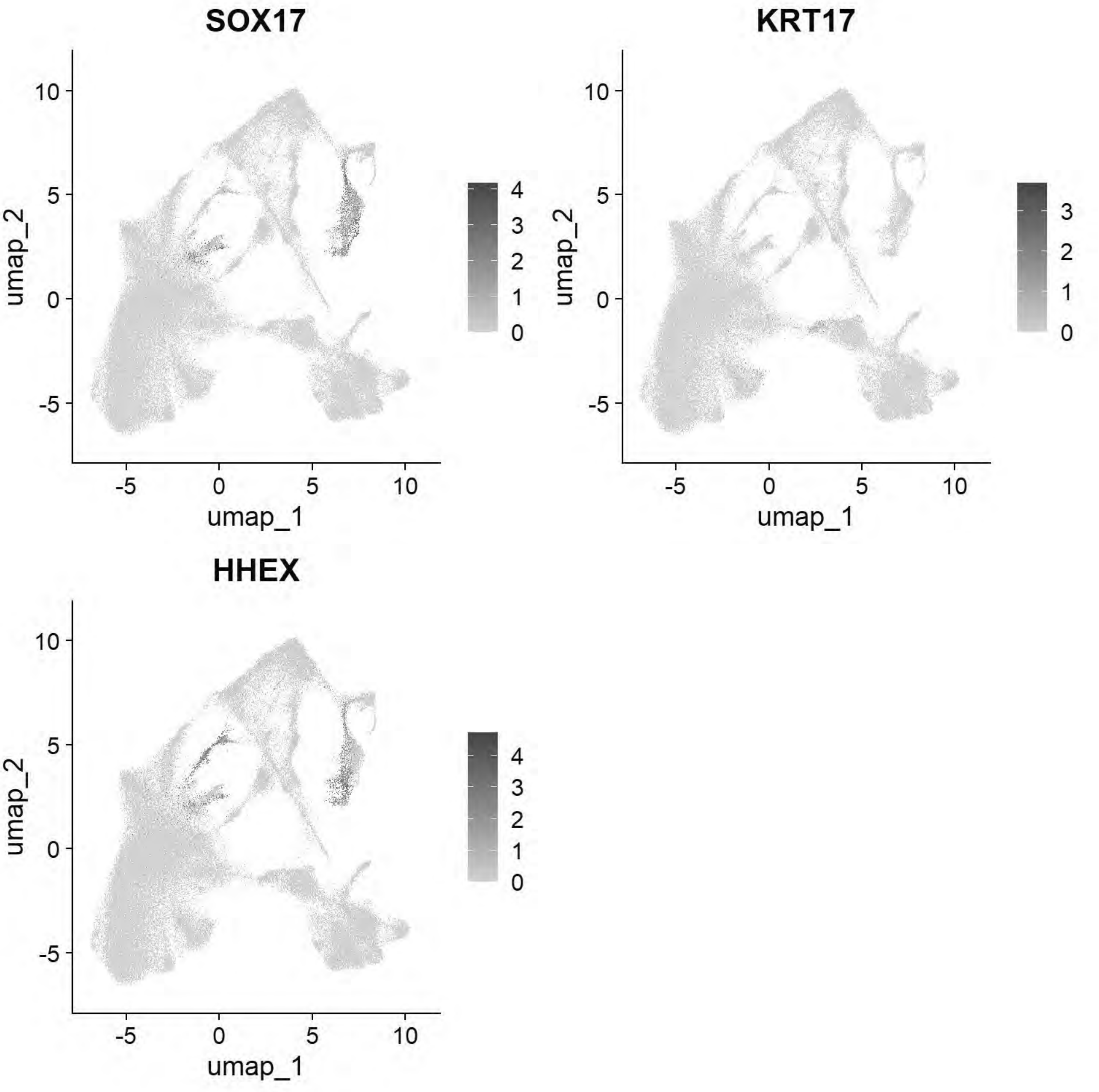


```
# Pajanoja: Mesoderm
```

```
FeaturePlot(chick, features = c("ALX1","TWIST1","HAND2","PITX2"))
```

```
## Rasterizing points since number of points exceeds 100,000.
## To disable this behavior set `raster=FALSE`
## Rasterizing points since number of points exceeds 100,000.
## To disable this behavior set `raster=FALSE`
## Rasterizing points since number of points exceeds 100,000.
## To disable this behavior set `raster=FALSE`
## Rasterizing points since number of points exceeds 100,000.
## To disable this behavior set `raster=FALSE`
```

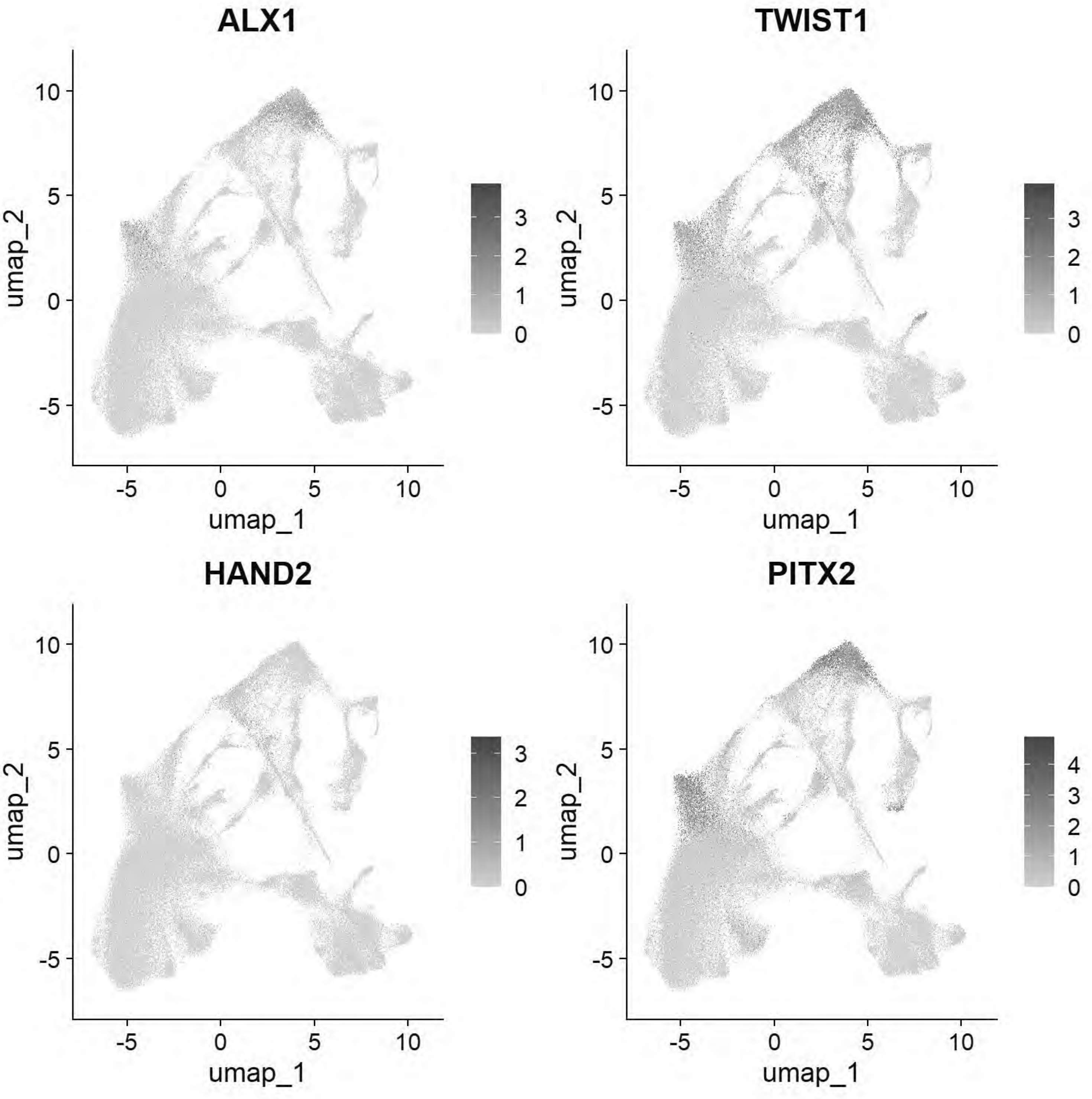

```
# Pajanoja: Notochord
FeaturePlot(chick, features = c("CHRD","TBXT","NOTO"))
```

```
## Rasterizing points since number of points exceeds 100,000.
## To disable this behavior set `raster=FALSE`
## Rasterizing points since number of points exceeds 100,000.
## To disable this behavior set `raster=FALSE`
## Rasterizing points since number of points exceeds 100,000.
## To disable this behavior set `raster=FALSE`
```

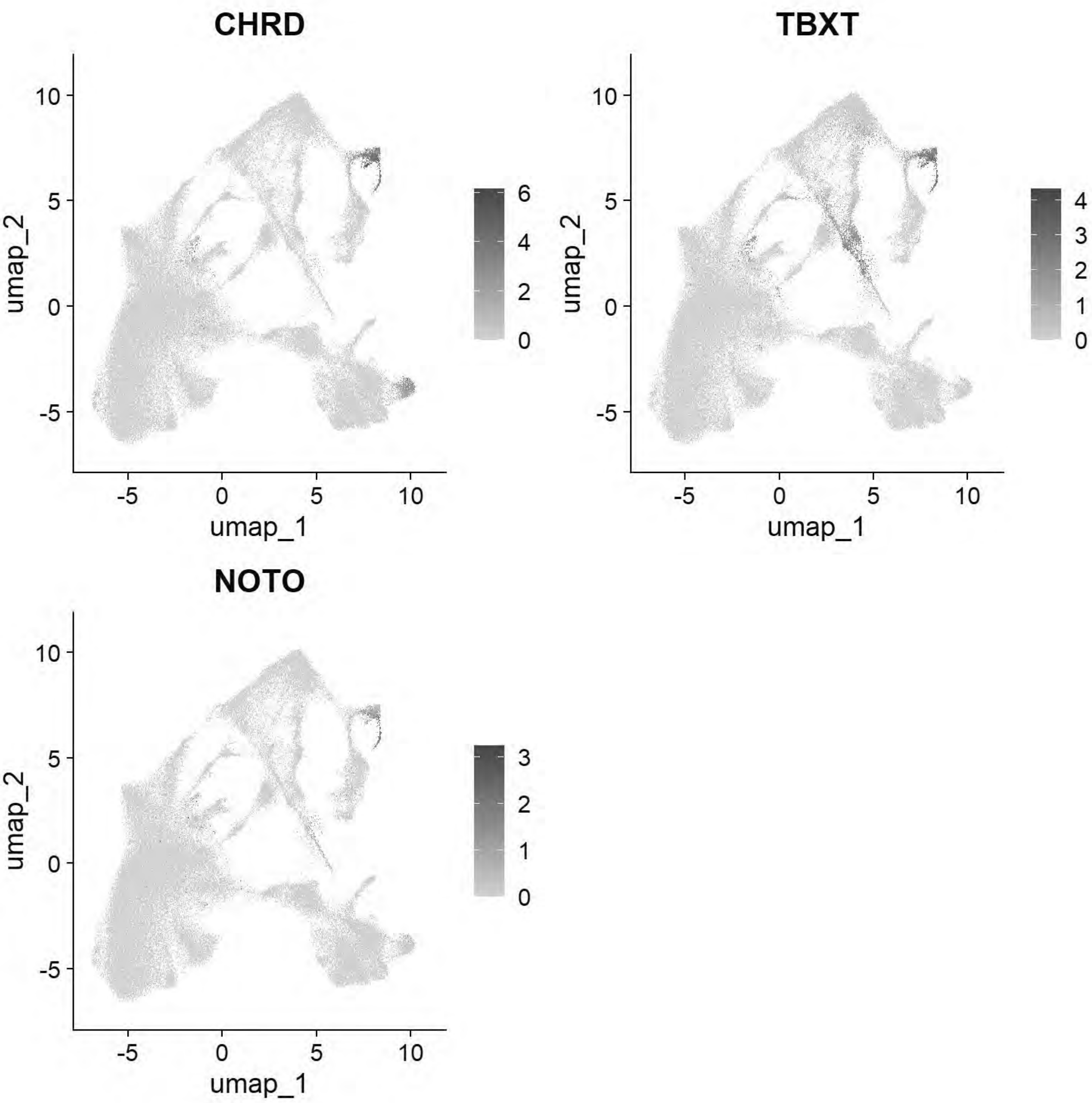

```
# Pajanoja: Ventral Neural Tube
FeaturePlot(chick, features=c("SHH","FOXA1","FOXA2"))
```

```
## Rasterizing points since number of points exceeds 100,000.
## To disable this behavior set `raster=FALSE`
## Rasterizing points since number of points exceeds 100,000.
## To disable this behavior set `raster=FALSE`
## Rasterizing points since number of points exceeds 100,000.
## To disable this behavior set `raster=FALSE`
```

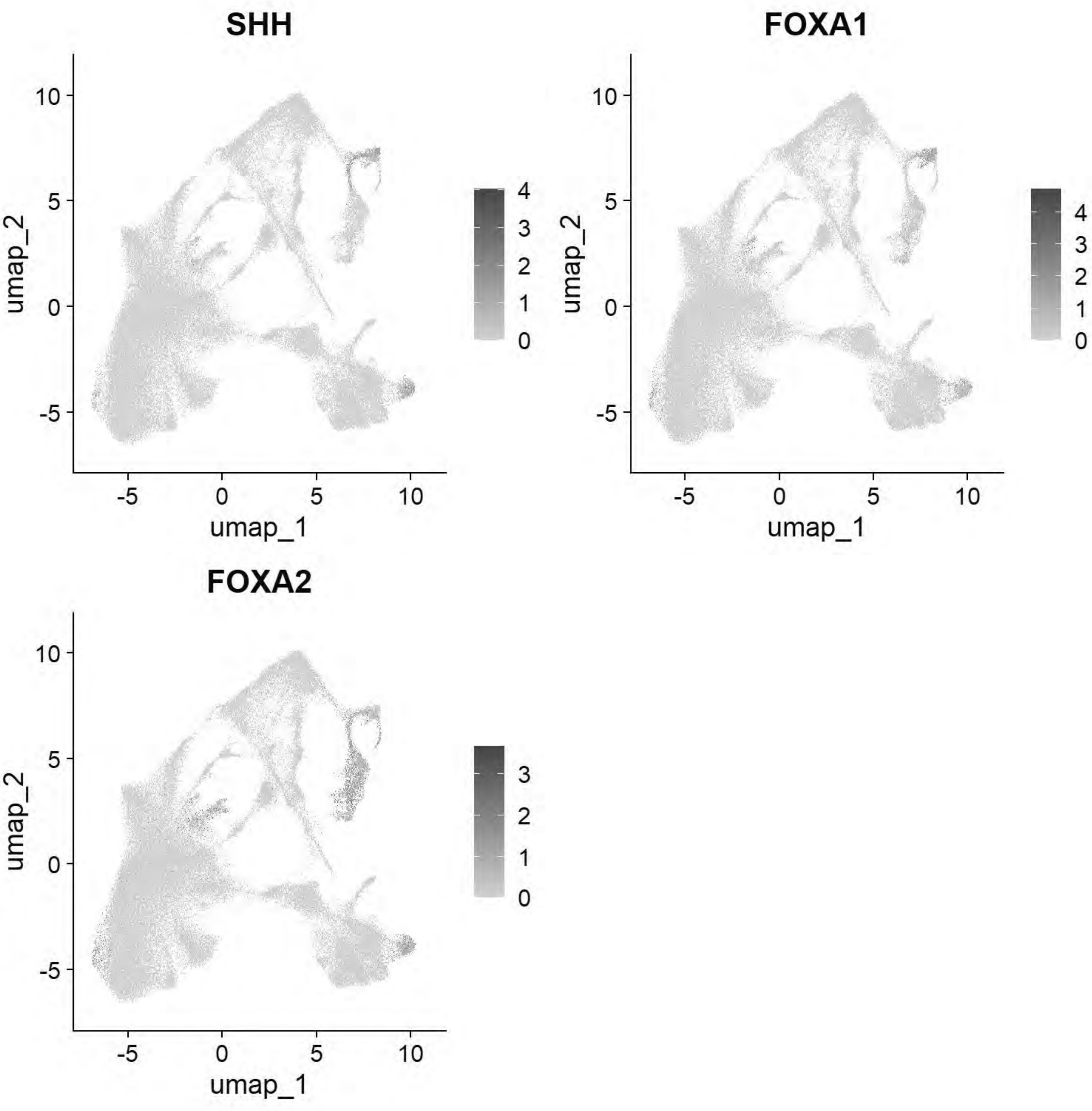

```
# Pajanoja: Neural Tube
FeaturePlot(chick, features=c("CDH2","HES5","MYCN","PAX2","NESTIN"))
```

```
## Warning: The following requested variables were not found: NESTIN
```

```
## Rasterizing points since number of points exceeds 100,000.
## To disable this behavior set `raster=FALSE`
## Rasterizing points since number of points exceeds 100,000.
## To disable this behavior set `raster=FALSE`
## Rasterizing points since number of points exceeds 100,000.
## To disable this behavior set `raster=FALSE`
## Rasterizing points since number of points exceeds 100,000.
## To disable this behavior set `raster=FALSE`
```

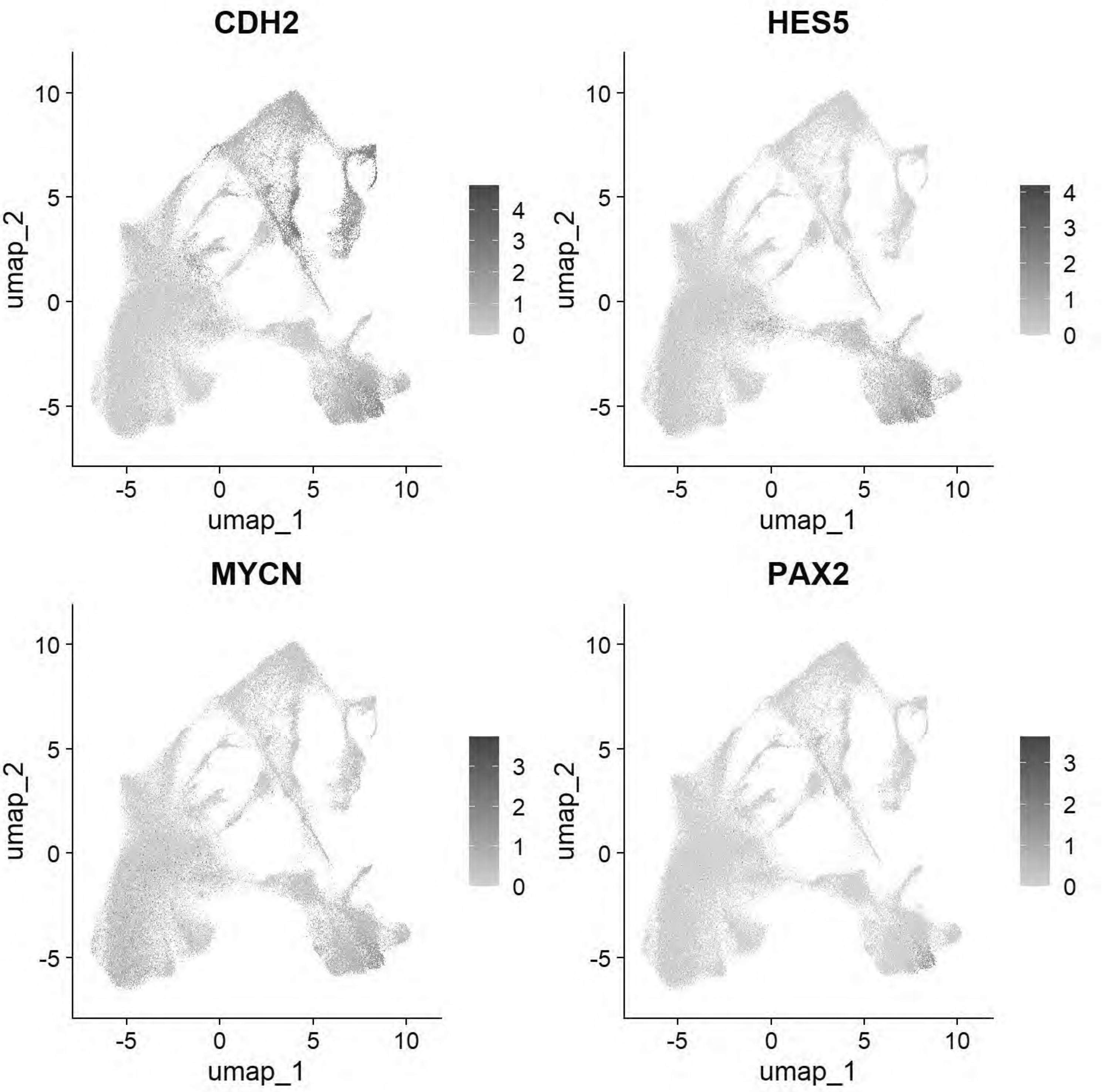


```
# Pajanoja: Non-Neural Ectoderm/Placode
FeaturePlot(chick, features=c("KRT24","KRT18","DLX5","SIX1","EYA2","TFAP2A"))
```

```
## Rasterizing points since number of points exceeds 100,000.
## To disable this behavior set `raster=FALSE`
## Rasterizing points since number of points exceeds 100,000.
## To disable this behavior set `raster=FALSE`
## Rasterizing points since number of points exceeds 100,000.
## To disable this behavior set `raster=FALSE`
## Rasterizing points since number of points exceeds 100,000.
```

```
## To disable this behavior set `raster=FALSE`
## Rasterizing points since number of points exceeds 100,000.
## To disable this behavior set `raster=FALSE`
## Rasterizing points since number of points exceeds 100,000.
## To disable this behavior set `raster=FALSE`
```

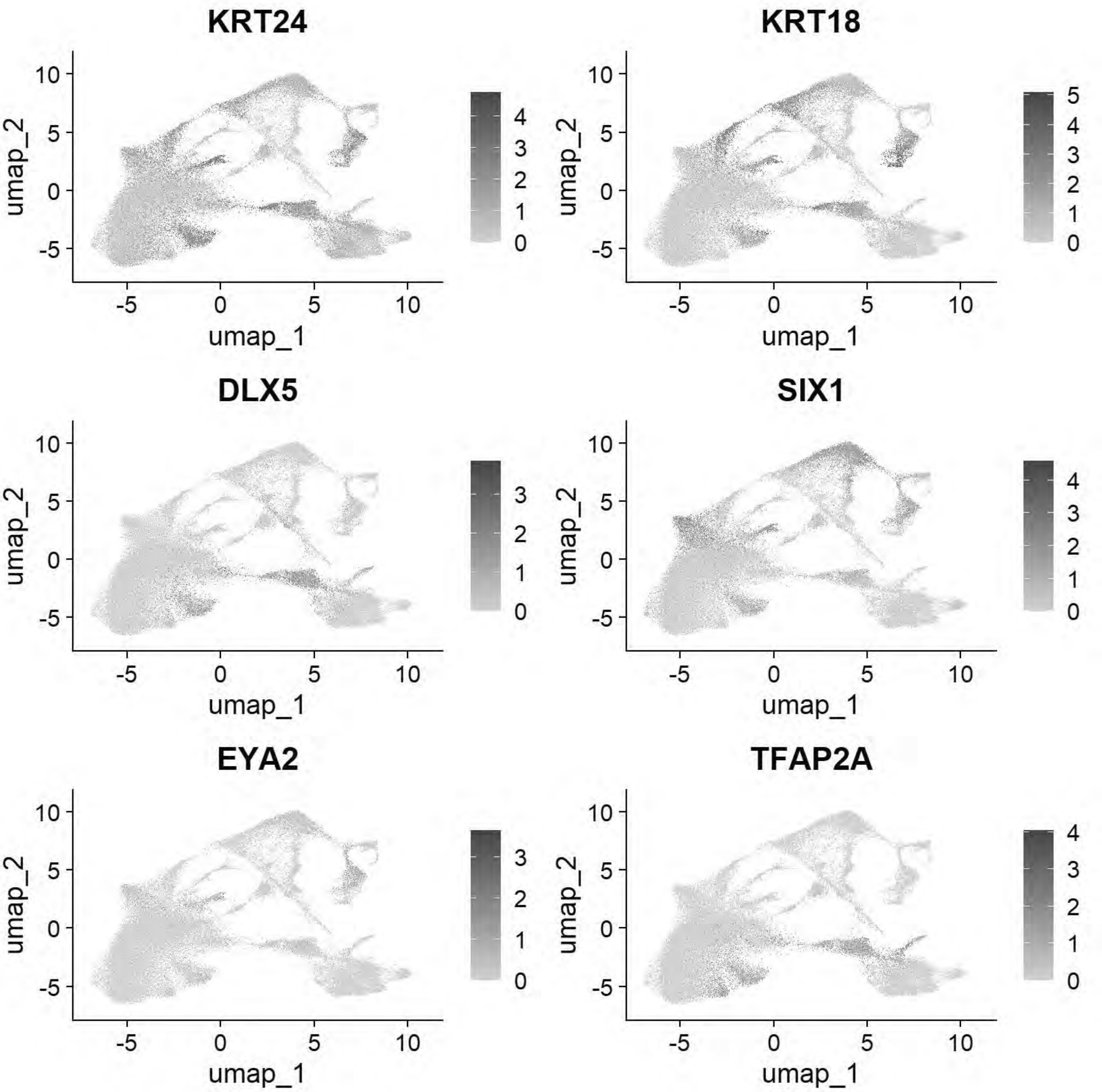


```
# Pajanoja: Neural crest
FeaturePlot(chick, features=c("FOXD3","TFAP2B","SOX9","ETS1","SNAI2"))
```

```
## Rasterizing points since number of points exceeds 100,000.
```

```
## To disable this behavior set `raster=FALSE`
## Rasterizing points since number of points exceeds 100,000.
## To disable this behavior set `raster=FALSE`
## Rasterizing points since number of points exceeds 100,000.
## To disable this behavior set `raster=FALSE`
## Rasterizing points since number of points exceeds 100,000.
## To disable this behavior set `raster=FALSE`
## Rasterizing points since number of points exceeds 100,000.
## To disable this behavior set `raster=FALSE`
```

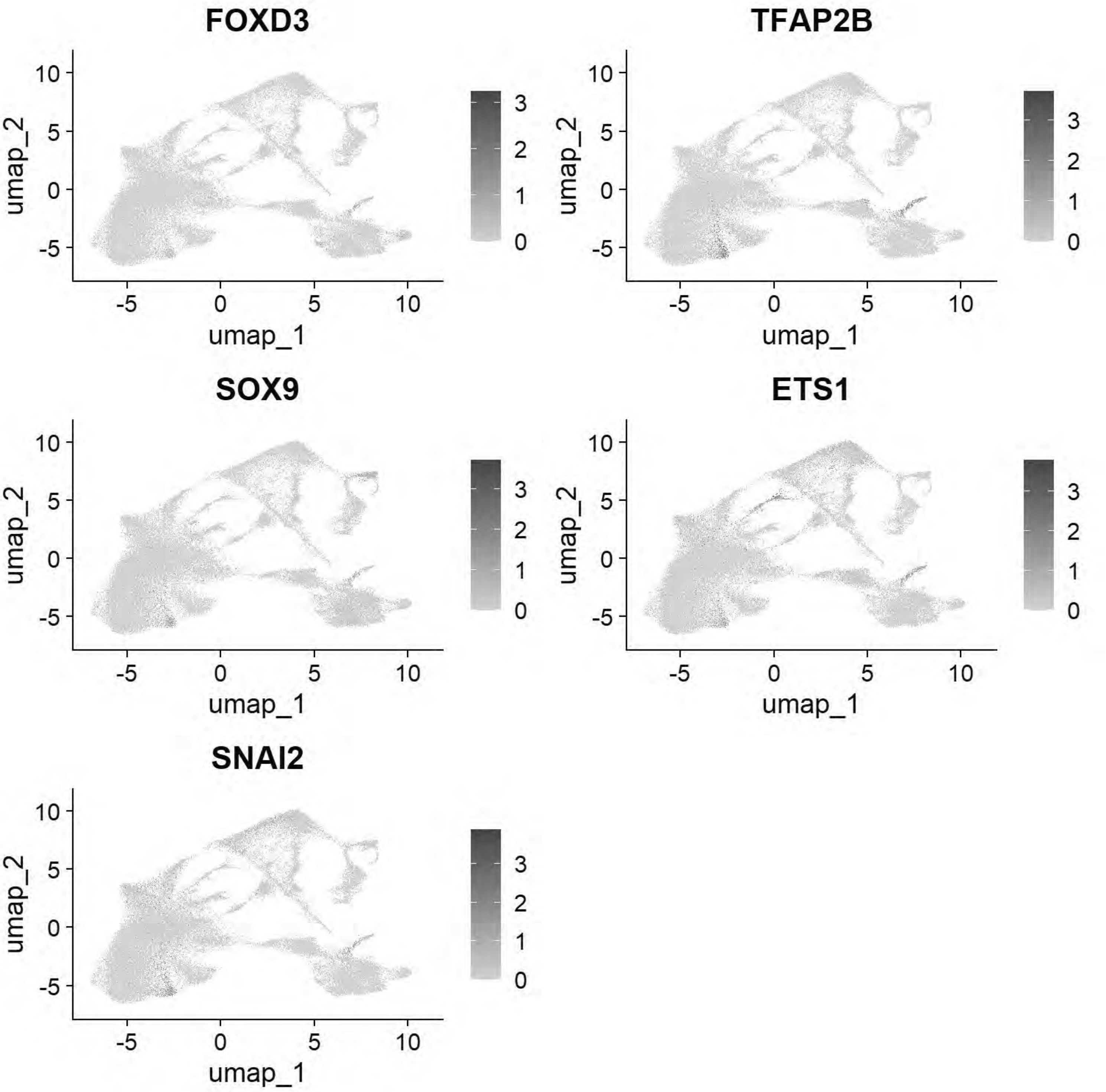

```
gc()
```

```
##              used   (Mb) gc trigger    (Mb)   max used    (Mb)
## Ncells   4630309  247.3   13651908   729.1   21331105  1139.3
## Vcells 820843221 6262.6 2298060473 17532.9 4488399353 34243.8
```

```
chick <- JoinLayers(chick)
chick.markers <- FindAllMarkers(chick, min.pct = 0.5, logfc.threshold = 0.5)
```

```
## Calculating cluster 0
```

```
## Calculating cluster 1
```

```
## Calculating cluster 2
```

```
## Calculating cluster 3
```

```
## Calculating cluster 4
```

```
## Calculating cluster 5
```

```
## Calculating cluster 6
```

```
## Calculating cluster 7
```

```
## Calculating cluster 8
```

```
## Calculating cluster 9
```

```
## Calculating cluster 10
```

```
## Calculating cluster 11
```

```
## Calculating cluster 12
```

```
## Calculating cluster 13
```

```
## Calculating cluster 14
```

```
## Calculating cluster 15
```

```
## Calculating cluster 16
```

```
## Calculating cluster 17
```

```
## Calculating cluster 18
```

```
## Calculating cluster 19
```

```
## Calculating cluster 20
```

```
## Calculating cluster 21
```

```
## Calculating cluster 22
```

```
## Calculating cluster 23
```

```
## Calculating cluster 24
```

```
## Calculating cluster 25
```

```
## Calculating cluster 26
```

```
## Warning: The following tests were not performed:
```

```
## Warning: When testing 22 versus all:
##  invalid class "dgCMatrix" object: 'i' slot has elements not in {0,...,Dim[1]-1}
```

```
chick.markers %>%
  group_by(cluster) %>%
  top_n(n = 5, wt = avg_log2FC) -> top5
DotPlot(chick, features = unique(top5$gene))+coord_flip()
```

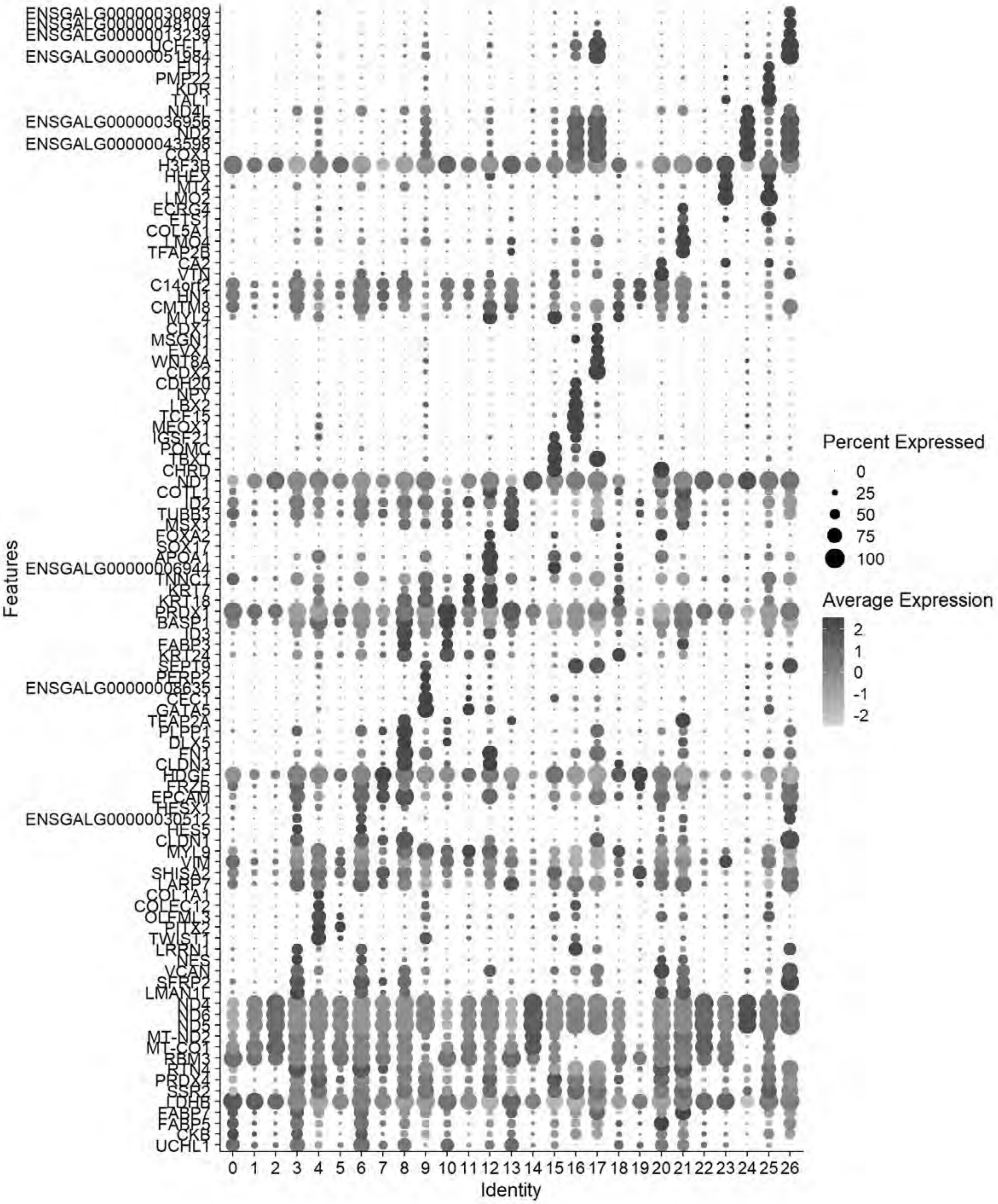
ENSGALG00000030809
ENSGALG00000048104
ENSGALG00000013239
ENSGALG00000051984
FLI1
PMP22
KDR
TAL1
ND4L
ENSGALG00000036956
ND2
ENSGALG00000043598
COX1
H3F3B
HHEX
MT4
LMO2
ECRG4
ETS1
COL5A1
LMO4
TFAP2B
CA2
VTN
C14orf2
HN1
CMTM8
MYL4
CDX1
MSGN1
EVX1
WNT8A
CDX2
CDH20
NPY
LBX2
TCF15
MEOX1
IGSF21
POMC
TBXT
CHRD
ND1
COTL1
ID2
TUBB3
MSX1
FOXA2
SOX17
APOA1
ENSGALG00000006944
TNNC1
KRT17
KRT18
PRDX1
BASP1
ID3
FABP3
KRT24
SEPT9
ENSGALG00000008635
CEC1
GATA5
TFAP2A
PLPP1
DLX5
FN1
CLDN3
HDGF
FRZB
EPCAM
HESX1
ENSGALG00000030512
HES5
CLDN1
MYL9
VIM
SHISA2
LARP7
COL1A1
COLEC12
OLFML3
PITX2
TWIST1
LRRN1
NES
VCAN
SFRP2
LMAN1L
ND4
ND6
ND5
MT-ND2
MT-CO1
RBM3
RTN4
PRDX4
SSR2
LDHB
FABP7
FABP5
CKB
UCHL1
Features
0 1 2 3 4 5 6 7 8 9 10 11 12 13 14 15 16 17 18 19 20 21 22 23 24 25 26
Identity
Percent Expressed
0
25
50
75
100
Average Expression
2
1
0
-1
-2

```
gc()
```

```
##              used    (Mb) gc trigger    (Mb)   max used    (Mb)
## Ncells    4719712   252.1   13651908   729.1   21331105  1139.3
## Vcells 1490119261 11368.8 3309383080 25248.6 4488399353 34243.8
```

```
#Filter low quality clusters (1, 2, 14, 22)
chick
```

```
## An object of class Seurat
## 18523 features across 107542 samples within 1 assay
## Active assay: RNA (18523 features, 2000 variable features)
##  3 layers present: data, counts, scale.data
##  3 dimensional reductions calculated: pca, harmony, umap
```

```
Idents(chick) <- "seurat_clusters"
chick=subset(chick, idents =c("1","2","14","22"), invert = TRUE)
chick
```

```
## An object of class Seurat
## 18523 features across 82092 samples within 1 assay
## Active assay: RNA (18523 features, 2000 variable features)
##  3 layers present: data, counts, scale.data
##  3 dimensional reductions calculated: pca, harmony, umap
```

```
chick[["RNA"]] <- split(chick[["RNA"]], f = chick$sample_id)
```

```
## Splitting 'counts', 'data' layers. Not splitting 'scale.data'. If you would like to split o
ther layers, set in `layers` argument.
```

```
chick
```

```
## An object of class Seurat
## 18523 features across 82092 samples within 1 assay
## Active assay: RNA (18523 features, 2000 variable features)
##  27 layers present: data.Pajanoja2023_HH5_1, data.Pajanoja2023_HH5_2, data.Pajanoja2023_HH7
_1, data.Pajanoja2023_HH7_2, data.Pajanoja2023_HH8_1, data.Pajanoja2023_HH8_2, data.Pajanoja20
23_HH9_1, data.Pajanoja2023_HH9_2, data.Pajanoja2025_HH11, data.Williams_HH4, data.Williams_HH
5, data.Williams_HH6, data.Williams_HH7, scale.data, counts.Pajanoja2023_HH5_1, counts.Pajanoj
a2023_HH5_2, counts.Pajanoja2023_HH7_1, counts.Pajanoja2023_HH7_2, counts.Pajanoja2023_HH8_1,
counts.Pajanoja2023_HH8_2, counts.Pajanoja2023_HH9_1, counts.Pajanoja2023_HH9_2, counts.Pajano
ja2025_HH11, counts.Williams_HH4, counts.Williams_HH5, counts.Williams_HH6, counts.Williams_HH
7
##  3 dimensional reductions calculated: pca, harmony, umap
```

```
chick <- NormalizeData(chick)
```

```
## Normalizing layer: counts.Pajanoja2023_HH5_1
```

```
## Normalizing layer: counts.Pajanoja2023_HH5_2
```

```
## Normalizing layer: counts.Pajanoja2023_HH7_1
```

```
## Normalizing layer: counts.Pajanoja2023_HH7_2
```

```
## Normalizing layer: counts.Pajanoja2023_HH8_1
```

```
## Normalizing layer: counts.Pajanoja2023_HH8_2
```

```
## Normalizing layer: counts.Pajanoja2023_HH9_1
```

```
## Normalizing layer: counts.Pajanoja2023_HH9_2
```

```
## Normalizing layer: counts.Pajanoja2025_HH11
```

```
## Normalizing layer: counts.Williams_HH4
```

```
## Normalizing layer: counts.Williams_HH5
```

```
## Normalizing layer: counts.Williams_HH6
```

```
## Normalizing layer: counts.Williams_HH7
```

```
chick <- CellCycleScoring(chick, s.features = cc.genes$s.genes, g2m.features = cc.genes$g2m.ge
nes)
```

```
## Warning: The following features are not present in the object: TYMS, CDCA7,
## PRIM1, MLF1IP, RAD51AP1, not searching for symbol synonyms
```

```
## Warning: The following features are not present in the object: MKI67, FAM64A,
## AURKB, KIF20B, CDC25C, CDCA2, CDCA8, PSRC1, CENPA, not searching for symbol
## synonyms
```

```
## Warning: The following features are not present in the object: TYMS, CDCA7,
## PRIM1, MLF1IP, RAD51AP1, not searching for symbol synonyms
```

```
## Warning: The following features are not present in the object: MKI67, FAM64A,
## AURKB, KIF20B, CDC25C, CDCA2, CDCA8, PSRC1, CENPA, not searching for symbol
## synonyms
```

```
## Warning: The following features are not present in the object: TYMS, CDCA7,
```

```
## PRIM1, MLF1IP, RAD51AP1, not searching for symbol synonyms
```

```
## Warning: The following features are not present in the object: MKI67, FAM64A,
## AURKB, KIF20B, CDC25C, CDCA2, CDCA8, PSRC1, CENPA, not searching for symbol
## synonyms
```

```
## Warning: The following features are not present in the object: TYMS, CDCA7,
## PRIM1, MLF1IP, RAD51AP1, not searching for symbol synonyms
```

```
## Warning: The following features are not present in the object: MKI67, FAM64A,
## AURKB, KIF20B, CDC25C, CDCA2, CDCA8, PSRC1, CENPA, not searching for symbol
## synonyms
```

```
## Warning: The following features are not present in the object: TYMS, CDCA7,
## PRIM1, MLF1IP, RAD51AP1, not searching for symbol synonyms
```

```
## Warning: The following features are not present in the object: MKI67, FAM64A,
## AURKB, KIF20B, CDC25C, CDCA2, CDCA8, PSRC1, CENPA, not searching for symbol
## synonyms
```

```
## Warning: The following features are not present in the object: TYMS, CDCA7,
## PRIM1, MLF1IP, RAD51AP1, not searching for symbol synonyms
```

```
## Warning: The following features are not present in the object: MKI67, FAM64A,
## AURKB, KIF20B, CDC25C, CDCA2, CDCA8, PSRC1, CENPA, not searching for symbol
## synonyms
```

```
## Warning: The following features are not present in the object: TYMS, CDCA7,
## PRIM1, MLF1IP, RAD51AP1, not searching for symbol synonyms
```

```
## Warning: The following features are not present in the object: MKI67, FAM64A,
## AURKB, KIF20B, CDC25C, CDCA2, CDCA8, PSRC1, CENPA, not searching for symbol
## synonyms
```

```
## Warning: The following features are not present in the object: TYMS, CDCA7,
## PRIM1, MLF1IP, RAD51AP1, not searching for symbol synonyms
```

```
## Warning: The following features are not present in the object: MKI67, FAM64A,
## AURKB, KIF20B, CDC25C, CDCA2, CDCA8, PSRC1, CENPA, not searching for symbol
## synonyms
```

```
## Warning: The following features are not present in the object: TYMS, CDCA7,
## PRIM1, MLF1IP, RAD51AP1, not searching for symbol synonyms
```

```
## Warning: The following features are not present in the object: MKI67, FAM64A,
```

```
## AURKB, KIF20B, CDC25C, CDCA2, CDCA8, PSRC1, CENPA, not searching for symbol
## synonyms
```

```
## Warning: The following features are not present in the object: TYMS, CDCA7,
## PRIM1, MLF1IP, RAD51AP1, not searching for symbol synonyms
```

```
## Warning: The following features are not present in the object: MKI67, FAM64A,
## AURKB, KIF20B, CDC25C, CDCA2, CDCA8, PSRC1, CENPA, not searching for symbol
## synonyms
```

```
## Warning: The following features are not present in the object: TYMS, CDCA7,
## PRIM1, MLF1IP, RAD51AP1, not searching for symbol synonyms
```

```
## Warning: The following features are not present in the object: MKI67, FAM64A,
## AURKB, KIF20B, CDC25C, CDCA2, CDCA8, PSRC1, CENPA, not searching for symbol
## synonyms
```

```
## Warning: The following features are not present in the object: TYMS, CDCA7,
## PRIM1, MLF1IP, RAD51AP1, not searching for symbol synonyms
```

```
## Warning: The following features are not present in the object: MKI67, FAM64A,
## AURKB, KIF20B, CDC25C, CDCA2, CDCA8, PSRC1, CENPA, not searching for symbol
## synonyms
```

```
## Warning: The following features are not present in the object: TYMS, CDCA7,
## PRIM1, MLF1IP, RAD51AP1, not searching for symbol synonyms
```

```
## Warning: The following features are not present in the object: MKI67, FAM64A,
## AURKB, KIF20B, CDC25C, CDCA2, CDCA8, PSRC1, CENPA, not searching for symbol
## synonyms
```

```
chick <- FindVariableFeatures(chick)
```

```
## Finding variable features for layer counts.Pajanoja2023_HH5_1
```

```
## Finding variable features for layer counts.Pajanoja2023_HH5_2
```

```
## Finding variable features for layer counts.Pajanoja2023_HH7_1
```

```
## Finding variable features for layer counts.Pajanoja2023_HH7_2
```

```
## Finding variable features for layer counts.Pajanoja2023_HH8_1
```

```
## Finding variable features for layer counts.Pajanoja2023_HH8_2
```

```
## Finding variable features for layer counts.Pajanoja2023_HH9_1
```

```
## Finding variable features for layer counts.Pajanoja2023_HH9_2
```

```
## Finding variable features for layer counts.Pajanoja2025_HH11
```

```
## Finding variable features for layer counts.Williams_HH4
```

```
## Finding variable features for layer counts.Williams_HH5
```

```
## Finding variable features for layer counts.Williams_HH6
```

```
## Finding variable features for layer counts.Williams_HH7
```

```
chick <- ScaleData(chick, vars.to.regress = c("percent.mt", "S.Score", "G2M.Score"))
```

```
## Regressing out percent.mt, S.Score, G2M.Score
```

```
## Centering and scaling data matrix
```

```
## Warning: Different features in new layer data than already exists for
## scale.data
```

```
chick <- RunPCA(chick, npcs = 50, verbose = FALSE)

# Harmony Integration
chick <- IntegrateLayers(object = chick, method = HarmonyIntegration, orig.reduction = "pca",
new.reduction = "harmony", assay = "RNA", group.by = "sample_id", verbose = TRUE)
```

```
## Transposing data matrix
```

```
## Using automatic lambda estimation
```

```
## Initializing state using k-means centroids initialization
```

```
## Harmony 1/10
```

```
## Harmony 2/10
```

```
## Harmony 3/10
```

```
## Harmony 4/10
```

```
## Harmony 5/10
```

```
## Harmony converged after 5 iterations
```

```
chick <- FindNeighbors(chick, reduction = "harmony", dims = 1:30)
```

```
## Computing nearest neighbor graph
```

```
## Computing SNN
```

```
chick <- RunUMAP(chick, reduction = "harmony", dims = 1:30)
```

```
## 22:34:24 UMAP embedding parameters a = 0.9922 b = 1.112
```

```
## 22:34:24 Read 82092 rows and found 30 numeric columns
```

```
## 22:34:24 Using Annoy for neighbor search, n_neighbors = 30
```

```
## 22:34:24 Building Annoy index with metric = cosine, n_trees = 50
```

```
## 0%   10   20   30   40   50   60   70   80   90   100%
```

```
## [----|----|----|----|----|----|----|----|----|----|
```

```
## **************************************************|
## 22:34:31 Writing NN index file to temp file C:\Users\RANEES~1\AppData\Local\Temp\RtmpqcWgVb
\file6a2c3e2a7894
## 22:34:31 Searching Annoy index using 1 thread, search_k = 3000
## 22:34:56 Annoy recall = 100%
## 22:34:56 Commencing smooth kNN distance calibration using 1 thread with target n_neighbors
= 30
## 22:35:00 Initializing from normalized Laplacian + noise (using RSpectra)
## 22:35:02 Commencing optimization for 200 epochs, with 3824716 positive edges
## 22:35:02 Using rng type: pcg
## 22:36:16 Optimization finished
```

```
DimPlot(chick)
```

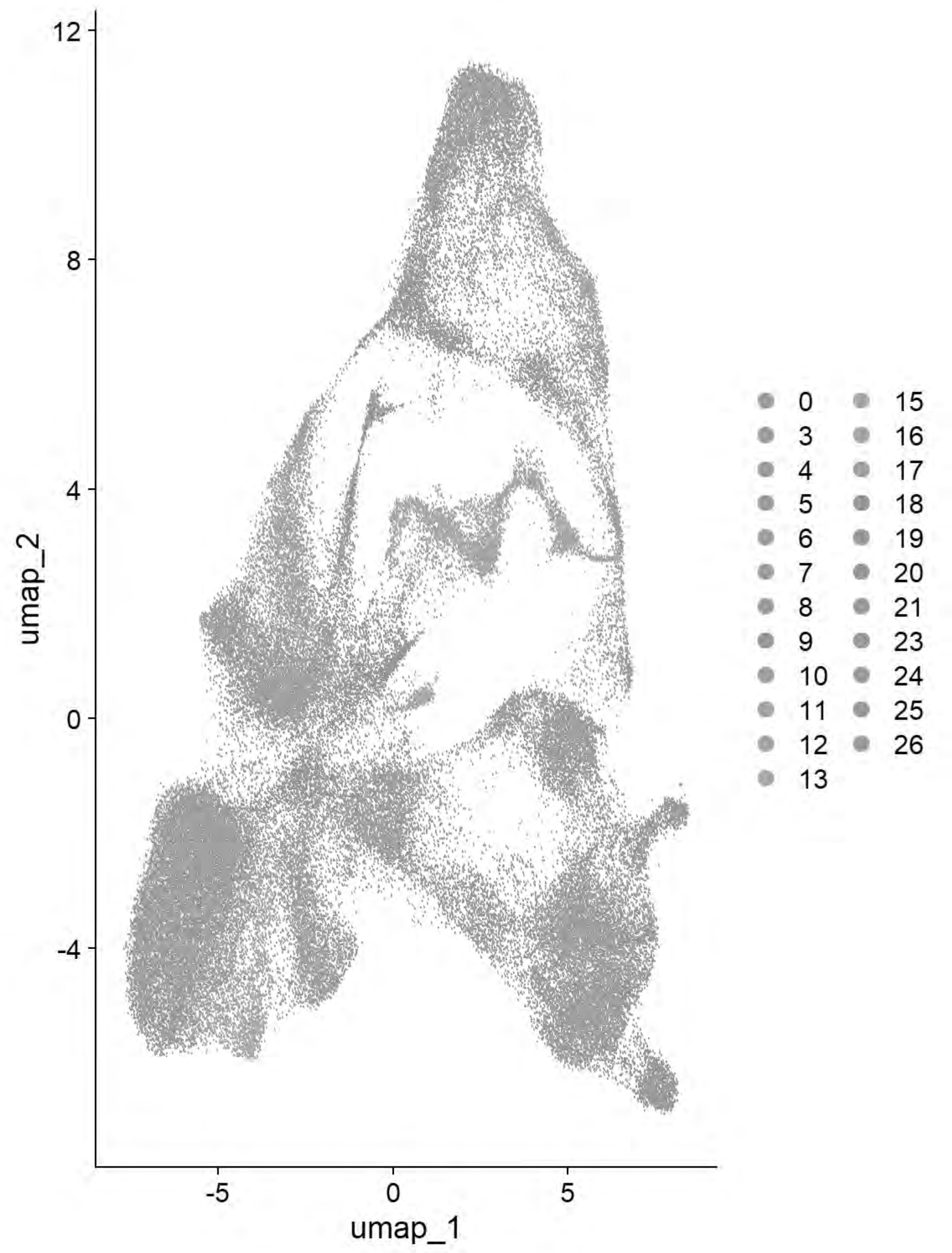


```
chick <- FindClusters(chick, resolution = 0.4)
```

```
## Modularity Optimizer version 1.3.0 by Ludo Waltman and Nees Jan van Eck
##
## Number of nodes: 82092
## Number of edges: 2530742
##
## Running Louvain algorithm...
## Maximum modularity in 10 random starts: 0.9325
```

```
## Number of communities: 16
## Elapsed time: 16 seconds
```

```
# Run QC Sanity Tester
qc_tester(chick)
```

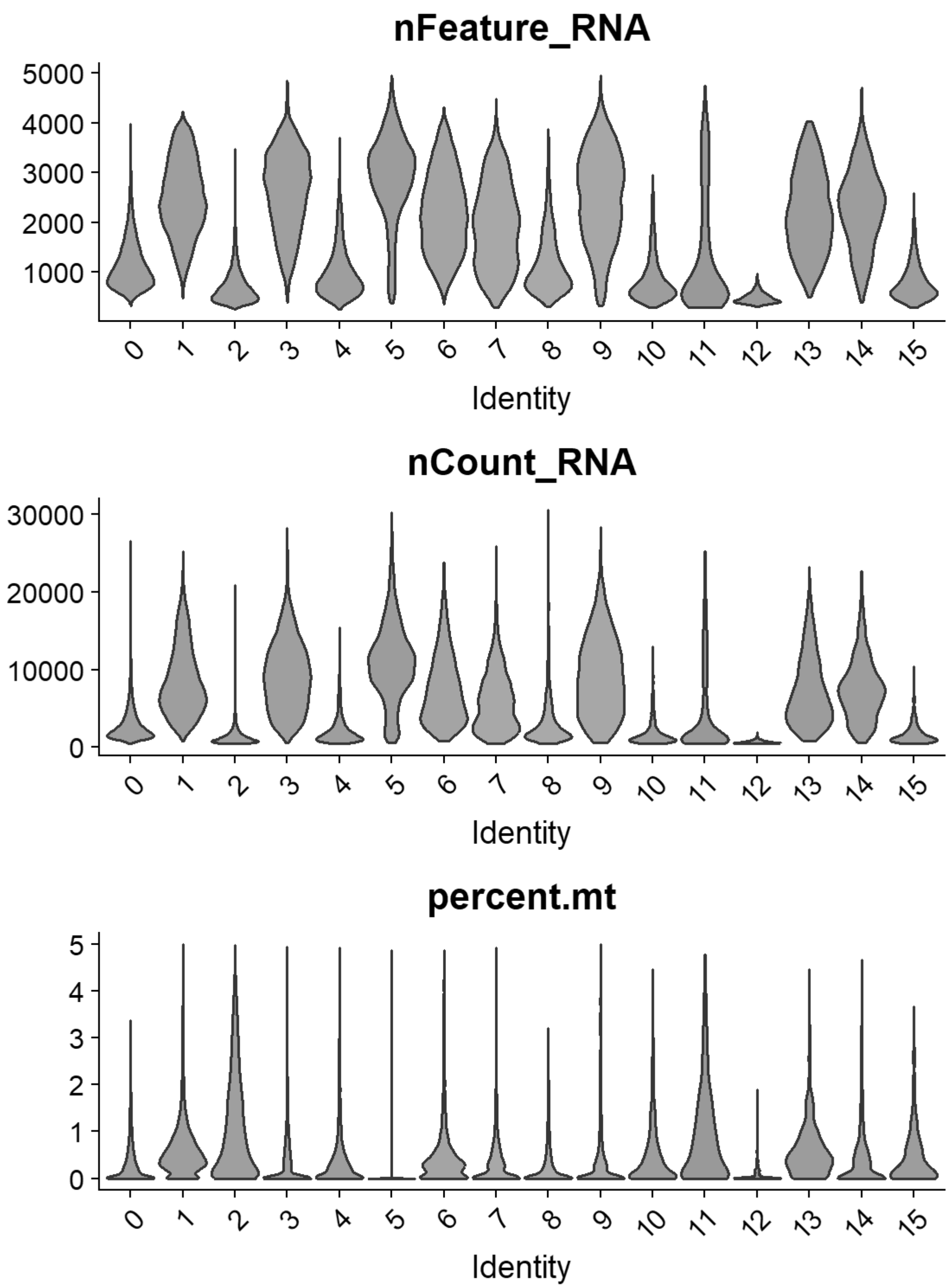


```
chick = FindClusters(chick, resolution= seq(0,2,0.1),n.start=10, verbose = FALSE)
p=clustree(chick, prefix= "RNA_snn_res.")
print(p)
```

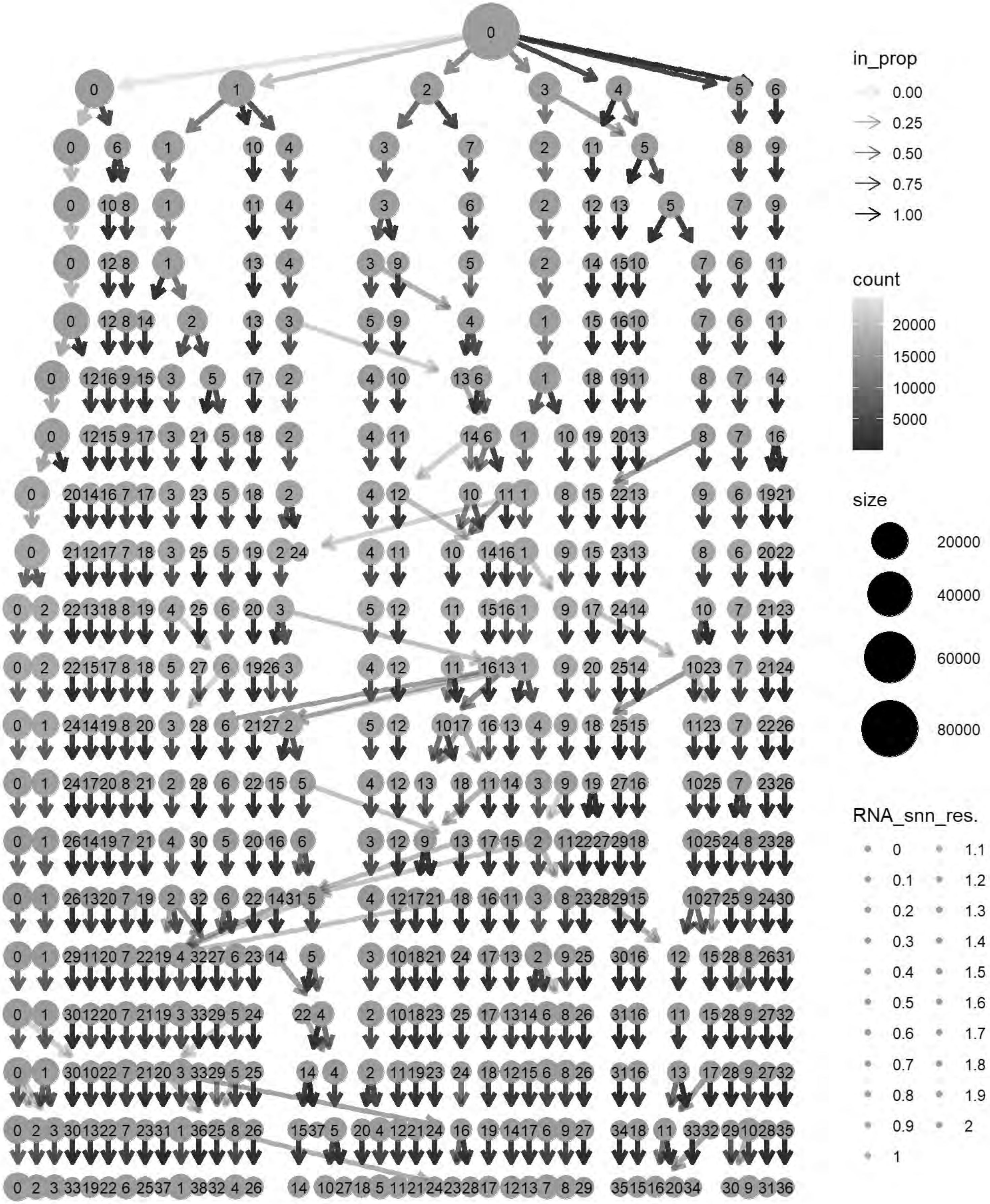
in_prop
0.00
0.25
0.50
0.75
1.00
count
20000
15000
10000
5000
size
20000
40000
60000
80000
RNA_snn_res.
0
0.1
0.2
0.3
0.4
0.5
0.6
0.7
0.8
0.9
1
1.1
1.2
1.3
1.4
1.5
1.6
1.7
1.8
1.9
2

```
cols= grep("^RNA_snn_res.",names(chick@meta.data))
chick@meta.data[cols] <- NULL
chick <- FindClusters(chick, resolution = 0.9, verbose = TRUE)
```

```
## Modularity Optimizer version 1.3.0 by Ludo Waltman and Nees Jan van Eck
##
## Number of nodes: 82092
## Number of edges: 2530742
##
## Running Louvain algorithm...
## Maximum modularity in 10 random starts: 0.8951
## Number of communities: 26
## Elapsed time: 17 seconds
```

```
p6=DimPlot(chick, label = TRUE)+NoLegend()
print(p6)
```

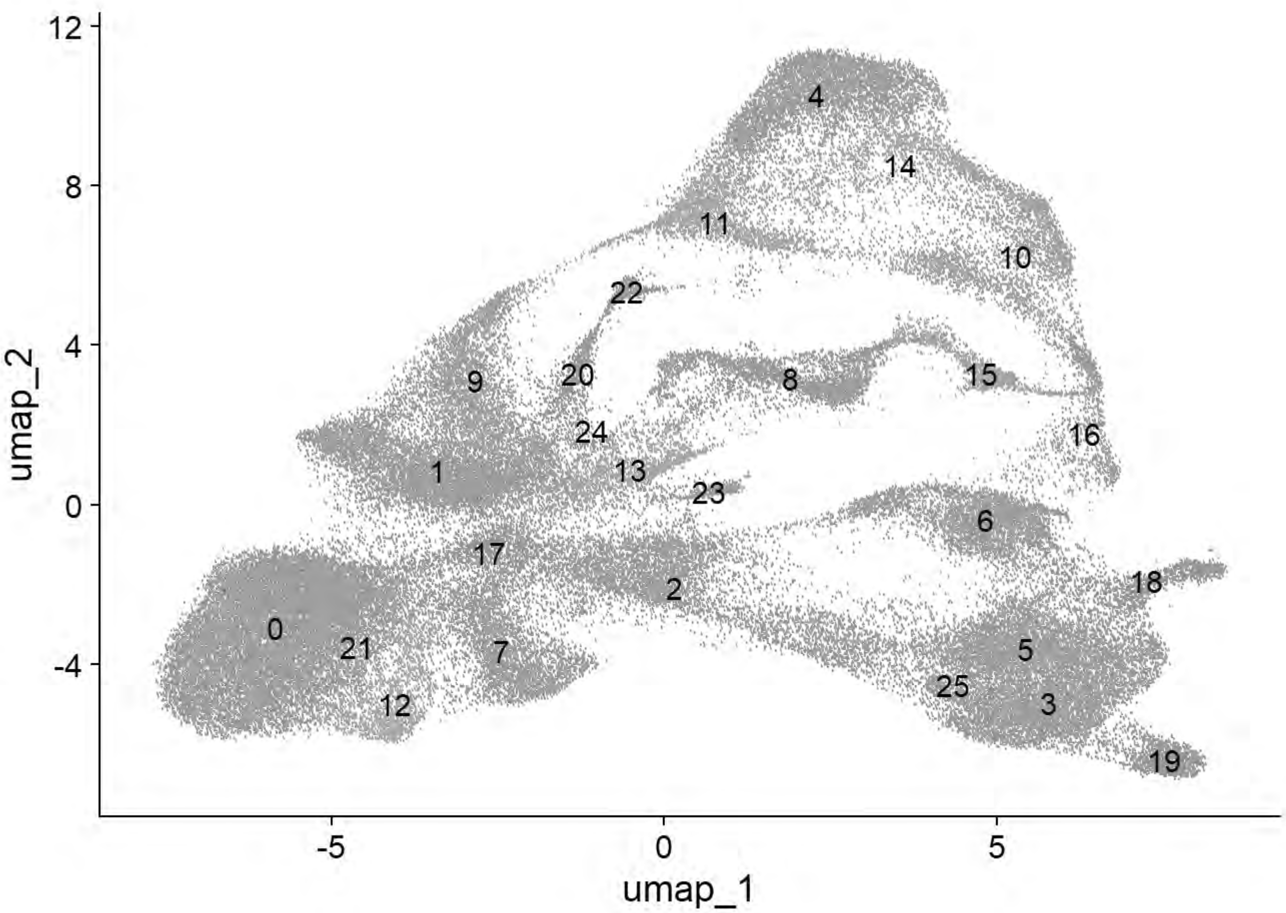


```
p6=DimPlot(chick, group.by = "Phase")
print(p6)
```

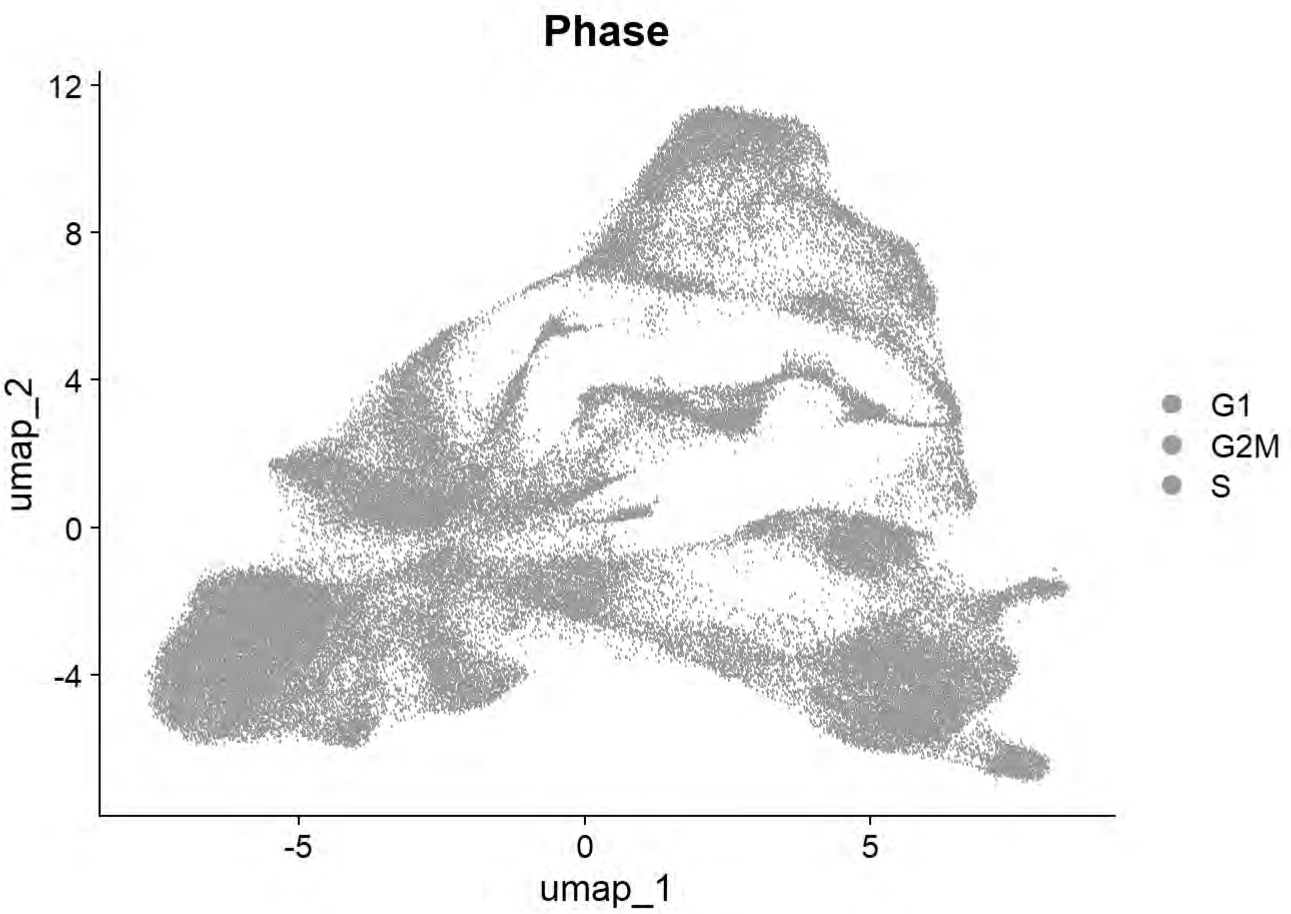


```
p6=DimPlot(chick, group.by = "sample_id")
print(p6)
```

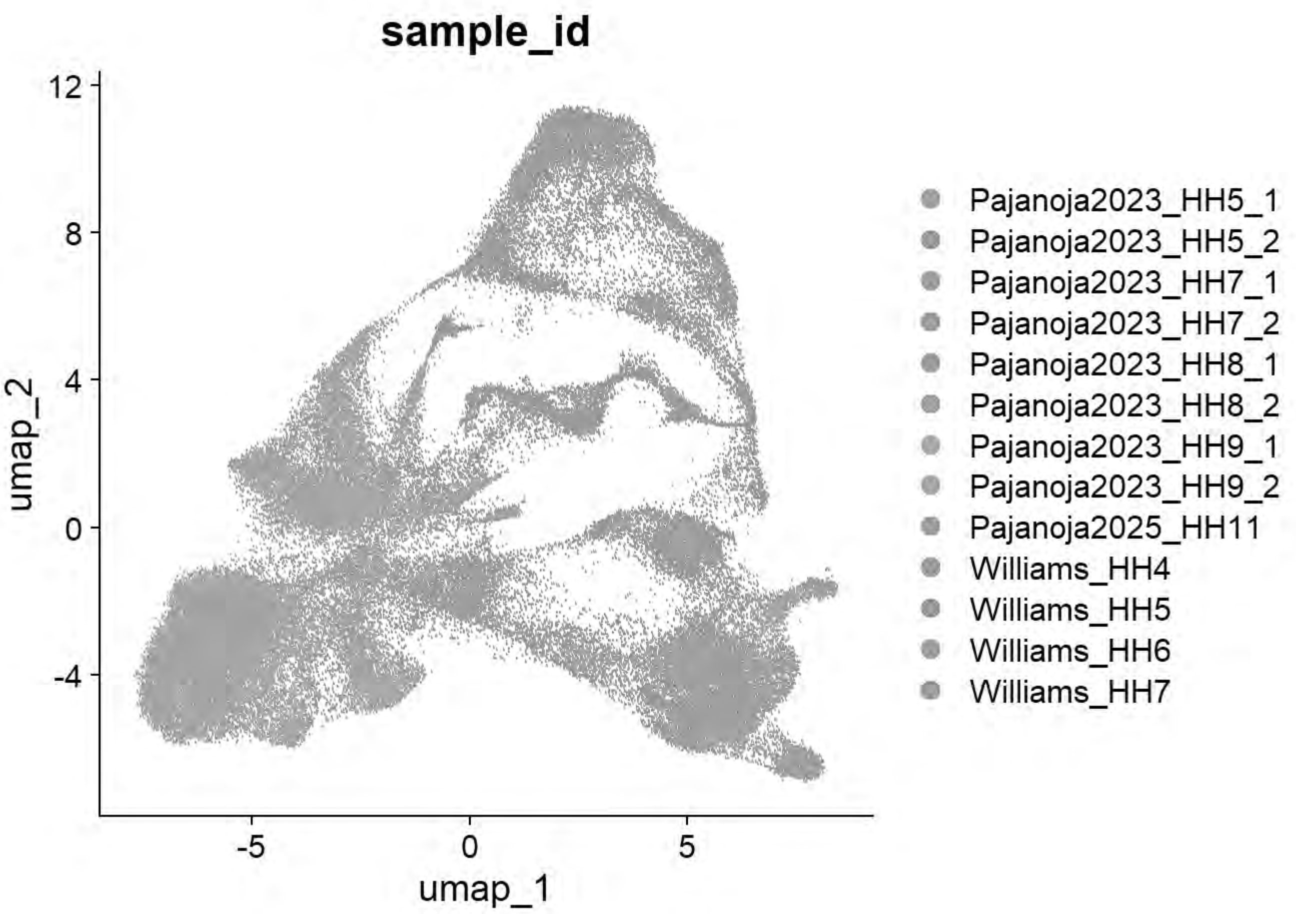


```
table(chick$seurat_clusters,chick$sample_id)
```

```
##
##      Pajanoja2023_HH5_1 Pajanoja2023_HH5_2 Pajanoja2023_HH7_1
##   0                1355                275                897
##   1                 379                476                304
##   2                1064               1109                740
##   3                1377                488                321
##   4                 544                289                294
##   5                 847                240                453
##   6                1286                392                269
##   7                 440                 65                163
##   8                 854                254                193
##   9                 122                113                153
##   10                  1                 12                  0
##   11                183                 57                116
##   12                  6                  0                 22
##   13                307                231                 74
##   14                 42                 50                  1
##   15                286                105                 71
##   16                 47                 29                 26
##   17                 34                135                309
##   18                 72                 16                 39
##   19                207                 67                 80
##   20                  3                  5                  4
```

```
##   21                 33                  5                 21
##   22                  1                  1                  6
##   23                 67                 75                 39
##   24                  7                  8                  1
##   25                  0                  0                  0
##
##      Pajanoja2023_HH7_2 Pajanoja2023_HH8_1 Pajanoja2023_HH8_2
##   0                 522               1120               3590
##   1                 444                585               2957
##   2                 879               1153                197
##   3                 635                745                268
##   4                 391                510                 53
##   5                 358                189                518
##   6                 319                544                117
##   7                 126                343                680
##   8                 160                409                 81
##   9                 238                498               1153
##   10                  1                  1                 53
##   11                193                340                176
##   12                 16                 83                308
##   13                109                191                210
##   14                  6                  1                  3
##   15                120                 94                 37
##   16                 24                 34                  4
##   17                220                460                  6
##   18                 62                164                109
##   19                153                163                 55
##   20                  8                 70                197
##   21                 14                 63                 67
##   22                  5                 53                 68
##   23                 63                 63                 69
##   24                  4                 17                  6
##   25                  1                  0                  0
##
##      Pajanoja2023_HH9_1 Pajanoja2023_HH9_2 Pajanoja2025_HH11 Williams_HH4
##   0                3451               3900              1573            8
##   1                1128               1466               247           32
##   2                 131                135               287          286
##   3                 505                256              1038          155
##   4                 517                363               652           78
##   5                 383                 60              1159          113
##   6                  16                118               297          290
##   7                 825                654               480           12
##   8                   6                 47               167          254
##   9                 492                242                41           26
##   10                  2                  2                68          234
##   11                 83                 49                23          148
##   12                536                606               200            0
##   13                229                202               115           10
##   14                  3                  2                11           44
##   15                 26                 29                48           32
##   16                  2                  0                25           93
##   17                 92                 75                42            3
##   18                140                106               569            3
```

```
##   19                 92                 73                248            9
##   20                215                241                 63            4
##   21                141                196                 39            0
##   22                 37                 48                 70           35
##   23                 48                 56                  1            1
##   24                  3                  1                  2           65
##   25                  0                  0                  5            1
##
##      Williams_HH5 Williams_HH6 Williams_HH7
##   0            12            5           14
##   1            72           23           55
##   2           118          101          158
##   3            66           16          139
##   4           370          510         1074
##   5            22           13          161
##   6            52          102          174
##   7             2            4            2
##   8           308          292          339
##   9             5           11           24
##   10          467         1128          966
##   11          109          452          342
##   12            0            0            0
##   13           17           15           24
##   14          241          487          828
##   15           39          183          400
##   16          188          388          555
##   17            0            0            1
##   18            2            3           35
##   19            3           17           79
##   20            2            9            0
##   21            0            3            1
##   22           19          116           58
##   23            4           10           17
##   24          105          120          107
##   25           31           64          174
```

```
qc_tester(chick)
```

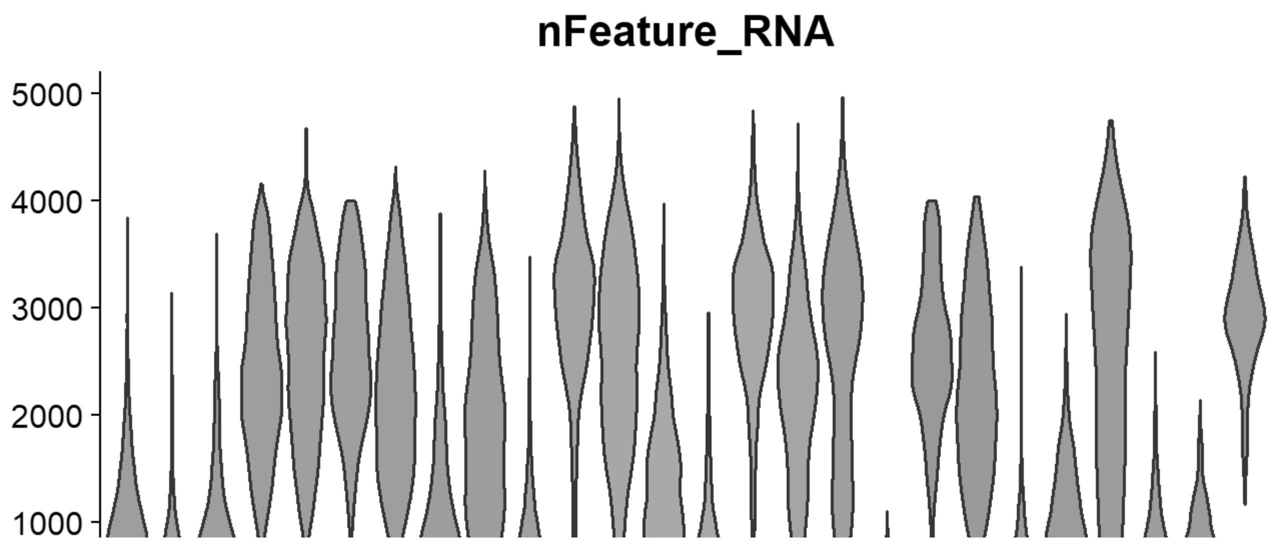

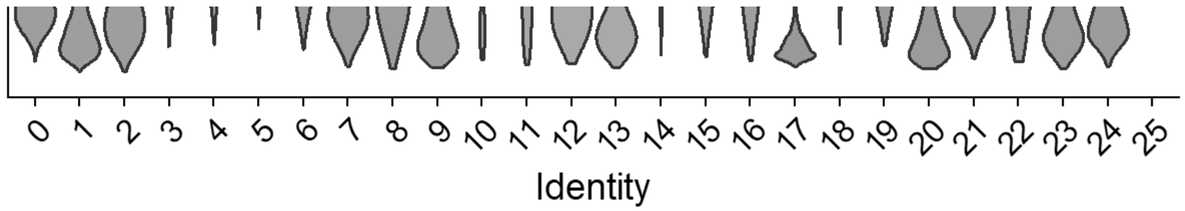


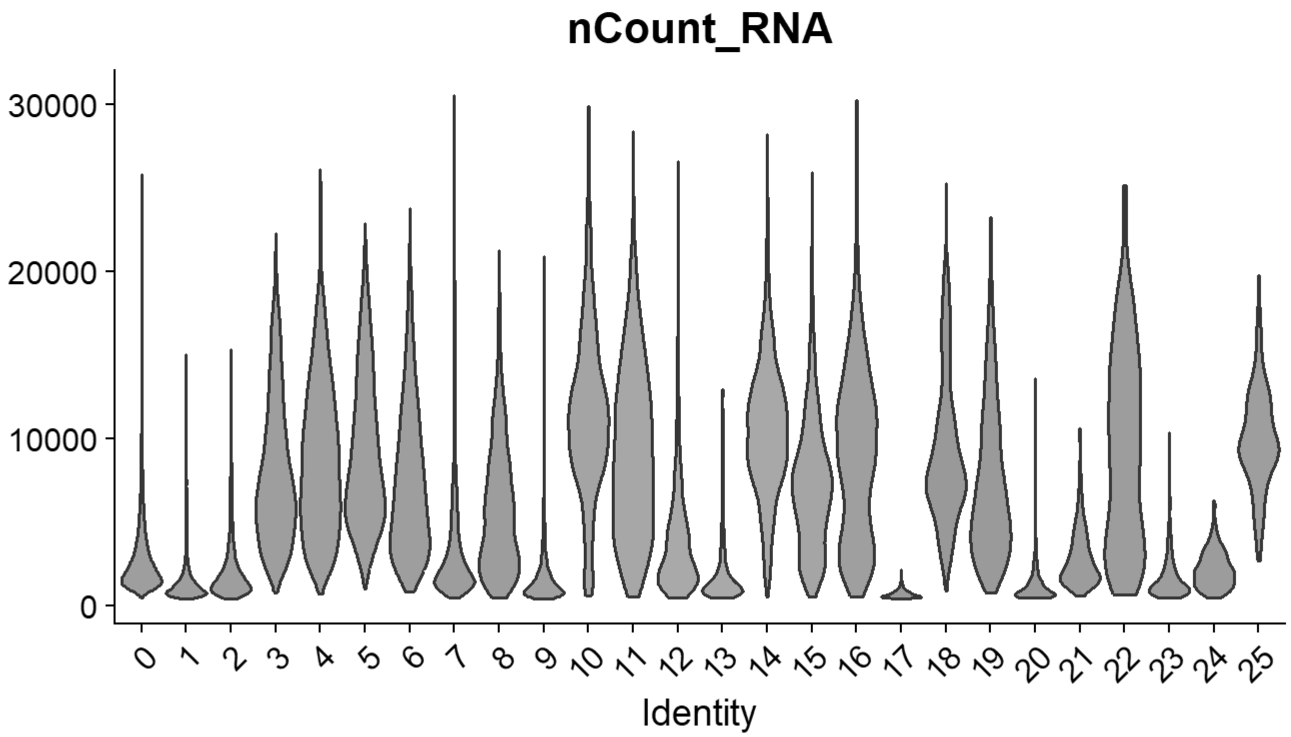


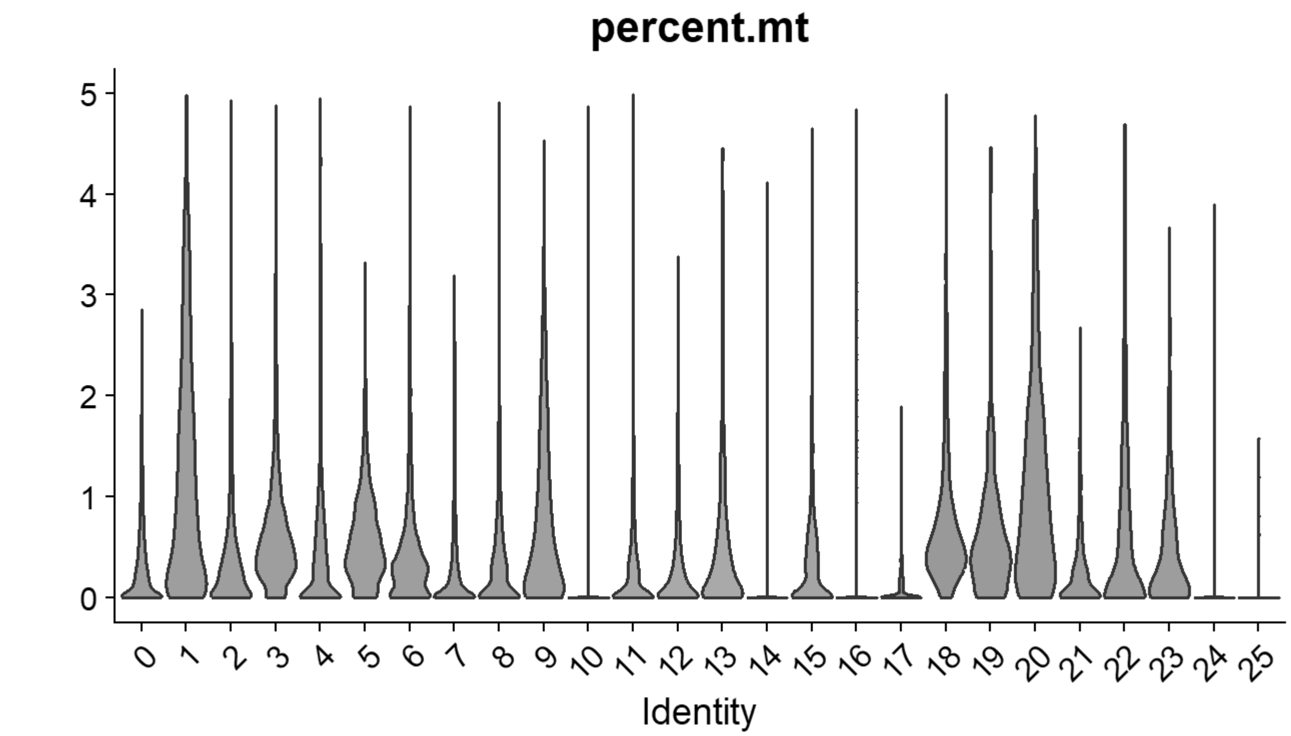


```
# Williams: Neural plate/tube
FeaturePlot(chick, features = c("CLDN1","FRZB","DNMT3A"))
```

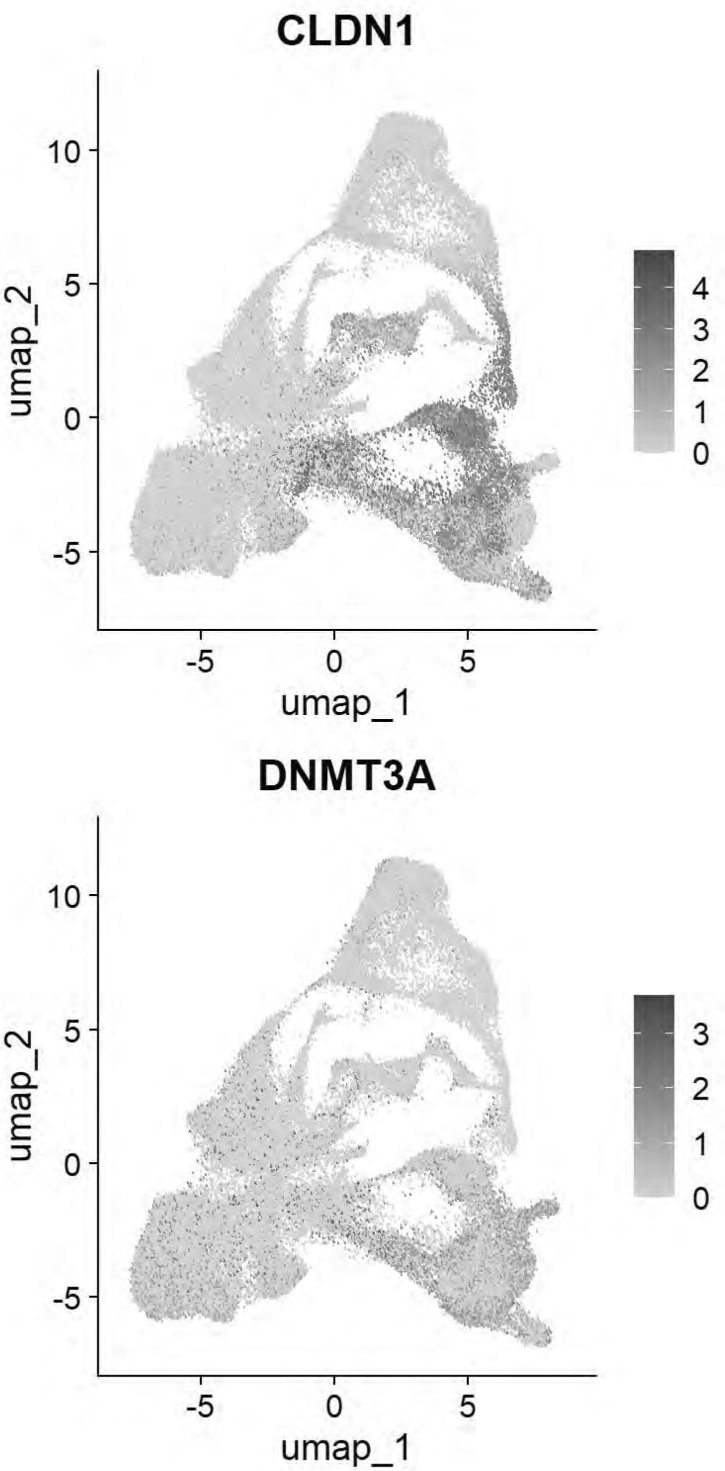


```r
# Williams: Non-Neural Ectoderm
FeaturePlot(chick, features = c("DLX5","TFAP2A","ASTL","PAX6"))
```

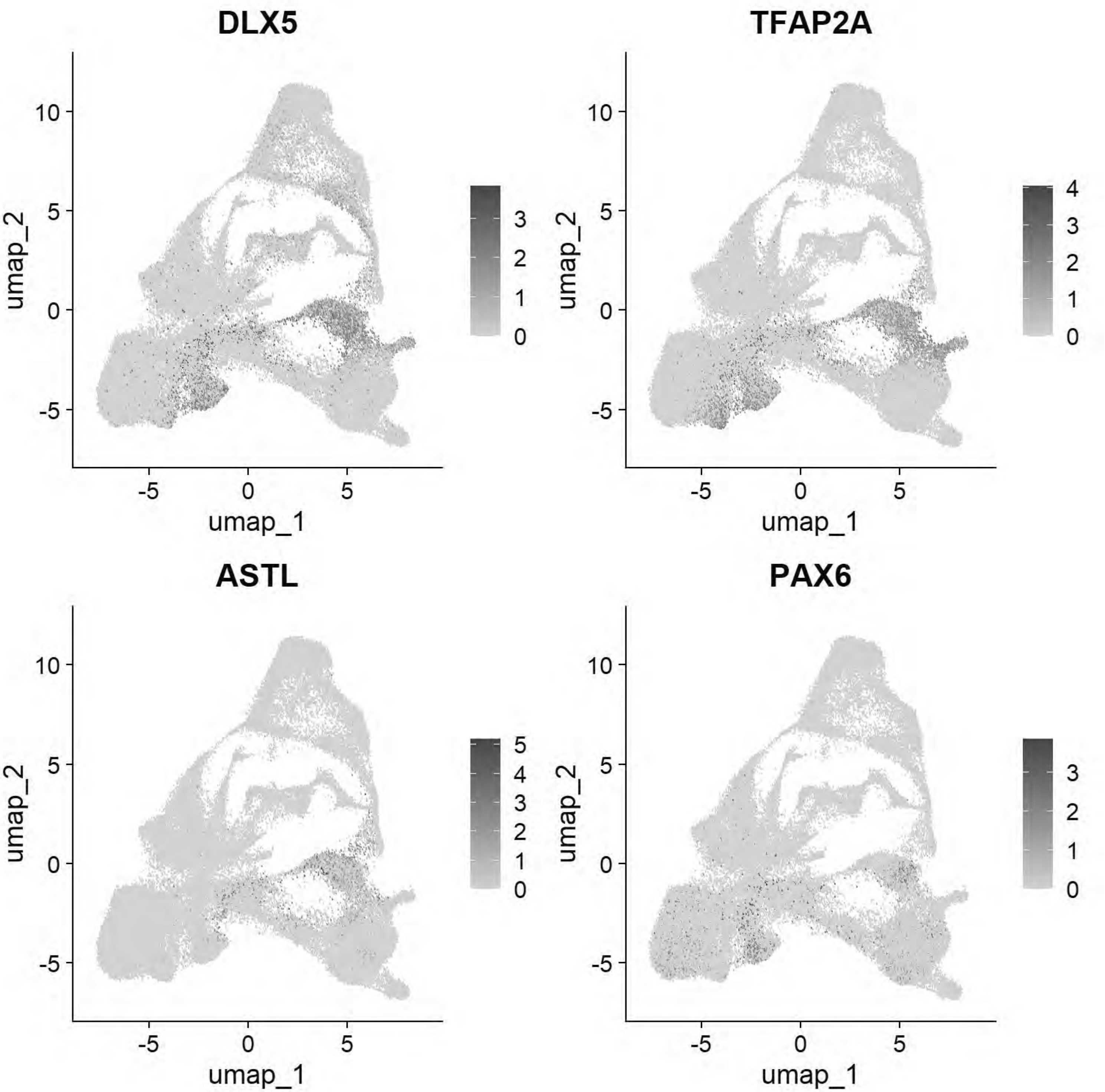


```r
# Williams: Ectoderm (Neural Plate)
FeaturePlot(chick, features = c("SOX2","SOX3","SOX21","FRZB","SFRP2"))
```

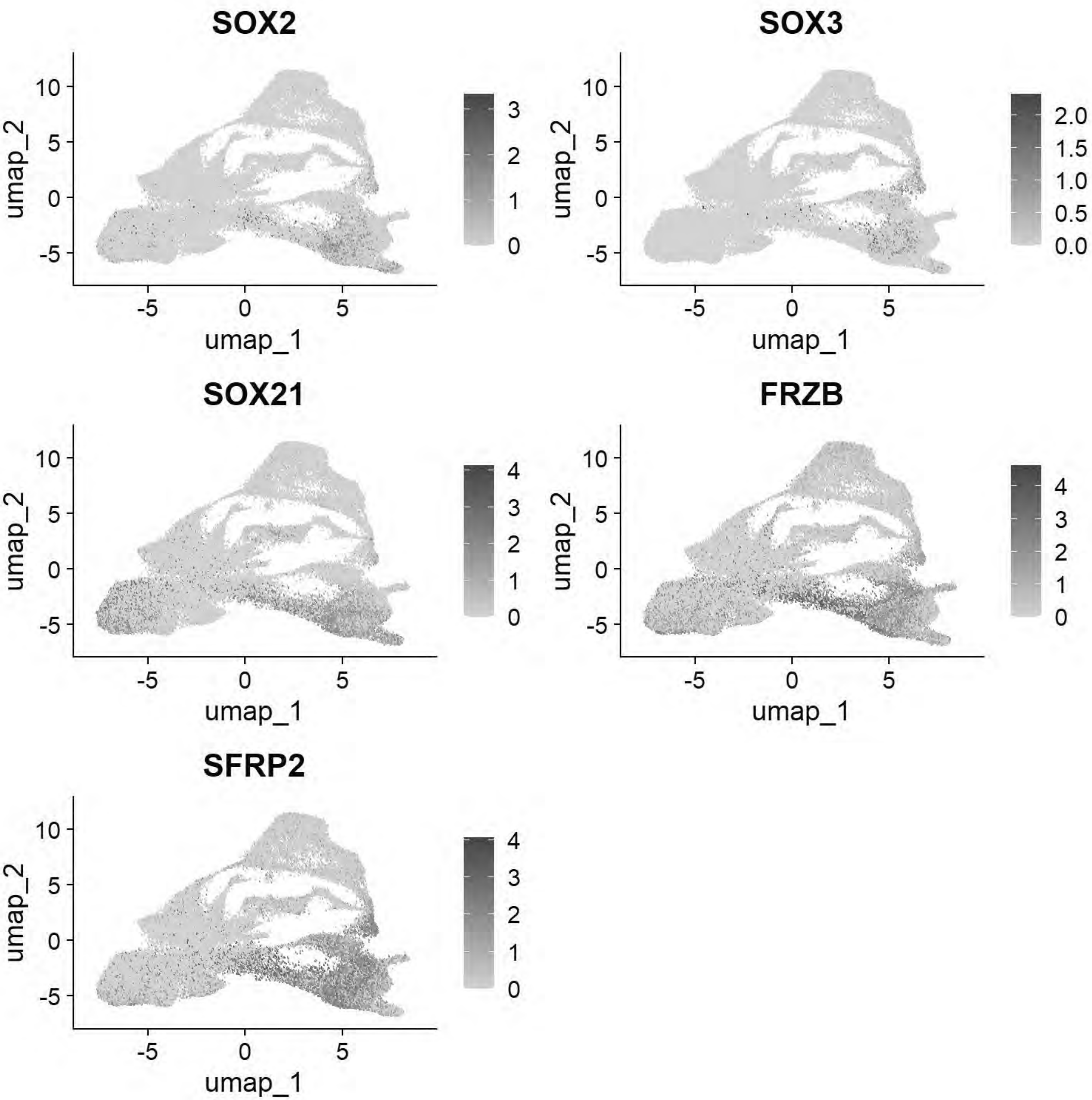


```
# Williams: Ectoderm (Neural Plate Border)
FeaturePlot(chick, features = c("PAX7","TFAP2A","DLX5","BMP4","MSX1","DRAXIN","TFAP2B"))
```

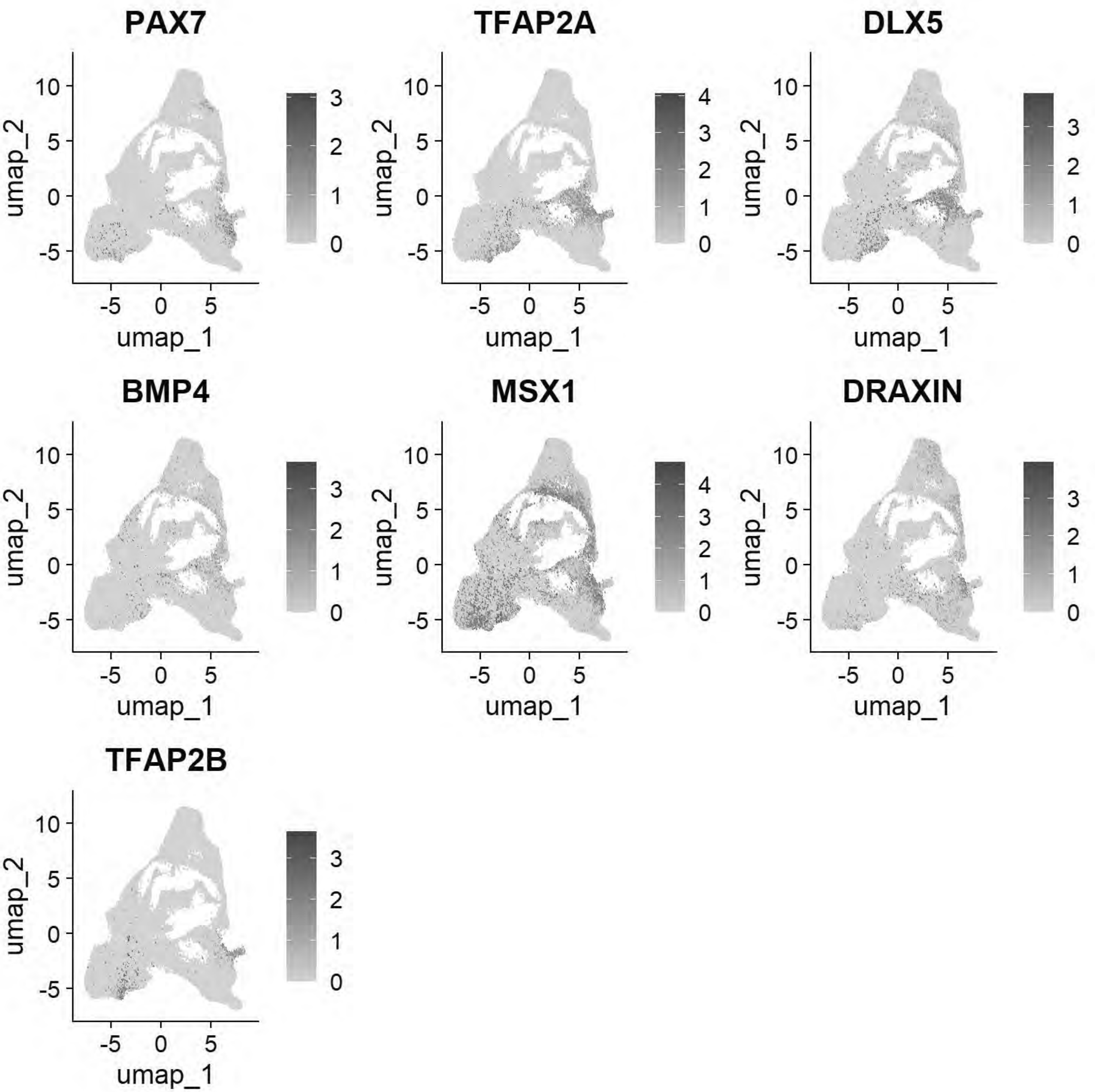


```
# Williams:Epiblast stem cell
FeaturePlot(chick, features = c("ID3","SALL4","TGIF1","ELAVL1"))
```

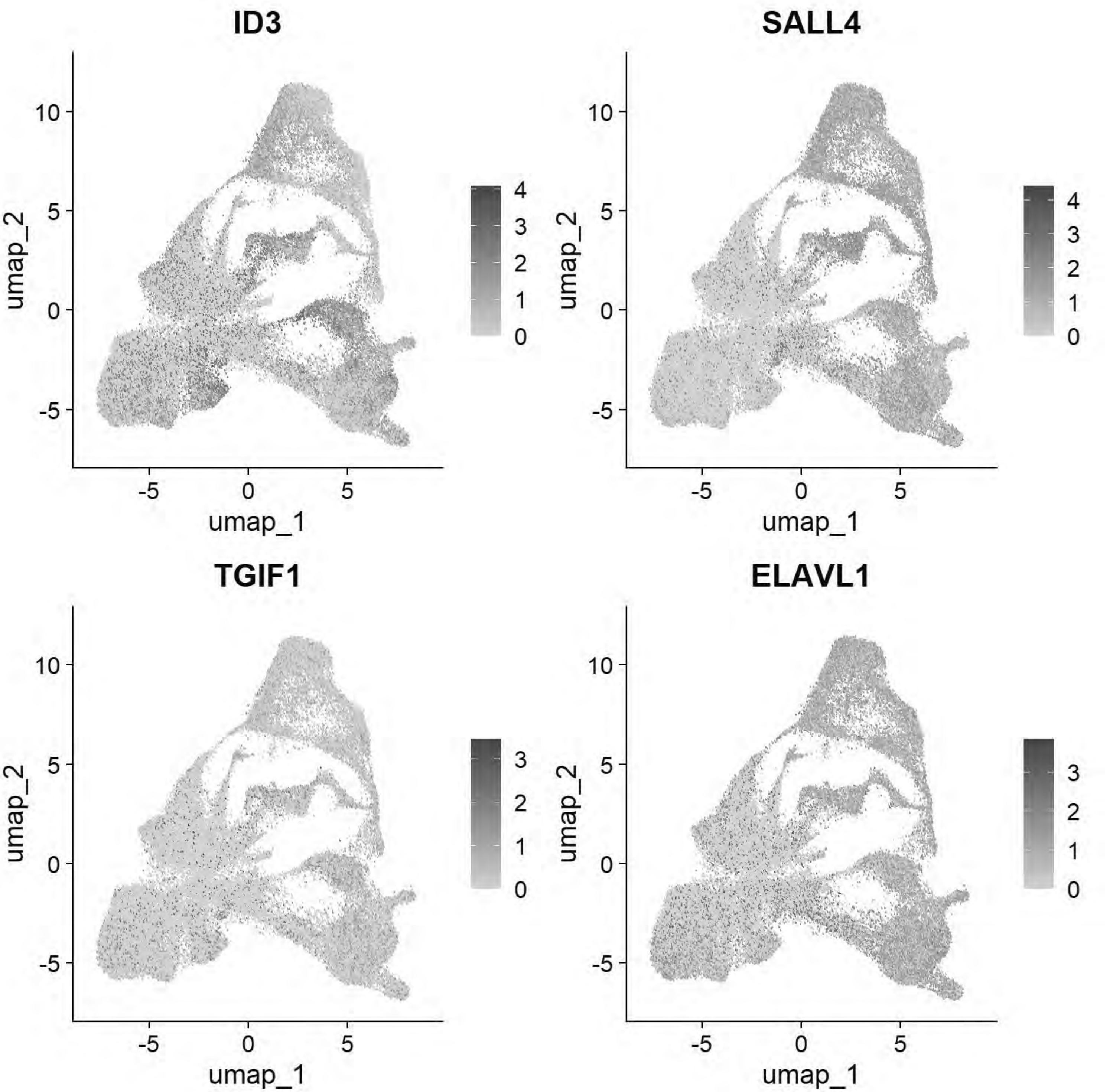


```
# Williams: Posterior Lateral Plate Mesoderm
FeaturePlot(chick, features = c("GATA2","HOXB5","CDX4"))
```

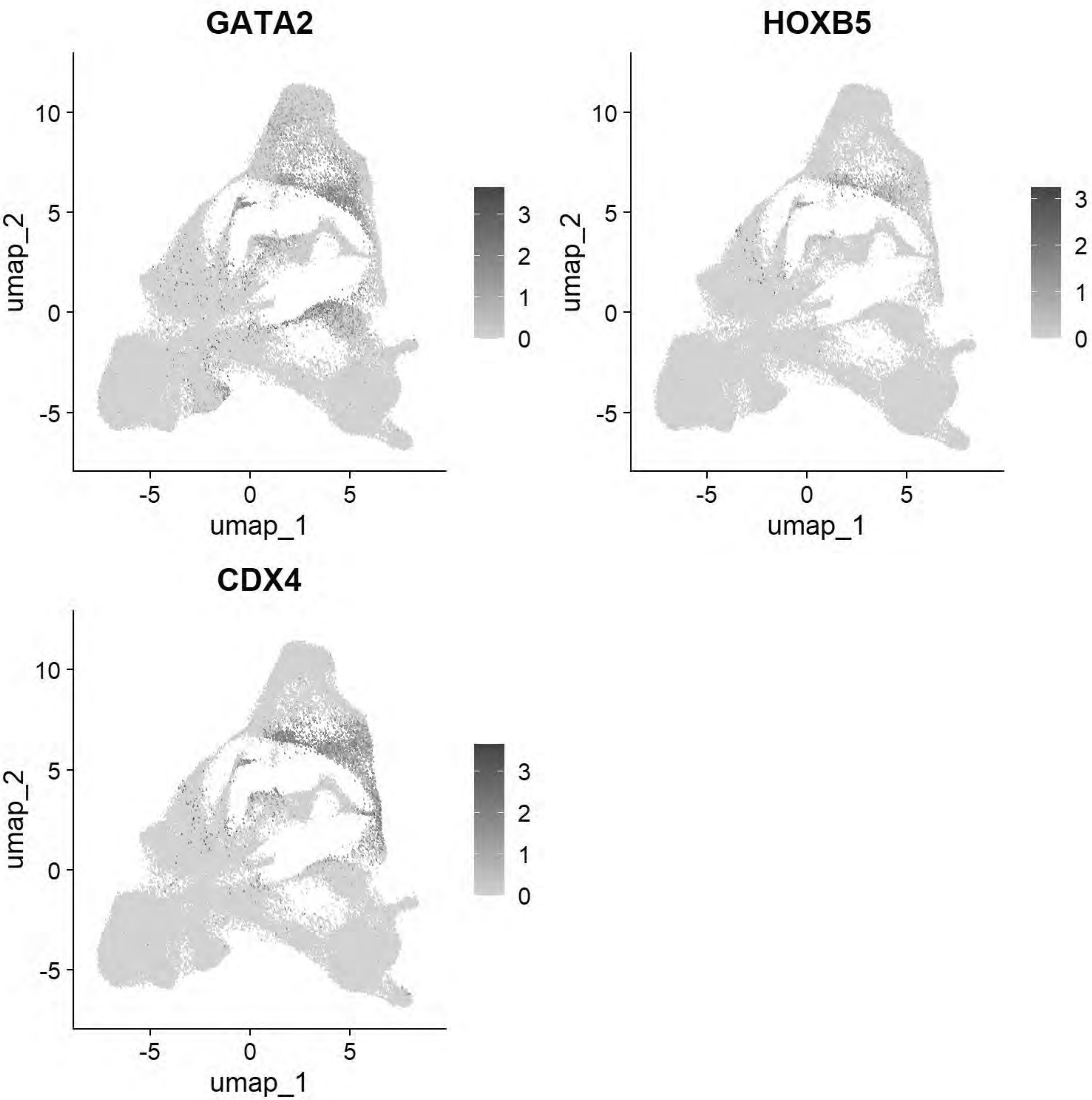


```
# Williams: Paraxial Mesoderm
FeaturePlot(chick, features = c("MSGN1","MESP1","MEOX1"))
```

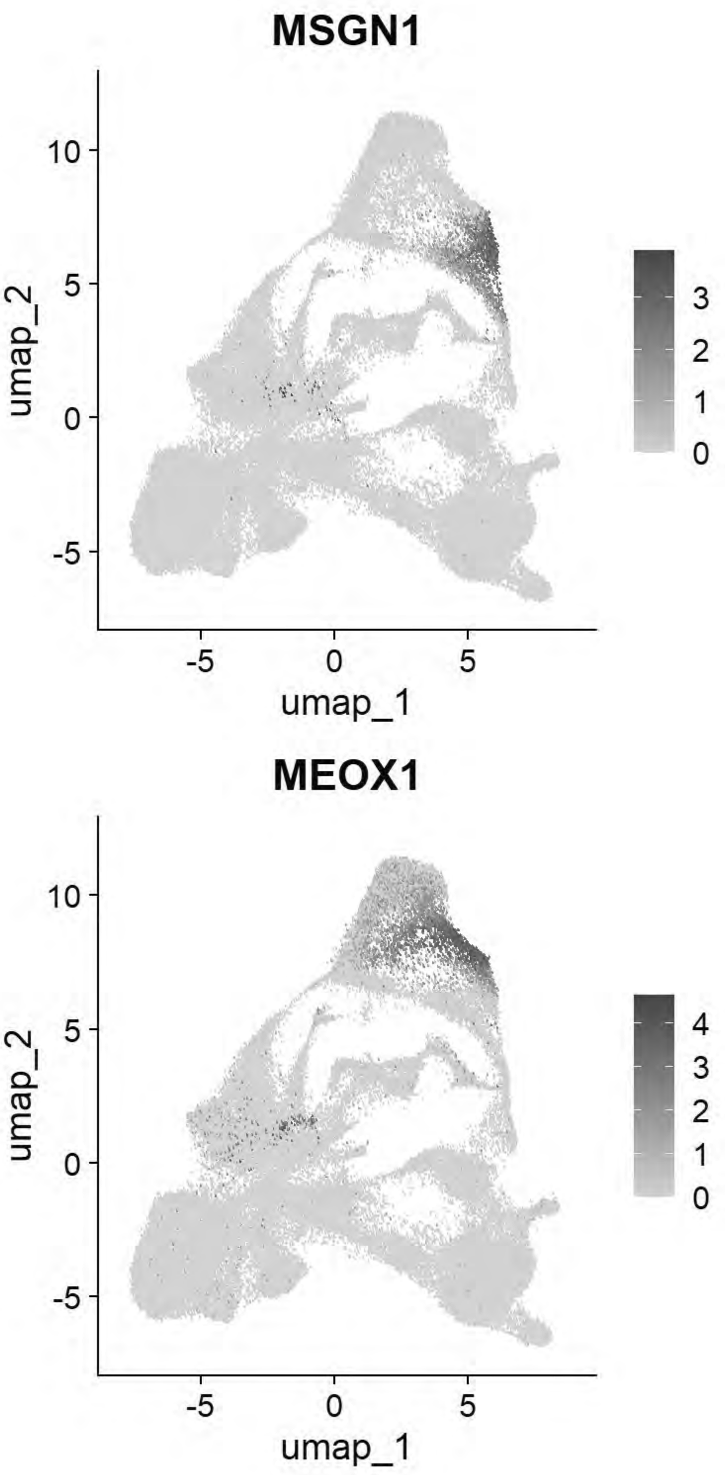


```
# Williams: Lateral Plate Mesoderm
FeaturePlot(chick, features = c("PITX2","ALX1","OLFML3","SIX1","TWIST1"))
```

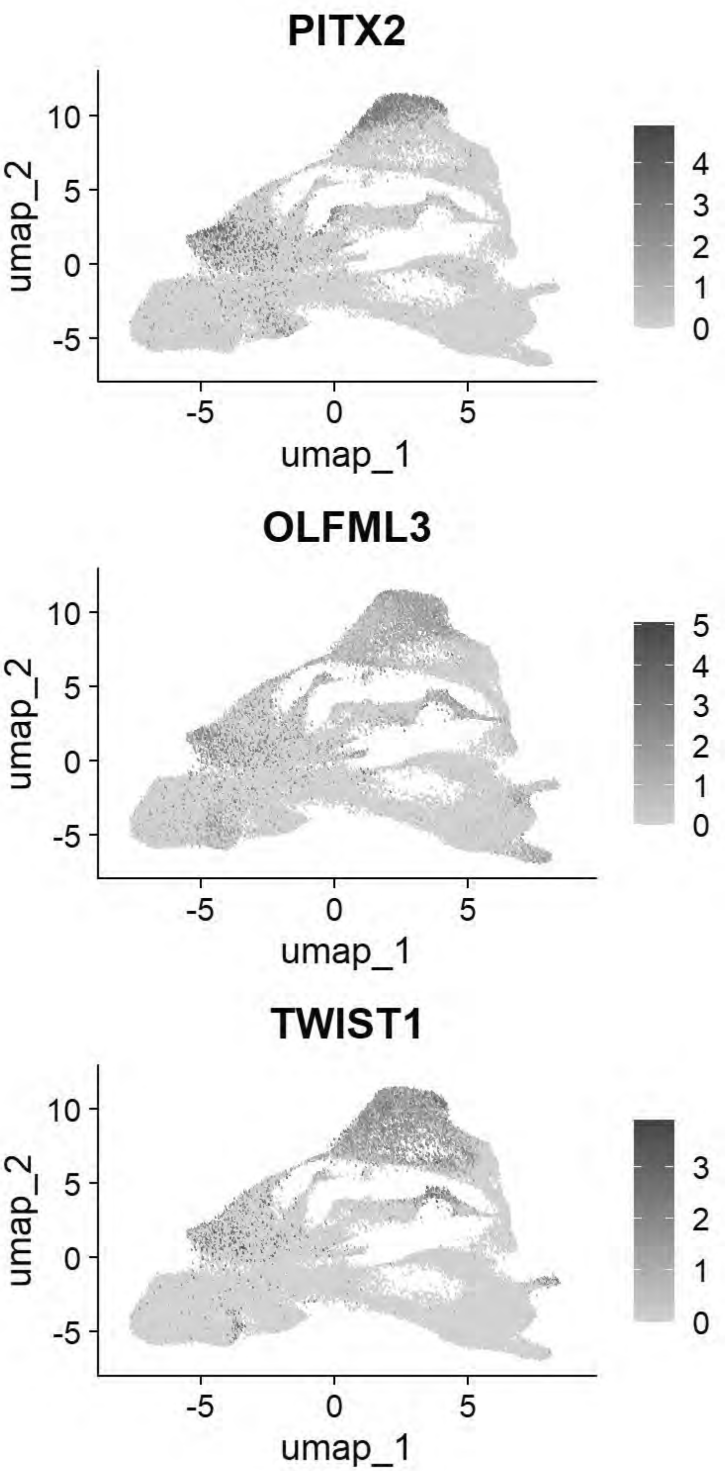


```
# Williams: Cardiac Mesoderm
FeaturePlot(chick, features = c("TCF21","GATA5","LMO2","ETS1","KDR"))
```

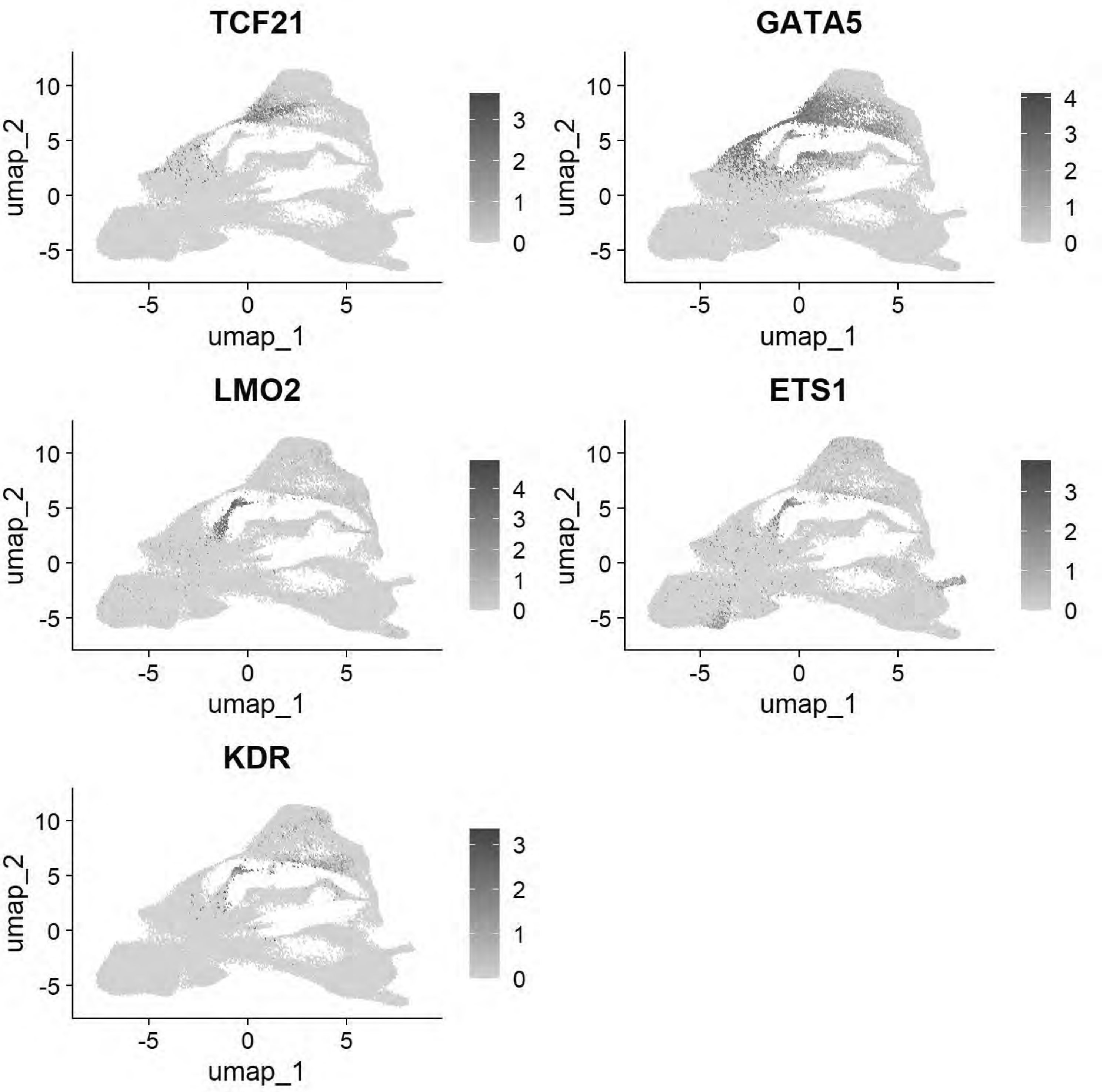


```
# Williams: Head Mesenchyme
FeaturePlot(chick, features = c("TCF21","GATA5","LMO2","ETS1","KDR"))
```

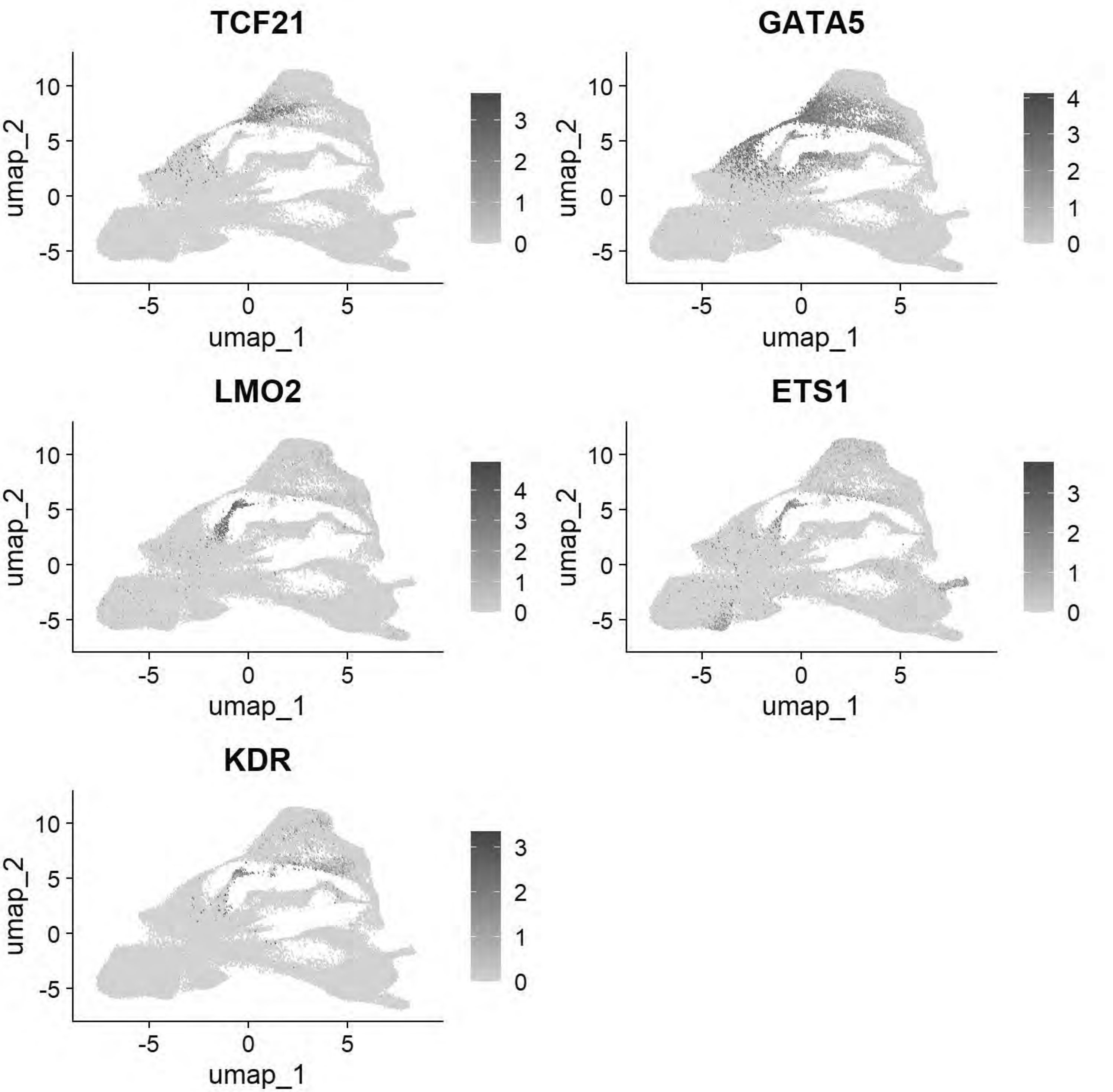


```
# Williams: Hensen's Node & Primitive Streak
FeaturePlot(chick, features = c("DLL1","FGF8","NOTO","CHRD"))
```

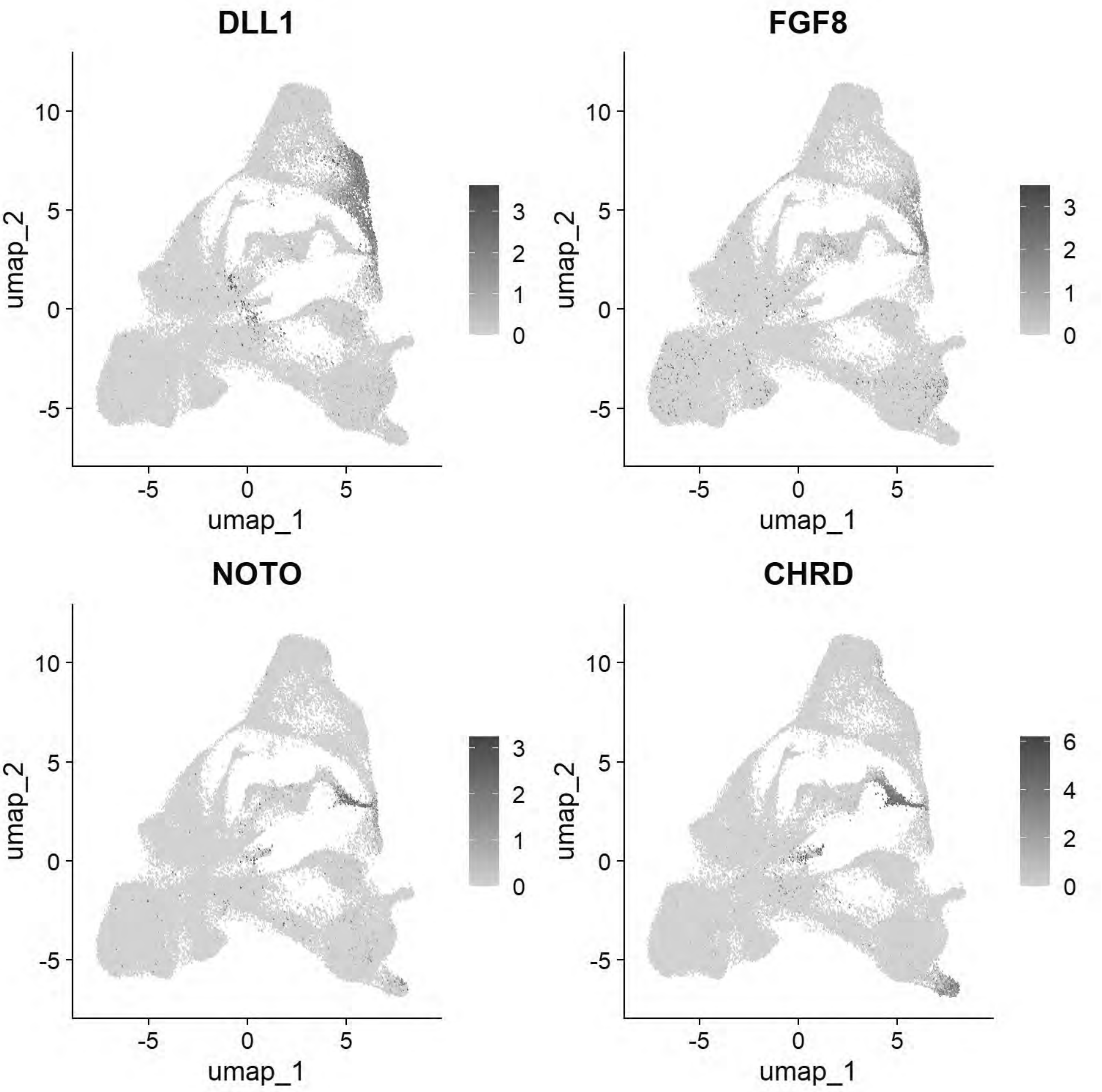


```
# Williams: Endoderm
FeaturePlot(chick, features = c("SOX17","FOXA2","CXCR4"))
```

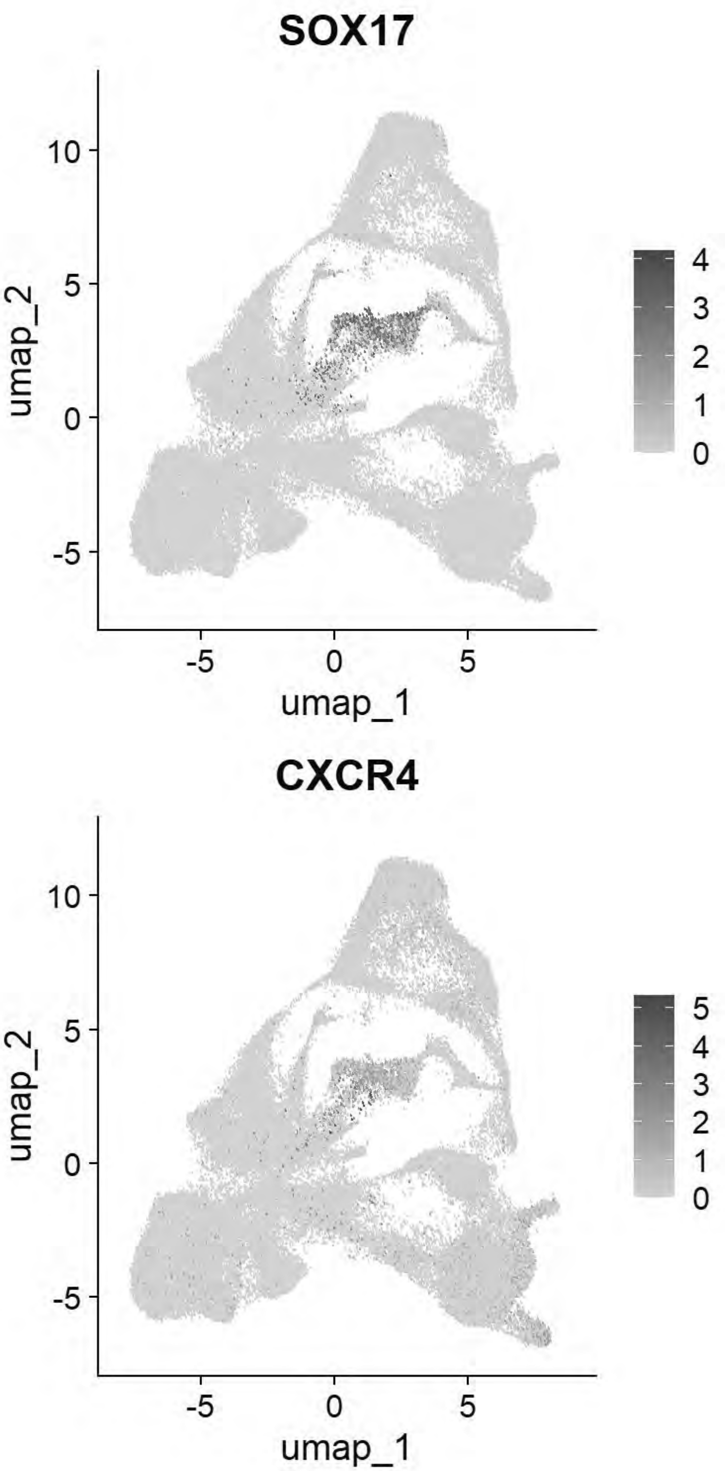


```
# Pajanoja: Ectoderm
FeaturePlot(chick, features = c("TFAP2A","DLX5","CLDN1","SOX2","NESTIN","MYCN"))
```

```
## Warning: The following requested variables were not found: NESTIN
```

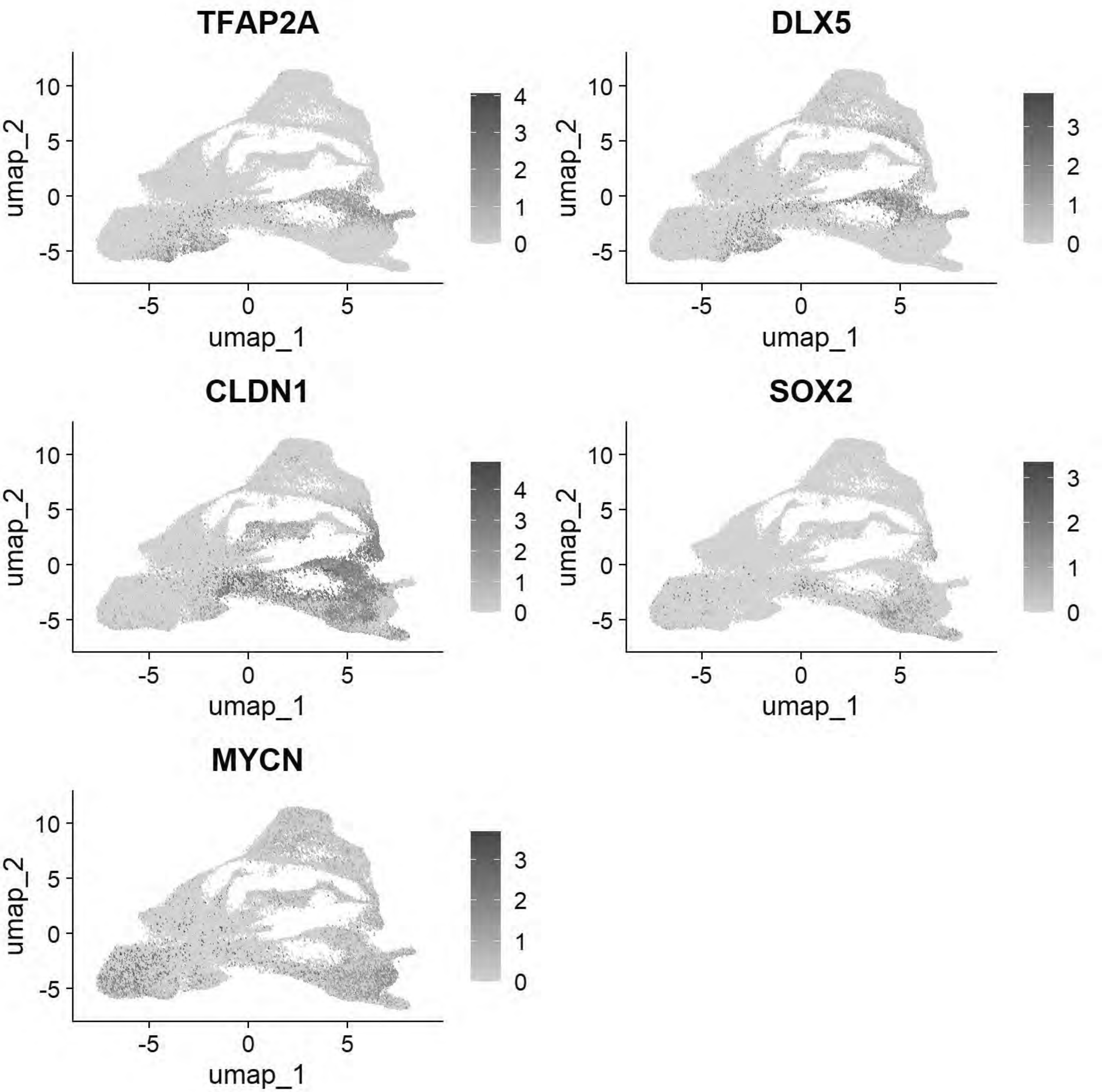


```
# Pajanoja: Endoderm
FeaturePlot(chick, features = c("SOX17","KRT17","HHEX"))
```

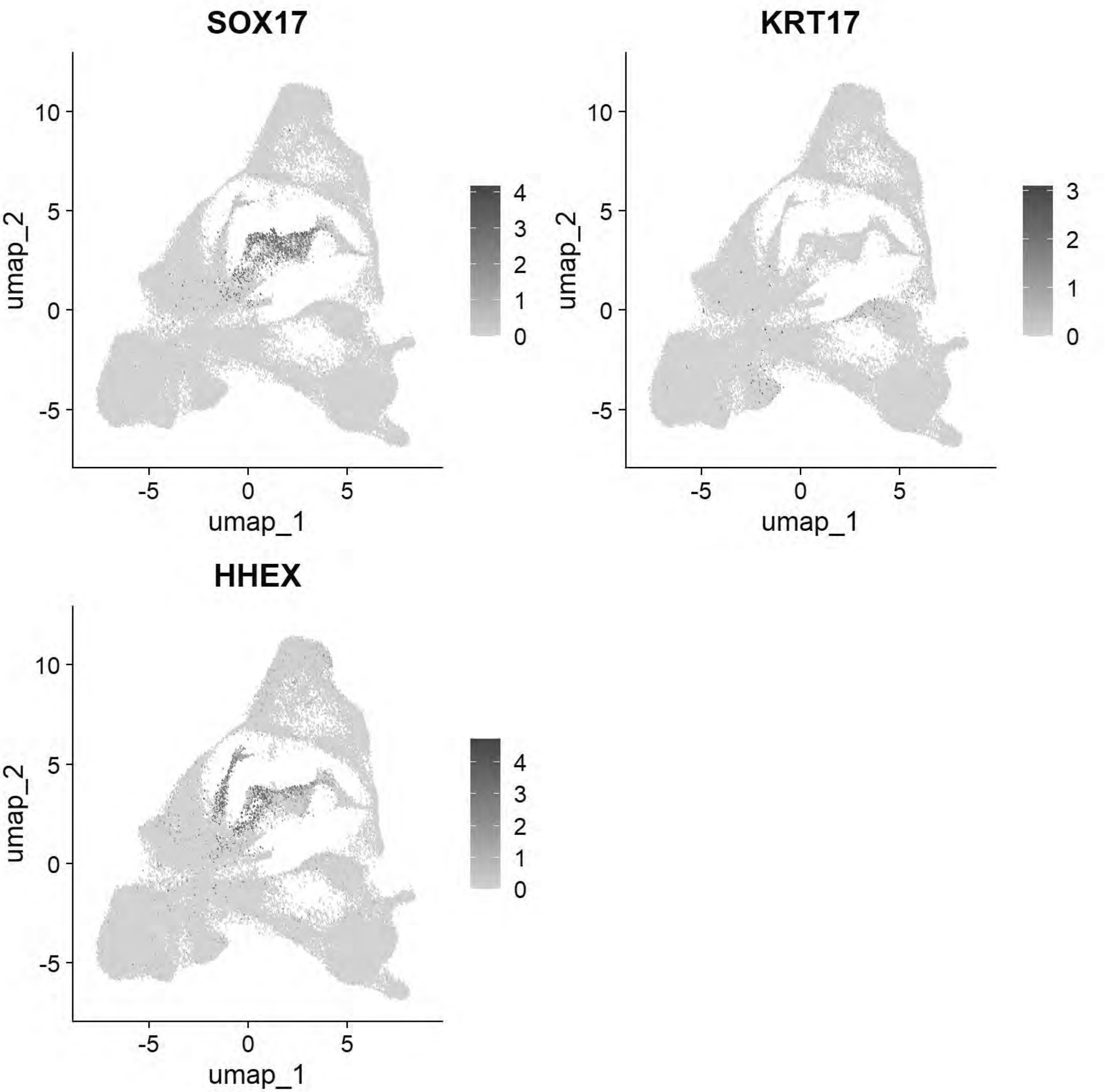


```
# Pajanoja: Mesoderm
FeaturePlot(chick, features = c("ALX1","TWIST1","HAND2","PITX2"))
```

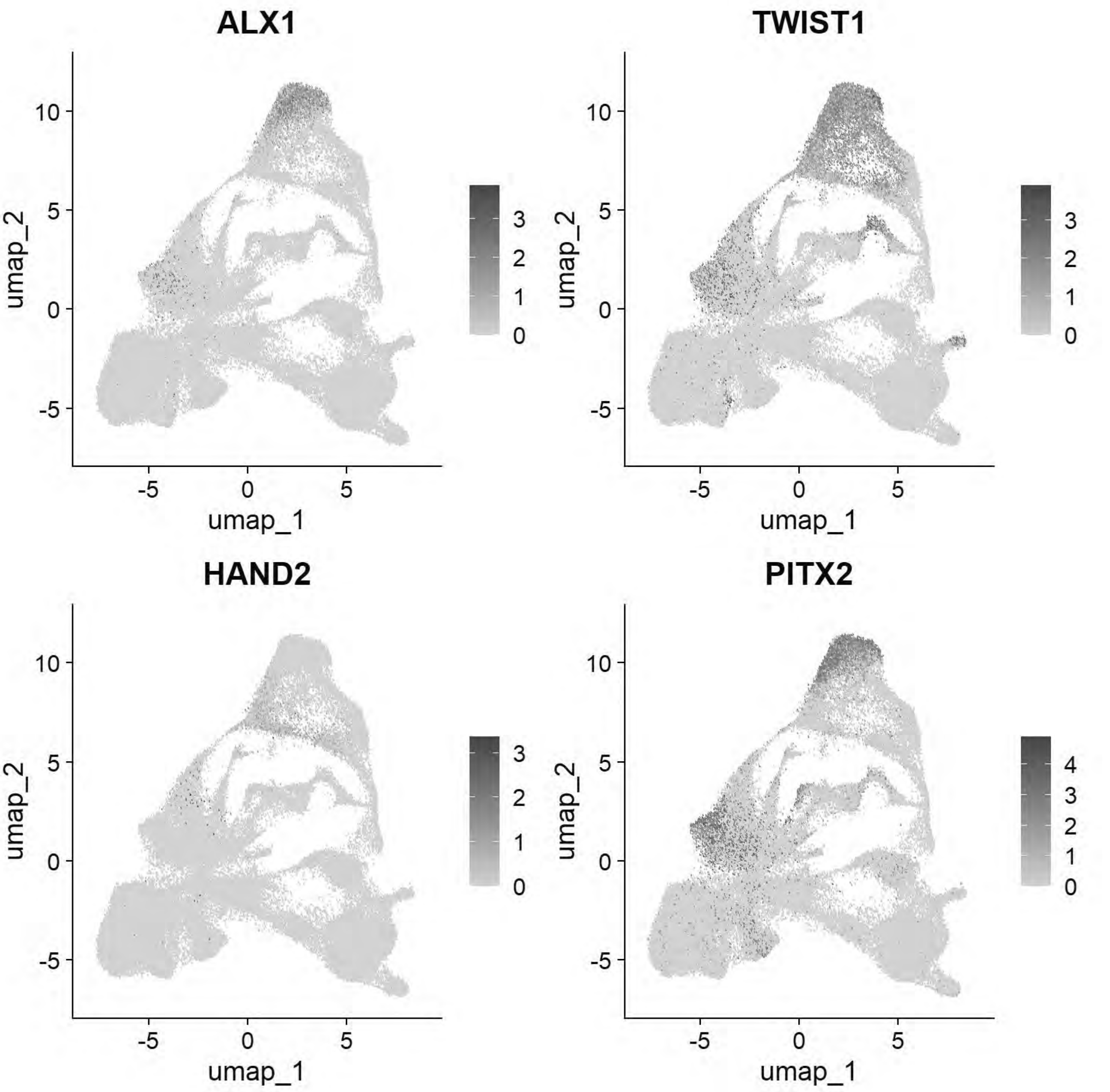


```
# Pajanoja: Notochord
FeaturePlot(chick, features = c("CHRD","TBXT","NOTO"))
```

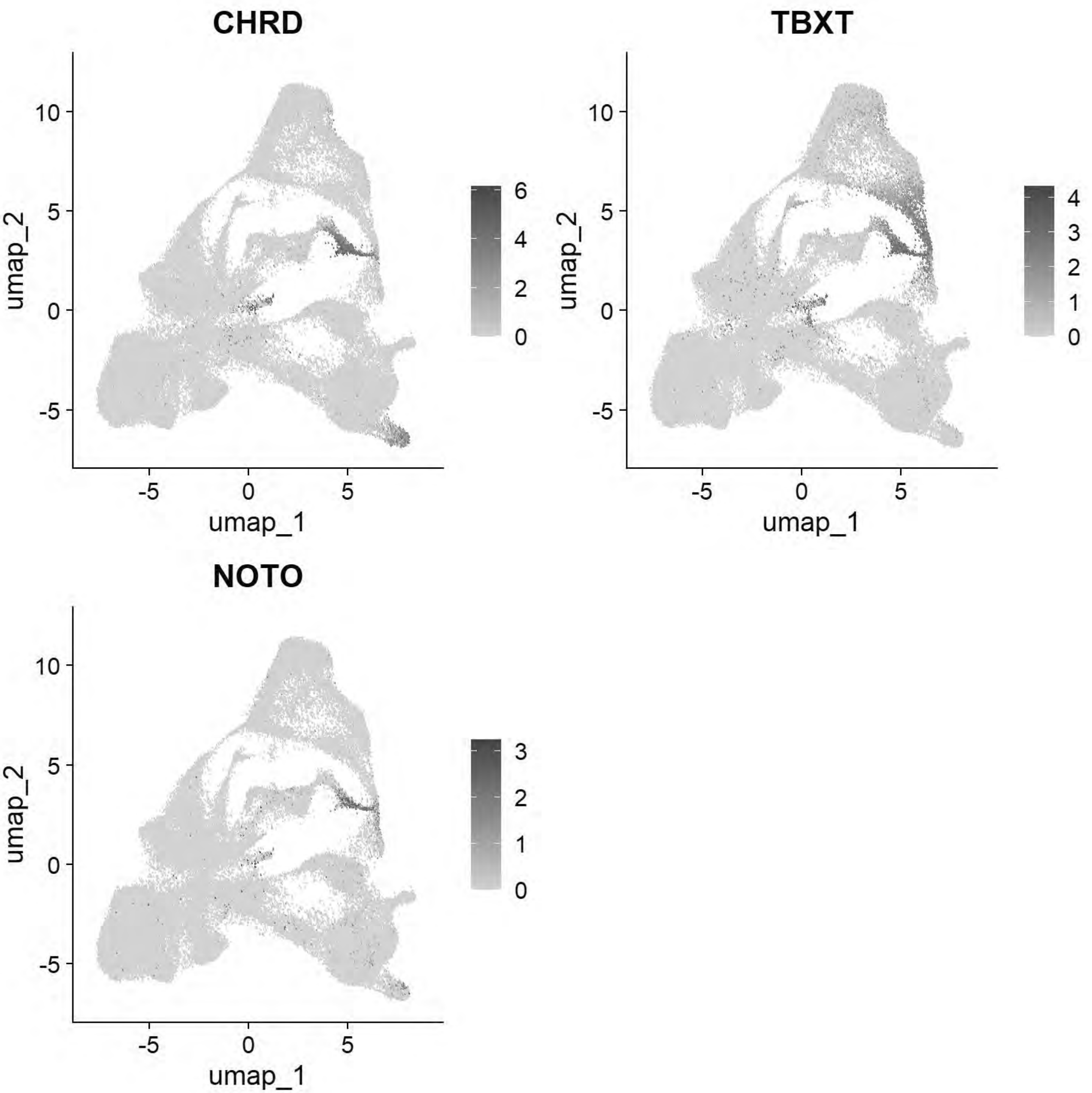


```
# Pajanoja: Ventral Neural Tube
FeaturePlot(chick, features=c("SHH","FOXA1","FOXA2"))
```

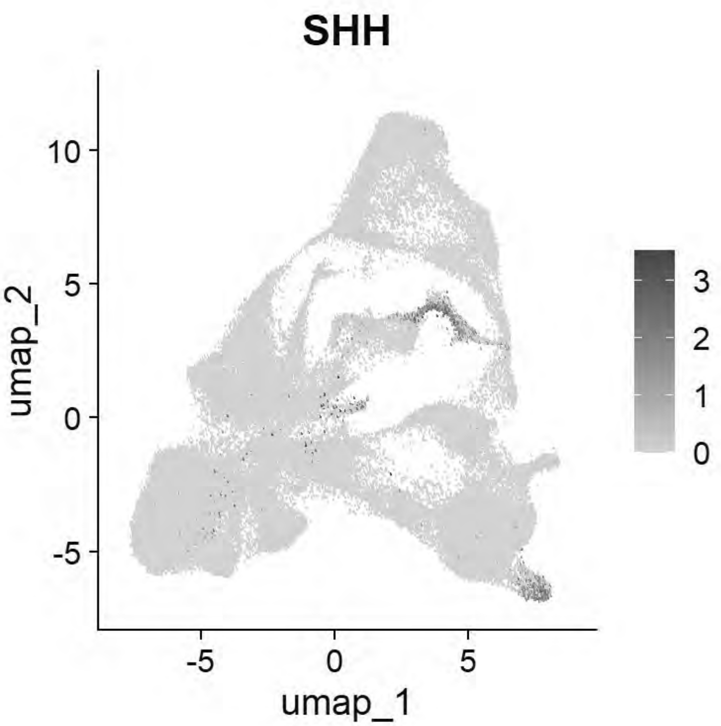


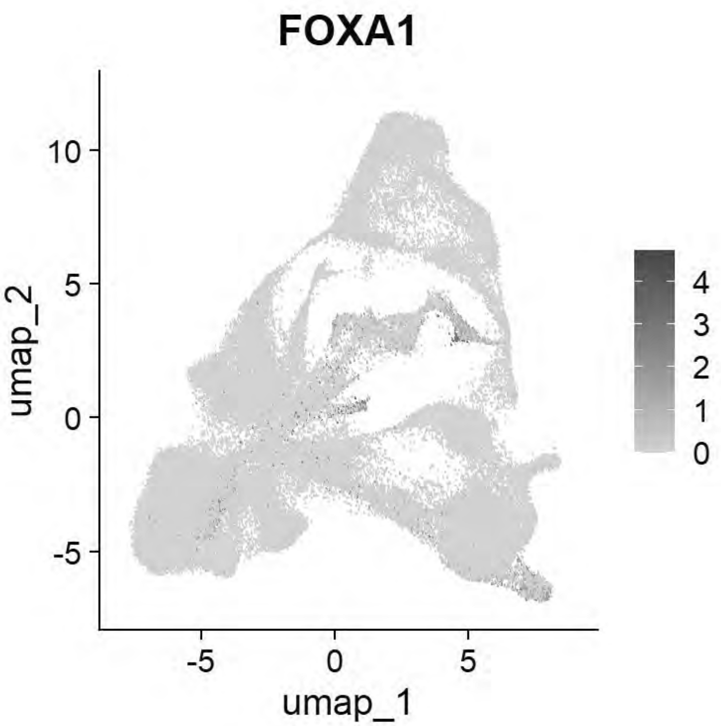


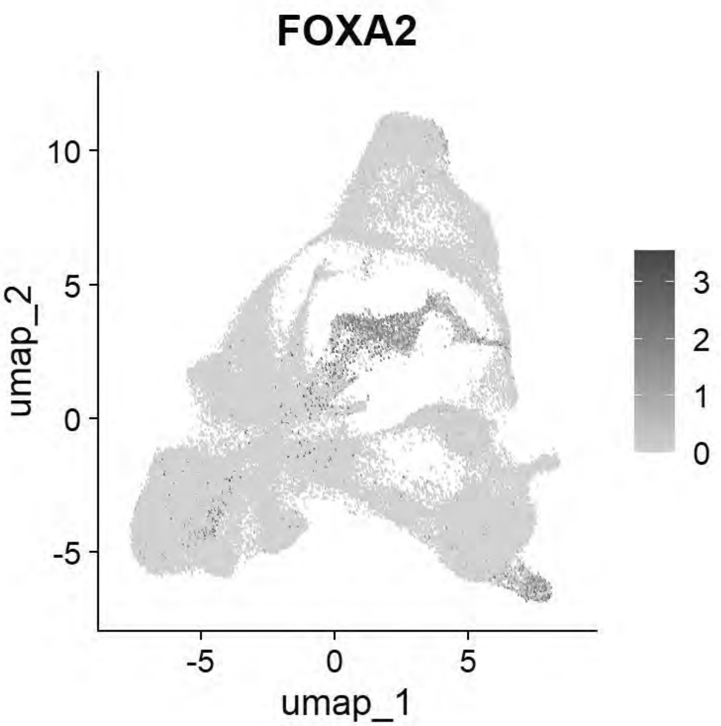


```
# Pajanoja: Neural Tube
FeaturePlot(chick, features=c("CDH2","HES5","MYCN","PAX2","NESTIN"))
```

```
## Warning: The following requested variables were not found: NESTIN
```

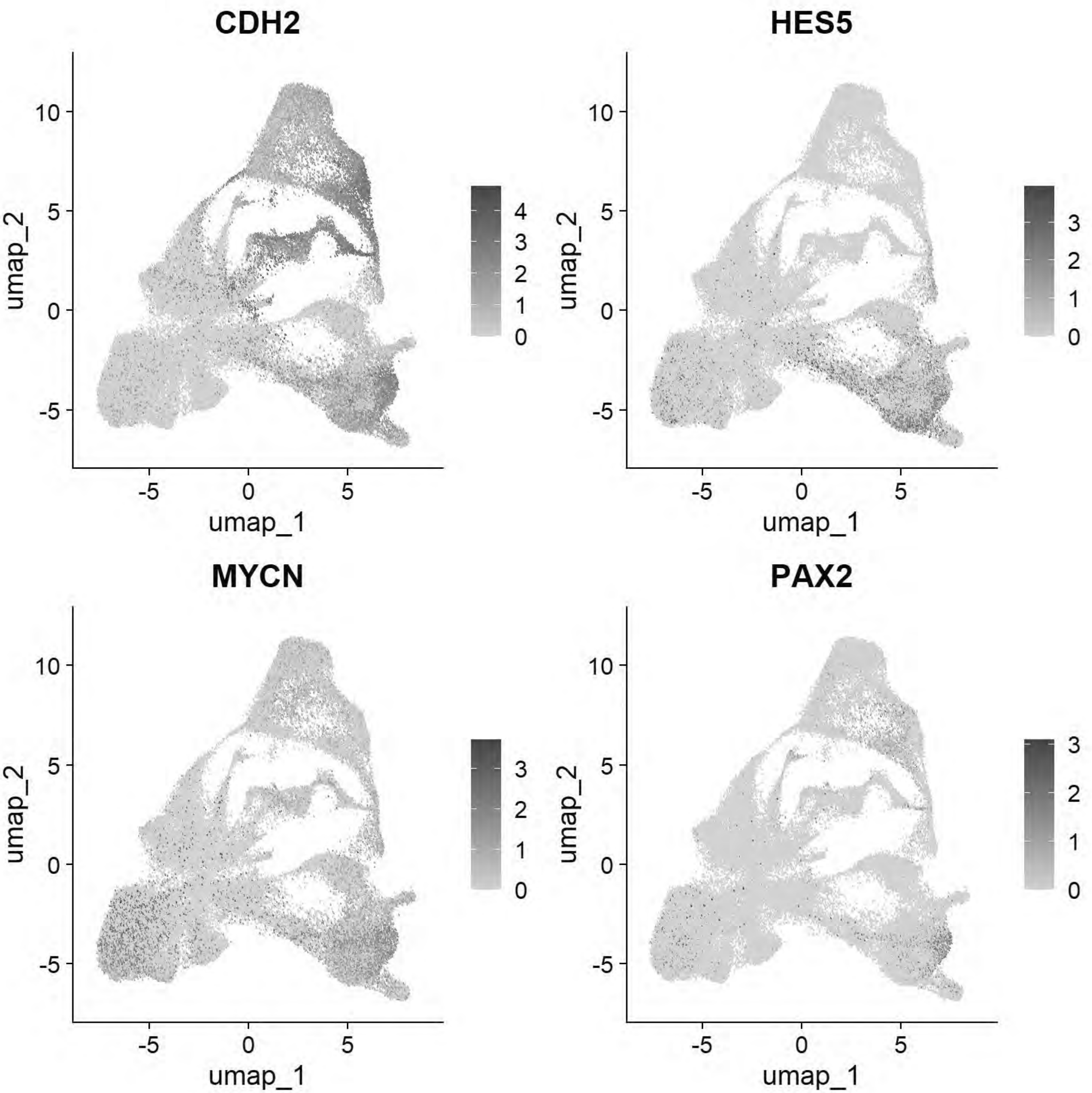


```
# Pajanoja: Non-Neural Ectoderm/Placode
FeaturePlot(chick, features=c("KRT24","KRT18","DLX5","SIX1","EYA2","TFAP2A"))
```

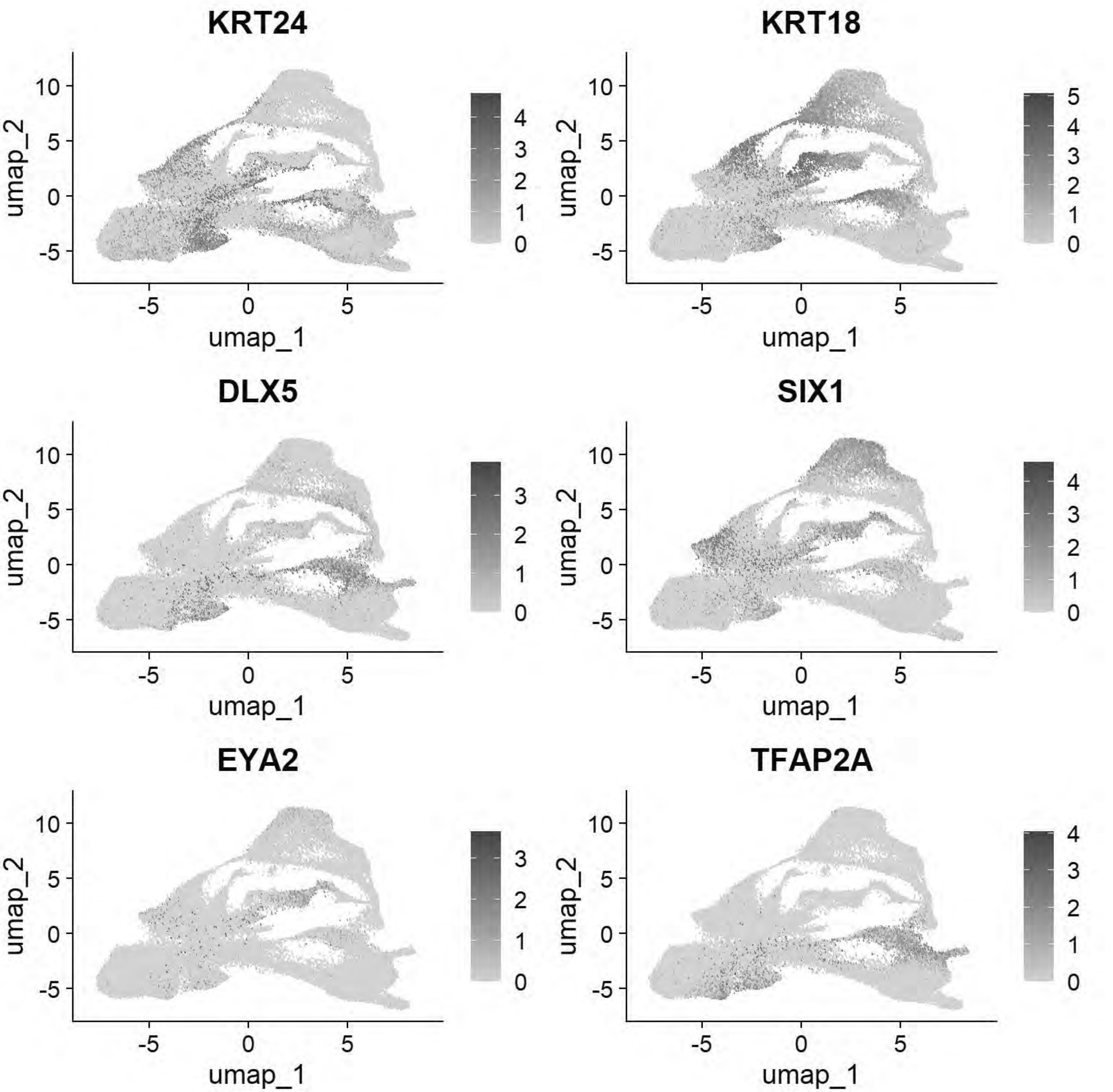


```
# Pajanoja: Neural crest
FeaturePlot(chick, features=c("FOXD3","TFAP2B","SOX9","ETS1","SNAI2"))
```

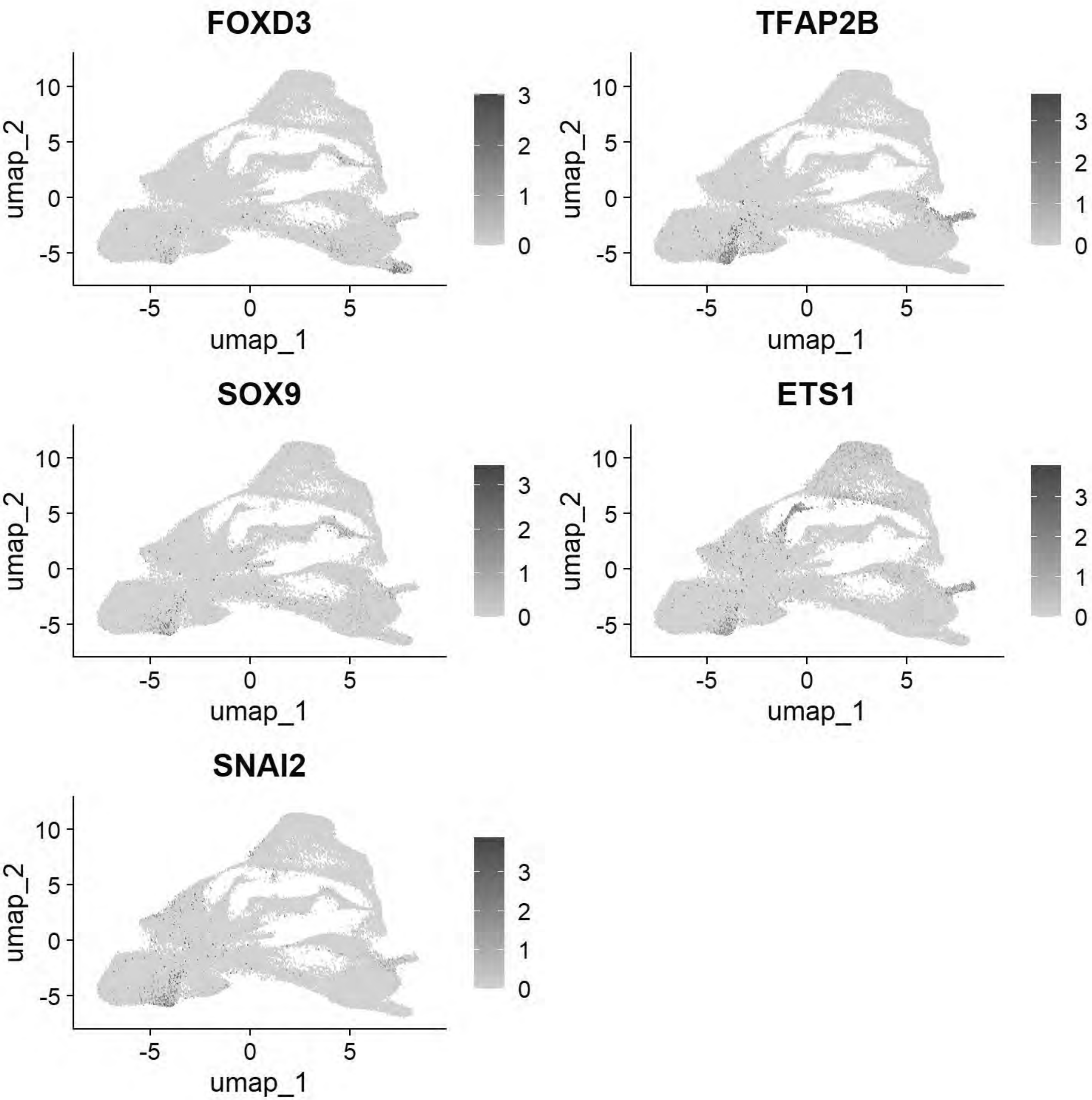


```
gc()
```

```
##              used   (Mb) gc trigger    (Mb)   max used    (Mb)
## Ncells   4802245  256.5   13651909   729.1   21331105  1139.3
## Vcells 730278216 5571.6 3177071757 24239.2 4488399353 34243.8
```

```
chick <- JoinLayers(chick)
chick.markers <- FindAllMarkers(chick, min.pct = 0.5, logfc.threshold = 0.5)
```

```
## Calculating cluster 0
```

```
## Calculating cluster 1
```

```
## Calculating cluster 2
```

```
## Calculating cluster 3
```

```
## Calculating cluster 4
```

```
## Calculating cluster 5
```

```
## Calculating cluster 6
```

```
## Calculating cluster 7
```

```
## Calculating cluster 8
```

```
## Calculating cluster 9
```

```
## Calculating cluster 10
```

```
## Calculating cluster 11
```

```
## Calculating cluster 12
```

```
## Calculating cluster 13
```

```
## Calculating cluster 14
```

```
## Calculating cluster 15
```

```
## Calculating cluster 16
```

```
## Calculating cluster 17
```

```
## Calculating cluster 18
```

```
## Calculating cluster 19
```

```
## Calculating cluster 20
```

```
## Calculating cluster 21
```

```
## Calculating cluster 22
```

```
## Calculating cluster 23
```

```
## Calculating cluster 24
```

```
## Calculating cluster 25
```

```
gc()
```

```
##              used   (Mb) gc trigger    (Mb)   max used    (Mb)
## Ncells    4803128  256.6   13651909   729.1   21331105  1139.3
## Vcells 1299709470 9916.0 3177071757 24239.2 4488399353 34243.8
```

```
chick.markers %>%
  group_by(cluster) %>%
  top_n(n = 5, wt = avg_log2FC) -> top5
DotPlot(chick, features = unique(top5$gene))+coord_flip()
```

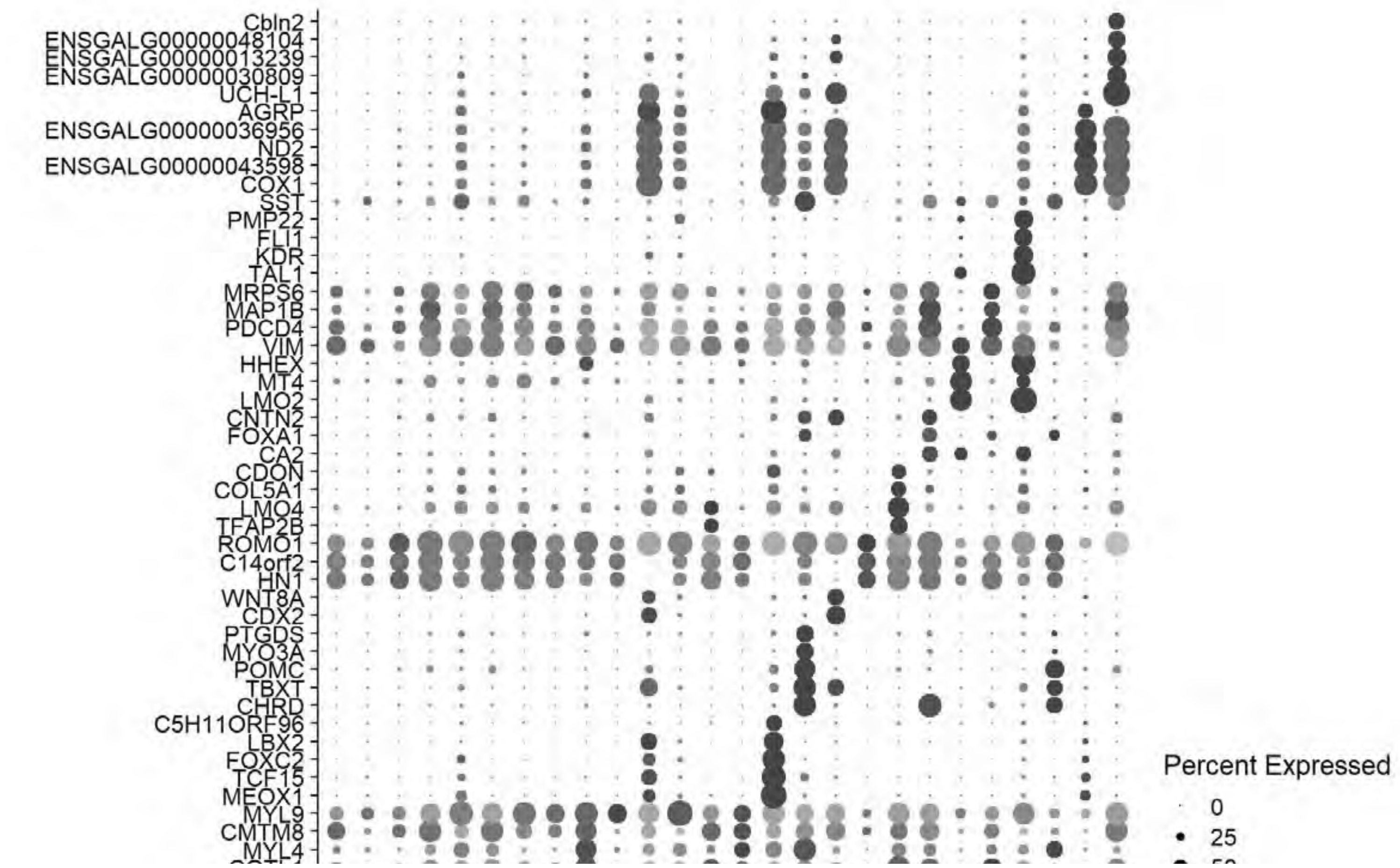

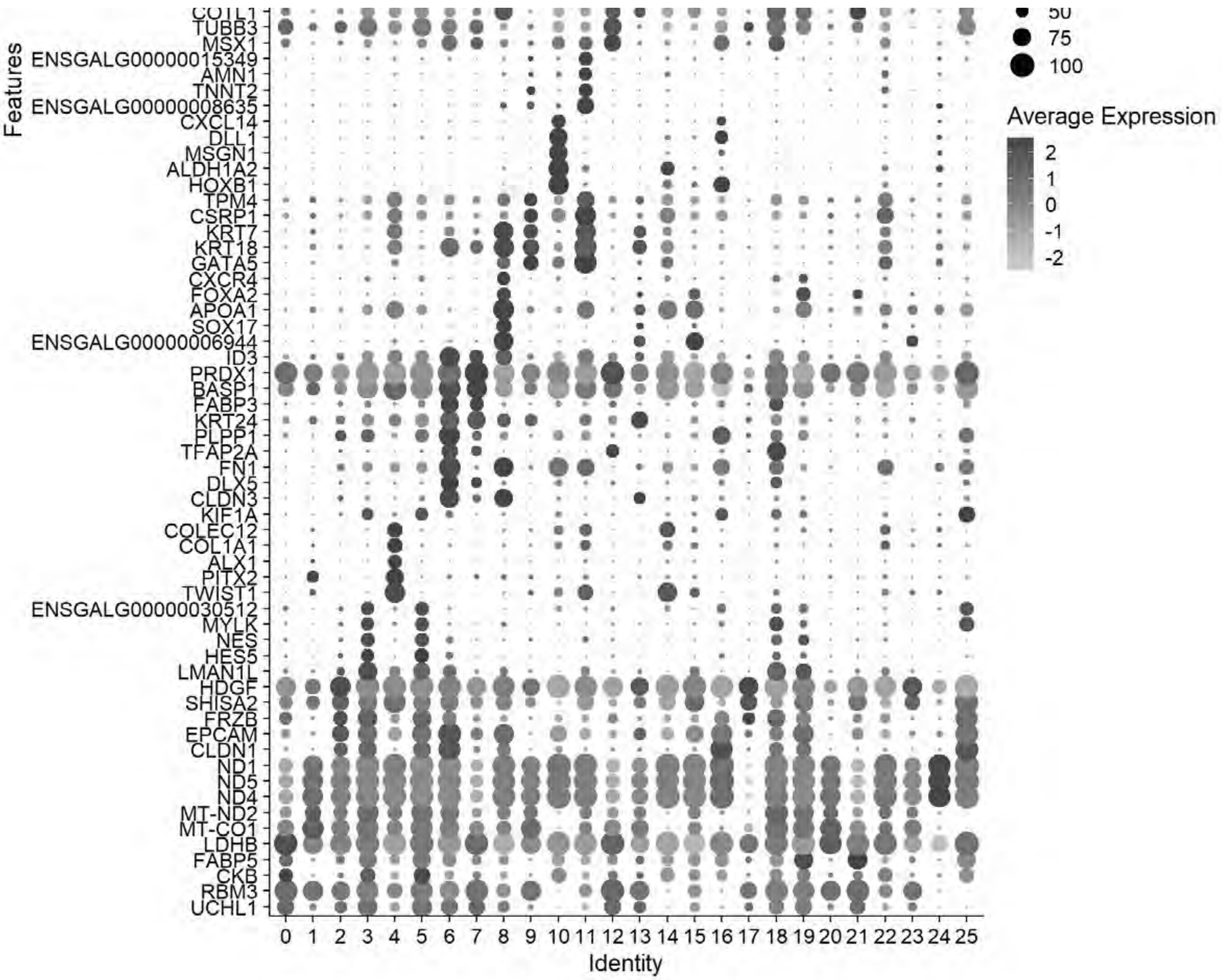


```
gc()
```

```
##              used   (Mb) gc trigger    (Mb)   max used    (Mb)
## Ncells    4764290  254.5   13651909   729.1   21331105  1139.3
## Vcells 1307955735 9979.0 3177071757 24239.2 4488399353 34243.8
```

```
#Annotation of Cell Types
cluster_map <- c("0"="Undecided Ectoderm 1",
                 "1"="Undecided Mesoderm 1",
                 "2"="Neural 1",
                 "3"="Def Neural 1",
                 "4"="Lateral Plate Mesoderm",
                 "5"="Def Neural 2",
                 "6"="NNE/Placode",
                 "7"="Undecided (NNE/Placode)",
                 "8"="Endoderm",
                 "9"="Undecided Mesoderm 2",
                 "10"="pLateral Plate Mesoderm",
```

```
                 "11"="Cardiac Mesoderm",
                 "12"="Neural Crest 1",
                 "13"="Mixed 1",
                 "14"="Paraxial Mesoderm",
                 "15"="Notochord",
                 "16"="Posterior mixed",
                 "17"="Neural 2",
                 "18"="Neural Crest 2",
                 "19"="vNeural Tube",
                 "20"="eHematogenic",
                 "21"="Undecided Ectoderm 2",
                 "22"="Hematogenic",
                 "23"="Mixed 2",
                 "24"="Mito-High",
                 "25"="Neural mixed" )
chick$celltype1 <- plyr::mapvalues(chick$seurat_clusters, from = names(cluster_map), to = clus
ter_map)

chick$celltype1 <- factor(chick$celltype1,
                          levels = c("Undecided Ectoderm 1","Undecided Ectoderm 2","Neural 1",
"Neural 2","Def Neural 1","Def Neural 2","Neural mixed",
                                     "Undecided (NNE/Placode)","NNE/Placode","Neural Crest 1",
"Neural Crest 2",
                                     "vNeural Tube","Posterior mixed","Notochord","Endoderm","
eHematogenic","Hematogenic","Mixed 1","Mixed 2","Mito-High",
                                     "Undecided Mesoderm 1","Undecided Mesoderm 2","Cardiac Me
soderm","Lateral Plate Mesoderm","Paraxial Mesoderm","pLateral Plate Mesoderm"))

#Find Markers & Export directly to CSV
chick <- JoinLayers(chick)
Idents(chick) <- "celltype1" # Setting active identity ensures markers are calculated by cell
type
global_markers <- FindAllMarkers(chick, min.pct = 0.5, logfc.threshold = 0.5)
```

```
## Calculating cluster Undecided Ectoderm 1
```

```
## Calculating cluster Undecided Ectoderm 2
```

```
## Calculating cluster Neural 1
```

```
## Calculating cluster Neural 2
```

```
## Calculating cluster Def Neural 1
```

```
## Calculating cluster Def Neural 2
```

```
## Calculating cluster Neural mixed
```

```
## Calculating cluster Undecided (NNE/Placode)
```

```
## Calculating cluster NNE/Placode
```

```
## Calculating cluster Neural Crest 1
```

```
## Calculating cluster Neural Crest 2
```

```
## Calculating cluster vNeural Tube
```

```
## Calculating cluster Posterior mixed
```

```
## Calculating cluster Notochord
```

```
## Calculating cluster Endoderm
```

```
## Calculating cluster eHematogenic
```

```
## Calculating cluster Hematogenic
```

```
## Calculating cluster Mixed 1
```

```
## Calculating cluster Mixed 2
```

```
## Calculating cluster Mito-High
```

```
## Calculating cluster Undecided Mesoderm 1
```

```
## Calculating cluster Undecided Mesoderm 2
```

```
## Calculating cluster Cardiac Mesoderm
```

```
## Calculating cluster Lateral Plate Mesoderm
```

```
## Calculating cluster Paraxial Mesoderm
```

```
## Calculating cluster pLateral Plate Mesoderm
```

```
write.csv(global_markers, "E:/Rogers Lab/Rogers NC EMT/Supplementary file 3_Global_Markers_Ann
otated.csv", row.names = FALSE)
```

```
# Generate and Save Summary Panel
# A) UMAP
cluster_colors <- c(
  "0"="#BFD3E6",
  "1"="#FDD0A2",
  "2"="#41B6C4",
  "3"="#2171B5",
  "4"="#CB181D",
  "5"="#1D91C0",
  "6"="#A1D99B",
  "7"="#C7E9C0",
  "8"="#FED976",
  "9"="#FC9272",
  "10"="#A50F15",
  "11"="#E31A1C",
  "12"="#8C96C6",
  "13"="#D9D9D9",
  "14"="#D94801",
  "15"="#DEEBF7",
  "16"="#C6DBEF",
  "17"="#7FCDBB",
  "18"="#8C6BB1",
  "19"="#08519C",
  "20"="#BD0026",
  "21"="#9EBCDA",
  "22"="#800026",
  "23"="#969696",
  "24"="#525252",
  "25"="#08306B"
)
celltype_colors <- c(
  "Undecided Ectoderm 1"="#BFD3E6",
  "Undecided Mesoderm 1"="#FDD0A2",
  "Neural 1"="#41B6C4",
  "Def Neural 1"="#2171B5",
  "Lateral Plate Mesoderm"="#CB181D",
  "Def Neural 2"="#1D91C0",
  "NNE/Placode"="#A1D99B",
  "Undecided (NNE/Placode)"="#C7E9C0",
  "Endoderm"="#FED976",
  "Undecided Mesoderm 2"="#FC9272",
  "pLateral Plate Mesoderm"="#A50F15",
  "Cardiac Mesoderm"="#E31A1C",
  "Neural Crest 1"="#8C96C6",
  "Mixed 1"="#D9D9D9",
  "Paraxial Mesoderm"="#D94801",
  "Notochord"="#DEEBF7",
  "Posterior mixed"="#C6DBEF",
  "Neural 2"="#7FCDBB",
  "Neural Crest 2"="#8C6BB1",
  "vNeural Tube"="#08519C",
  "eHematogenic"="#BD0026",
  "Undecided Ectoderm 2"="#9EBCDA",
  "Hematogenic"="#800026",
```

```
  "Mixed 2"="#969696",
  "Mito-High"="#525252",
  "Neural mixed"="#08306B")


p_umap <- DimPlot(chick, group.by = "celltype1",cols = celltype_colors, label = FALSE, repel =
 TRUE) + ggtitle(NULL)+
            NoLegend()
p_umap
```

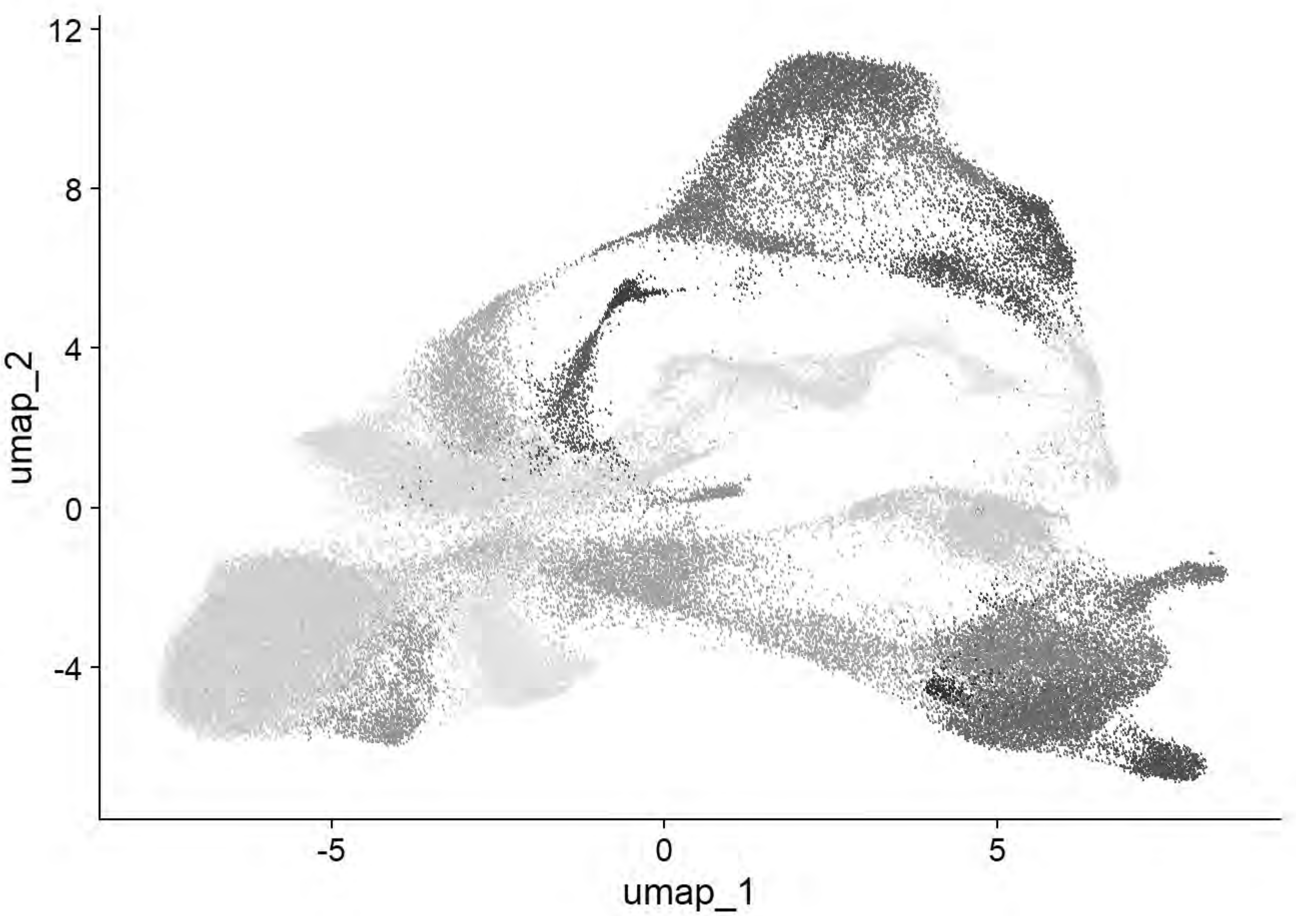


```
ggsave("E:/Rogers Lab/Rogers NC EMT/Figures/Fig_Global_Summary_UMAP1.png", plot = p_umap, widt
h = 6, height = 6, dpi = 600)

chick$stage <- factor(chick$stage,
                          levels = c("HH4","HH5","HH6","HH7","HH8","HH9","HH11"))
p_umap <- DimPlot(chick,cols = c("#C7E9B4","#7FCDBB","#41B6C4","#1D91C0","#225EA8","#253494","
#081D58"), group.by = "stage")+NoAxes()+ggtitle(NULL)
p_umap
```

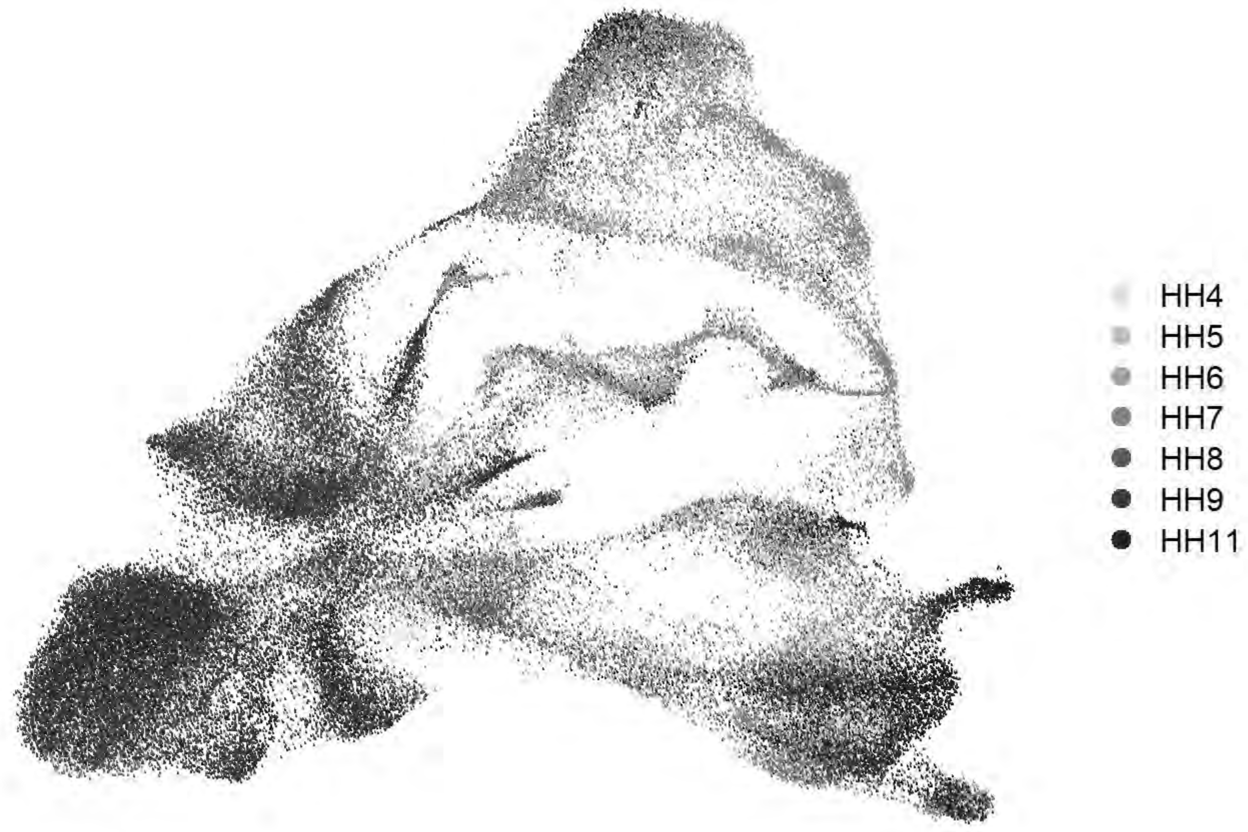


```
ggsave("E:/Rogers Lab/Rogers NC EMT/Figures/Fig_Global_Summary_UMAP2.png", plot = p_umap, widt
h = 7, height = 6, dpi = 600)

# B) Table
cptbl <- table(chick@meta.data$celltype1,chick@meta.data$stage)
cptbl <- cbind(cptbl, TOTAL = rowSums(cptbl))
colnames(cptbl)[length(cptbl[1,])] <- "TOTAL"

tg = gridExtra::tableGrob(cptbl)
h = grid::convertHeight(sum(tg$heights), "in", TRUE)
w = grid::convertWidth(sum(tg$widths), "in", TRUE)
ggsave("E:/Rogers Lab/Rogers NC EMT/Figures/Fig_Global_Summary_Table.png", tg, width=w, height
=h, device = 'png', dpi =600)

# C) DotPlot (Top 5 markers per cluster)
top5 <- global_markers %>% group_by(cluster) %>% top_n(n = 5, wt = avg_log2FC)
p_dot=DotPlot(chick, features = unique(top5$gene), group.by = "celltype1") + # cluster.idents
dendrograms the y-axis
  coord_flip() +
  scale_size_continuous(range = c(1, 10)) +
  scale_color_gradientn(colors = c("grey90", "#313695", "#d73027"))+
  theme_light() +
  theme(
    axis.text.x = element_text(angle = 45, hjust = 1, color = "black", size = 12),
    axis.text.y = element_text(face = "italic", color = "black", size = 12),
    axis.title.x = element_blank(),
```

```
    axis.title.y = element_blank(),
    panel.grid.major = element_line(color = "grey90")
  )
```

```
## Scale for size is already present.
## Adding another scale for size, which will replace the existing scale.
```

```
## Scale for colour is already present.
## Adding another scale for colour, which will replace the existing scale.
```

```
p_dot
```

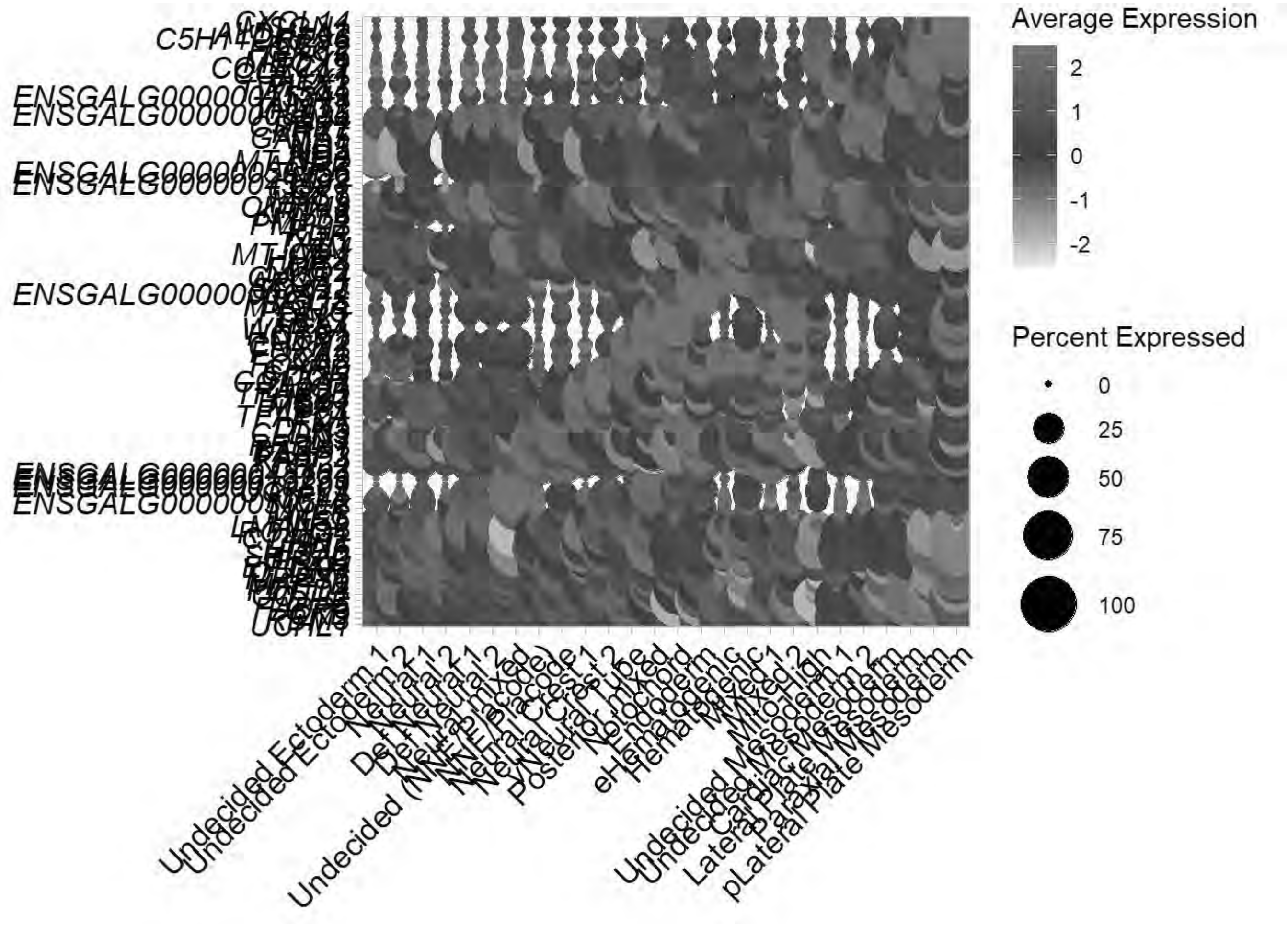


```
ggsave("E:/Rogers Lab/Rogers NC EMT/Figures/Fig_Global_Summary_Dotplot.png", plot = p_dot, wid
th = 13, height = 17, dpi = 600)

saveRDS(chick, "E:\\Rogers Lab\\Rogers NC EMT\\Global_chick_v2.rds")
```

### Global Tubulin Mapping

```
tub_subunit_map <- c("TUBA1A"="A","TUBA1B"="A","TUBA1C"="A","TUBA3E"="A","TUBA8B"="A","TUBAL3"
="A",
       "TUBB"="B","TUBB1"="B","TUBB2A"="B","TUBB2B"="B","TUBB3"="B","TUBB4B"="B","TUBB6"="B",
```

```
       "TUBG1"="G")
family_text_colors <- c(
  "A" = "#6A3D9A",
  "B" = "#008B8B",
  "G" = "#B15928"
)

# Sort
tub_in_data <- c("TUBA1A","TUBA1B","TUBA1C","TUBA3E","TUBA8B","TUBAL3",
       "TUBB","TUBB1","TUBB2A","TUBB2B","TUBB3","TUBB4B","TUBB6",
       "TUBG1")

sorted_tub_df <- data.frame(Gene = tub_in_data) %>%
  mutate(Family = tub_subunit_map[Gene]) %>%
  filter(!is.na(Family)) %>%
  mutate(Family_Factor = factor(Family, levels = c("A", "B", "G"))) %>%
  arrange(Family_Factor, Gene)

organized_tub <- sorted_tub_df$Gene
axis_text_colors <- family_text_colors[sorted_tub_df$Family]

# Plot
Idents(chick) <- "celltype1"
p <- DotPlot(chick, features = rev(organized_tub)) +
  coord_flip() +
  scale_color_gradientn(colors = c("grey90", "#313695", "#d73027")) +
  scale_size_continuous(range = c(1, 10)) +
  theme_light() +
  theme(
    axis.text.x = element_text(angle = 60, hjust = 1, color = "black", size = 12),
    axis.text.y = element_text(face = "italic", color = rev(axis_text_colors), size = 12),
    axis.title.x = element_blank(),
    axis.title.y = element_blank(),
    panel.grid.major = element_line(color = "grey90")
  )
```

```
## Scale for colour is already present.
## Adding another scale for colour, which will replace the existing scale.
## Scale for size is already present.
## Adding another scale for size, which will replace the existing scale.
```

```
p
```

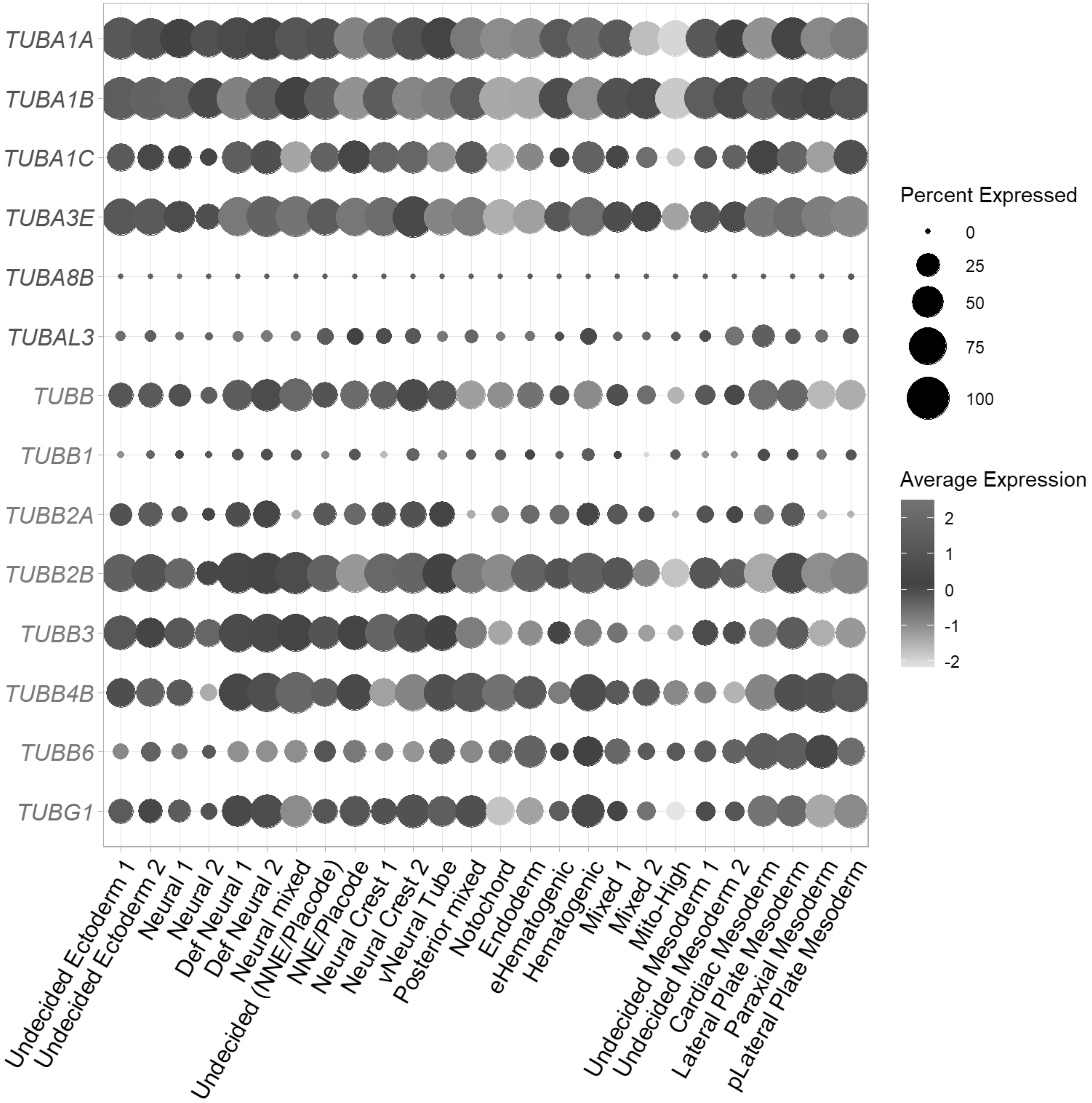


```
ggsave(filename = paste0("E:/Rogers Lab/Rogers NC EMT/Figures/Fig_Global_tub_categorization.pn
g"), plot = p, device = 'png', width = 13, height = 7)

f1=FeaturePlot(chick, features=c("TUBA1A"))+NoAxes()+scale_color_gradientn(colors = c("grey90"
, "#2EACBD", "#08306B"), limit=c(0,6))
```

```
## Scale for colour is already present.
## Adding another scale for colour, which will replace the existing scale.
```

```
f2=FeaturePlot(chick, features=c("TUBA1B"))+NoAxes()+scale_color_gradientn(colors=c("grey90",
"#2EACBD", "#08306B"), limit=c(0,6))
```

```
## Scale for colour is already present.
## Adding another scale for colour, which will replace the existing scale.
```

```
f3=FeaturePlot(chick, features=c("TUBB3"))+NoAxes()+scale_color_gradientn(colors=c("grey90", "
#2EACBD", "#08306B"), limit=c(0,6))
```

```
## Scale for colour is already present.
## Adding another scale for colour, which will replace the existing scale.
```

```
f4=FeaturePlot(chick, features=c("TUBB2A"))+NoAxes()+scale_color_gradientn(colors=c("grey90",
"#2EACBD", "#08306B"), limit=c(0,6))
```

```
## Scale for colour is already present.
## Adding another scale for colour, which will replace the existing scale.
```

```
f5=FeaturePlot(chick, features=c("TUBB2B"))+NoAxes()+scale_color_gradientn(colors=c("grey90",
"#2EACBD", "#08306B"), limit=c(0,6))
```

```
## Scale for colour is already present.
## Adding another scale for colour, which will replace the existing scale.
```

```
f6=FeaturePlot(chick, features=c("TUBB4B"))+NoAxes()+scale_color_gradientn(colors=c("grey90",
"#2EACBD", "#08306B"), limit=c(0,6))
```

```
## Scale for colour is already present.
## Adding another scale for colour, which will replace the existing scale.
```

```
f= ggarrange(plotlist = list(f1,f2,f3,f4,f5,f6), ncol = 3, nrow = 2)
f
```

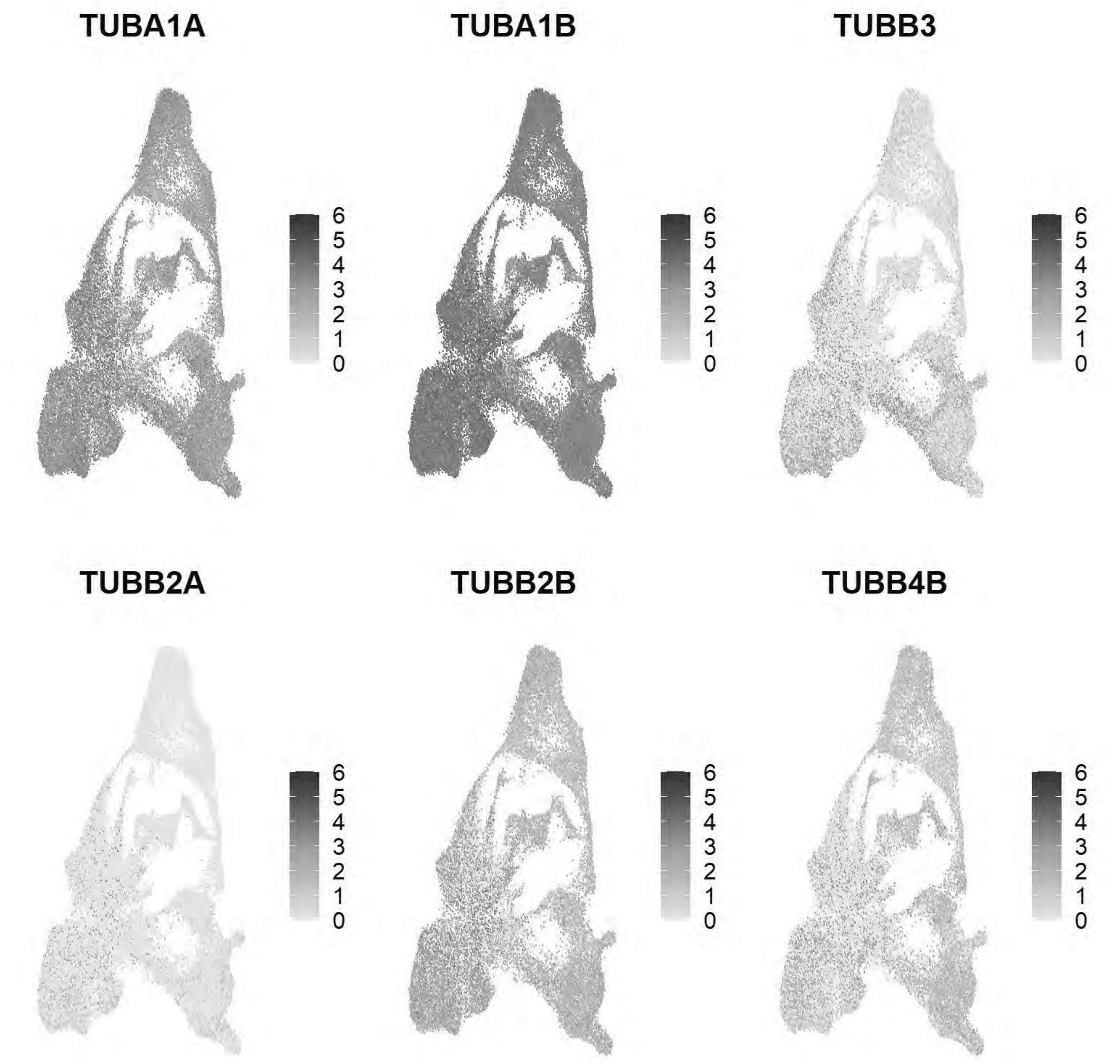


```
ggsave(f, filename = "E:/Rogers Lab/Rogers NC EMT/Figures/Fig_Global_selecttubs_FeaturePlot1.p
ng", width = 10.5, height = 6, device='png', dpi=600)

f1=FeaturePlot(chick, features=c("TUBAL3"))+NoAxes()+scale_color_gradientn(colors = c("grey90"
, "#2EACBD", "#08306B"), limit=c(0,6))
```

```
## Scale for colour is already present.
## Adding another scale for colour, which will replace the existing scale.
```

```
f2=FeaturePlot(chick, features=c("TUBB6"))+NoAxes()+scale_color_gradientn(colors=c("grey90", "
#2EACBD", "#08306B"), limit=c(0,6))
```

```
## Scale for colour is already present.
## Adding another scale for colour, which will replace the existing scale.
```

```
f3=FeaturePlot(chick, features=c("TUBG1"))+NoAxes()+scale_color_gradientn(colors=c("grey90", "
#2EACBD", "#08306B"), limit=c(0,6))
```

```
## Scale for colour is already present.
## Adding another scale for colour, which will replace the existing scale.
```

```
f= ggarrange(plotlist = list(f1,f2,f3), ncol = 1, nrow = 3)
f
```

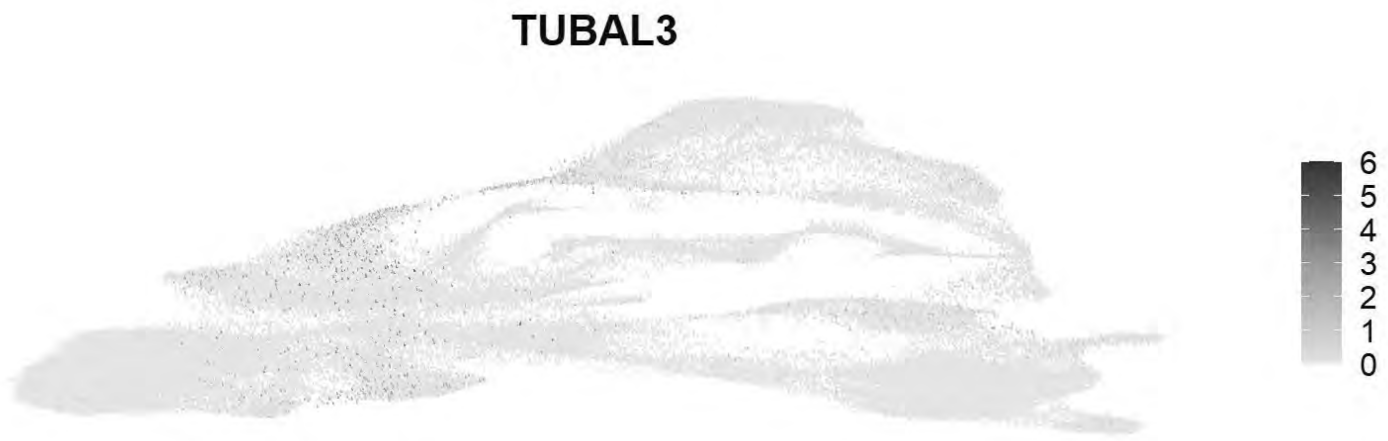


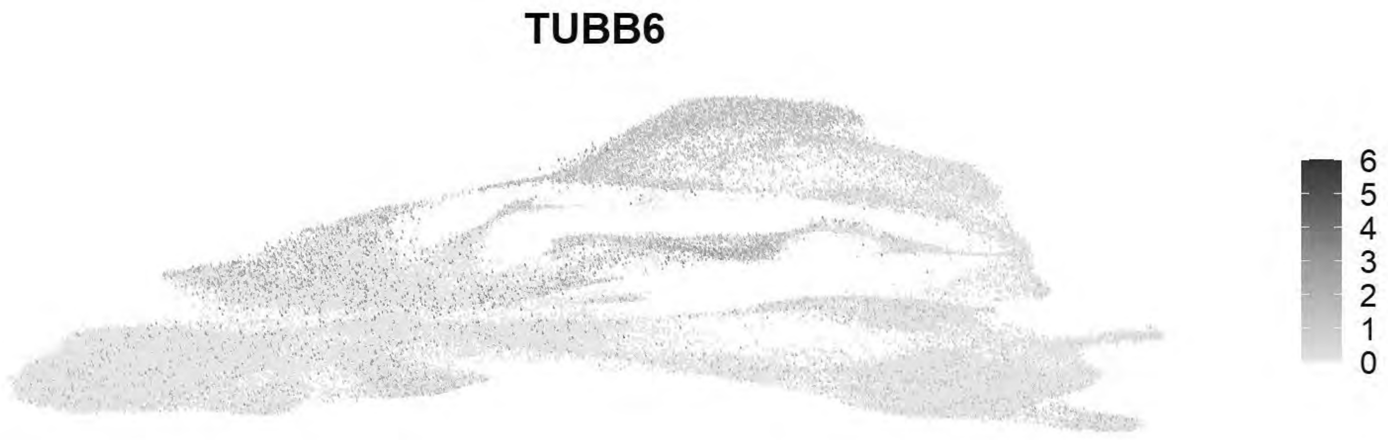


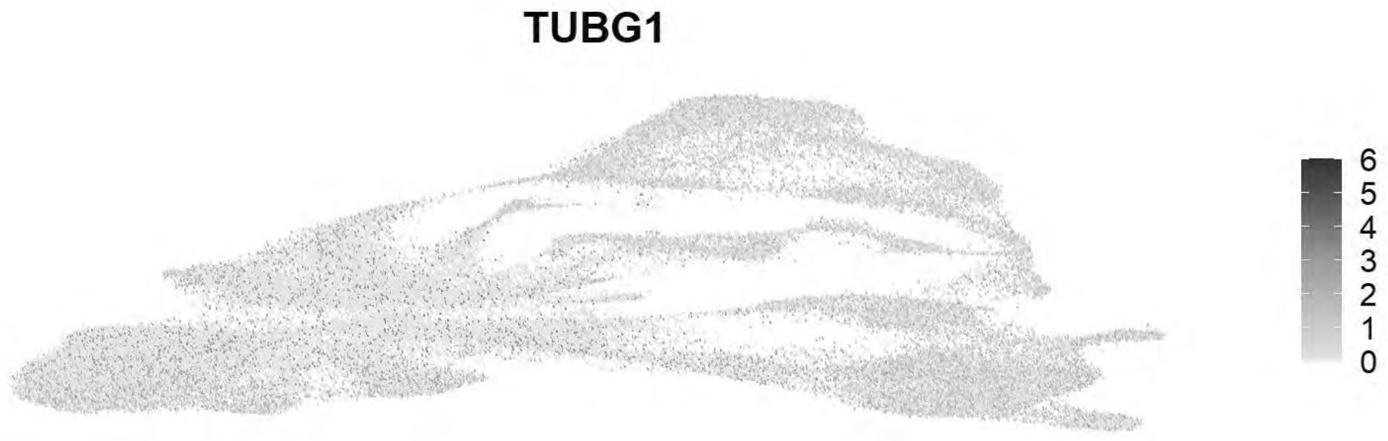


```
ggsave(f, filename = "E:/Rogers Lab/Rogers NC EMT/Figures/Fig_Global_selecttubs_FeaturePlot2.p
ng", width = 3.5, height = 9, device='png', dpi=600)

gc()
```

```
##              used    (Mb) gc trigger    (Mb)   max used    (Mb)
## Ncells    5152931   275.2   13651909   729.1   21331105  1139.3
## Vcells 1321437610 10081.8 3660127464 27924.6 4574774951 34902.8
```

```
chick <- AddModuleScore(chick, features = list(c("TUBA1A","TUBA1B","TUBA1C","TUBA3E","TUBA8B",
"TUBAL3",
       "TUBB","TUBB1","TUBB2A","TUBB2B","TUBB3","TUBB4B","TUBB6",
       "TUBG1")), name = "Global_TUB_Score")
chick <- AddModuleScore(chick, features = list(c("TUBA1A","TUBA1B","TUBA1C","TUBA3E","TUBA8B",
"TUBAL3")), name = "Global_TUBB_Score")
chick <- AddModuleScore(chick, features = list(c("TUBB","TUBB1","TUBB2A","TUBB2B","TUBB3","TUB
B4B","TUBB6")), name = "Global_TUBA_Score")
f1=FeaturePlot(chick, features=c("Global_TUB_Score1"))+NoAxes()+scale_color_gradientn(colors =
 c("grey90", "#313695", "#d73027"), limit=c(0,2.5))
```

```
## Scale for colour is already present.
## Adding another scale for colour, which will replace the existing scale.
```

```
f2=FeaturePlot(chick, features=c("Global_TUBA_Score1"))+NoAxes()+scale_color_gradientn(colors=
c("grey90", "#313695", "#d73027"), limit=c(0,2.5))
```

```
## Scale for colour is already present.
## Adding another scale for colour, which will replace the existing scale.
```

```
f3=FeaturePlot(chick, features=c("Global_TUBB_Score1"))+NoAxes()+scale_color_gradientn(colors=
c("grey90", "#313695", "#d73027"), limit=c(0,2.5))
```

```
## Scale for colour is already present.
## Adding another scale for colour, which will replace the existing scale.
```

```
f= ggarrange(plotlist = list(f1,f2,f3), ncol = 3)
f
```

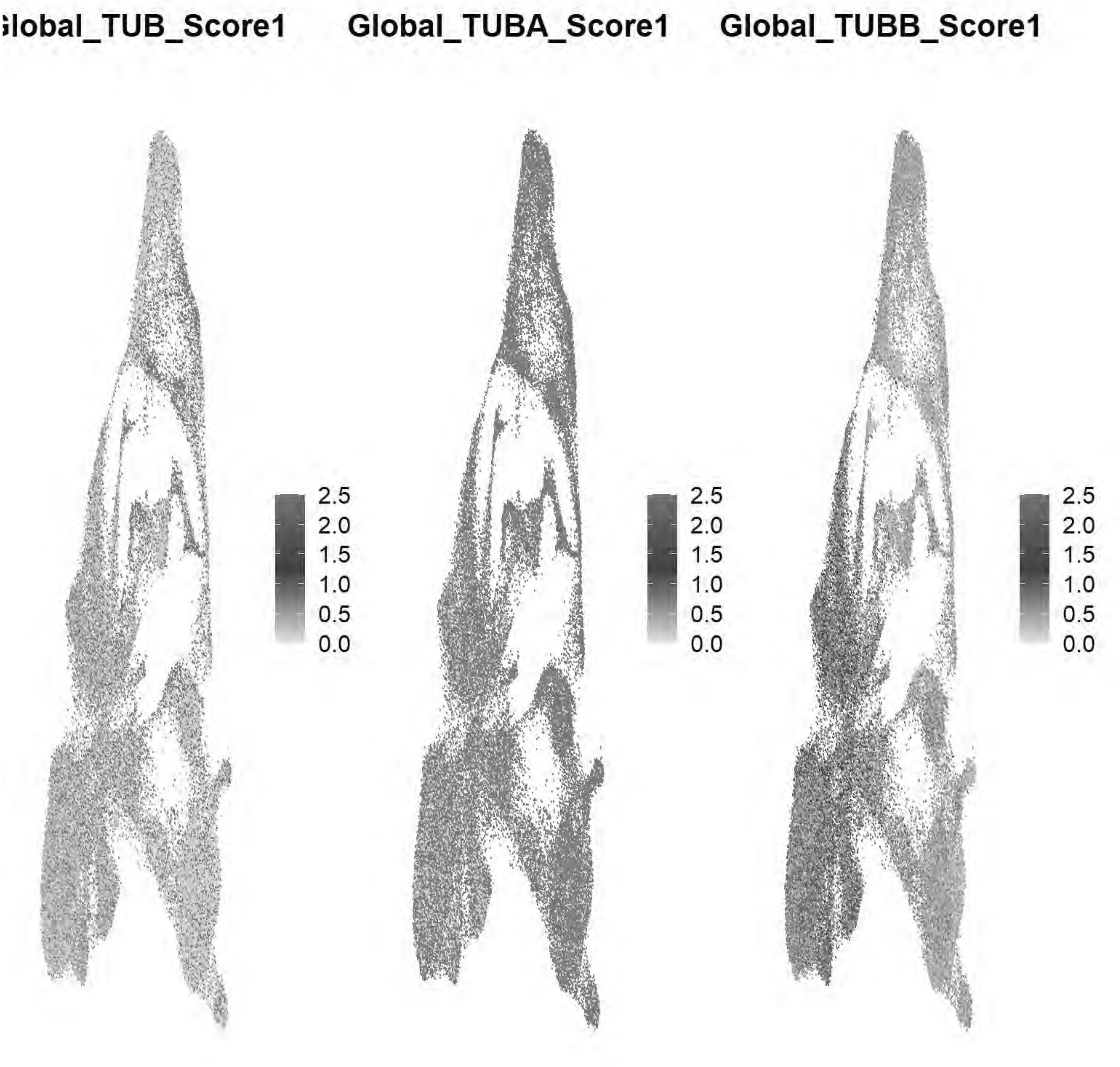


```
ggsave(f, filename = "E:/Rogers Lab/Rogers NC EMT/Figures/Fig_Global_tubscore_FeaturePlot.png"
, width = 10.5, height = 3, device='png', dpi=600)

f1=VlnPlot(chick, features=c("Global_TUB_Score1"), pt.size=0,sort = "increasing", cols = cellt
ype_colors)+NoLegend()+ggtitle(NULL)+
  theme(axis.text.x = element_text(angle = 60, hjust = 1, color = "black", size = 12),axis.tit
le.x = element_blank(),axis.title.y = element_blank())
f2=VlnPlot(chick, features=c("Global_TUBA_Score1"),pt.size=0,sort = "increasing", cols = cellt
ype_colors)+NoLegend()+ggtitle(NULL)+
  theme(axis.text.x = element_text(angle = 60, hjust = 1, color = "black", size = 12),axis.tit
```

```
le.x = element_blank(),axis.title.y = element_blank())
f3=VlnPlot(chick, features=c("Global_TUBB_Score1"),pt.size = 0,sort = "increasing", cols = cel
ltype_colors)+NoLegend()+ggtitle(NULL)+
  theme(axis.text.x = element_text(angle = 60, hjust = 1, color = "black", size = 12),axis.tit
le.x = element_blank(),axis.title.y = element_blank())
f= ggarrange(plotlist = list(f1,f2,f3), ncol = 1)
f
```

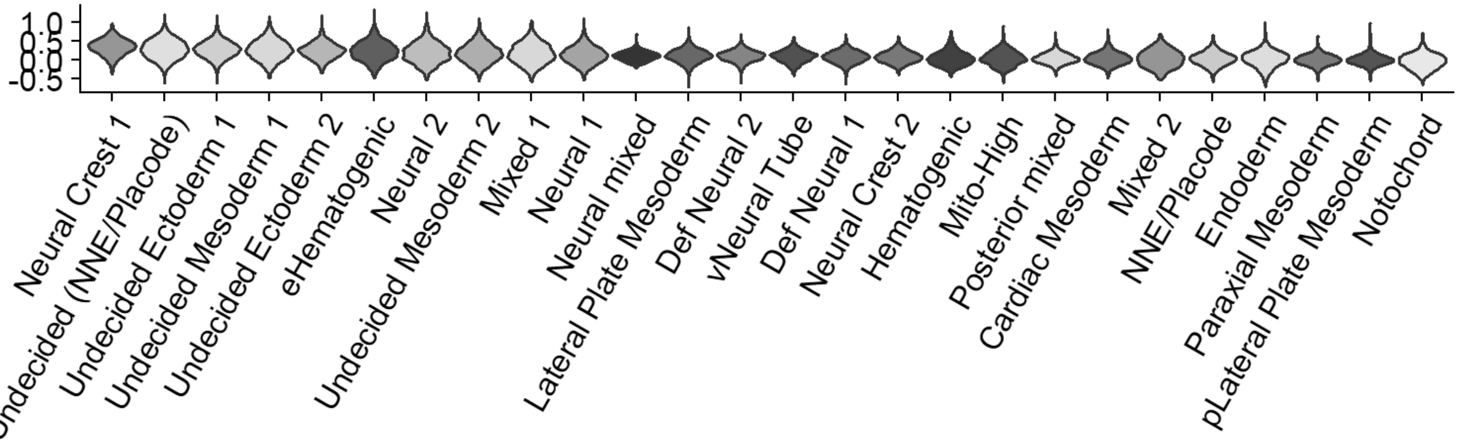


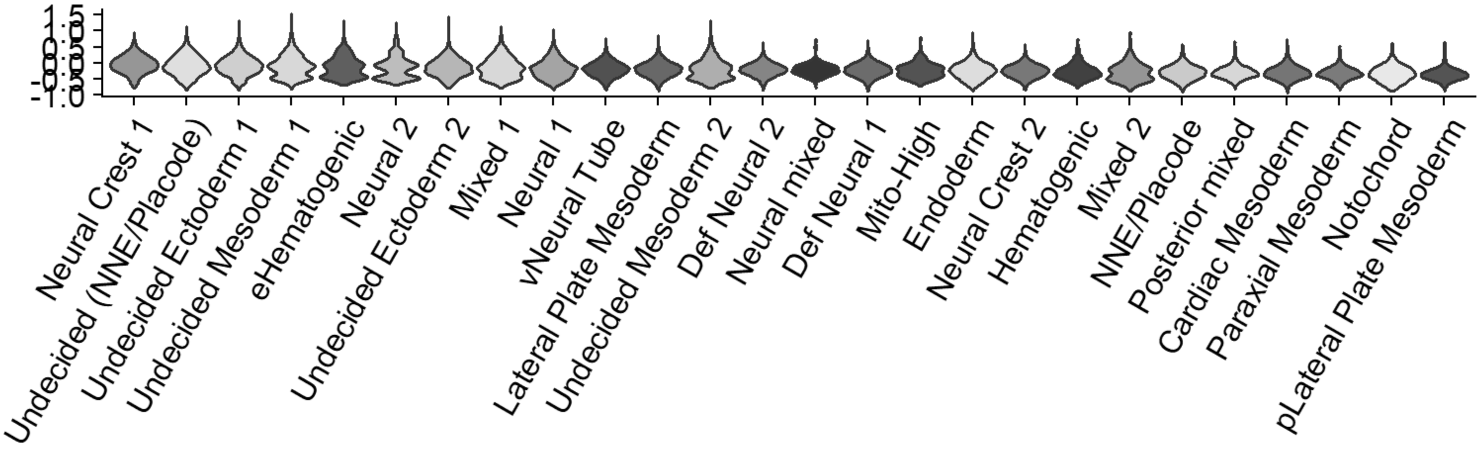


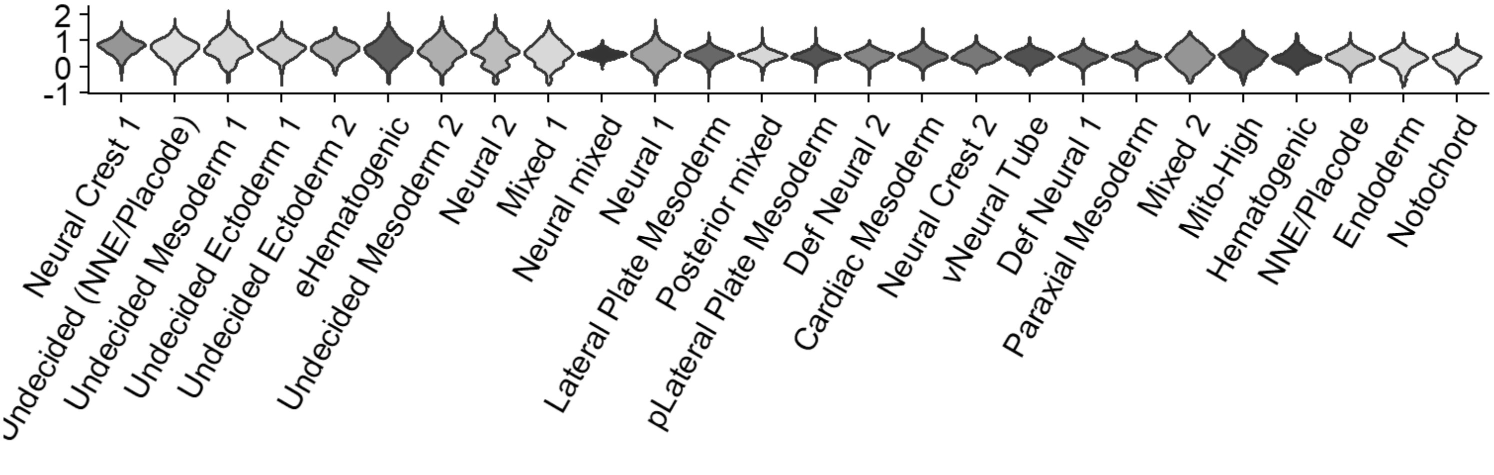


```
ggsave(f, filename = "E:/Rogers Lab/Rogers NC EMT/Figures/Fig_Global_tubscore_VlnPlot.png", wi
dth = 8.5, height = 10, device='png', dpi=600)
```

### Global Dynein Mapping

```
dyn_subunit_map <- c(
  "DYNC1H1"="C1", "DYNC1I1"="C1","DYNC1I2"="C1","DYNC1LI1"="C1","DYNC1LI2"="C1",
  "DYNLL1"="C1+C2","DYNLL2"="C1+C2","DYNLRB1"="C1+C2","DYNLRB2"="C1+C2","DYNLT1"="C1+C2","DYNL
T3"="C1+C2",
  "DYNLT2"="C2","DYNC2H1"="C2","DYNC2I1"="C2","DYNC2I2"="C2","DYNC2LI1"="C2")

# 2. A vibrant, high-saturation categorical palette
family_text_colors <- c(
  "C1"        = "#008B8B",
  "C1+C2" = "#B15928",
  "C2"        = "#6A3D9A"
)

# 3. Sort your dataset's dyn
dyn_in_data <- rownames(chick)[grep("^DYN", toupper(rownames(chick)))]

sorted_dyn_df <- data.frame(Gene = dyn_in_data) %>%
  mutate(Family = dyn_subunit_map[Gene]) %>%
  filter(!is.na(Family)) %>%
  mutate(Family_Factor = factor(Family, levels = c("C1", "C1+C2", "C2"))) %>%
  arrange(Family_Factor, Gene)

organized_dyn <- sorted_dyn_df$Gene
axis_text_colors <- family_text_colors[sorted_dyn_df$Family]

# 4. Plot
Idents(chick) <- "celltype1"

p <- DotPlot(chick, features = rev(organized_dyn)) +
  coord_flip() +

  # NEW GRADIENT: Cool to Warm (Light grey -> Deep Navy -> Crimson)
  # This provides massive contrast against both the white background and the colored text
  scale_color_gradientn(colors = c("grey90", "#313695", "#d73027")) +

  scale_size_continuous(range = c(1, 10)) +
  theme_light() +
  theme(
    axis.text.x = element_text(angle = 60, hjust = 1, color = "black", size = 12),
    axis.text.y = element_text(face = "italic", color = rev(axis_text_colors), size = 12),
    axis.title.x = element_blank(),
    axis.title.y = element_blank(),
    panel.grid.major = element_line(color = "grey90")
  )
```

```
## Scale for colour is already present.
## Adding another scale for colour, which will replace the existing scale.
## Scale for size is already present.
## Adding another scale for size, which will replace the existing scale.
```

```
p
```

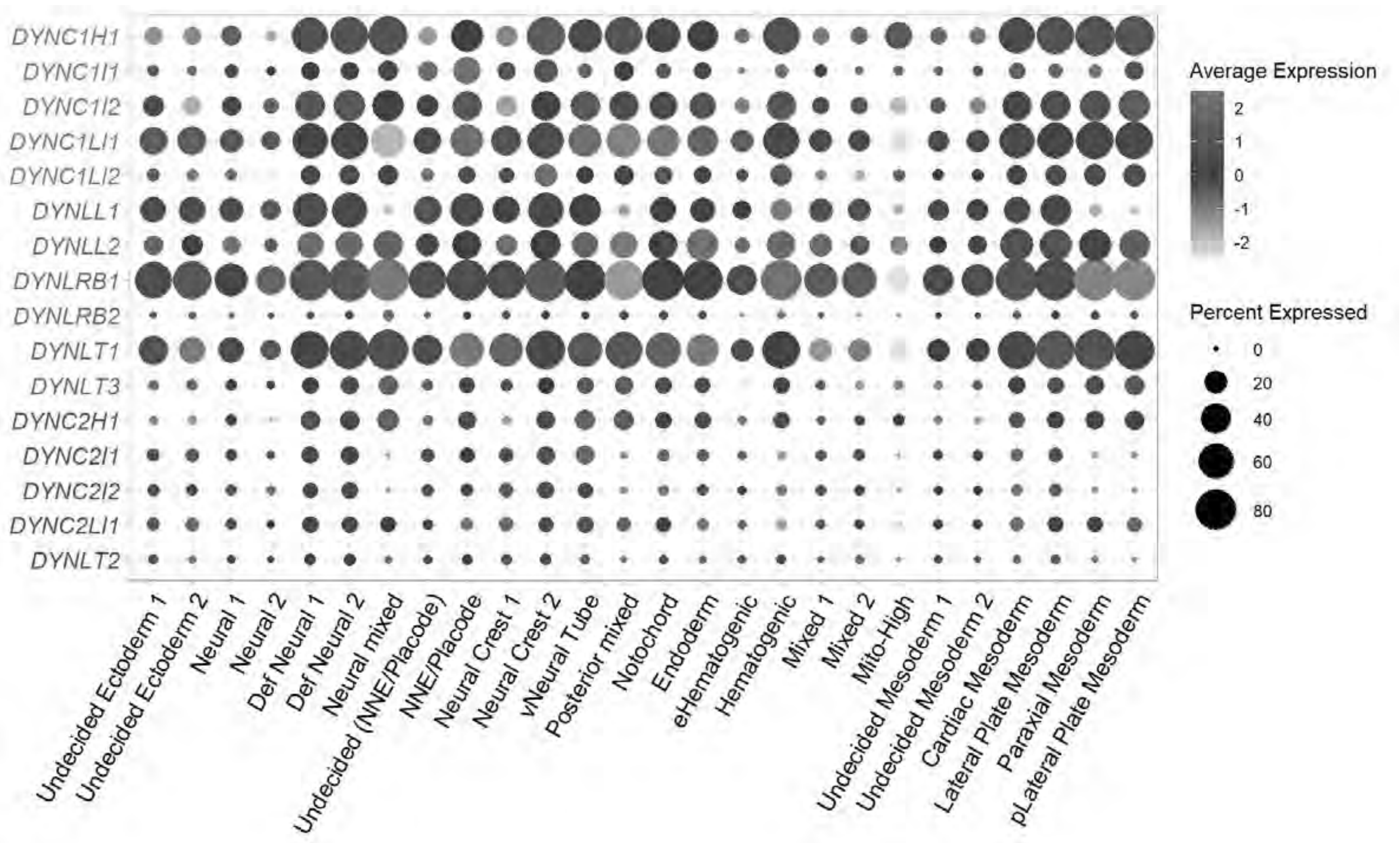


```
ggsave(filename = paste0("E:/Rogers Lab/Rogers NC EMT/Figures/Fig_Global_dyn_dotplot.png"), pl
ot = p, device = 'png', width = 10, height = 6)
```

### Global Kinesin Mapping

```
#Gene to superfamily mapping
gene_superfamily_map <- c(
  "KIF2A"="M", "KIF2B"="M", "KIF2C"="M",
  "KIFC1"="C", "KIFC2"="C", "KIFC3"="C",
  "KIF5A"="N-1", "KIF5B"="N-1", "KIF5C"="N-1",
  "KIF6"="N-2", "KIF7"="N-2", "KIF8"="N-2", "KIF9"="N-2", "KIF11"="N-2",
  "KIF1A"="N-3", "KIF1B"="N-3", "KIF1C"="N-3", "KIF13A"="N-3", "KIF13B"="N-3", "KIF14"="N-3",
"KIF16A"="N-3", "KIF16B"="N-3",
  "KIF3A"="N-4", "KIF3B"="N-4", "KIF3C"="N-4", "KIF17"="N-4",
  "KIF4A"="N-5", "KIF4B"="N-5", "KIF21A"="N-5", "KIF21B"="N-5",
  "KIF20A"="N-6", "KIF20B"="N-6", "KIF23"="N-6",
  "KIF10"="N-7",
  "KIF18A"="N-8", "KIF18B"="N-8", "KIF19A"="N-8", "KIF19B"="N-8", "KIF22"="N-8",
  "KIF12"="N-9",
  "KIF15"="N-10",
  "KIF24"="N-11", "KIF25"="N-11", "KIF26A"="N-11", "KIF26B"="N-11"
)
```

```
#
family_text_colors <- c(
  "C"    = "#1A1A1A", # Crisp Black
  "M"    = "#E31A1C", # Bright Red
  "N-1"  = "#33A02C", # Strong Green
  "N-2"  = "#1F78B4", # Strong Blue
  "N-3"  = "#FF7F00", # Vibrant Orange
  "N-4"  = "#6A3D9A", # Vibrant Purple
  "N-5"  = "#008B8B", # Dark Cyan
  "N-6"  = "#B15928", # Burnt Orange
  "N-7"  = "#E7298A", # Hot Pink
  "N-8"  = "#00BA38", # Bright Lime/Green
  "N-9"  = "#619CFF", # Bright Sky Blue
  "N-10" = "#B8860B", # Goldenrod
  "N-11" = "#8B4513"  # Saddle Brown
)

# Sort
kifs_in_data <- rownames(chick)[grep("^KIF", toupper(rownames(chick)))]

sorted_kif_df <- data.frame(Gene = kifs_in_data) %>%
  mutate(Family = gene_superfamily_map[Gene]) %>%
  filter(!is.na(Family)) %>%
  mutate(Family_Factor = factor(Family, levels = c("C", "M", "N-1", "N-2", "N-3", "N-4", "N-5"
, "N-6", "N-7", "N-8", "N-9", "N-10", "N-11"))) %>%
  arrange(Family_Factor, Gene)

organized_kifs <- sorted_kif_df$Gene
axis_text_colors <- family_text_colors[sorted_kif_df$Family]

# Plot
Idents(chick) <- "celltype1"

p <- DotPlot(chick, features = rev(organized_kifs)) +
  coord_flip() +
  scale_color_gradientn(colors = c("grey90", "#313695", "#d73027")) +

  scale_size_continuous(range = c(1, 10)) +
  theme_light() +
  theme(
    axis.text.x = element_text(angle = 60, hjust = 1, color = "black", size = 12),
    axis.text.y = element_text(face = "italic", color = rev(axis_text_colors), size = 12),
    axis.title.x = element_blank(),
    axis.title.y = element_blank(),
    panel.grid.major = element_line(color = "grey90")
  )
```

```
## Scale for colour is already present.
## Adding another scale for colour, which will replace the existing scale.
## Scale for size is already present.
## Adding another scale for size, which will replace the existing scale.
```

```
p
```

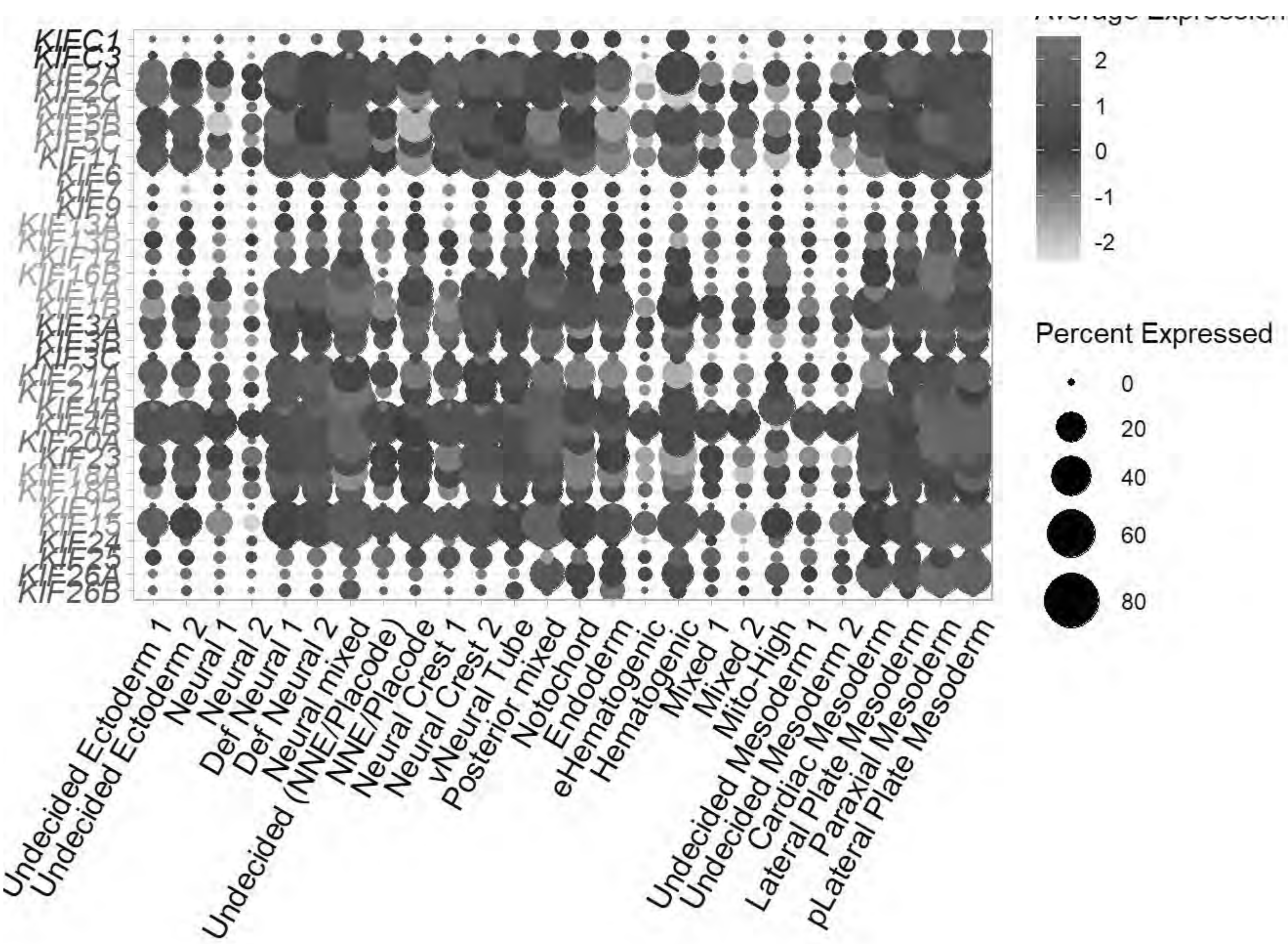


```
ggsave(filename = paste0("E:/Rogers Lab/Rogers NC EMT/Figures/Fig_Global_kif_dotplot.png"), pl
ot = p, device = 'png', width = 10, height = 10)
```

### 4. Extracting the Ectoderm & Stage Subsets relevant to NC EMT

### Stage Analysis: HH8

```
#chick=readRDS("E:\\Rogers Lab\\NC EMT\\Global_chick_v2.rds")
chick
```

```
## An object of class Seurat
## 18523 features across 82092 samples within 1 assay
## Active assay: RNA (18523 features, 2000 variable features)
##  3 layers present: data, counts, scale.data
##  3 dimensional reductions calculated: pca, harmony, umap
```

```
Idents(chick) <- "stage"
chick=subset(chick, idents =c("HH8"))
```

```
chick
```

```
## An object of class Seurat
## 18523 features across 18875 samples within 1 assay
## Active assay: RNA (18523 features, 2000 variable features)
##  3 layers present: data, counts, scale.data
##  3 dimensional reductions calculated: pca, harmony, umap
```

```
gc()
```

```
##              used    (Mb) gc trigger    (Mb)   max used    (Mb)
## Ncells    5009499   267.6   13651909   729.1   21331105  1139.3
## Vcells 1428472570 10898.4 3660127464 27924.6 4574774951 34902.8
```

```
chick <- NormalizeData(chick)
```

```
## Normalizing layer: counts
```

```
chick <- CellCycleScoring(chick, s.features = cc.genes$s.genes, g2m.features = cc.genes$g2m.ge
nes, set.ident = TRUE)
chick <- FindVariableFeatures(chick)
```

```
## Finding variable features for layer counts
```

```
chick <- ScaleData(chick, vars.to.regress = c("percent.mt","G2M.Score","S.Score"))
```

```
## Regressing out percent.mt, G2M.Score, S.Score
```

```
## Centering and scaling data matrix
```

```
chick <- RunPCA(chick, npcs = 50, verbose = FALSE)

ElbowPlot(chick, ndims= 50, reduction = "pca")
```

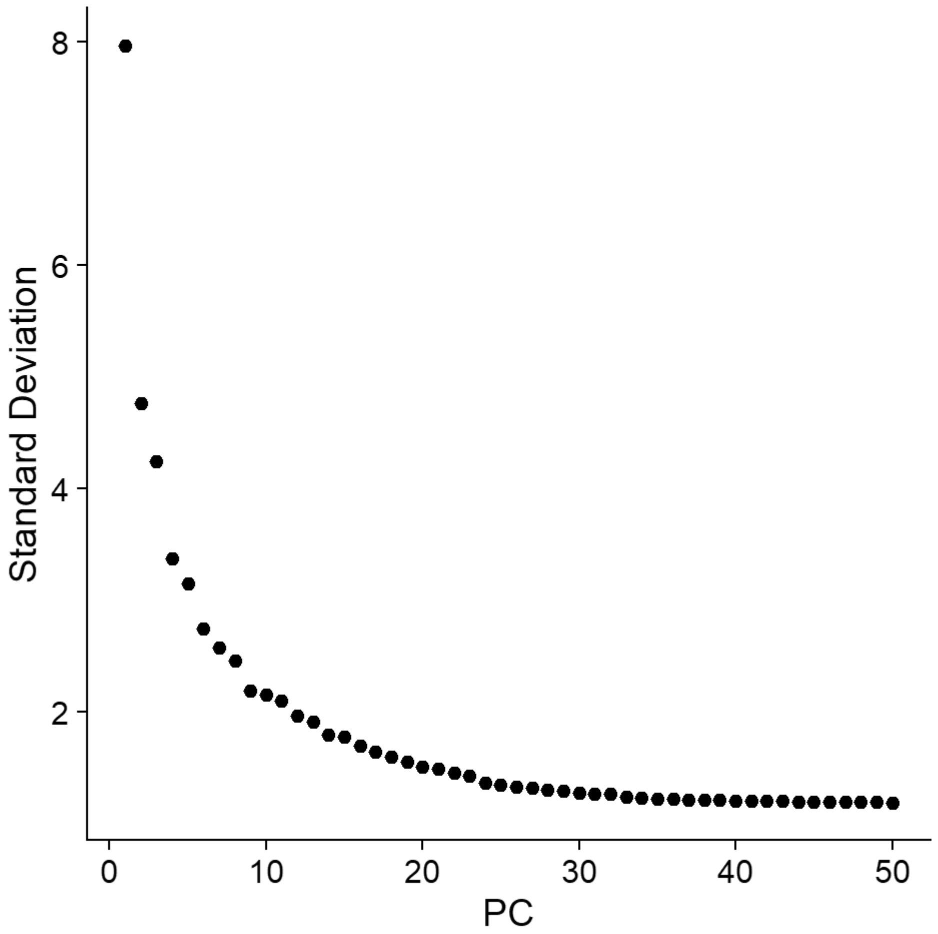


```
gc()
```

```
##              used    (Mb) gc trigger    (Mb)   max used    (Mb)
## Ncells    5031235   268.7   13651909   729.1   21331105  1139.3
## Vcells 1428619858 10899.6 3660127464 27924.6 4574774951 34902.8
```

```
chick[["RNA"]] <- split(chick[["RNA"]], f = chick$sample_id)
```

```
## Splitting 'counts', 'data' layers. Not splitting 'scale.data'. If you would like to split o
ther layers, set in `layers` argument.
```

```
chick <- IntegrateLayers(object=chick, method=HarmonyIntegration, orig.reduction="pca",
                       new.reduction="harmony", assay="RNA", group.by="stage", verbose=TRUE)
```

```
## Transposing data matrix
```

```
## Using automatic lambda estimation
```

```
## Initializing state using k-means centroids initialization
```

```
## Harmony 1/10
```

```
## Harmony 2/10
```

```
## Harmony converged after 2 iterations
```

```
chick <- FindNeighbors(chick, reduction="harmony", dims=1:30, nn.method="rann", k.param=20)
```

```
## Computing nearest neighbor graph
```

```
## Computing SNN
```

```
chick <- RunUMAP(chick, reduction="harmony", dims=1:30)
```

```
## 23:10:08 UMAP embedding parameters a = 0.9922 b = 1.112
```

```
## 23:10:08 Read 18875 rows and found 30 numeric columns
```

```
## 23:10:08 Using Annoy for neighbor search, n_neighbors = 30
```

```
## 23:10:08 Building Annoy index with metric = cosine, n_trees = 50
```

```
## 0%   10   20   30   40   50   60   70   80   90   100%
```

```
## [----|----|----|----|----|----|----|----|----|----|
```

```
## **************************************************|
## 23:10:09 Writing NN index file to temp file C:\Users\RANEES~1\AppData\Local\Temp\RtmpqcWgVb
\file6a2c309551
## 23:10:09 Searching Annoy index using 1 thread, search_k = 3000
## 23:10:14 Annoy recall = 100%
## 23:10:14 Commencing smooth kNN distance calibration using 1 thread with target n_neighbors
= 30
## 23:10:15 Initializing from normalized Laplacian + noise (using RSpectra)
## 23:10:16 Commencing optimization for 200 epochs, with 855018 positive edges
## 23:10:16 Using rng type: pcg
## 23:10:32 Optimization finished
```

```
chick <- FindClusters(chick, resolution=0.4)
```

```
## Modularity Optimizer version 1.3.0 by Ludo Waltman and Nees Jan van Eck
##
## Number of nodes: 18875
## Number of edges: 682194
##
```

```
## Running Louvain algorithm...
## Maximum modularity in 10 random starts: 0.9136
## Number of communities: 13
## Elapsed time: 2 seconds
```

```
DimPlot(chick)
```

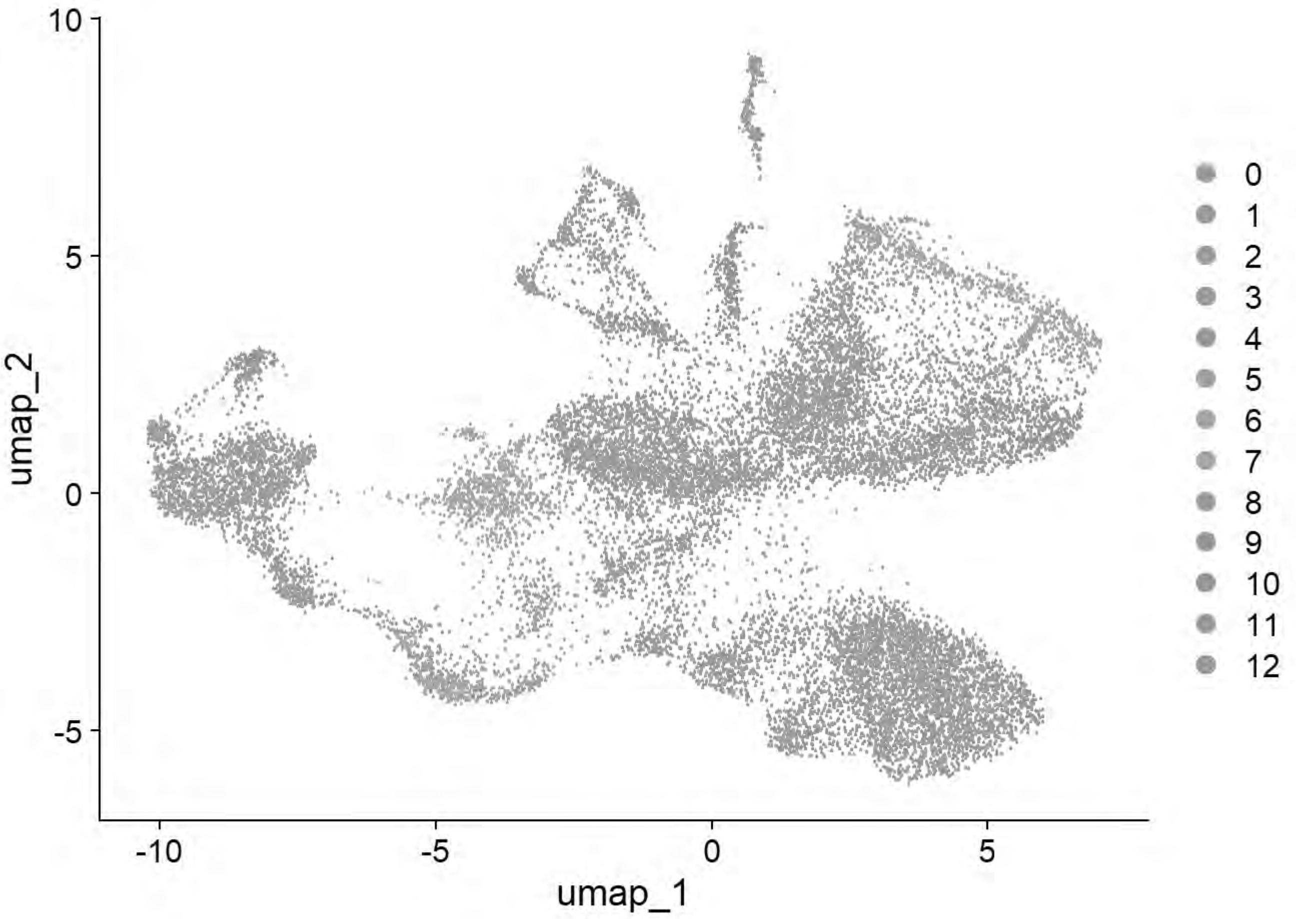


```
DimPlot(chick, group.by = "sample_id")
```

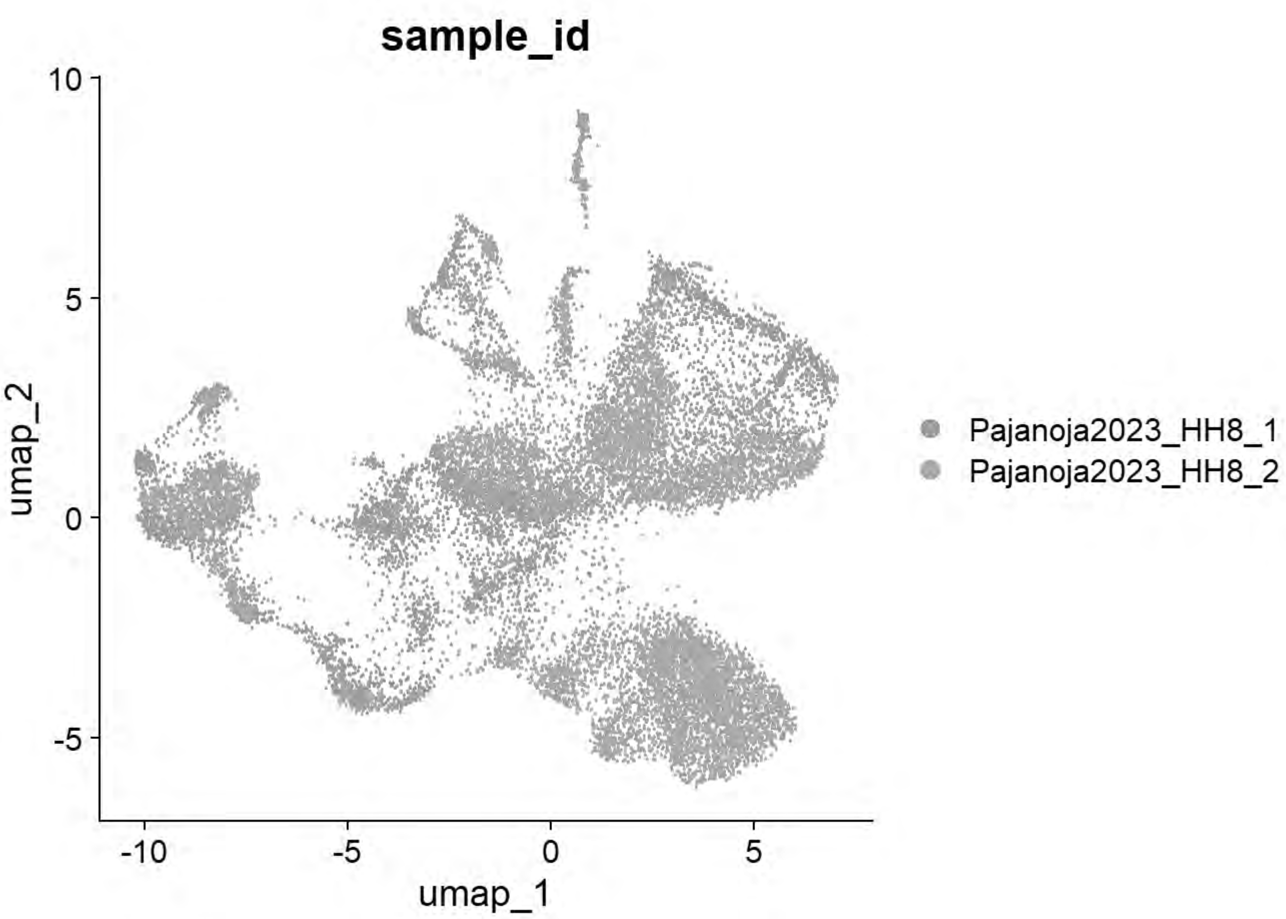


```
chick = FindClusters(chick, resolution= seq(0,2,0.1),n.start=10)
```

```
## Modularity Optimizer version 1.3.0 by Ludo Waltman and Nees Jan van Eck
##
## Number of nodes: 18875
## Number of edges: 682194
##
## Running Louvain algorithm...
## Maximum modularity in 10 random starts: 1.0000
## Number of communities: 1
## Elapsed time: 1 seconds
## Modularity Optimizer version 1.3.0 by Ludo Waltman and Nees Jan van Eck
##
## Number of nodes: 18875
## Number of edges: 682194
##
## Running Louvain algorithm...
## Maximum modularity in 10 random starts: 0.9593
## Number of communities: 8
## Elapsed time: 2 seconds
## Modularity Optimizer version 1.3.0 by Ludo Waltman and Nees Jan van Eck
##
## Number of nodes: 18875
## Number of edges: 682194
##
```

```
## Running Louvain algorithm...
## Maximum modularity in 10 random starts: 0.9414
## Number of communities: 10
## Elapsed time: 2 seconds
## Modularity Optimizer version 1.3.0 by Ludo Waltman and Nees Jan van Eck
##
## Number of nodes: 18875
## Number of edges: 682194
##
## Running Louvain algorithm...
## Maximum modularity in 10 random starts: 0.9268
## Number of communities: 11
## Elapsed time: 2 seconds
## Modularity Optimizer version 1.3.0 by Ludo Waltman and Nees Jan van Eck
##
## Number of nodes: 18875
## Number of edges: 682194
##
## Running Louvain algorithm...
## Maximum modularity in 10 random starts: 0.9136
## Number of communities: 13
## Elapsed time: 2 seconds
## Modularity Optimizer version 1.3.0 by Ludo Waltman and Nees Jan van Eck
##
## Number of nodes: 18875
## Number of edges: 682194
##
## Running Louvain algorithm...
## Maximum modularity in 10 random starts: 0.9017
## Number of communities: 13
## Elapsed time: 2 seconds
## Modularity Optimizer version 1.3.0 by Ludo Waltman and Nees Jan van Eck
##
## Number of nodes: 18875
## Number of edges: 682194
##
## Running Louvain algorithm...
## Maximum modularity in 10 random starts: 0.8911
## Number of communities: 15
## Elapsed time: 2 seconds
## Modularity Optimizer version 1.3.0 by Ludo Waltman and Nees Jan van Eck
##
## Number of nodes: 18875
## Number of edges: 682194
##
## Running Louvain algorithm...
## Maximum modularity in 10 random starts: 0.8817
## Number of communities: 17
## Elapsed time: 2 seconds
## Modularity Optimizer version 1.3.0 by Ludo Waltman and Nees Jan van Eck
##
## Number of nodes: 18875
## Number of edges: 682194
##
```

```
## Running Louvain algorithm...
## Maximum modularity in 10 random starts: 0.8726
## Number of communities: 18
## Elapsed time: 2 seconds
## Modularity Optimizer version 1.3.0 by Ludo Waltman and Nees Jan van Eck
##
## Number of nodes: 18875
## Number of edges: 682194
##
## Running Louvain algorithm...
## Maximum modularity in 10 random starts: 0.8633
## Number of communities: 18
## Elapsed time: 2 seconds
## Modularity Optimizer version 1.3.0 by Ludo Waltman and Nees Jan van Eck
##
## Number of nodes: 18875
## Number of edges: 682194
##
## Running Louvain algorithm...
## Maximum modularity in 10 random starts: 0.8555
## Number of communities: 21
## Elapsed time: 2 seconds
## Modularity Optimizer version 1.3.0 by Ludo Waltman and Nees Jan van Eck
##
## Number of nodes: 18875
## Number of edges: 682194
##
## Running Louvain algorithm...
## Maximum modularity in 10 random starts: 0.8483
## Number of communities: 23
## Elapsed time: 2 seconds
## Modularity Optimizer version 1.3.0 by Ludo Waltman and Nees Jan van Eck
##
## Number of nodes: 18875
## Number of edges: 682194
##
## Running Louvain algorithm...
## Maximum modularity in 10 random starts: 0.8424
## Number of communities: 23
## Elapsed time: 2 seconds
## Modularity Optimizer version 1.3.0 by Ludo Waltman and Nees Jan van Eck
##
## Number of nodes: 18875
## Number of edges: 682194
##
## Running Louvain algorithm...
## Maximum modularity in 10 random starts: 0.8360
## Number of communities: 23
## Elapsed time: 2 seconds
## Modularity Optimizer version 1.3.0 by Ludo Waltman and Nees Jan van Eck
##
## Number of nodes: 18875
## Number of edges: 682194
##
```

```
## Running Louvain algorithm...
## Maximum modularity in 10 random starts: 0.8296
## Number of communities: 23
## Elapsed time: 2 seconds
## Modularity Optimizer version 1.3.0 by Ludo Waltman and Nees Jan van Eck
##
## Number of nodes: 18875
## Number of edges: 682194
##
## Running Louvain algorithm...
## Maximum modularity in 10 random starts: 0.8228
## Number of communities: 25
## Elapsed time: 2 seconds
## Modularity Optimizer version 1.3.0 by Ludo Waltman and Nees Jan van Eck
##
## Number of nodes: 18875
## Number of edges: 682194
##
## Running Louvain algorithm...
## Maximum modularity in 10 random starts: 0.8172
## Number of communities: 28
## Elapsed time: 2 seconds
## Modularity Optimizer version 1.3.0 by Ludo Waltman and Nees Jan van Eck
##
## Number of nodes: 18875
## Number of edges: 682194
##
## Running Louvain algorithm...
## Maximum modularity in 10 random starts: 0.8116
## Number of communities: 28
## Elapsed time: 2 seconds
## Modularity Optimizer version 1.3.0 by Ludo Waltman and Nees Jan van Eck
##
## Number of nodes: 18875
## Number of edges: 682194
##
## Running Louvain algorithm...
## Maximum modularity in 10 random starts: 0.8060
## Number of communities: 28
## Elapsed time: 2 seconds
## Modularity Optimizer version 1.3.0 by Ludo Waltman and Nees Jan van Eck
##
## Number of nodes: 18875
## Number of edges: 682194
##
## Running Louvain algorithm...
## Maximum modularity in 10 random starts: 0.8004
## Number of communities: 28
## Elapsed time: 2 seconds
## Modularity Optimizer version 1.3.0 by Ludo Waltman and Nees Jan van Eck
##
## Number of nodes: 18875
## Number of edges: 682194
##
```

```
## Running Louvain algorithm...
## Maximum modularity in 10 random starts: 0.7948
## Number of communities: 28
## Elapsed time: 2 seconds
```

```
p=clustree(chick, prefix= "RNA_snn_res.")
print(p)
```

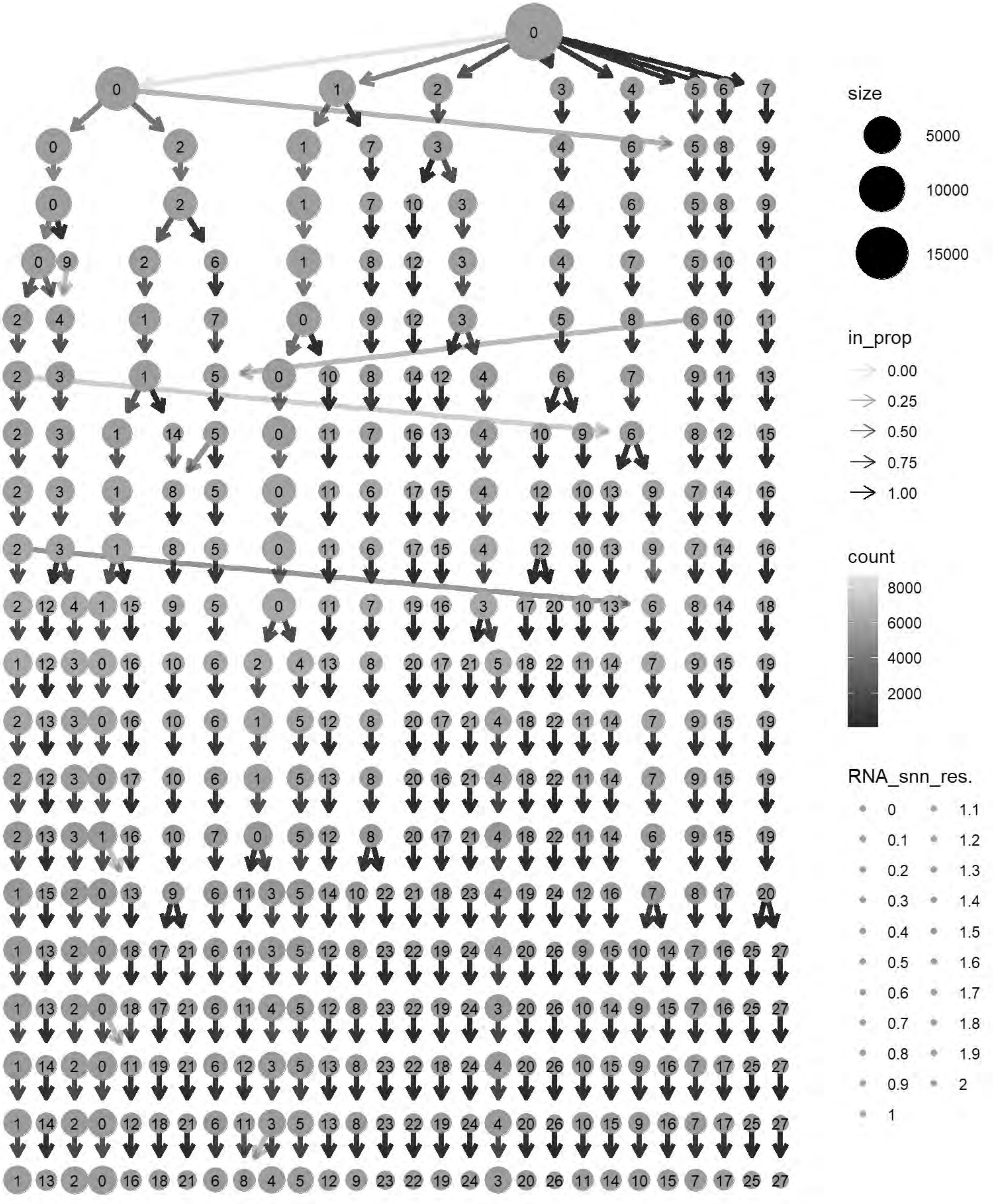
size
5000
10000
15000
in_prop
0.00
0.25
0.50
0.75
1.00
count
8000
6000
4000
2000
RNA_snn_res.
0
0.1
0.2
0.3
0.4
0.5
0.6
0.7
0.8
0.9
1
1.1
1.2
1.3
1.4
1.5
1.6
1.7
1.8
1.9
2

```
cols= grep("^RNA_snn_res.",names(chick@meta.data))
chick@meta.data[cols] <- NULL
chick <- FindClusters(chick, resolution = 0.3, verbose = TRUE)
```

```
## Modularity Optimizer version 1.3.0 by Ludo Waltman and Nees Jan van Eck
##
## Number of nodes: 18875
## Number of edges: 682194
##
## Running Louvain algorithm...
## Maximum modularity in 10 random starts: 0.9268
## Number of communities: 11
## Elapsed time: 2 seconds
```

```
p6=DimPlot(chick, label = TRUE)+NoLegend()
print(p6)
```

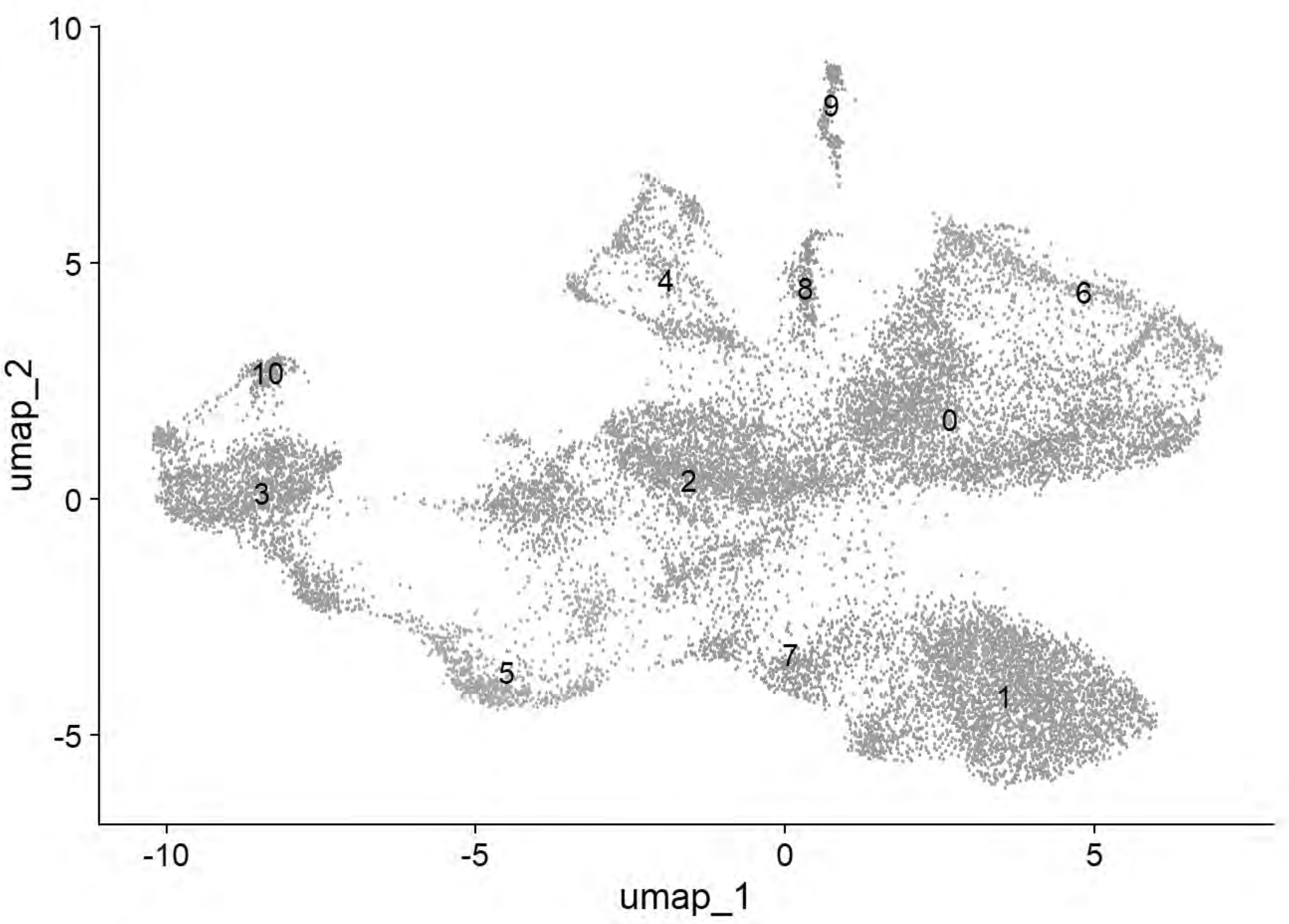


```
p6=DimPlot(chick, group.by = "Phase")
print(p6)
```

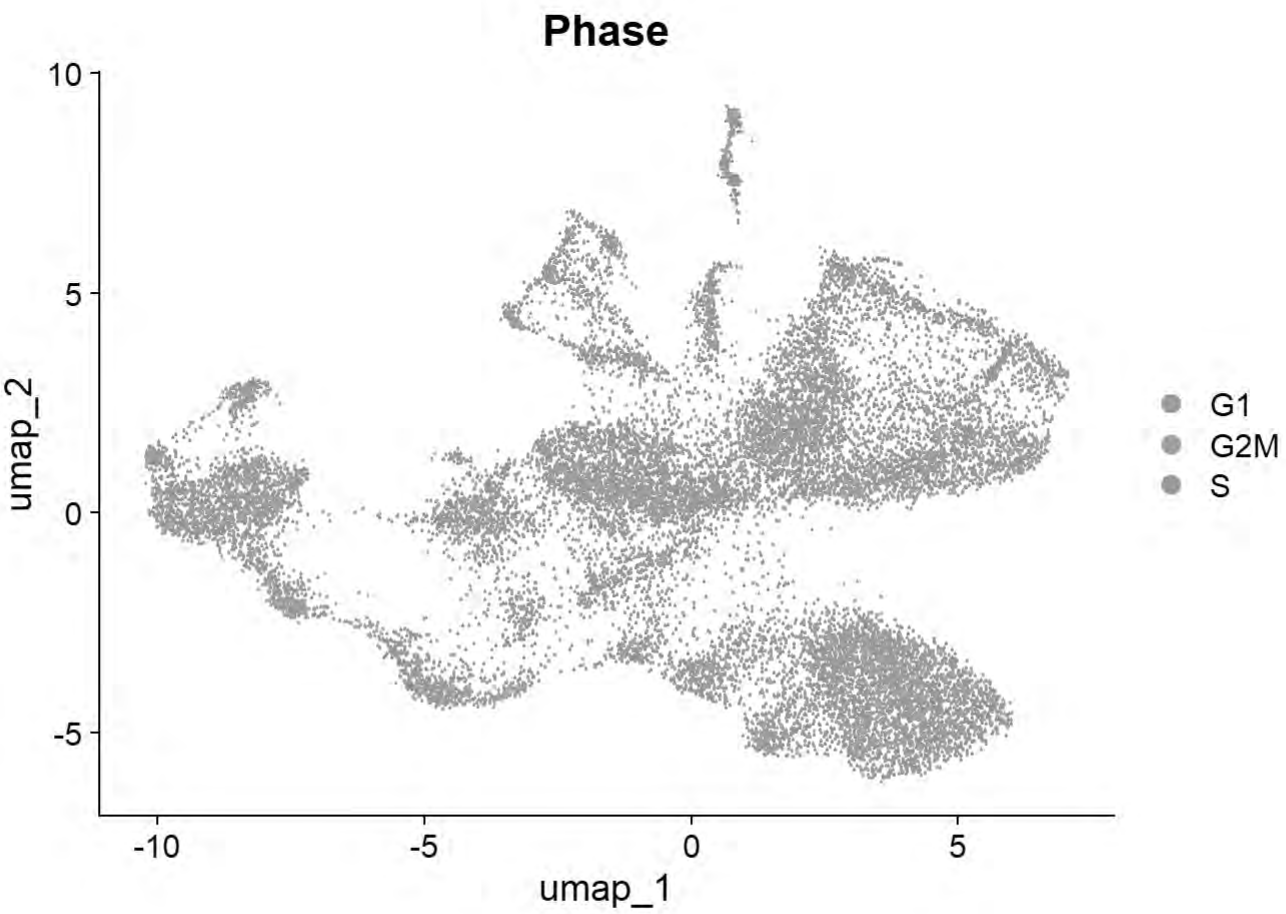


```
p6=DimPlot(chick, group.by = "sample_id")
print(p6)
```

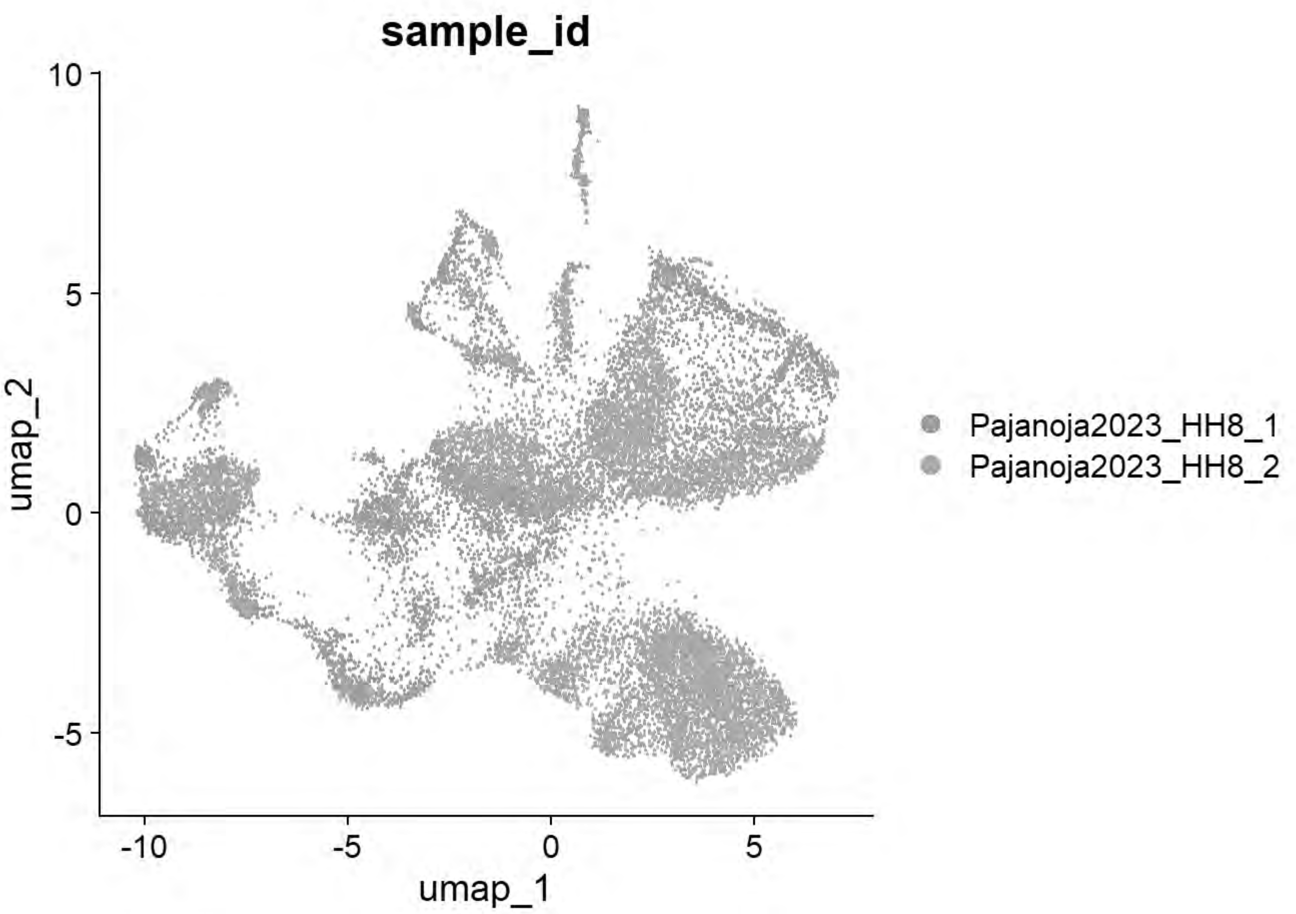


```
# Williams: Neural plate/tube
FeaturePlot(chick, features = c("CLDN1","FRZB","DNMT3A"))
```

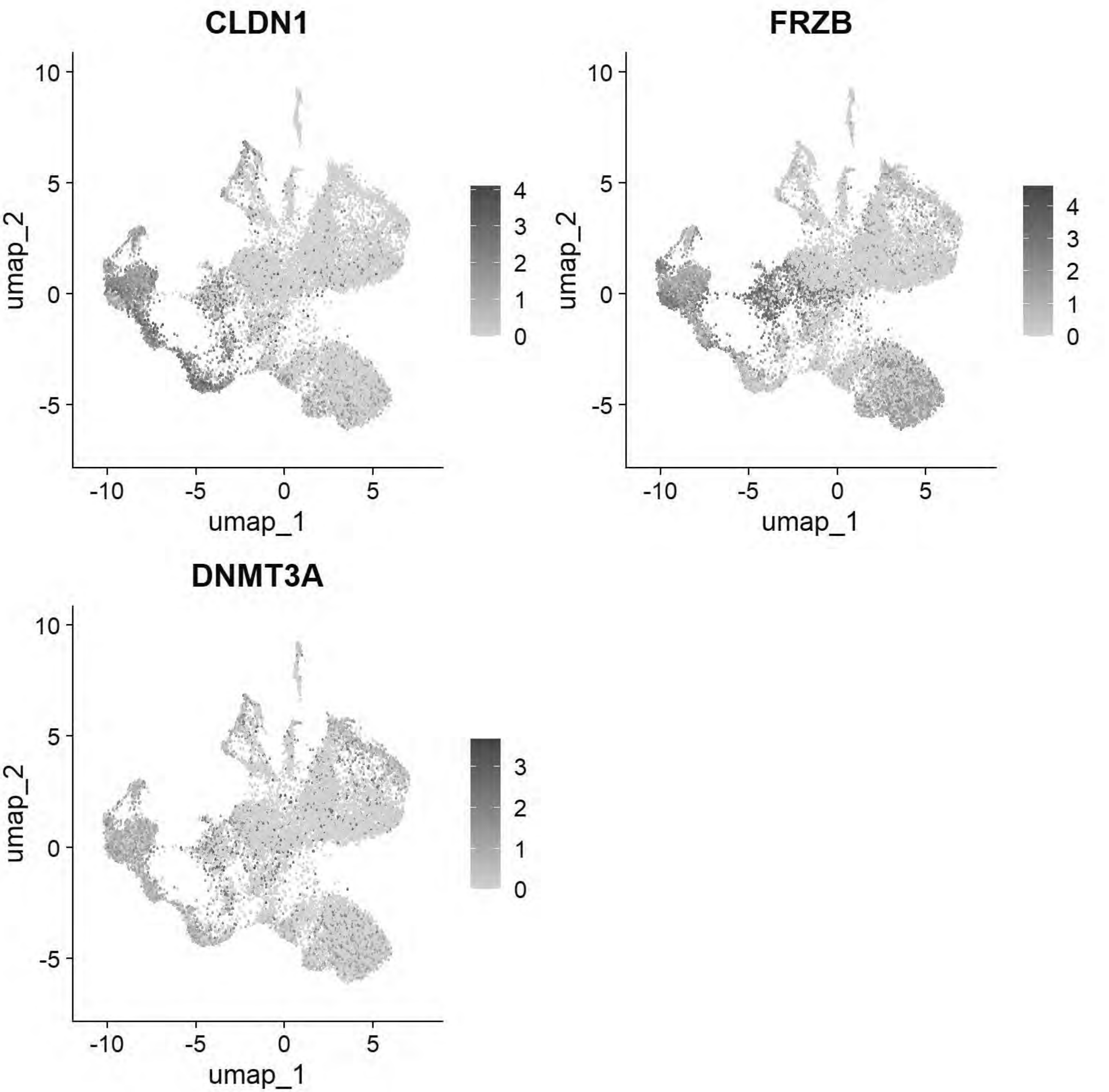


```
# Williams: Non-Neural Ectoderm
FeaturePlot(chick, features = c("DLX5","TFAP2A","ASTL","PAX6"))
```

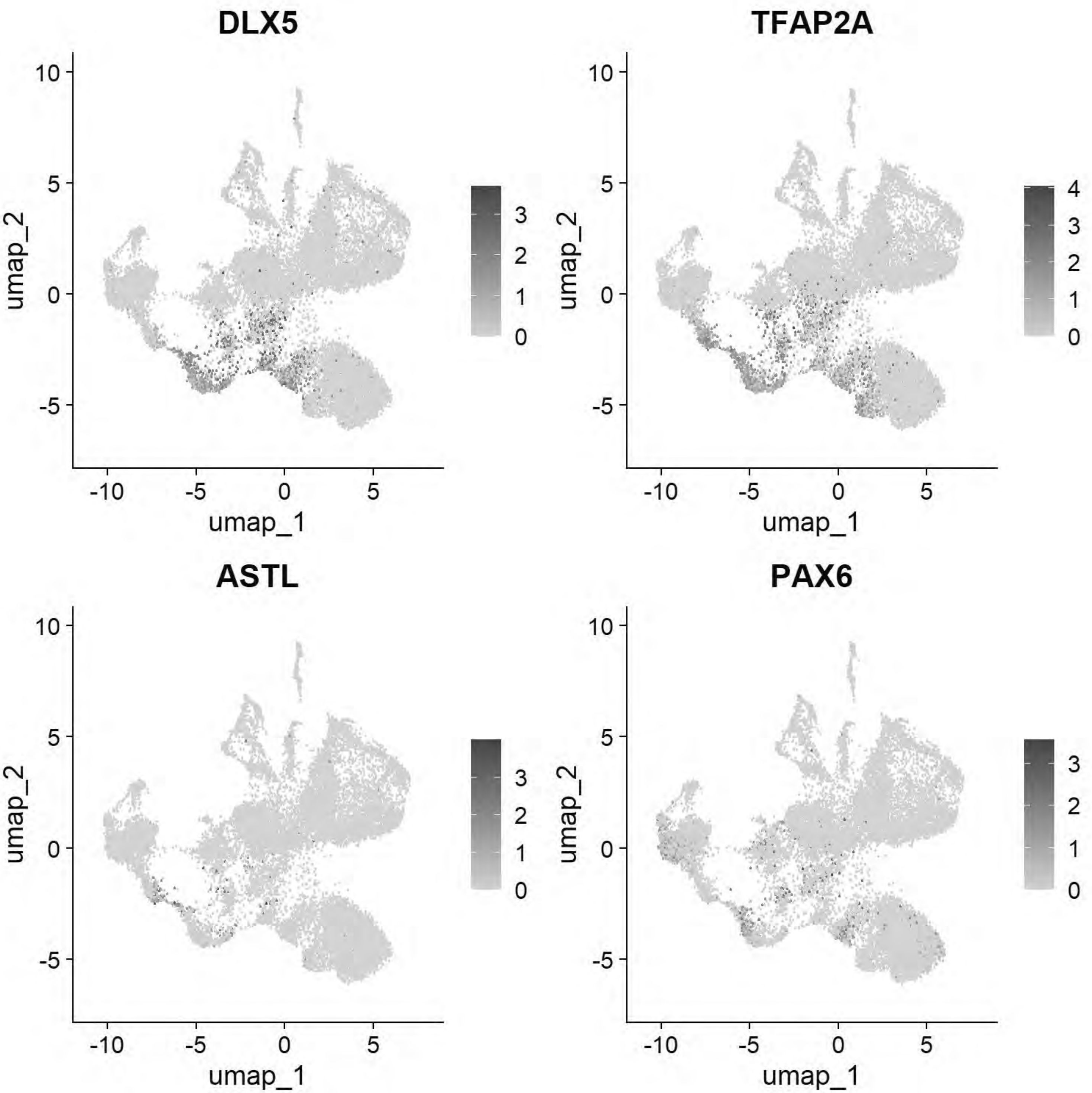


```
# Williams: Ectoderm (Neural Plate)
FeaturePlot(chick, features = c("SOX2","SOX3","SOX21","FRZB","SFRP2"))
```

```
## Warning: All cells have the same value (0) of "SOX3"
```

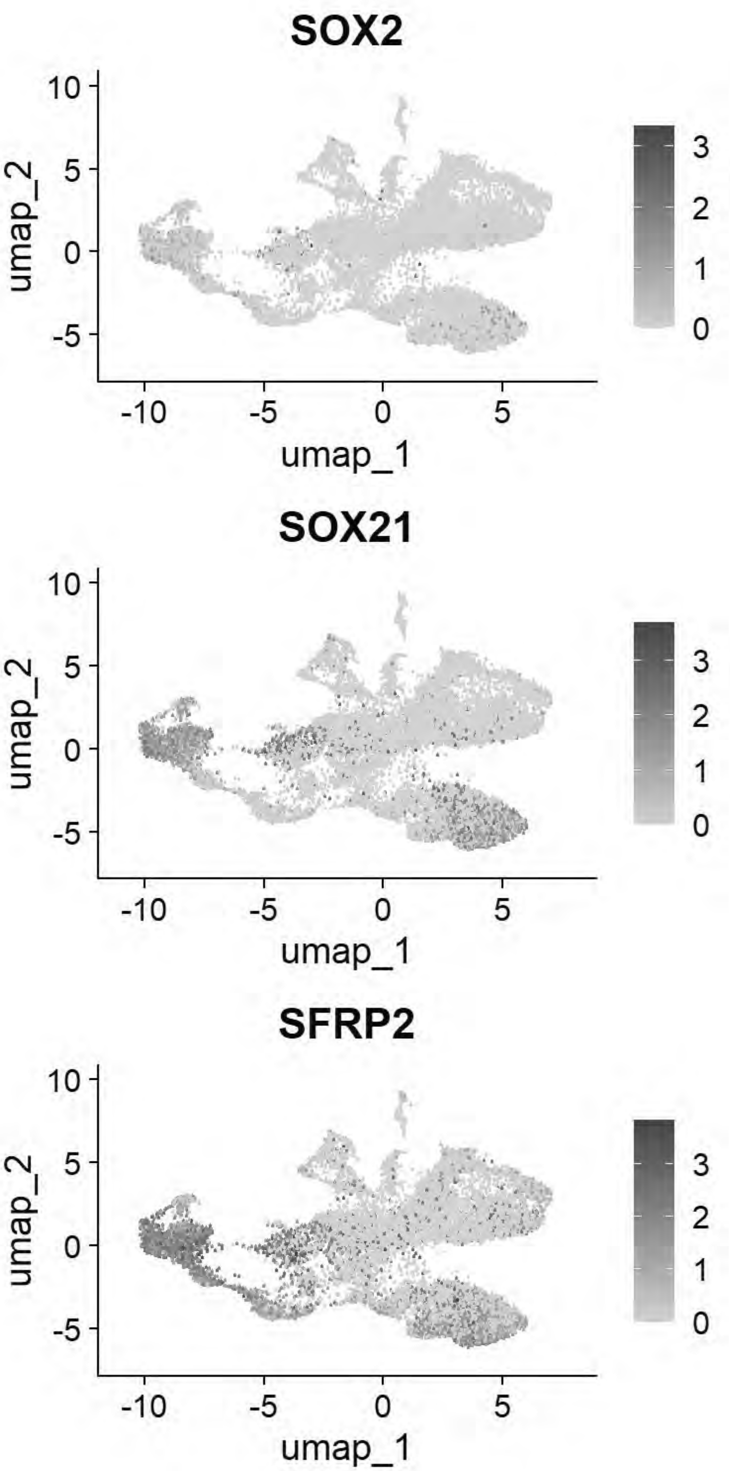


```
# Williams: Ectoderm (Neural Plate Border)
FeaturePlot(chick, features = c("PAX7","TFAP2A","DLX5","BMP4","MSX1","DRAXIN","TFAP2B"))
```

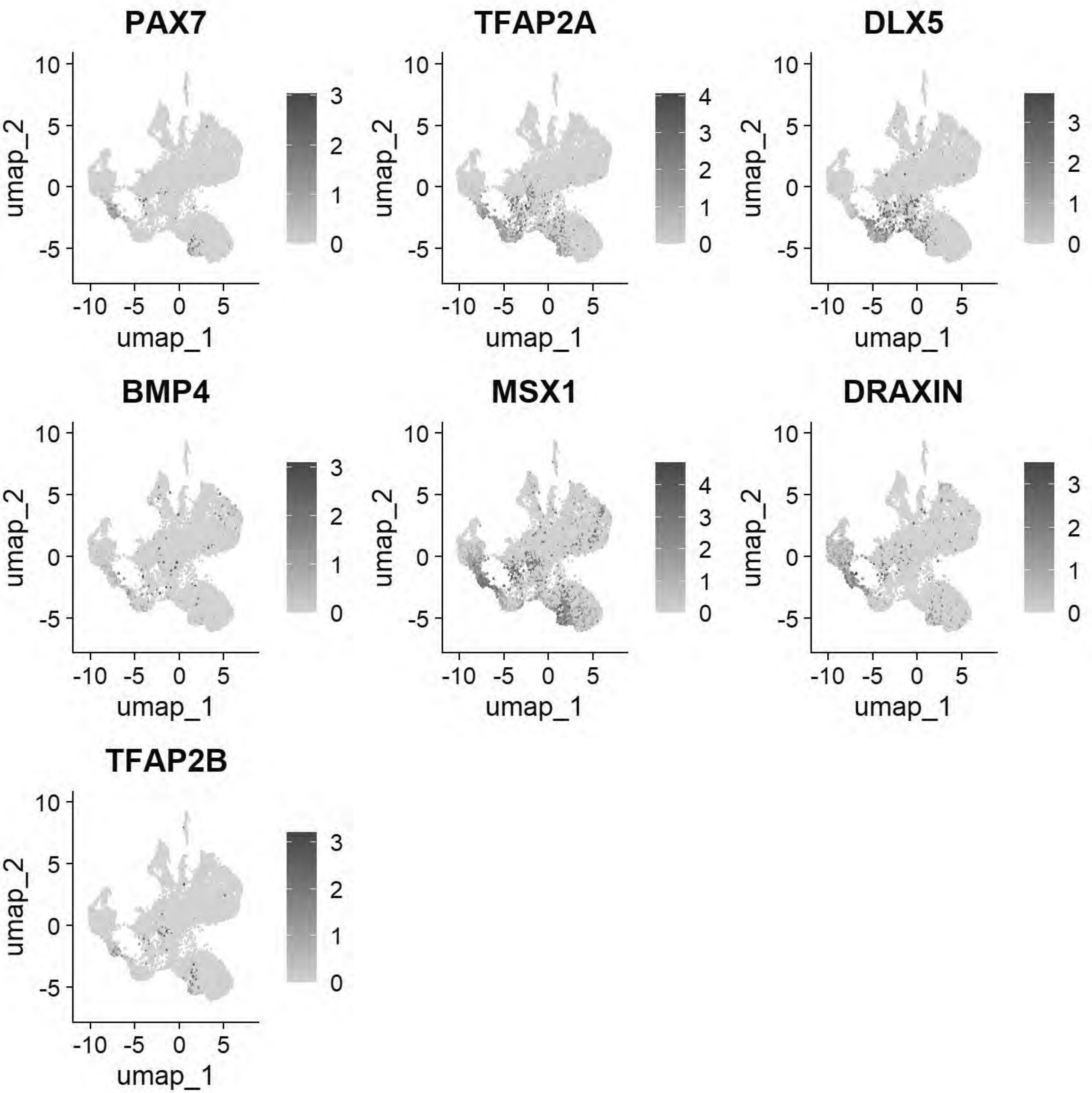


```
# Williams: Posterior Lateral Plate Mesoderm
FeaturePlot(chick, features = c("GATA2","HOXB5","CDX4"))
```

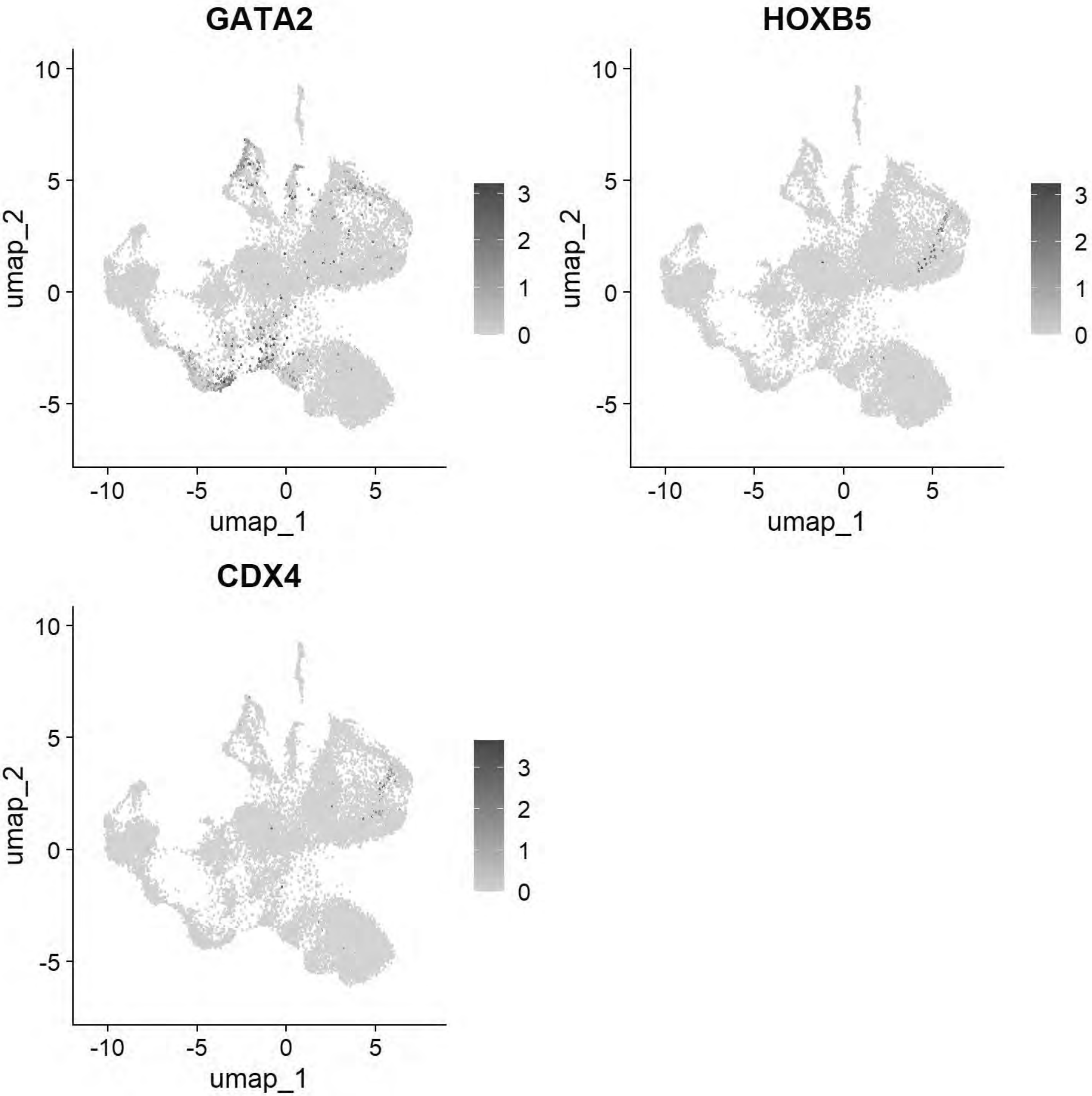


```r
# Williams: Paraxial Mesoderm
FeaturePlot(chick, features = c("MSGN1","MESP1","MEOX1"))
```

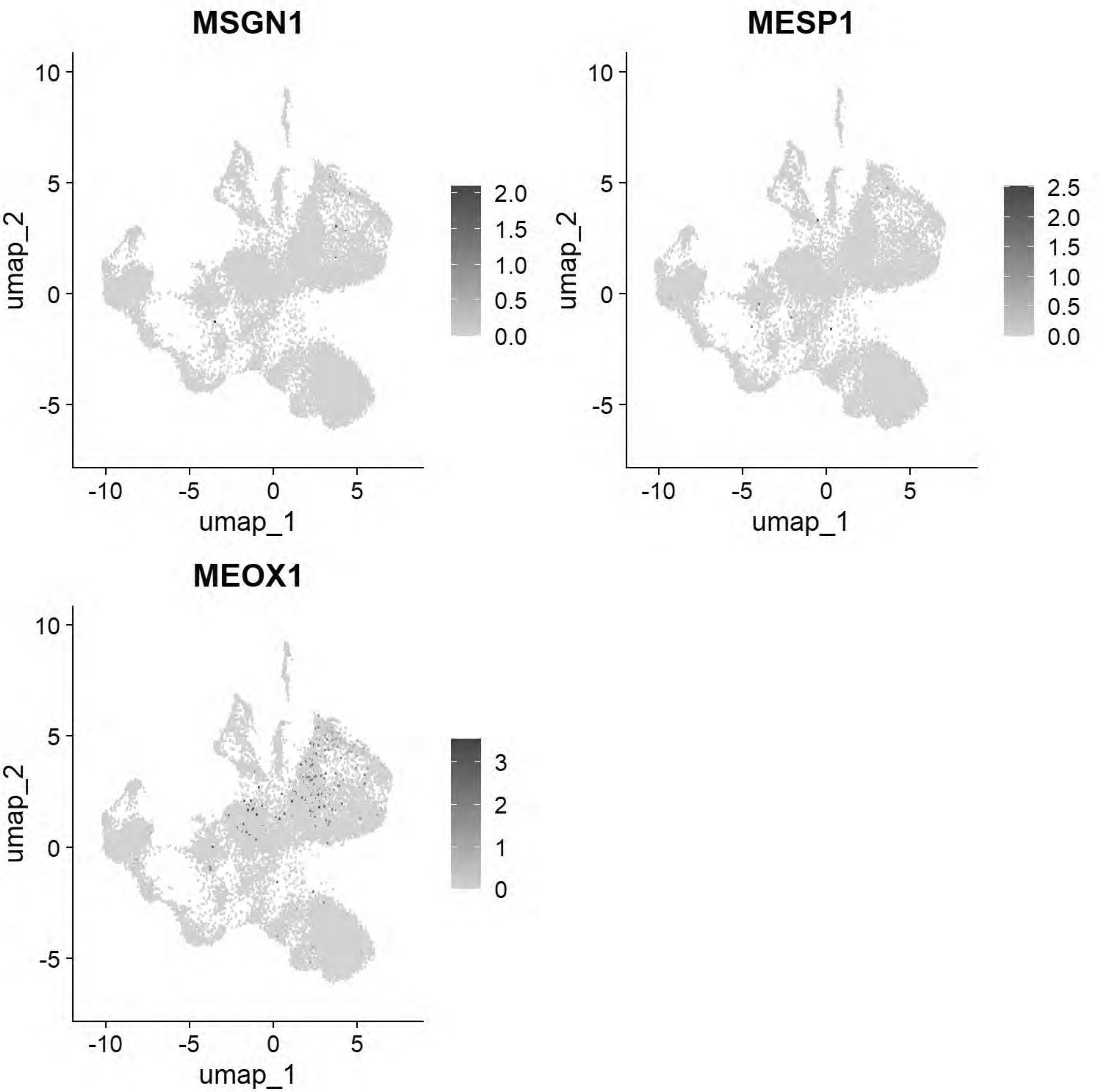


```r
# Williams: Lateral Plate Mesoderm
FeaturePlot(chick, features = c("PITX2","ALX1","OLFML3","SIX1","TWIST1"))
```

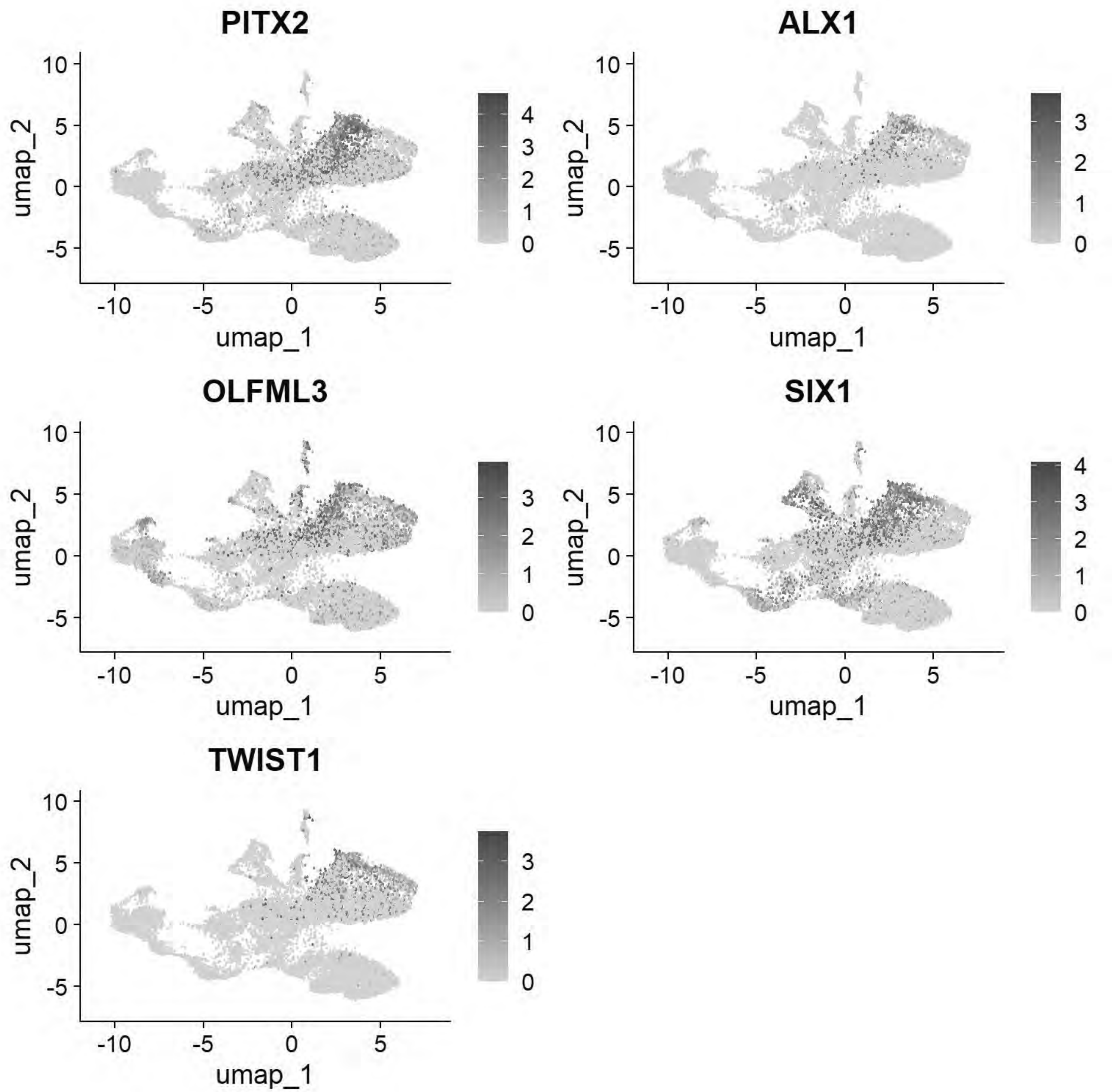


```
# Williams: Head Mesenchyme
FeaturePlot(chick, features = c("TCF21","GATA5","LMO2","ETS1","KDR"))
```

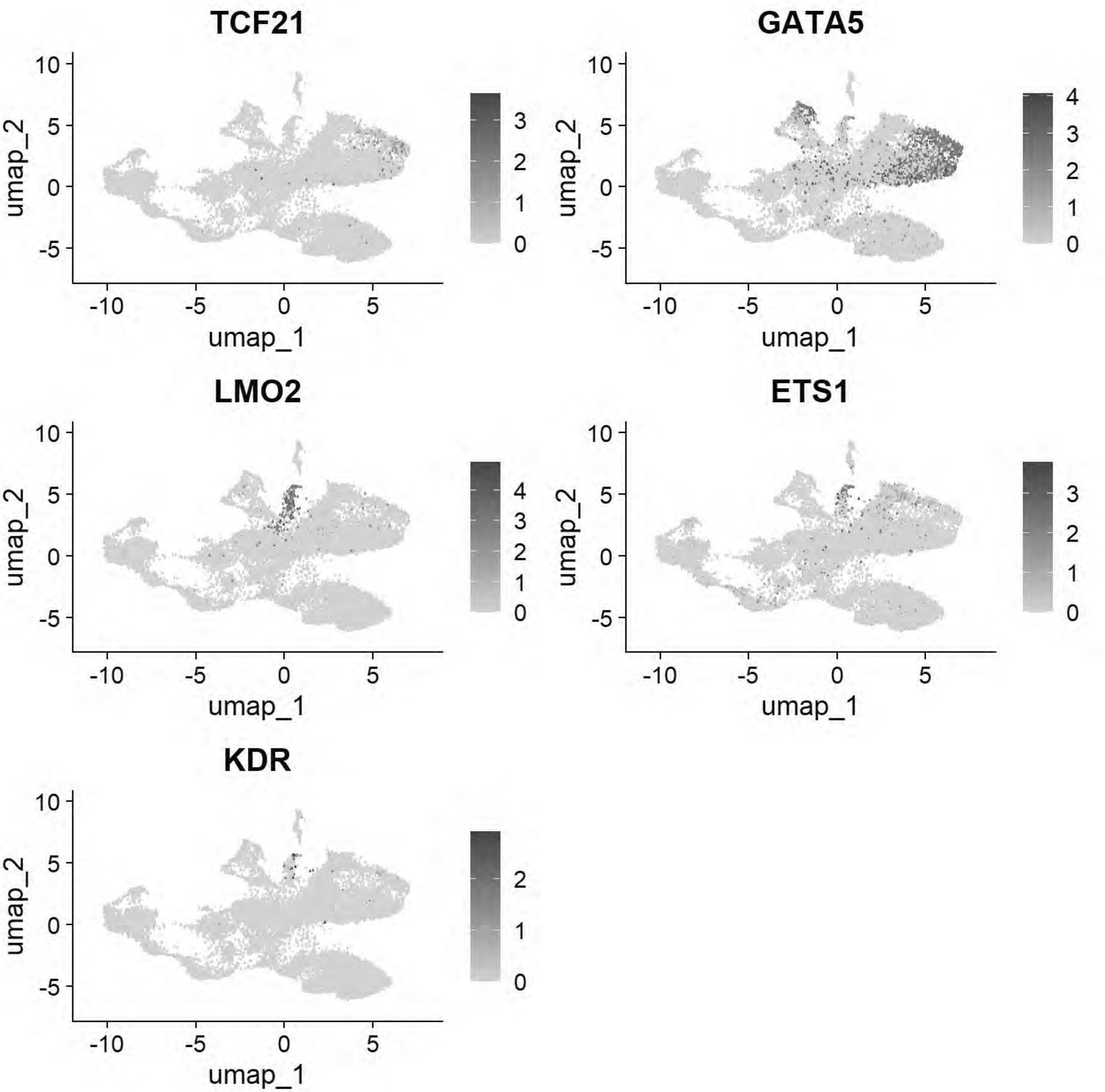


```
# Williams: Hensen's Node & Primitive Streak
FeaturePlot(chick, features = c("DLL1","FGF8","NOTO","CHRD"))
```

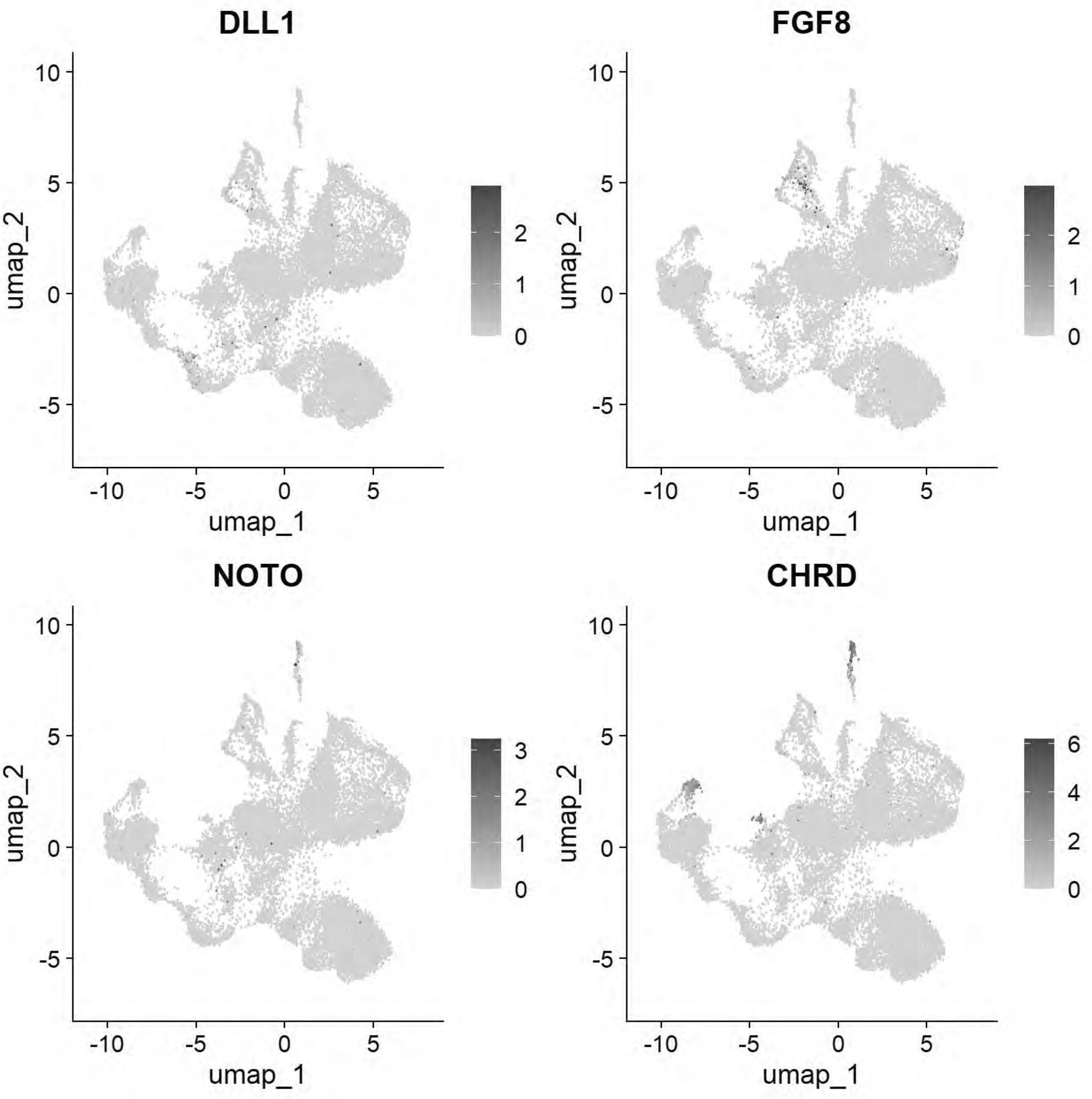


```r
# Williams: Endoderm
FeaturePlot(chick, features = c("SOX17","FOXA2","CXCR4"))
```

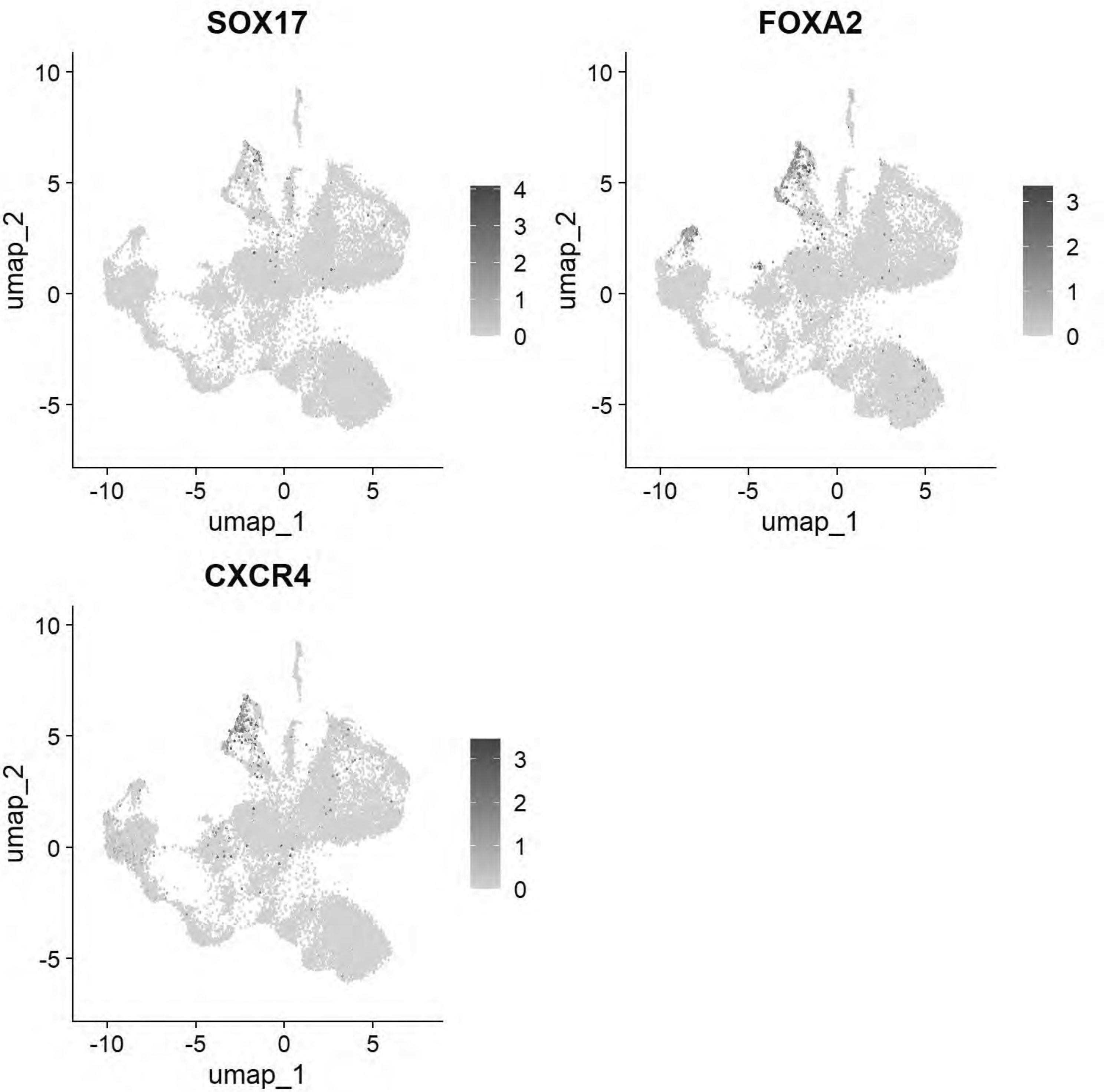


```
# Pajanoja: Ectoderm
FeaturePlot(chick, features = c("TFAP2A","DLX5","CLDN1","SOX2","NESTIN","MYCN"))
```

```
## Warning: The following requested variables were not found: NESTIN
```

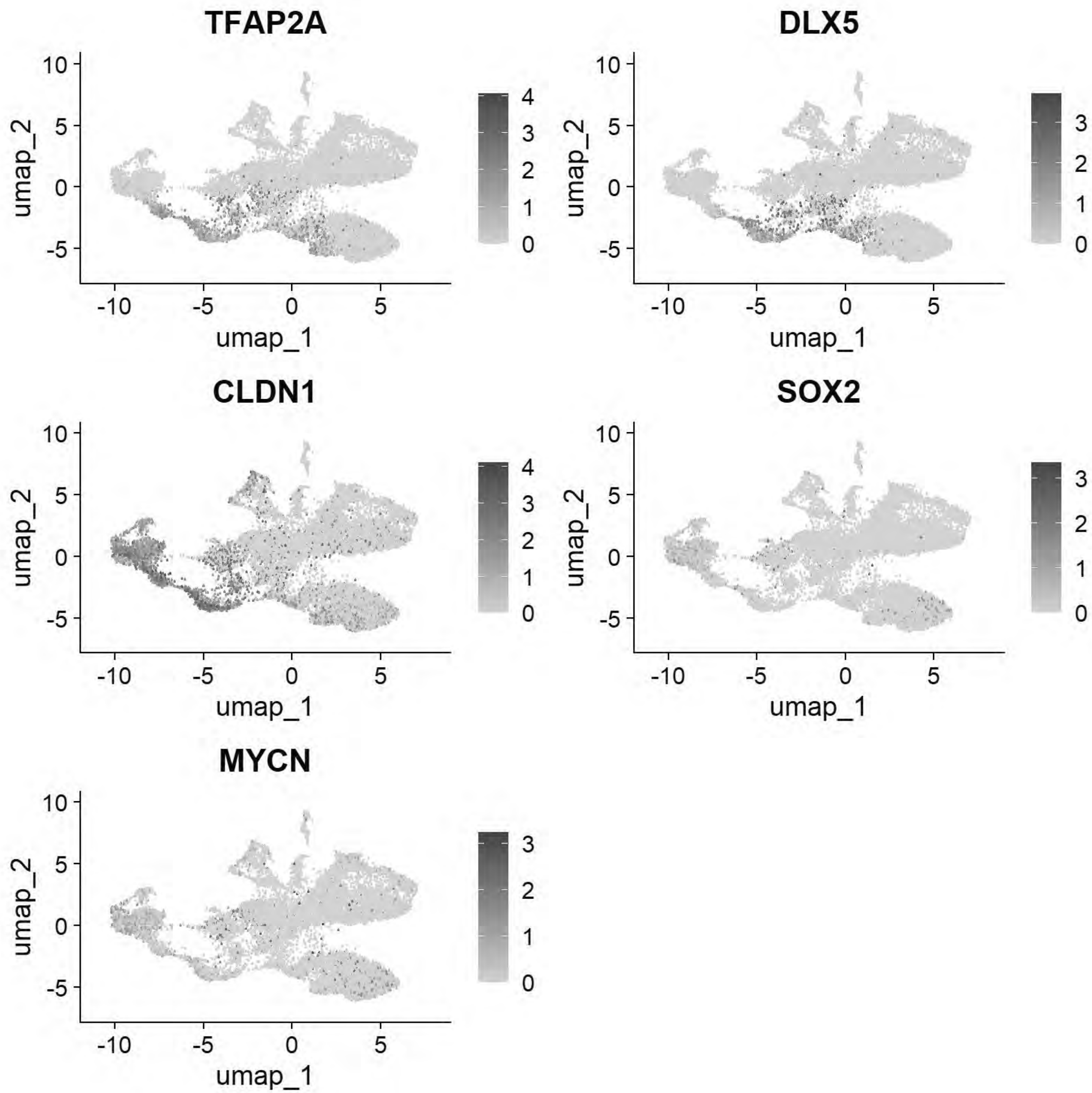


```
# Pajanoja: Endoderm
FeaturePlot(chick, features = c("SOX17","KRT17","HHEX"))
```

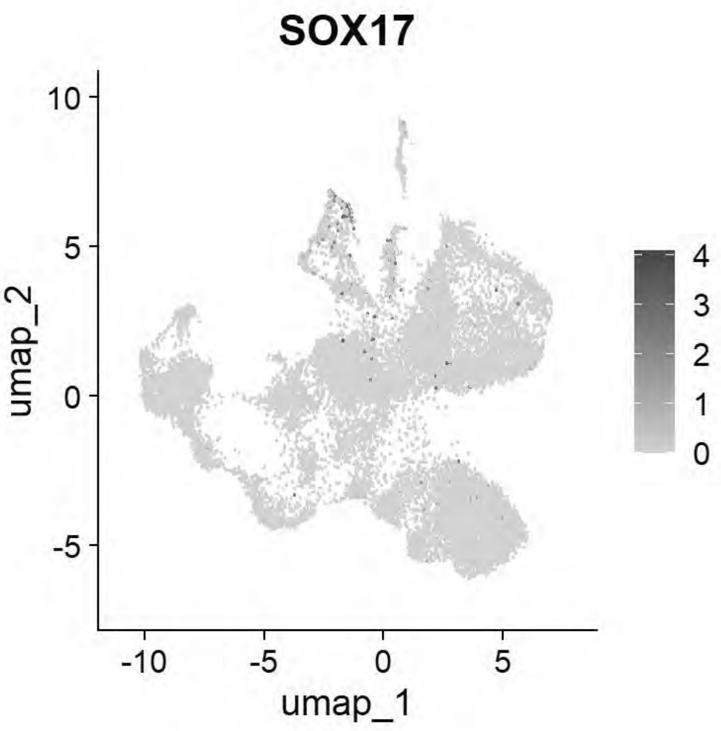


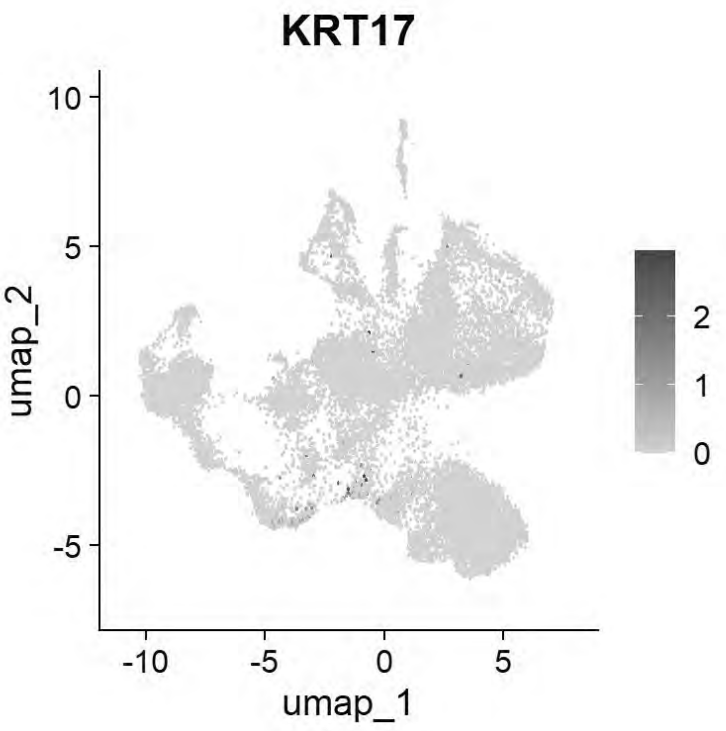


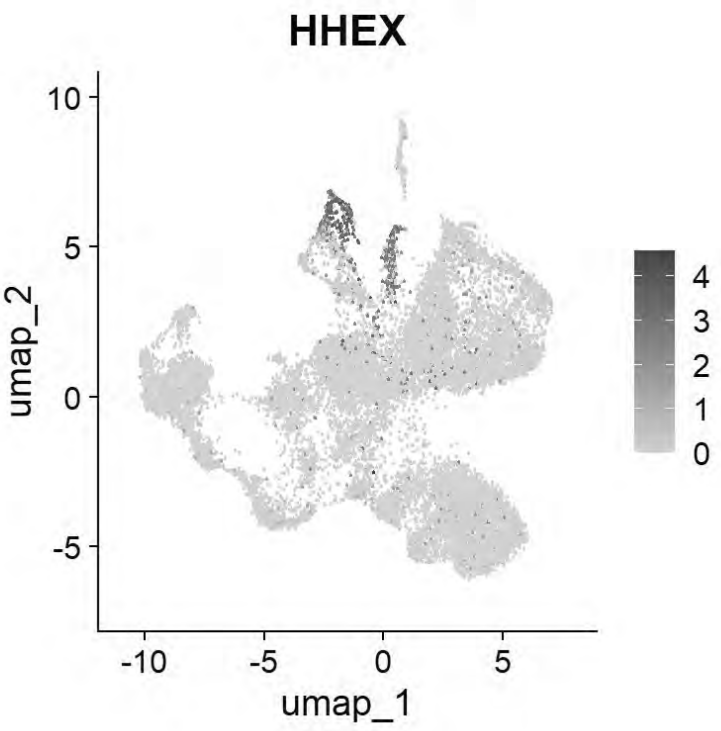


```
# Pajanoja: Mesoderm
FeaturePlot(chick, features = c("ALX1","TWIST1","HAND2","PITX2"))
```

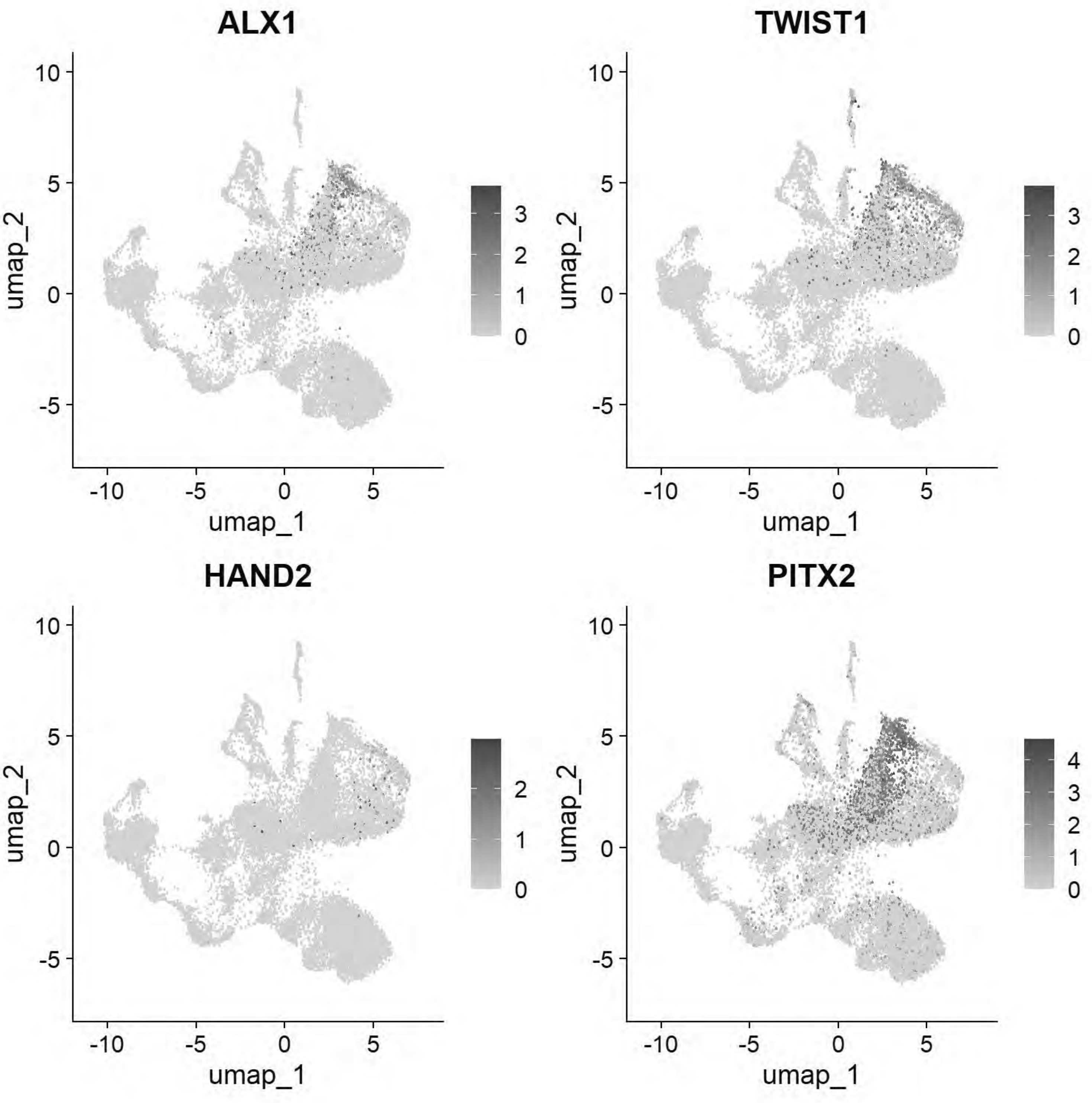


```
# Pajanoja: Notochord
FeaturePlot(chick, features = c("CHRD","TBXT","NOTO"))
```

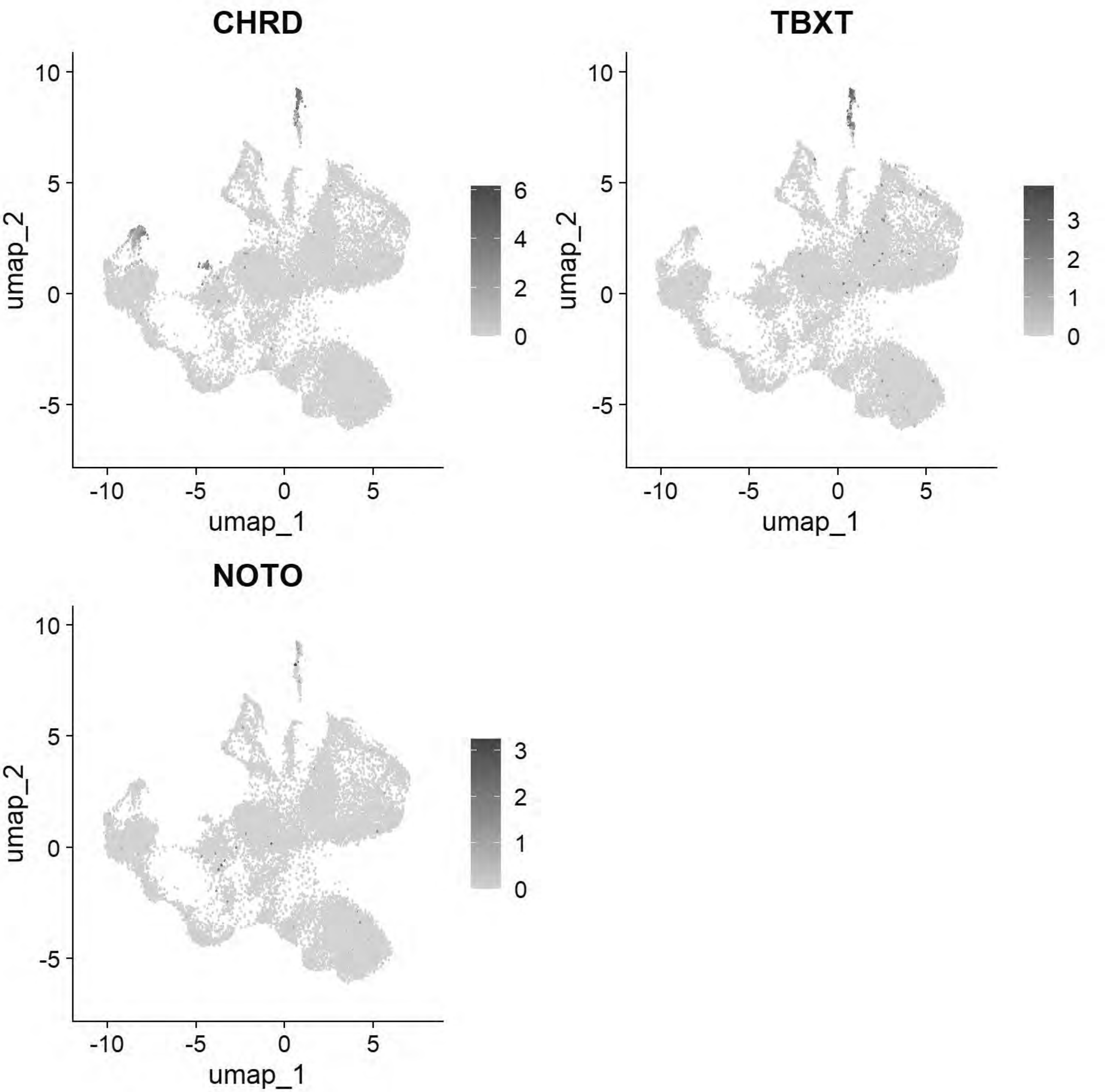


```
gc()
```

```
##             used   (Mb) gc trigger    (Mb)   max used    (Mb)
## Ncells   5088213  271.8   13651909   729.1   21331105  1139.3
## Vcells 857323722 6540.9 2342481578 17871.8 4574774951 34902.8
```

```
chick <- JoinLayers(chick)
chick.markers <- FindAllMarkers(chick, min.pct = 0.5, logfc.threshold = 0.5)
```

```
## Calculating cluster 0
```

```
## Calculating cluster 1
```

```
## Calculating cluster 2
```

```
## Calculating cluster 3
```

```
## Calculating cluster 4
```

```
## Calculating cluster 5
```

```
## Calculating cluster 6
```

```
## Calculating cluster 7
```

```
## Calculating cluster 8
```

```
## Calculating cluster 9
```

```
## Calculating cluster 10
```

```
chick.markers %>%
  group_by(cluster) %>%
  top_n(n = 5, wt = avg_log2FC) -> top5
DotPlot(chick, features = unique(top5$gene))+coord_flip()
```

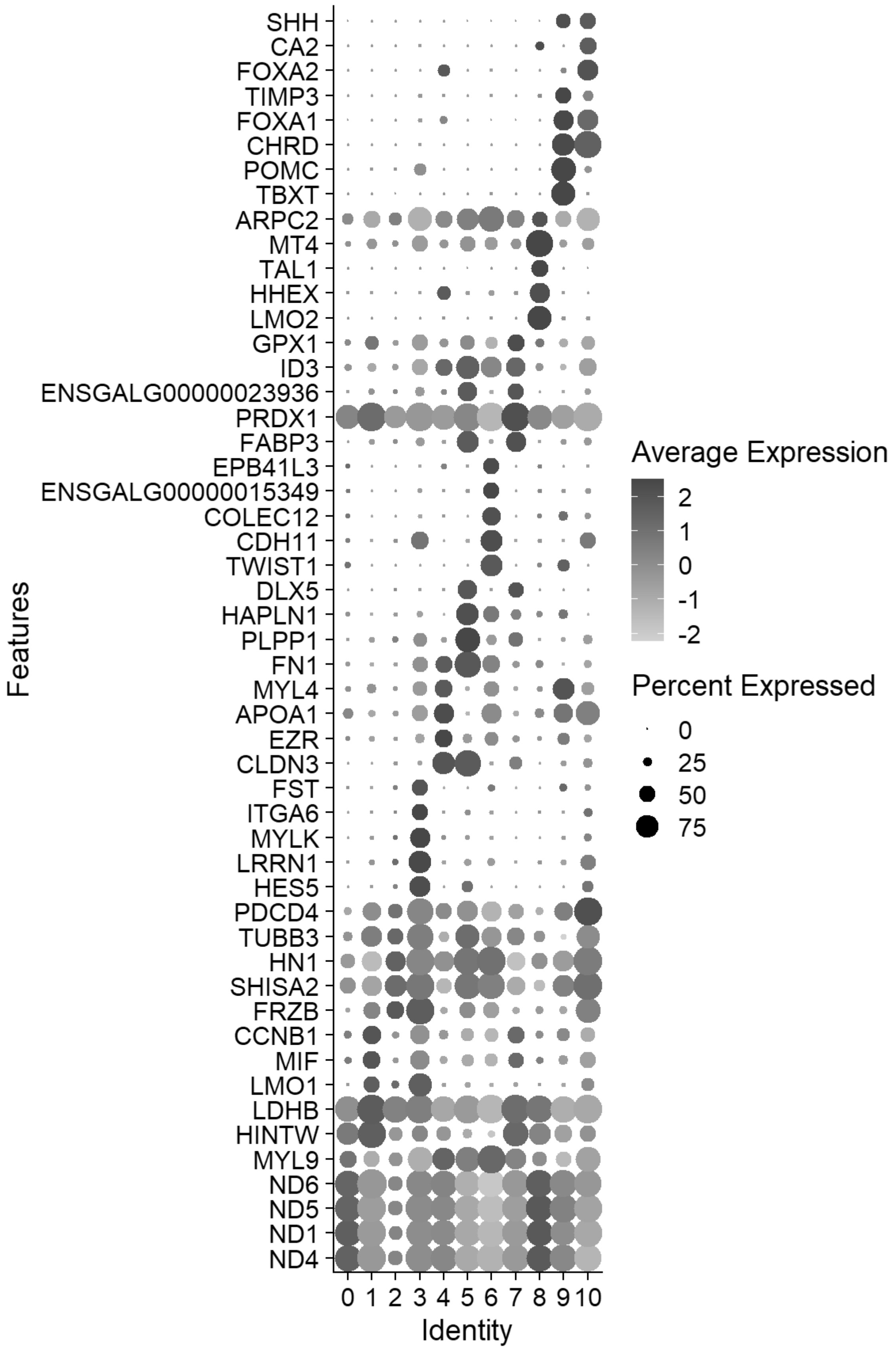


```
gc()
```

```
##              used   (Mb) gc trigger    (Mb)   max used    (Mb)
## Ncells   5048755  269.7   13651909   729.1   21331105  1139.3
```

```
## Vcells 970378524 7403.5 2342481578 17871.8 4574774951 34902.8
```

```
cluster_map <- c("0"="Mesoderm 1",
                 "1"="Undecided Ectoderm 1",
                 "2"="Undecided Ectoderm 2",
                 "3"="Neural",
                 "4"="Endoderm",
                 "5"="NNE/Placode 1",
                 "6"="Mesoderm 2",
                 "7"="NNE/Placode 2",
                 "8"="Hematogenic",
                 "9"="Notochord",
                 "10"="vNeural Tube")
chick$celltype1 <- plyr::mapvalues(chick$seurat_clusters, from = names(cluster_map), to = clus
ter_map)

chick <- JoinLayers(chick)
Idents(chick) <- "celltype1" # Setting active identity ensures markers are calculated by cell
type
global_markers <- FindAllMarkers(chick, min.pct = 0.5, logfc.threshold = 0.5)
```

```
## Calculating cluster Mesoderm 1
```

```
## Calculating cluster Undecided Ectoderm 1
```

```
## Calculating cluster Undecided Ectoderm 2
```

```
## Calculating cluster Neural
```

```
## Calculating cluster Endoderm
```

```
## Calculating cluster NNE/Placode 1
```

```
## Calculating cluster Mesoderm 2
```

```
## Calculating cluster NNE/Placode 2
```

```
## Calculating cluster Hematogenic
```

```
## Calculating cluster Notochord
```

```
## Calculating cluster vNeural Tube
```

```
write.csv(global_markers, "E:/Rogers Lab/Rogers NC EMT/Supplementary file 4_HH8_Markers_Annota
ted.csv", row.names = FALSE)
```

```
celltype_colors <- c("Mesoderm 1"="#FC9272",
                     "Undecided Ectoderm 1"="#BFD3E6",
                   "Undecided Ectoderm 2"="#9EBCDA",
                 "Neural"="#1D91C0",
                 "Endoderm"="#FED976",
                 "NNE/Placode 1"="#A1D99B",
                 "Mesoderm 2"="#CB181D",
                 "NNE/Placode 2"="#C7E9C0",
                 "Hematogenic"="#BD0026",
                 "Notochord"="#DEEBF7",
                 "vNeural Tube"="#08519C")

p_umap <- DimPlot(chick, group.by = "celltype1",cols = celltype_colors, label = TRUE, repel =
TRUE) + ggtitle(NULL)+
            NoLegend()
p_umap
```

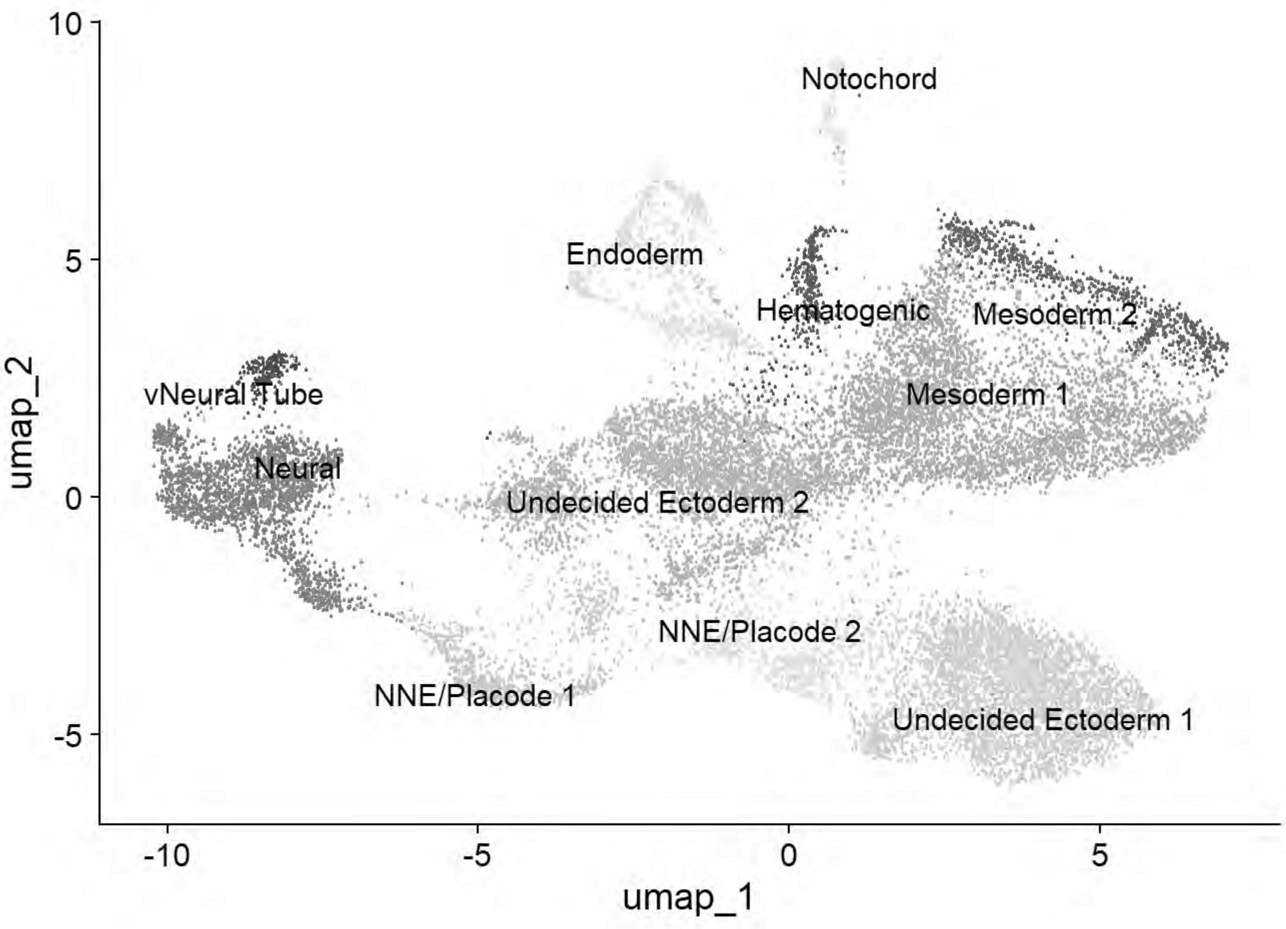


```
ggsave("E:/Rogers Lab/Rogers NC EMT/Figures/Fig_HH8_Summary_UMAP1.png", plot = p_umap, width =
 6, height = 6, dpi = 600)


cptbl <- table(chick@meta.data$celltype1,chick@meta.data$stage)
cptbl <- cbind(cptbl, TOTAL = rowSums(cptbl))
colnames(cptbl)[length(cptbl[1,])] <- "TOTAL"
```

```
tg = gridExtra::tableGrob(cptbl)
h = grid::convertHeight(sum(tg$heights), "in", TRUE)
w = grid::convertWidth(sum(tg$widths), "in", TRUE)
ggsave("E:/Rogers Lab/Rogers NC EMT/Figures/Fig_HH8_Summary_Table.png", tg, width=w, height=h,
 device = 'png', dpi =600)

top5 <- global_markers %>% group_by(cluster) %>% top_n(n = 5, wt = avg_log2FC)
p_dot=DotPlot(chick, features = unique(top5$gene), group.by = "celltype1") + # cluster.idents
dendrograms the y-axis
  coord_flip() +
  scale_size_continuous(range = c(1, 10)) +
  scale_color_gradientn(colors = c("grey90", "#313695", "#d73027"))+
  theme_light() +
  theme(
    axis.text.x = element_text(angle = 45, hjust = 1, color = "black", size = 12),
    axis.text.y = element_text(face = "italic", color = "black", size = 12),
    axis.title.x = element_blank(),
    axis.title.y = element_blank(),
    panel.grid.major = element_line(color = "grey90")
  )
```

```
## Scale for size is already present.
## Adding another scale for size, which will replace the existing scale.
```

```
## Scale for colour is already present.
## Adding another scale for colour, which will replace the existing scale.
```

```
p_dot
```

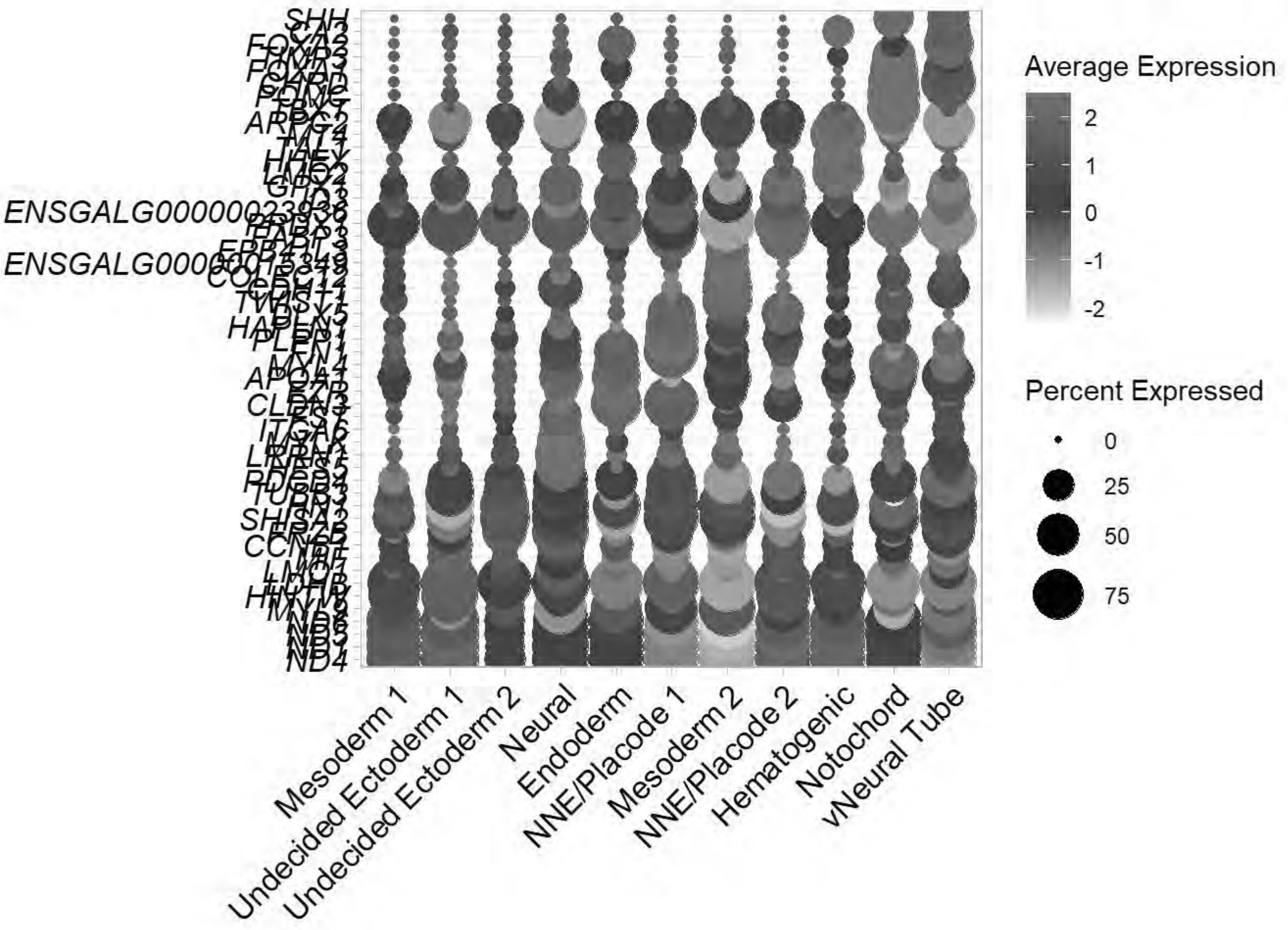


```
ggsave("E:/Rogers Lab/Rogers NC EMT/Figures/Fig_HH8_Summary_Dotplot.png", plot = p_dot, width
= 7, height = 10, dpi = 600)
```

```
saveRDS(chick, "E:\\Rogers Lab\\Rogers NC EMT\\HH8_chick_v2.rds")

Idents(chick)<-"celltype1"
chick=subset(chick, idents =c("Undecided Ectoderm 1","Undecided Ectoderm 2","Neural","NNE/Plac
ode 1","NNE/Placode 2","vNeural Tube"))
chick
```

```
## An object of class Seurat
## 18523 features across 11869 samples within 1 assay
## Active assay: RNA (18523 features, 2000 variable features)
##  3 layers present: data, counts, scale.data
##  3 dimensional reductions calculated: pca, harmony, umap
```

```
saveRDS(chick,"E:\\Rogers Lab\\Rogers NC EMT\\HH8_chick_NC_v2.rds")
gc()
```

```
##              used   (Mb) gc trigger    (Mb)   max used    (Mb)
## Ncells    4923647  263.0   13651909   729.1   21331105  1139.3
## Vcells 1042071010 7950.4 2342481578 17871.8 4574774951 34902.8
```

### Stage Analysis: HH9

```
chick=readRDS("E:\\Rogers Lab\\Rogers NC EMT\\Global_chick_v2.rds")
chick
```

```
## An object of class Seurat
## 18523 features across 82092 samples within 1 assay
## Active assay: RNA (18523 features, 2000 variable features)
##  3 layers present: data, counts, scale.data
##  3 dimensional reductions calculated: pca, harmony, umap
```

```
Idents(chick) <- "stage"
chick=subset(chick, idents =c("HH9"))
chick
```

```
## An object of class Seurat
## 18523 features across 18030 samples within 1 assay
## Active assay: RNA (18523 features, 2000 variable features)
##  3 layers present: data, counts, scale.data
##  3 dimensional reductions calculated: pca, harmony, umap
```

```
gc()
```

```
##              used   (Mb) gc trigger    (Mb)   max used    (Mb)
## Ncells    4923926  263.0   13651909   729.1   21331105  1139.3
## Vcells 1076741337 8214.9 2342481578 17871.8 4574774951 34902.8
```

```
chick <- NormalizeData(chick)
```

```
## Normalizing layer: counts
```

```
chick <- CellCycleScoring(chick, s.features = cc.genes$s.genes, g2m.features = cc.genes$g2m.ge
nes, set.ident = TRUE)
chick <- FindVariableFeatures(chick)
```

```
## Finding variable features for layer counts
```

```
chick <- ScaleData(chick, vars.to.regress = c("percent.mt","G2M.Score","S.Score"))
```

```
## Regressing out percent.mt, G2M.Score, S.Score
```

```
## Centering and scaling data matrix
```

```
chick <- RunPCA(chick, npcs = 50, verbose = FALSE)
```

```
ElbowPlot(chick, ndims= 50, reduction = "pca")
```

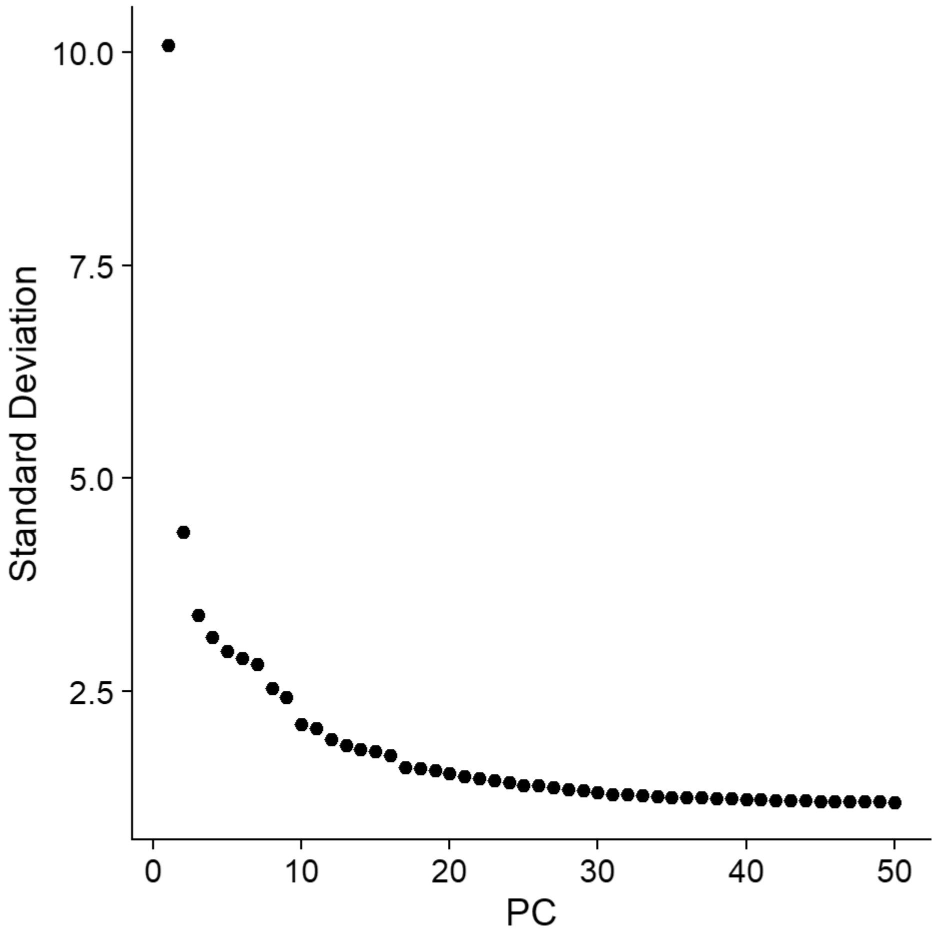


```
gc()
```

```
##              used   (Mb) gc trigger    (Mb)   max used    (Mb)
## Ncells    4945202  264.2   13651909   729.1   21331105  1139.3
## Vcells 1076886854 8216.0 2342481578 17871.8 4574774951 34902.8
```

```
chick[["RNA"]] <- split(chick[["RNA"]], f = chick$sample_id)
```

```
## Splitting 'counts', 'data' layers. Not splitting 'scale.data'. If you would like to split o
ther layers, set in `layers` argument.
```

```
chick <- IntegrateLayers(object=chick, method=HarmonyIntegration, orig.reduction="pca",
                       new.reduction="harmony", assay="RNA", group.by="stage", verbose=TRUE)
```

```
## Transposing data matrix
```

```
## Using automatic lambda estimation
```

```
## Initializing state using k-means centroids initialization
```

```
## Harmony 1/10
```

```
## Harmony 2/10
```

```
## Harmony 3/10
```

```
## Harmony converged after 3 iterations
```

```
chick <- FindNeighbors(chick, reduction="harmony", dims=1:30, nn.method="rann", k.param=20)
```

```
## Computing nearest neighbor graph
```

```
## Computing SNN
```

```
chick <- RunUMAP(chick, reduction="harmony", dims=1:30)
```

```
## 23:16:51 UMAP embedding parameters a = 0.9922 b = 1.112
```

```
## 23:16:51 Read 18030 rows and found 30 numeric columns
```

```
## 23:16:51 Using Annoy for neighbor search, n_neighbors = 30
```

```
## 23:16:51 Building Annoy index with metric = cosine, n_trees = 50
```

```
## 0%   10   20   30   40   50   60   70   80   90   100%
```

```
## [----|----|----|----|----|----|----|----|----|----|
```

```
## **************************************************|
## 23:16:52 Writing NN index file to temp file C:\Users\RANEES~1\AppData\Local\Temp\RtmpqcWgVb
\file6a2c15237ada
## 23:16:52 Searching Annoy index using 1 thread, search_k = 3000
## 23:16:56 Annoy recall = 100%
## 23:16:57 Commencing smooth kNN distance calibration using 1 thread with target n_neighbors
= 30
## 23:16:58 Initializing from normalized Laplacian + noise (using RSpectra)
## 23:16:58 Commencing optimization for 200 epochs, with 828650 positive edges
## 23:16:58 Using rng type: pcg
## 23:17:14 Optimization finished
```

```
chick <- FindClusters(chick, resolution=0.4)
```

```
## Modularity Optimizer version 1.3.0 by Ludo Waltman and Nees Jan van Eck
##
## Number of nodes: 18030
## Number of edges: 654017
##
## Running Louvain algorithm...
## Maximum modularity in 10 random starts: 0.9006
## Number of communities: 15
## Elapsed time: 2 seconds
```

```
DimPlot(chick)
```

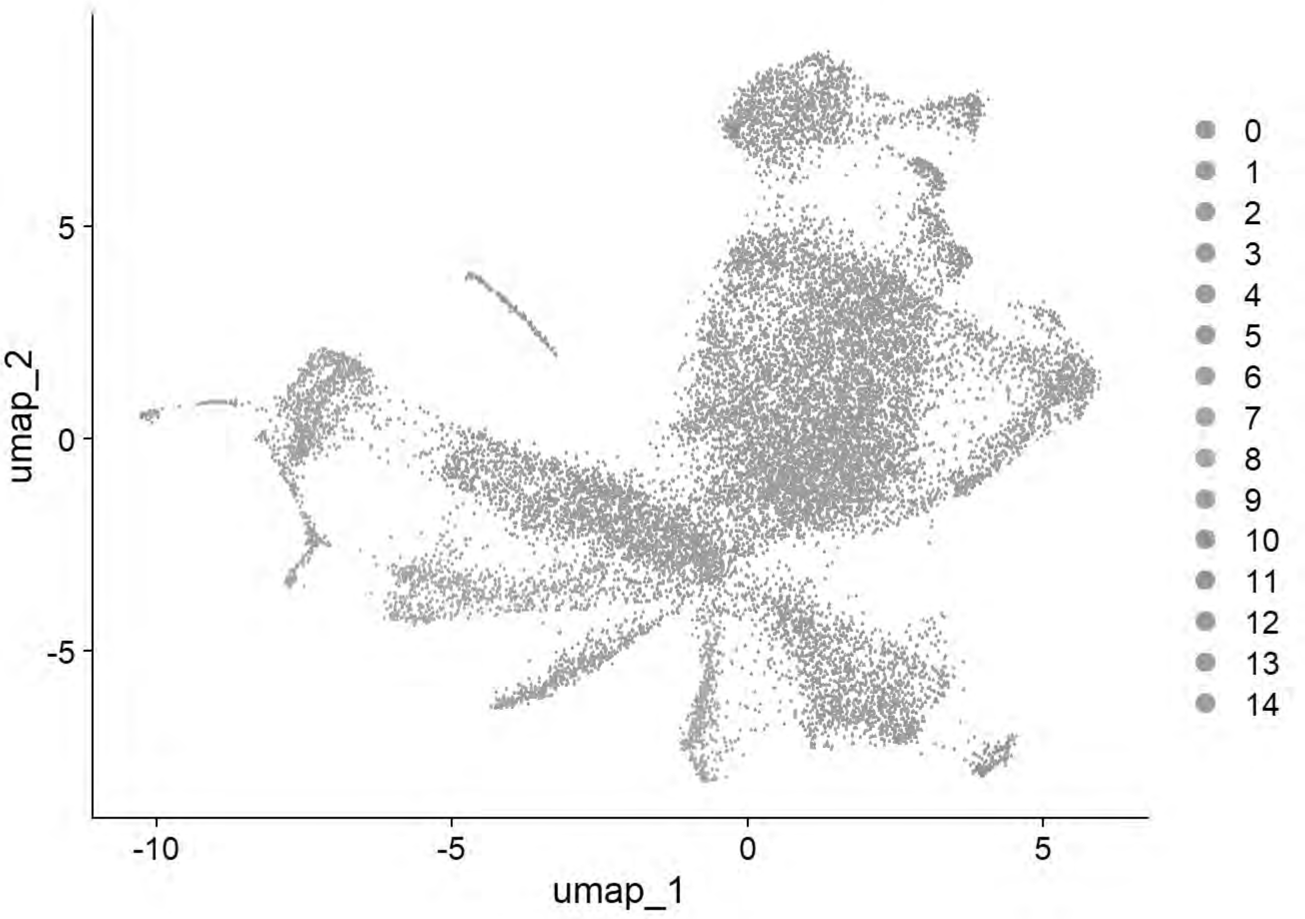


```
DimPlot(chick, group.by = "sample_id")
```

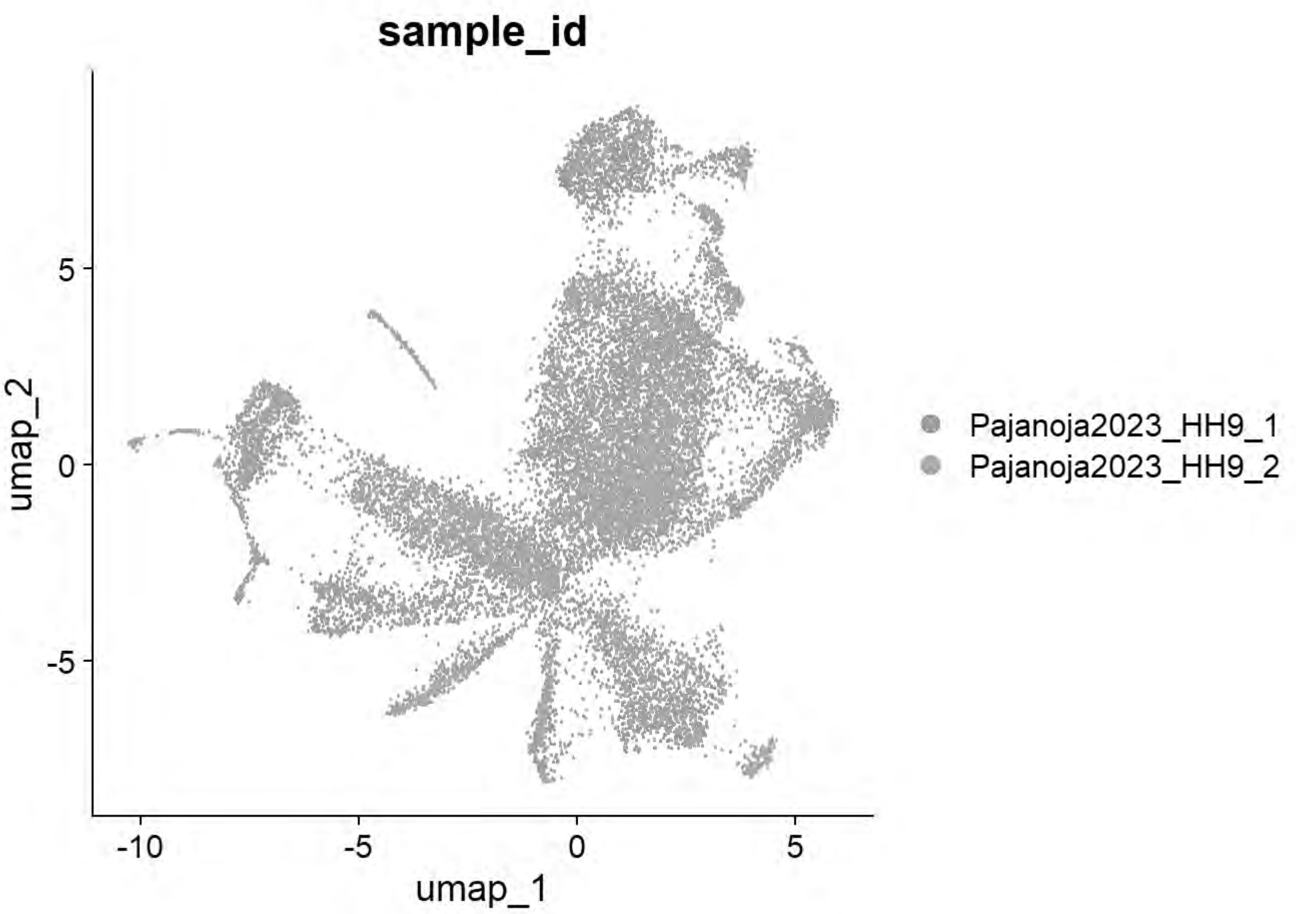


```
chick = FindClusters(chick, resolution= seq(0,2,0.1),n.start=10)
```

```
## Modularity Optimizer version 1.3.0 by Ludo Waltman and Nees Jan van Eck
##
## Number of nodes: 18030
## Number of edges: 654017
##
## Running Louvain algorithm...
## Maximum modularity in 10 random starts: 1.0000
## Number of communities: 1
## Elapsed time: 2 seconds
## Modularity Optimizer version 1.3.0 by Ludo Waltman and Nees Jan van Eck
##
## Number of nodes: 18030
## Number of edges: 654017
##
## Running Louvain algorithm...
## Maximum modularity in 10 random starts: 0.9576
## Number of communities: 6
## Elapsed time: 2 seconds
## Modularity Optimizer version 1.3.0 by Ludo Waltman and Nees Jan van Eck
##
## Number of nodes: 18030
## Number of edges: 654017
##
```

```
## Running Louvain algorithm...
## Maximum modularity in 10 random starts: 0.9349
## Number of communities: 11
## Elapsed time: 2 seconds
## Modularity Optimizer version 1.3.0 by Ludo Waltman and Nees Jan van Eck
##
## Number of nodes: 18030
## Number of edges: 654017
##
## Running Louvain algorithm...
## Maximum modularity in 10 random starts: 0.9162
## Number of communities: 14
## Elapsed time: 2 seconds
## Modularity Optimizer version 1.3.0 by Ludo Waltman and Nees Jan van Eck
##
## Number of nodes: 18030
## Number of edges: 654017
##
## Running Louvain algorithm...
## Maximum modularity in 10 random starts: 0.9006
## Number of communities: 15
## Elapsed time: 2 seconds
## Modularity Optimizer version 1.3.0 by Ludo Waltman and Nees Jan van Eck
##
## Number of nodes: 18030
## Number of edges: 654017
##
## Running Louvain algorithm...
## Maximum modularity in 10 random starts: 0.8879
## Number of communities: 17
## Elapsed time: 2 seconds
## Modularity Optimizer version 1.3.0 by Ludo Waltman and Nees Jan van Eck
##
## Number of nodes: 18030
## Number of edges: 654017
##
## Running Louvain algorithm...
## Maximum modularity in 10 random starts: 0.8776
## Number of communities: 18
## Elapsed time: 2 seconds
## Modularity Optimizer version 1.3.0 by Ludo Waltman and Nees Jan van Eck
##
## Number of nodes: 18030
## Number of edges: 654017
##
## Running Louvain algorithm...
## Maximum modularity in 10 random starts: 0.8670
## Number of communities: 18
## Elapsed time: 2 seconds
## Modularity Optimizer version 1.3.0 by Ludo Waltman and Nees Jan van Eck
##
## Number of nodes: 18030
## Number of edges: 654017
##
```

```
## Running Louvain algorithm...
## Maximum modularity in 10 random starts: 0.8567
## Number of communities: 19
## Elapsed time: 2 seconds
## Modularity Optimizer version 1.3.0 by Ludo Waltman and Nees Jan van Eck
##
## Number of nodes: 18030
## Number of edges: 654017
##
## Running Louvain algorithm...
## Maximum modularity in 10 random starts: 0.8488
## Number of communities: 20
## Elapsed time: 2 seconds
## Modularity Optimizer version 1.3.0 by Ludo Waltman and Nees Jan van Eck
##
## Number of nodes: 18030
## Number of edges: 654017
##
## Running Louvain algorithm...
## Maximum modularity in 10 random starts: 0.8406
## Number of communities: 20
## Elapsed time: 2 seconds
## Modularity Optimizer version 1.3.0 by Ludo Waltman and Nees Jan van Eck
##
## Number of nodes: 18030
## Number of edges: 654017
##
## Running Louvain algorithm...
## Maximum modularity in 10 random starts: 0.8323
## Number of communities: 20
## Elapsed time: 2 seconds
## Modularity Optimizer version 1.3.0 by Ludo Waltman and Nees Jan van Eck
##
## Number of nodes: 18030
## Number of edges: 654017
##
## Running Louvain algorithm...
## Maximum modularity in 10 random starts: 0.8247
## Number of communities: 22
## Elapsed time: 2 seconds
## Modularity Optimizer version 1.3.0 by Ludo Waltman and Nees Jan van Eck
##
## Number of nodes: 18030
## Number of edges: 654017
##
## Running Louvain algorithm...
## Maximum modularity in 10 random starts: 0.8173
## Number of communities: 22
## Elapsed time: 2 seconds
## Modularity Optimizer version 1.3.0 by Ludo Waltman and Nees Jan van Eck
##
## Number of nodes: 18030
## Number of edges: 654017
##
```

```
## Running Louvain algorithm...
## Maximum modularity in 10 random starts: 0.8099
## Number of communities: 22
## Elapsed time: 2 seconds
## Modularity Optimizer version 1.3.0 by Ludo Waltman and Nees Jan van Eck
##
## Number of nodes: 18030
## Number of edges: 654017
##
## Running Louvain algorithm...
## Maximum modularity in 10 random starts: 0.8043
## Number of communities: 24
## Elapsed time: 2 seconds
## Modularity Optimizer version 1.3.0 by Ludo Waltman and Nees Jan van Eck
##
## Number of nodes: 18030
## Number of edges: 654017
##
## Running Louvain algorithm...
## Maximum modularity in 10 random starts: 0.7983
## Number of communities: 24
## Elapsed time: 2 seconds
## Modularity Optimizer version 1.3.0 by Ludo Waltman and Nees Jan van Eck
##
## Number of nodes: 18030
## Number of edges: 654017
##
## Running Louvain algorithm...
## Maximum modularity in 10 random starts: 0.7915
## Number of communities: 26
## Elapsed time: 2 seconds
## Modularity Optimizer version 1.3.0 by Ludo Waltman and Nees Jan van Eck
##
## Number of nodes: 18030
## Number of edges: 654017
##
## Running Louvain algorithm...
## Maximum modularity in 10 random starts: 0.7856
## Number of communities: 26
## Elapsed time: 2 seconds
## Modularity Optimizer version 1.3.0 by Ludo Waltman and Nees Jan van Eck
##
## Number of nodes: 18030
## Number of edges: 654017
##
## Running Louvain algorithm...
## Maximum modularity in 10 random starts: 0.7809
## Number of communities: 27
## Elapsed time: 2 seconds
## Modularity Optimizer version 1.3.0 by Ludo Waltman and Nees Jan van Eck
##
## Number of nodes: 18030
## Number of edges: 654017
##
```

```
## Running Louvain algorithm...
## Maximum modularity in 10 random starts: 0.7754
## Number of communities: 27
## Elapsed time: 2 seconds
```

```
p=clustree(chick, prefix= "RNA_snn_res.")
print(p)
```

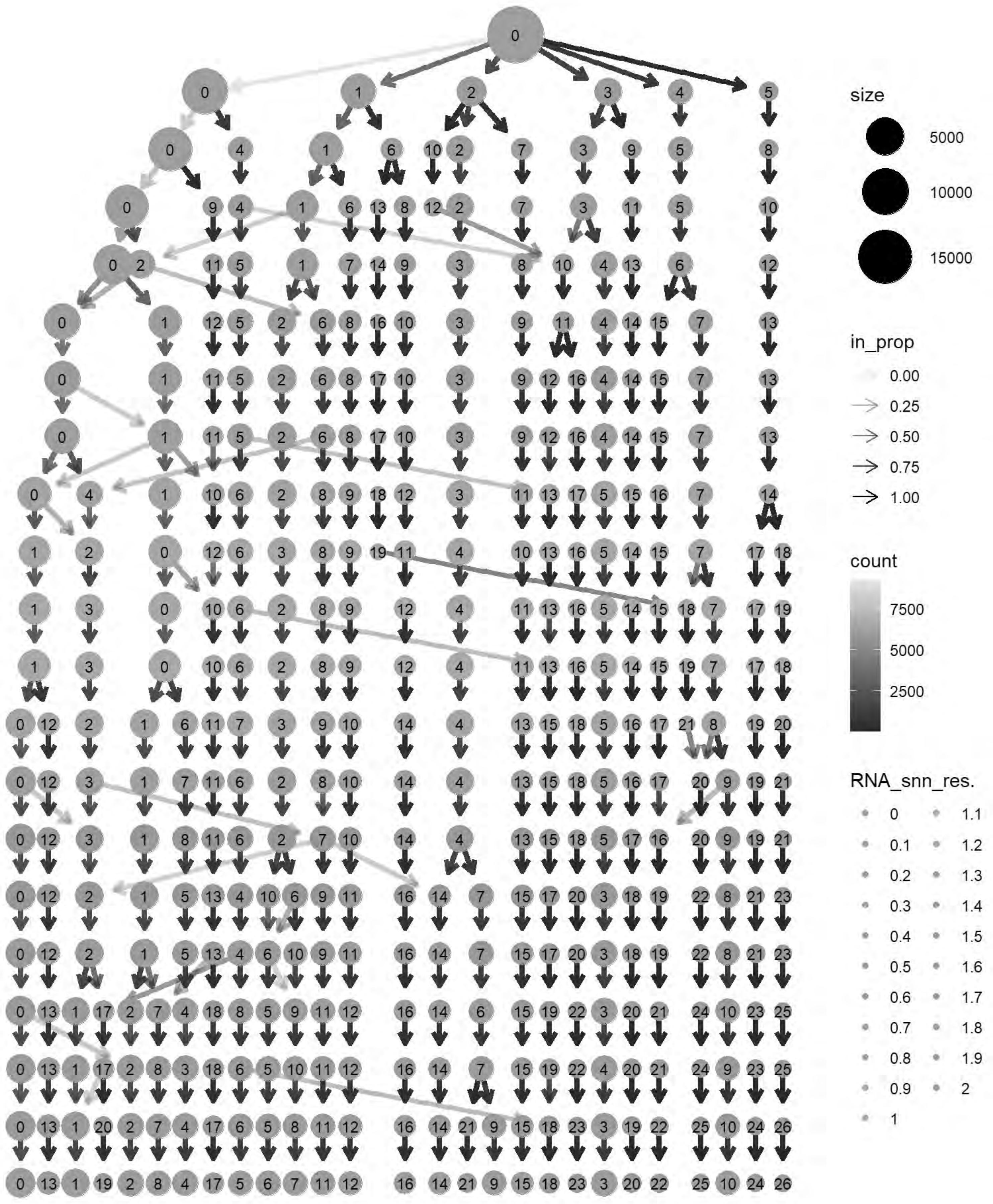
size
5000
10000
15000
in_prop
0.00
0.25
0.50
0.75
1.00
count
7500
5000
2500
RNA_snn_res.
0
0.1
0.2
0.3
0.4
0.5
0.6
0.7
0.8
0.9
1
1.1
1.2
1.3
1.4
1.5
1.6
1.7
1.8
1.9
2

```
cols= grep("^RNA_snn_res.",names(chick@meta.data))
chick@meta.data[cols] <- NULL
chick <- FindClusters(chick, resolution = 0.5, verbose = TRUE)
```

```
## Modularity Optimizer version 1.3.0 by Ludo Waltman and Nees Jan van Eck
##
## Number of nodes: 18030
## Number of edges: 654017
##
## Running Louvain algorithm...
## Maximum modularity in 10 random starts: 0.8879
## Number of communities: 17
## Elapsed time: 2 seconds
```

```
p6=DimPlot(chick, label = TRUE)+NoLegend()
print(p6)
```

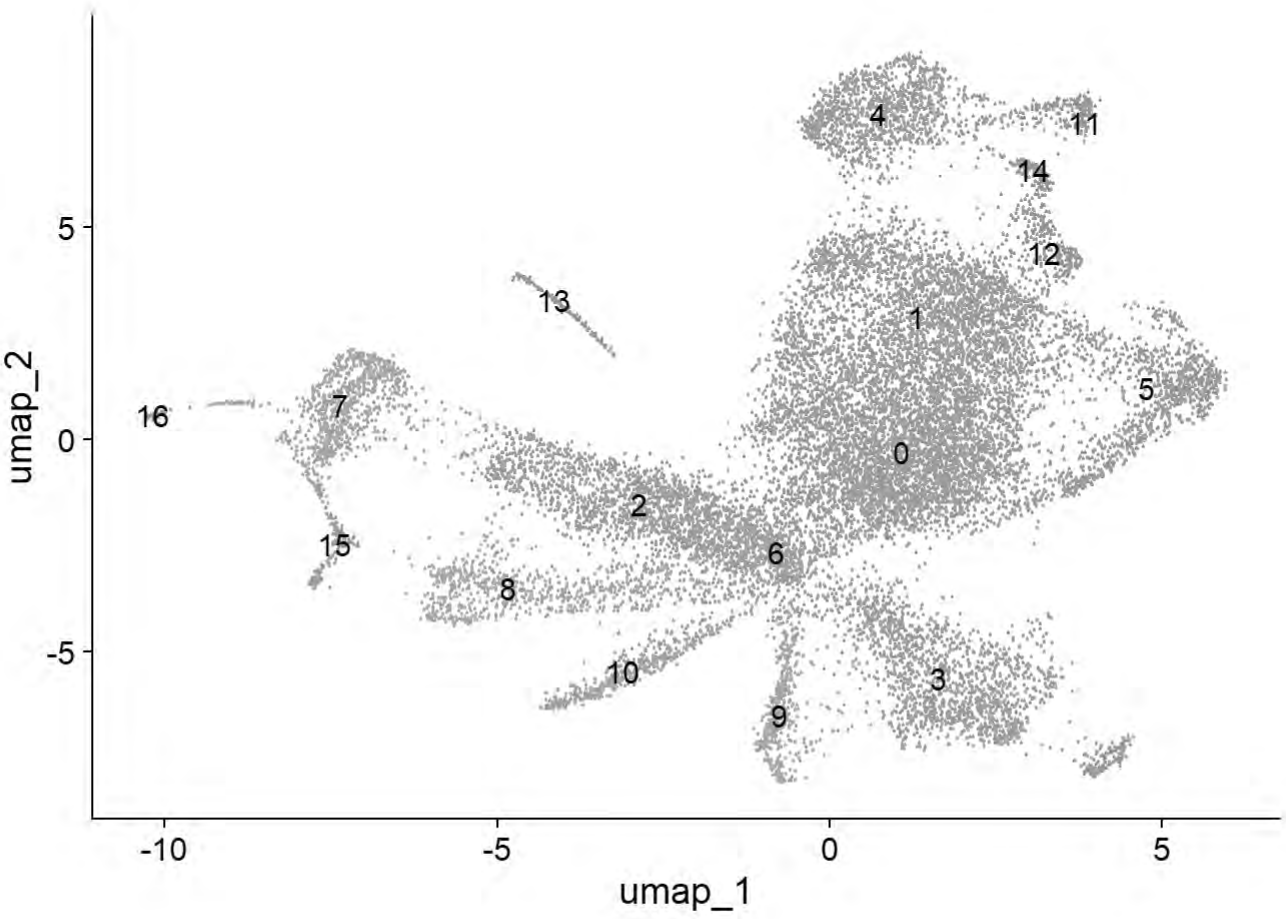


```
p6=DimPlot(chick, group.by = "Phase")
print(p6)
```

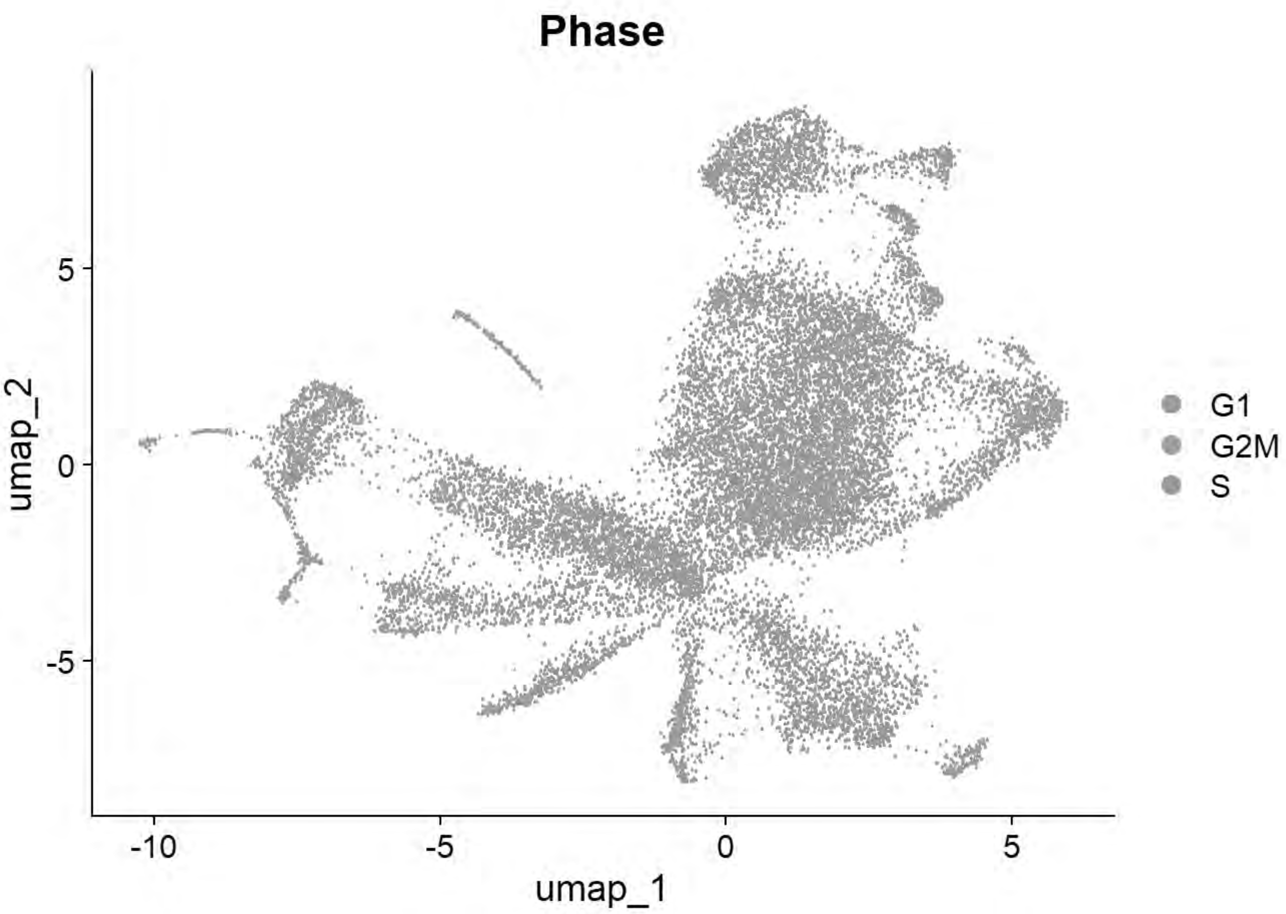


```
p6=DimPlot(chick, group.by = "sample_id")
print(p6)
```

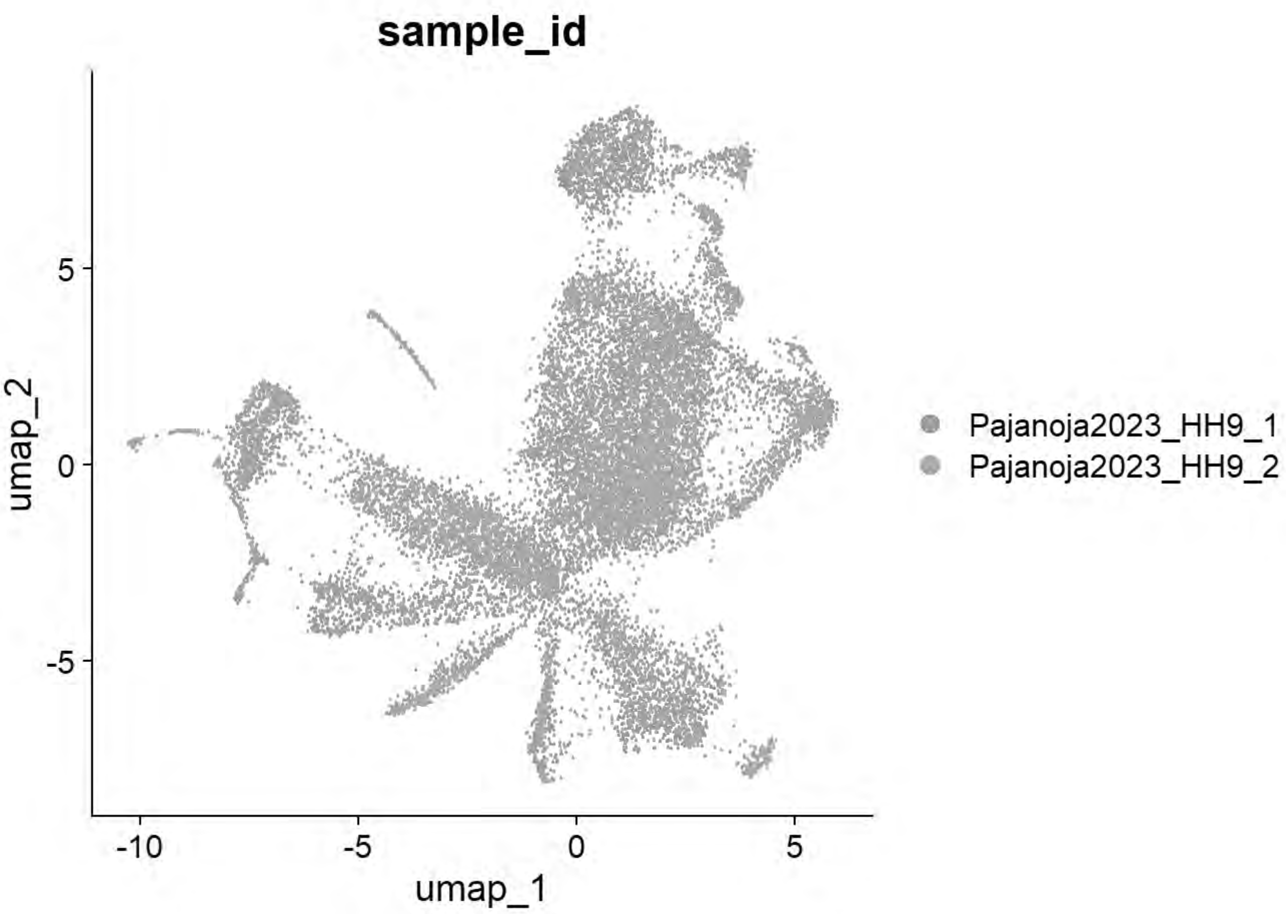


```
# Williams: Neural plate/tube
FeaturePlot(chick, features = c("CLDN1","FRZB","DNMT3A"))
```

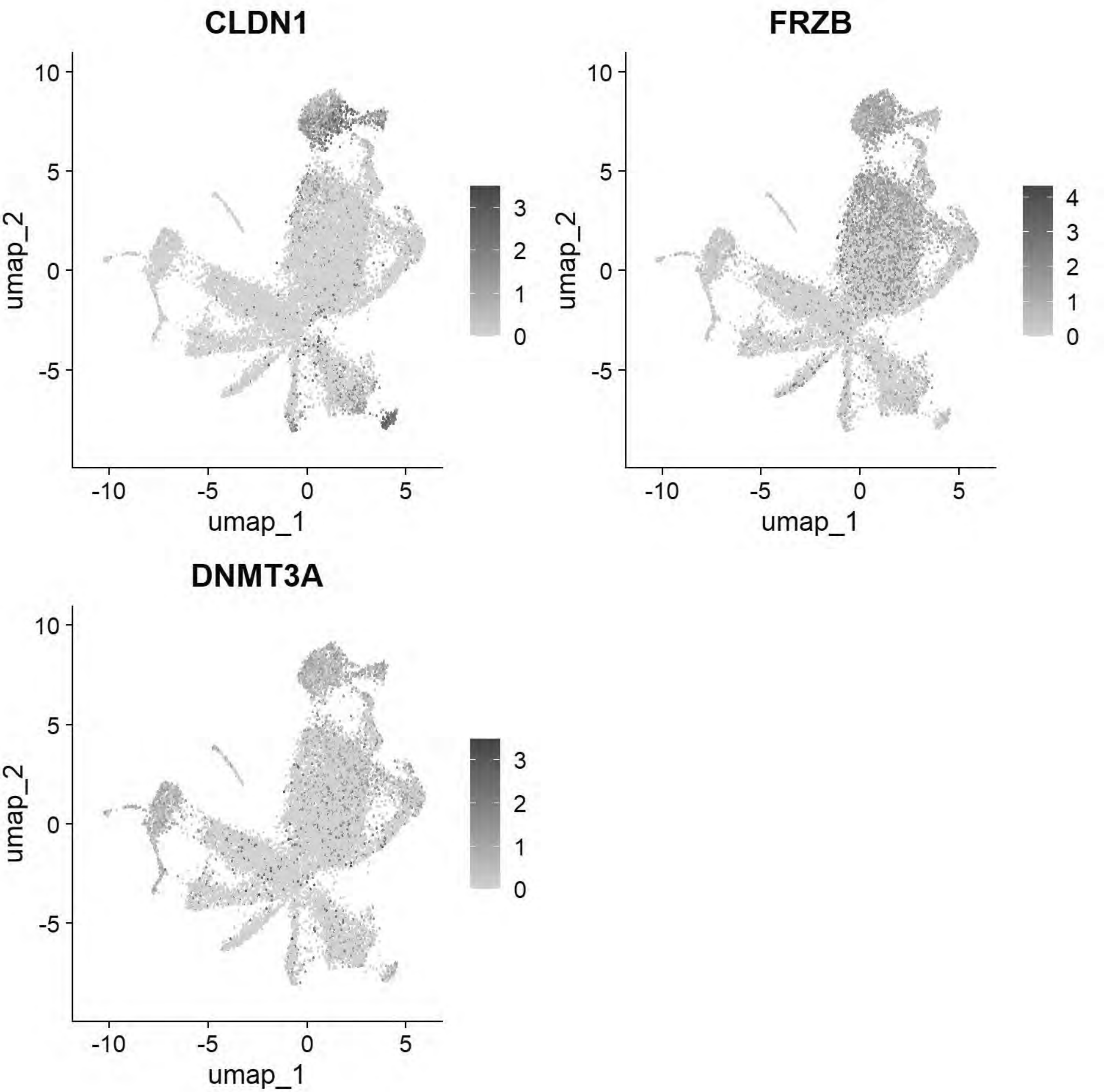


```
# Williams: Non-Neural Ectoderm
FeaturePlot(chick, features = c("DLX5","TFAP2A","ASTL","PAX6"))
```

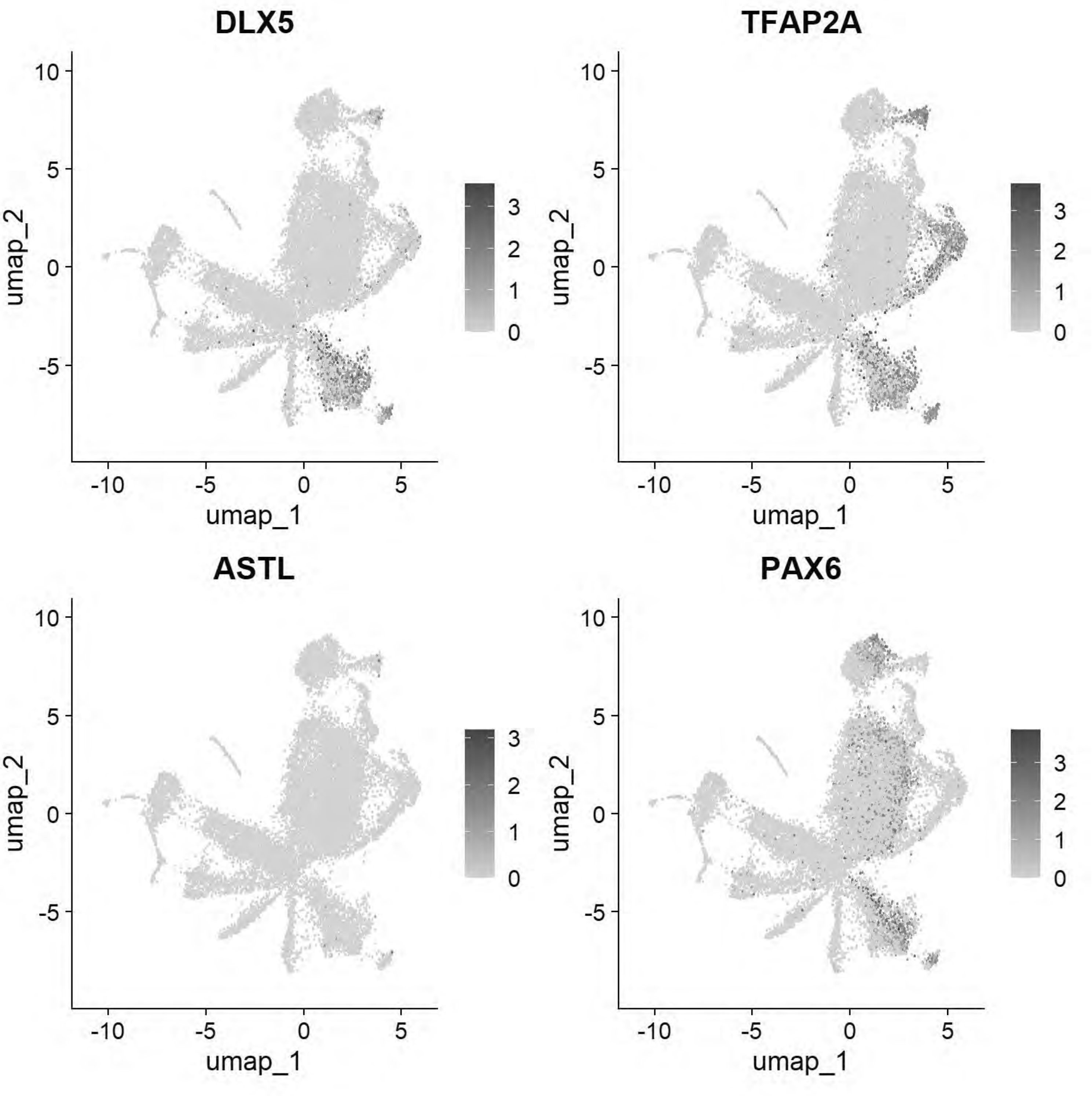


```r
# Williams: Ectoderm (Neural Plate)
FeaturePlot(chick, features = c("SOX2","SOX3","SOX21","FRZB","SFRP2"))
```

```
## Warning: All cells have the same value (0) of "SOX3"
```

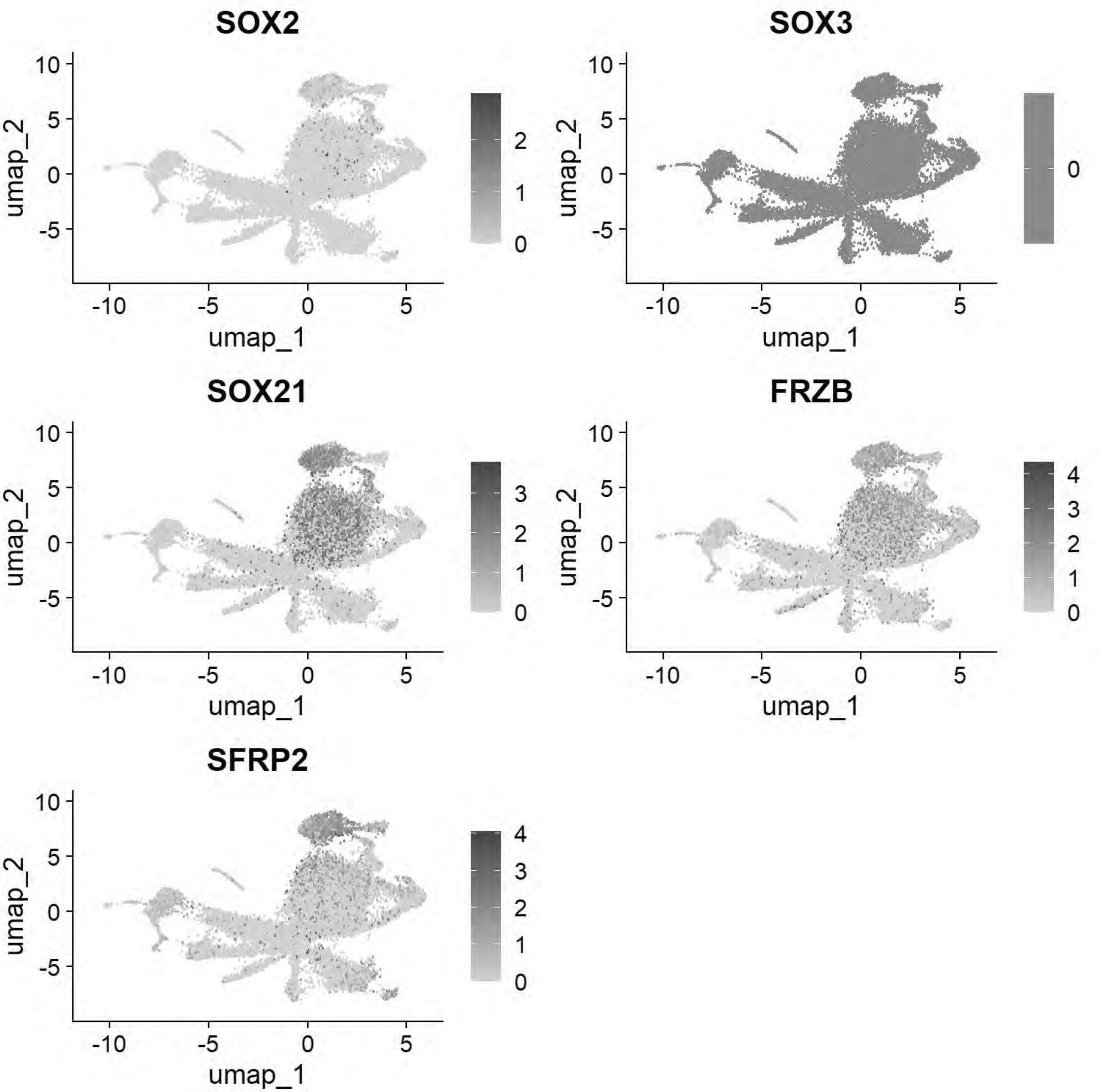


```
# Williams: Ectoderm (Neural Plate Border)
FeaturePlot(chick, features = c("PAX7","TFAP2A","DLX5","BMP4","MSX1","DRAXIN","TFAP2B"))
```

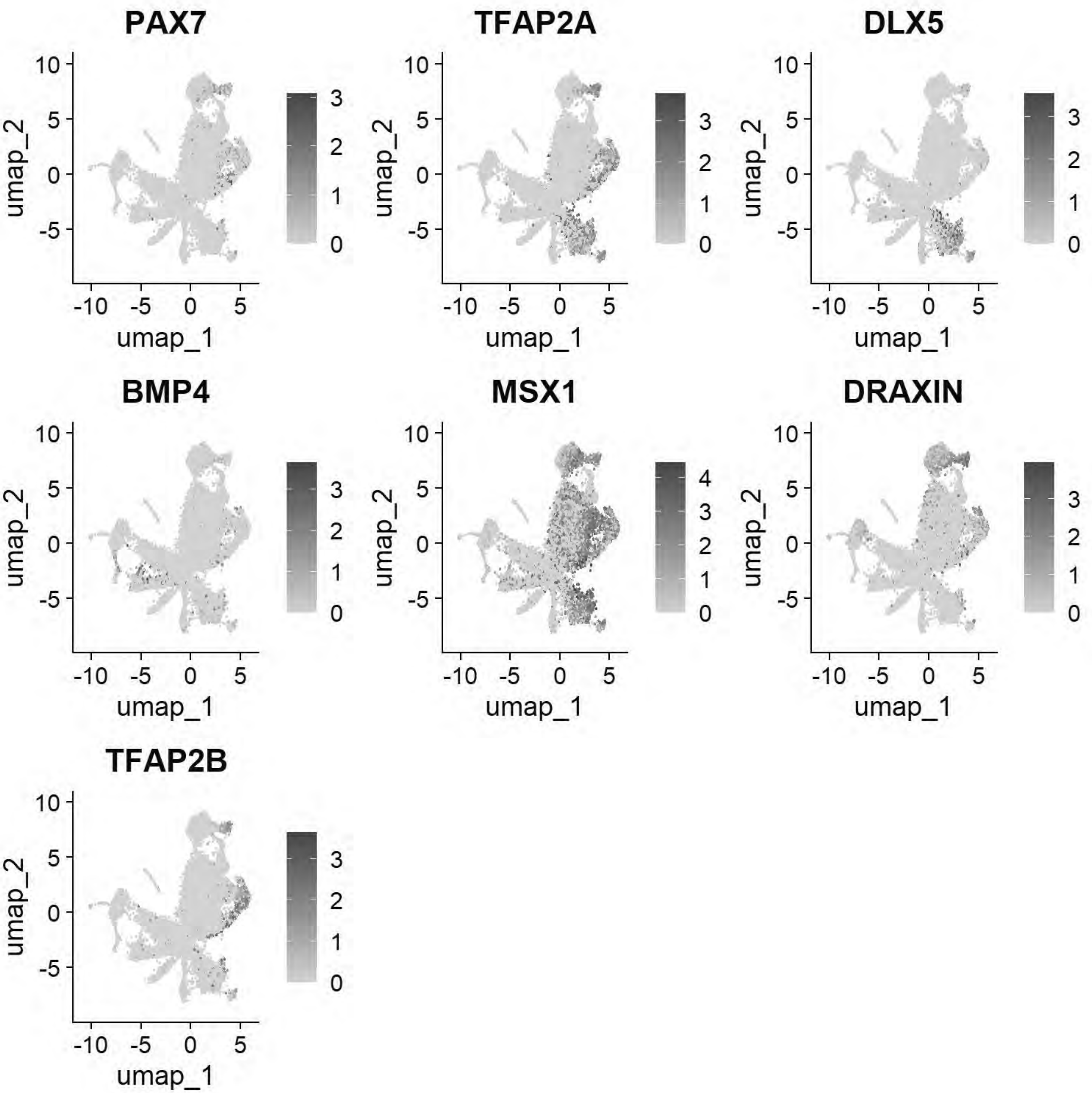


```
# Williams: Posterior Lateral Plate Mesoderm
FeaturePlot(chick, features = c("GATA2","HOXB5","CDX4"))
```

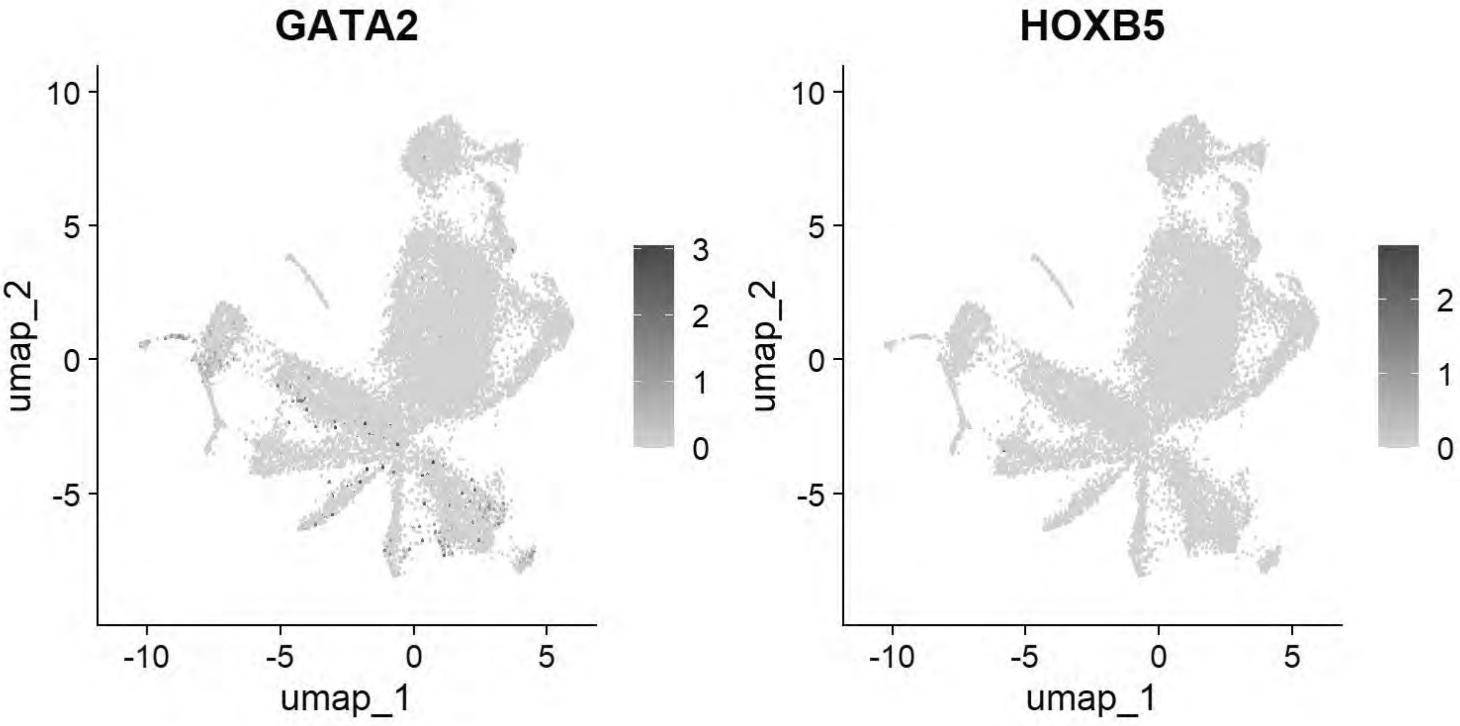


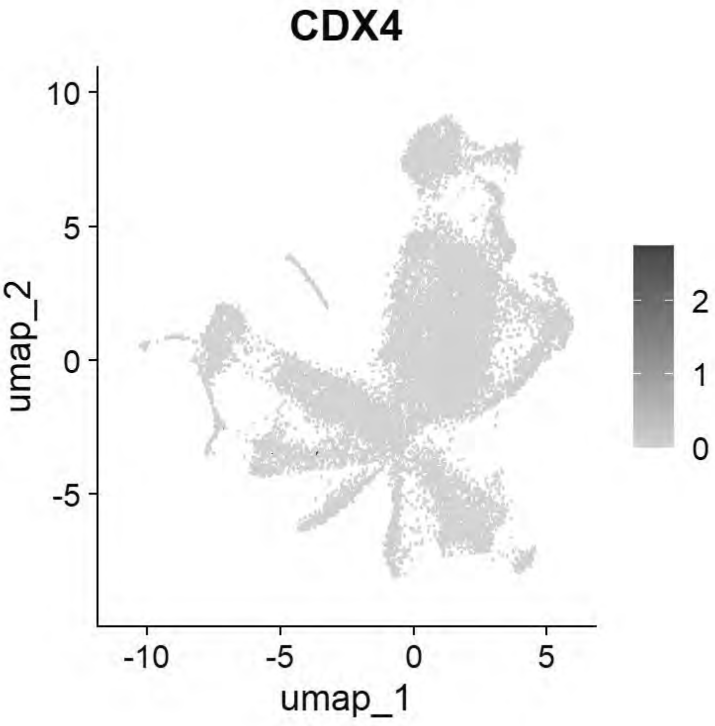


```
# Williams: Paraxial Mesoderm
FeaturePlot(chick, features = c("MSGN1","MESP1","MEOX1"))
```

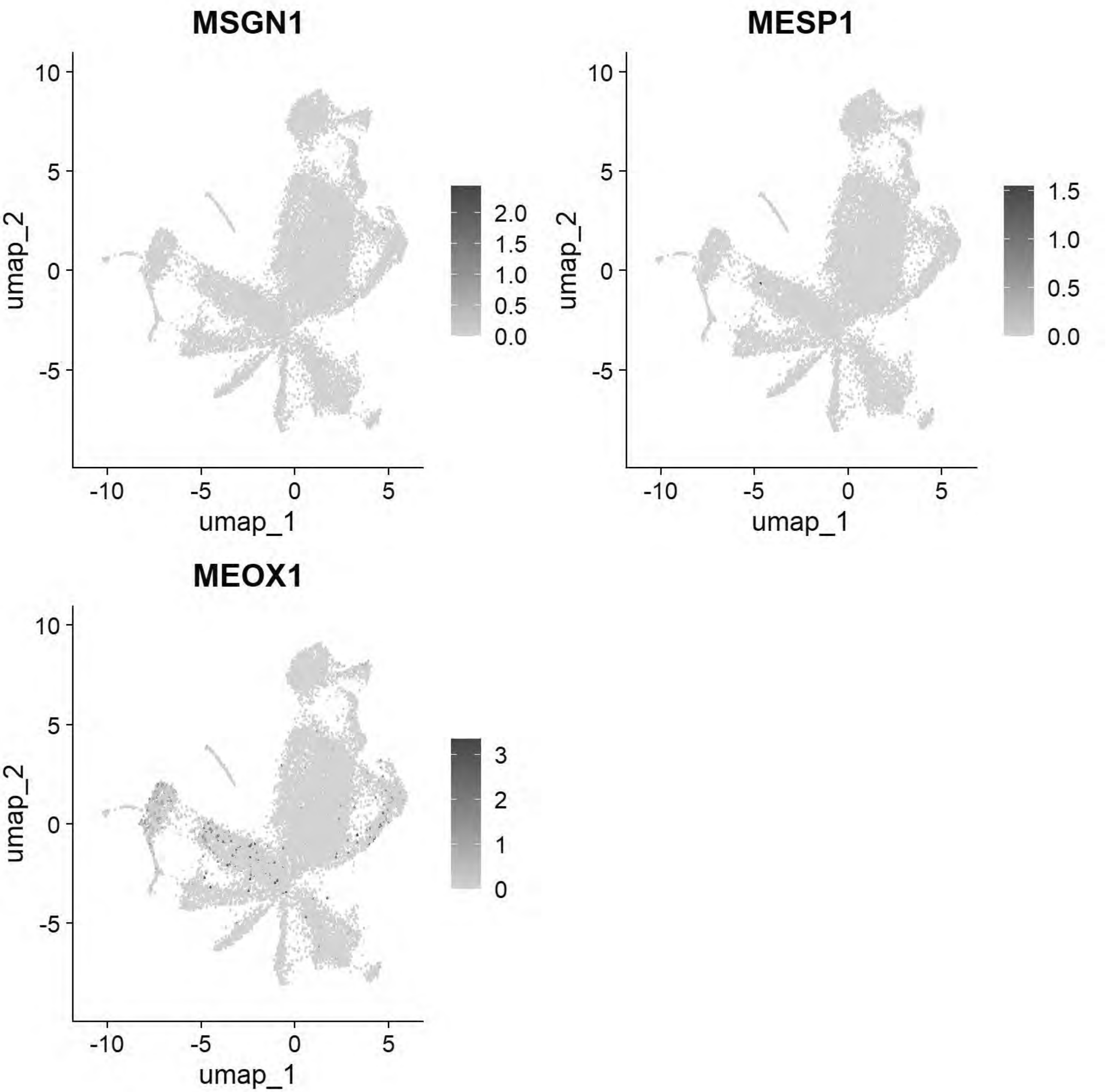


```
# Williams: Lateral Plate Mesoderm
FeaturePlot(chick, features = c("PITX2","ALX1","OLFML3","SIX1","TWIST1"))
```

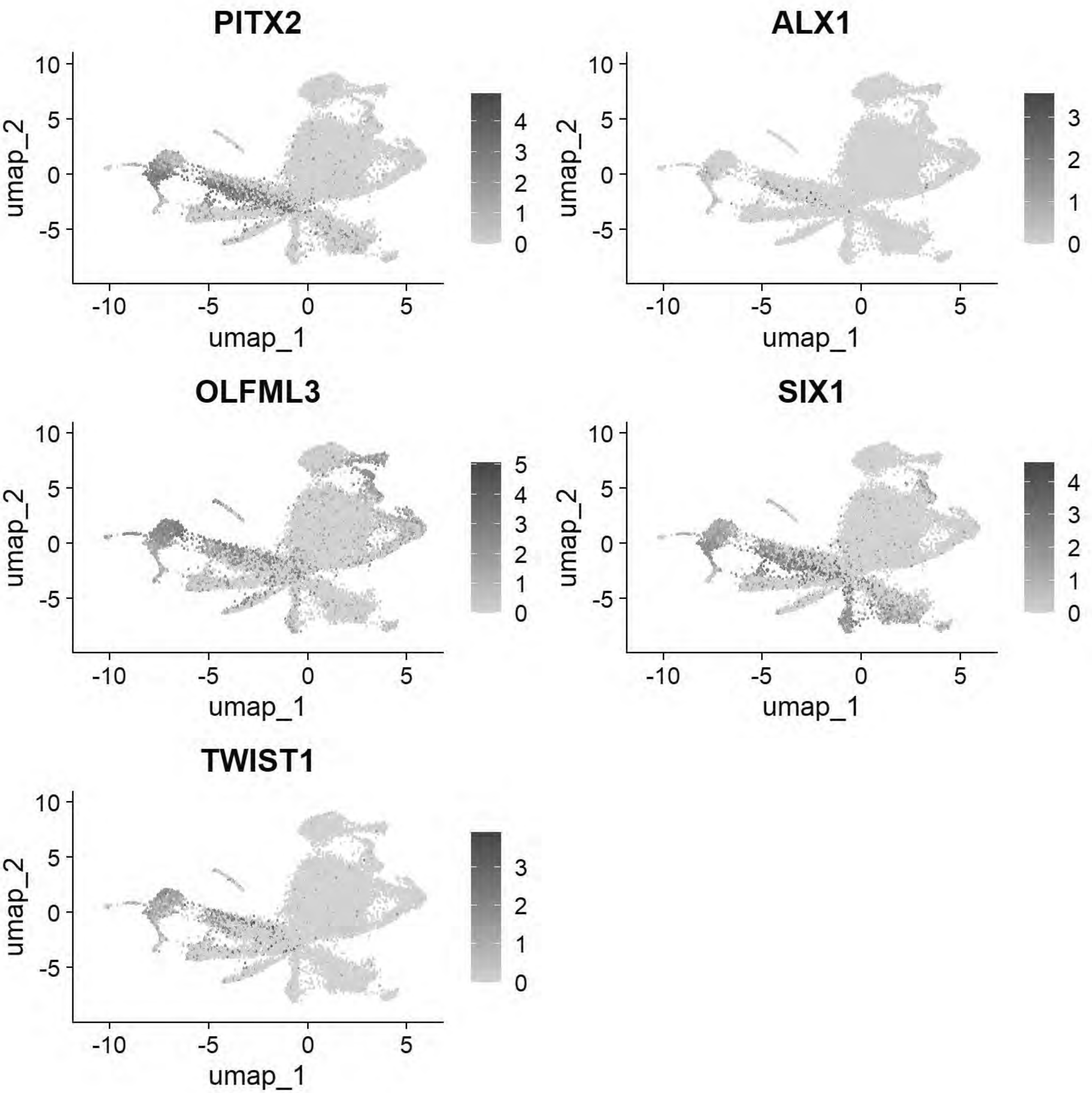


```
# Williams: Head Mesenchyme
FeaturePlot(chick, features = c("TCF21","GATA5","LMO2","ETS1","KDR"))
```

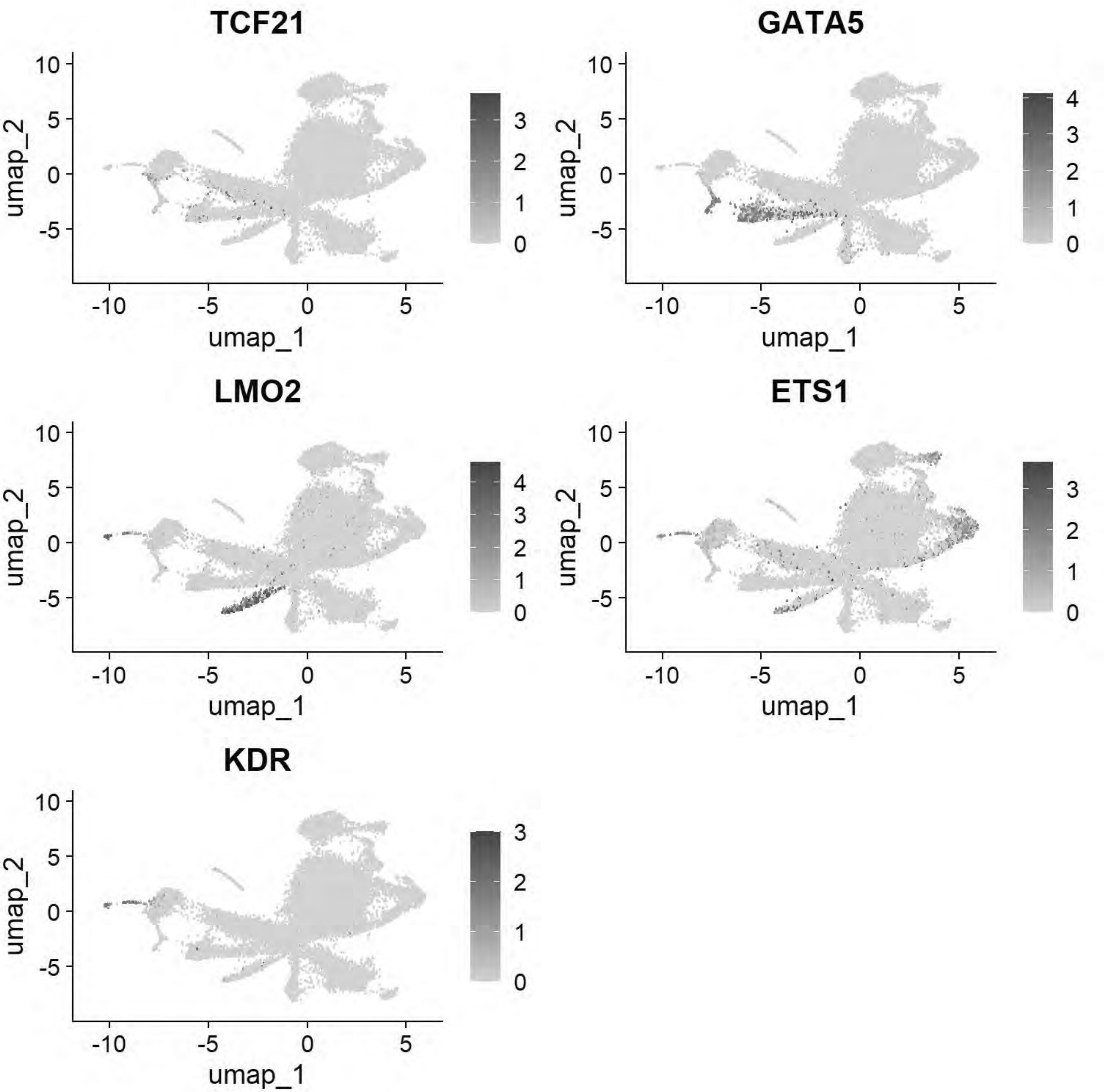


```
# Williams: Hensen's Node & Primitive Streak
FeaturePlot(chick, features = c("DLL1","FGF8","NOTO","CHRD"))
```

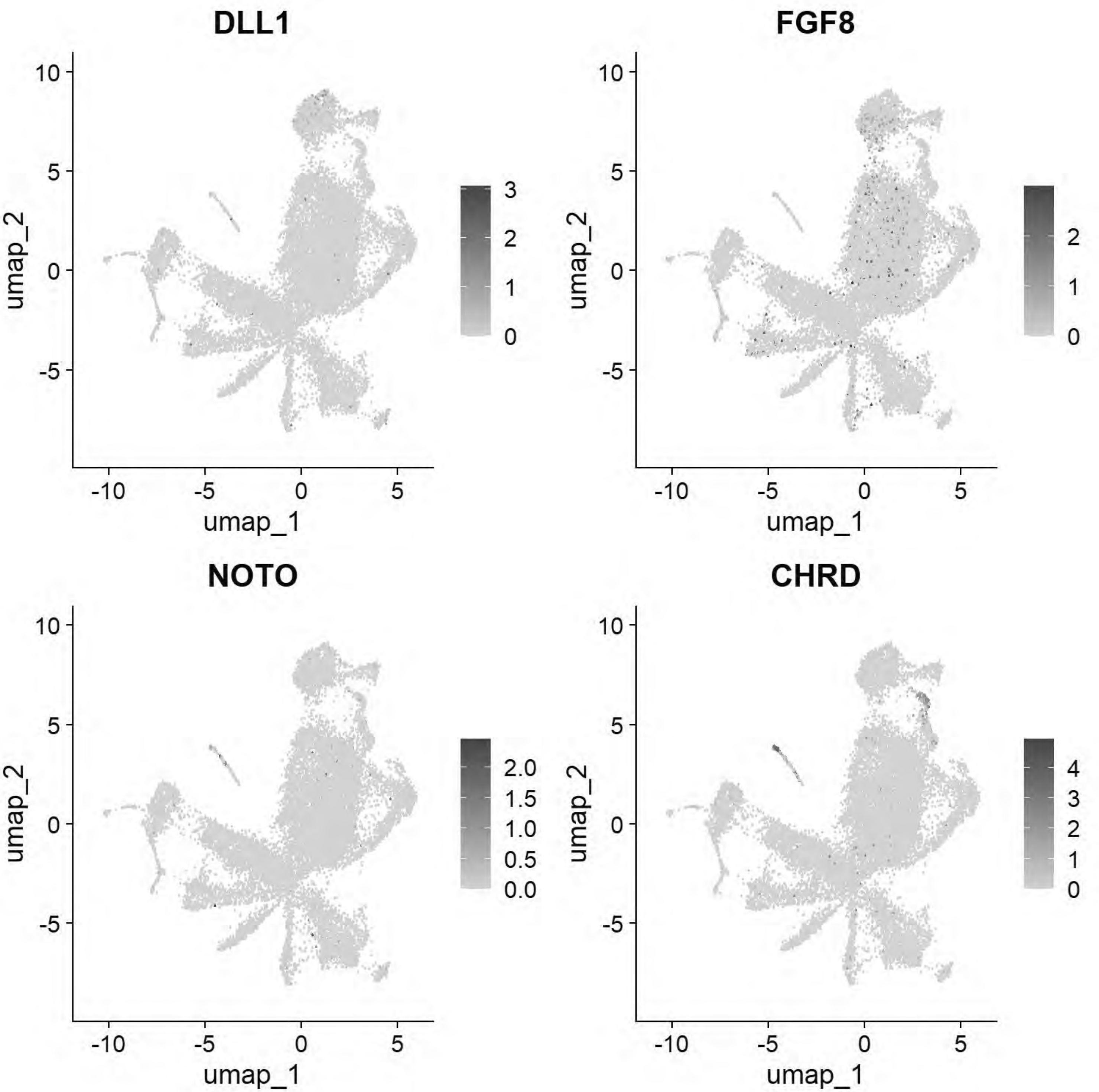


```
# Williams: Endoderm
FeaturePlot(chick, features = c("SOX17","FOXA2","CXCR4"))
```

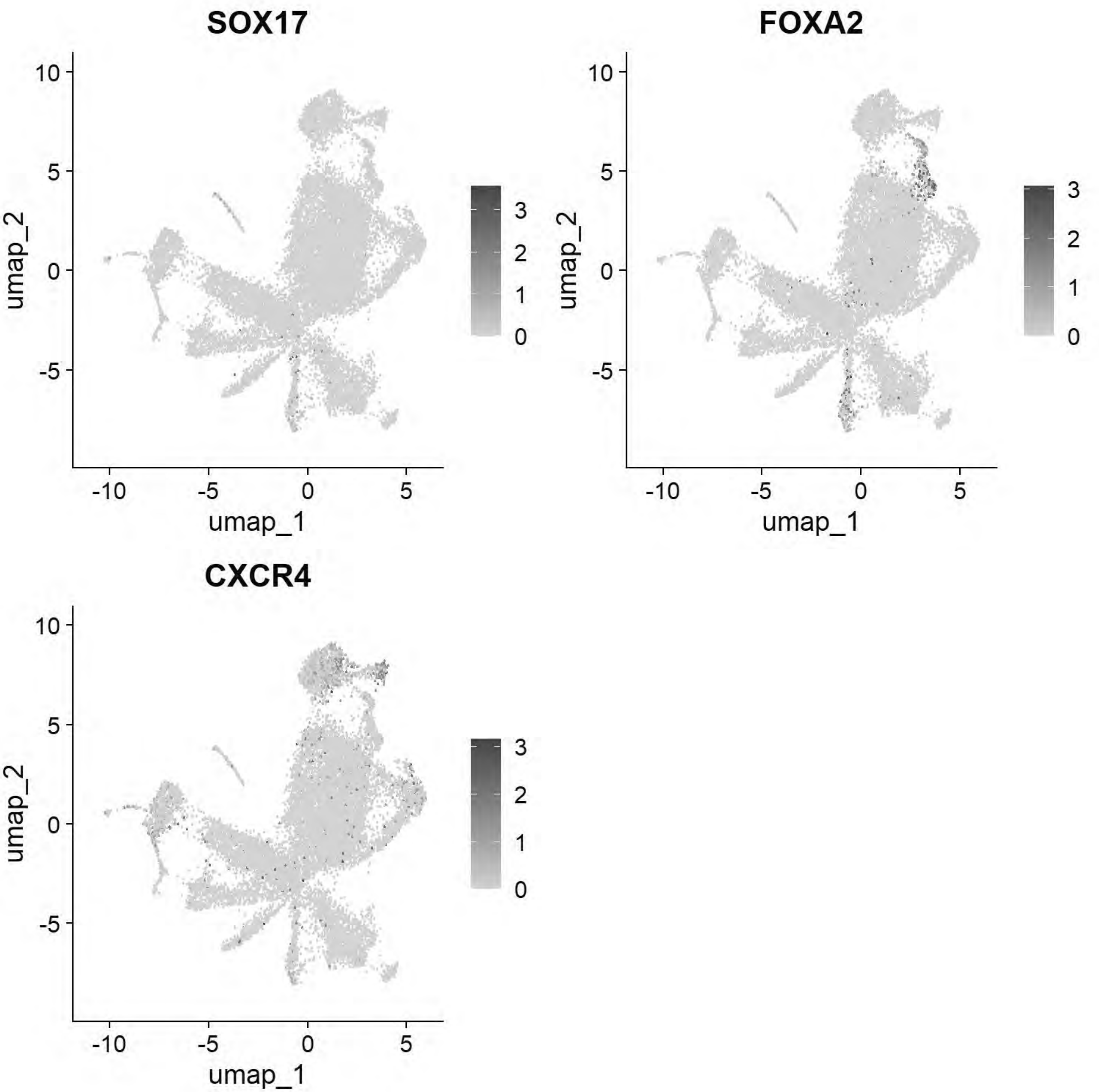


```
# Pajanoja: Ectoderm
FeaturePlot(chick, features = c("TFAP2A","DLX5","CLDN1","SOX2","NESTIN","MYCN"))
```

```
## Warning: The following requested variables were not found: NESTIN
```

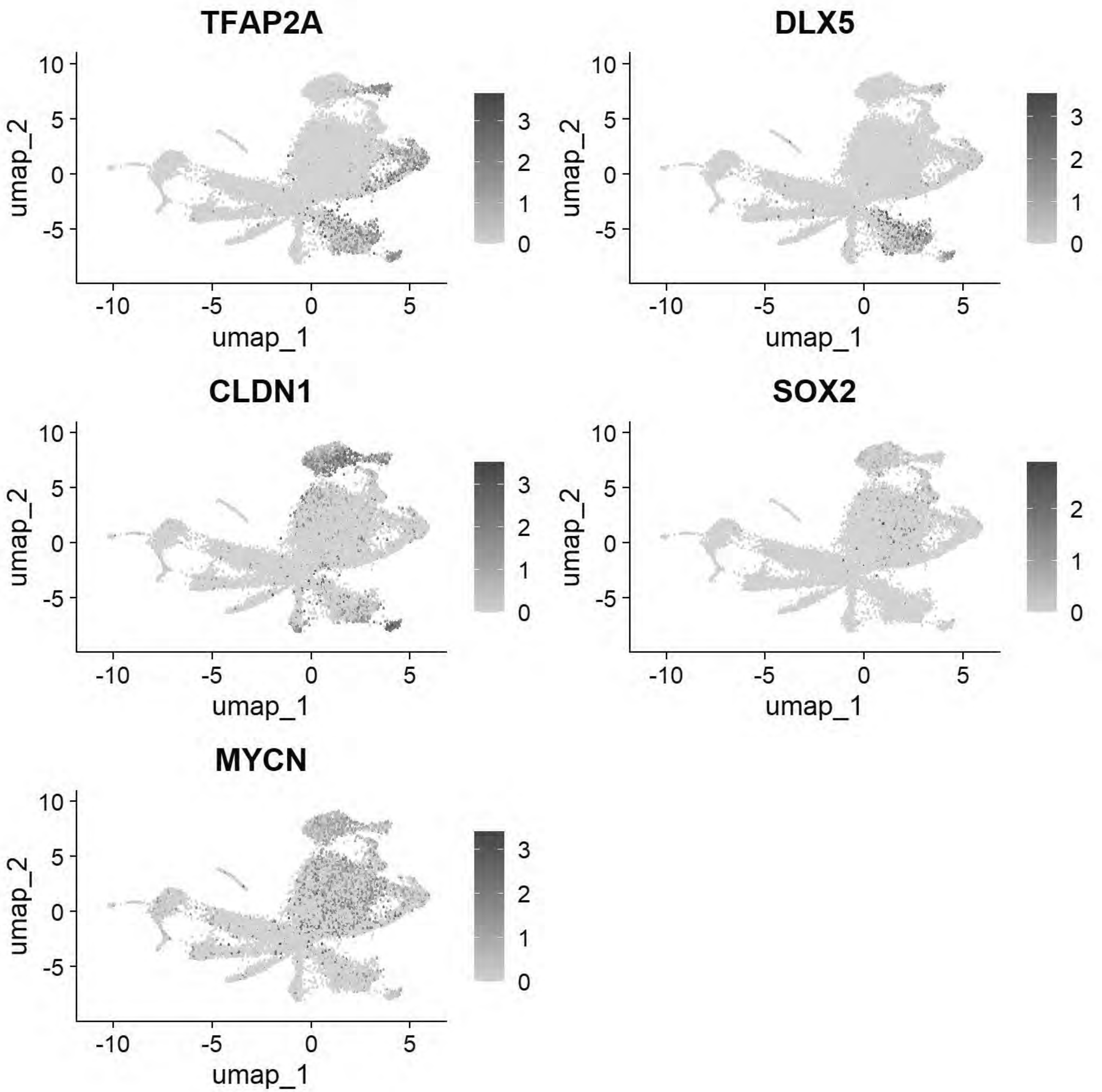


```r
# Pajanoja: Endoderm
FeaturePlot(chick, features = c("SOX17","KRT17","HHEX"))
```

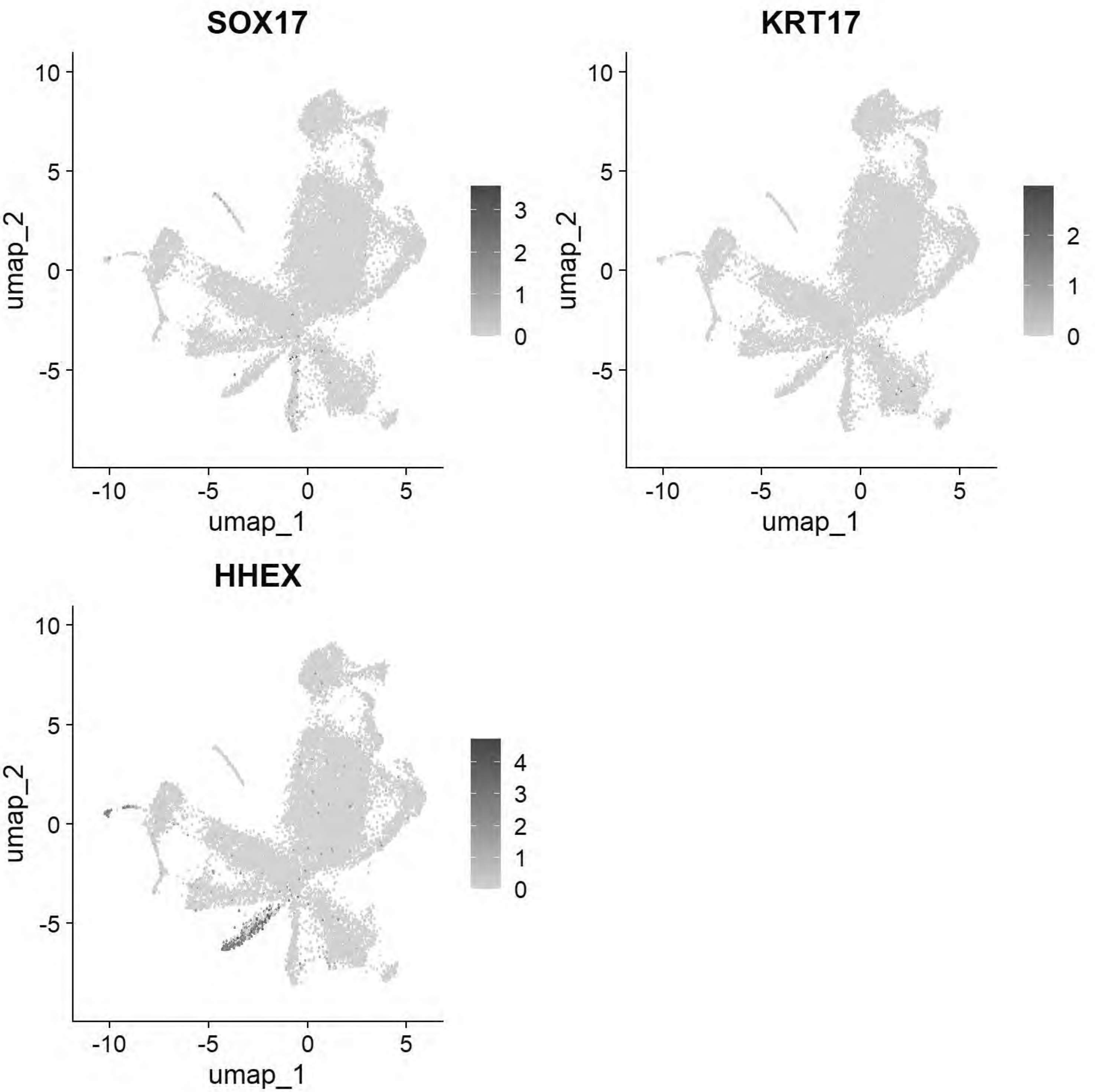


```
# Pajanoja: Mesoderm
FeaturePlot(chick, features = c("ALX1","TWIST1","HAND2","PITX2"))
```

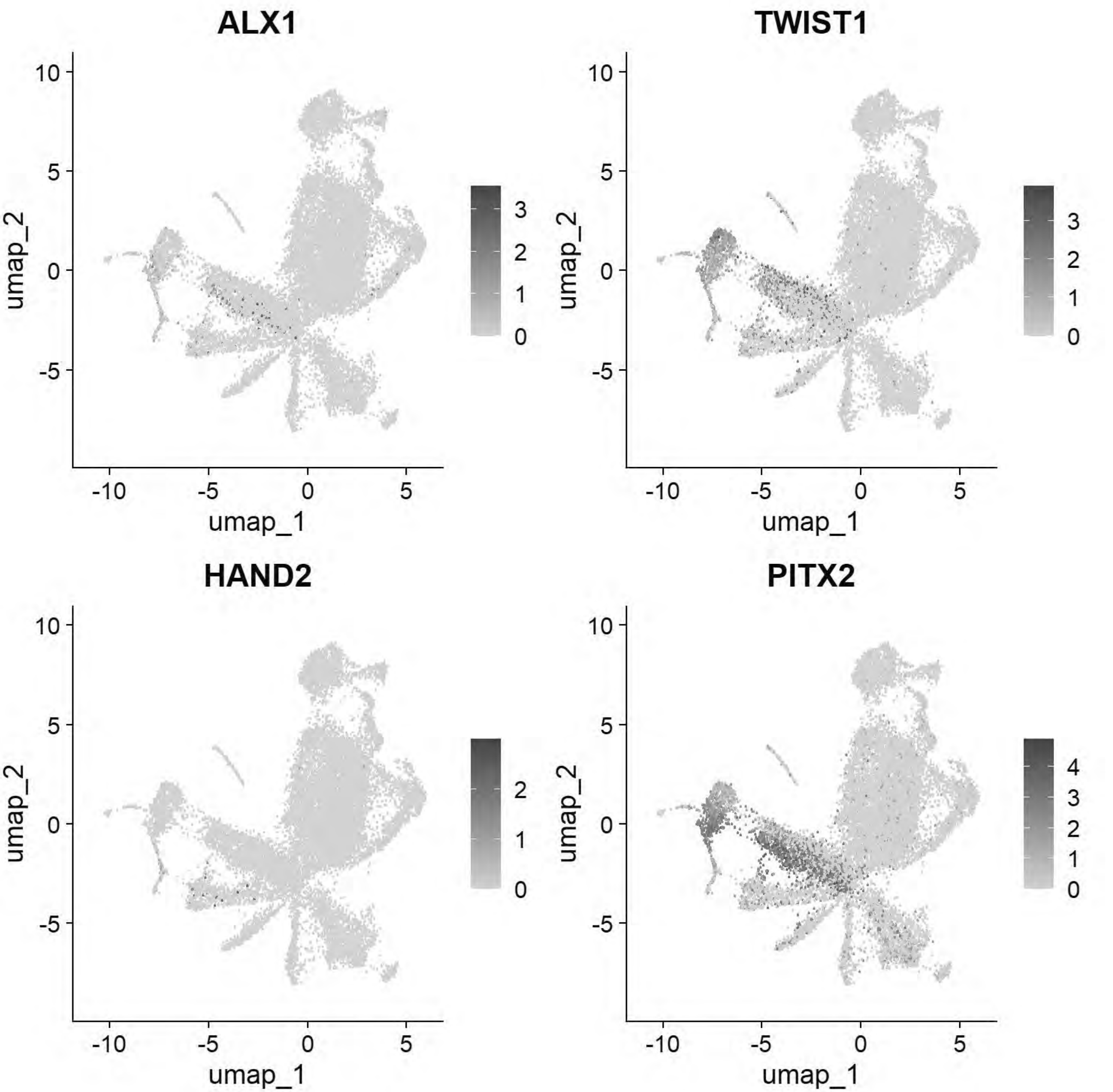


```
# Pajanoja: Notochord
FeaturePlot(chick, features = c("CHRD","TBXT","NOTO"))
```

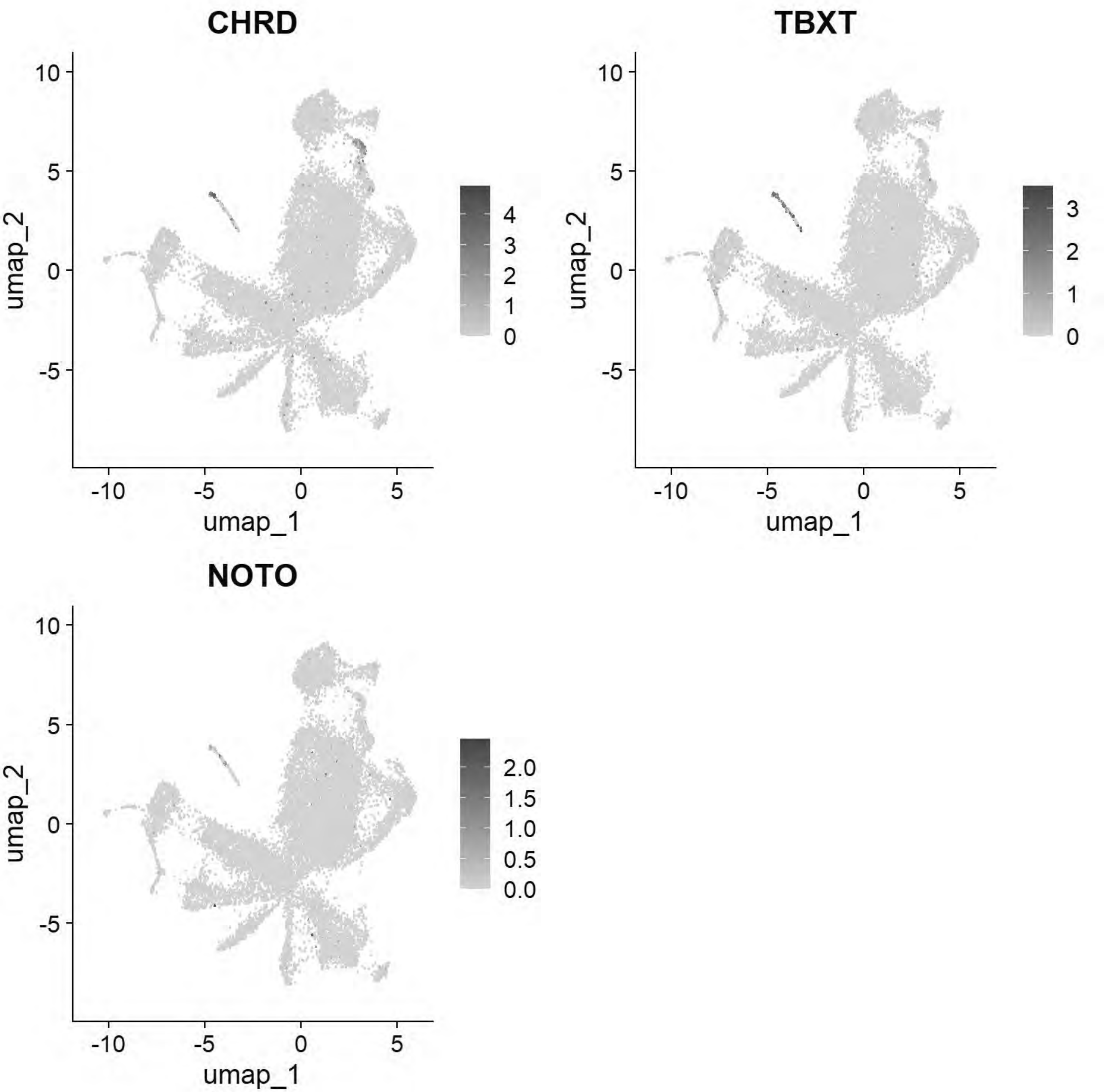


```
# Pajanoja: Ventral Neural Tube
FeaturePlot(chick, features=c("SHH","FOXA1","FOXA2"))
```

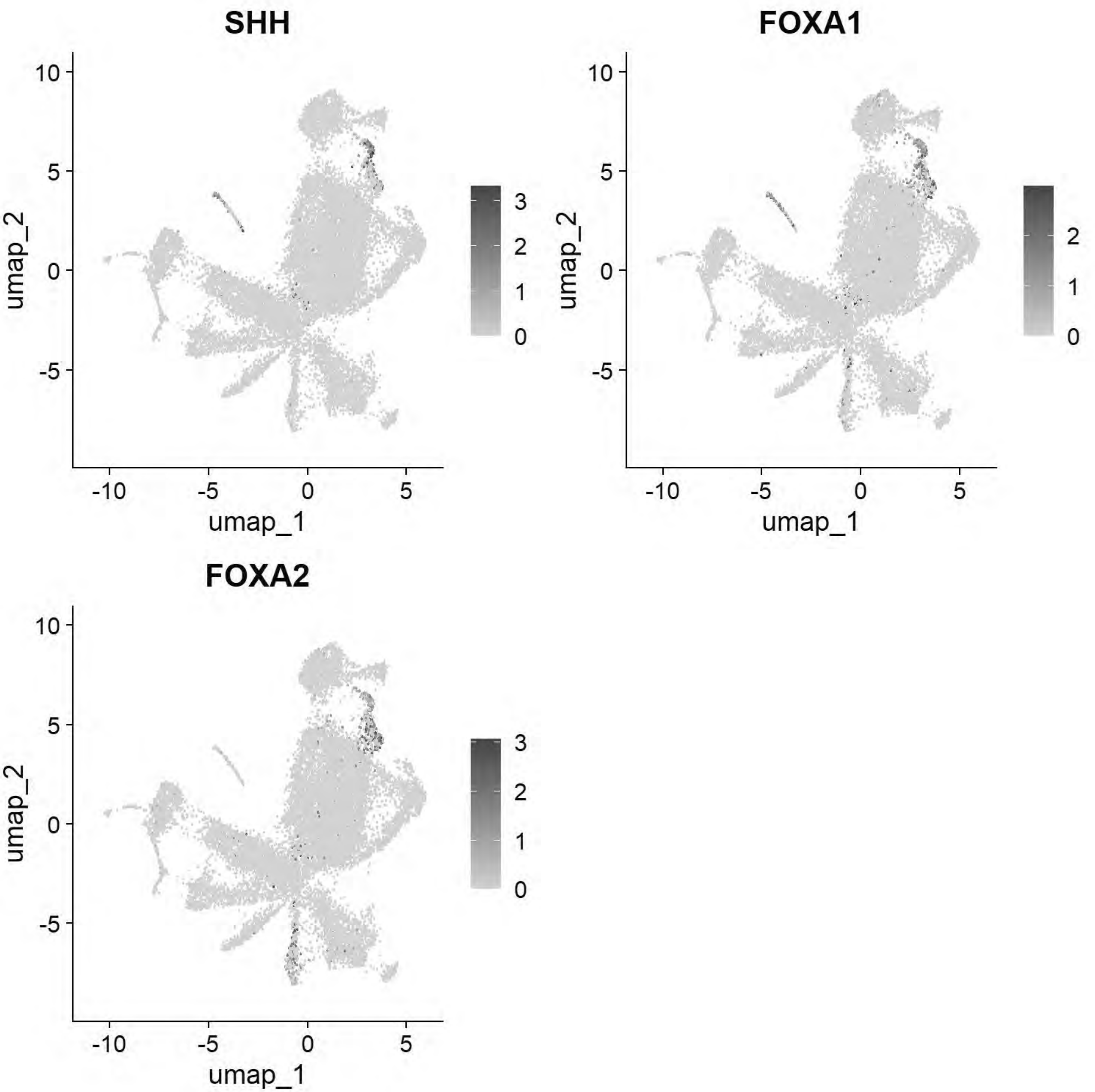


```
# Pajanoja: Neural Tube
FeaturePlot(chick, features=c("CDH2","HES5","MYCN","PAX2","NESTIN"))
```

```
## Warning: The following requested variables were not found: NESTIN
```

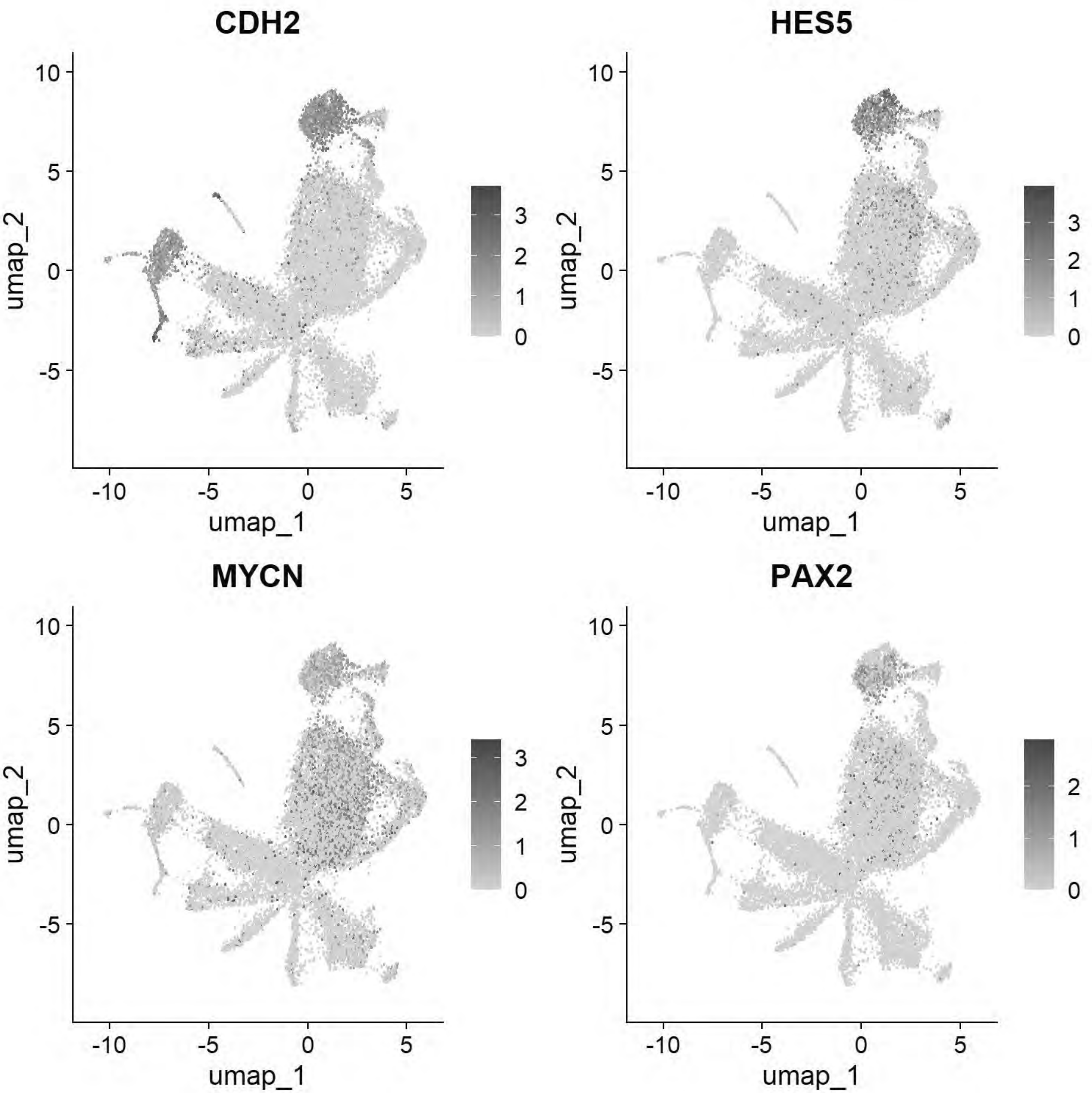


```
# Pajanoja: Non-Neural Ectoderm/Placode
FeaturePlot(chick, features=c("KRT24","KRT18","DLX5","SIX1","EYA2","TFAP2A"))
```

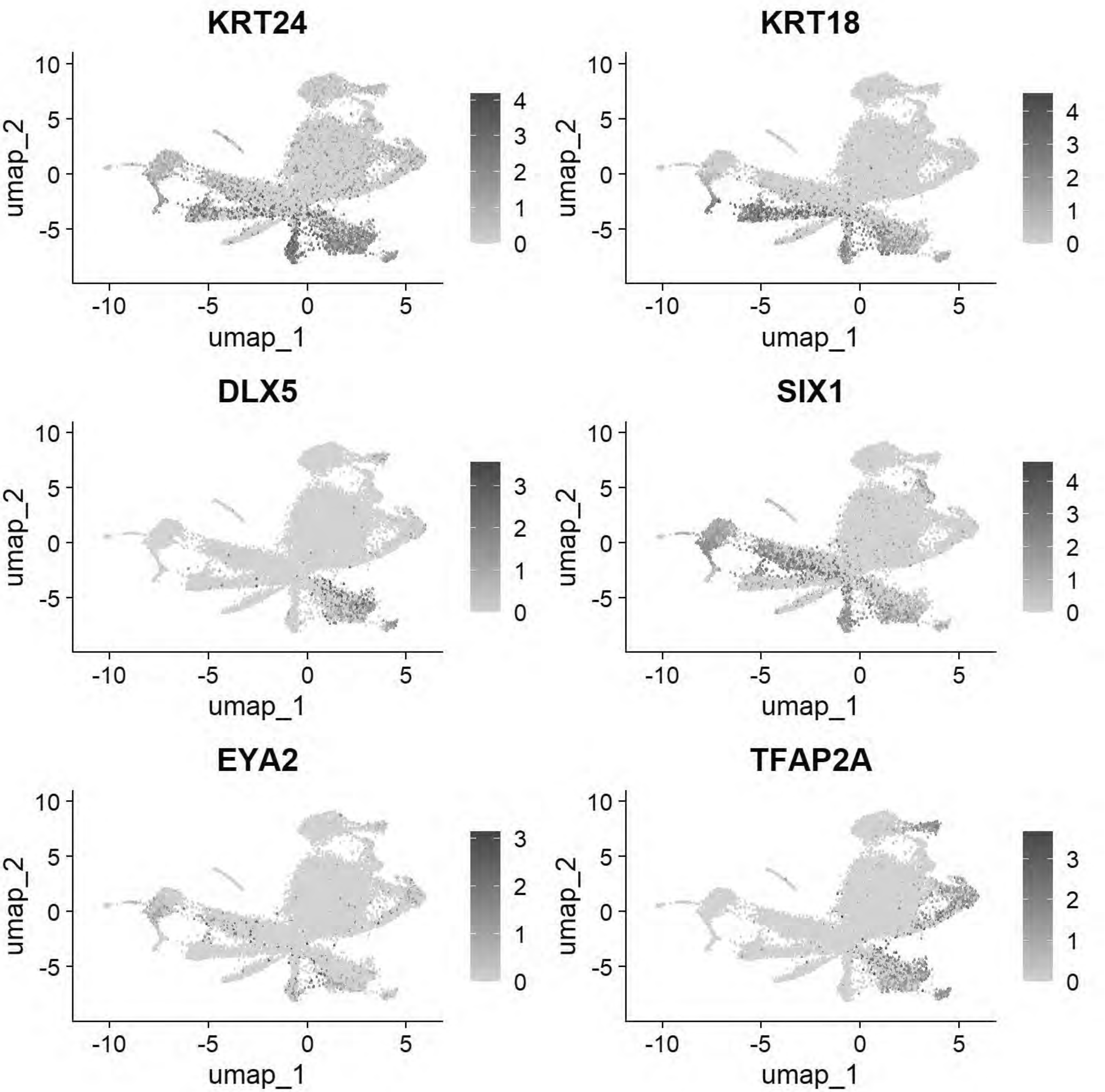


```
# Pajanoja: Neural crest
FeaturePlot(chick, features=c("FOXD3","TFAP2B","SOX9","ETS1","SNAI2"))
```

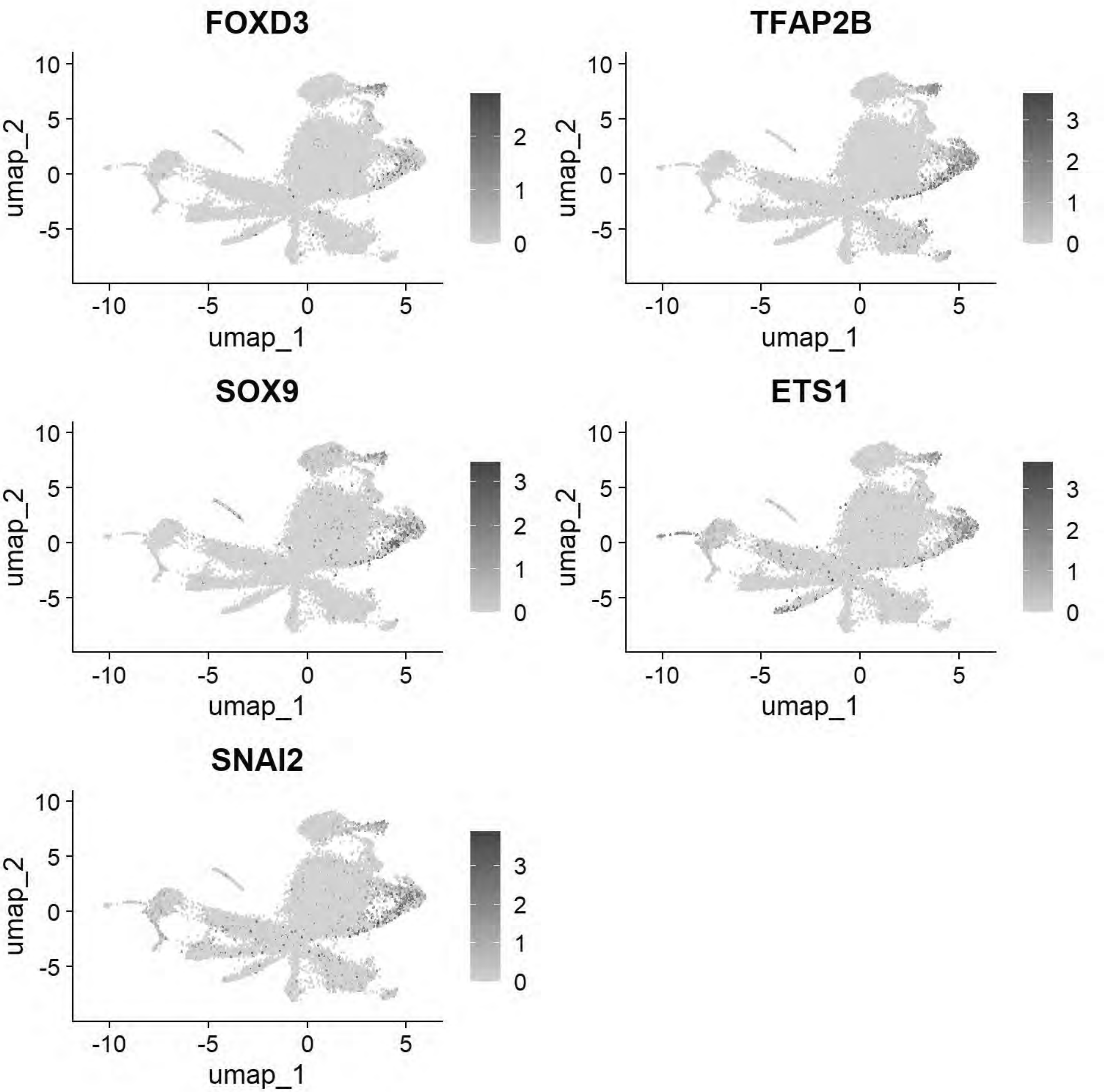


```r
gc()
```

```
##              used   (Mb) gc trigger    (Mb)   max used    (Mb)
## Ncells   5076964  271.2   13651909   729.1   21331105  1139.3
## Vcells 966224058 7371.8 2342481578 17871.8 4574774951 34902.8
```

```r
chick <- JoinLayers(chick)
chick.markers <- FindAllMarkers(chick, min.pct = 0.5, logfc.threshold = 0.5)
```

```
## Calculating cluster 0
```

```
## Calculating cluster 1
```

```
## Calculating cluster 2
```

```
## Calculating cluster 3
```

```
## Calculating cluster 4
```

```
## Calculating cluster 5
```

```
## Calculating cluster 6
```

```
## Calculating cluster 7
```

```
## Calculating cluster 8
```

```
## Calculating cluster 9
```

```
## Calculating cluster 10
```

```
## Calculating cluster 11
```

```
## Calculating cluster 12
```

```
## Calculating cluster 13
```

```
## Calculating cluster 14
```

```
## Calculating cluster 15
```

```
## Calculating cluster 16
```

```
chick.markers %>%
  group_by(cluster) %>%
  top_n(n = 5, wt = avg_log2FC) -> top5
DotPlot(chick, features = unique(top5$gene))+coord_flip()
```

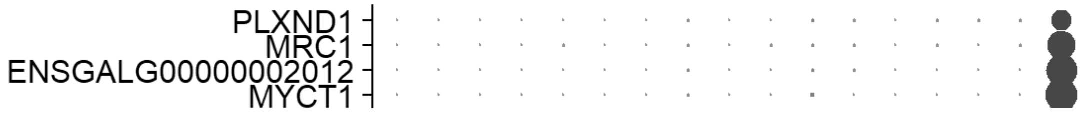

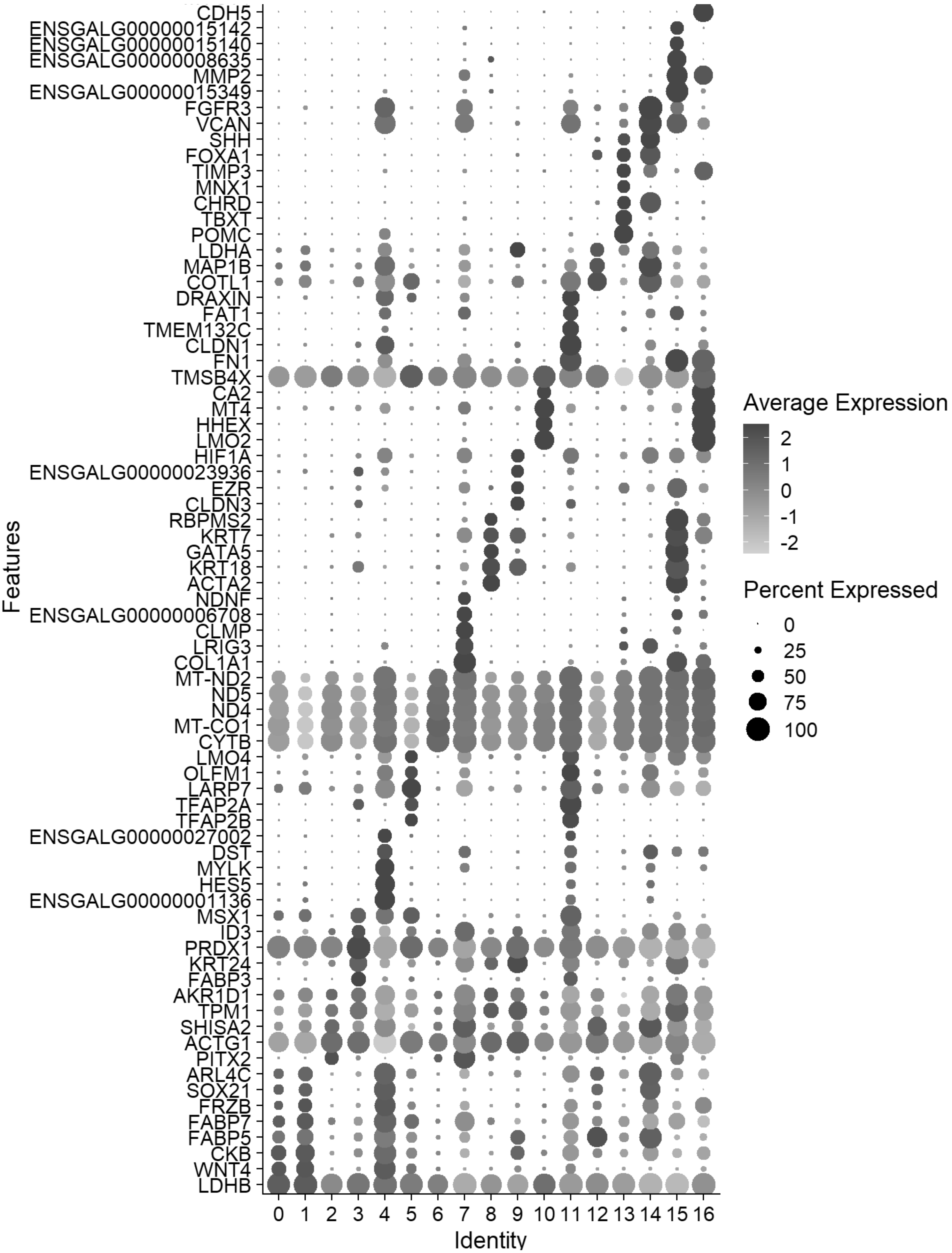
CDH5
ENSGALG00000015142
ENSGALG00000015140
ENSGALG00000008635
MMP2
ENSGALG00000015349
FGFR3
VCAN
SHH
FOXA1
TIMP3
MNX1
CHRD
TBXT
POMC
LDHA
MAP1B
COTL1
DRAXIN
FAT1
TMEM132C
CLDN1
FN1
TMSB4X
CA2
MT4
HHEX
LMO2
HIF1A
ENSGALG00000023936
EZR
CLDN3
RBPMS2
KRT7
GATA5
KRT18
ACTA2
NDNF
ENSGALG00000006708
CLMP
LRIG3
COL1A1
MT-ND2
ND5
ND4
MT-CO1
CYTB
LMO4
OLFM1
LARP7
TFAP2A
TFAP2B
ENSGALG00000027002
DST
MYLK
HES5
ENSGALG00000001136
MSX1
ID3
PRDX1
KRT24
FABP3
AKR1D1
TPM1
SHISA2
ACTG1
PITX2
ARL4C
SOX21
FRZB
FABP7
FABP5
CKB
WNT4
LDHB
Features
0 1 2 3 4 5 6 7 8 9 10 11 12 13 14 15 16
Identity
Average Expression
2
1
0
-1
-2
Percent Expressed
0
25
50
75
100

```
gc()
```

```
##              used   (Mb) gc trigger    (Mb)   max used    (Mb)
## Ncells    4981207  266.1   13651909   729.1   21331105  1139.3
## Vcells 1078020221 8224.7 2342481578 17871.8 4574774951 34902.8
```

```
cluster_map <- c("0"="Neural 1",
                 "1"="Neural 2",
                 "2"="Mesoderm 1",
                 "3"="NNE",
                 "4"="Def Neural 1",
                 "5"="Neural Crest 1",
                 "6"="Mixed",
                 "7"="Mesoderm 2",
                 "8"="Cardiac Mesoderm 1",
                 "9"="Placode",
                 "10"="Hematogenic",
                 "11"="Neural Crest 2",
                 "12"="Def Neural 2",
                 "13"="Notochord",
                 "14"="vNeural Tube",
                 "15"="Cardiac Mesoderm 2",
                 "16"="Vascular"
                 )
chick$celltype1 <- plyr::mapvalues(chick$seurat_clusters, from = names(cluster_map), to = clus
ter_map)

chick <- JoinLayers(chick)
Idents(chick) <- "celltype1" # Setting active identity ensures markers are calculated by cell
type
global_markers <- FindAllMarkers(chick, min.pct = 0.5, logfc.threshold = 0.5)
```

```
## Calculating cluster Neural 1
```

```
## Calculating cluster Neural 2
```

```
## Calculating cluster Mesoderm 1
```

```
## Calculating cluster NNE
```

```
## Calculating cluster Def Neural 1
```

```
## Calculating cluster Neural Crest 1
```

```
## Calculating cluster Mixed
```

```
## Calculating cluster Mesoderm 2
```

```
## Calculating cluster Cardiac Mesoderm 1
```

```
## Calculating cluster Placode
```

```
## Calculating cluster Hematogenic
```

```
## Calculating cluster Neural Crest 2
```

```
## Calculating cluster Def Neural 2
```

```
## Calculating cluster Notochord
```

```
## Calculating cluster vNeural Tube
```

```
## Calculating cluster Cardiac Mesoderm 2
```

```
## Calculating cluster Vascular
```

```
write.csv(global_markers, "E:/Rogers Lab/Rogers NC EMT/Supplementary file 5_HH9_Markers_Annotated.csv", row.names = FALSE)

celltype_colors <- c("Neural 1"="#7FCDBB",
                 "Neural 2"="#41B6C4",
                 "Mesoderm 1"="#FC9272",
                 "NNE"="#A1D99B",
                 "Def Neural 1"="#2171B5",
                 "Neural Crest 1"="#8C96C6",
                 "Mixed"="#D9D9D9",
                 "Mesoderm 2"="#CB181D",
                 "Cardiac Mesoderm 1"="#E31A1C",
                 "Placode"="#41AB5D",
                 "Hematogenic"="#BD0026",
                 "Neural Crest 2"="#8C6BB1",
                 "Def Neural 2"="#1D91C0",
                 "Notochord"="#DEEBF7",
                 "vNeural Tube"="#08519C",
                 "Cardiac Mesoderm 2"="#A50F15",
                 "Vascular"="#800026")

p_umap <- DimPlot(chick, group.by = "celltype1",cols = celltype_colors, label = TRUE, repel = TRUE) + ggtitle(NULL)+
            NoLegend()
p_umap
```

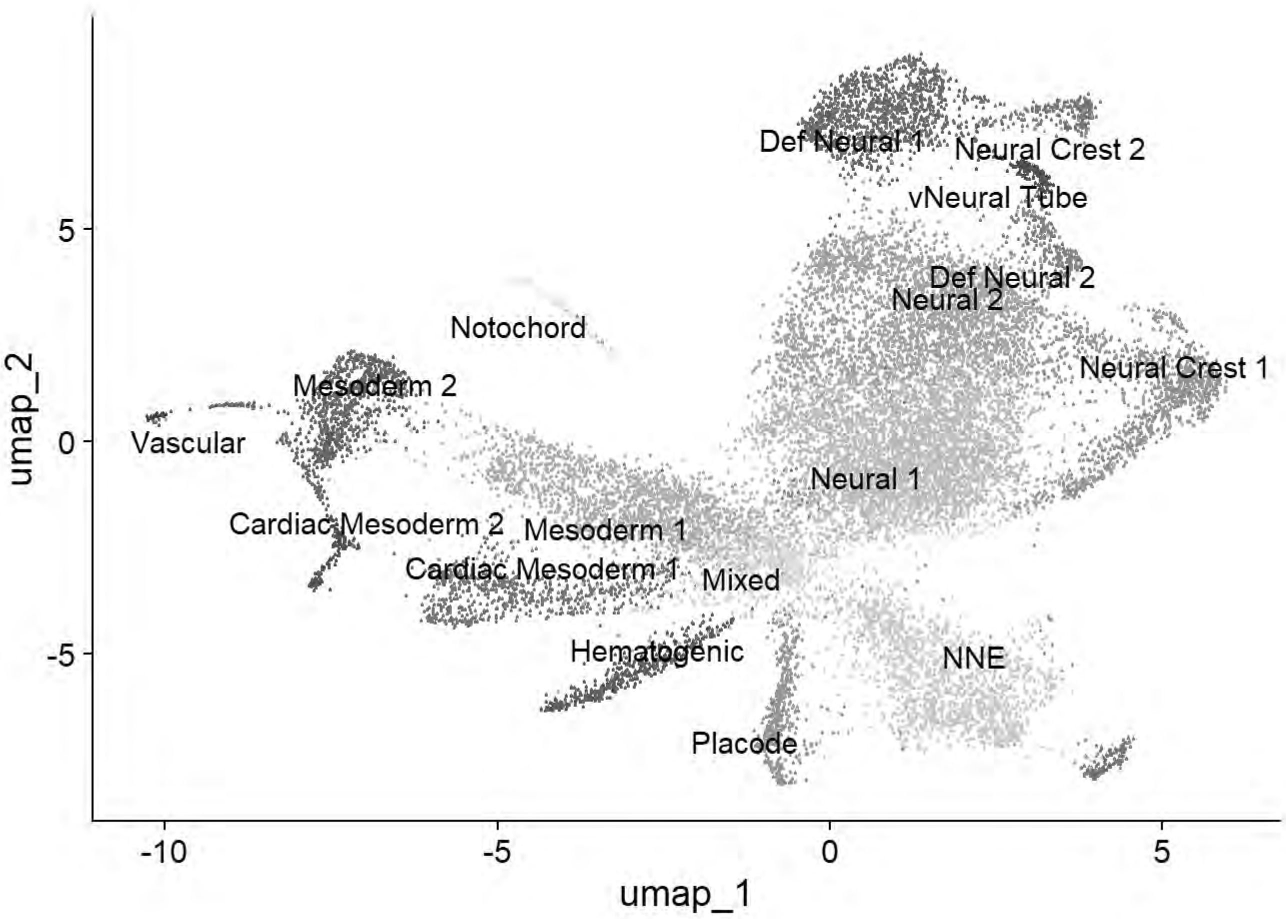


```
ggsave("E:/Rogers Lab/Rogers NC EMT/Figures/Fig_HH9_Summary_UMAP1.png", plot = p_umap, width =
 6, height = 6, dpi = 600)

cptbl <- table(chick@meta.data$celltype1,chick@meta.data$stage)
cptbl <- cbind(cptbl, TOTAL = rowSums(cptbl))
colnames(cptbl)[length(cptbl[1,])] <- "TOTAL"

tg = gridExtra::tableGrob(cptbl)
h = grid::convertHeight(sum(tg$heights), "in", TRUE)
w = grid::convertWidth(sum(tg$widths), "in", TRUE)
ggsave("E:/Rogers Lab/Rogers NC EMT/Figures/Fig_HH9_Summary_Table.png", tg, width=w, height=h,
 device = 'png', dpi =600)

top5 <- global_markers %>% group_by(cluster) %>% top_n(n = 5, wt = avg_log2FC)
p_dot=DotPlot(chick, features = unique(top5$gene), group.by = "celltype1") + # cluster.idents
dendrograms the y-axis
  coord_flip() +
  scale_size_continuous(range = c(1, 10)) +
  scale_color_gradientn(colors = c("grey90", "#313695", "#d73027"))+
  theme_light() +
  theme(
    axis.text.x = element_text(angle = 45, hjust = 1, color = "black", size = 12),
    axis.text.y = element_text(face = "italic", color = "black", size = 12),
    axis.title.x = element_blank(),
    axis.title.y = element_blank(),
    panel.grid.major = element_line(color = "grey90")
```

```
)
```

```
## Scale for size is already present.
## Adding another scale for size, which will replace the existing scale.
```

```
## Scale for colour is already present.
## Adding another scale for colour, which will replace the existing scale.
```

```
p_dot
```

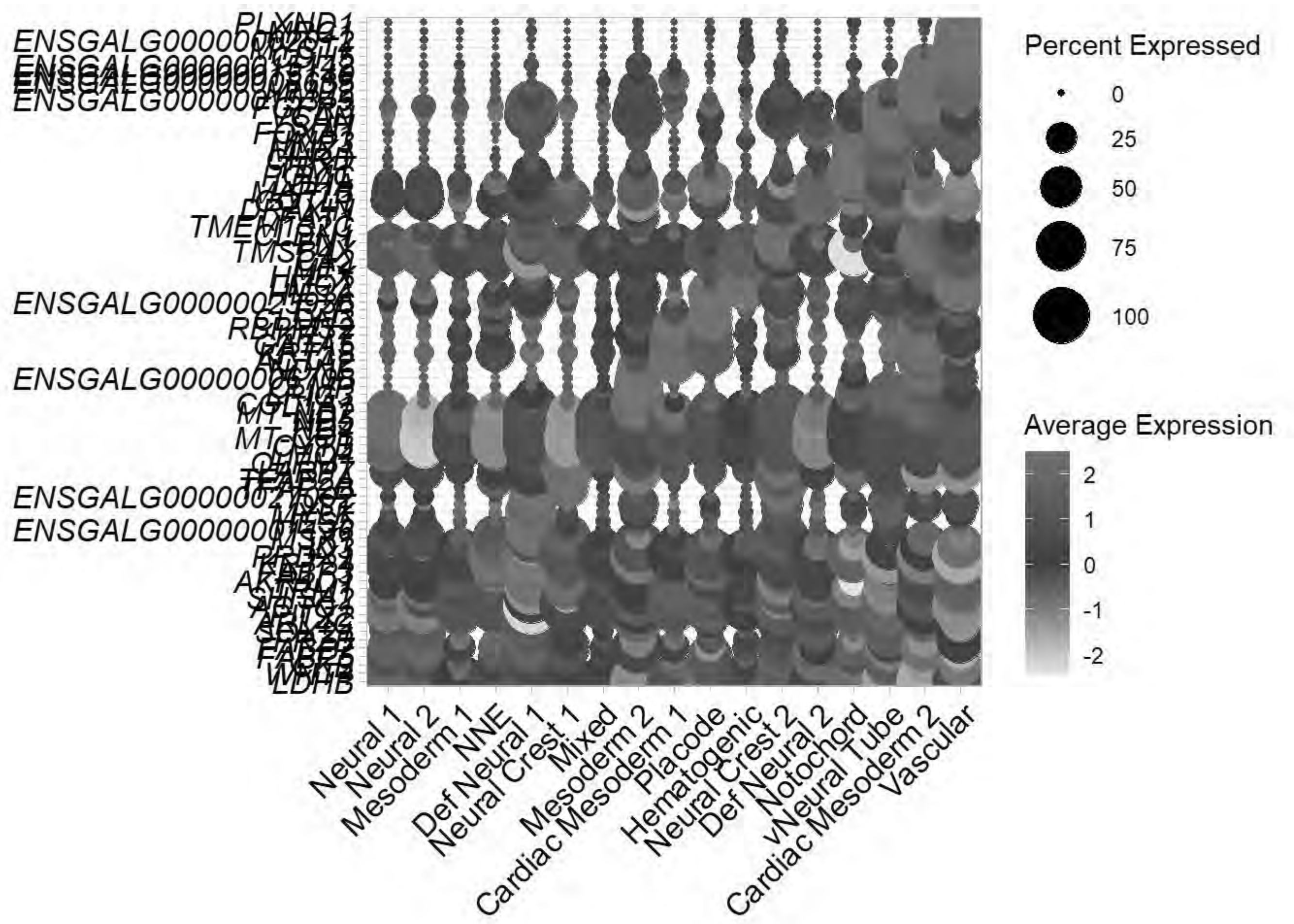


```
ggsave("E:/Rogers Lab/Rogers NC EMT/Figures/Fig_HH9_Summary_Dotplot.png", plot = p_dot, width
= 10, height = 16, dpi = 600)
```

```
saveRDS(chick, "E:\\Rogers Lab\\Rogers NC EMT\\HH9_chick_v2.rds")

Idents(chick)<-"celltype1"
chick=subset(chick, idents =c("Neural 1","Neural 2","NNE","Def Neural 1","Neural Crest 1","Pla
code","Neural Crest 2","Def Neural 2","vNeural Tube"))
chick
```

```
## An object of class Seurat
## 18523 features across 12769 samples within 1 assay
## Active assay: RNA (18523 features, 2000 variable features)
```

```
##  3 layers present: data, counts, scale.data
##  3 dimensional reductions calculated: pca, harmony, umap
```

```
saveRDS(chick,"E:\\Rogers Lab\\Rogers NC EMT\\HH9_chick_NC_v2.rds")
gc()
```

```
##              used   (Mb) gc trigger    (Mb)   max used    (Mb)
## Ncells    4929464  263.3   13651909   729.1   21331105  1139.3
## Vcells 1043414189 7960.7 2342481578 17871.8 4574774951 34902.8
```

### Stage Analysis: HH11

```
chick=readRDS("E:\\Rogers Lab\\Rogers NC EMT\\Global_chick_v2.rds")
chick
```

```
## An object of class Seurat 
## 18523 features across 82092 samples within 1 assay 
## Active assay: RNA (18523 features, 2000 variable features)
##  3 layers present: data, counts, scale.data
##  3 dimensional reductions calculated: pca, harmony, umap
```

```
Idents(chick) <- "stage"
chick=subset(chick, idents =c("HH11"))
chick
```

```
## An object of class Seurat 
## 18523 features across 7470 samples within 1 assay 
## Active assay: RNA (18523 features, 2000 variable features)
##  3 layers present: data, counts, scale.data
##  3 dimensional reductions calculated: pca, harmony, umap
```

```
gc()
```

```
##              used   (Mb) gc trigger    (Mb)   max used    (Mb)
## Ncells    4929739  263.3   13651909   729.1   21331105  1139.3
## Vcells 1010726229 7711.3 2342481578 17871.8 4574774951 34902.8
```

```
chick <- NormalizeData(chick)
```

```
## Normalizing layer: counts
```

```
chick <- CellCycleScoring(chick, s.features = cc.genes$s.genes, g2m.features = cc.genes$g2m.ge
nes, set.ident = TRUE)
chick <- FindVariableFeatures(chick)
```

```
## Finding variable features for layer counts
```

```
chick <- ScaleData(chick, vars.to.regress = c("percent.mt","G2M.Score","S.Score"))
```

```
## Regressing out percent.mt, G2M.Score, S.Score
```

```
## Centering and scaling data matrix
```

```
chick <- RunPCA(chick, npcs = 50, verbose = FALSE)

ElbowPlot(chick, ndims= 50, reduction = "pca")
```

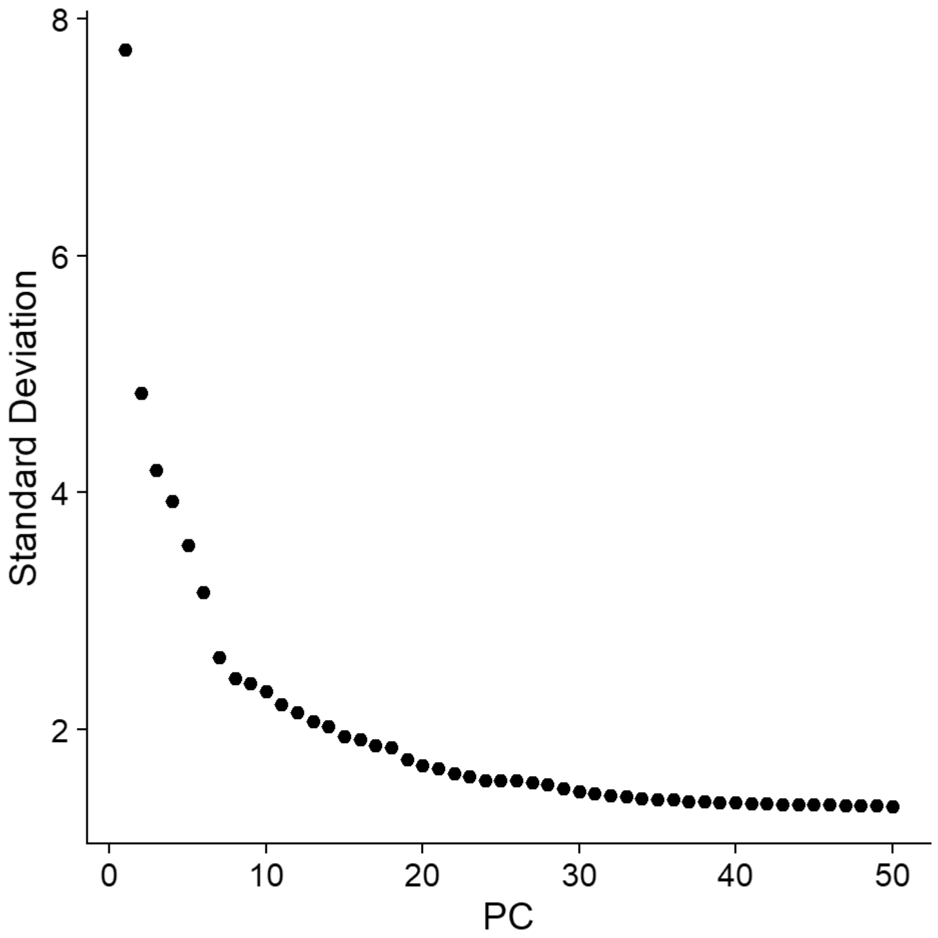


```
gc()
```

```
##              used   (Mb) gc trigger    (Mb)   max used    (Mb)
## Ncells    4951022  264.5   13651909   729.1   21331105  1139.3
## Vcells 1010873936 7712.4 2342481578 17871.8 4574774951 34902.8
```

```
chick <- FindNeighbors(chick, reduction="pca", dims=1:30, nn.method="rann", k.param=20)
```

```
## Computing nearest neighbor graph
```

```
## Computing SNN
```

```
chick <- RunUMAP(chick, reduction="pca", dims=1:30)
```

```
## 23:23:23 UMAP embedding parameters a = 0.9922 b = 1.112
```

```
## 23:23:23 Read 7470 rows and found 30 numeric columns
```

```
## 23:23:23 Using Annoy for neighbor search, n_neighbors = 30
```

```
## 23:23:23 Building Annoy index with metric = cosine, n_trees = 50
```

```
## 0%   10   20   30   40   50   60   70   80   90   100%
```

```
## [----|----|----|----|----|----|----|----|----|----|
```

```
## **************************************************|
## 23:23:24 Writing NN index file to temp file C:\Users\RANEES~1\AppData\Local\Temp\RtmpqcWgVb
\file6a2c75af234d
## 23:23:24 Searching Annoy index using 1 thread, search_k = 3000
## 23:23:25 Annoy recall = 100%
## 23:23:26 Commencing smooth kNN distance calibration using 1 thread with target n_neighbors
= 30
## 23:23:26 Initializing from normalized Laplacian + noise (using RSpectra)
## 23:23:26 Commencing optimization for 500 epochs, with 325366 positive edges
## 23:23:26 Using rng type: pcg
## 23:23:43 Optimization finished
```

```
chick <- FindClusters(chick, resolution=0.4)
```

```
## Modularity Optimizer version 1.3.0 by Ludo Waltman and Nees Jan van Eck
##
## Number of nodes: 7470
## Number of edges: 279475
##
## Running Louvain algorithm...
## Maximum modularity in 10 random starts: 0.9114
## Number of communities: 13
## Elapsed time: 0 seconds
```

```
DimPlot(chick)
```

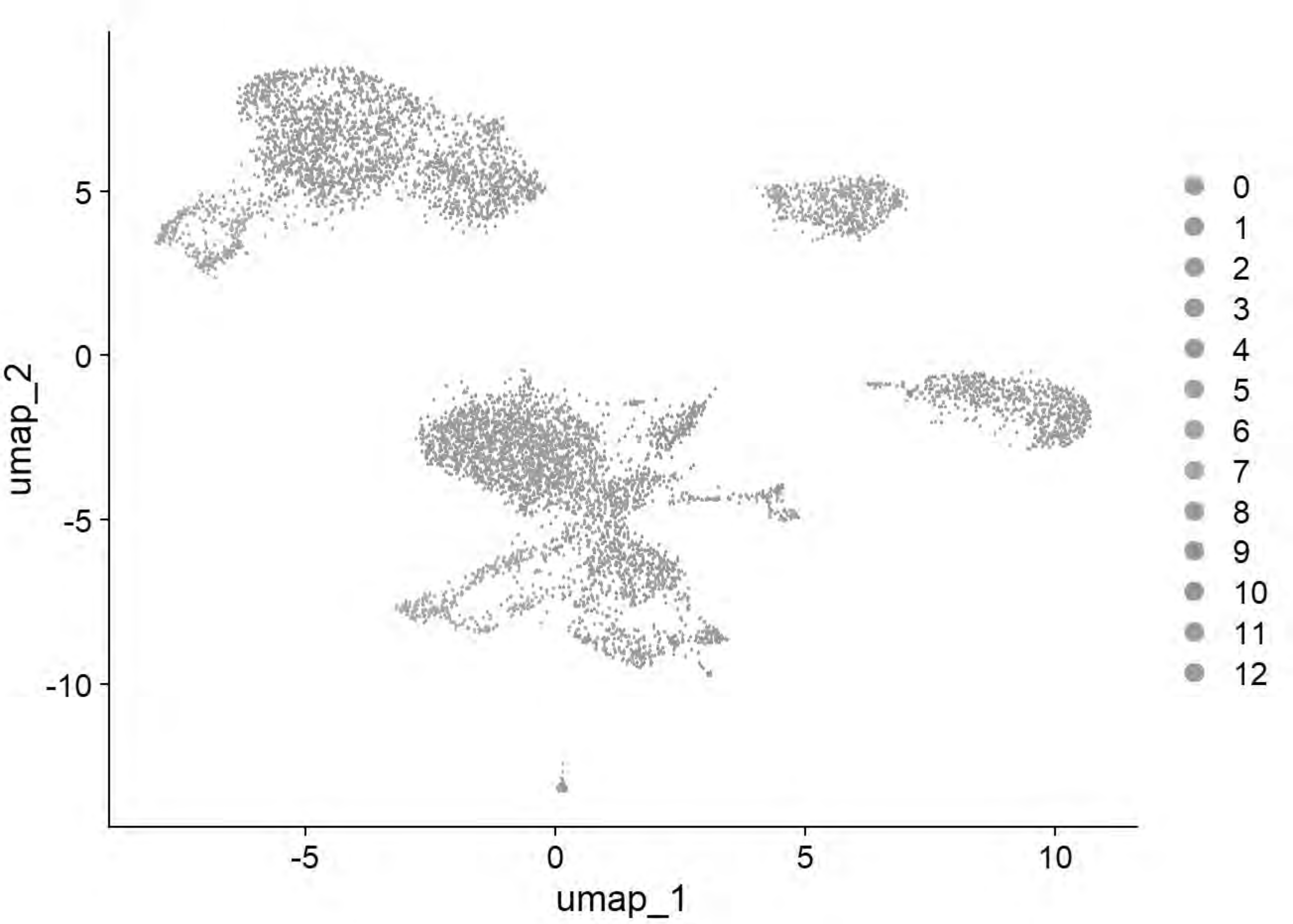


```
DimPlot(chick, group.by = "sample_id")
```

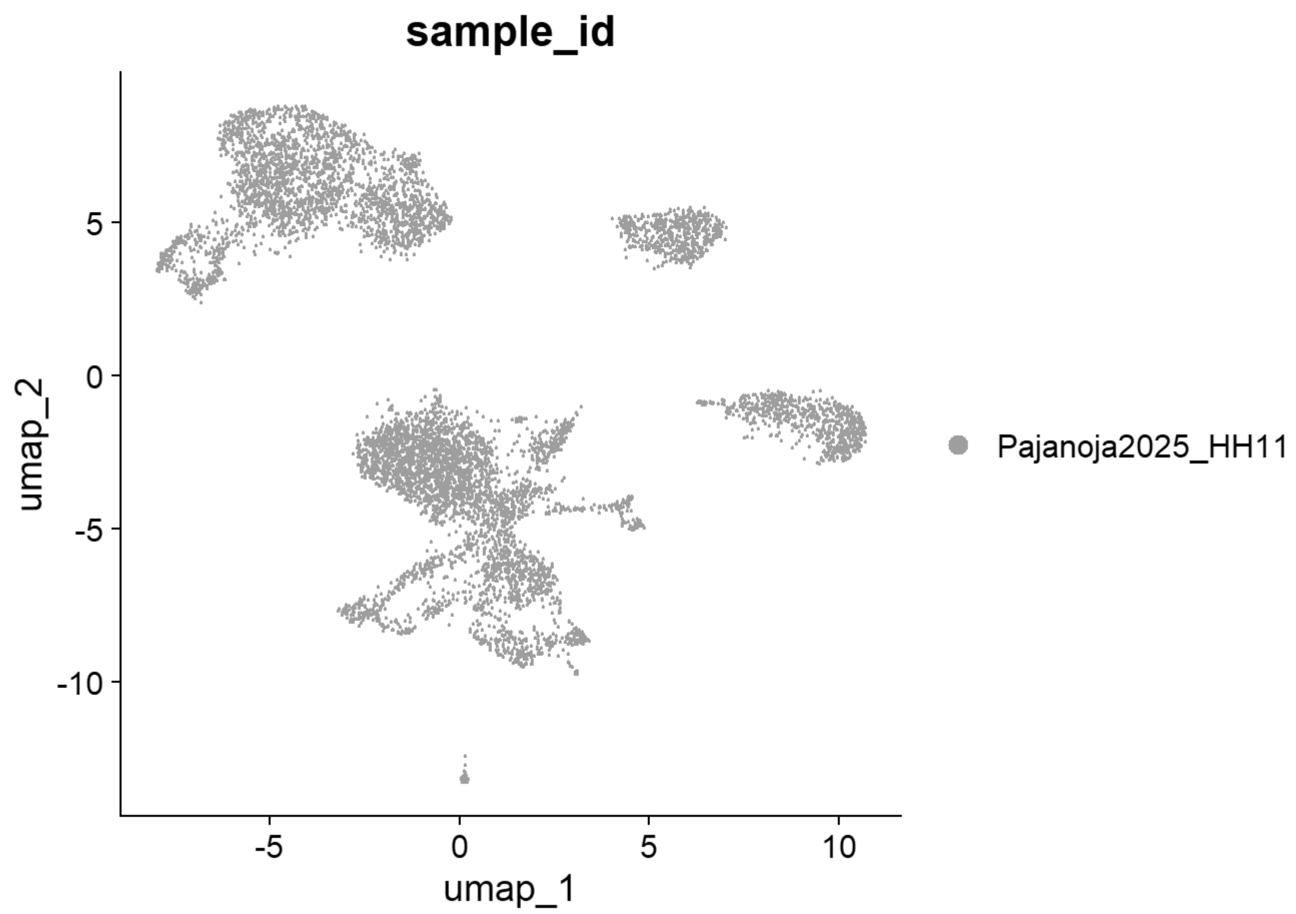


```
chick = FindClusters(chick, resolution= seq(0,2,0.1),n.start=10)
```

```
## Modularity Optimizer version 1.3.0 by Ludo Waltman and Nees Jan van Eck
##
## Number of nodes: 7470
## Number of edges: 279475
##
## Running Louvain algorithm...
## Maximum modularity in 10 random starts: 1.0000
## Number of communities: 1
## Elapsed time: 0 seconds
## Modularity Optimizer version 1.3.0 by Ludo Waltman and Nees Jan van Eck
##
## Number of nodes: 7470
## Number of edges: 279475
##
## Running Louvain algorithm...
## Maximum modularity in 10 random starts: 0.9632
## Number of communities: 7
## Elapsed time: 0 seconds
## Modularity Optimizer version 1.3.0 by Ludo Waltman and Nees Jan van Eck
##
## Number of nodes: 7470
## Number of edges: 279475
##
```

```
## Running Louvain algorithm...
## Maximum modularity in 10 random starts: 0.9417
## Number of communities: 10
## Elapsed time: 0 seconds
## Modularity Optimizer version 1.3.0 by Ludo Waltman and Nees Jan van Eck
##
## Number of nodes: 7470
## Number of edges: 279475
##
## Running Louvain algorithm...
## Maximum modularity in 10 random starts: 0.9259
## Number of communities: 11
## Elapsed time: 0 seconds
## Modularity Optimizer version 1.3.0 by Ludo Waltman and Nees Jan van Eck
##
## Number of nodes: 7470
## Number of edges: 279475
##
## Running Louvain algorithm...
## Maximum modularity in 10 random starts: 0.9114
## Number of communities: 13
## Elapsed time: 0 seconds
## Modularity Optimizer version 1.3.0 by Ludo Waltman and Nees Jan van Eck
##
## Number of nodes: 7470
## Number of edges: 279475
##
## Running Louvain algorithm...
## Maximum modularity in 10 random starts: 0.8979
## Number of communities: 12
## Elapsed time: 0 seconds
## Modularity Optimizer version 1.3.0 by Ludo Waltman and Nees Jan van Eck
##
## Number of nodes: 7470
## Number of edges: 279475
##
## Running Louvain algorithm...
## Maximum modularity in 10 random starts: 0.8845
## Number of communities: 12
## Elapsed time: 0 seconds
## Modularity Optimizer version 1.3.0 by Ludo Waltman and Nees Jan van Eck
##
## Number of nodes: 7470
## Number of edges: 279475
##
## Running Louvain algorithm...
## Maximum modularity in 10 random starts: 0.8713
## Number of communities: 13
## Elapsed time: 0 seconds
## Modularity Optimizer version 1.3.0 by Ludo Waltman and Nees Jan van Eck
##
## Number of nodes: 7470
## Number of edges: 279475
##
```

```
## Running Louvain algorithm...
## Maximum modularity in 10 random starts: 0.8584
## Number of communities: 14
## Elapsed time: 0 seconds
## Modularity Optimizer version 1.3.0 by Ludo Waltman and Nees Jan van Eck
##
## Number of nodes: 7470
## Number of edges: 279475
##
## Running Louvain algorithm...
## Maximum modularity in 10 random starts: 0.8457
## Number of communities: 14
## Elapsed time: 0 seconds
## Modularity Optimizer version 1.3.0 by Ludo Waltman and Nees Jan van Eck
##
## Number of nodes: 7470
## Number of edges: 279475
##
## Running Louvain algorithm...
## Maximum modularity in 10 random starts: 0.8338
## Number of communities: 15
## Elapsed time: 0 seconds
## Modularity Optimizer version 1.3.0 by Ludo Waltman and Nees Jan van Eck
##
## Number of nodes: 7470
## Number of edges: 279475
##
## Running Louvain algorithm...
## Maximum modularity in 10 random starts: 0.8225
## Number of communities: 16
## Elapsed time: 0 seconds
## Modularity Optimizer version 1.3.0 by Ludo Waltman and Nees Jan van Eck
##
## Number of nodes: 7470
## Number of edges: 279475
##
## Running Louvain algorithm...
## Maximum modularity in 10 random starts: 0.8119
## Number of communities: 17
## Elapsed time: 0 seconds
## Modularity Optimizer version 1.3.0 by Ludo Waltman and Nees Jan van Eck
##
## Number of nodes: 7470
## Number of edges: 279475
##
## Running Louvain algorithm...
## Maximum modularity in 10 random starts: 0.8035
## Number of communities: 19
## Elapsed time: 0 seconds
## Modularity Optimizer version 1.3.0 by Ludo Waltman and Nees Jan van Eck
##
## Number of nodes: 7470
## Number of edges: 279475
##
```

```
## Running Louvain algorithm...
## Maximum modularity in 10 random starts: 0.7958
## Number of communities: 19
## Elapsed time: 0 seconds
## Modularity Optimizer version 1.3.0 by Ludo Waltman and Nees Jan van Eck
##
## Number of nodes: 7470
## Number of edges: 279475
##
## Running Louvain algorithm...
## Maximum modularity in 10 random starts: 0.7891
## Number of communities: 18
## Elapsed time: 0 seconds
## Modularity Optimizer version 1.3.0 by Ludo Waltman and Nees Jan van Eck
##
## Number of nodes: 7470
## Number of edges: 279475
##
## Running Louvain algorithm...
## Maximum modularity in 10 random starts: 0.7817
## Number of communities: 19
## Elapsed time: 0 seconds
## Modularity Optimizer version 1.3.0 by Ludo Waltman and Nees Jan van Eck
##
## Number of nodes: 7470
## Number of edges: 279475
##
## Running Louvain algorithm...
## Maximum modularity in 10 random starts: 0.7744
## Number of communities: 19
## Elapsed time: 0 seconds
## Modularity Optimizer version 1.3.0 by Ludo Waltman and Nees Jan van Eck
##
## Number of nodes: 7470
## Number of edges: 279475
##
## Running Louvain algorithm...
## Maximum modularity in 10 random starts: 0.7676
## Number of communities: 21
## Elapsed time: 0 seconds
## Modularity Optimizer version 1.3.0 by Ludo Waltman and Nees Jan van Eck
##
## Number of nodes: 7470
## Number of edges: 279475
##
## Running Louvain algorithm...
## Maximum modularity in 10 random starts: 0.7609
## Number of communities: 21
## Elapsed time: 0 seconds
## Modularity Optimizer version 1.3.0 by Ludo Waltman and Nees Jan van Eck
##
## Number of nodes: 7470
## Number of edges: 279475
##
```

```
## Running Louvain algorithm...
## Maximum modularity in 10 random starts: 0.7544
## Number of communities: 22
## Elapsed time: 0 seconds
```

```
p=clustree(chick, prefix= "RNA_snn_res.")
print(p)
```

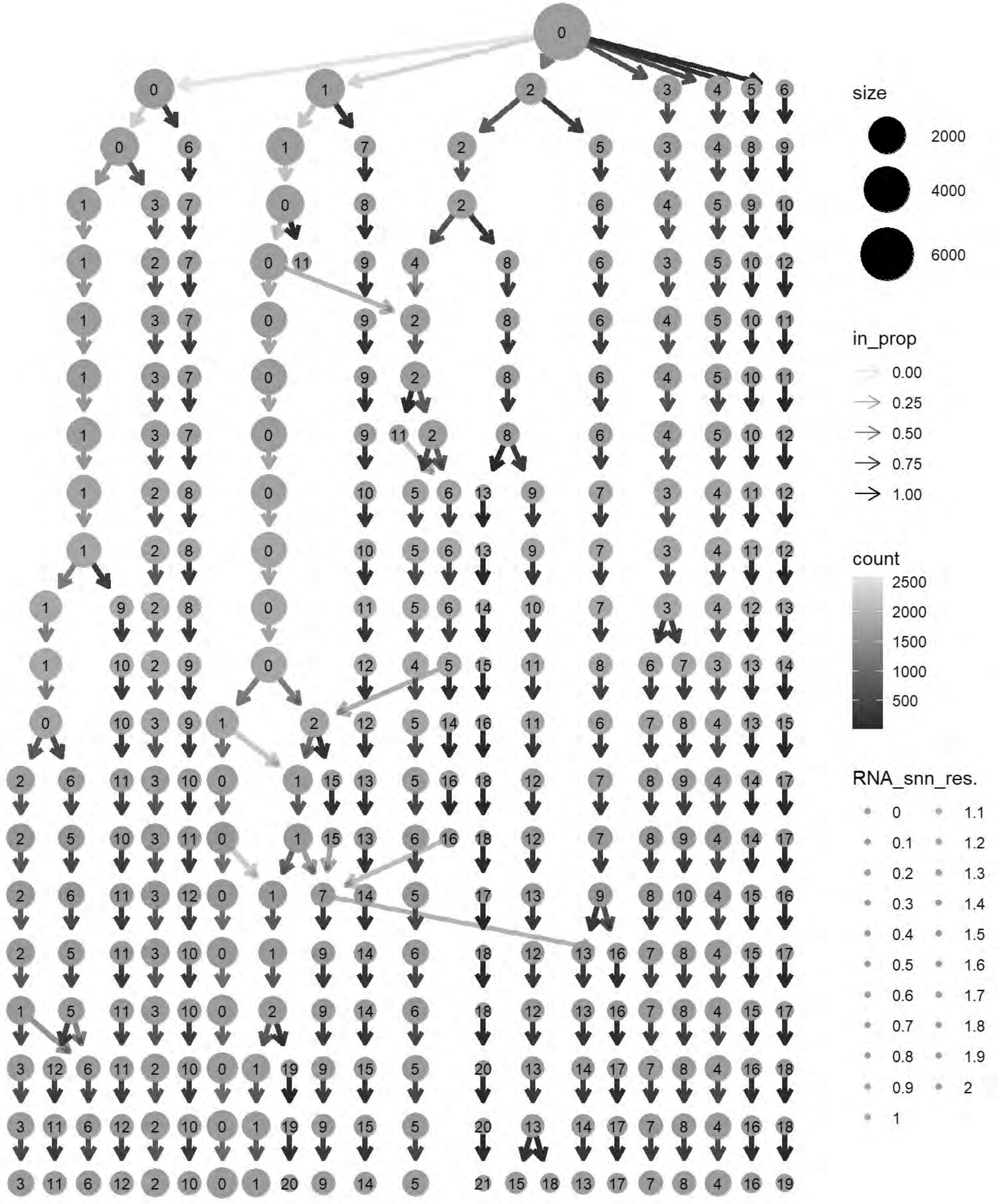
size
2000
4000
6000
in_prop
0.00
0.25
0.50
0.75
1.00
count
2500
2000
1500
1000
500
RNA_snn_res.
0
0.1
0.2
0.3
0.4
0.5
0.6
0.7
0.8
0.9
1
1.1
1.2
1.3
1.4
1.5
1.6
1.7
1.8
1.9
2

```
cols= grep("^RNA_snn_res.",names(chick@meta.data))
chick@meta.data[cols] <- NULL
chick <- FindClusters(chick, resolution = 0.5, verbose = TRUE)
```

```
## Modularity Optimizer version 1.3.0 by Ludo Waltman and Nees Jan van Eck
##
## Number of nodes: 7470
## Number of edges: 279475
##
## Running Louvain algorithm...
## Maximum modularity in 10 random starts: 0.8979
## Number of communities: 12
## Elapsed time: 0 seconds
```

```
p6=DimPlot(chick, label = TRUE)+NoLegend()
print(p6)
```

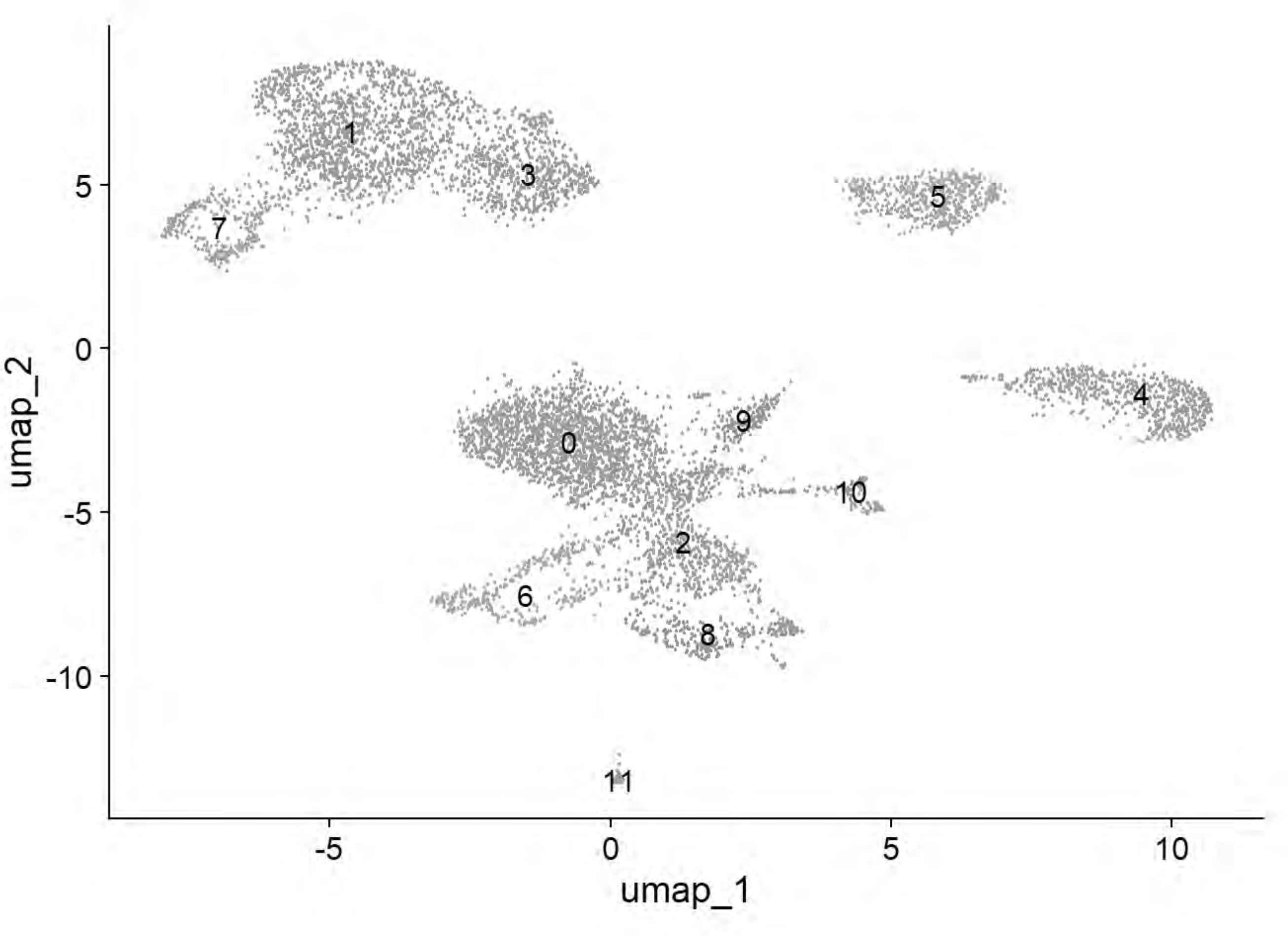


```
p6=DimPlot(chick, group.by = "Phase")
print(p6)
```

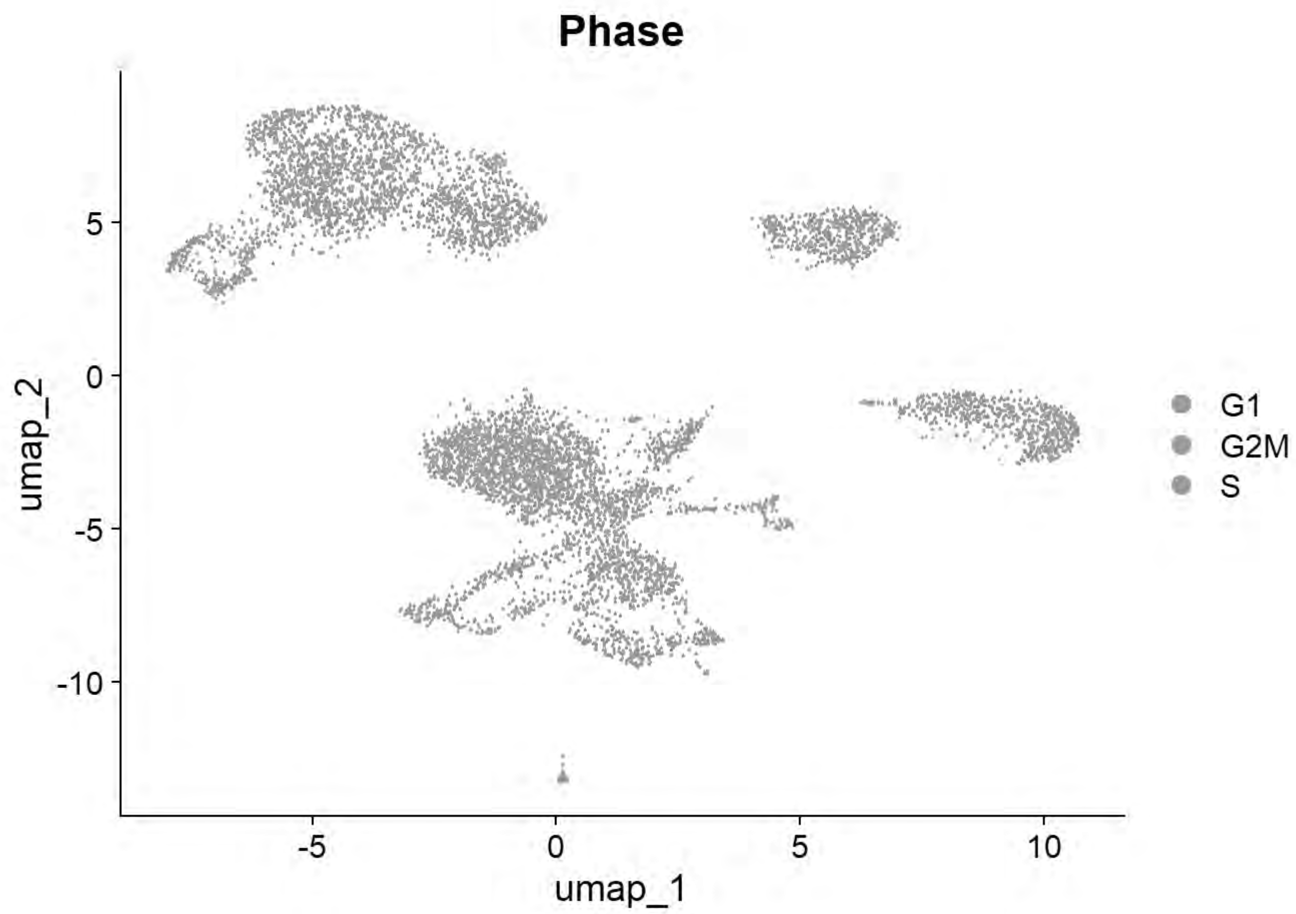


```
p6=DimPlot(chick, group.by = "sample_id")
print(p6)
```

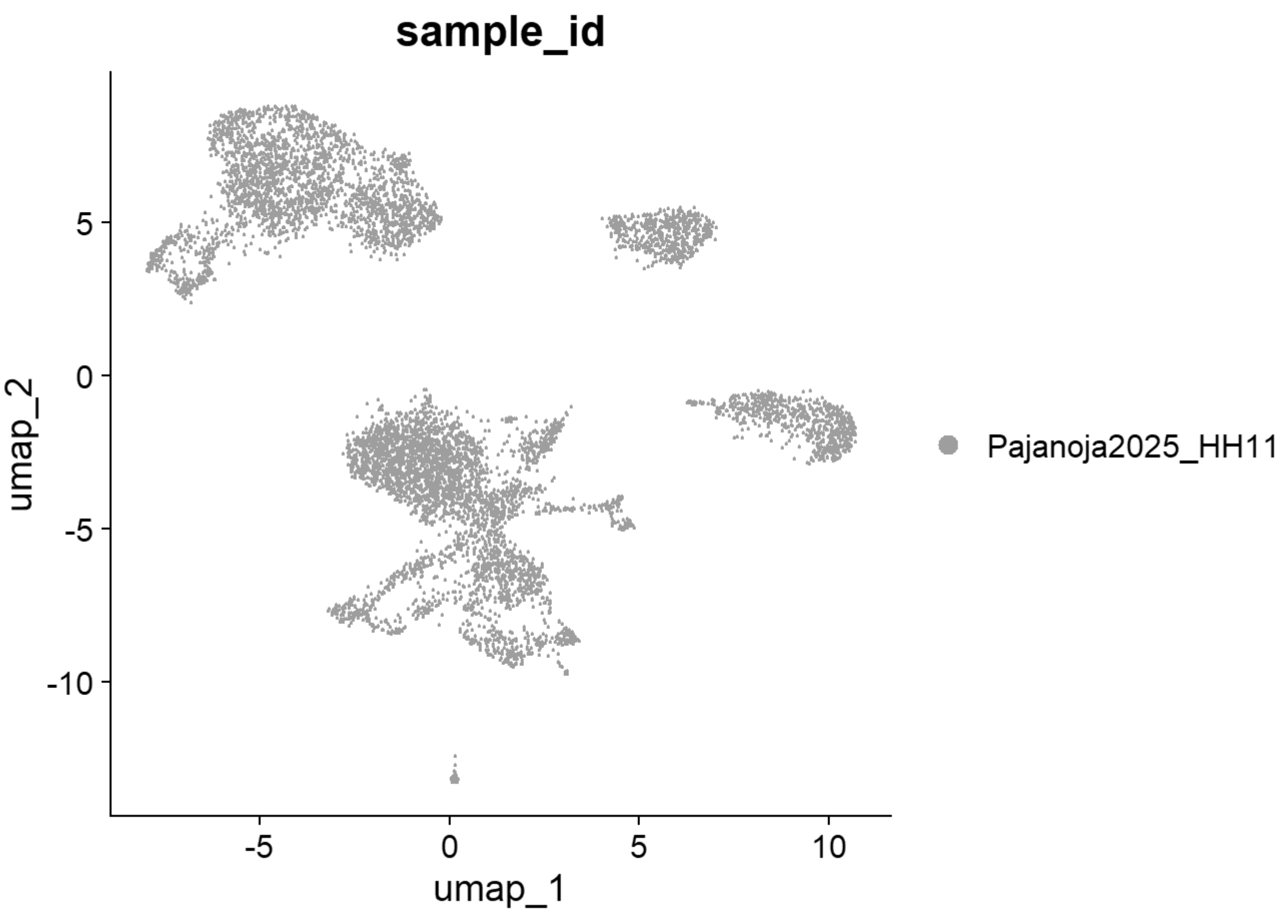


```
# Williams: Neural plate/tube
FeaturePlot(chick, features = c("CLDN1","FRZB","DNMT3A"))
```

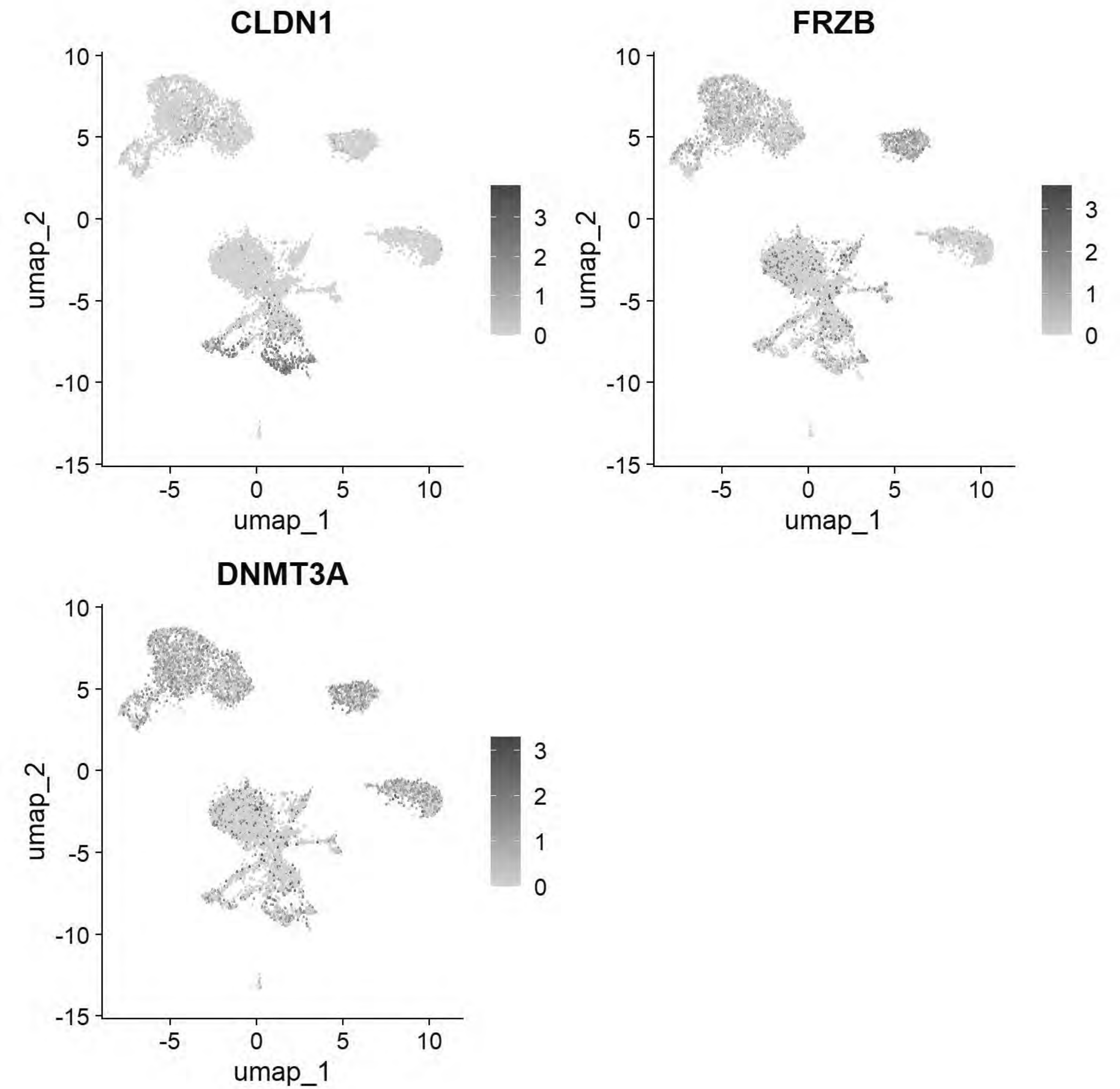


```
# Williams: Non-Neural Ectoderm
FeaturePlot(chick, features = c("DLX5","TFAP2A","ASTL","PAX6"))
```

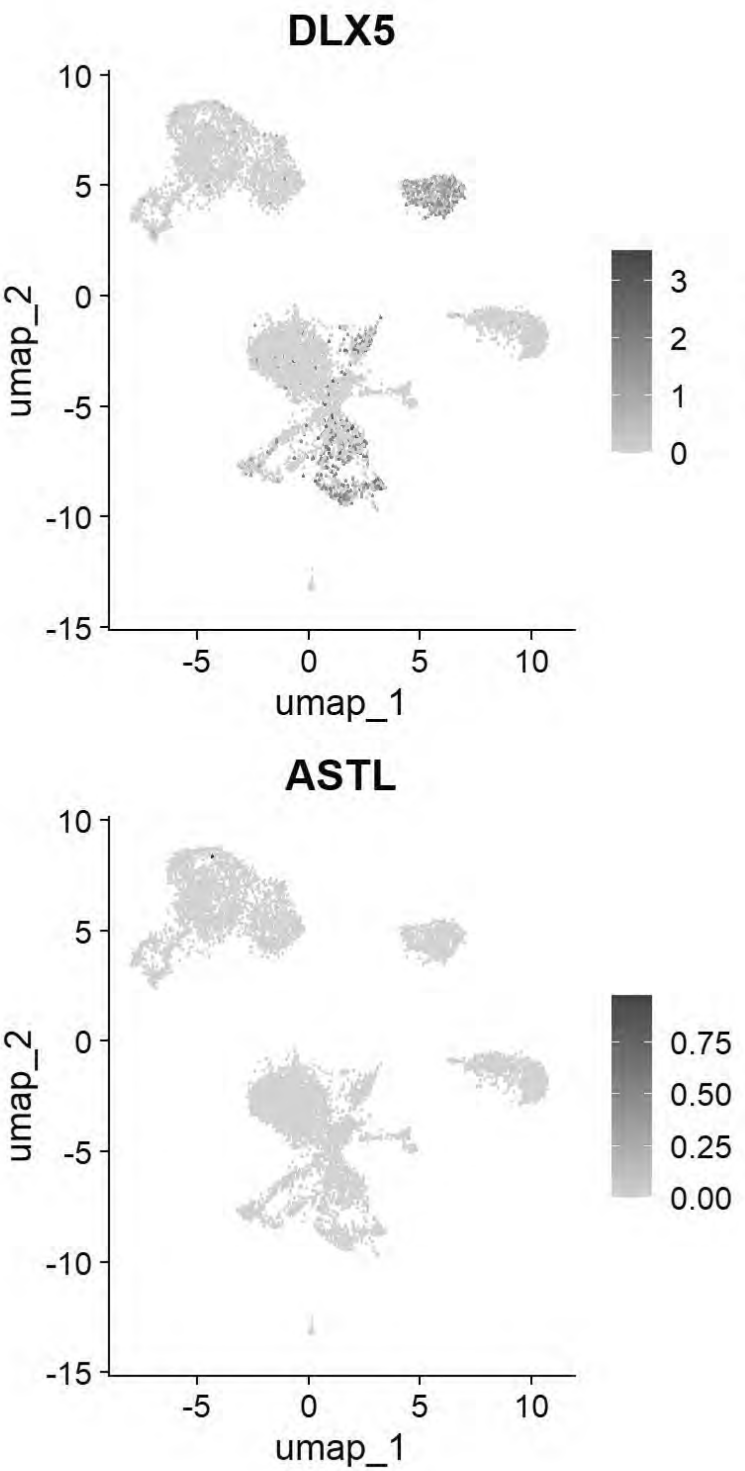


```
# Williams: Ectoderm (Neural Plate)
FeaturePlot(chick, features = c("SOX2","SOX3","SOX21","FRZB","SFRP2"))
```

```
## Warning: All cells have the same value (0) of "SOX3"
```

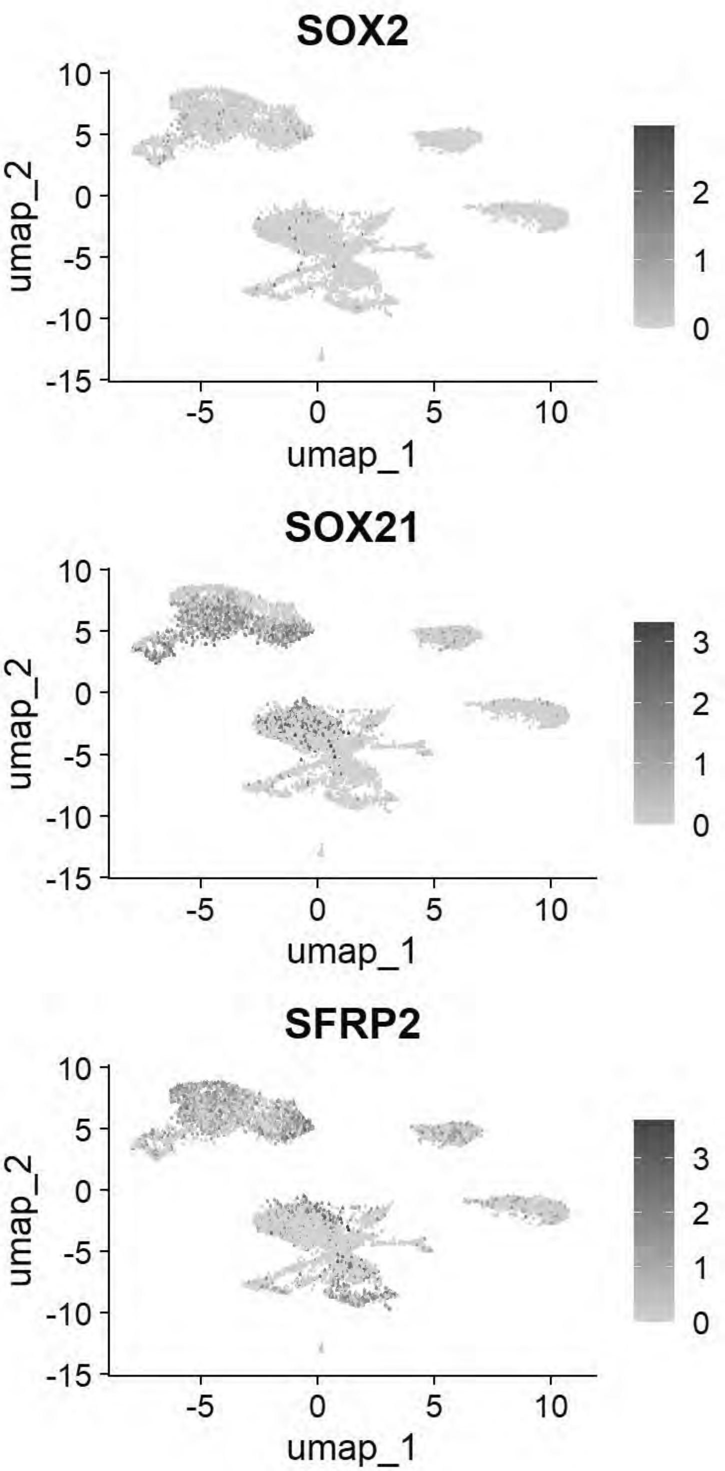


```r
# Williams: Ectoderm (Neural Plate Border)
FeaturePlot(chick, features = c("PAX7","TFAP2A","DLX5","BMP4","MSX1","DRAXIN","TFAP2B"))
```

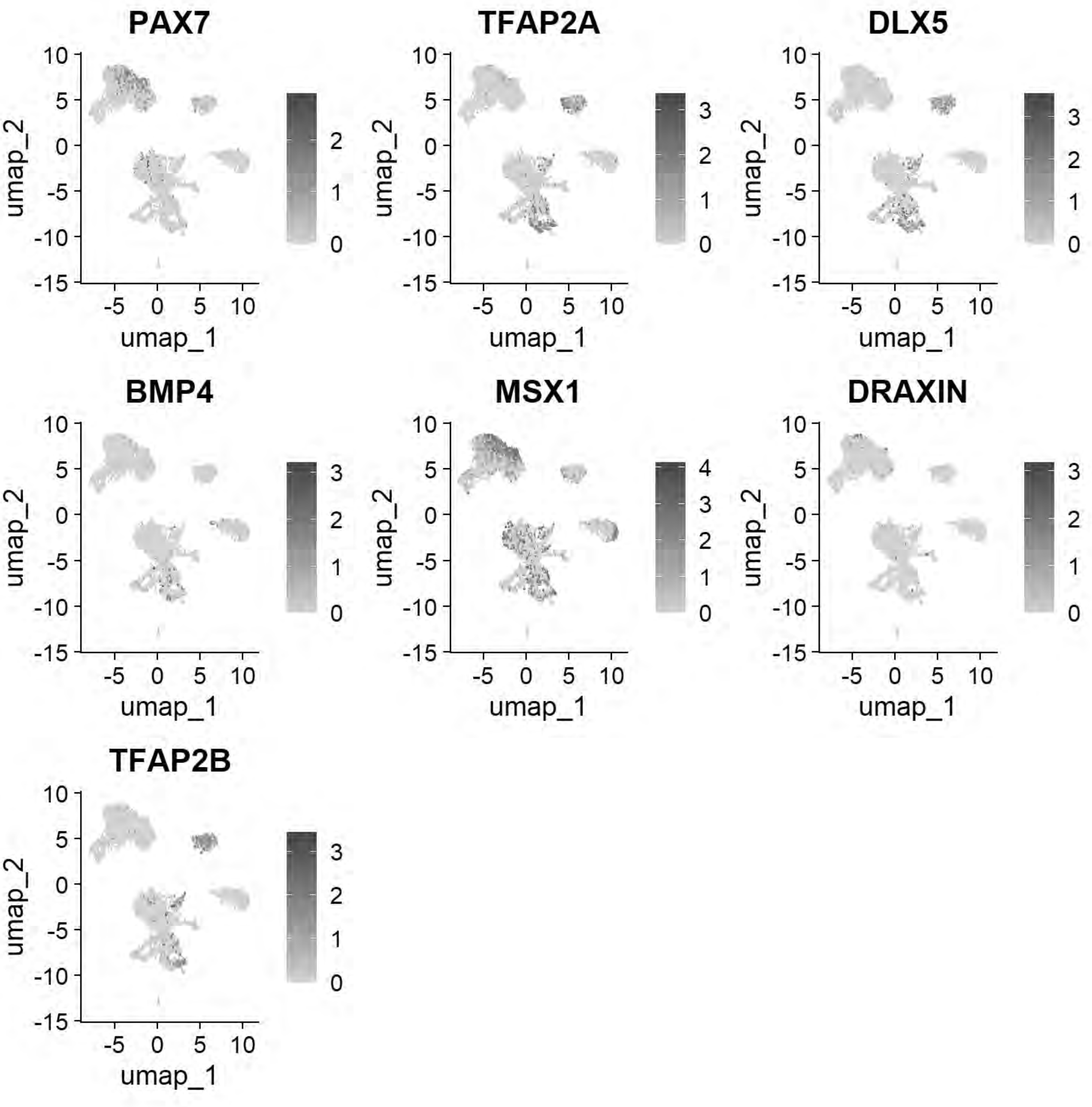


```
# Williams: Posterior Lateral Plate Mesoderm
FeaturePlot(chick, features = c("GATA2","HOXB5","CDX4"))
```

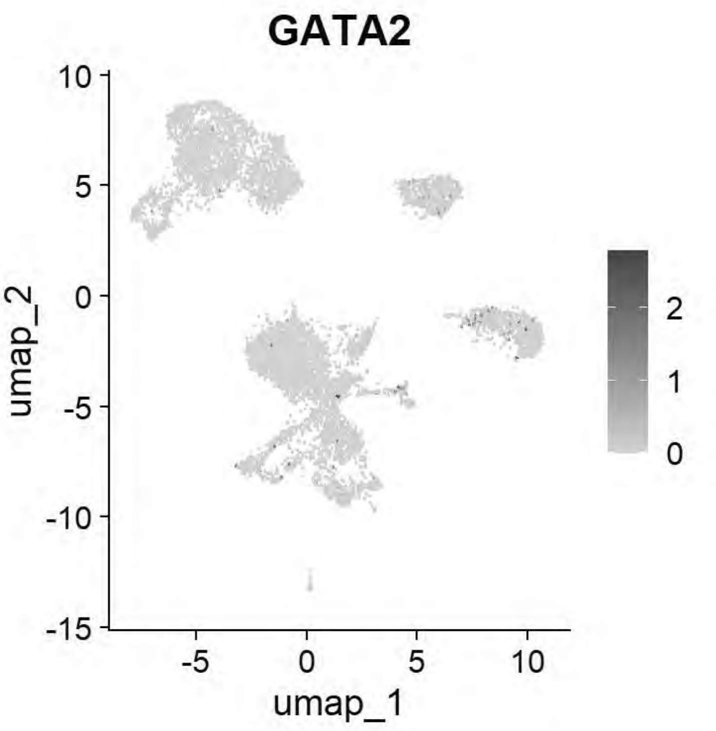


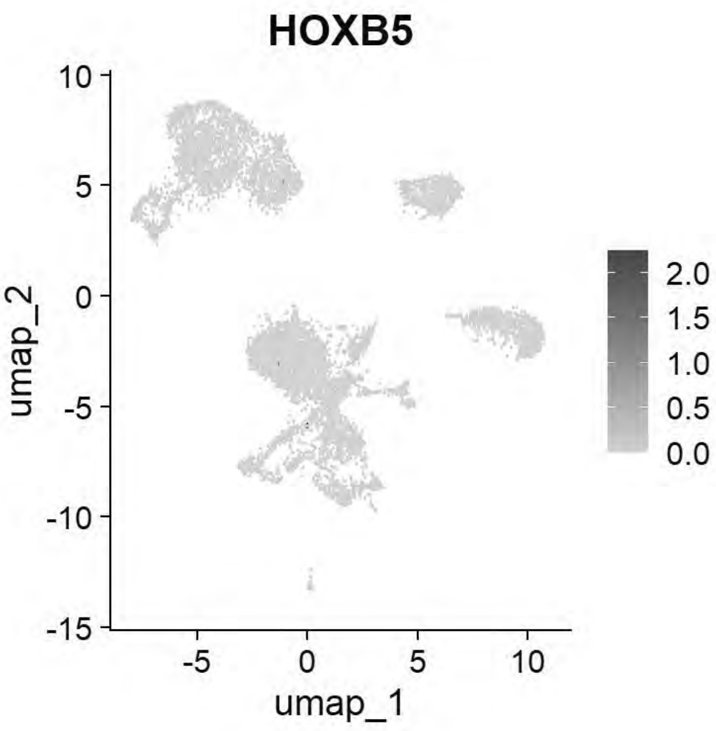


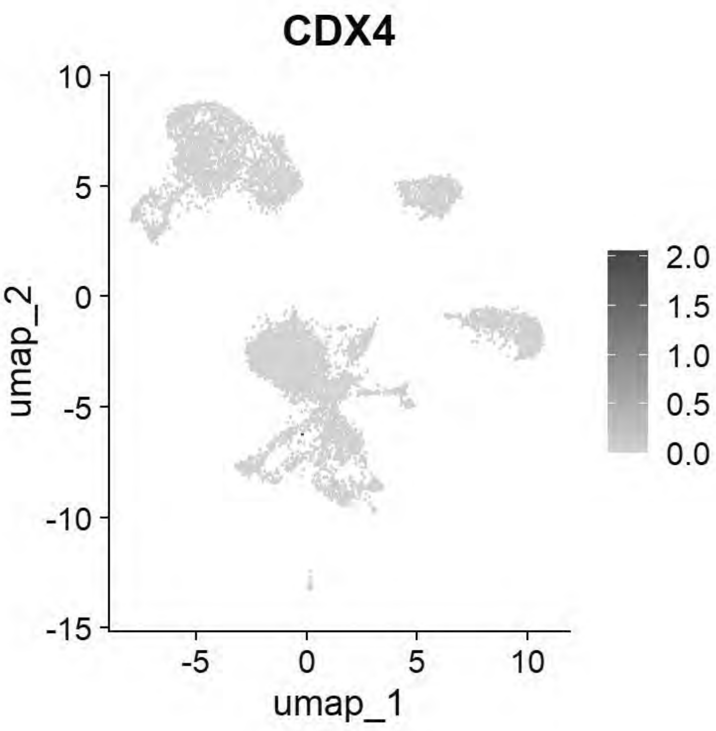


```
# Williams: Paraxial Mesoderm
FeaturePlot(chick, features = c("MSGN1","MESP1","MEOX1"))
```

```
## Warning: All cells have the same value (0) of "MESP1"
```

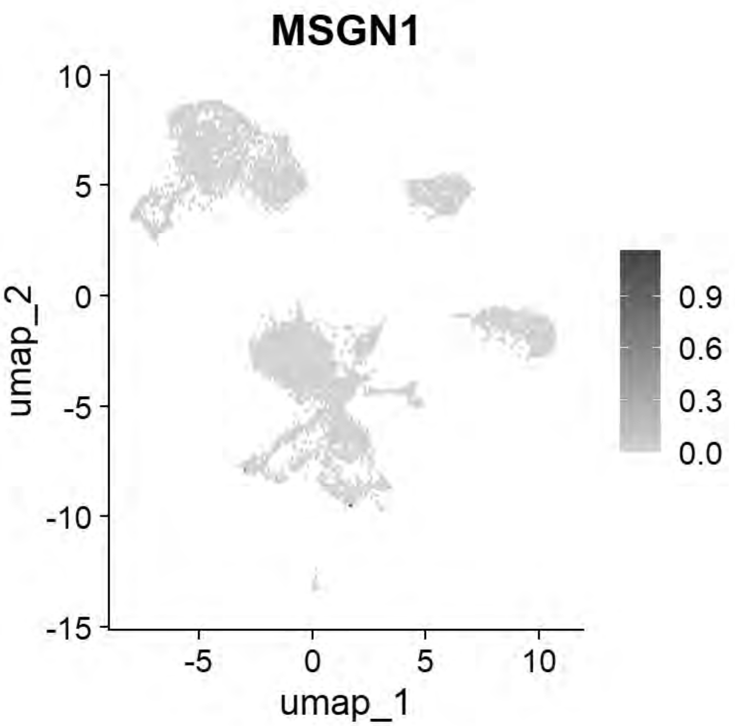


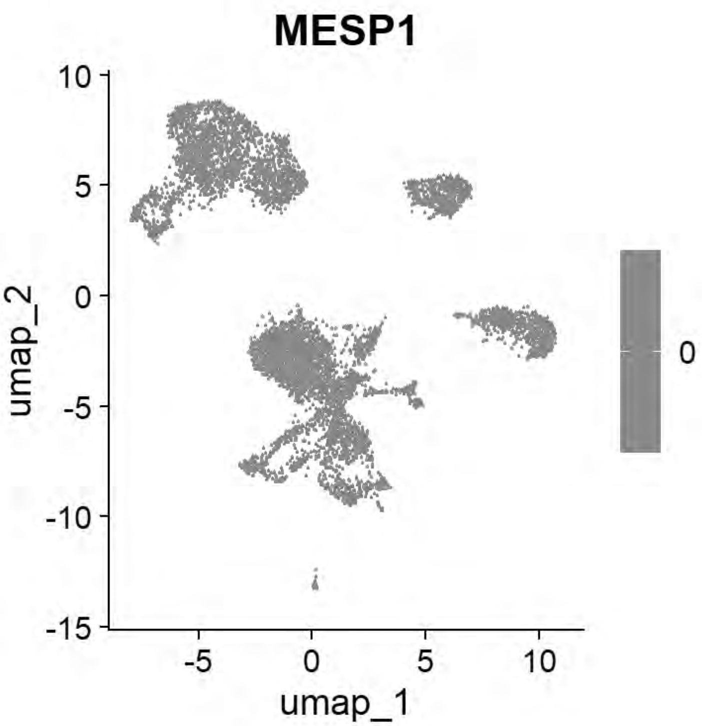


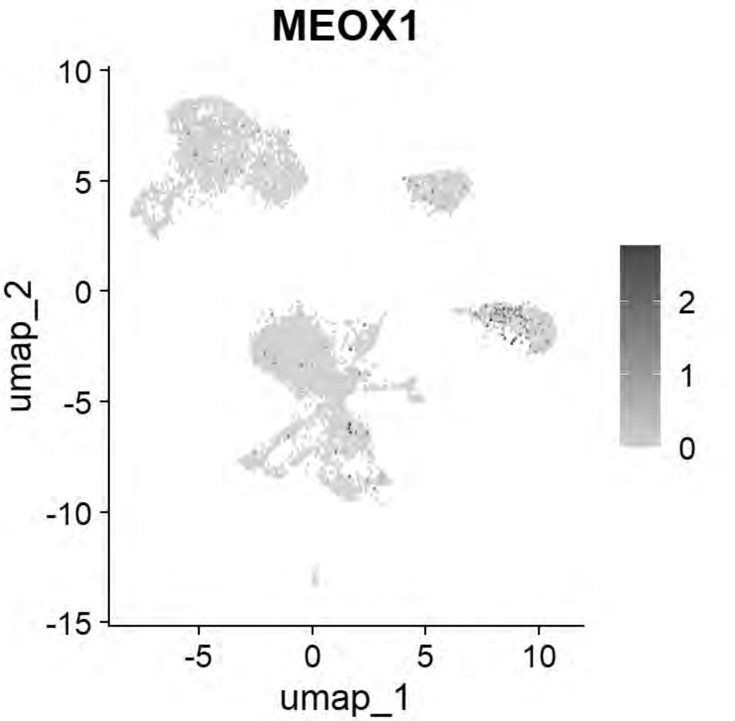


```
# Williams: Lateral Plate Mesoderm
FeaturePlot(chick, features = c("PITX2","ALX1","OLFML3","SIX1","TWIST1"))
```

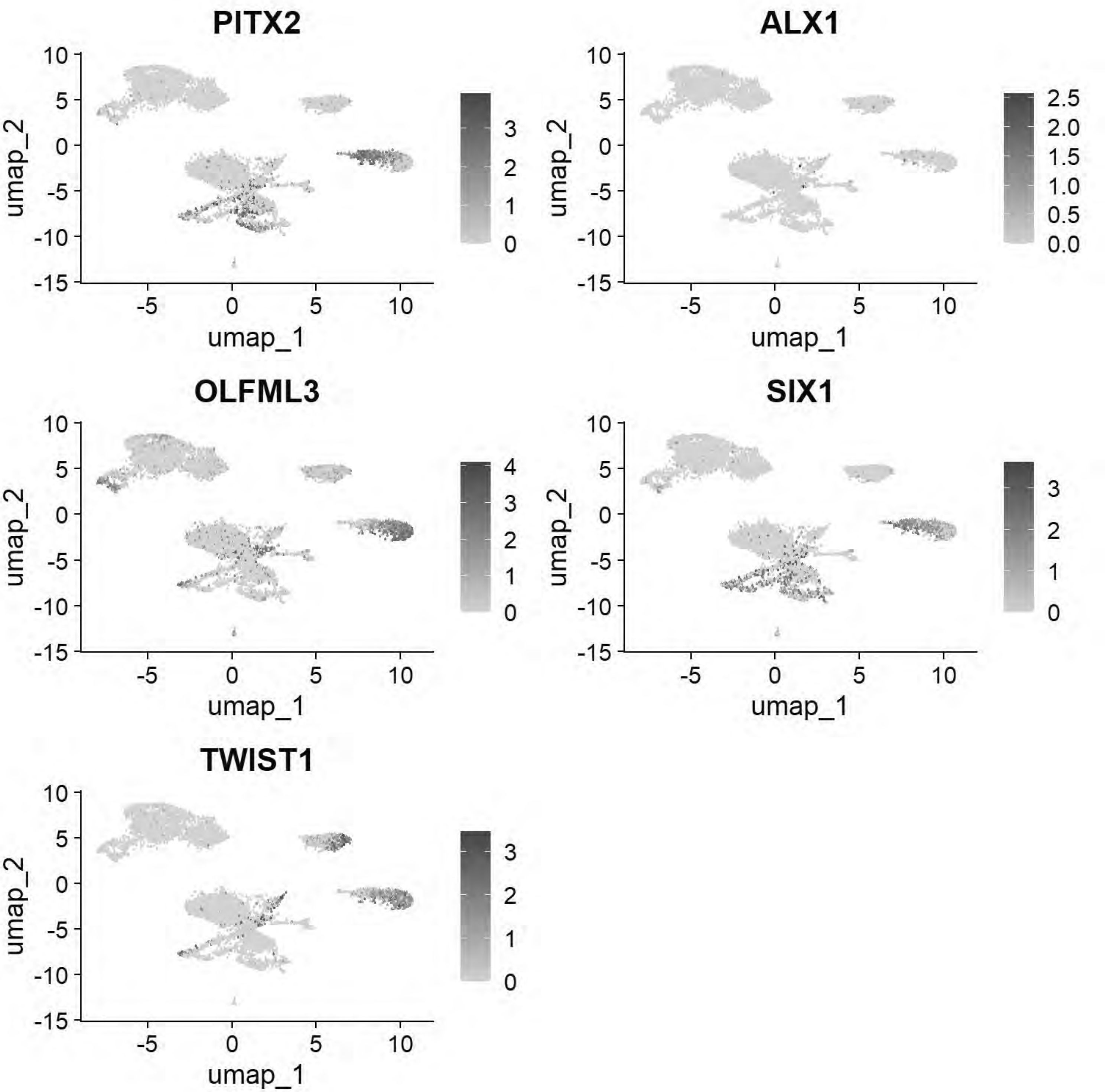


```
# Williams: Head Mesenchyme
FeaturePlot(chick, features = c("TCF21","GATA5","LMO2","ETS1","KDR"))
```

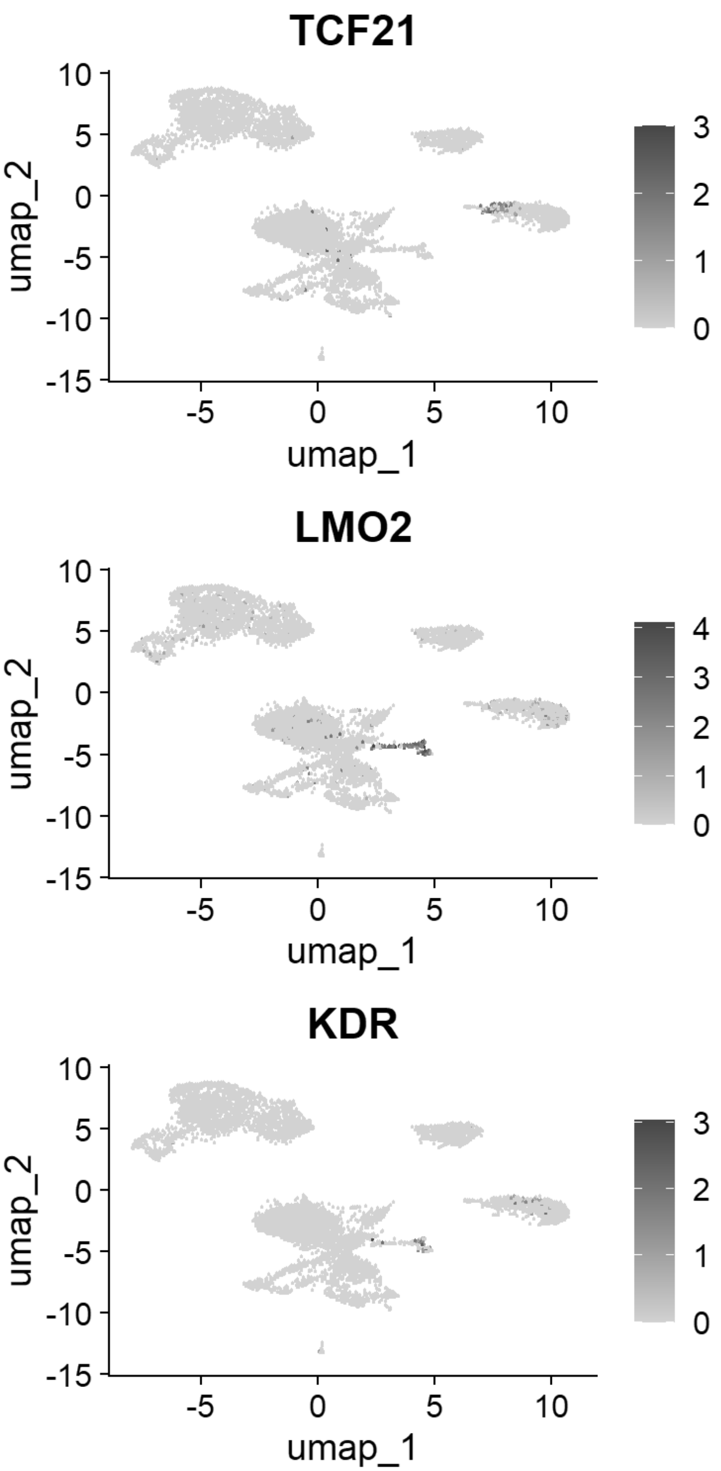


```
# Williams: Hensen's Node & Primitive Streak
FeaturePlot(chick, features = c("DLL1","FGF8","NOTO","CHRD"))
```

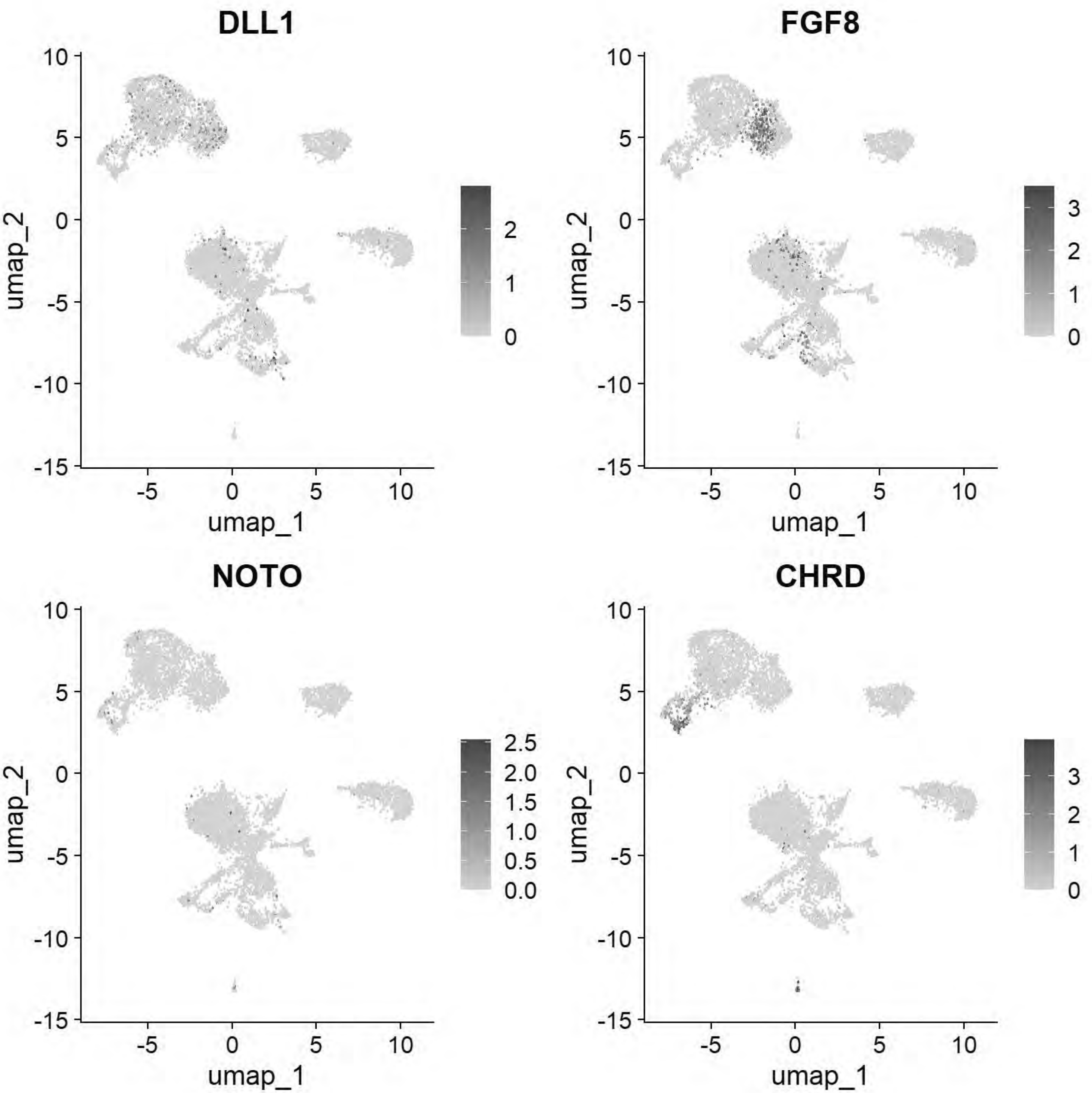


```
# Williams: Endoderm
FeaturePlot(chick, features = c("SOX17","FOXA2","CXCR4"))
```

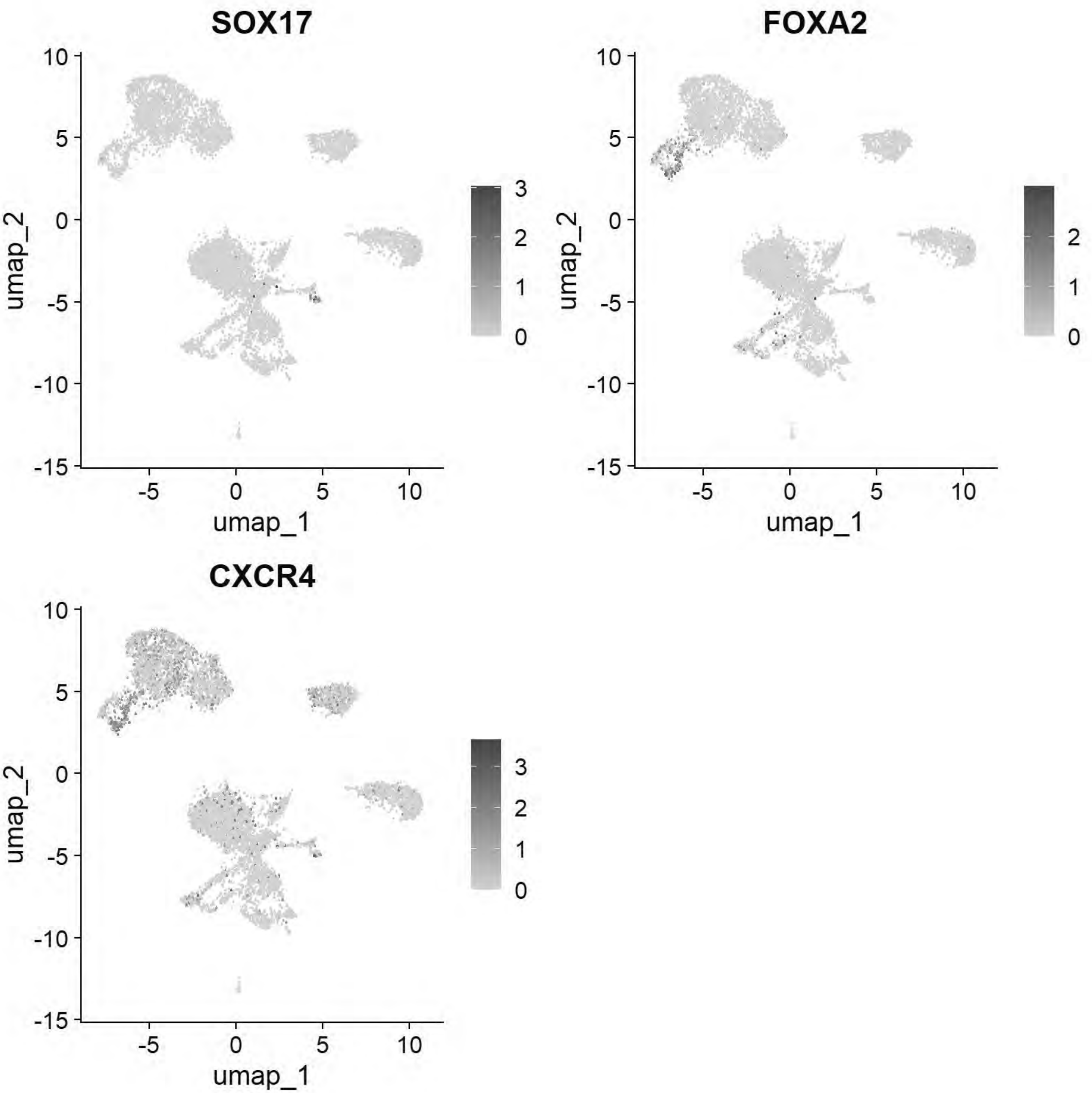


```
# Pajanoja: Ectoderm
FeaturePlot(chick, features = c("TFAP2A","DLX5","CLDN1","SOX2","NESTIN","MYCN"))
```

```
## Warning: The following requested variables were not found: NESTIN
```

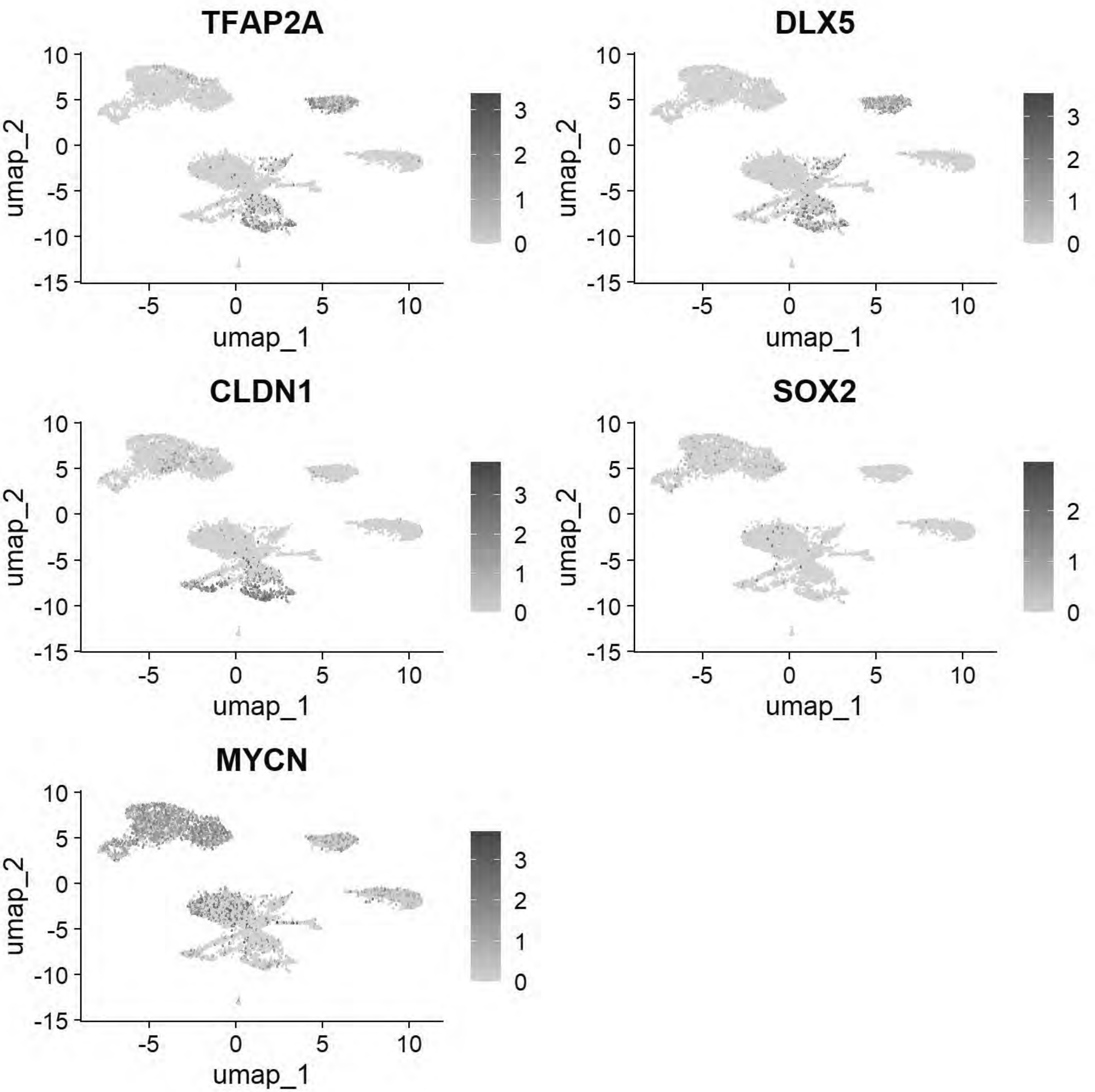


```
# Pajanoja: Endoderm
FeaturePlot(chick, features = c("SOX17","KRT17","HHEX"))
```

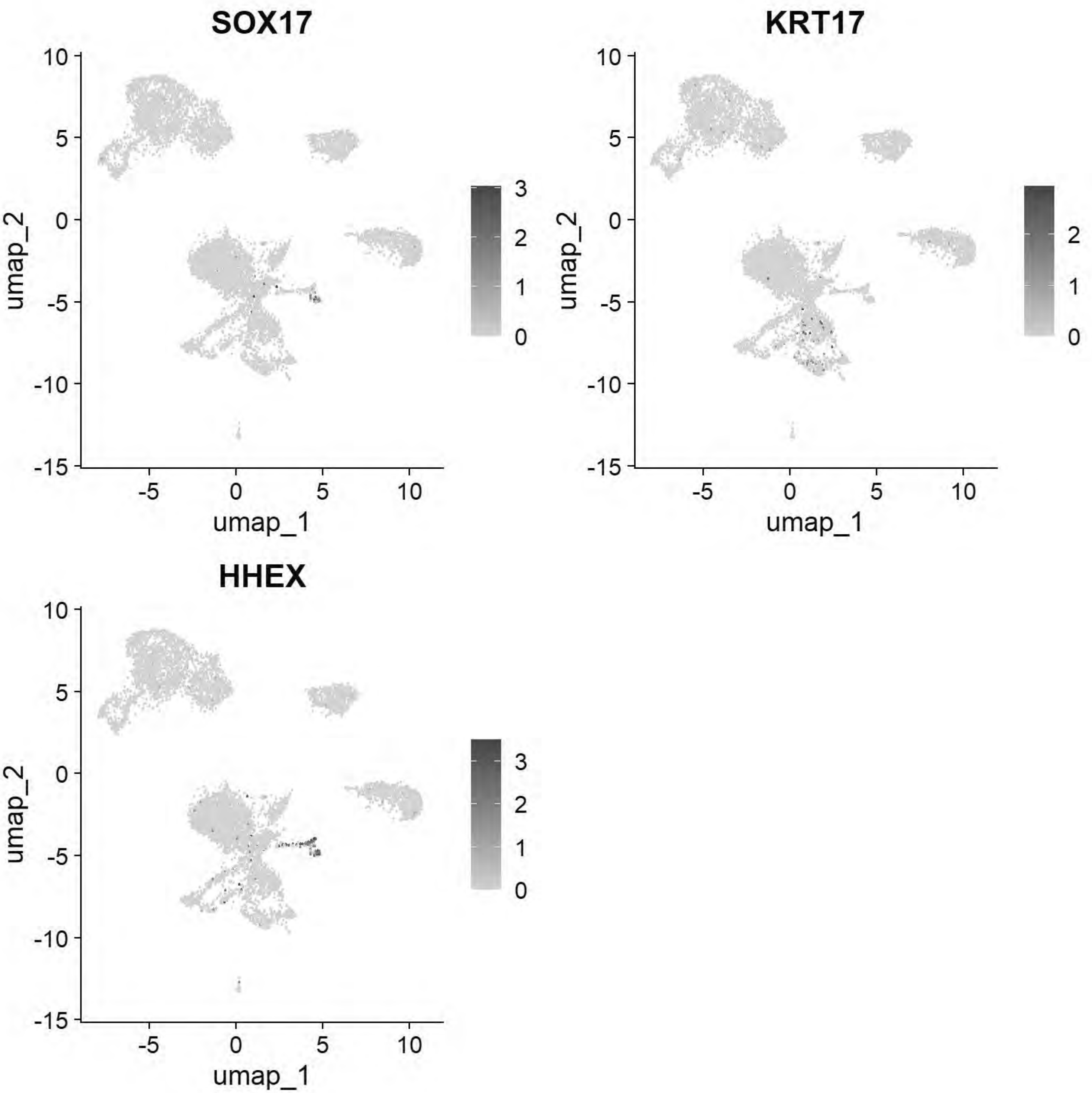


```
# Pajanoja: Mesoderm
FeaturePlot(chick, features = c("ALX1","TWIST1","HAND2","PITX2"))
```

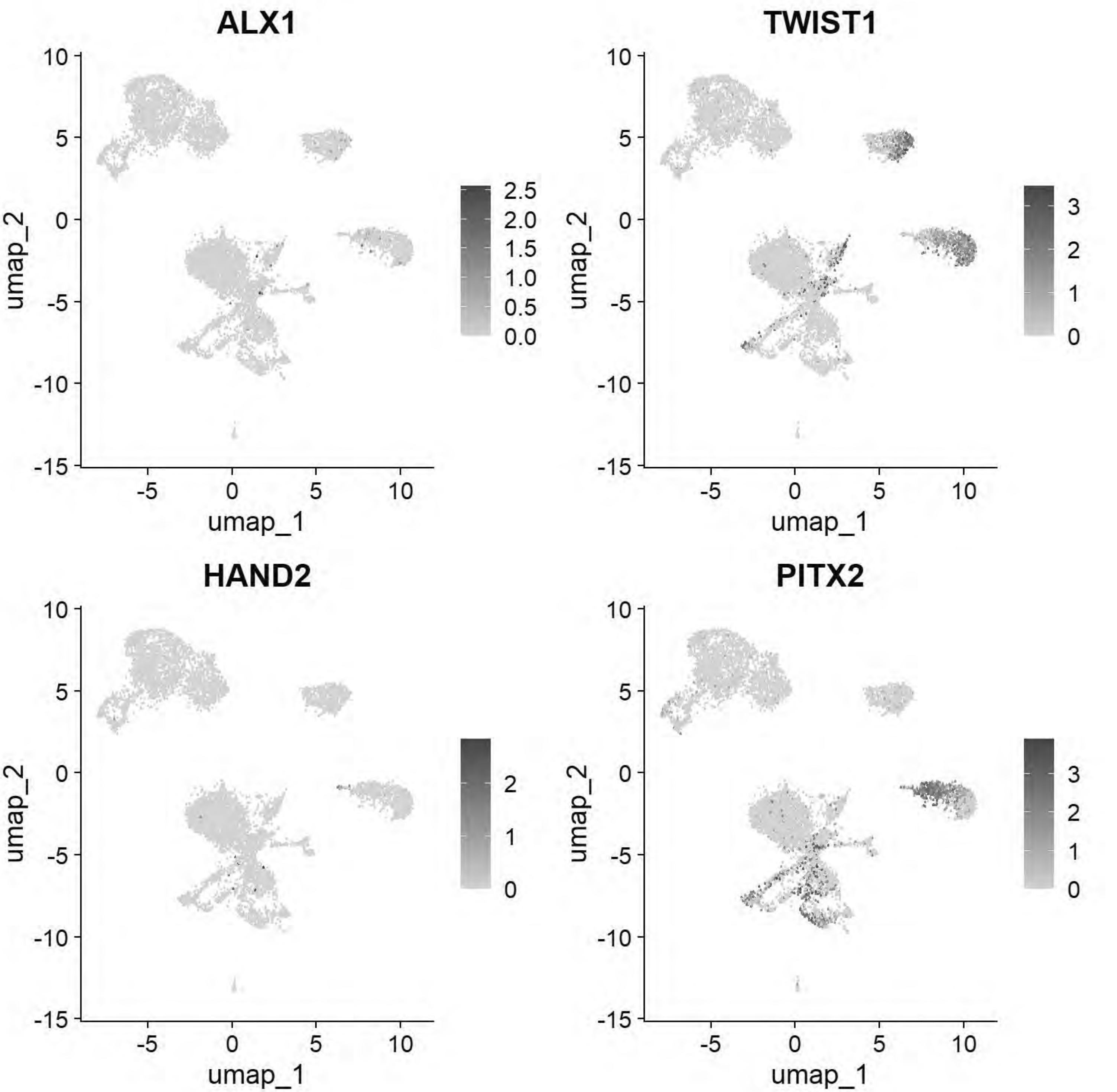


```
# Pajanoja: Notochord
FeaturePlot(chick, features = c("CHRD","TBXT","NOTO"))
```

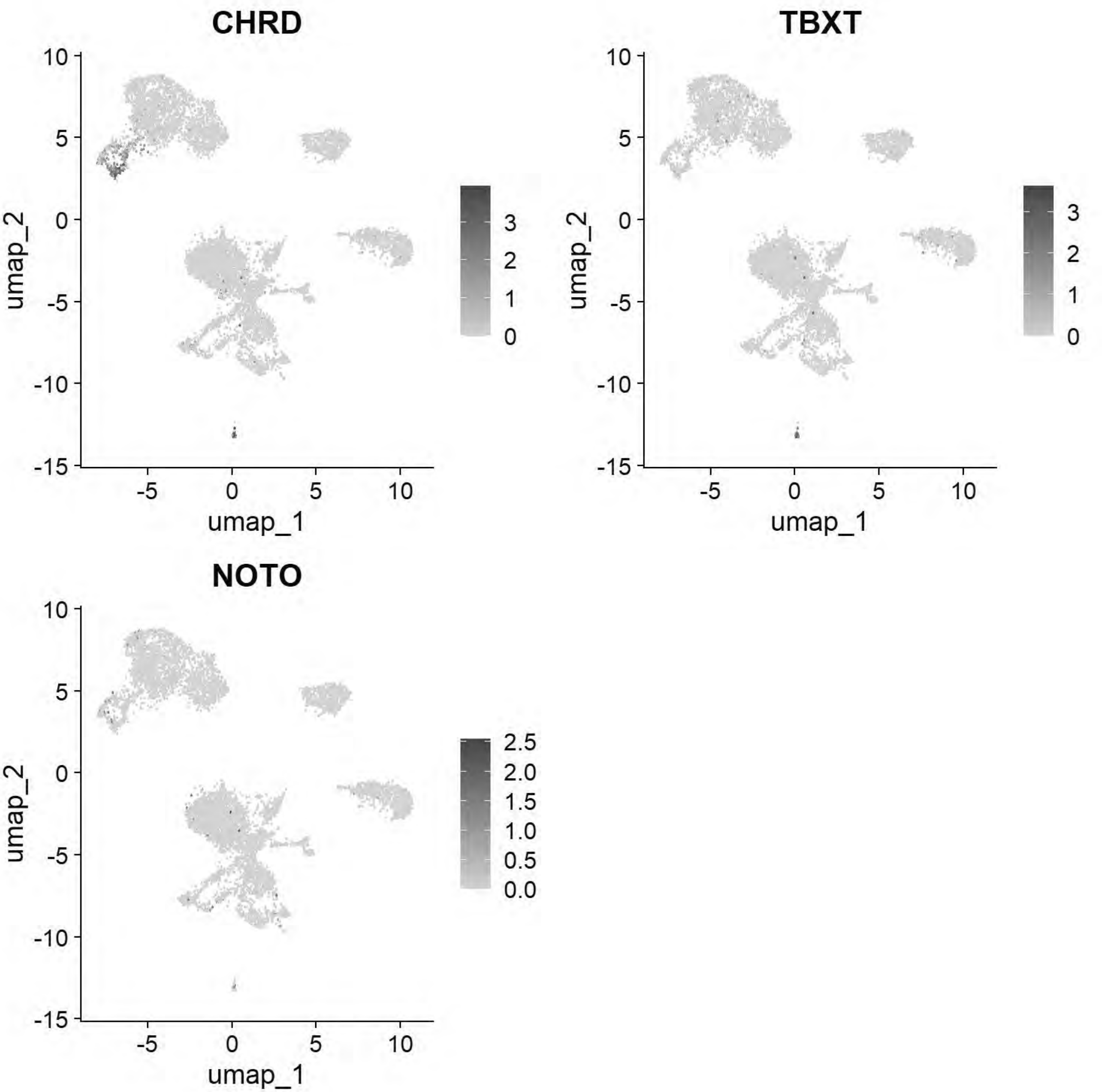


```
# Pajanoja: Ventral Neural Tube
FeaturePlot(chick, features=c("SHH","FOXA1","FOXA2"))
```

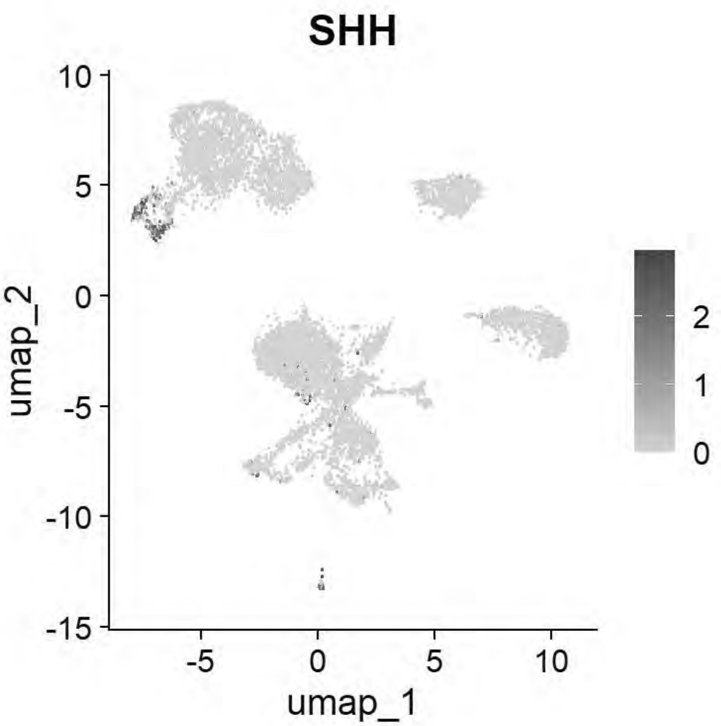


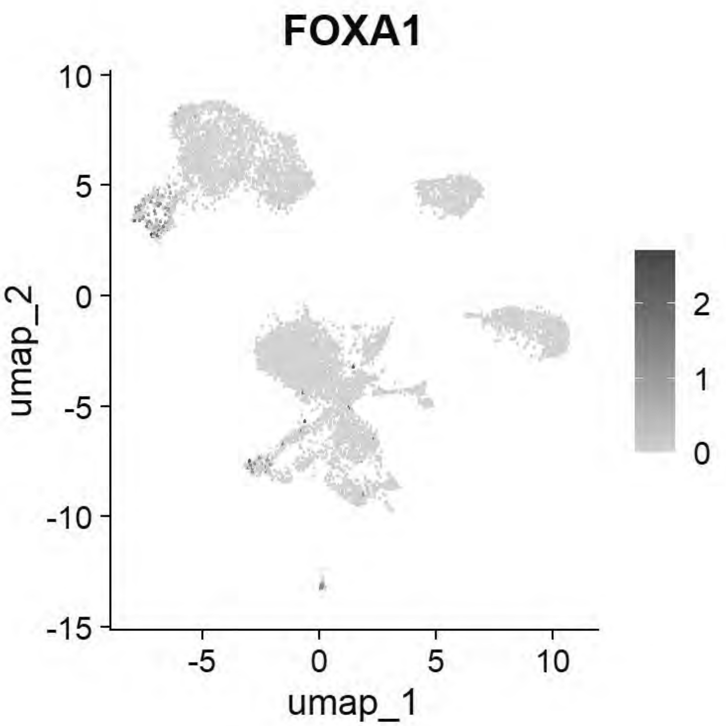


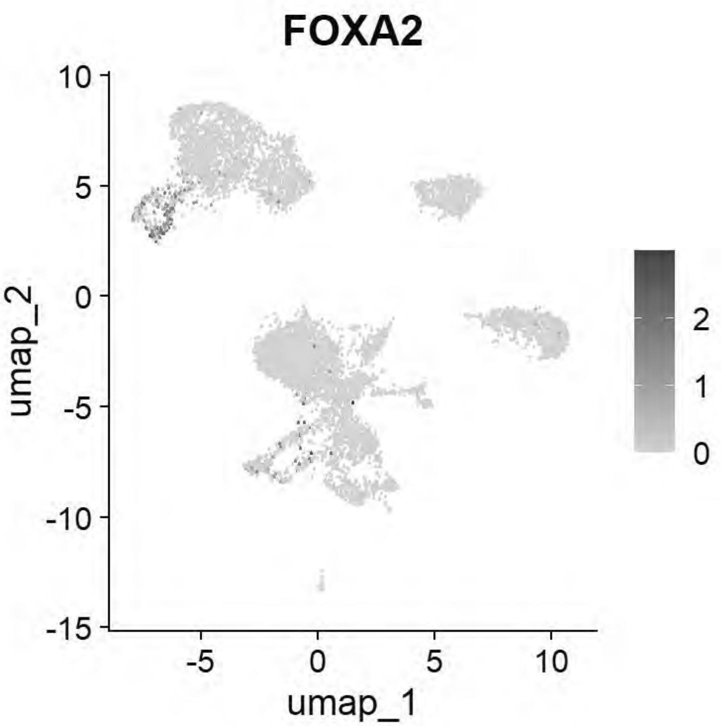


```
# Pajanoja: Neural Tube
FeaturePlot(chick, features=c("CDH2","HES5","MYCN","PAX2","NESTIN"))
```

```
## Warning: The following requested variables were not found: NESTIN
```

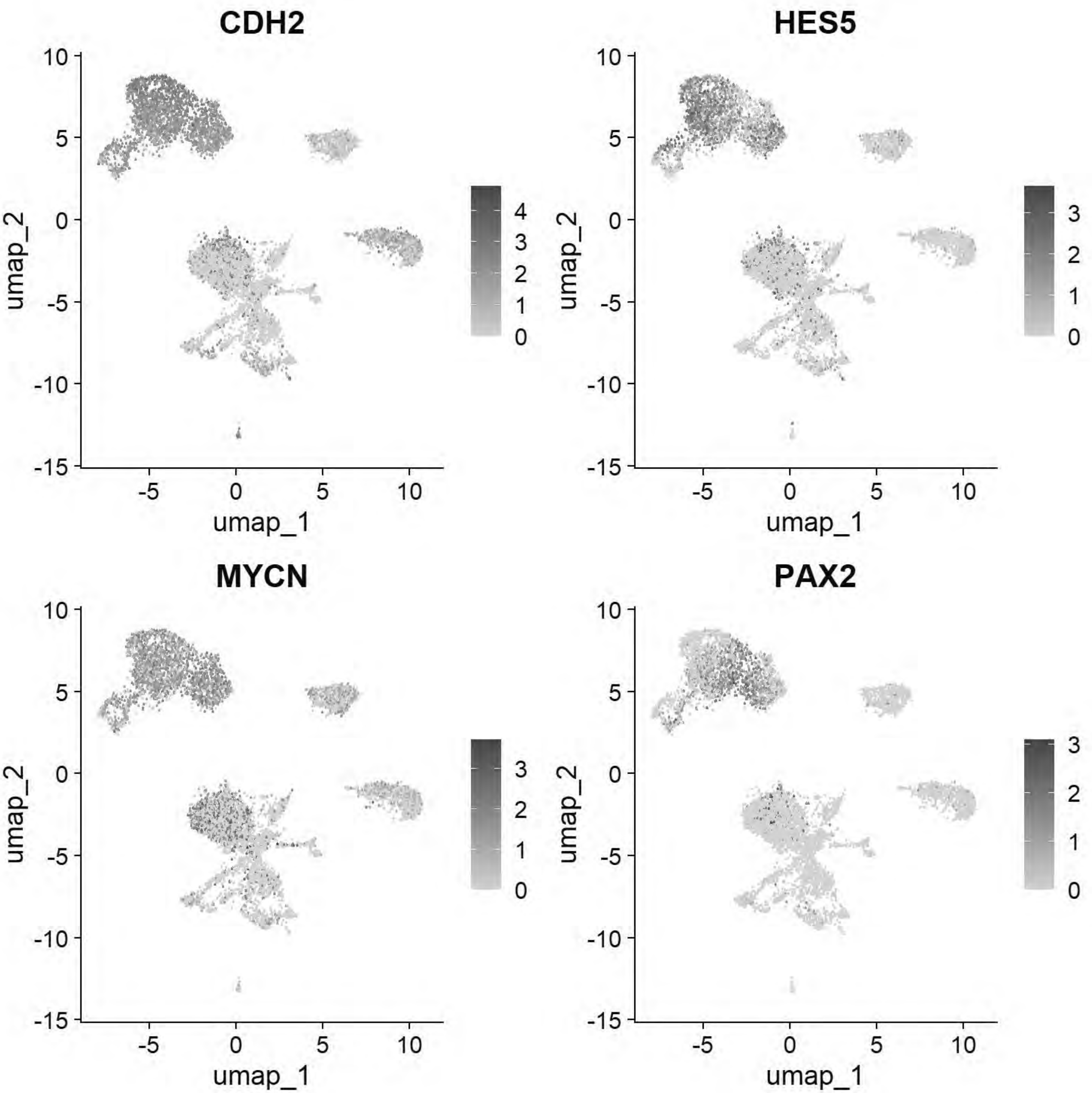


```
# Pajanoja: Non-Neural Ectoderm/Placode
FeaturePlot(chick, features=c("KRT24","KRT18","DLX5","SIX1","EYA2","TFAP2A"))
```

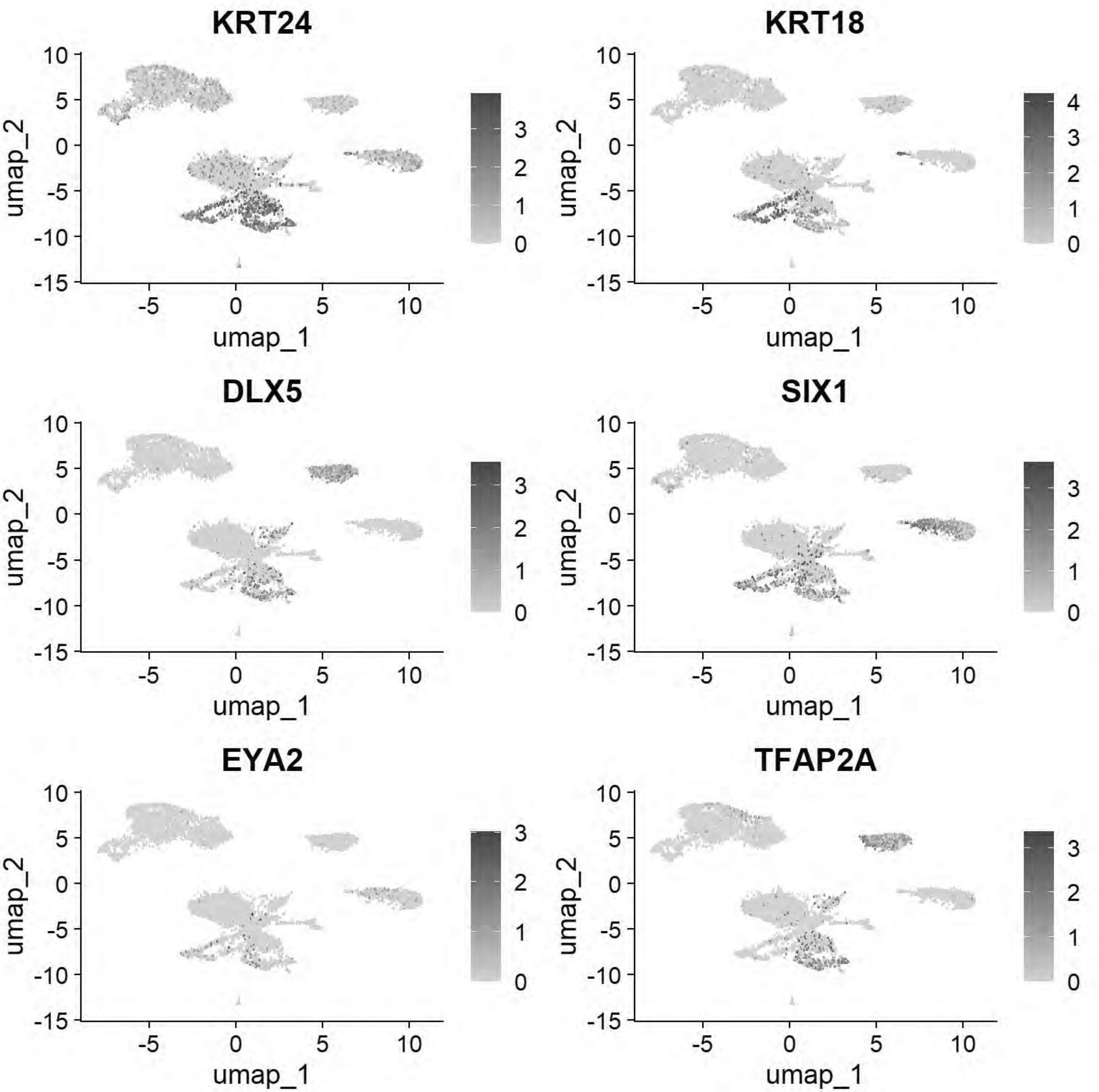


```
# Pajanoja: Neural crest
FeaturePlot(chick, features=c("FOXD3","TFAP2B","SOX9","ETS1","SNAI2"))
```

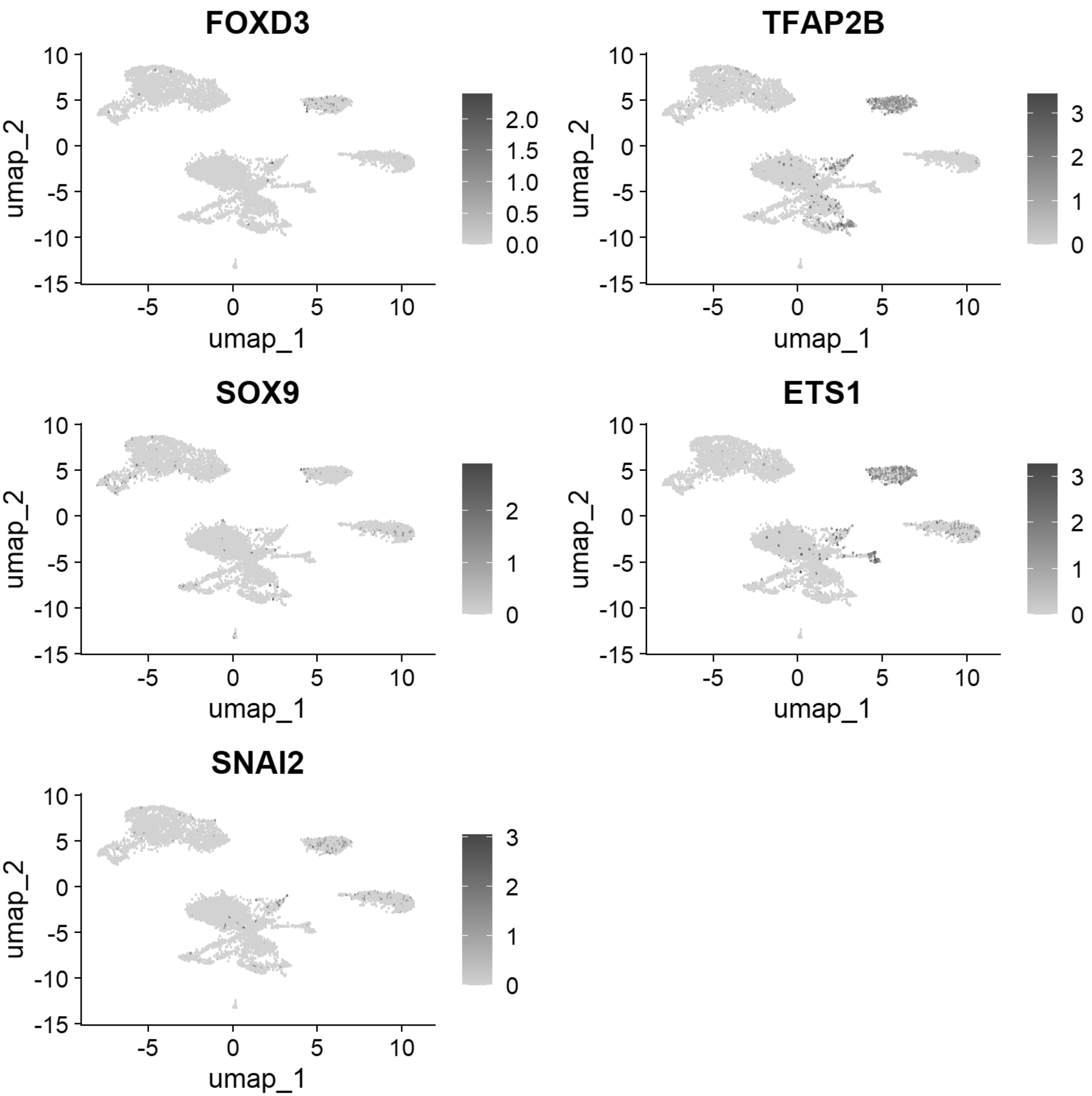


```
gc()
```

```
##              used   (Mb) gc trigger    (Mb)   max used    (Mb)
## Ncells   5082726  271.5   13651909   729.1   21331105  1139.3
## Vcells 900642110 6871.4 2342481578 17871.8 4574774951 34902.8
```

```
chick <- JoinLayers(chick)
chick.markers <- FindAllMarkers(chick, min.pct = 0.5, logfc.threshold = 0.5)
```

```
## Calculating cluster 0
```

```
## Calculating cluster 1
```

```
## Calculating cluster 2
```

```
## Calculating cluster 3
```

```
## Calculating cluster 4
```

```
## Calculating cluster 5
```

```
## Calculating cluster 6
```

```
## Calculating cluster 7
```

```
## Calculating cluster 8
```

```
## Calculating cluster 9
```

```
## Calculating cluster 10
```

```
## Calculating cluster 11
```

```
chick.markers %>%
  group_by(cluster) %>%
  top_n(n = 5, wt = avg_log2FC) -> top5
DotPlot(chick, features = unique(top5$gene))+coord_flip()
```

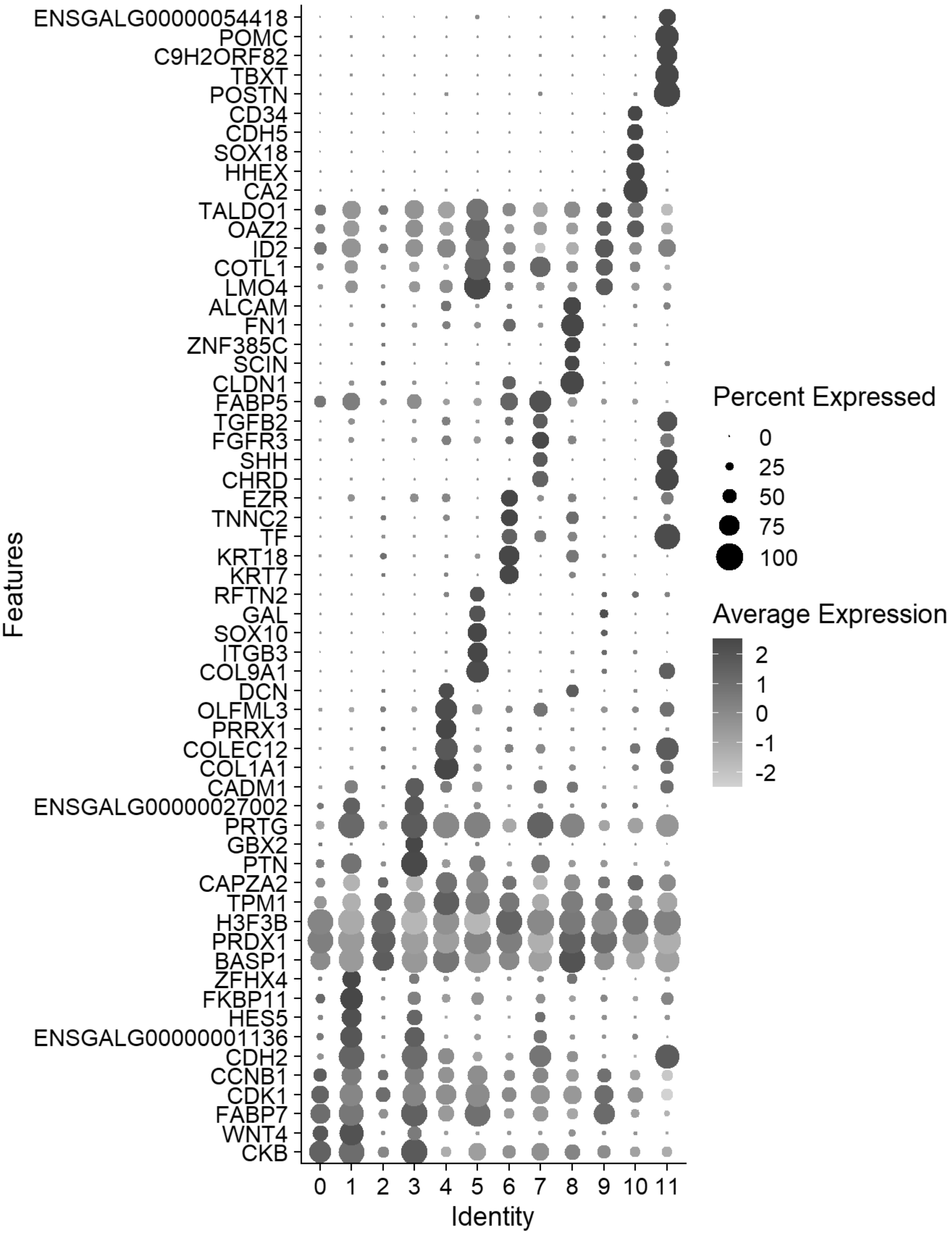


```
gc()
```

```
##             used   (Mb) gc trigger    (Mb)   max used    (Mb)
## Ncells   4972325  265.6   13651909   729.1   21331105  1139.3
```

```
## Vcells 951016888 7255.7 2342481578 17871.8 4574774951 34902.8
```

```
cluster_map <- c("0"="Neural",
                 "1"="Def Neural 1",
                 "2"="Ectoderm Mixed",
                 "3"="Def Neural 2",
                 "4"="Lateral Plate Mesoderm",
                 "5"="Neural Crest 1",
                 "6"="Placode",
                 "7"="vNeural Tube",
                 "8"="NNE",
                 "9"="Neural Crest 2",
                 "10"="Hematogenic",
                 "11"="Notochord")
chick$celltype1 <- plyr::mapvalues(chick$seurat_clusters, from = names(cluster_map), to = clus
ter_map)

chick <- JoinLayers(chick)
Idents(chick) <- "celltype1" # Setting active identity ensures markers are calculated by cell
type
global_markers <- FindAllMarkers(chick, min.pct = 0.5, logfc.threshold = 0.5)
```

```
## Calculating cluster Neural
```

```
## Calculating cluster Def Neural 1
```

```
## Calculating cluster Ectoderm Mixed
```

```
## Calculating cluster Def Neural 2
```

```
## Calculating cluster Lateral Plate Mesoderm
```

```
## Calculating cluster Neural Crest 1
```

```
## Calculating cluster Placode
```

```
## Calculating cluster vNeural Tube
```

```
## Calculating cluster NNE
```

```
## Calculating cluster Neural Crest 2
```

```
## Calculating cluster Hematogenic
```

```
## Calculating cluster Notochord
```

```
write.csv(global_markers, "E:/Rogers Lab/Rogers NC EMT/Supplementary file 6_HH11_Markers_Annotated.csv", row.names = FALSE)

celltype_colors <- c("Neural"="#41B6C4",
                 "Def Neural 1"="#2171B5",
                 "Ectoderm Mixed"="#BFD3E6",
                 "Def Neural 2"="#1D91C0",
                 "Lateral Plate Mesoderm"="#CB181D",
                 "Neural Crest 1"="#8C6BB1",
                 "Placode"="#41AB5D",
                 "vNeural Tube"="#08519C",
                 "NNE"="#A1D99B",
                 "Neural Crest 2"="#8C96C6",
                 "Hematogenic"="#BD0026",
                 "Notochord"="#DEEBF7")

p_umap <- DimPlot(chick, group.by = "celltype1",cols = celltype_colors, label = TRUE, repel = TRUE) + ggtitle(NULL)+
            NoLegend()
p_umap
```

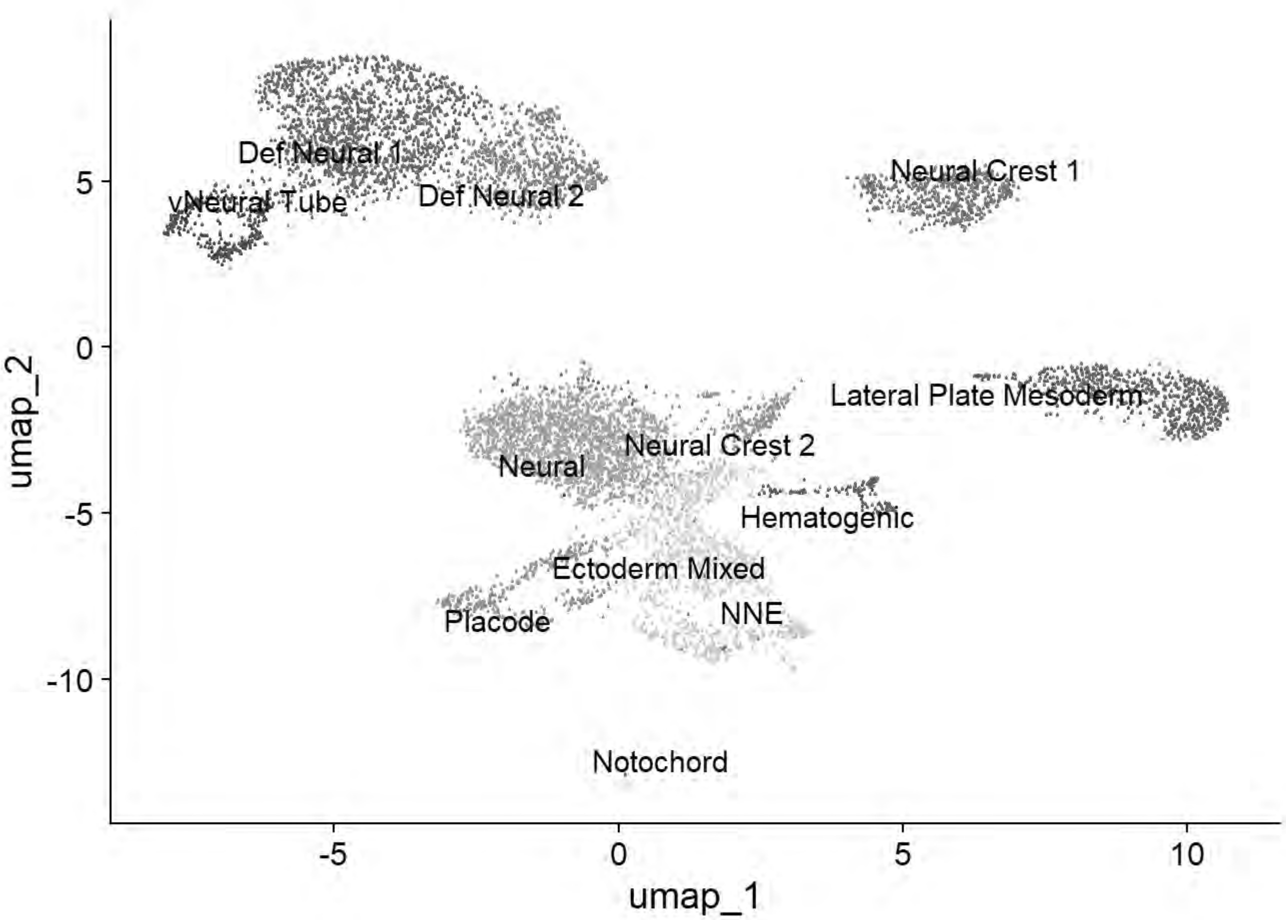


```
ggsave("E:/Rogers Lab/Rogers NC EMT/Figures/Fig_HH11_Summary_UMAP1.png", plot = p_umap, width = 6, height = 6, dpi = 600)

cptbl <- table(chick@meta.data$celltype1,chick@meta.data$stage)
```

```
cptbl <- cbind(cptbl, TOTAL = rowSums(cptbl))
colnames(cptbl)[length(cptbl[1,])] <- "TOTAL"

tg = gridExtra::tableGrob(cptbl)
h = grid::convertHeight(sum(tg$heights), "in", TRUE)
w = grid::convertWidth(sum(tg$widths), "in", TRUE)
ggsave("E:/Rogers Lab/Rogers NC EMT/Figures/Fig_HH11_Summary_Table.png", tg, width=w, height=h
, device = 'png', dpi =600)

top5 <- global_markers %>% group_by(cluster) %>% top_n(n = 5, wt = avg_log2FC)
p_dot=DotPlot(chick, features = unique(top5$gene), group.by = "celltype1") + # cluster.idents
dendrograms the y-axis
  coord_flip() +
  scale_size_continuous(range = c(1, 10)) +
  scale_color_gradientn(colors = c("grey90", "#313695", "#d73027"))+
  theme_light() +
  theme(
    axis.text.x = element_text(angle = 45, hjust = 1, color = "black", size = 12),
    axis.text.y = element_text(face = "italic", color = "black", size = 12),
    axis.title.x = element_blank(),
    axis.title.y = element_blank(),
    panel.grid.major = element_line(color = "grey90")
  )
```

```
## Scale for size is already present.
## Adding another scale for size, which will replace the existing scale.
```

```
## Scale for colour is already present.
## Adding another scale for colour, which will replace the existing scale.
```

```
p_dot
```

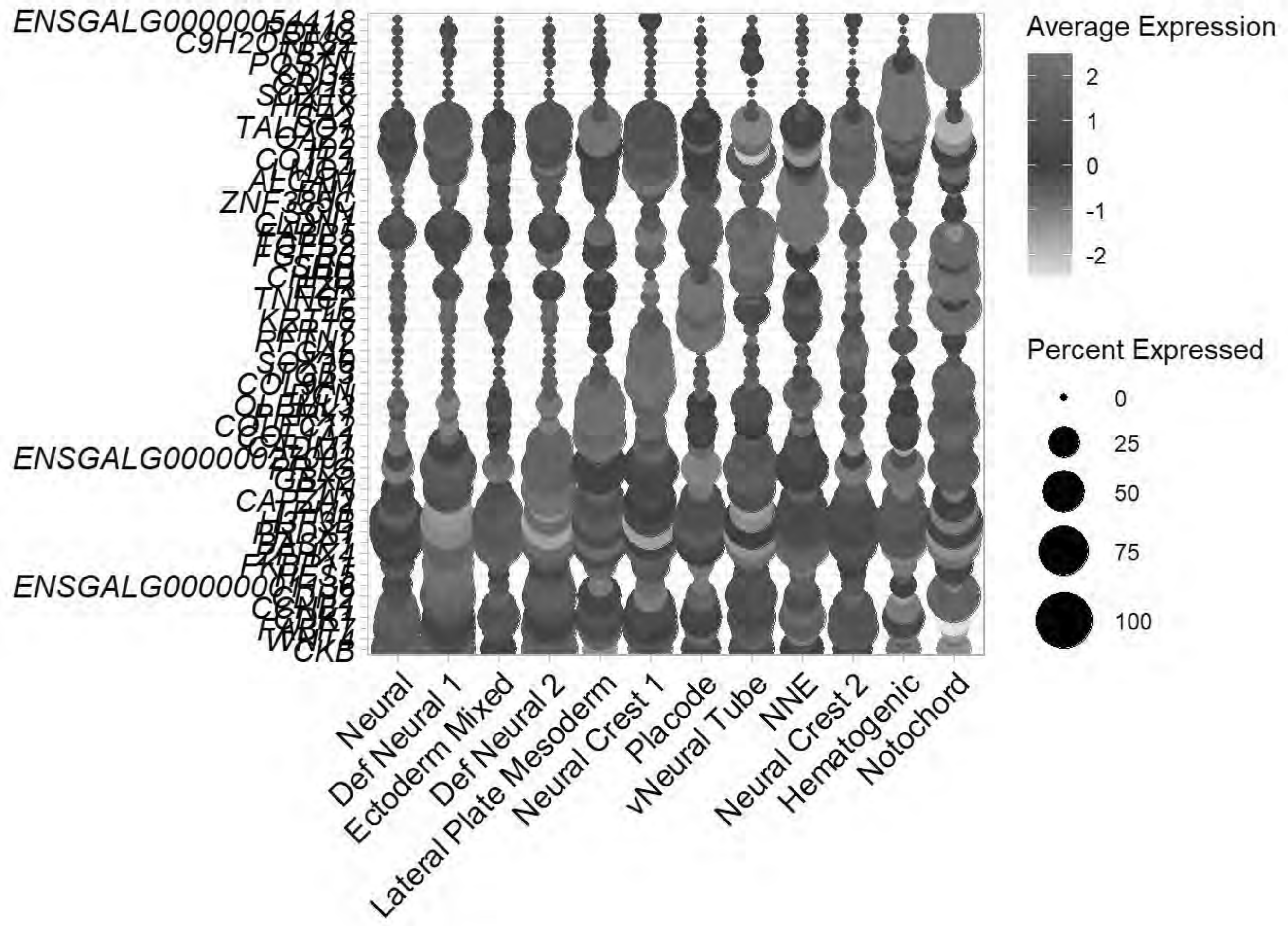


```
ggsave("E:/Rogers Lab/Rogers NC EMT/Figures/Fig_HH11_Summary_Dotplot.png", plot = p_dot, width
 = 7, height = 10, dpi = 600)
```

```
saveRDS(chick, "E:\\Rogers Lab\\Rogers NC EMT\\HH11_chick_v2.rds")

Idents(chick)<-"celltype1"
chick=subset(chick, idents =c("Neural","Def Neural 1","Ectoderm Mixed","Def Neural 2","Neural
Crest 1","Placode","vNeural Tube","NNE","Neural Crest 2","Hematogenic","Notochord"))
chick
```

```
## An object of class Seurat
## 18523 features across 6799 samples within 1 assay
## Active assay: RNA (18523 features, 2000 variable features)
##  3 layers present: data, counts, scale.data
##  3 dimensional reductions calculated: pca, harmony, umap
```

```
saveRDS(chick,"E:\\Rogers Lab\\Rogers NC EMT\\HH11_chick_NC_v2.rds")
gc()
```

```
##             used   (Mb) gc trigger    (Mb)   max used    (Mb)
## Ncells   4915453  262.6   13651909   729.1   21331105  1139.3
## Vcells 880271653 6716.0 2342481578 17871.8 4574774951 34902.8
```

### 4. Neural Crest EMT Final Integration (HH8, HH9, HH11)

```
rm(chick)
gc()
```

```
##             used   (Mb) gc trigger    (Mb)   max used    (Mb)
## Ncells   4915050  262.5   13651909   729.1   21331105  1139.3
## Vcells 832162507 6348.9 2342481578 17871.8 4574774951 34902.8
```

```
hh8=readRDS("E:\\Rogers Lab\\Rogers NC EMT\\HH8_chick_NC_v2.rds")
hh9=readRDS("E:\\Rogers Lab\\Rogers NC EMT\\HH9_chick_NC_v2.rds")
hh11=readRDS("E:\\Rogers Lab\\Rogers NC EMT\\HH11_chick_NC_v2.rds")

chick <- merge(hh8, y = c(hh9,hh11))
chick
```

```
## An object of class Seurat
## 18523 features across 31437 samples within 1 assay
## Active assay: RNA (18523 features, 2000 variable features)
##  9 layers present: data.1, data.2, data.3, counts.1, scale.data.1, counts.2, scale.data.2,
counts.3, scale.data.3
```

```
chick <- JoinLayers(chick)
chick[["RNA"]] <- split(chick[["RNA"]], f = chick$sample_id)
```

```
## Splitting 'counts', 'data' layers. Not splitting 'scale.data'. If you would like to split o
ther layers, set in `layers` argument.
```

```
chick
```

```
## An object of class Seurat
## 18523 features across 31437 samples within 1 assay
## Active assay: RNA (18523 features, 2000 variable features)
##  11 layers present: counts.Pajanoja2023_HH8_1, counts.Pajanoja2023_HH8_2, counts.Pajanoja20
23_HH9_1, counts.Pajanoja2023_HH9_2, counts.Pajanoja2025_HH11, scale.data, data.Pajanoja2023_H
H8_1, data.Pajanoja2023_HH8_2, data.Pajanoja2023_HH9_1, data.Pajanoja2023_HH9_2, data.Pajanoja
2025_HH11
```

```
rm(hh8,hh9,hh11)
```

```
chick <- NormalizeData(chick)
```

```
## Normalizing layer: counts.Pajanoja2023_HH8_1
```

```
## Normalizing layer: counts.Pajanoja2023_HH8_2
```

```
## Normalizing layer: counts.Pajanoja2023_HH9_1
```

```
## Normalizing layer: counts.Pajanoja2023_HH9_2
```

```
## Normalizing layer: counts.Pajanoja2025_HH11
```

```
chick <- CellCycleScoring(chick, s.features = cc.genes$s.genes, g2m.features = cc.genes$g2m.ge
nes)
```

```
## Warning: The following features are not present in the object: TYMS, CDCA7,
## PRIM1, MLF1IP, RAD51AP1, not searching for symbol synonyms
```

```
## Warning: The following features are not present in the object: MKI67, FAM64A,
## AURKB, KIF20B, CDC25C, CDCA2, CDCA8, PSRC1, CENPA, not searching for symbol
## synonyms
```

```
## Warning: The following features are not present in the object: TYMS, CDCA7,
## PRIM1, MLF1IP, RAD51AP1, not searching for symbol synonyms
```

```
## Warning: The following features are not present in the object: MKI67, FAM64A,
## AURKB, KIF20B, CDC25C, CDCA2, CDCA8, PSRC1, CENPA, not searching for symbol
## synonyms
```

```
## Warning: The following features are not present in the object: TYMS, CDCA7,
## PRIM1, MLF1IP, RAD51AP1, not searching for symbol synonyms
```

```
## Warning: The following features are not present in the object: MKI67, FAM64A,
## AURKB, KIF20B, CDC25C, CDCA2, CDCA8, PSRC1, CENPA, not searching for symbol
## synonyms
```

```
## Warning: The following features are not present in the object: TYMS, CDCA7,
## PRIM1, MLF1IP, RAD51AP1, not searching for symbol synonyms
```

```
## Warning: The following features are not present in the object: MKI67, FAM64A,
## AURKB, KIF20B, CDC25C, CDCA2, CDCA8, PSRC1, CENPA, not searching for symbol
## synonyms
```

```
## Warning: The following features are not present in the object: TYMS, CDCA7,
## PRIM1, MLF1IP, RAD51AP1, not searching for symbol synonyms
```

```
## Warning: The following features are not present in the object: MKI67, FAM64A,
## AURKB, KIF20B, CDC25C, CDCA2, CDCA8, PSRC1, CENPA, not searching for symbol
```

```
## synonyms
```

```
chick <- FindVariableFeatures(chick)
```

```
## Finding variable features for layer counts.Pajanoja2023_HH8_1
```

```
## Finding variable features for layer counts.Pajanoja2023_HH8_2
```

```
## Finding variable features for layer counts.Pajanoja2023_HH9_1
```

```
## Finding variable features for layer counts.Pajanoja2023_HH9_2
```

```
## Finding variable features for layer counts.Pajanoja2025_HH11
```

```
chick <- ScaleData(chick, vars.to.regress = c("percent.mt", "S.Score", "G2M.Score"))
```

```
## Regressing out percent.mt, S.Score, G2M.Score
```

```
## Centering and scaling data matrix
```

```
## Warning: Different features in new layer data than already exists for
## scale.data
```

```
chick <- RunPCA(chick, npcs = 50, verbose = FALSE)

chick <- IntegrateLayers(object = chick, method = HarmonyIntegration, orig.reduction = "pca",
new.reduction = "harmony", assay = "RNA", group.by = "sample_id", verbose = TRUE)
```

```
## Transposing data matrix
```

```
## Using automatic lambda estimation
```

```
## Initializing state using k-means centroids initialization
```

```
## Harmony 1/10
```

```
## Harmony 2/10
```

```
## Harmony 3/10
```

```
## Harmony 4/10
```

```
## Harmony converged after 4 iterations
```

```
chick <- FindNeighbors(chick, reduction = "harmony", dims = 1:30)
```

```
## Computing nearest neighbor graph
```

```
## Computing SNN
```

```
chick <- RunUMAP(chick, reduction = "harmony", dims = 1:30)
```

```
## 23:28:55 UMAP embedding parameters a = 0.9922 b = 1.112
```

```
## 23:28:55 Read 31437 rows and found 30 numeric columns
```

```
## 23:28:55 Using Annoy for neighbor search, n_neighbors = 30
```

```
## 23:28:55 Building Annoy index with metric = cosine, n_trees = 50
```

```
## 0%   10   20   30   40   50   60   70   80   90   100%
```

```
## [----|----|----|----|----|----|----|----|----|----|
```

```
## **************************************************|
## 23:28:58 Writing NN index file to temp file C:\Users\RANEES~1\AppData\Local\Temp\RtmpqcWgVb
\file6a2c54a2190c
## 23:28:58 Searching Annoy index using 1 thread, search_k = 3000
## 23:29:06 Annoy recall = 100%
## 23:29:07 Commencing smooth kNN distance calibration using 1 thread with target n_neighbors
= 30
## 23:29:08 Initializing from normalized Laplacian + noise (using RSpectra)
## 23:29:09 Commencing optimization for 200 epochs, with 1482326 positive edges
## 23:29:09 Using rng type: pcg
## 23:29:37 Optimization finished
```

```
DimPlot(chick)
```

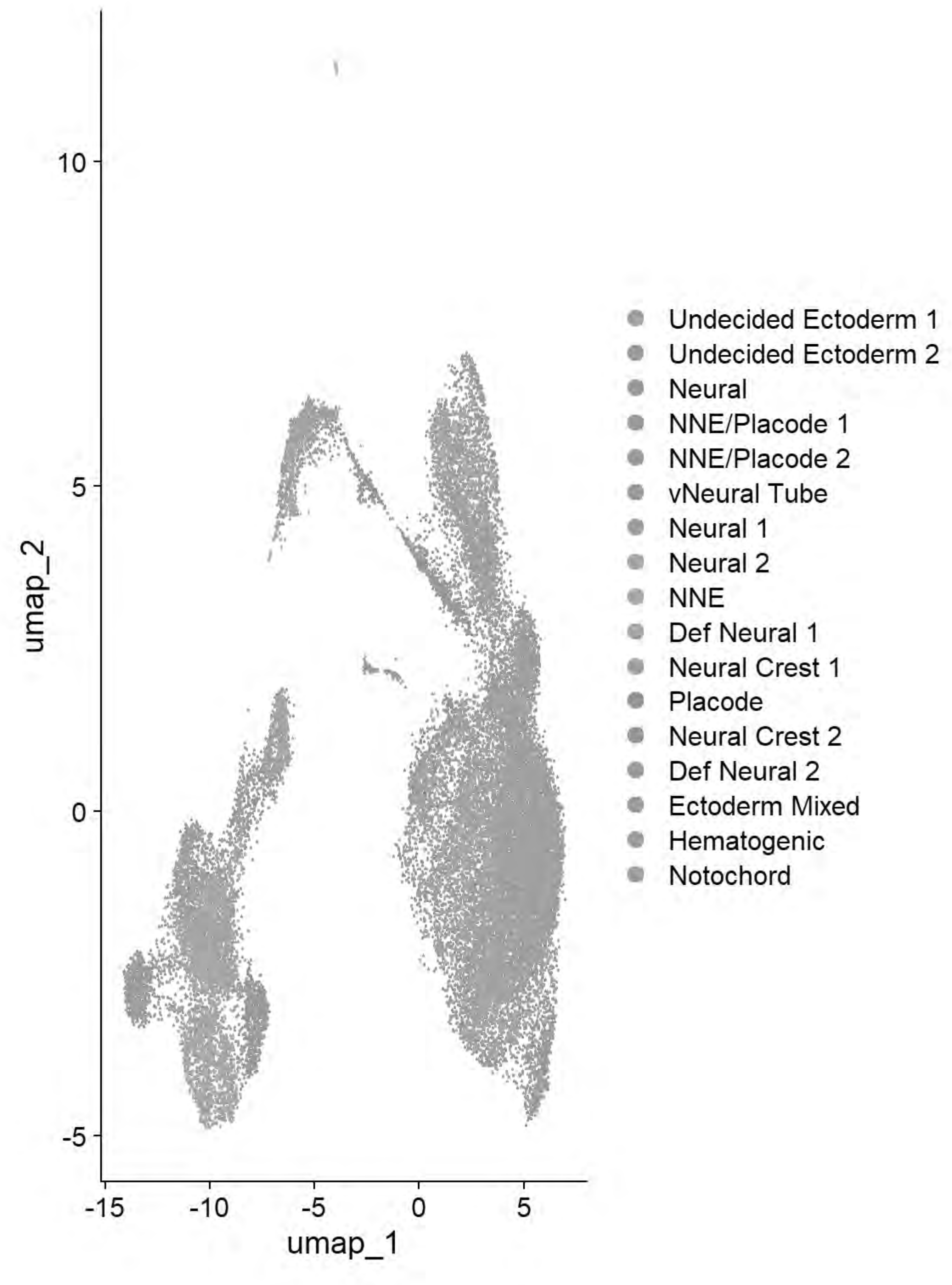


```
chick <- FindClusters(chick, resolution = 0.4)
```

```
## Modularity Optimizer version 1.3.0 by Ludo Waltman and Nees Jan van Eck
##
## Number of nodes: 31437
## Number of edges: 1015587
##
## Running Louvain algorithm...
## Maximum modularity in 10 random starts: 0.8980
```

```
## Number of communities: 14
## Elapsed time: 5 seconds
```

```
qc_tester(chick)
```

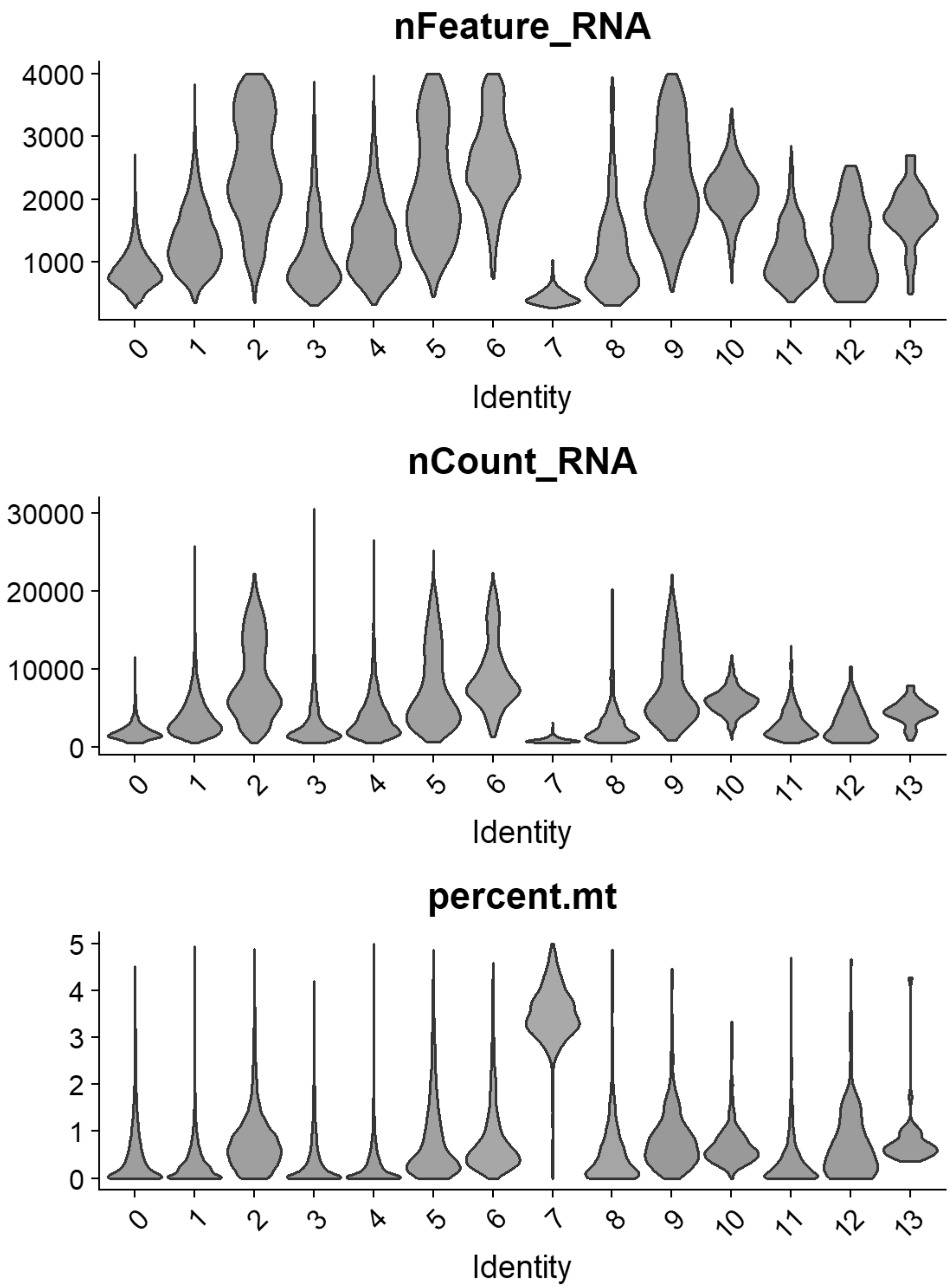


```
chick = FindClusters(chick, resolution= seq(0,2,0.1),n.start=10, verbose = FALSE)
p=clustree(chick, prefix= "RNA_snn_res.")
print(p)
```

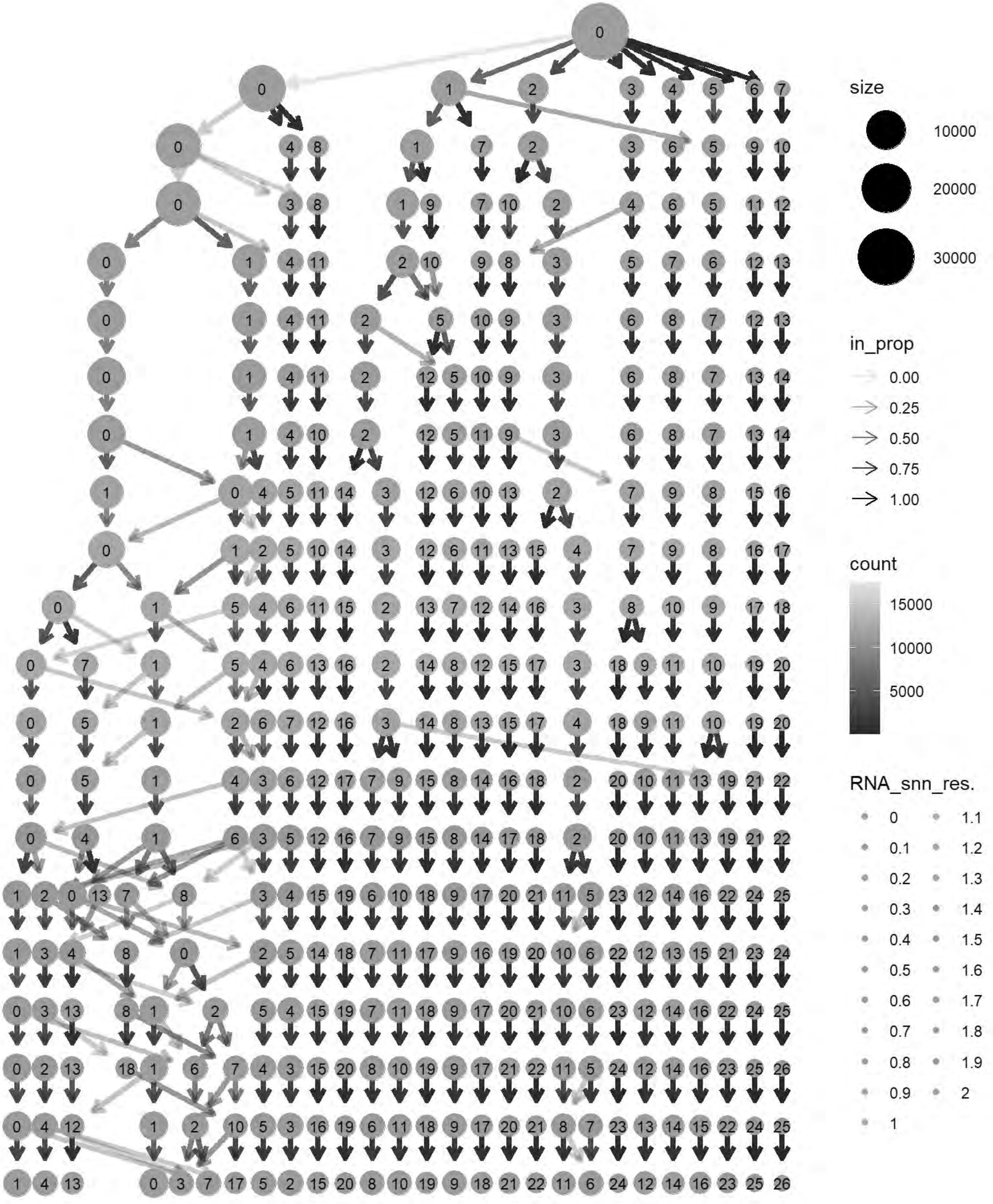
size
10000
20000
30000
in_prop
0.00
0.25
0.50
0.75
1.00
count
15000
10000
5000
RNA_snn_res.
0
0.1
0.2
0.3
0.4
0.5
0.6
0.7
0.8
0.9
1
1.1
1.2
1.3
1.4
1.5
1.6
1.7
1.8
1.9
2

```
cols= grep("^RNA_snn_res.",names(chick@meta.data))
chick@meta.data[cols] <- NULL
chick <- FindClusters(chick, resolution = 0.6, verbose = TRUE)
```

```
## Modularity Optimizer version 1.3.0 by Ludo Waltman and Nees Jan van Eck
##
## Number of nodes: 31437
## Number of edges: 1015587
##
## Running Louvain algorithm...
## Maximum modularity in 10 random starts: 0.8734
## Number of communities: 15
## Elapsed time: 4 seconds
```

```
p6=DimPlot(chick, label = TRUE)+NoLegend()
print(p6)
```

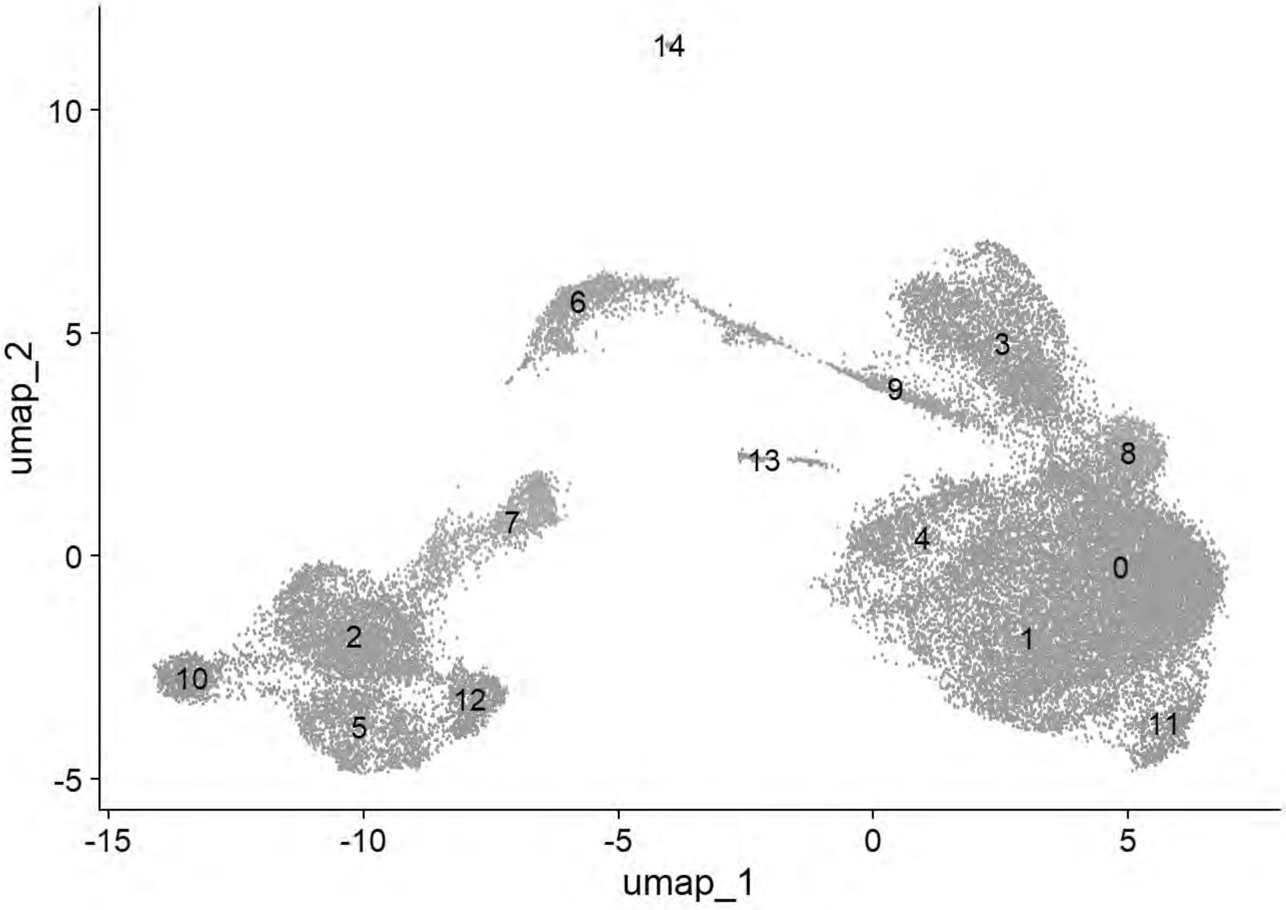


```
p6=DimPlot(chick, group.by = "Phase")
print(p6)
```

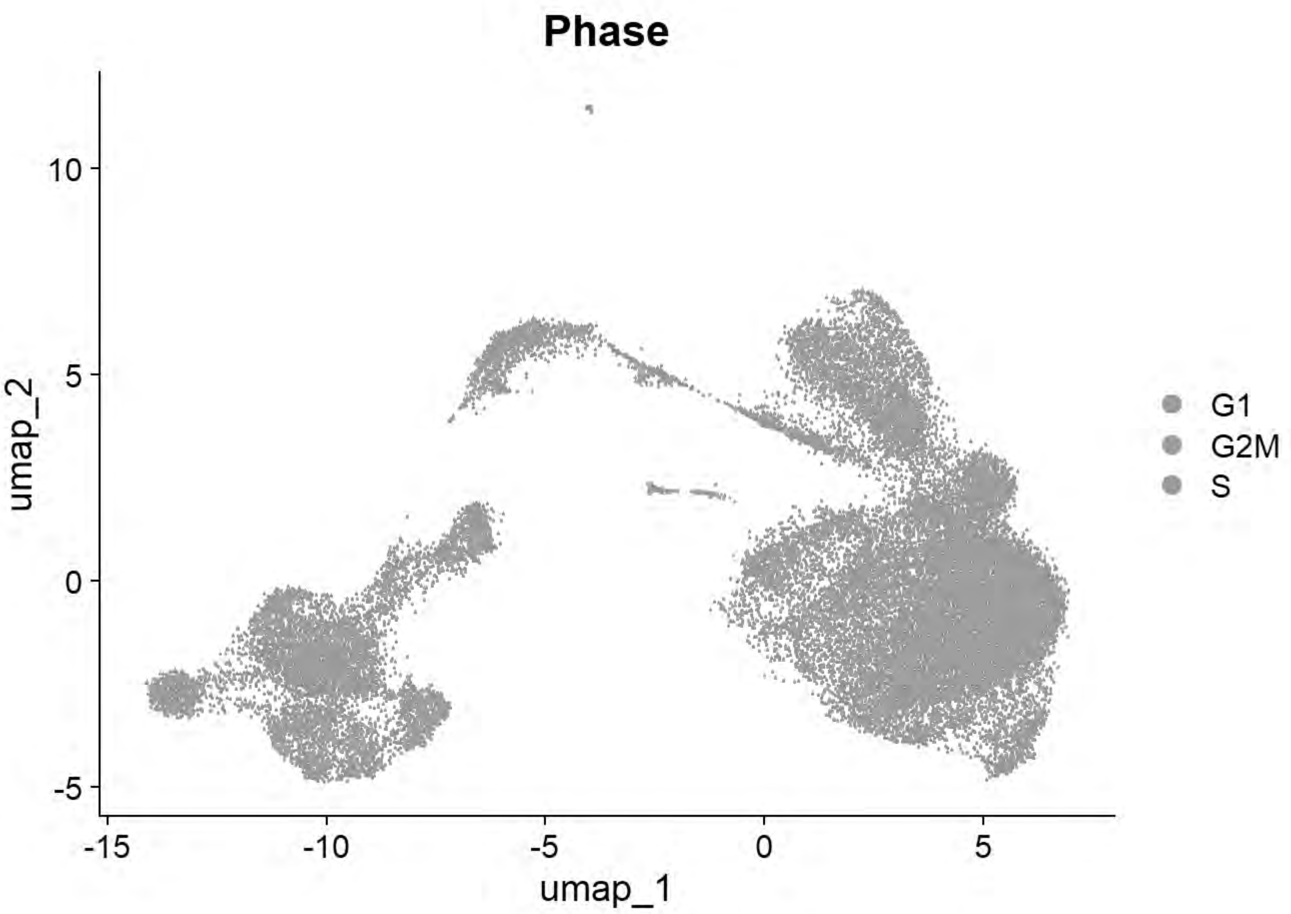


```
p6=DimPlot(chick, group.by = "sample_id")
print(p6)
```

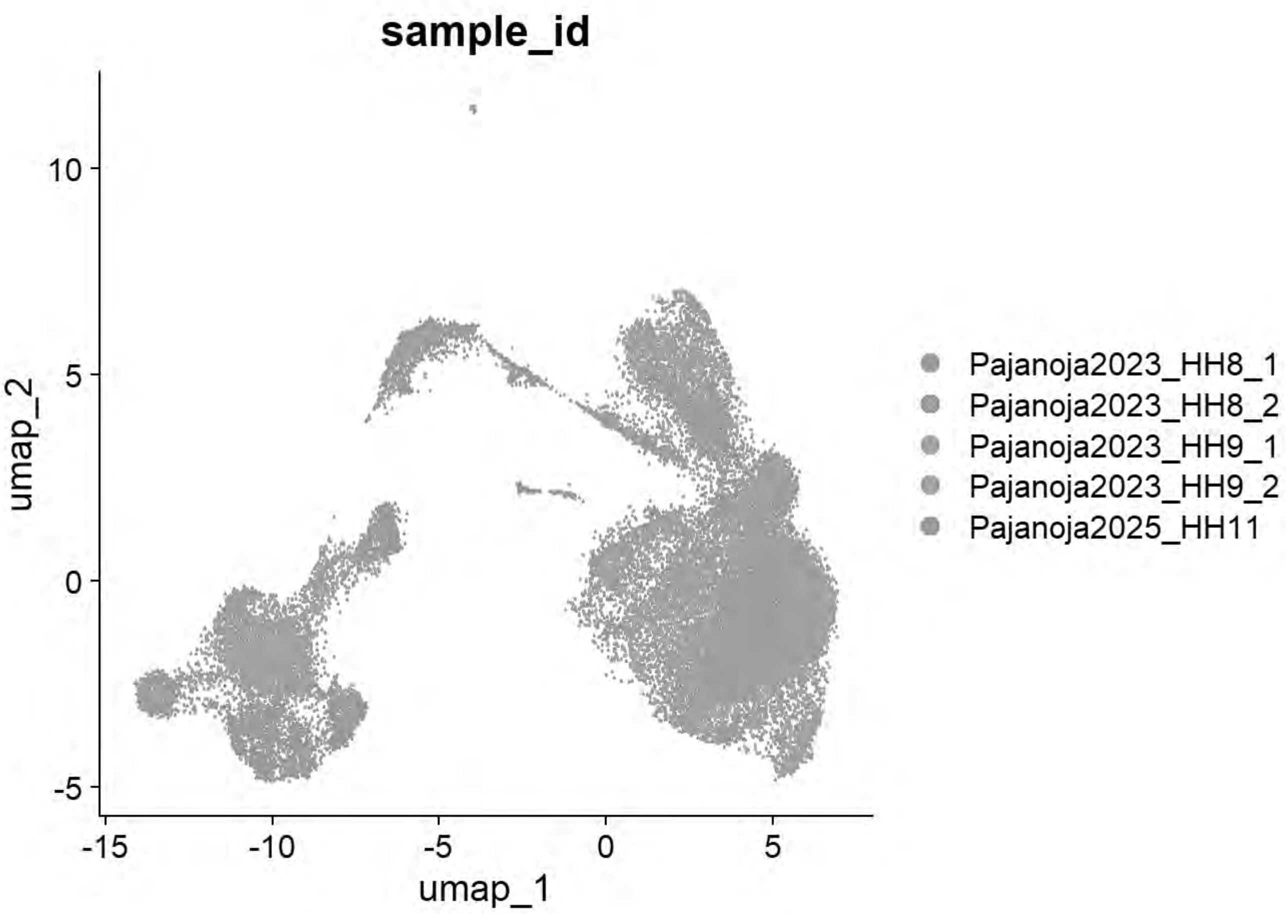


```
table(chick$seurat_clusters,chick$sample_id)
```

```
## 
##      Pajanoja2023_HH8_1 Pajanoja2023_HH8_2 Pajanoja2023_HH9_1
##   0                 969               1527               2402
##   1                 829               2210               1106
##   2                1338                774                879
##   3                 610                720                862
##   4                 169                342                519
##   5                   3                  0                  2
##   6                 580                100                 17
##   7                 184                115                140
##   8                   8                848                  1
##   9                  33                 25                233
##   10                180                 60                103
##   11                110                124                175
##   12                  0                  1                  1
##   13                  5                  5                  9
##   14                  0                  0                  0
## 
##      Pajanoja2023_HH9_2 Pajanoja2025_HH11
##   0                2715              1309
##   1                1221               634
##   2                 316               337
##   3                 671               576
```

```
##   4                  614                221
##   5                    4               1395
##   6                  118                336
##   7                  105                557
##   8                    4                  9
##   9                  246                268
##   10                  73                286
##   11                 230                 51
##   12                   0                661
##   13                   3                119
##   14                   0                 40
```

```
qc_tester(chick)
```

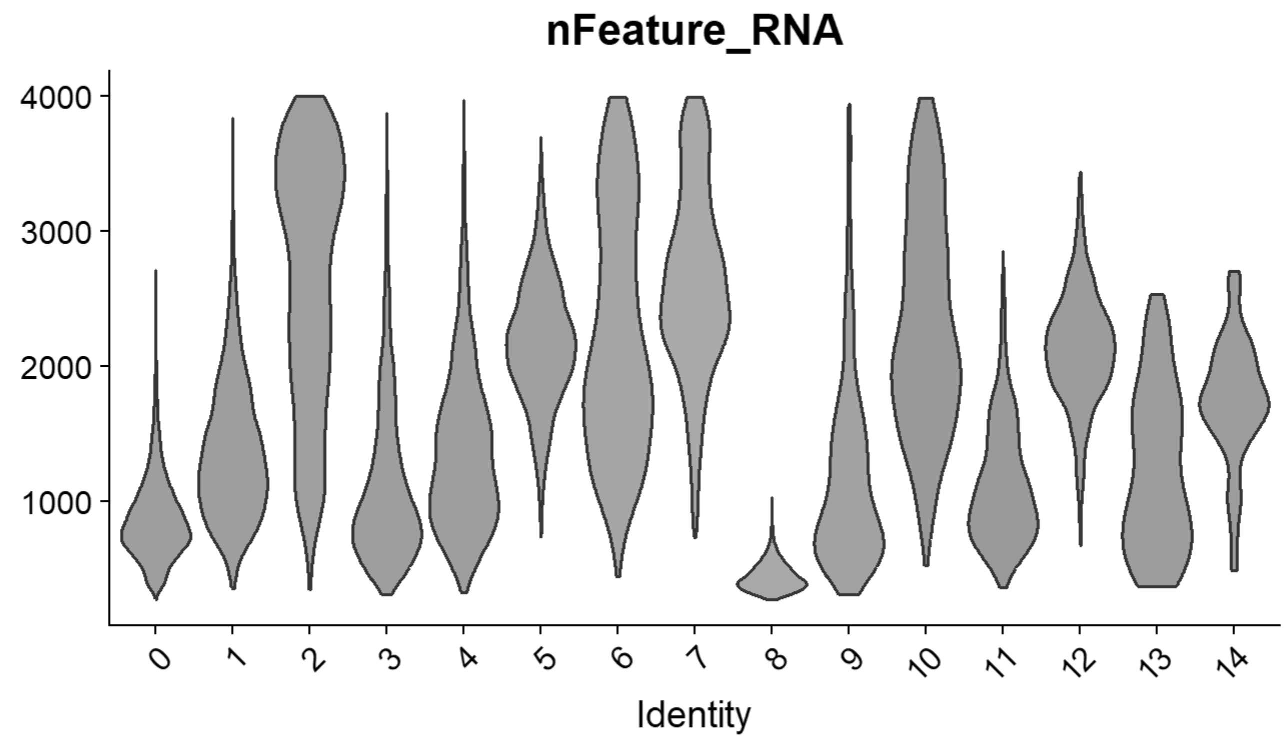


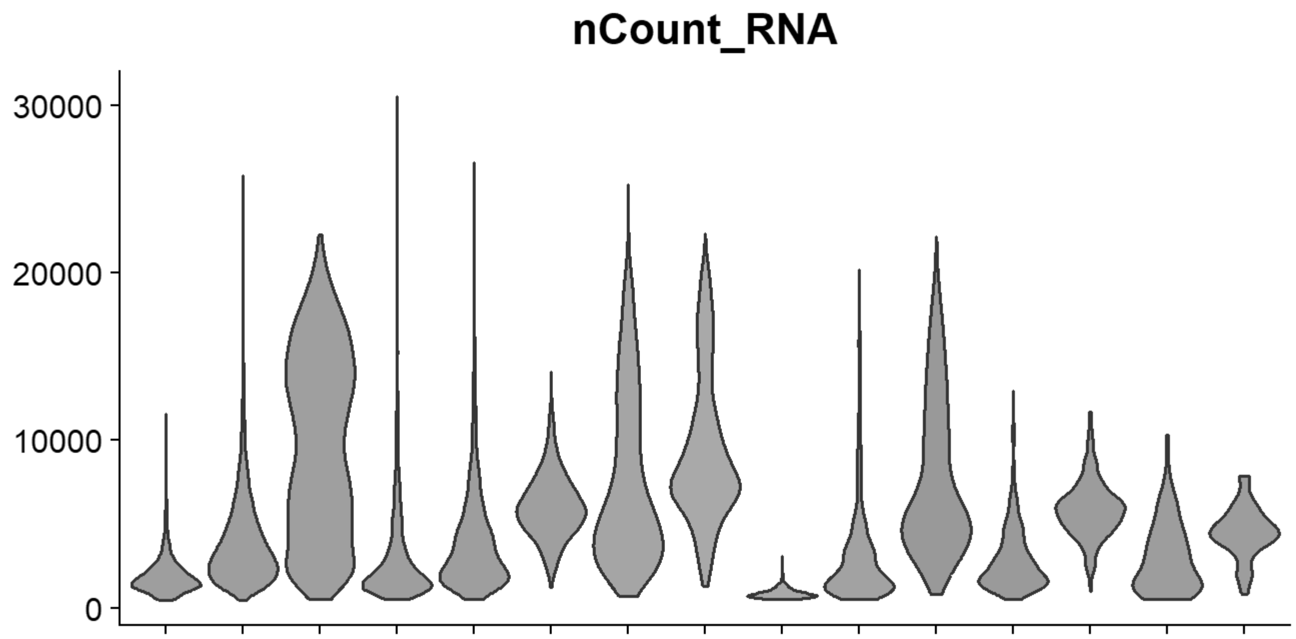

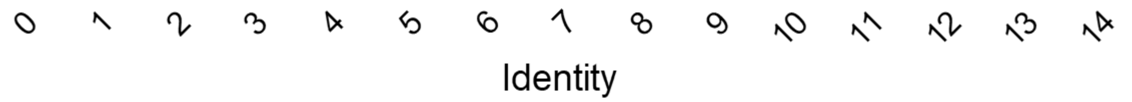


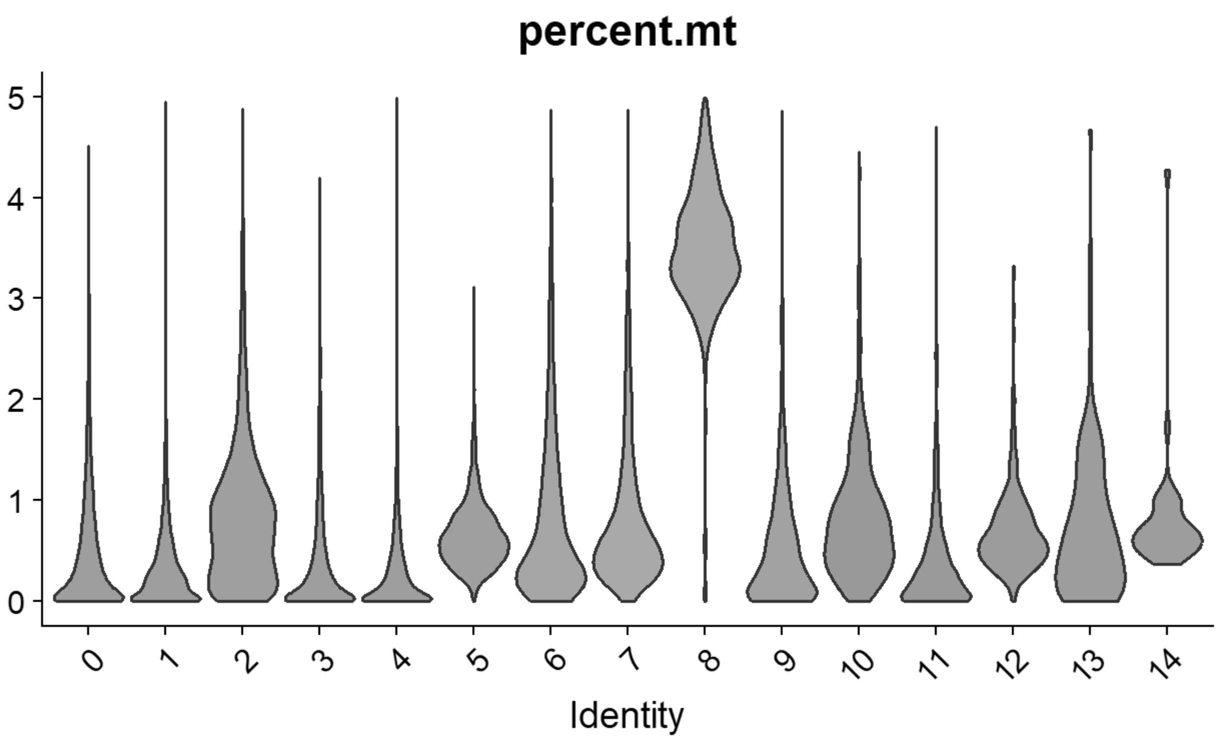


```
# Williams: Neural plate/tube
FeaturePlot(chick, features = c("CLDN1","FRZB","DNMT3A"))
```

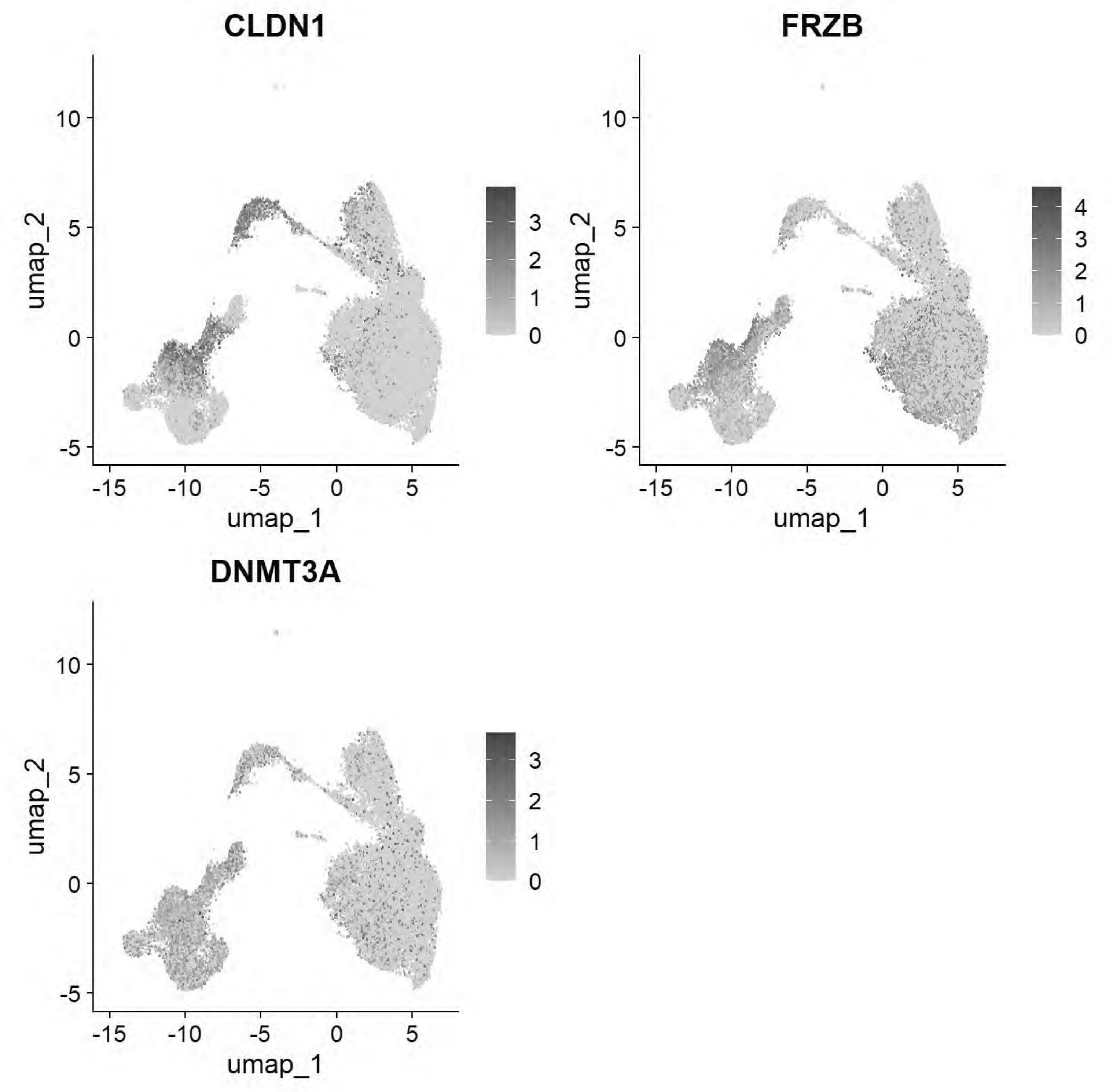


```r
# Williams: Non-Neural Ectoderm
FeaturePlot(chick, features = c("DLX5","TFAP2A","ASTL","PAX6"))
```

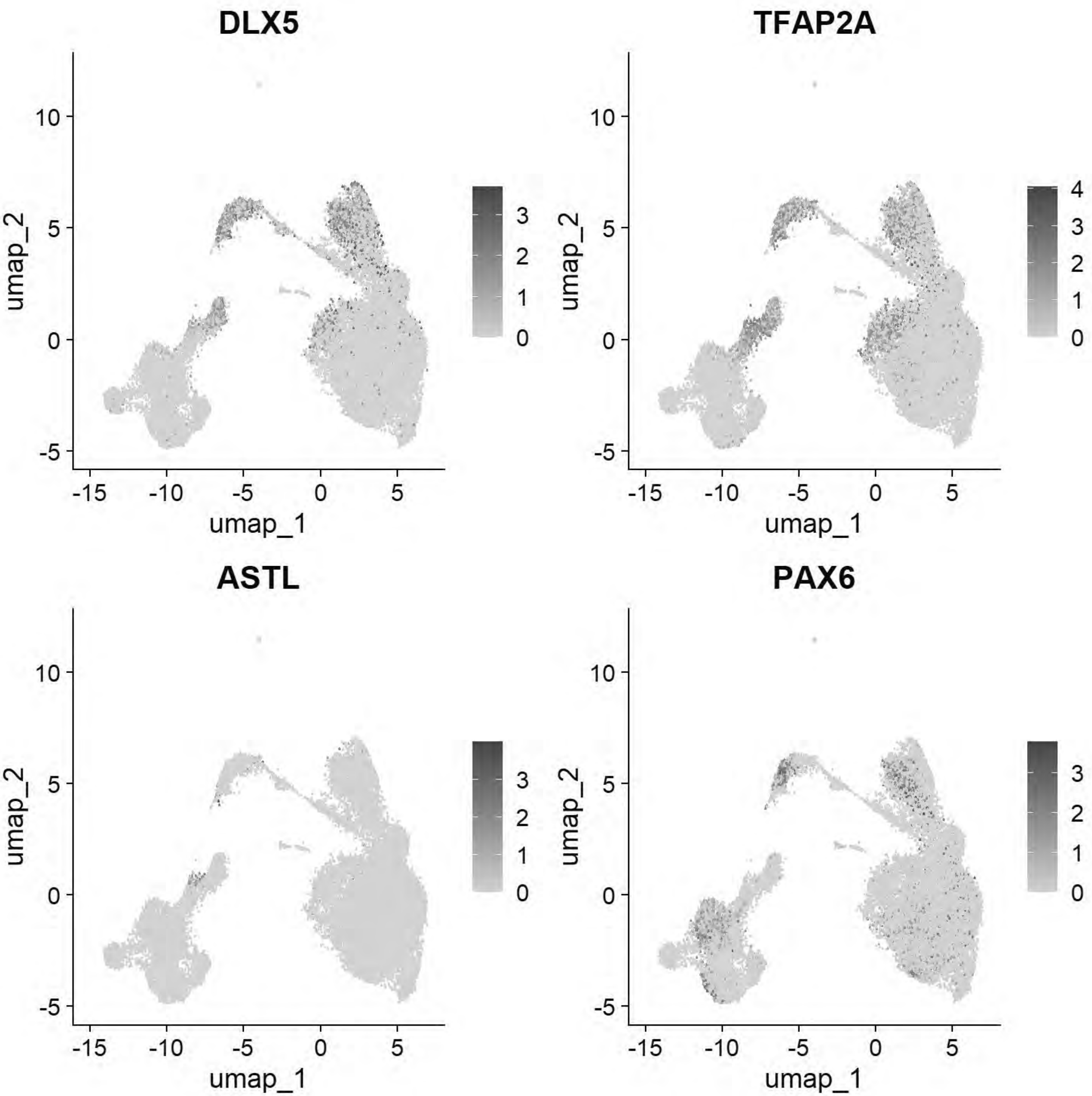


```
# Williams: Ectoderm (Neural Plate)
FeaturePlot(chick, features = c("SOX2","SOX3","SOX21","FRZB","SFRP2"))
```

```
## Warning: All cells have the same value (0) of "SOX3"
```

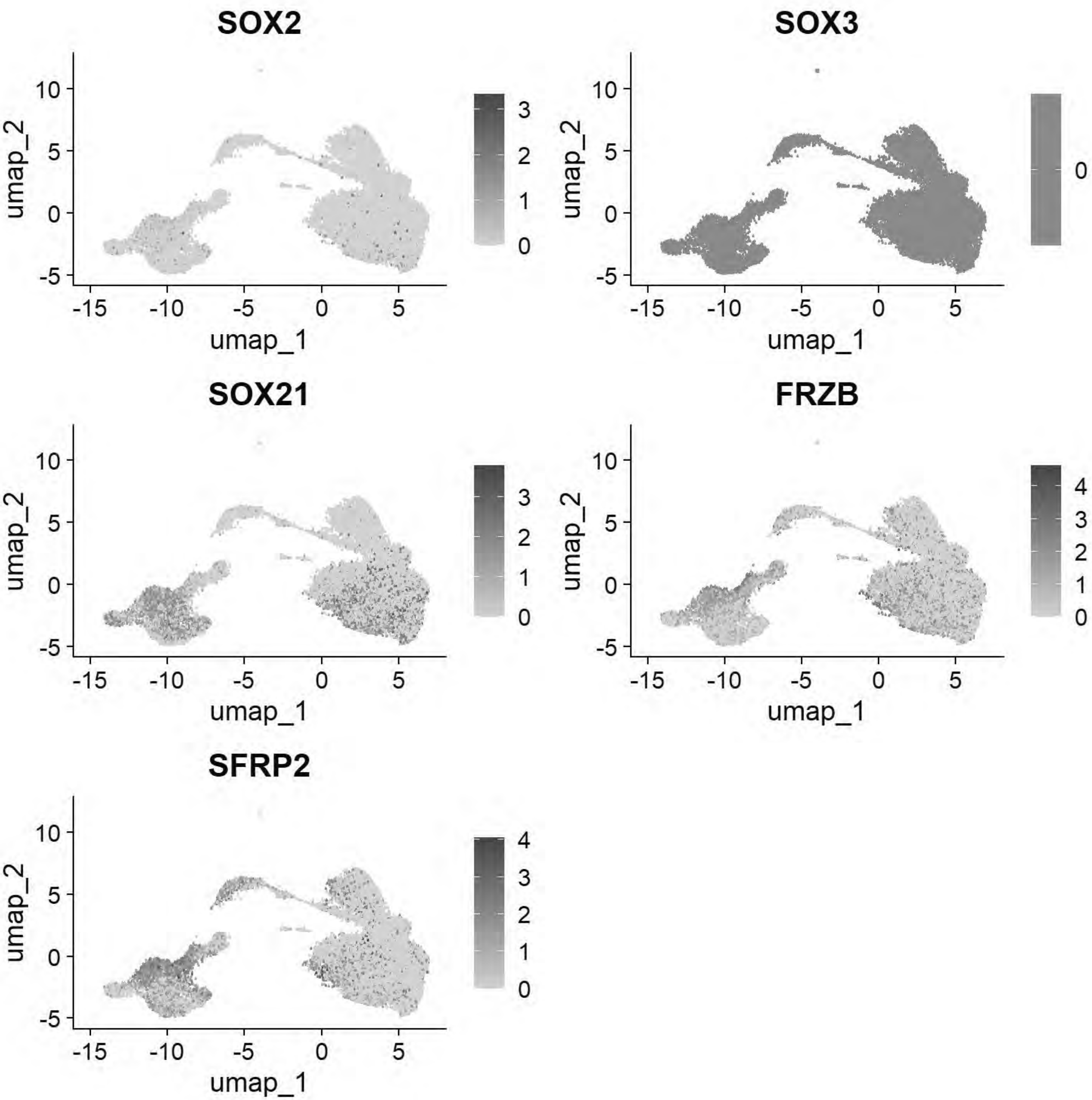


```
# Williams: Ectoderm (Neural Plate Border)
FeaturePlot(chick, features = c("PAX7","TFAP2A","DLX5","BMP4","MSX1","DRAXIN","TFAP2B"))
```

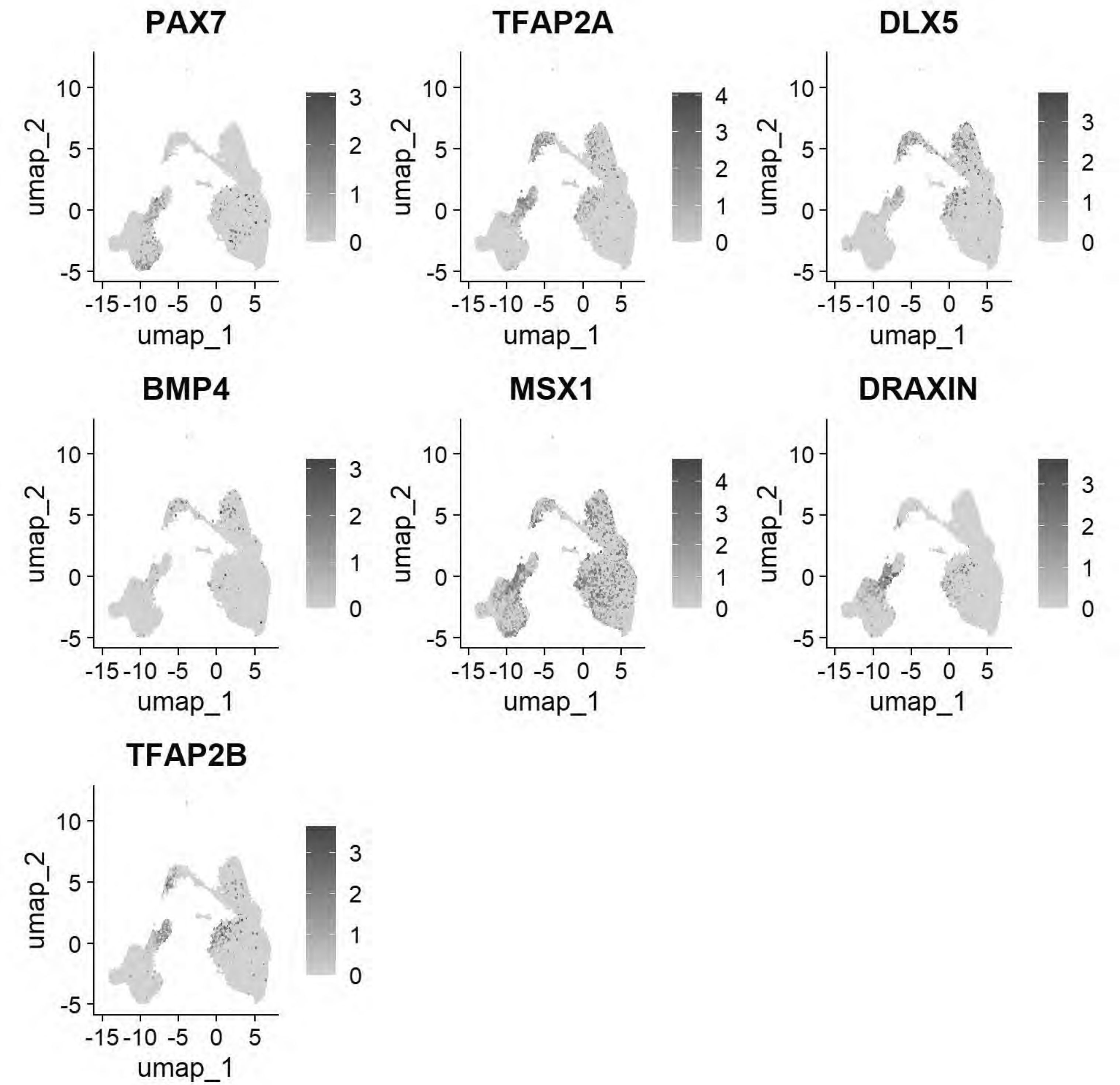


```
# Williams:Epiblast stem cell
FeaturePlot(chick, features = c("ID3","SALL4","TGIF1","ELAVL1"))
```

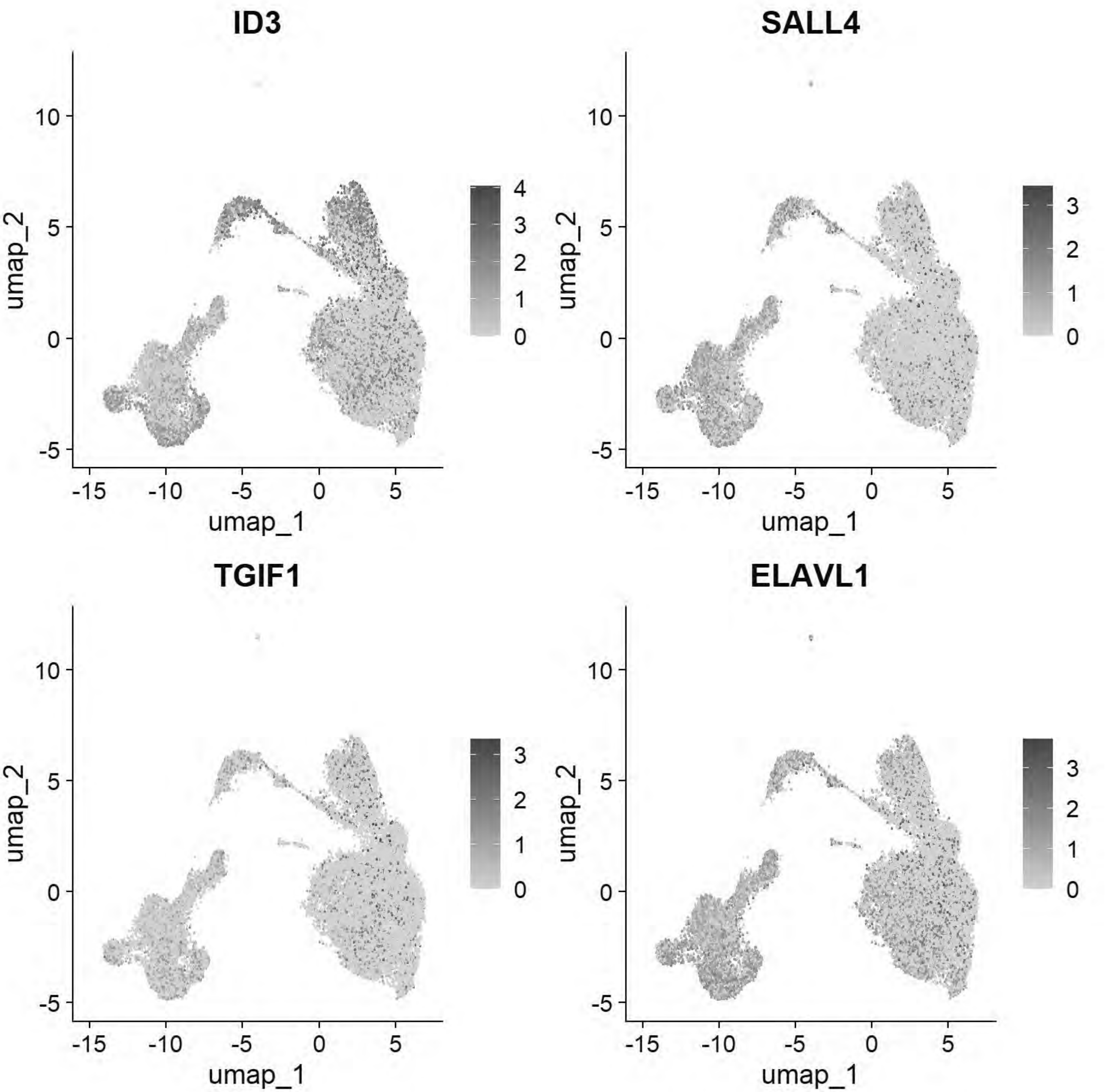


```
# Williams: Posterior Lateral Plate Mesoderm
FeaturePlot(chick, features = c("GATA2","HOXB5","CDX4"))
```

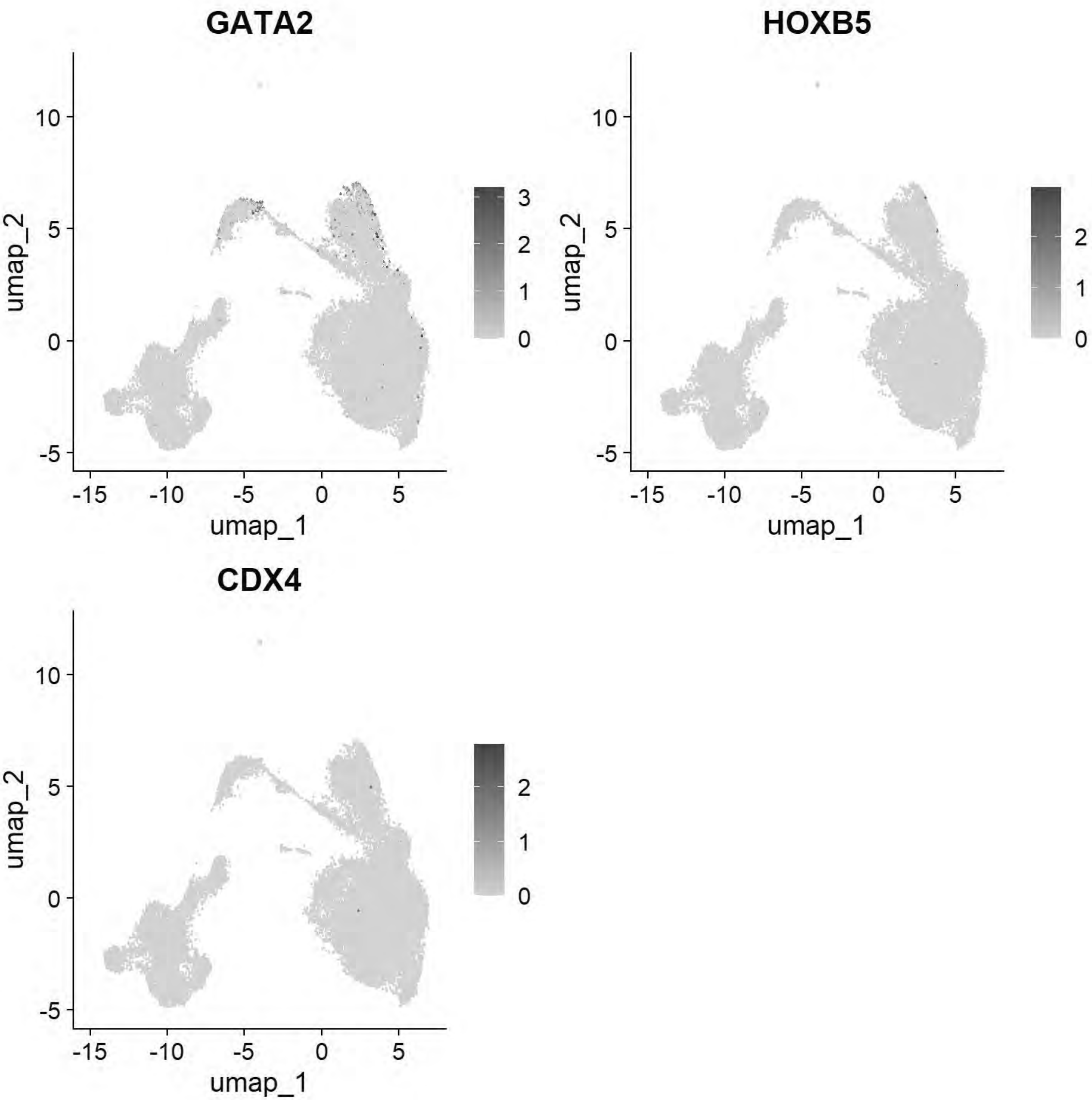


```
# Williams: Paraxial Mesoderm
FeaturePlot(chick, features = c("MSGN1","MESP1","MEOX1"))
```

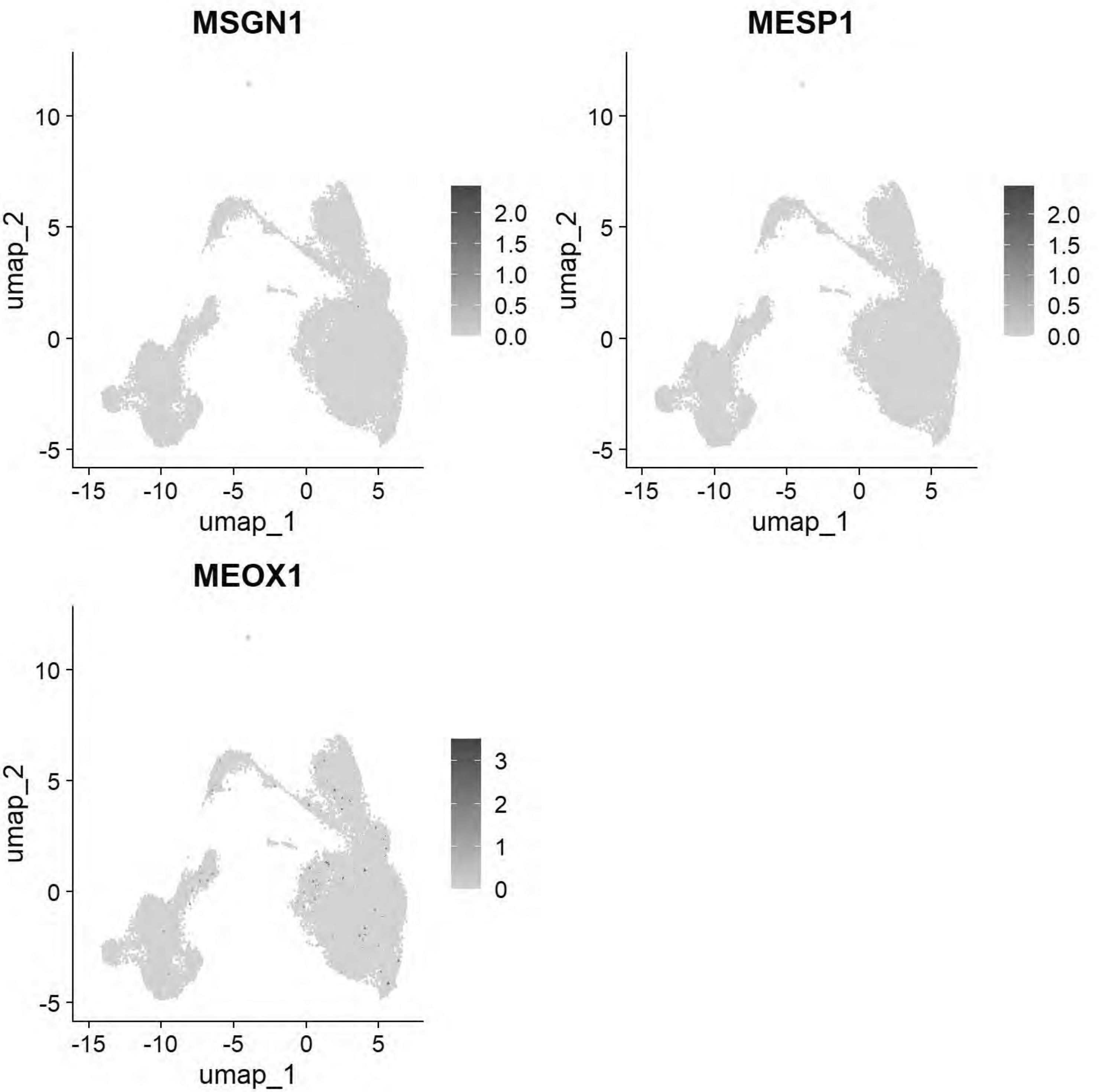


```
# Williams: Lateral Plate Mesoderm
FeaturePlot(chick, features = c("PITX2","ALX1","OLFML3","SIX1","TWIST1"))
```

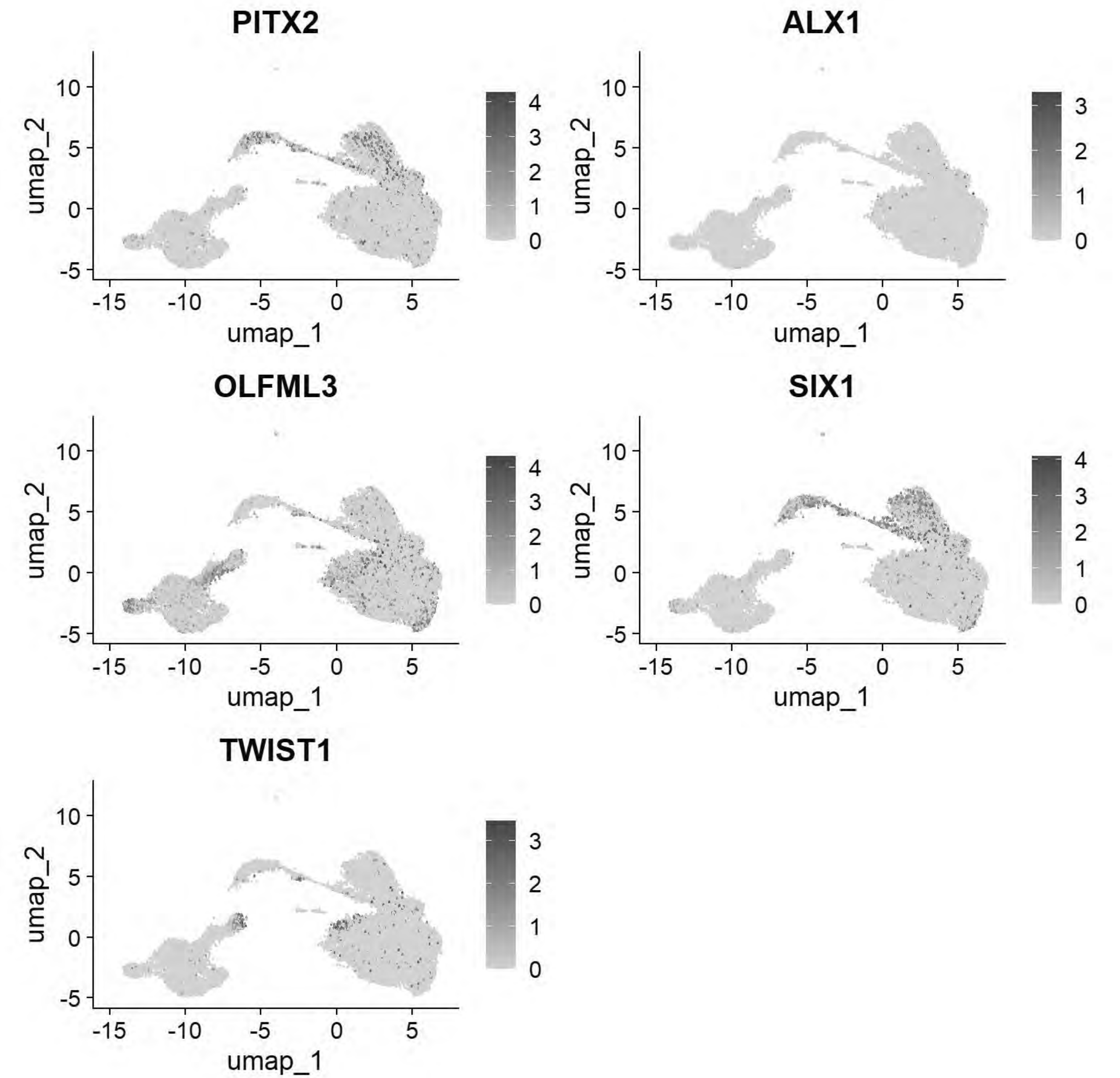


```
# Williams: Cardiac Mesoderm
FeaturePlot(chick, features = c("TCF21","GATA5","LMO2","ETS1","KDR"))
```

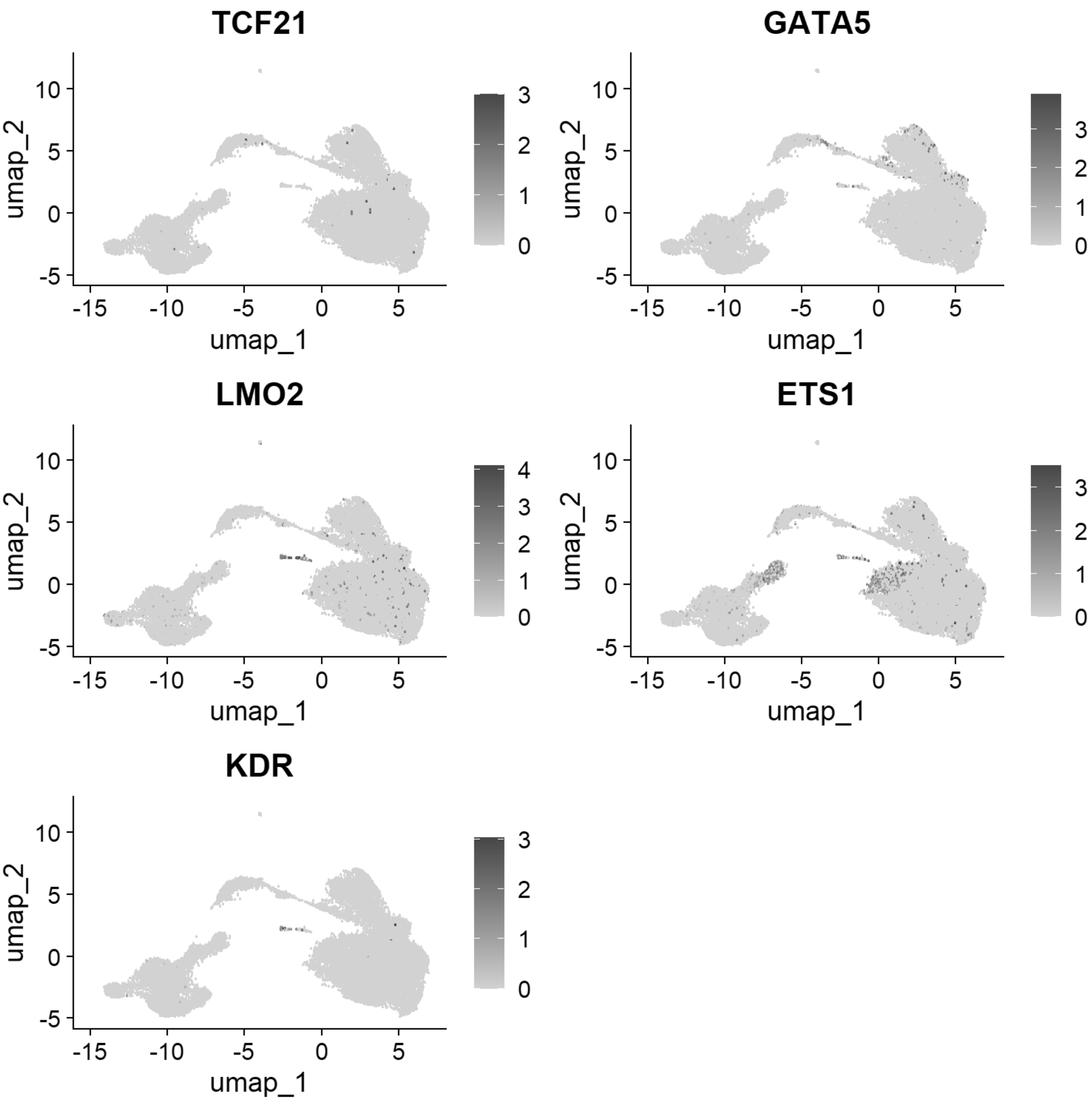


```
# Williams: Head Mesenchyme
FeaturePlot(chick, features = c("TCF21","GATA5","LMO2","ETS1","KDR"))
```

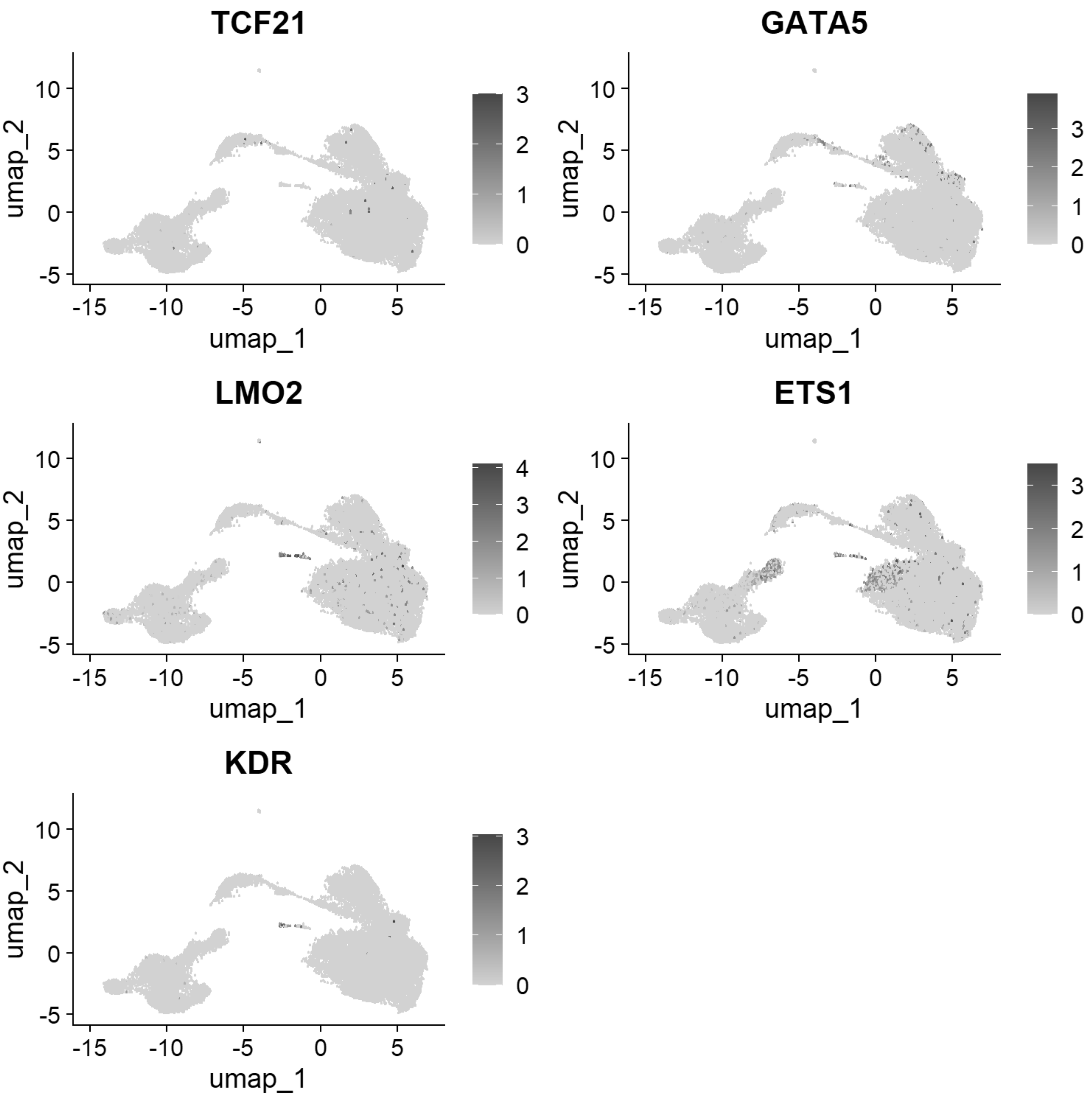


```r
# Williams: Hensen's Node & Primitive Streak
FeaturePlot(chick, features = c("DLL1","FGF8","NOTO","CHRD"))
```

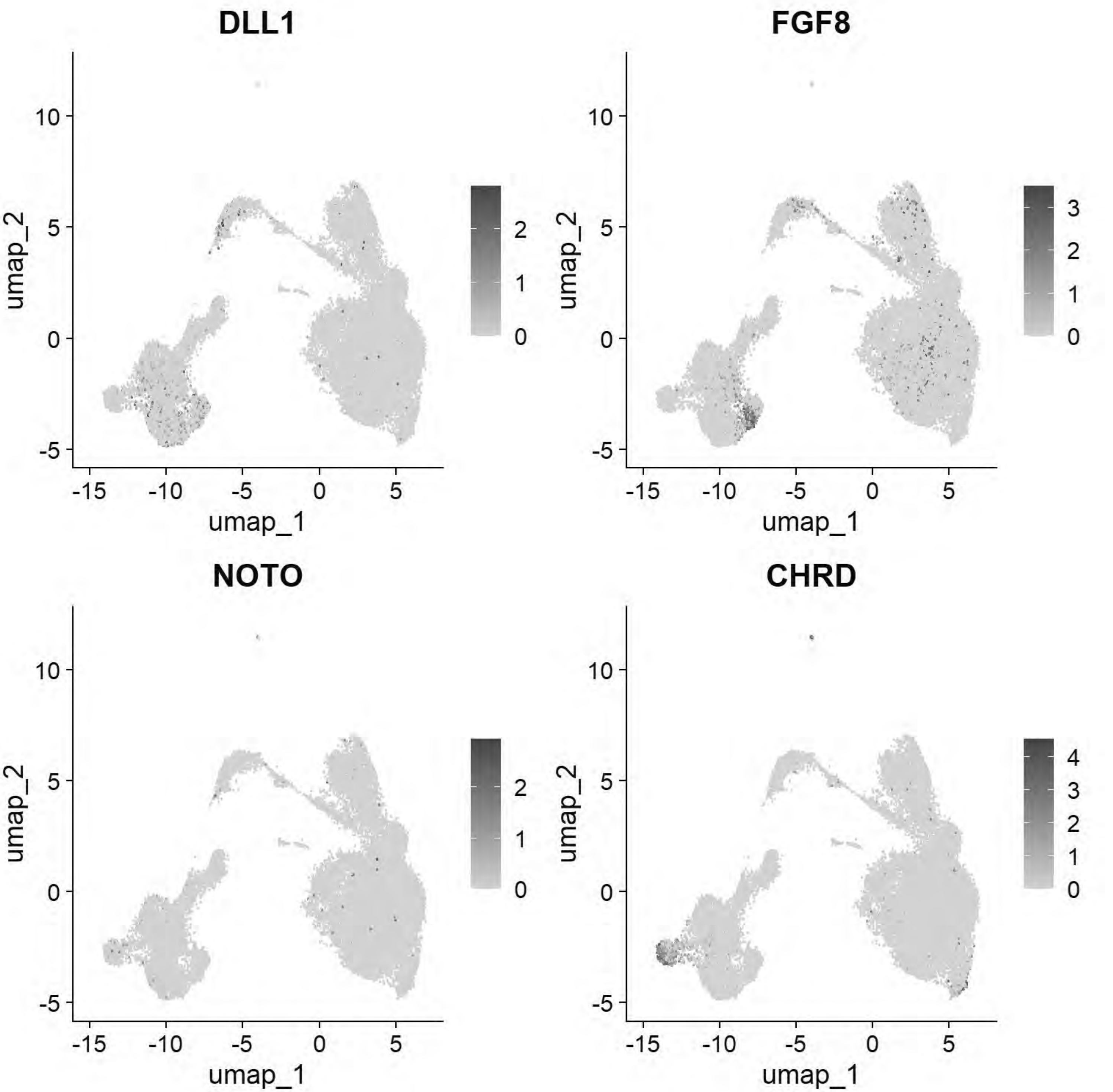


```
# Williams: Endoderm
FeaturePlot(chick, features = c("SOX17","FOXA2","CXCR4"))
```

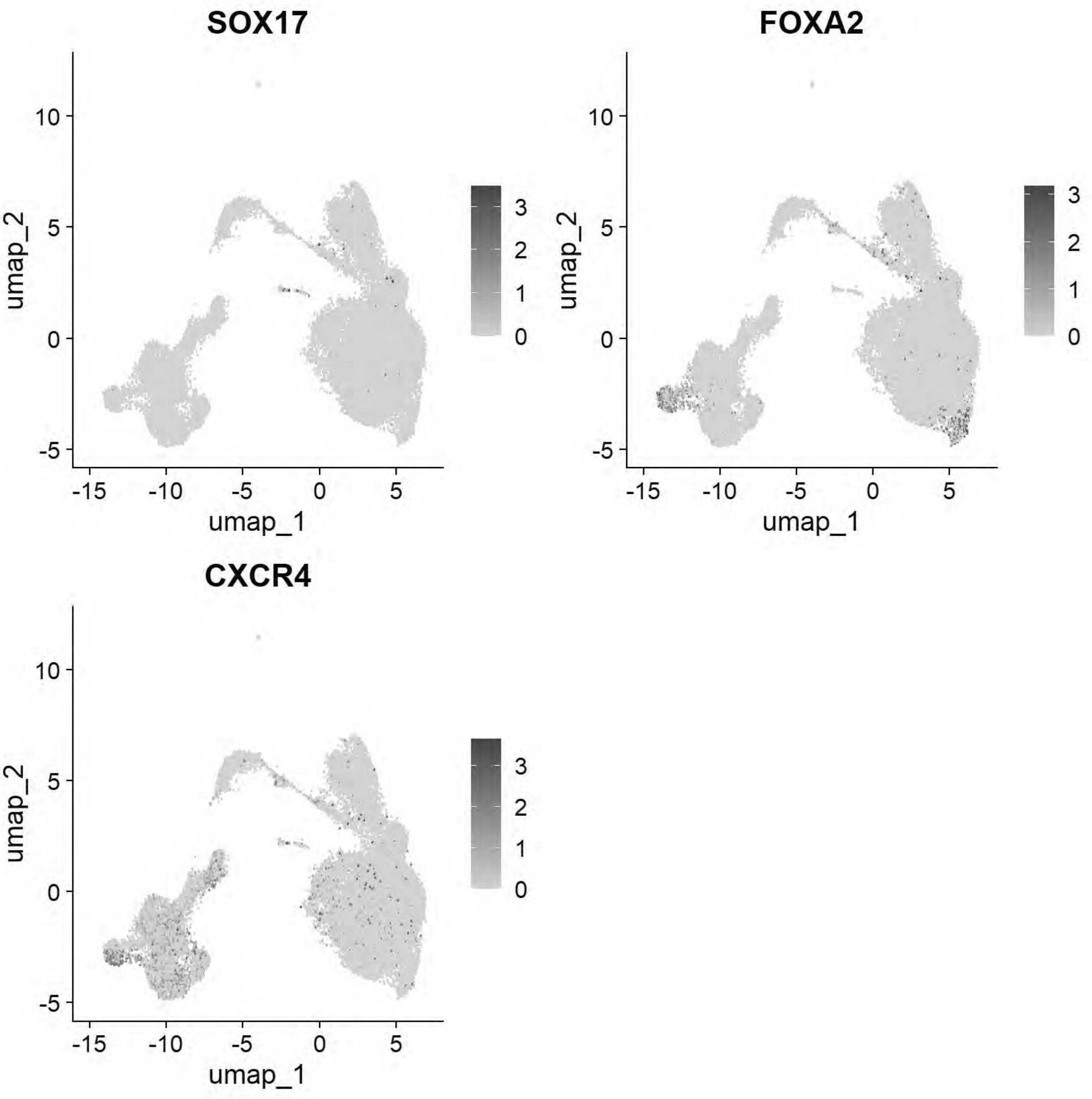


```
# Pajanoja: Ectoderm
FeaturePlot(chick, features = c("TFAP2A","DLX5","CLDN1","SOX2","NESTIN","MYCN"))
```

```
## Warning: The following requested variables were not found: NESTIN
```

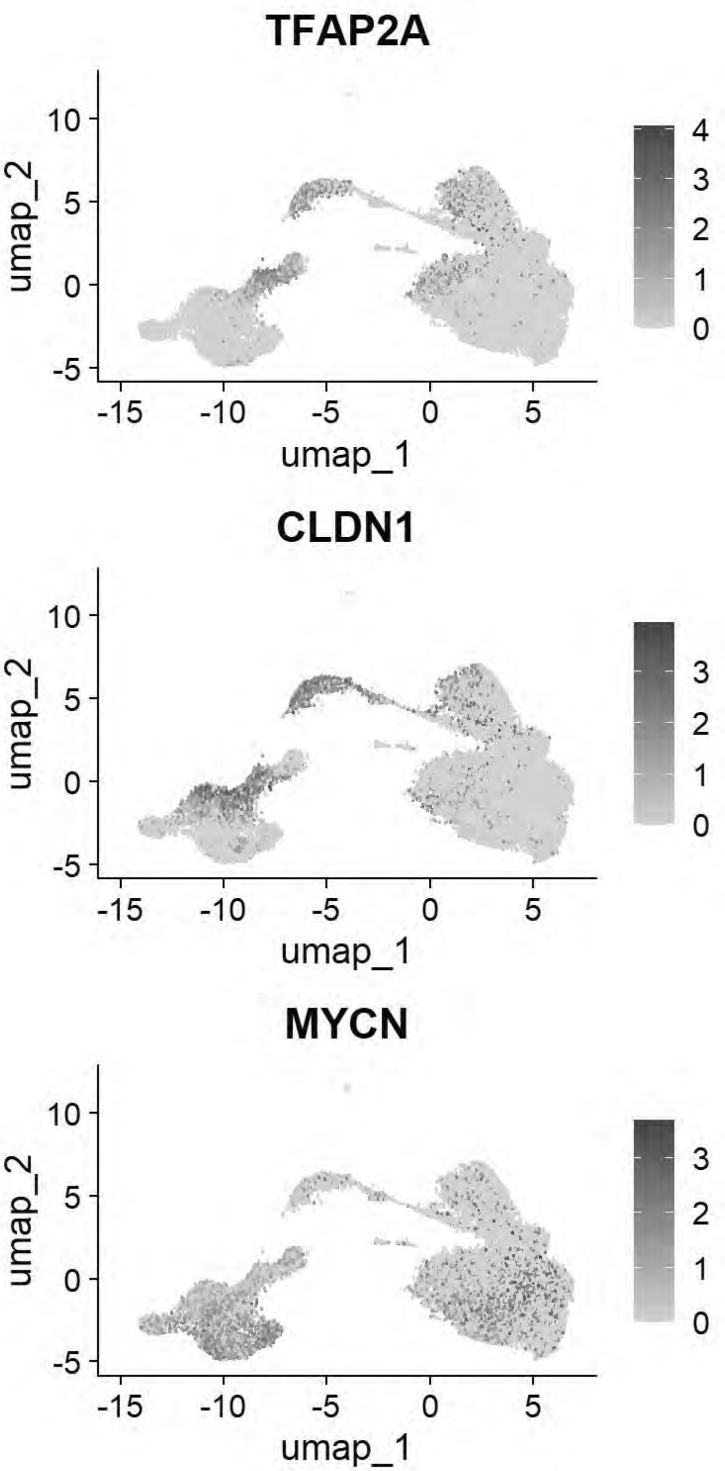


```
# Pajanoja: Endoderm
FeaturePlot(chick, features = c("SOX17","KRT17","HHEX"))
```

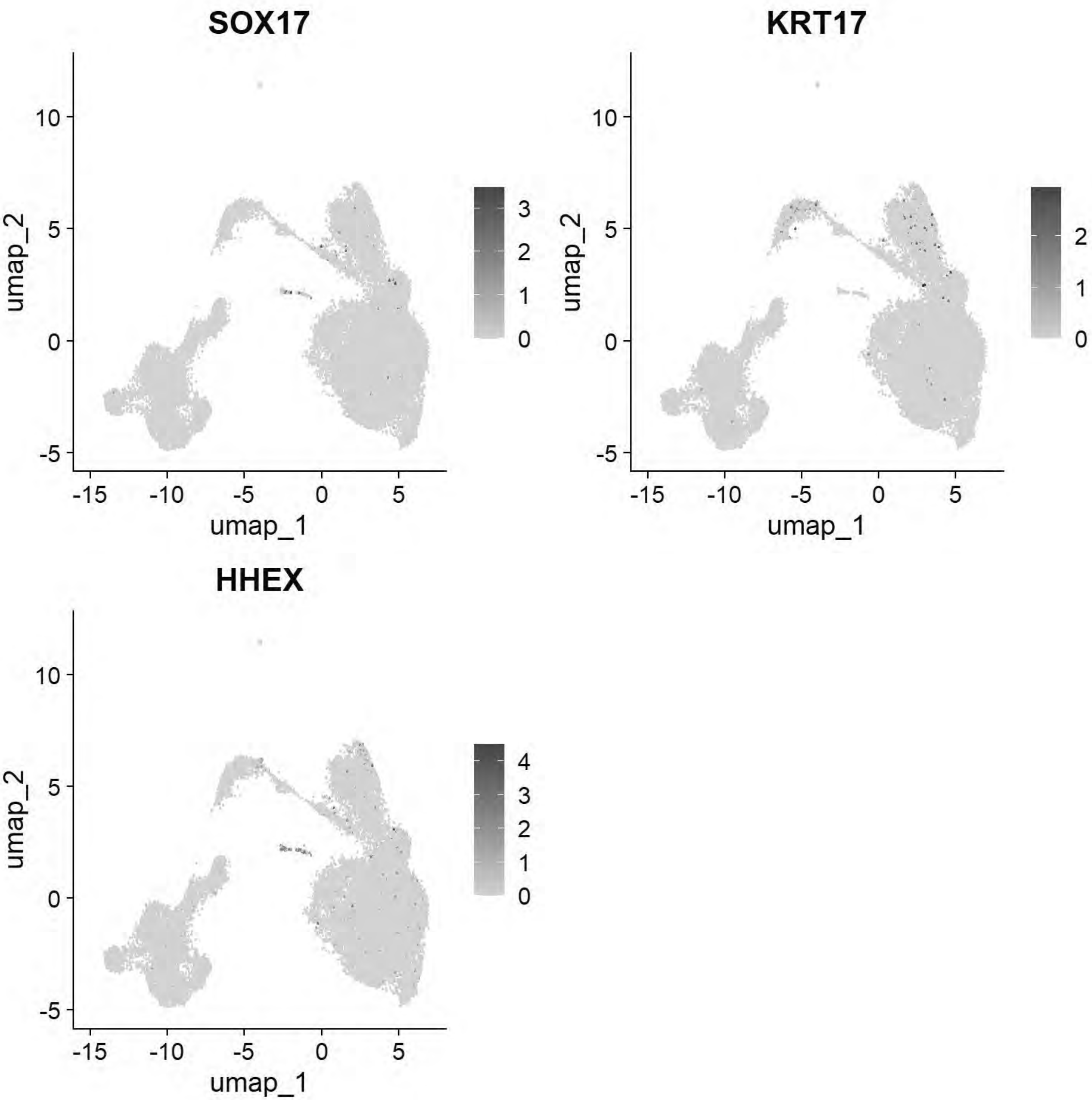


```r
# Pajanoja: Mesoderm
FeaturePlot(chick, features = c("ALX1","TWIST1","HAND2","PITX2"))
```

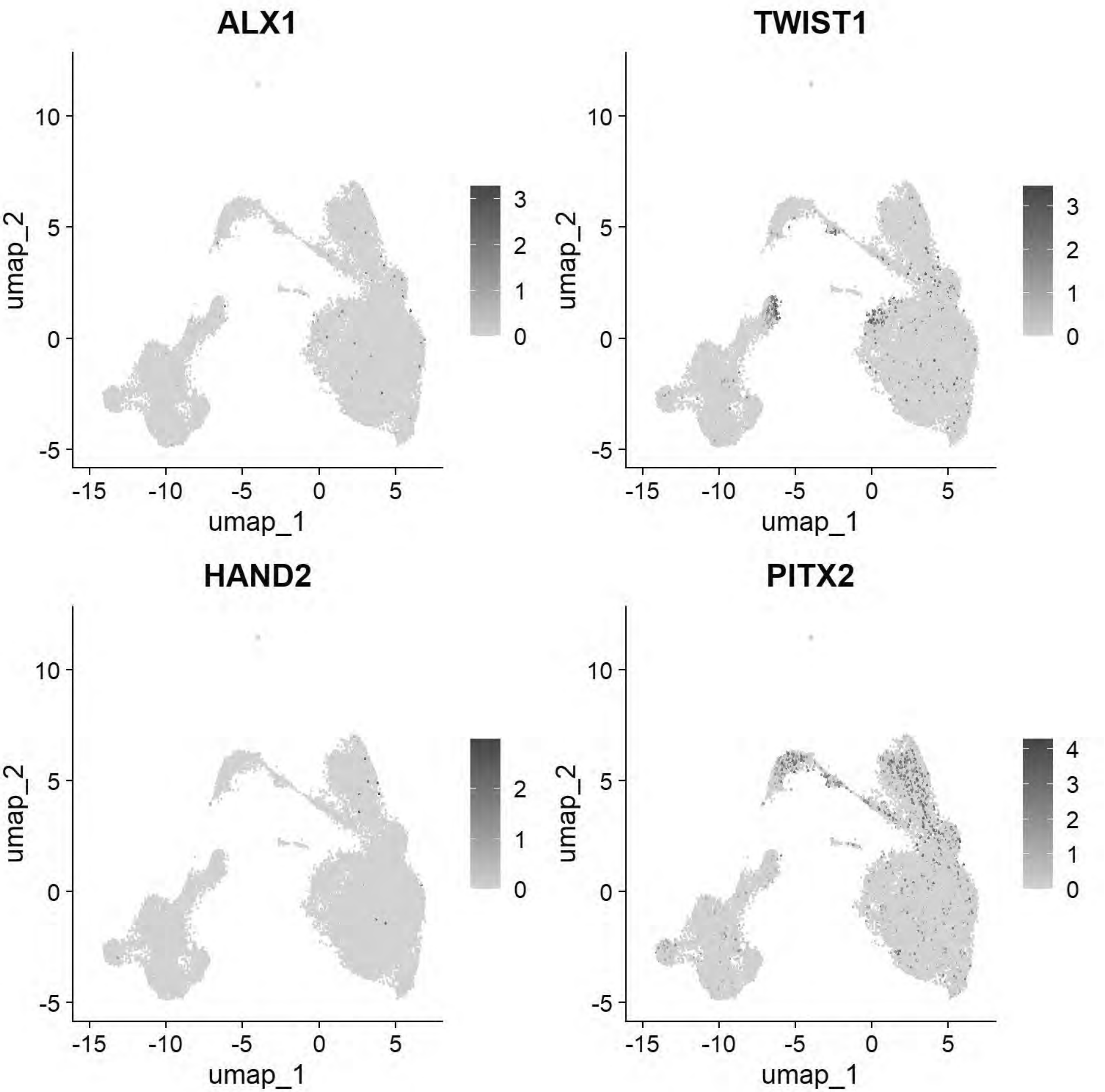


```
# Pajanoja: Notochord
FeaturePlot(chick, features = c("CHRD","TBXT","NOTO"))
```

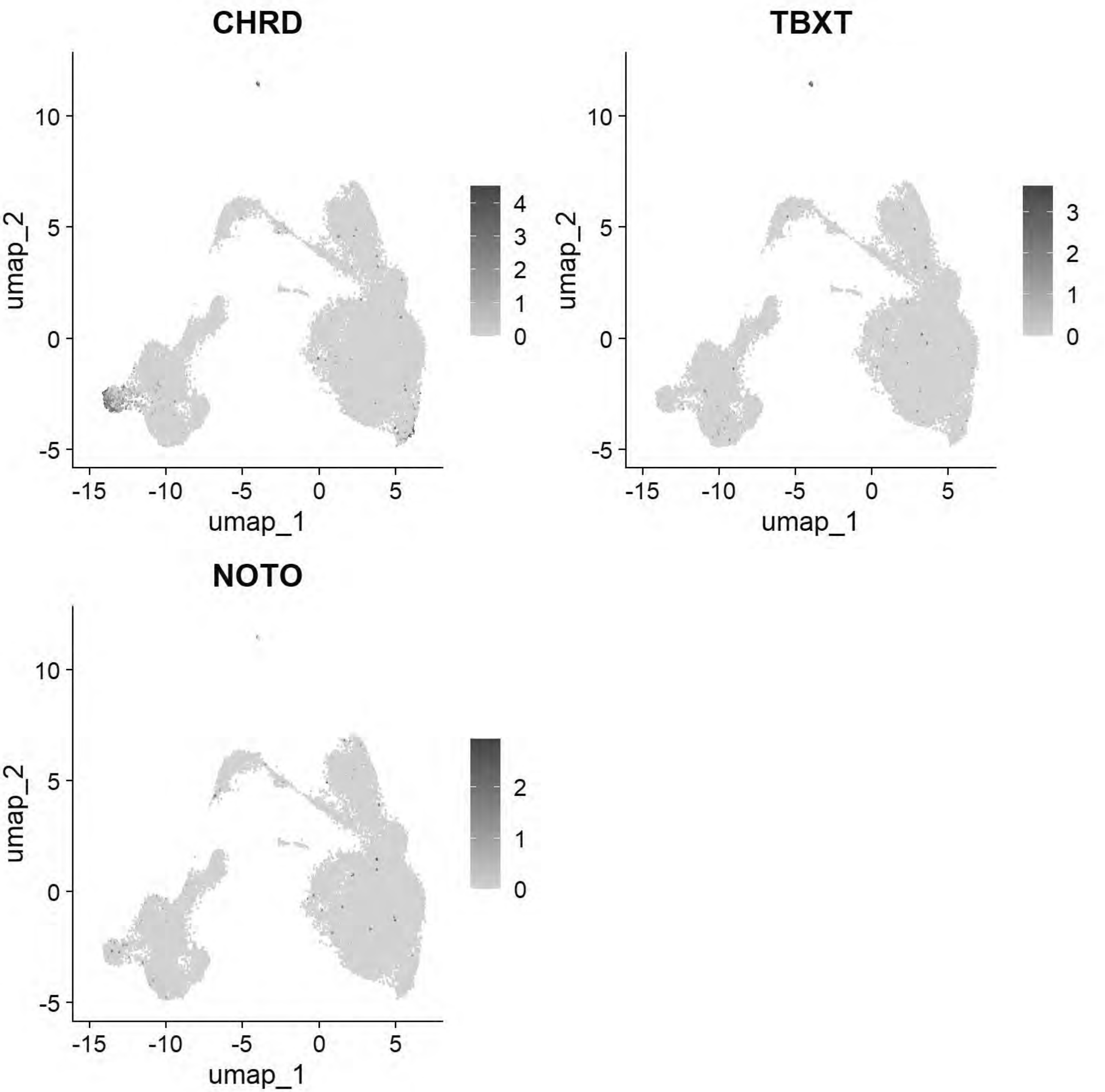


```
# Pajanoja: Ventral Neural Tube
FeaturePlot(chick, features=c("SHH","FOXA1","FOXA2"))
```

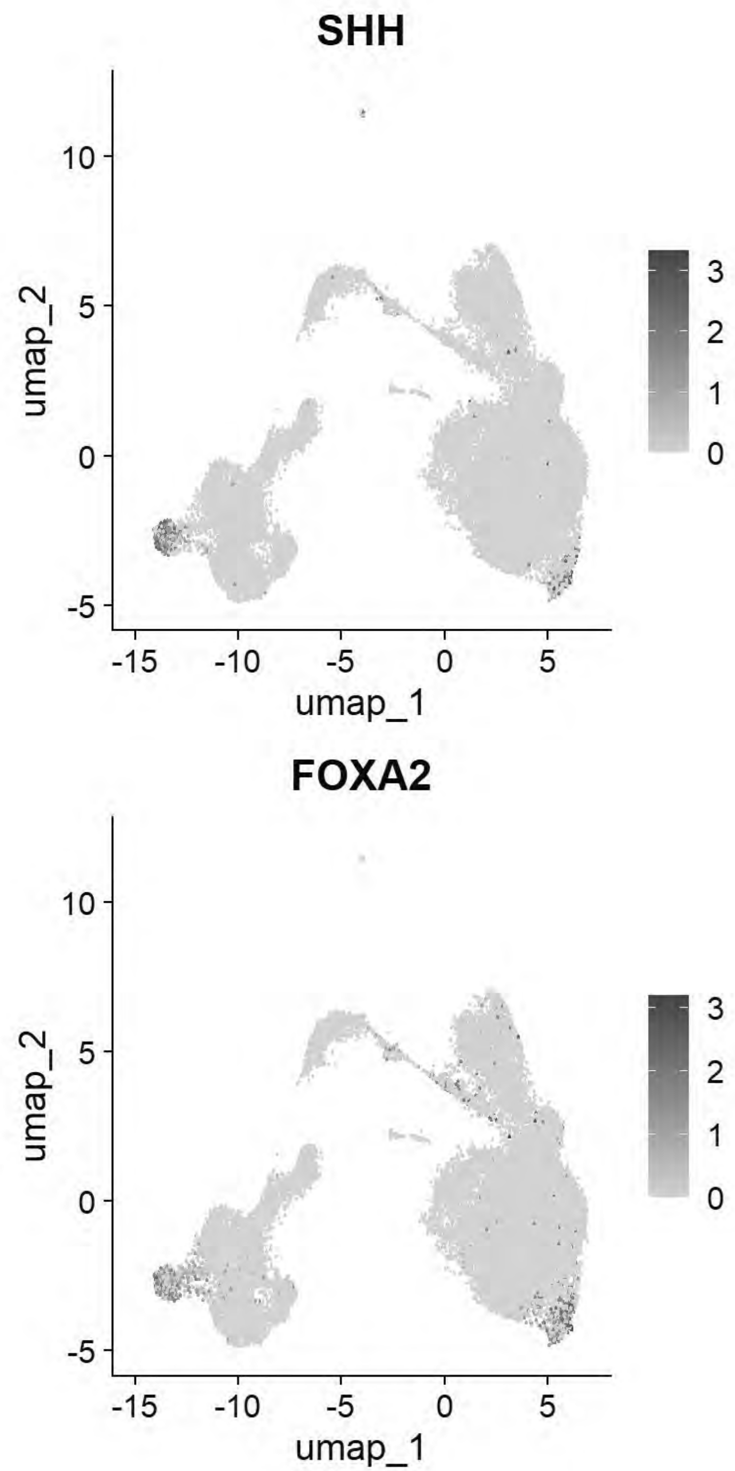


```
# Pajanoja: Neural Tube
FeaturePlot(chick, features=c("CDH2","HES5","MYCN","PAX2","NESTIN"))
```

```
## Warning: The following requested variables were not found: NESTIN
```

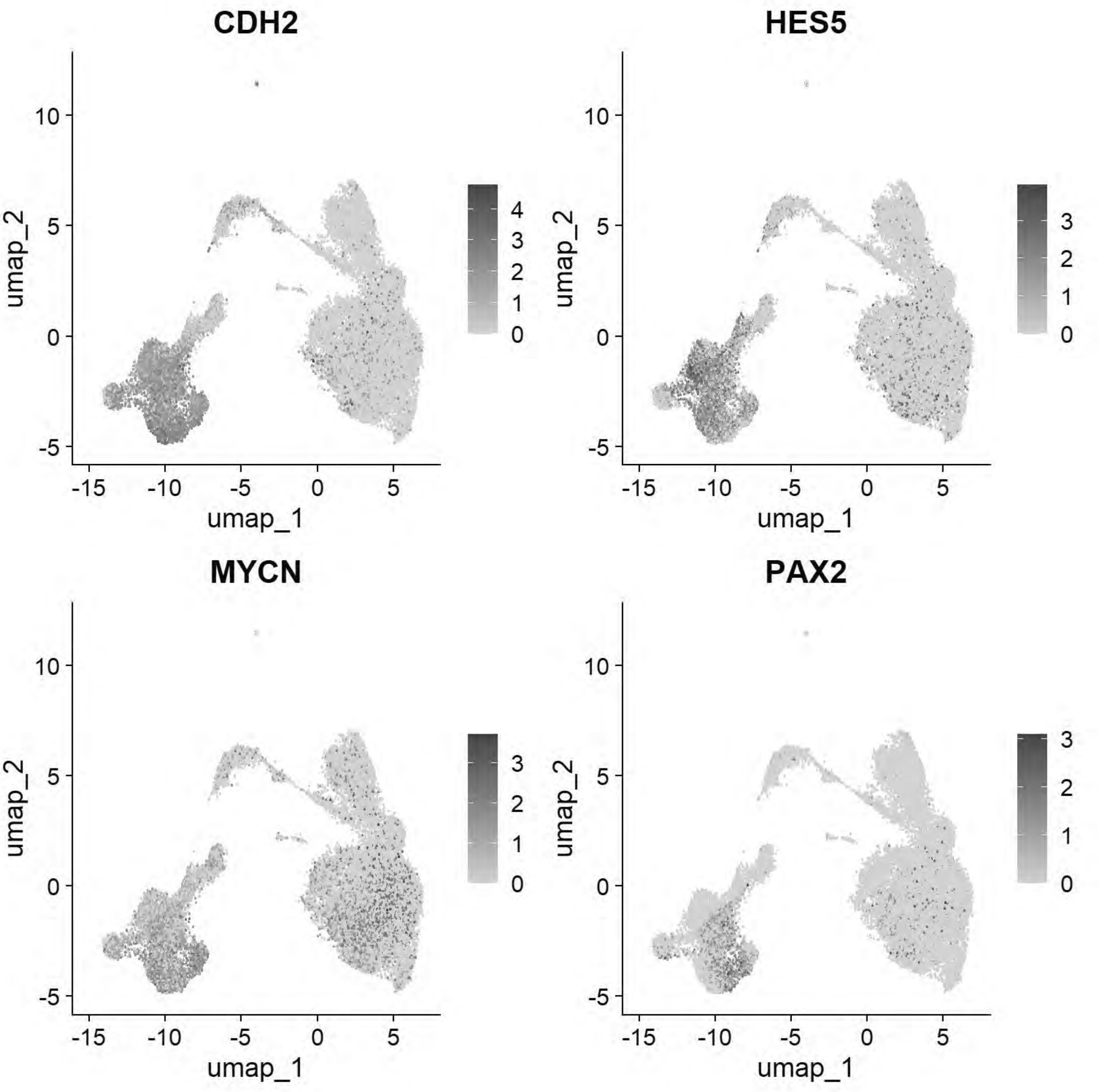


```
# Pajanoja: Non-Neural Ectoderm/Placode
FeaturePlot(chick, features=c("KRT24","KRT18","DLX5","SIX1","EYA2","TFAP2A"))
```

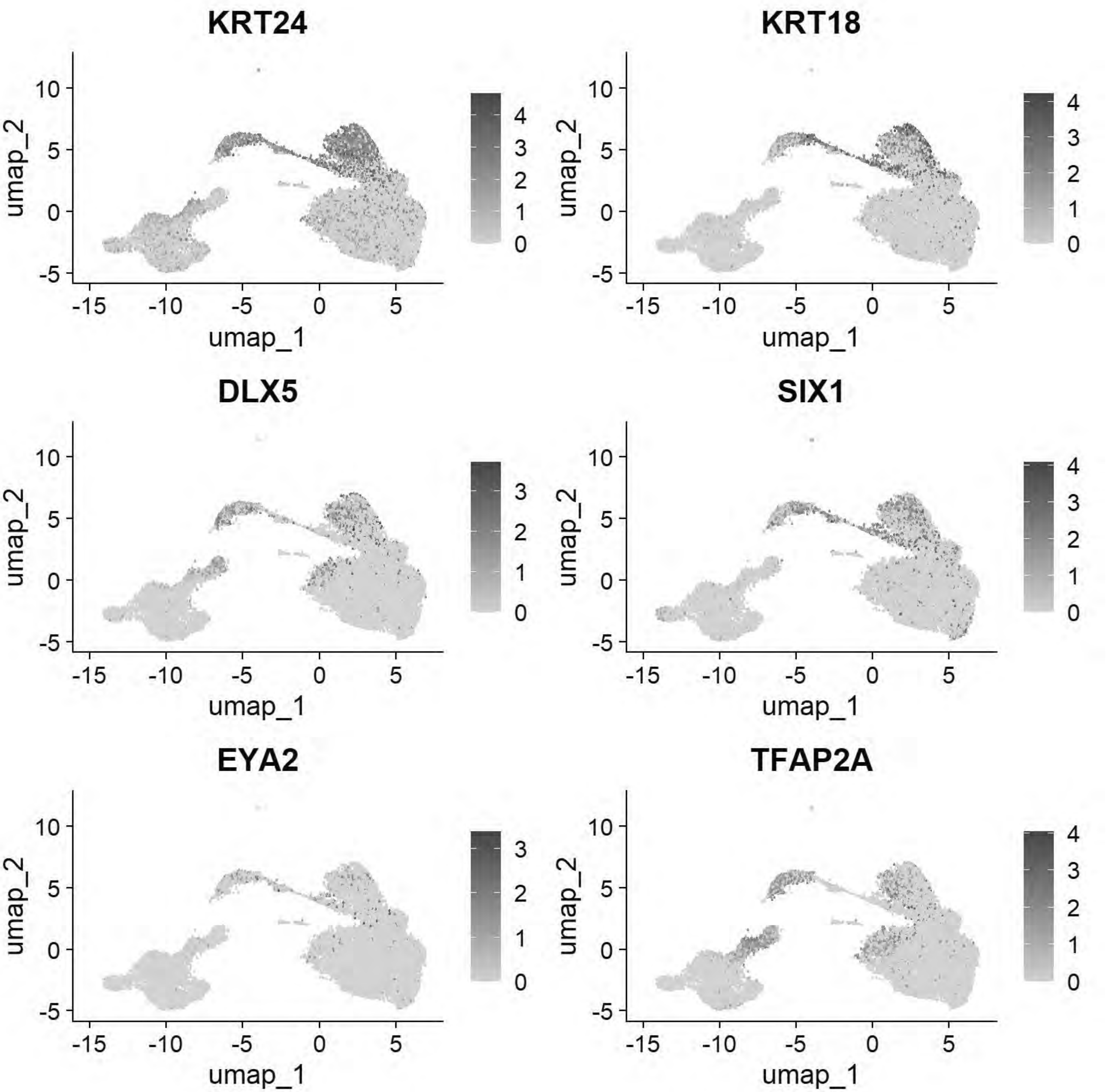


```
# Pajanoja: Neural crest
FeaturePlot(chick, features=c("FOXD3","TFAP2B","SOX9","ETS1","SNAI2"))
```

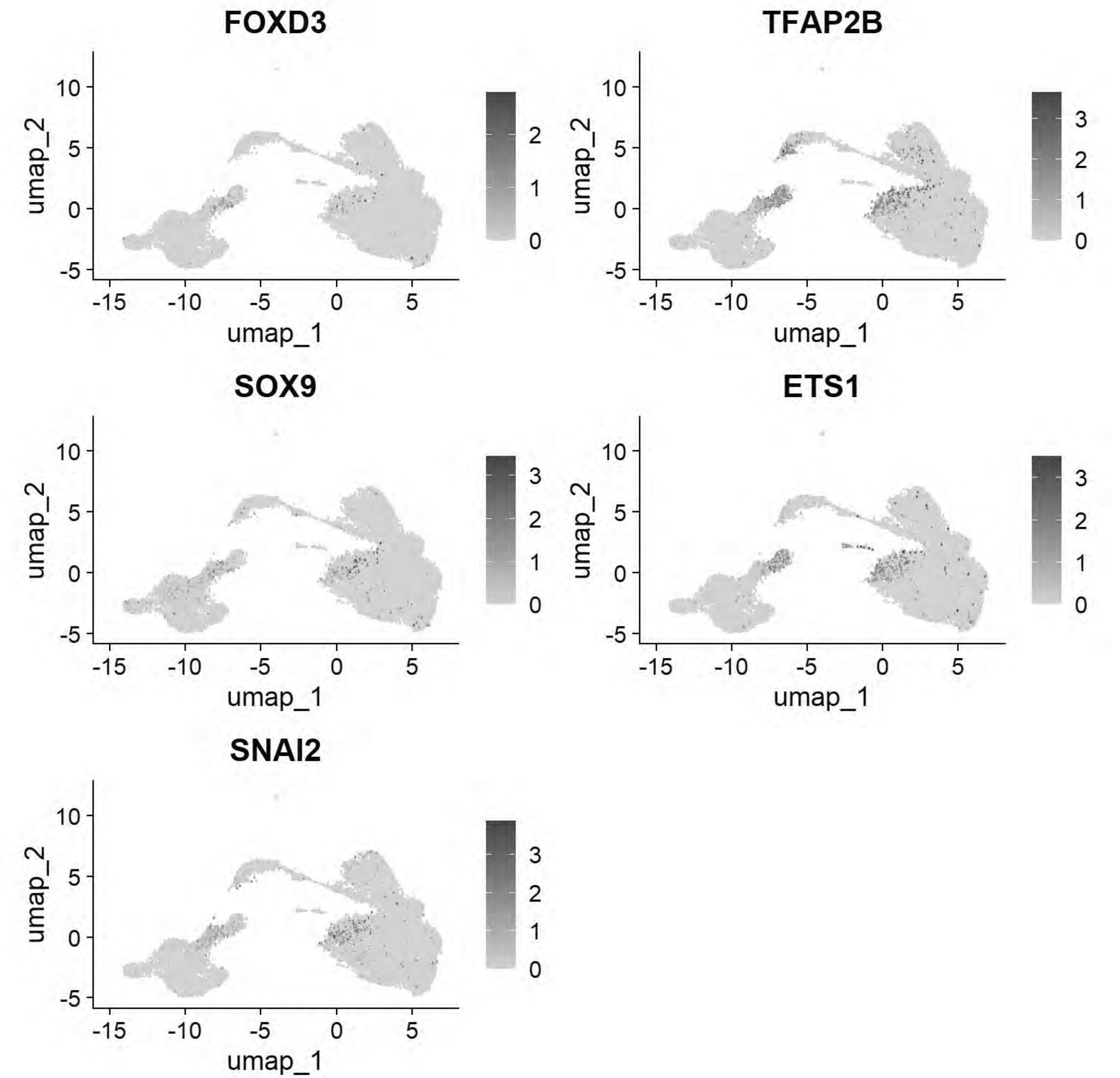


```
gc()
```

```
##             used   (Mb) gc trigger    (Mb)   max used    (Mb)
## Ncells   5069170  270.8   13651909   729.1   21331105  1139.3
## Vcells 998530263 7618.2 2342481578 17871.8 4574774951 34902.8
```

```
chick <- JoinLayers(chick)
chick.markers <- FindAllMarkers(chick, min.pct = 0.5, logfc.threshold = 0.5)
```

```
## Calculating cluster 0
```

```
## Calculating cluster 1
```

```
## Calculating cluster 2
```

```
## Calculating cluster 3
```

```
## Calculating cluster 4
```

```
## Calculating cluster 5
```

```
## Calculating cluster 6
```

```
## Calculating cluster 7
```

```
## Calculating cluster 8
```

```
## Calculating cluster 9
```

```
## Calculating cluster 10
```

```
## Calculating cluster 11
```

```
## Calculating cluster 12
```

```
## Calculating cluster 13
```

```
## Calculating cluster 14
```

```
gc()
```

```
##               used   (Mb) gc trigger    (Mb)   max used    (Mb)
## Ncells    5070863  270.9   13651909   729.1   21331105  1139.3
## Vcells 1199780791 9153.7 2342481578 17871.8 4574774951 34902.8
```

```
chick.markers %>%
  group_by(cluster) %>%
  top_n(n = 5, wt = avg_log2FC) -> top5
DotPlot(chick, features = unique(top5$gene))+coord_flip()
```

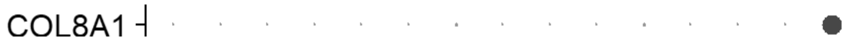

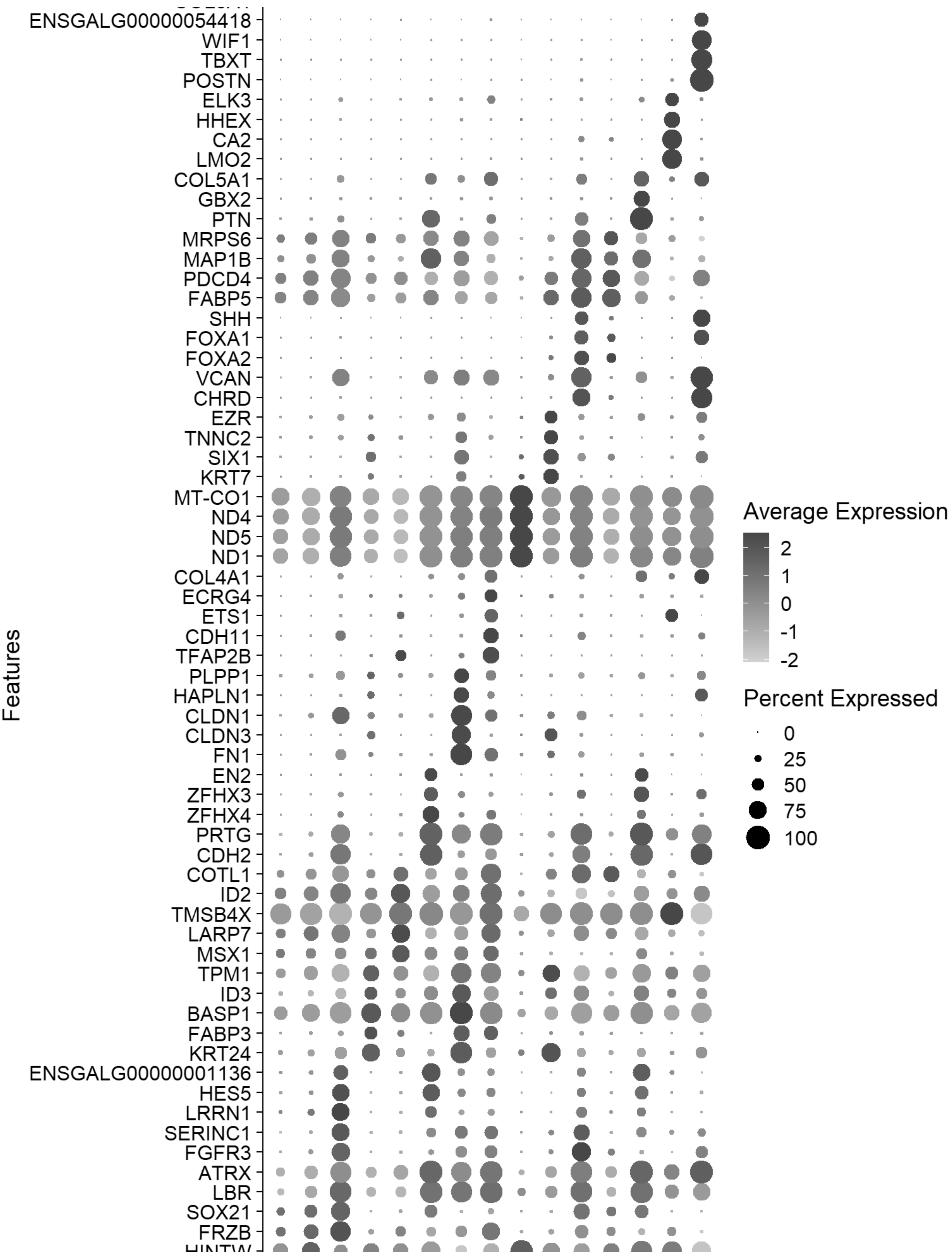
ENSGALG00000054418
WIF1
TBXT
POSTN
ELK3
HHEX
CA2
LMO2
COL5A1
GBX2
PTN
MRPS6
MAP1B
PDCD4
FABP5
SHH
FOXA1
FOXA2
VCAN
CHRD
EZR
TNNC2
SIX1
KRT7
MT-CO1
ND4
ND5
ND1
COL4A1
ECRG4
ETS1
CDH11
TFAP2B
PLPP1
HAPLN1
CLDN1
CLDN3
FN1
EN2
ZFHX3
ZFHX4
PRTG
CDH2
COTL1
ID2
TMSB4X
LARP7
MSX1
TPM1
ID3
BASP1
FABP3
KRT24
ENSGALG00000001136
HES5
LRRN1
SERINC1
FGFR3
ATRX
LBR
SOX21
FRZB
Features
Average Expression
2
1
0
-1
-2
Percent Expressed
0
25
50
75
100

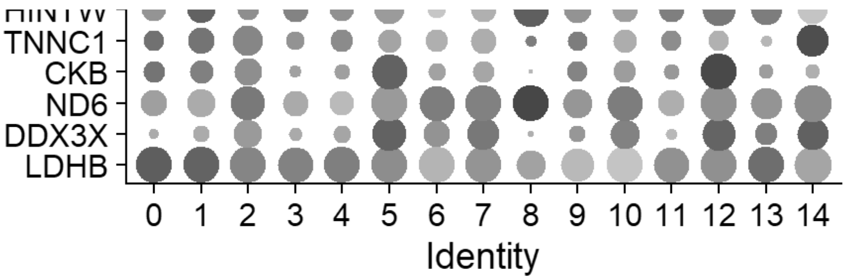


```
gc()
```

```
##               used   (Mb) gc trigger    (Mb)   max used    (Mb)
## Ncells    4984289  266.2   13651909   729.1   21331105  1139.3
## Vcells 1201721386 9168.5 2342481578 17871.8 4574774951 34902.8
```

```
FeaturePlot(chick, features = c("PAX7","SNAI2","SOX9","SOX10"))
```

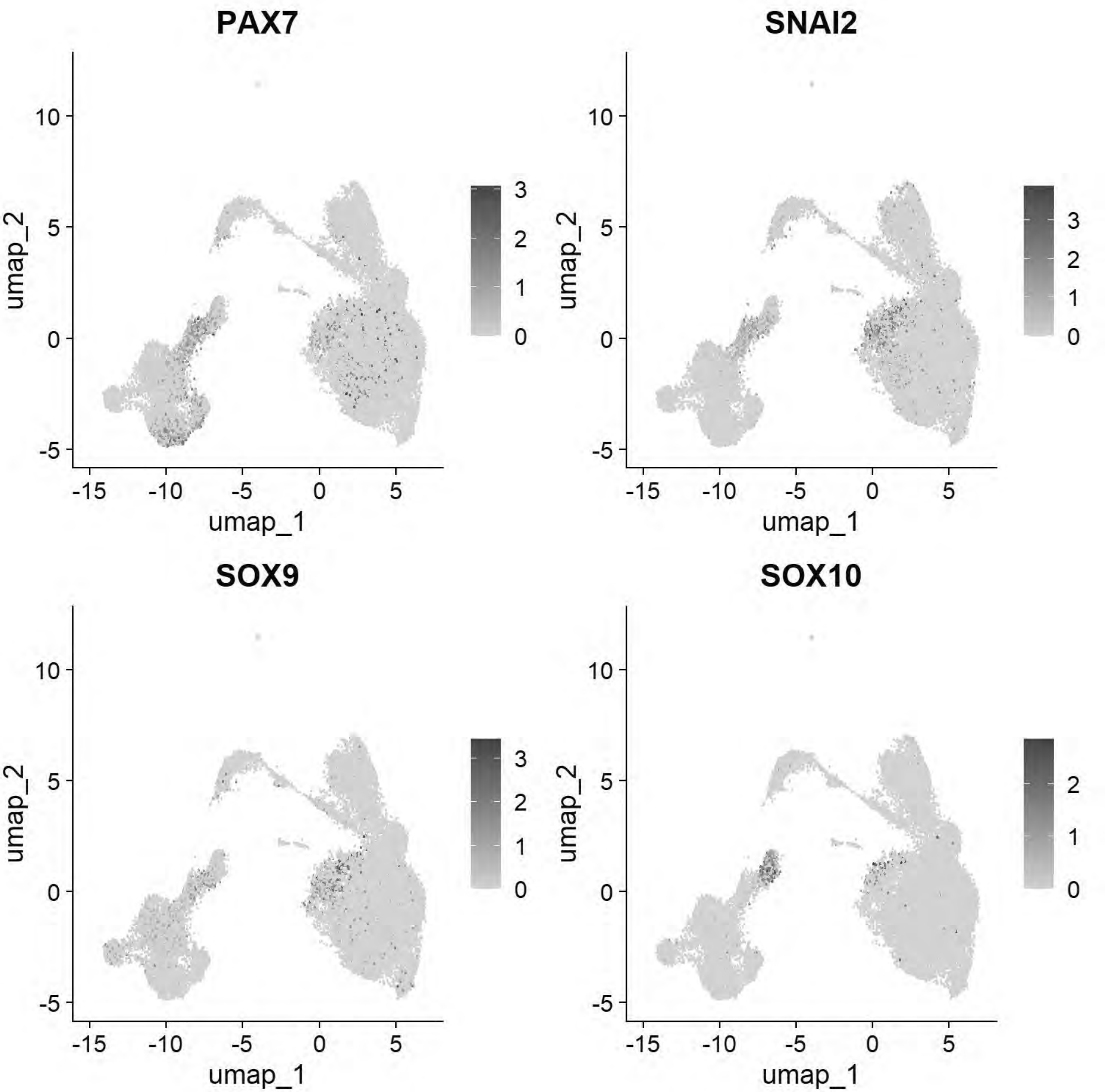


```
FeaturePlot(chick, features = c("SHH","DRAXIN","CDH6","SOX2"))
```

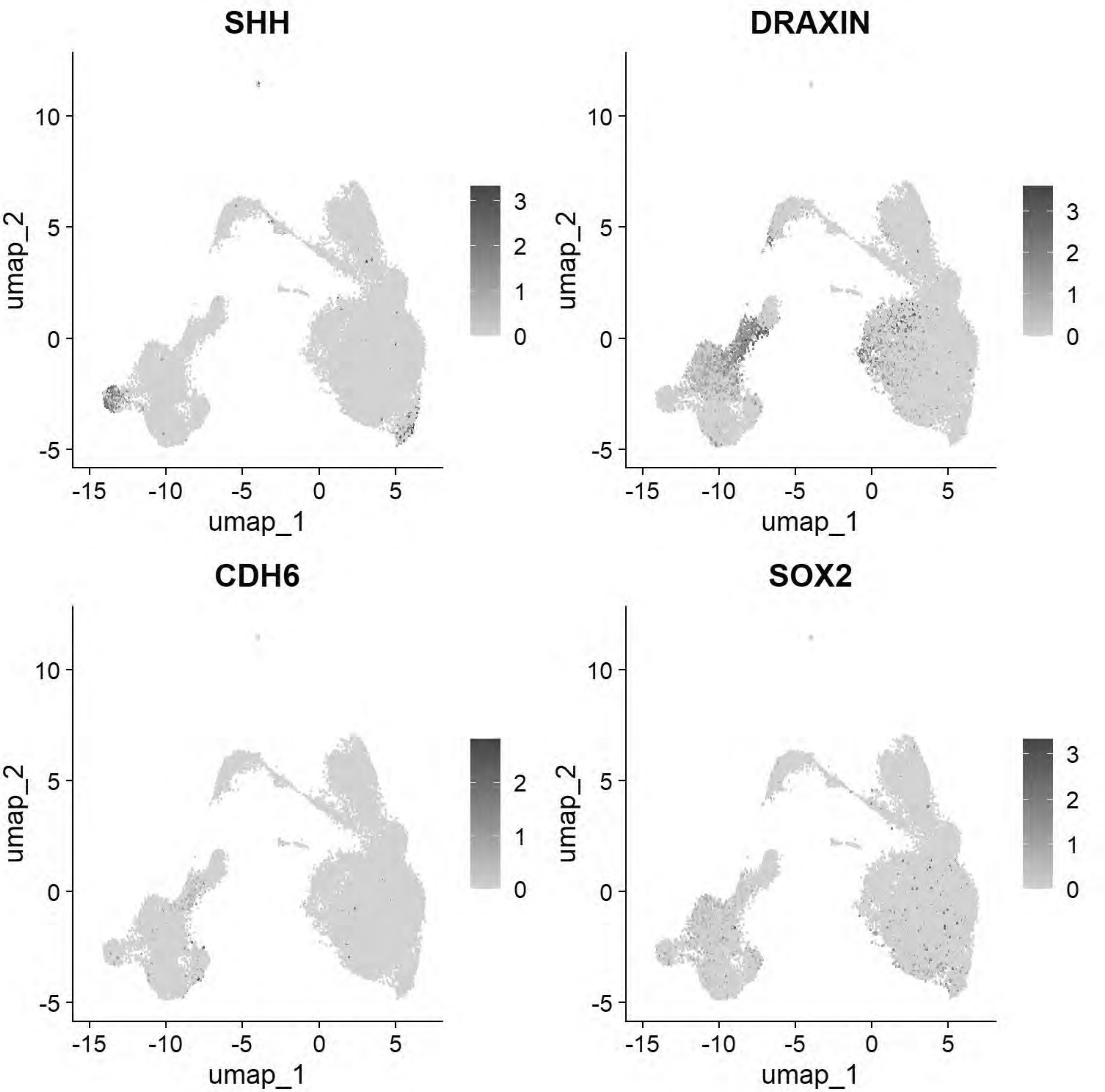


```
FeaturePlot(chick, features = c("PAX7","SHH","PAX6","OLIG2"))
```

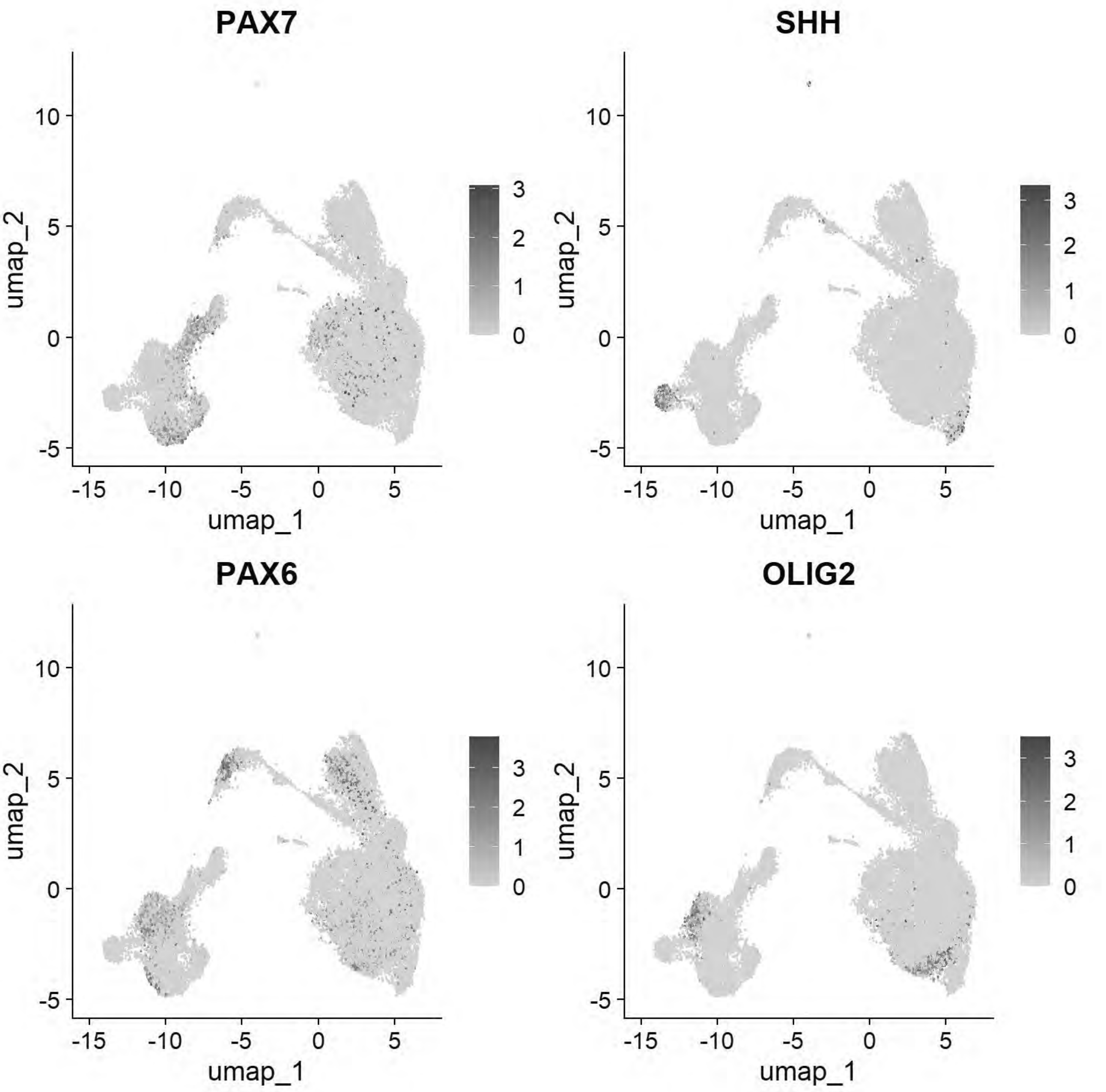


```
FeaturePlot(chick, features = c("PAX6","PAX2","PAX7","OLIG2"))
```

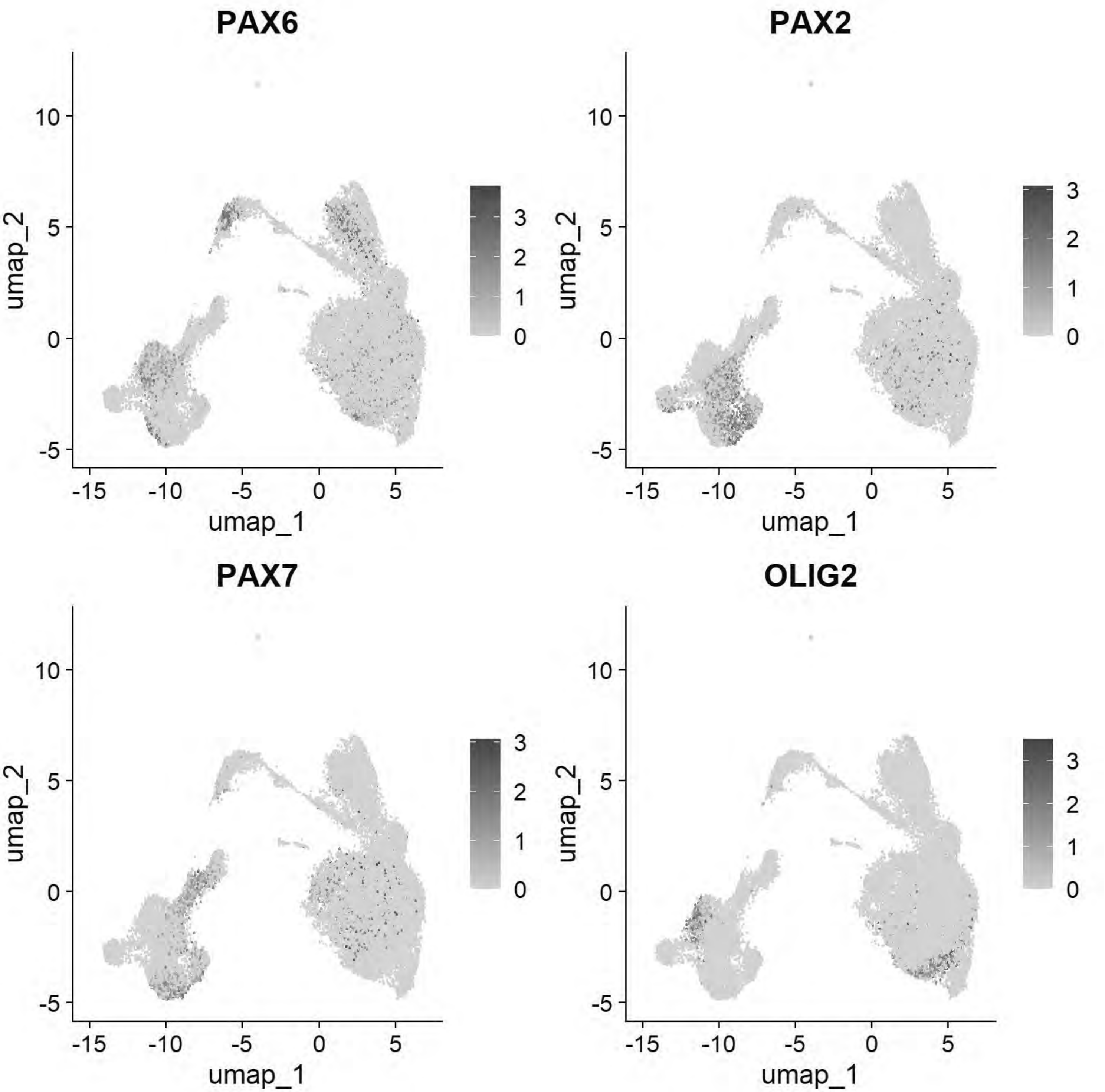


```
DimPlot(chick, group.by = "stage")
```

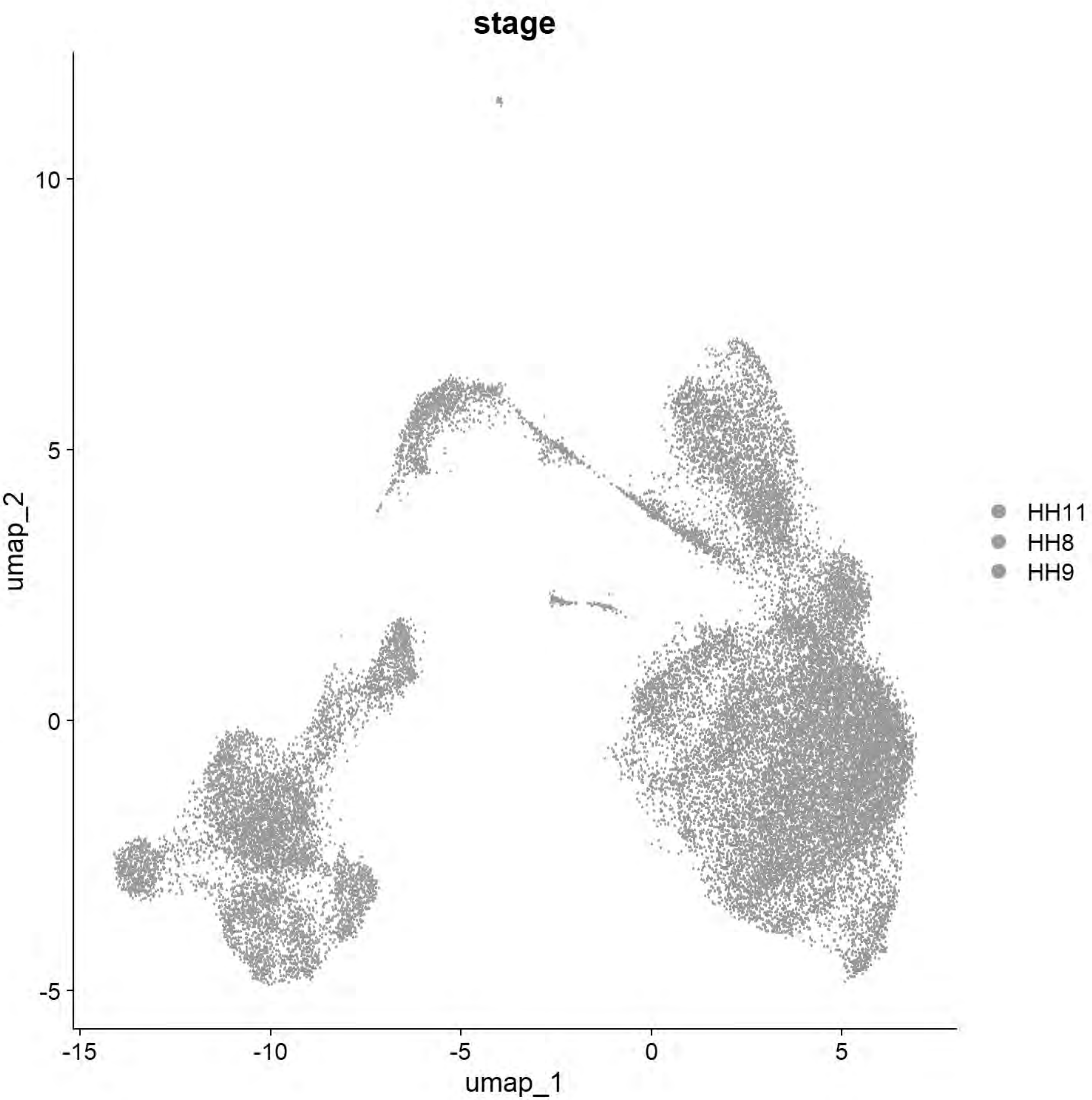


```
#FeaturePlot(chick, features = c("PAX7","TFAP2A","DLX5","BMP4","MSX1","DRAXIN","SOX10"))
```

```
chick
```

```
## An object of class Seurat
## 18523 features across 31437 samples within 1 assay
## Active assay: RNA (18523 features, 2000 variable features)
##  3 layers present: data, counts, scale.data
##  3 dimensional reductions calculated: pca, harmony, umap
```

```
Idents(chick) <- "seurat_clusters"
#eliminating for low quality
chick <- subset(chick, idents = c("8","13","14"), invert = TRUE)
chick
```

```
## An object of class Seurat
## 18523 features across 30386 samples within 1 assay
## Active assay: RNA (18523 features, 2000 variable features)
##  3 layers present: data, counts, scale.data
##  3 dimensional reductions calculated: pca, harmony, umap
```

```
chick[["RNA"]] <- split(chick[["RNA"]], f = chick$sample_id)
```

```
## Splitting 'counts', 'data' layers. Not splitting 'scale.data'. If you would like to split o
ther layers, set in `layers` argument.
```

```
chick <- NormalizeData(chick)
```

```
## Normalizing layer: counts.Pajanoja2023_HH8_1
```

```
## Normalizing layer: counts.Pajanoja2023_HH8_2
```

```
## Normalizing layer: counts.Pajanoja2023_HH9_1
```

```
## Normalizing layer: counts.Pajanoja2023_HH9_2
```

```
## Normalizing layer: counts.Pajanoja2025_HH11
```

```
chick <- CellCycleScoring(chick, s.features = cc.genes$s.genes, g2m.features = cc.genes$g2m.ge
nes)
```

```
## Warning: The following features are not present in the object: TYMS, CDCA7,
## PRIM1, MLF1IP, RAD51AP1, not searching for symbol synonyms
```

```
## Warning: The following features are not present in the object: MKI67, FAM64A,
## AURKB, KIF20B, CDC25C, CDCA2, CDCA8, PSRC1, CENPA, not searching for symbol
## synonyms
```

```
## Warning: The following features are not present in the object: TYMS, CDCA7,
## PRIM1, MLF1IP, RAD51AP1, not searching for symbol synonyms
```

```
## Warning: The following features are not present in the object: MKI67, FAM64A,
## AURKB, KIF20B, CDC25C, CDCA2, CDCA8, PSRC1, CENPA, not searching for symbol
```

```
## synonyms
```

```
## Warning: The following features are not present in the object: TYMS, CDCA7,
## PRIM1, MLF1IP, RAD51AP1, not searching for symbol synonyms
```

```
## Warning: The following features are not present in the object: MKI67, FAM64A,
## AURKB, KIF20B, CDC25C, CDCA2, CDCA8, PSRC1, CENPA, not searching for symbol
## synonyms
```

```
## Warning: The following features are not present in the object: TYMS, CDCA7,
## PRIM1, MLF1IP, RAD51AP1, not searching for symbol synonyms
```

```
## Warning: The following features are not present in the object: MKI67, FAM64A,
## AURKB, KIF20B, CDC25C, CDCA2, CDCA8, PSRC1, CENPA, not searching for symbol
## synonyms
```

```
## Warning: The following features are not present in the object: TYMS, CDCA7,
## PRIM1, MLF1IP, RAD51AP1, not searching for symbol synonyms
```

```
## Warning: The following features are not present in the object: MKI67, FAM64A,
## AURKB, KIF20B, CDC25C, CDCA2, CDCA8, PSRC1, CENPA, not searching for symbol
## synonyms
```

```
chick <- FindVariableFeatures(chick)
```

```
## Finding variable features for layer counts.Pajanoja2023_HH8_1
```

```
## Finding variable features for layer counts.Pajanoja2023_HH8_2
```

```
## Finding variable features for layer counts.Pajanoja2023_HH9_1
```

```
## Finding variable features for layer counts.Pajanoja2023_HH9_2
```

```
## Finding variable features for layer counts.Pajanoja2025_HH11
```

```
chick <- ScaleData(chick, vars.to.regress = c("percent.mt", "S.Score", "G2M.Score"))
```

```
## Regressing out percent.mt, S.Score, G2M.Score
```

```
## Centering and scaling data matrix
```

```
## Warning: Different features in new layer data than already exists for
## scale.data
```

```
chick <- RunPCA(chick, npcs = 50, verbose = FALSE)

chick <- IntegrateLayers(object = chick, method = HarmonyIntegration, orig.reduction = "pca",
new.reduction = "harmony", assay = "RNA", group.by = "sample_id", verbose = TRUE)
```

```
## Transposing data matrix
```

```
## Using automatic lambda estimation
```

```
## Initializing state using k-means centroids initialization
```

```
## Harmony 1/10
```

```
## Harmony 2/10
```

```
## Harmony 3/10
```

```
## Harmony 4/10
```

```
## Harmony converged after 4 iterations
```

```
chick <- FindNeighbors(chick, reduction = "harmony", dims = 1:30)
```

```
## Computing nearest neighbor graph
```

```
## Computing SNN
```

```
chick <- RunUMAP(chick, reduction = "harmony", dims = 1:30)
```

```
## 23:39:17 UMAP embedding parameters a = 0.9922 b = 1.112
```

```
## 23:39:17 Read 30386 rows and found 30 numeric columns
```

```
## 23:39:17 Using Annoy for neighbor search, n_neighbors = 30
```

```
## 23:39:17 Building Annoy index with metric = cosine, n_trees = 50
```

```
## 0%   10   20   30   40   50   60   70   80   90   100%
```

```
## [----|----|----|----|----|----|----|----|----|----|
```

```
## **************************************************|
## 23:39:20 Writing NN index file to temp file C:\Users\RANEES~1\AppData\Local\Temp\RtmpqcWgVb
\file6a2c61136471
## 23:39:20 Searching Annoy index using 1 thread, search_k = 3000
## 23:39:28 Annoy recall = 100%
## 23:39:28 Commencing smooth kNN distance calibration using 1 thread with target n_neighbors
= 30
## 23:39:30 Initializing from normalized Laplacian + noise (using RSpectra)
## 23:39:30 Commencing optimization for 200 epochs, with 1442294 positive edges
## 23:39:30 Using rng type: pcg
## 23:39:57 Optimization finished
```

```
DimPlot(chick)
```

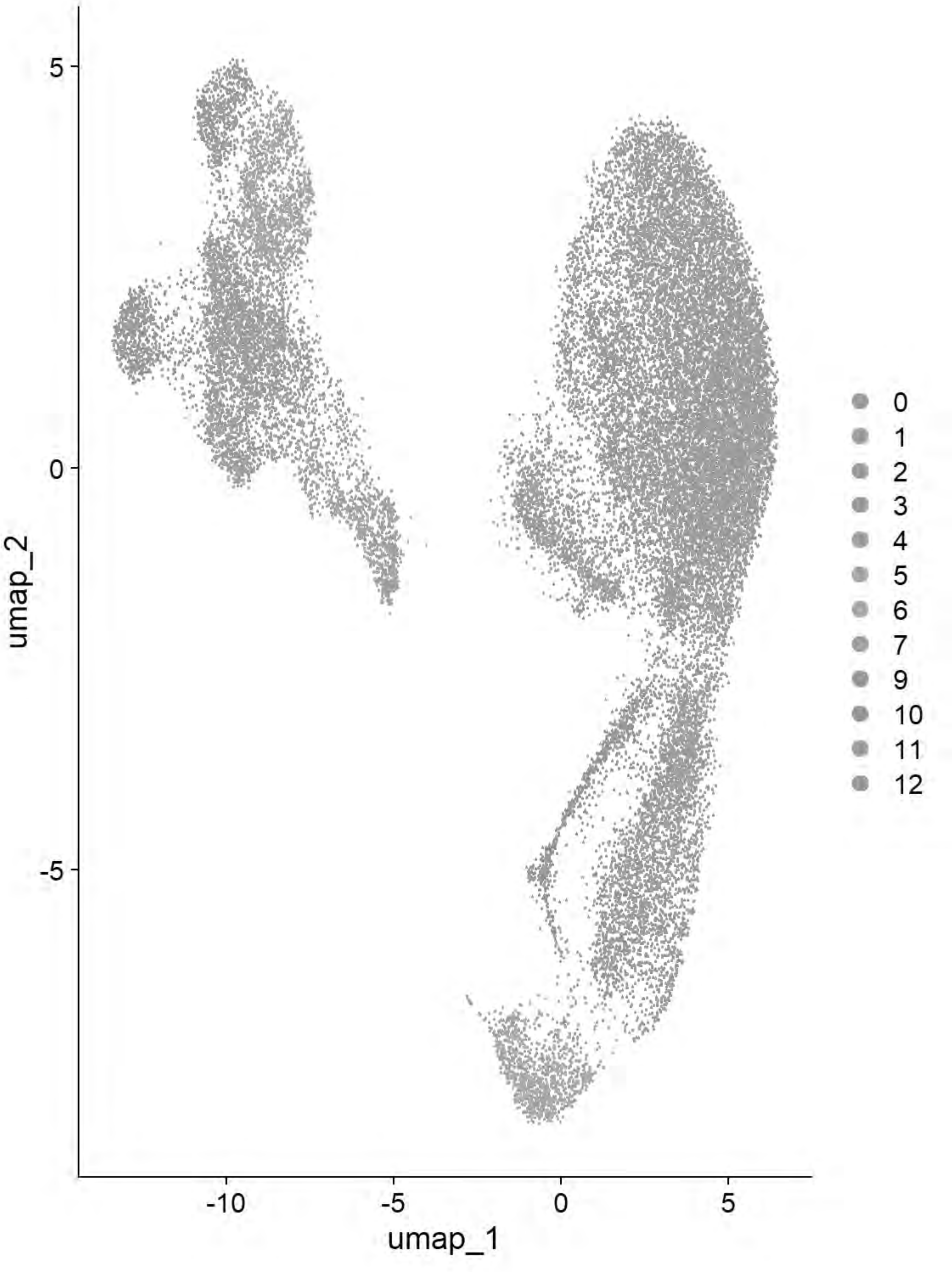


```
chick <- FindClusters(chick, resolution = 0.4)
```

```
## Modularity Optimizer version 1.3.0 by Ludo Waltman and Nees Jan van Eck
##
## Number of nodes: 30386
## Number of edges: 972783
##
## Running Louvain algorithm...
## Maximum modularity in 10 random starts: 0.8942
```

```
## Number of communities: 13
## Elapsed time: 4 seconds
```

```
## 1 singletons identified. 12 final clusters.
```

```
qc_tester(chick)
```

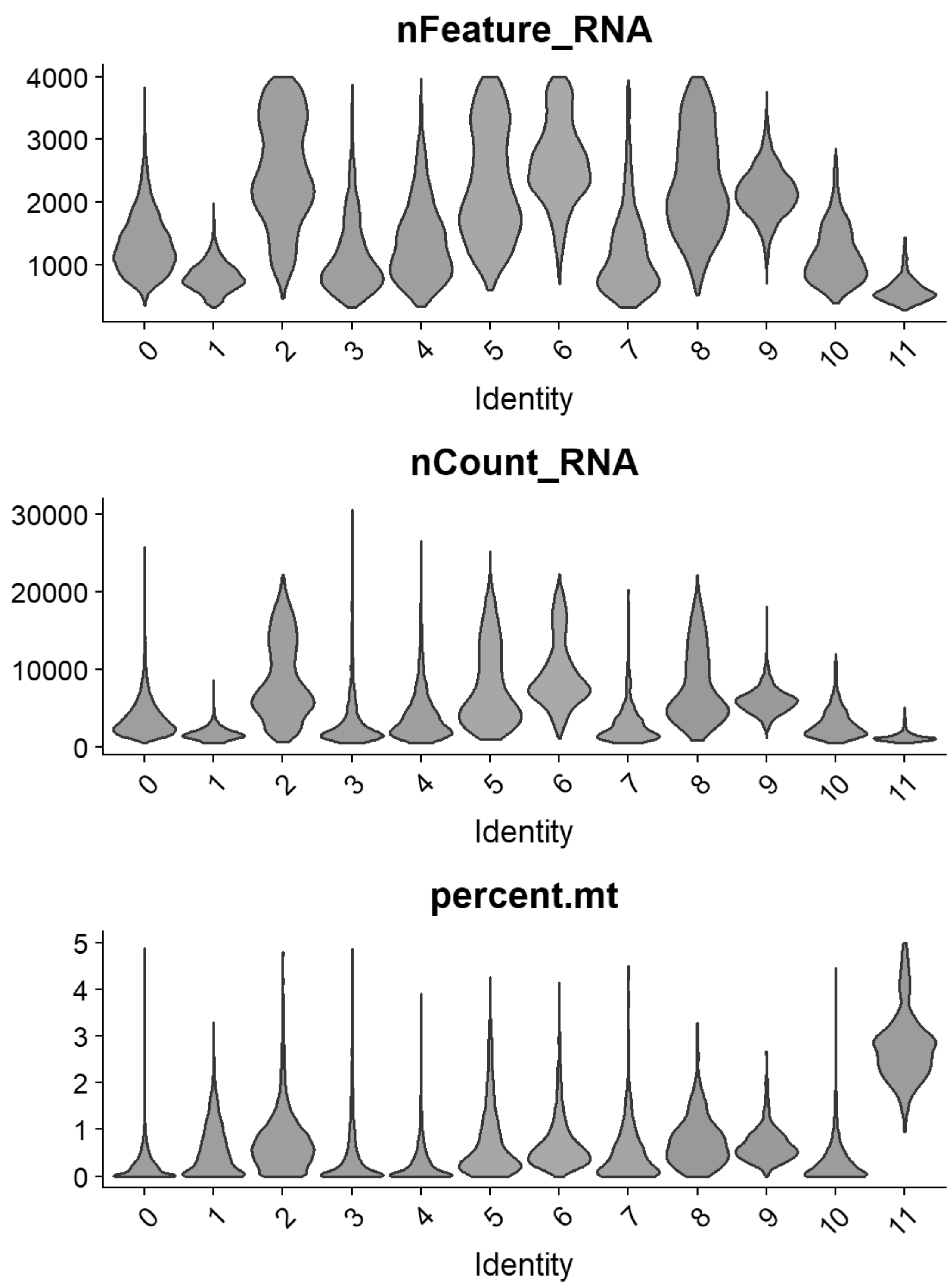


```
chick = FindClusters(chick, resolution= seq(0,2,0.1),n.start=10, verbose = FALSE)
p=clustree(chick, prefix= "RNA_snn_res.")
```

```
print(p)
```

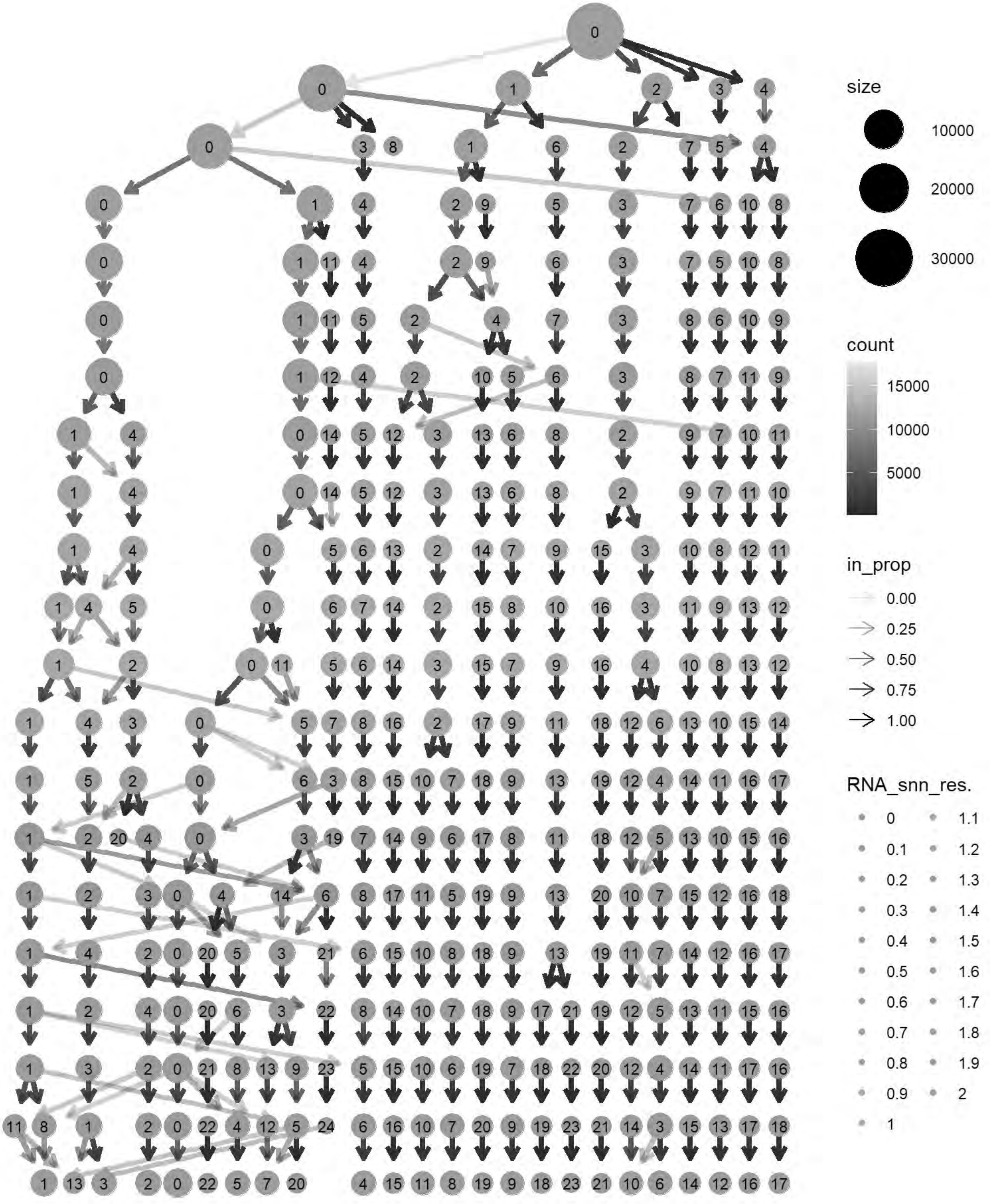
size
10000
20000
30000
count
15000
10000
5000
in_prop
0.00
0.25
0.50
0.75
1.00
RNA_snn_res.
0
0.1
0.2
0.3
0.4
0.5
0.6
0.7
0.8
0.9
1
1.1
1.2
1.3
1.4
1.5
1.6
1.7
1.8
1.9
2

```
cols= grep("^RNA_snn_res.",names(chick@meta.data))
chick@meta.data[cols] <- NULL
chick <- FindClusters(chick, resolution = 0.7, verbose = TRUE)
```

```
## Modularity Optimizer version 1.3.0 by Ludo Waltman and Nees Jan van Eck
##
## Number of nodes: 30386
## Number of edges: 972783
##
## Running Louvain algorithm...
## Maximum modularity in 10 random starts: 0.8568
## Number of communities: 16
## Elapsed time: 5 seconds
```

```
## 1 singletons identified. 15 final clusters.
```

```
p6=DimPlot(chick, label = TRUE)+NoLegend()
print(p6)
```

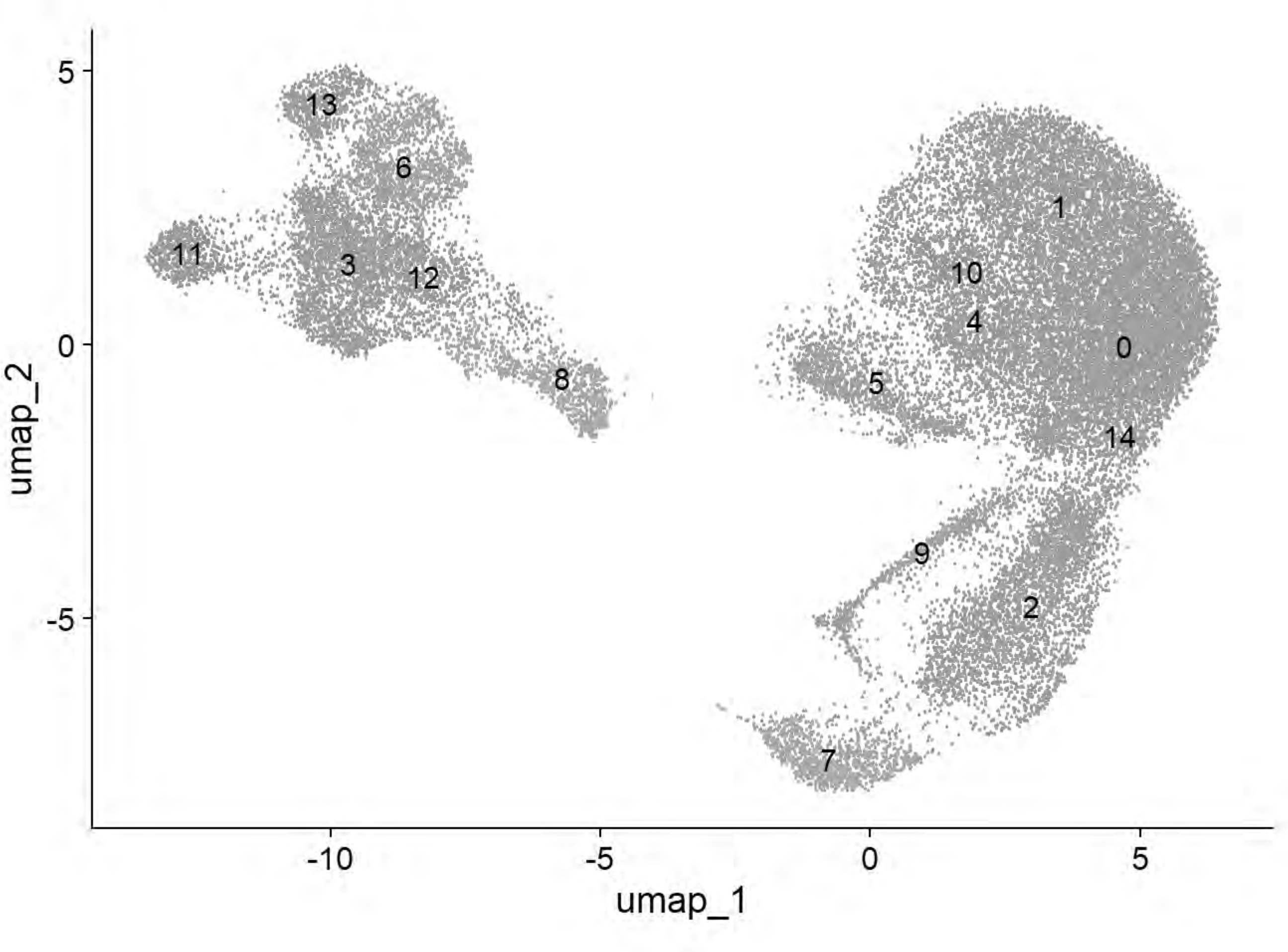


```
p6=DimPlot(chick, group.by = "Phase")
print(p6)
```

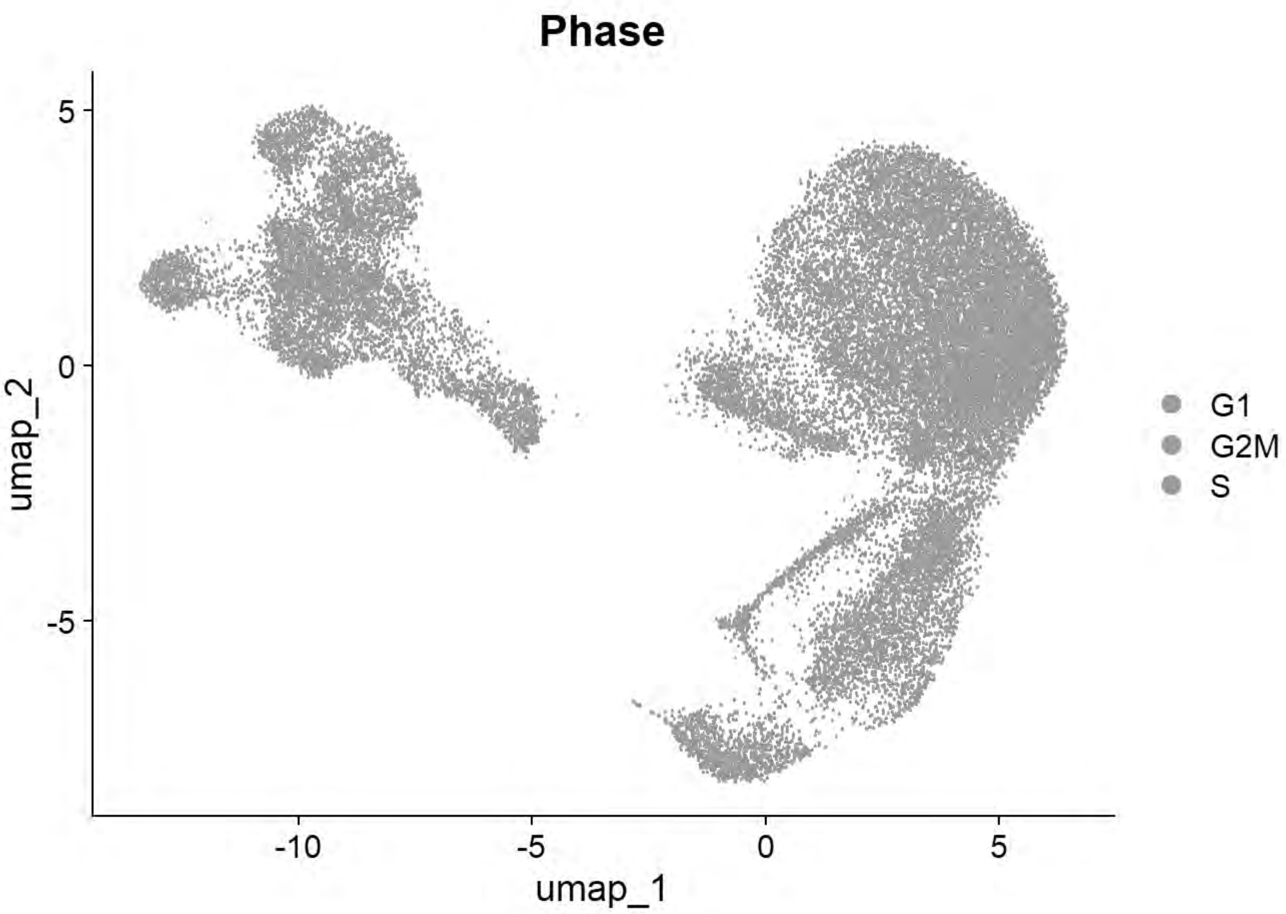


```
p6=DimPlot(chick, group.by = "sample_id")
print(p6)
```

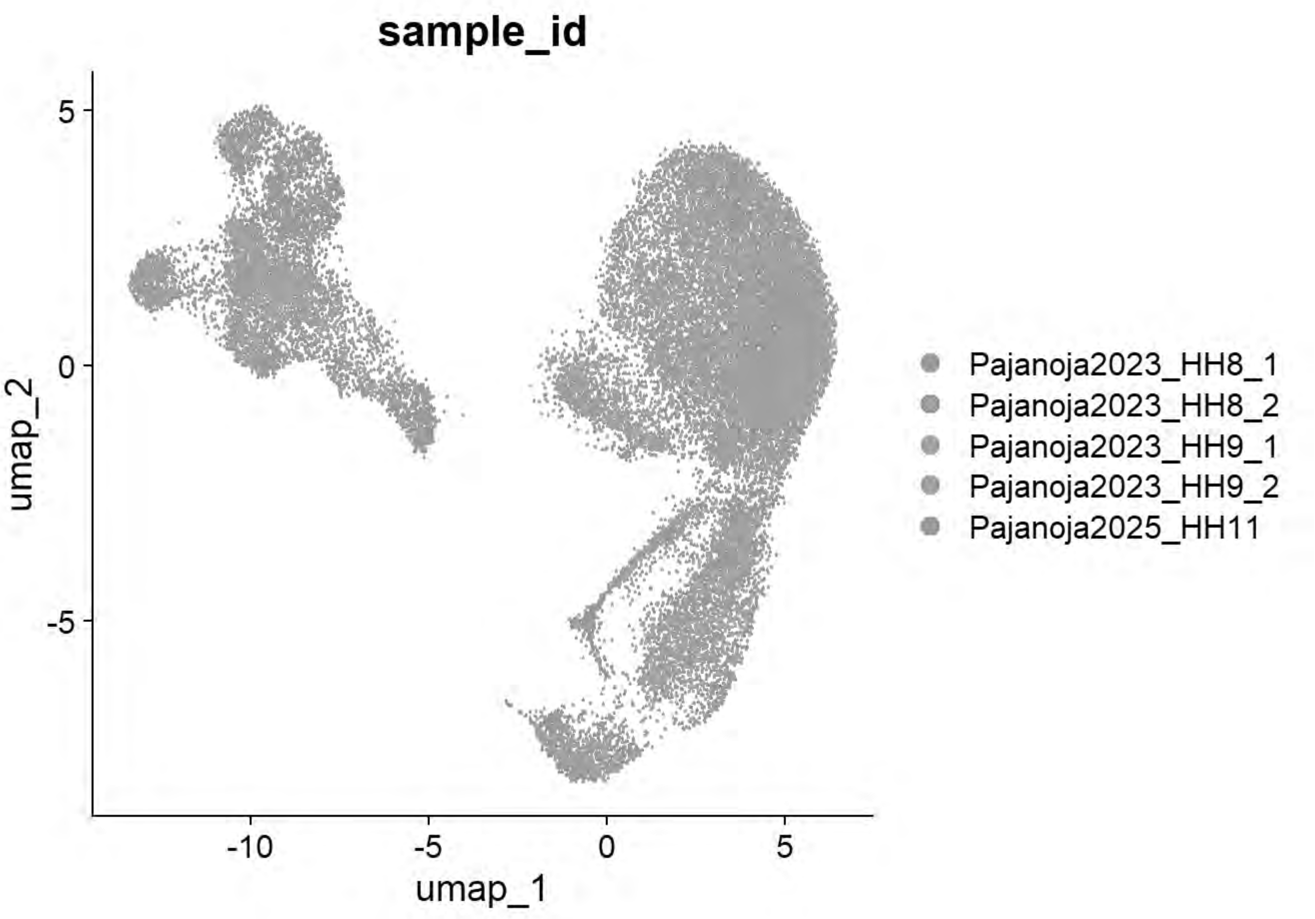


```
table(chick$seurat_clusters,chick$sample_id)
```

```
##
##      Pajanoja2023_HH8_1 Pajanoja2023_HH8_2 Pajanoja2023_HH9_1
##   0                 782               1463               1739
##   1                 515               1488               1433
##   2                 629                669                866
##   3                 974                717                839
##   4                 479                633                379
##   5                  94                247                491
##   6                   4                  1                  3
##   7                 550                 97                 21
##   8                 167                 97                120
##   9                  34                 26                216
##   10                115                125                177
##   11                195                 59                101
##   12                462                 47                 41
##   13                  0                  0                  2
##   14                  5                329                 11
##
##      Pajanoja2023_HH9_2 Pajanoja2025_HH11
##   0                2037               880
##   1                1430               882
##   2                 676               583
##   3                 279               250
```

```
##   4                  523                233
##   5                  567                190
##   6                    4               1376
##   7                  110                301
##   8                   82                546
##   9                  238                273
##   10                 228                 51
##   11                  70                269
##   12                  40                 72
##   13                   0                654
##   14                  29                 71
```

```
qc_tester(chick)
```

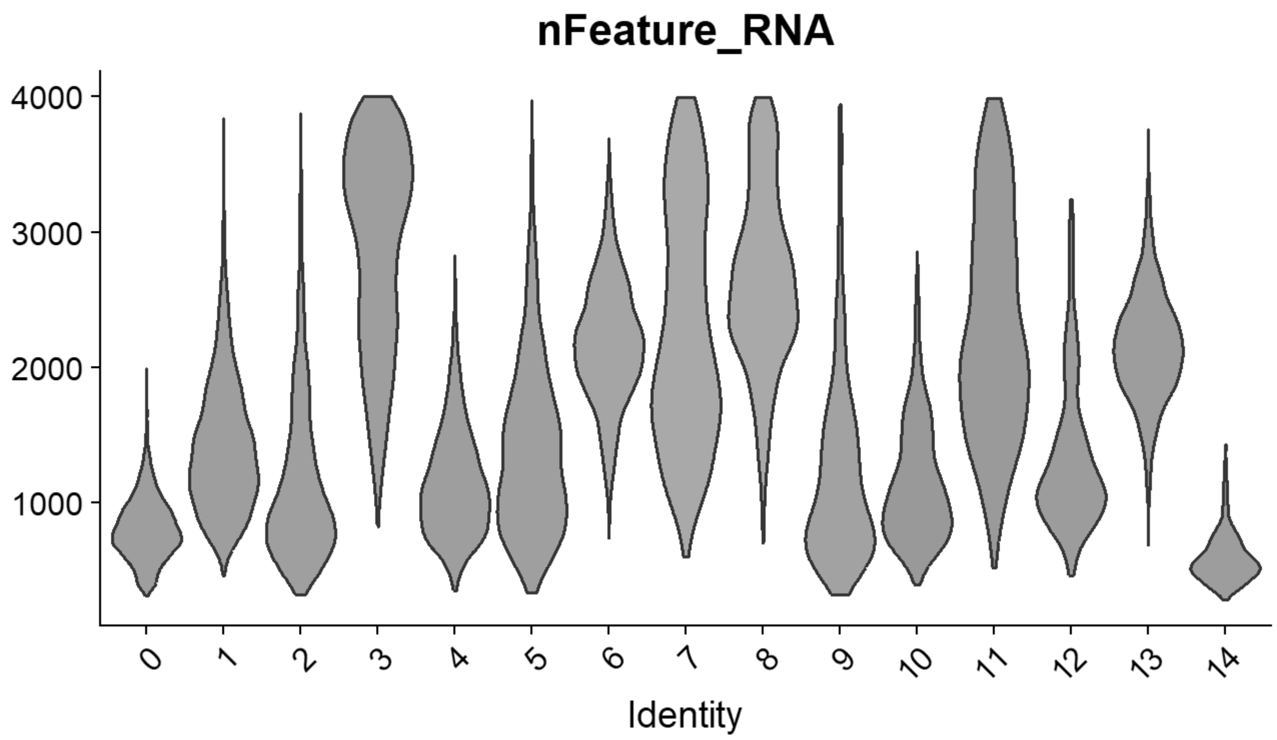


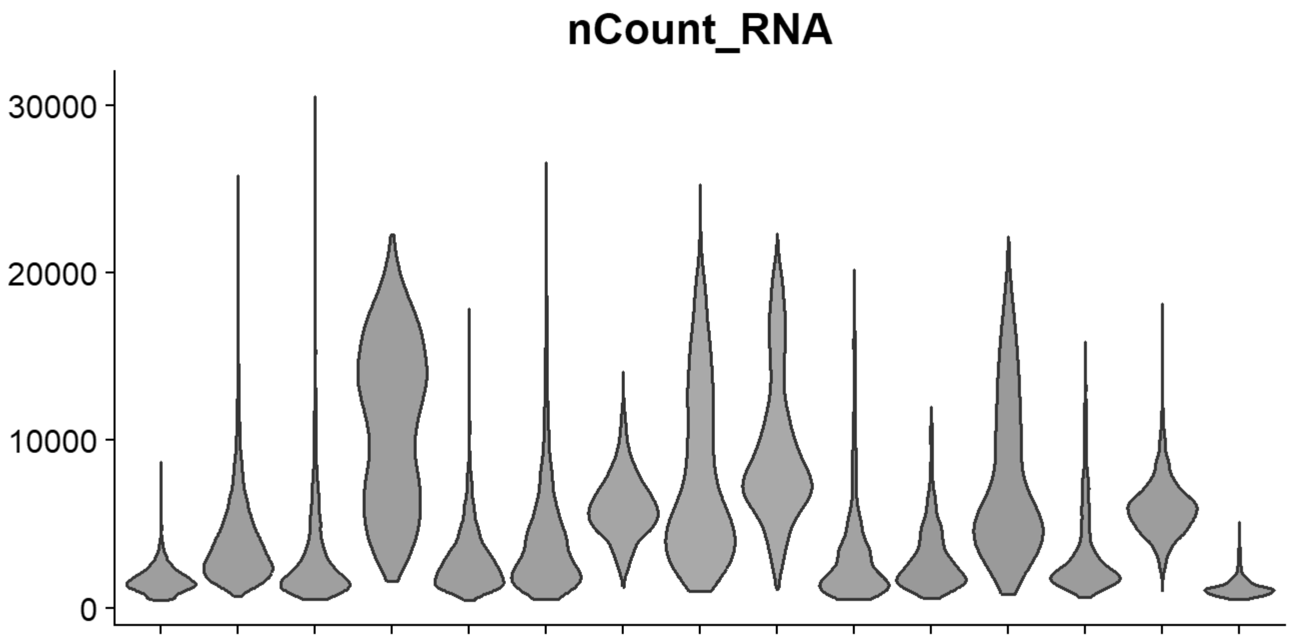

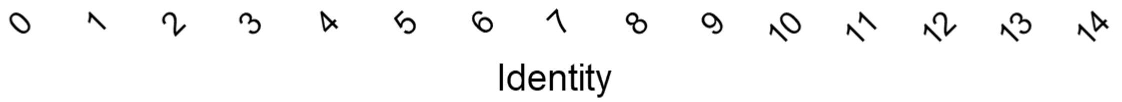


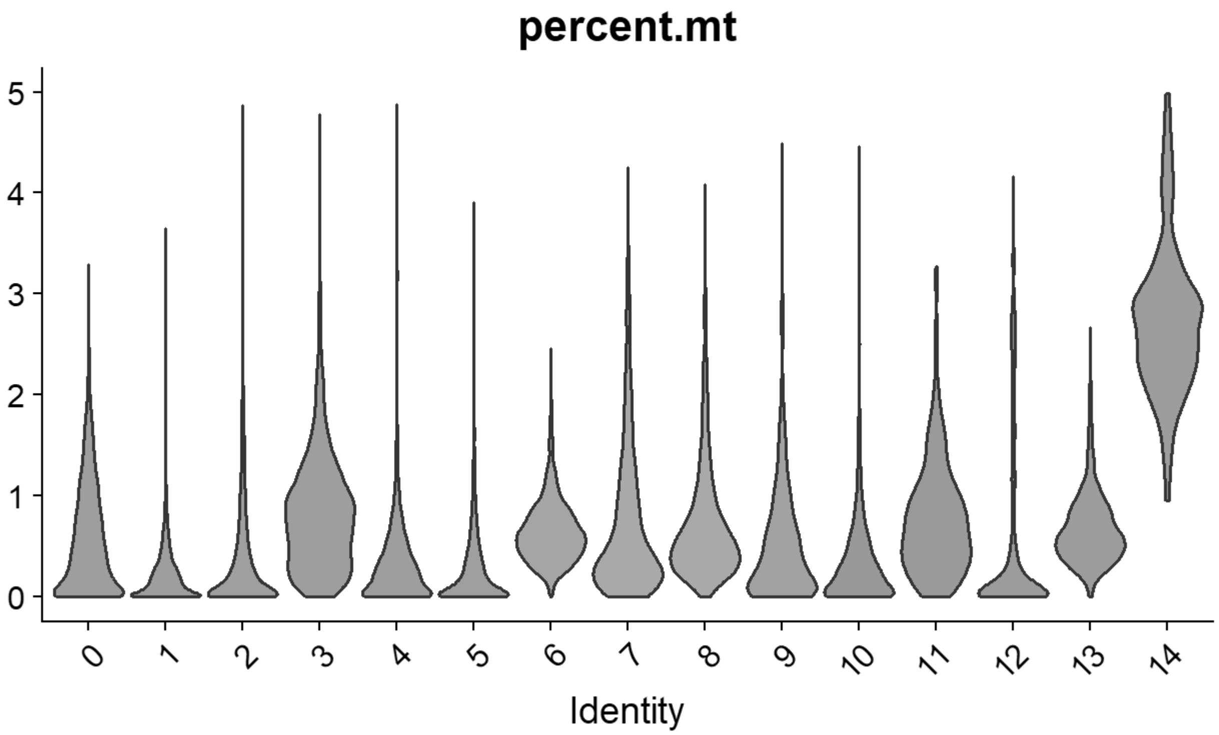


```
# Williams: Neural plate/tube
FeaturePlot(chick, features = c("CLDN1","FRZB","DNMT3A"))
```

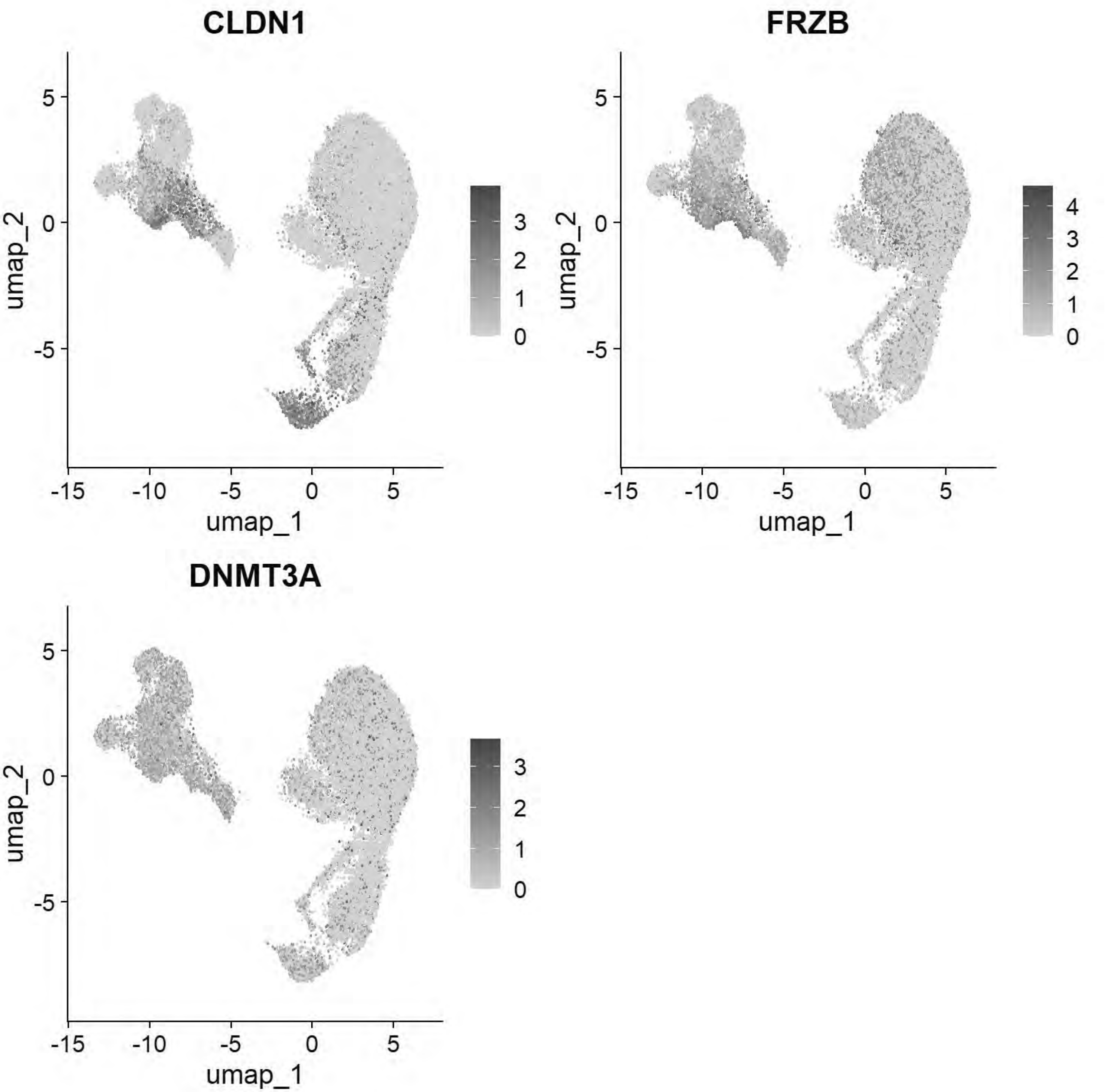


```
# Williams: Non-Neural Ectoderm
FeaturePlot(chick, features = c("DLX5","TFAP2A","ASTL","PAX6"))
```

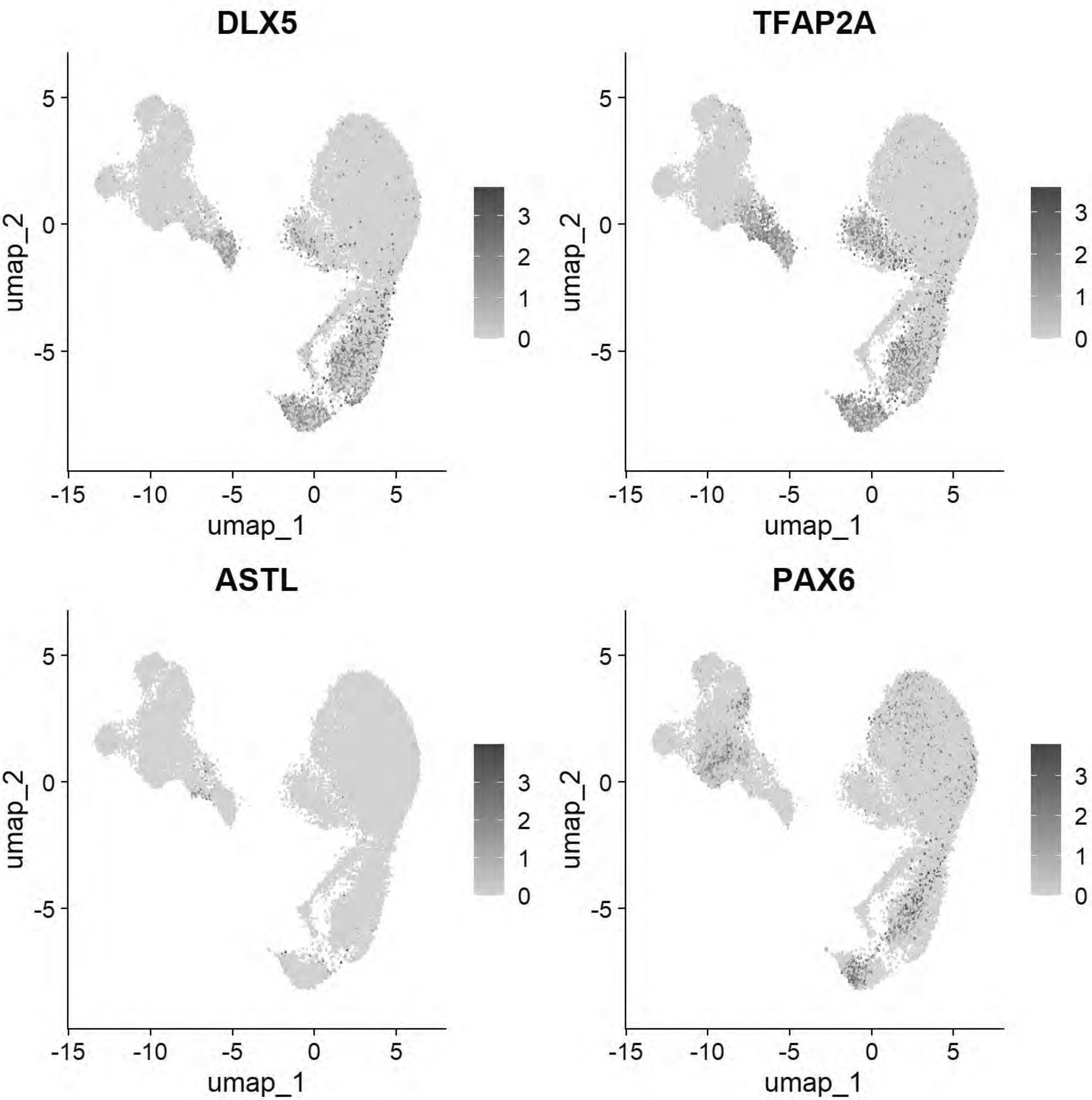


```
# Williams: Ectoderm (Neural Plate)
FeaturePlot(chick, features = c("SOX2","SOX3","SOX21","FRZB","SFRP2"))
```

```
## Warning: All cells have the same value (0) of "SOX3"
```

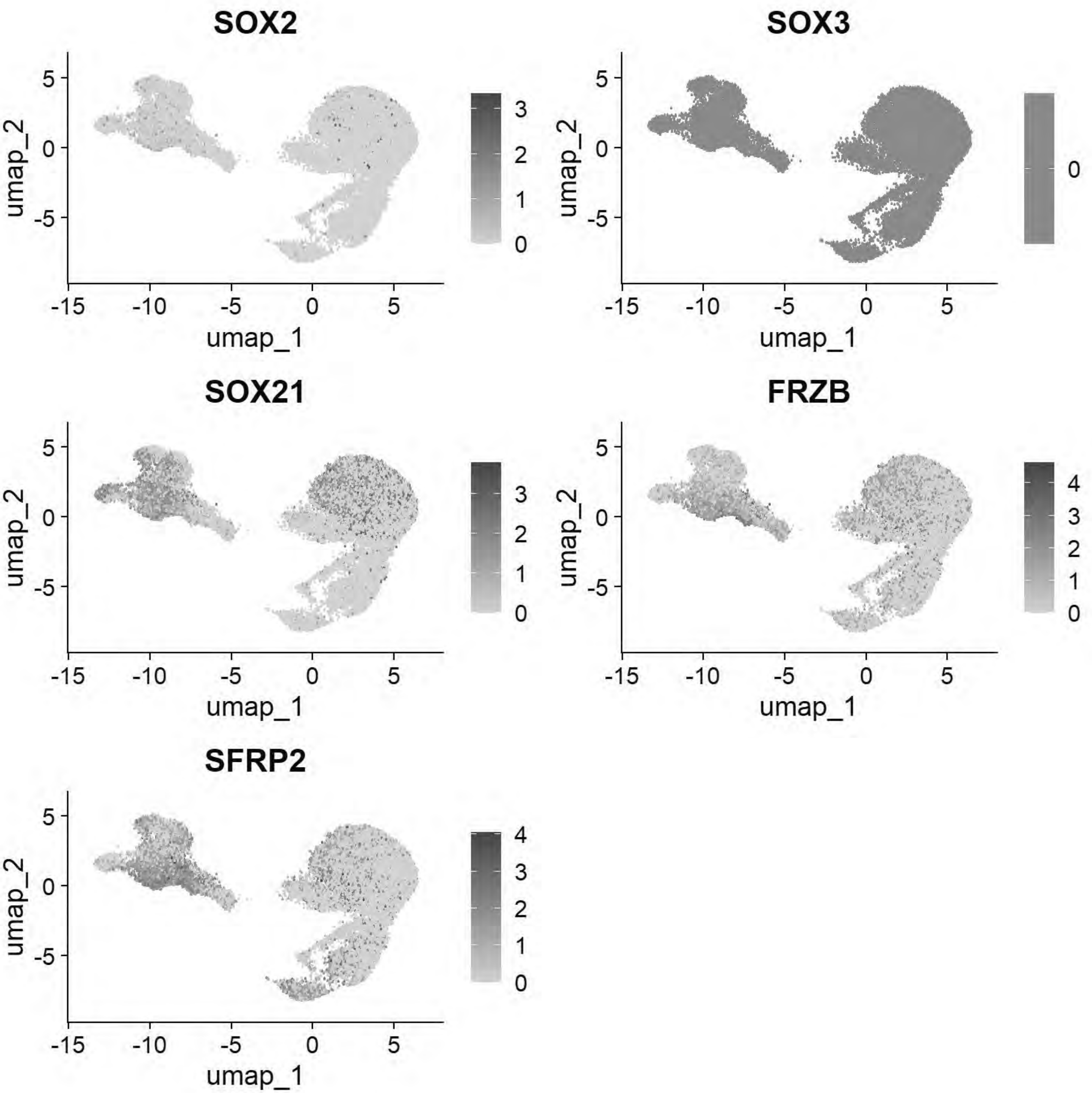


```
# Williams: Ectoderm (Neural Plate Border)
FeaturePlot(chick, features = c("PAX7","TFAP2A","DLX5","BMP4","MSX1","DRAXIN","TFAP2B"))
```

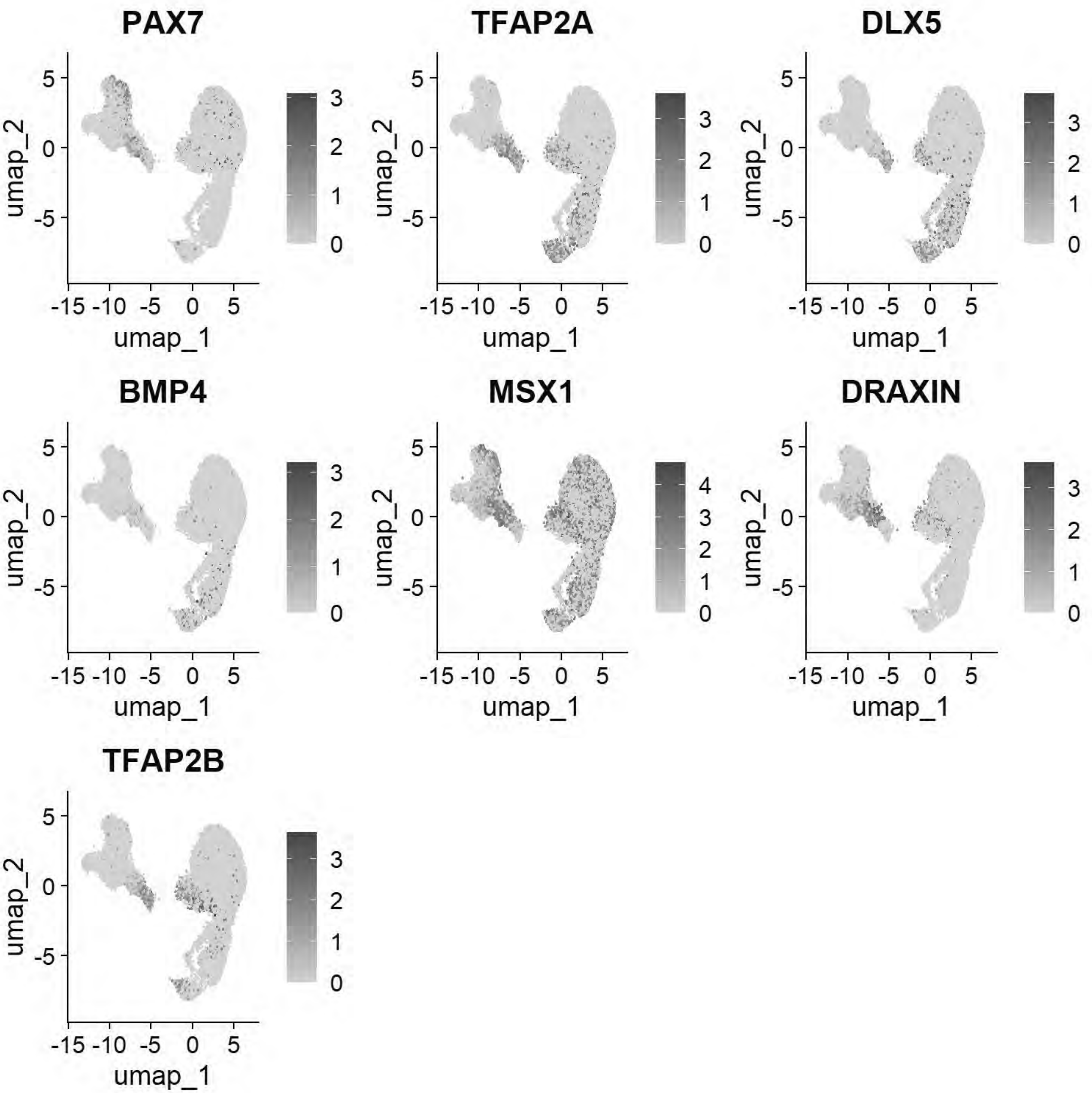


```
# Williams:Epiblast stem cell
FeaturePlot(chick, features = c("ID3","SALL4","TGIF1","ELAVL1"))
```

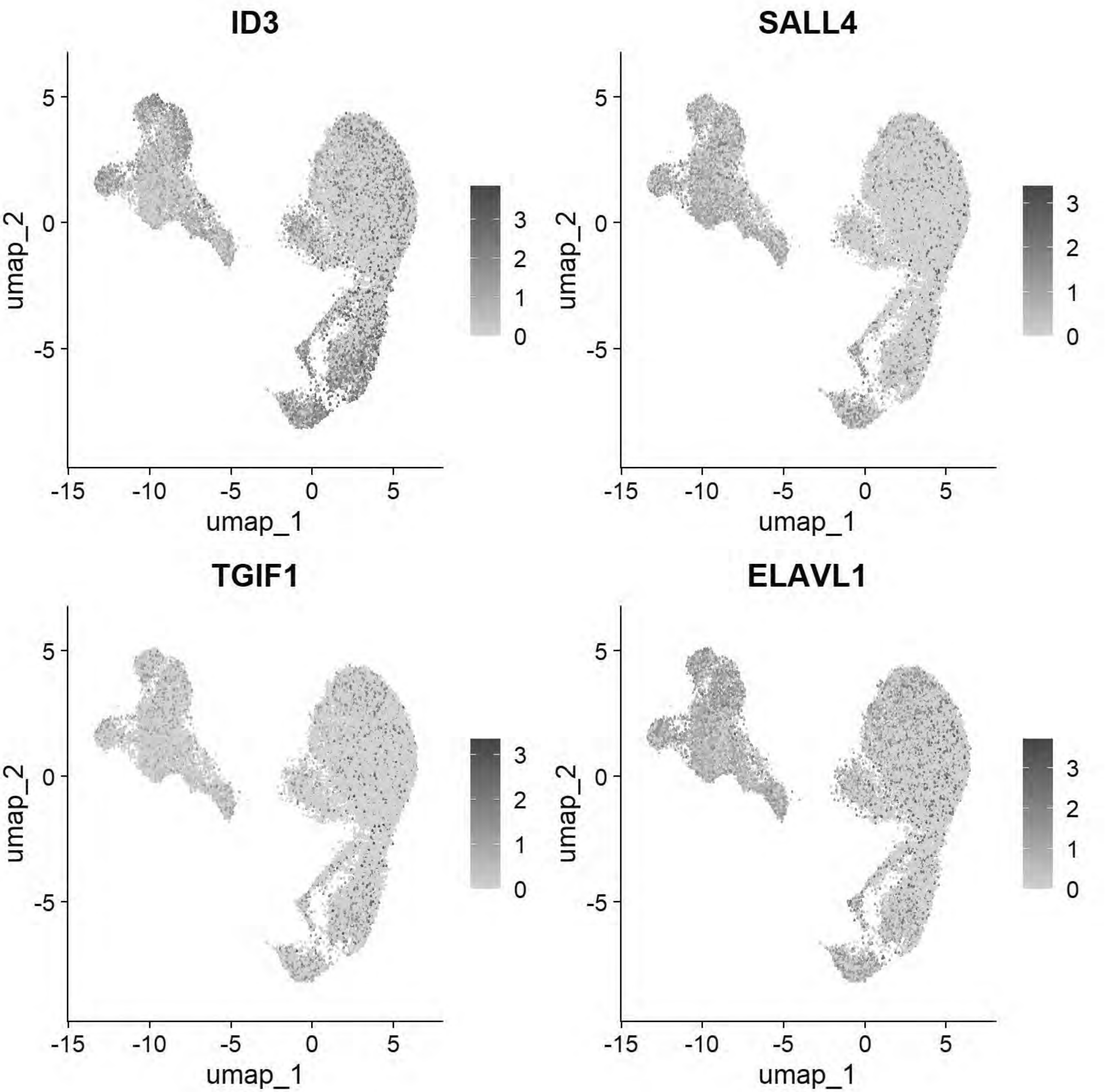


```
# Williams: Posterior Lateral Plate Mesoderm
FeaturePlot(chick, features = c("GATA2","HOXB5","CDX4"))
```

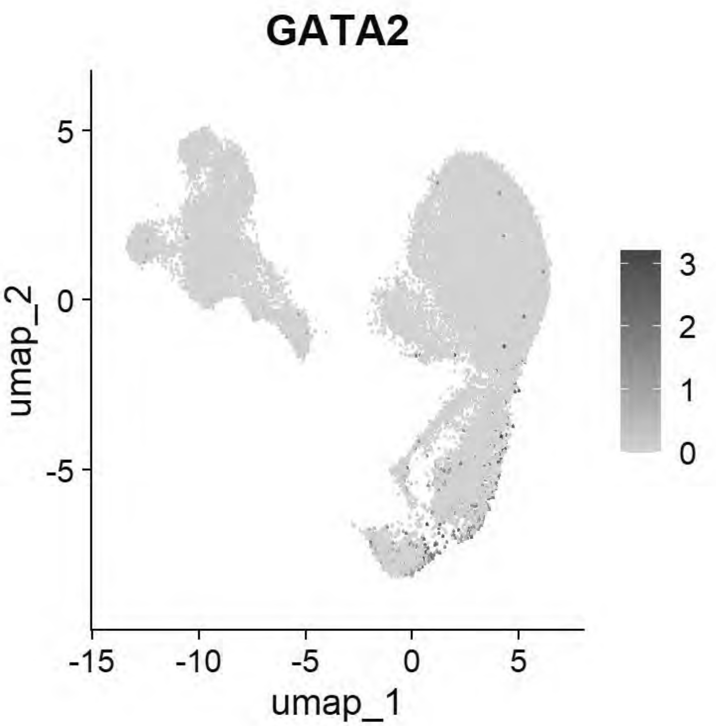


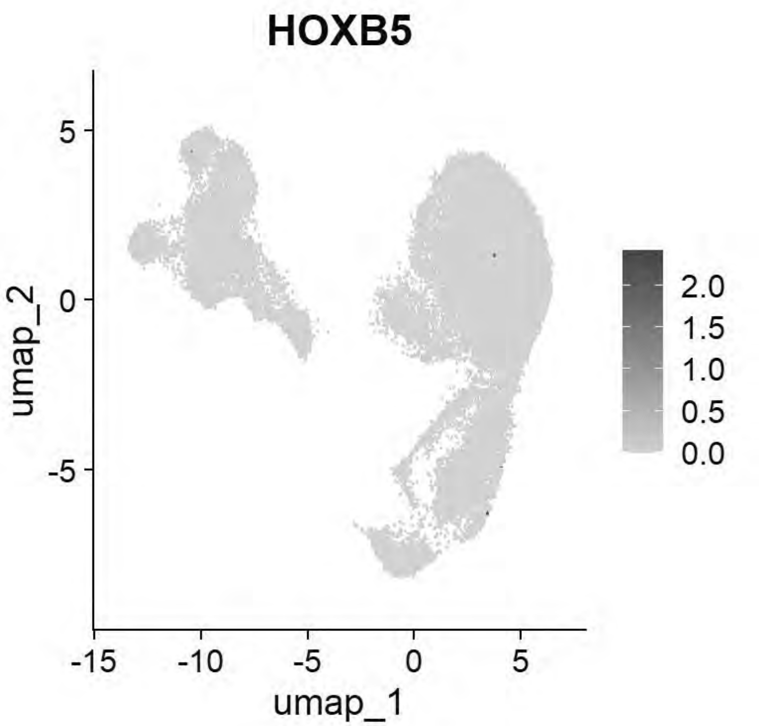


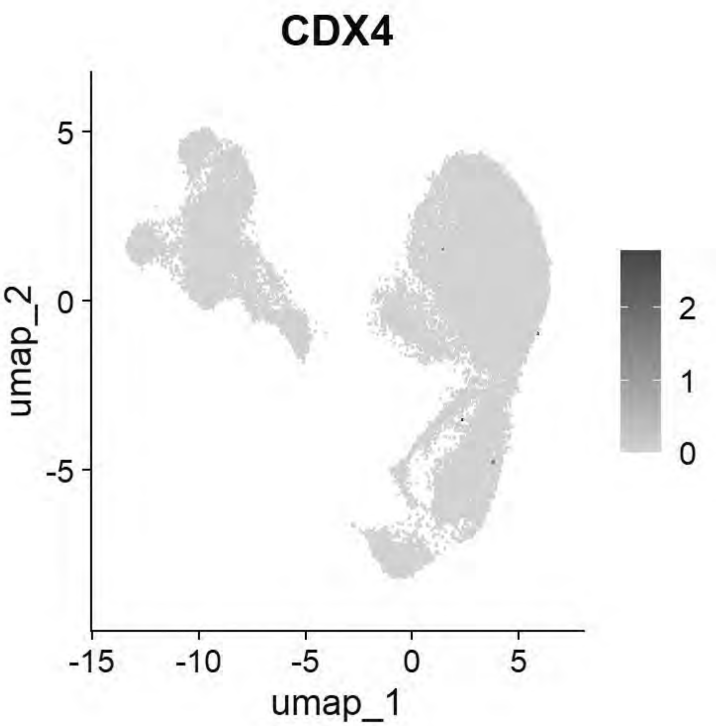


```
# Williams: Paraxial Mesoderm
FeaturePlot(chick, features = c("MSGN1","MESP1","MEOX1"))
```

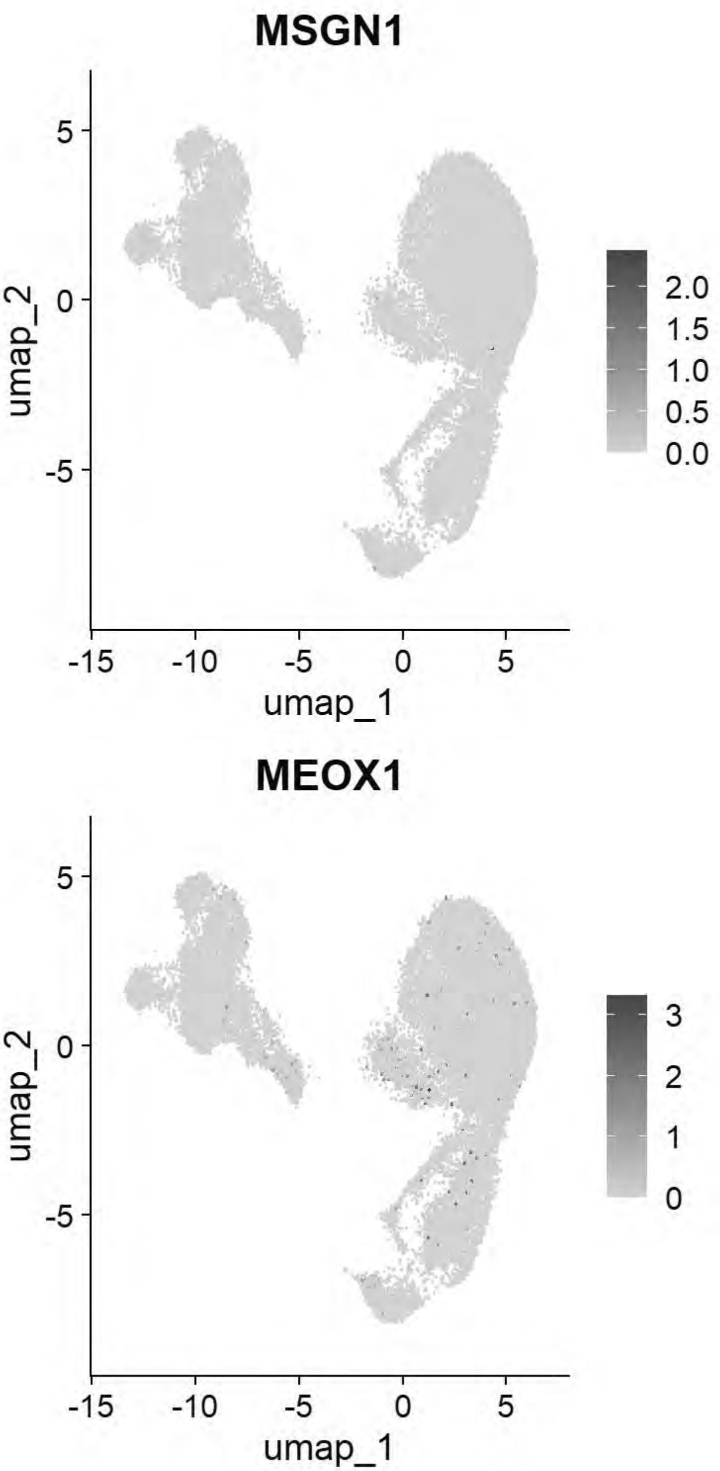


```r
# Williams: Lateral Plate Mesoderm
FeaturePlot(chick, features = c("PITX2","ALX1","OLFML3","SIX1","TWIST1"))
```

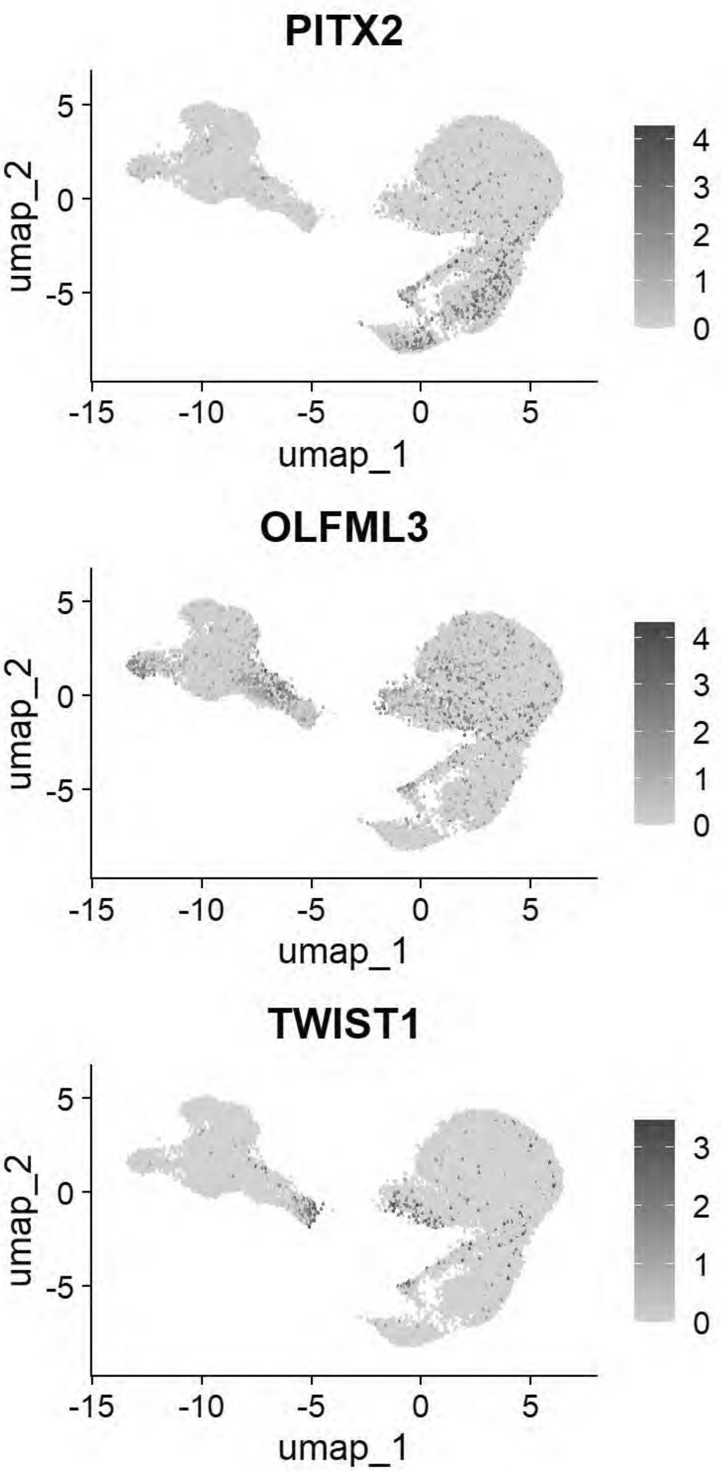


```
# Williams: Cardiac Mesoderm
FeaturePlot(chick, features = c("TCF21","GATA5","LMO2","ETS1","KDR"))
```

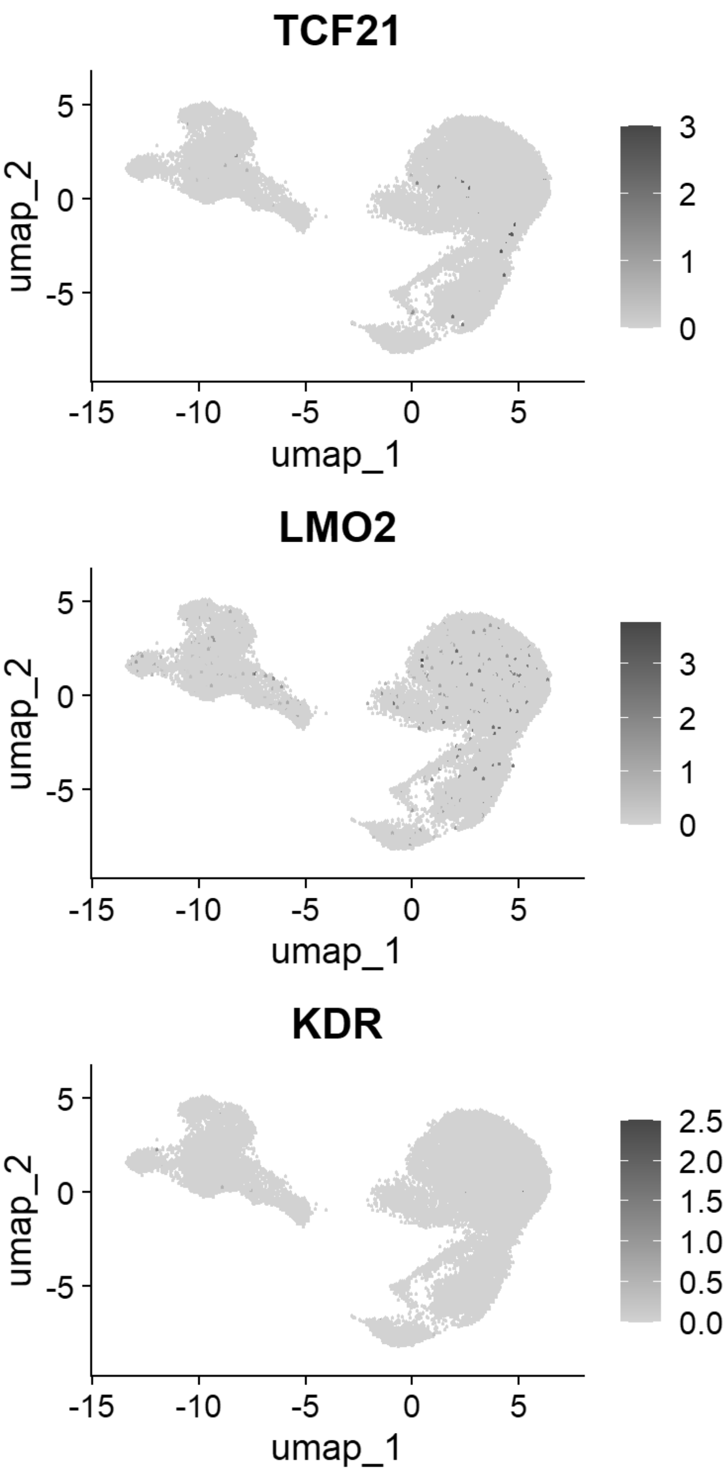


```
# Williams: Head Mesenchyme
FeaturePlot(chick, features = c("TCF21","GATA5","LMO2","ETS1","KDR"))
```

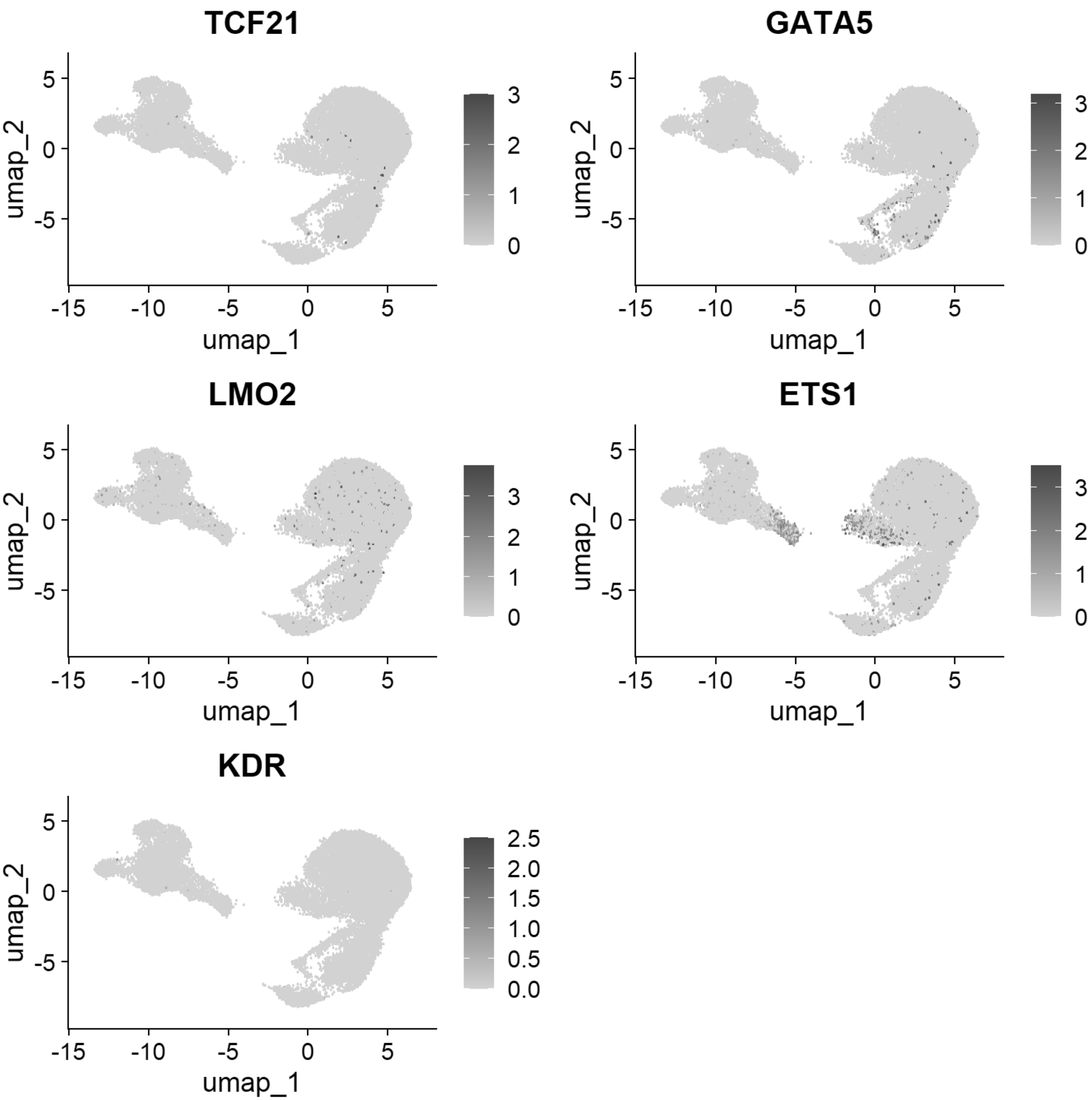


```
# Williams: Hensen's Node & Primitive Streak
FeaturePlot(chick, features = c("DLL1","FGF8","NOTO","CHRD"))
```

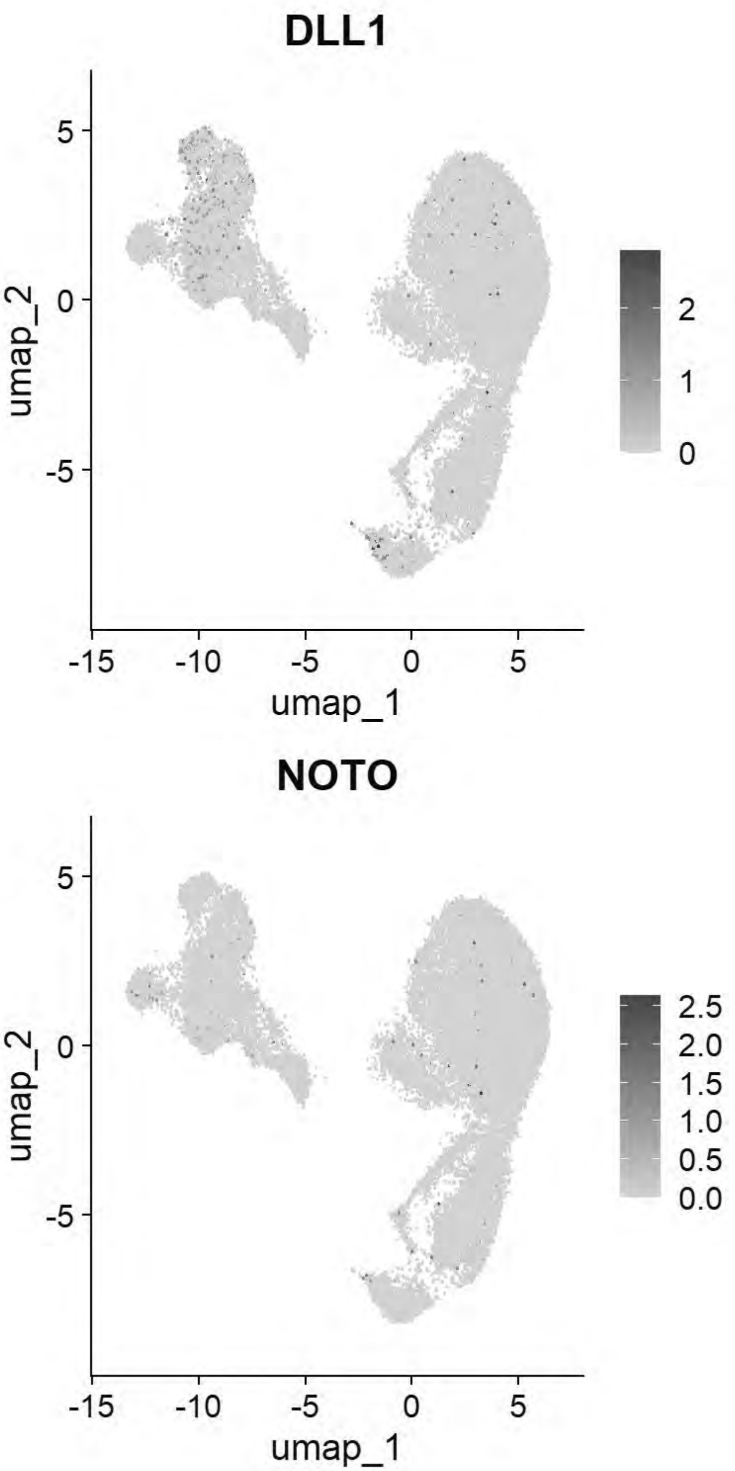


```
# Williams: Endoderm
FeaturePlot(chick, features = c("SOX17","FOXA2","CXCR4"))
```

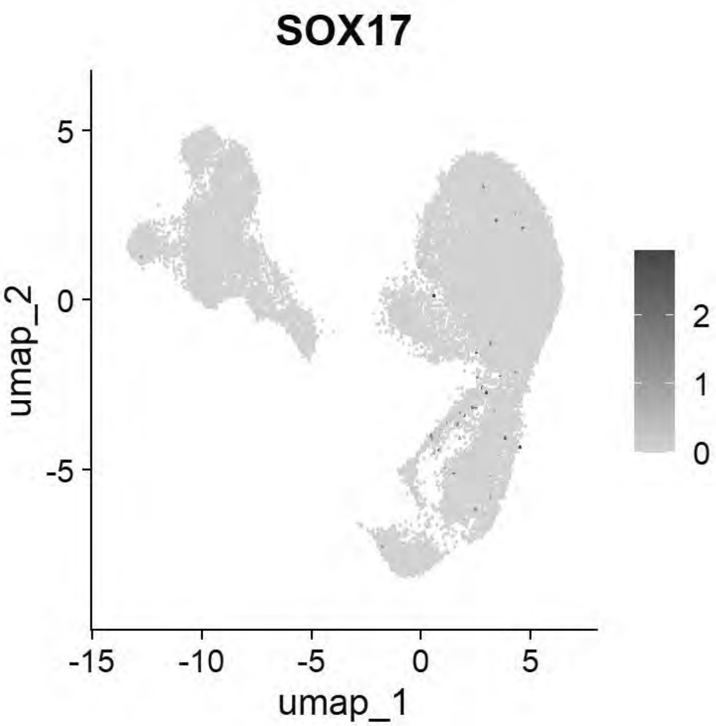


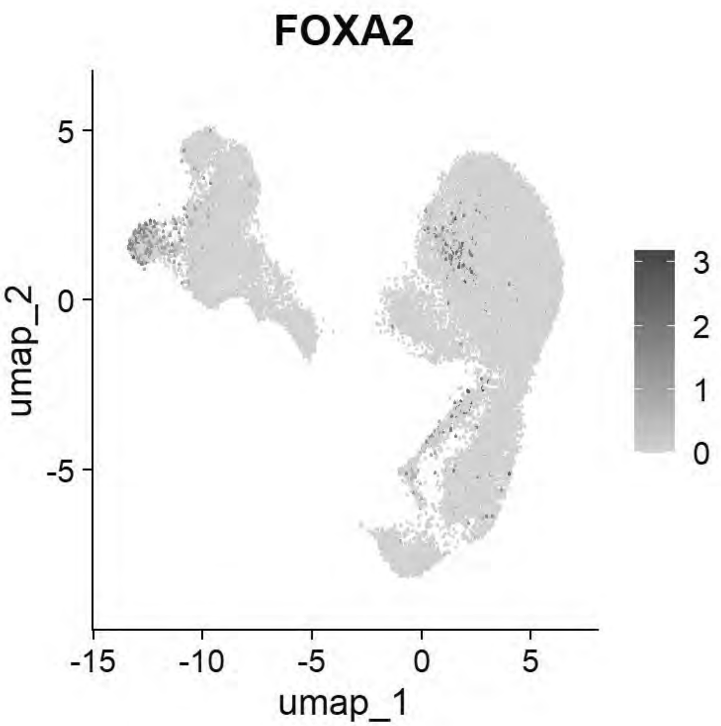


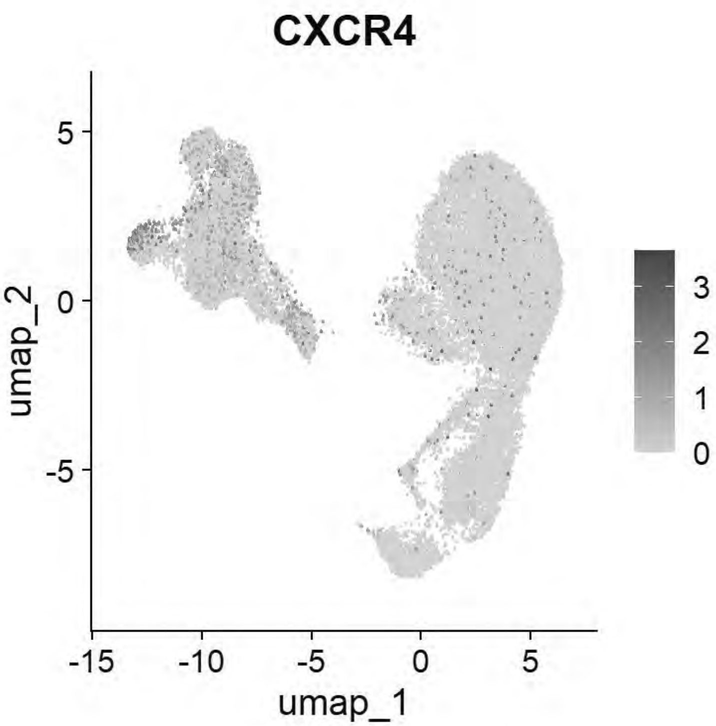


```
# Pajanoja: Ectoderm
FeaturePlot(chick, features = c("TFAP2A","DLX5","CLDN1","SOX2","NESTIN","MYCN"))
```

```
## Warning: The following requested variables were not found: NESTIN
```

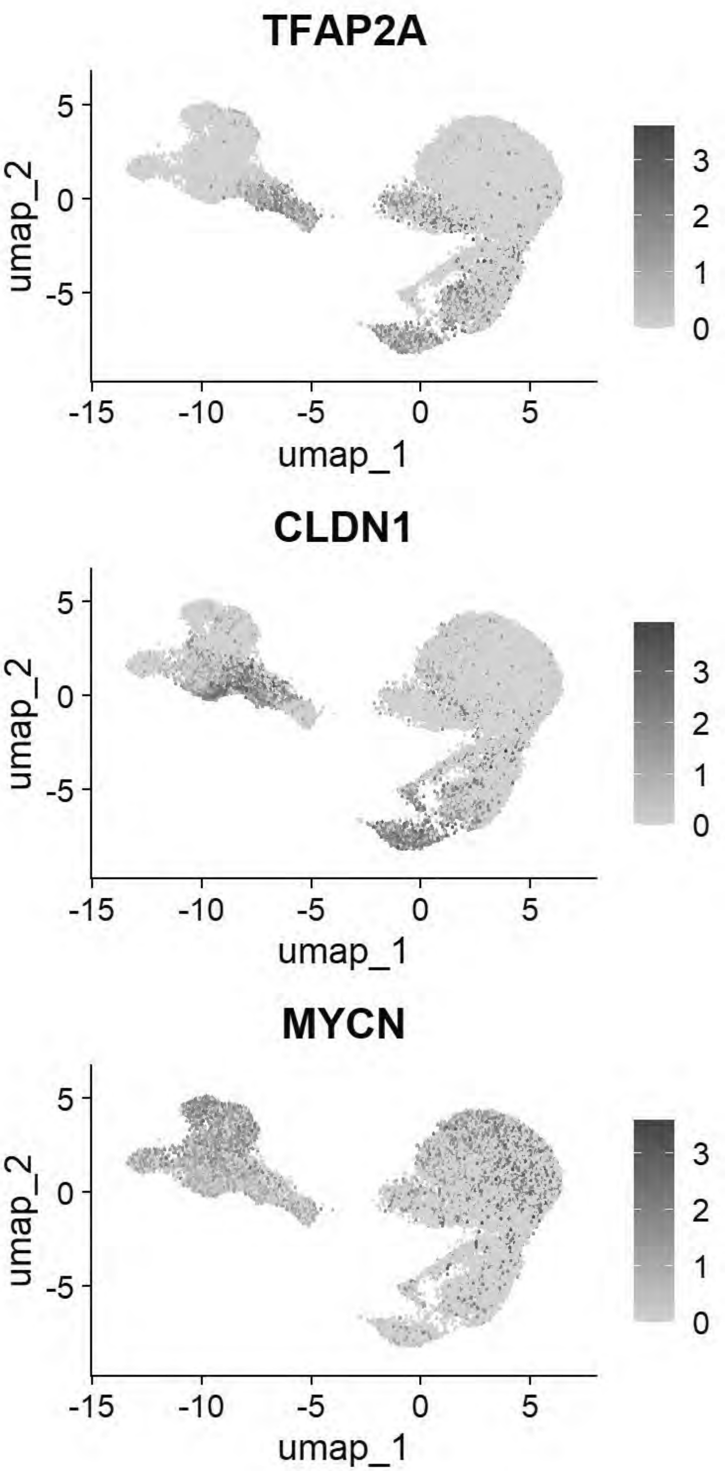


```
# Pajanoja: Endoderm
FeaturePlot(chick, features = c("SOX17","KRT17","HHEX"))
```

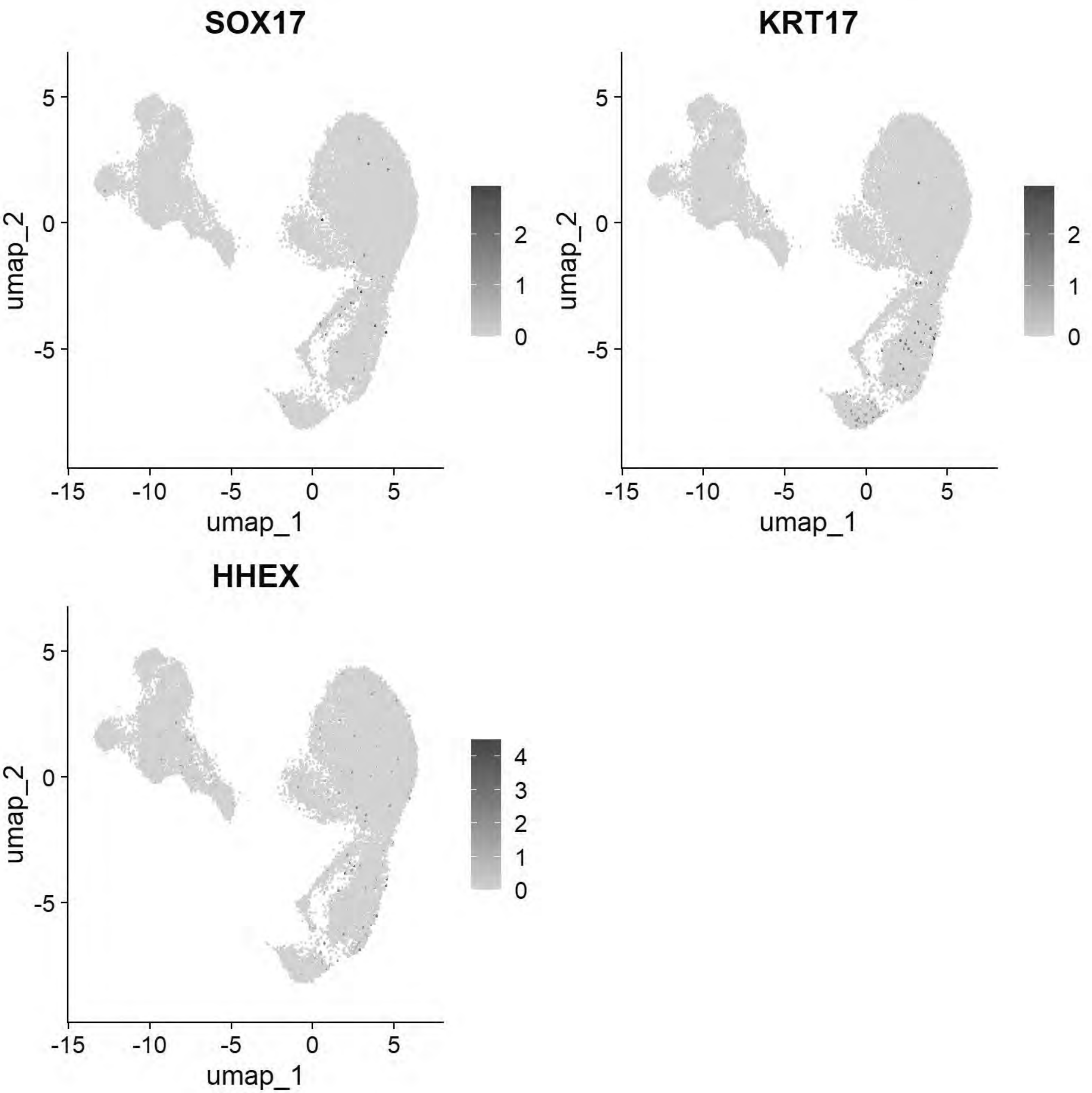


```
# Pajanoja: Mesoderm
FeaturePlot(chick, features = c("ALX1","TWIST1","HAND2","PITX2"))
```

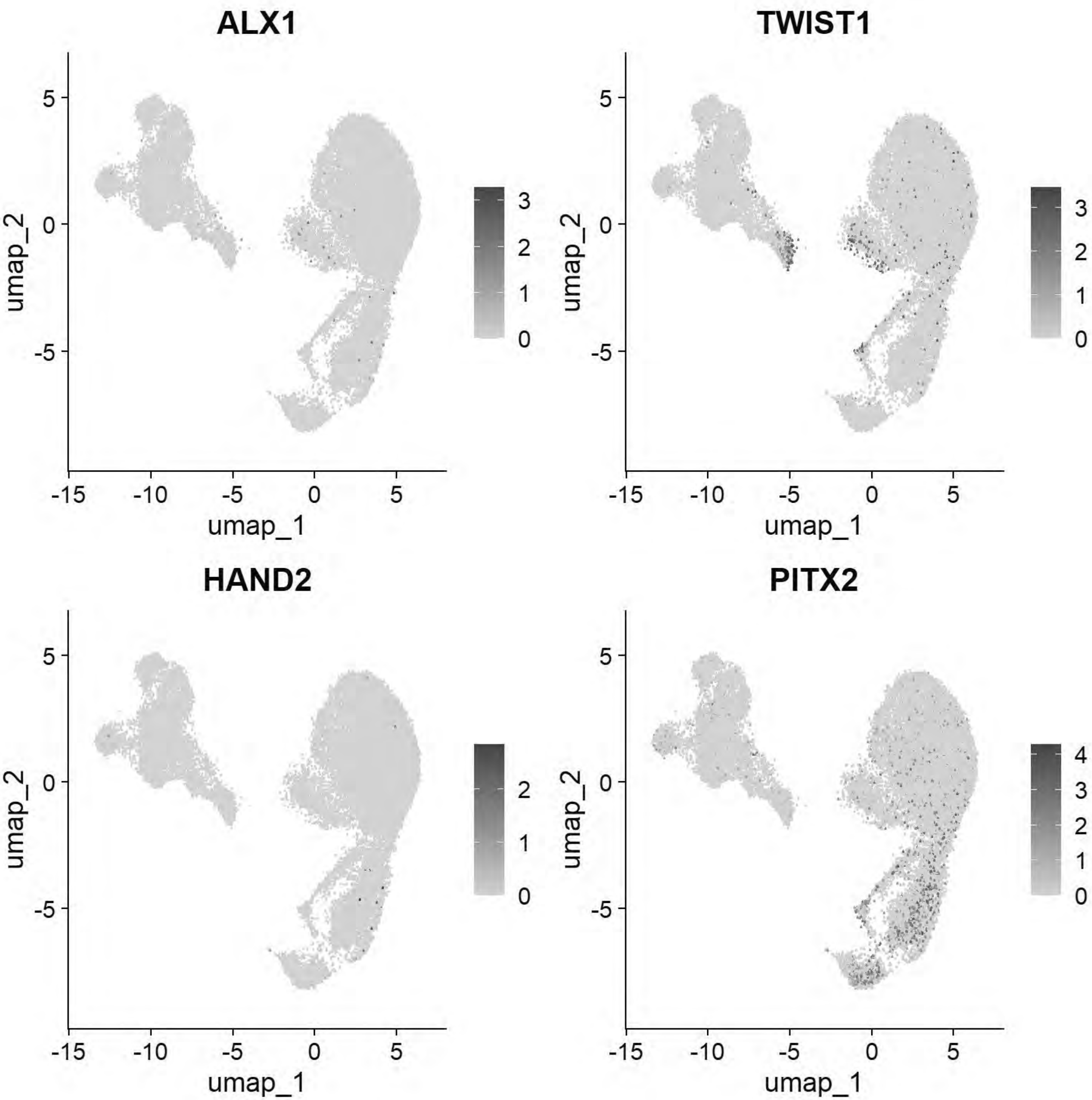


```
# Pajanoja: Notochord
FeaturePlot(chick, features = c("CHRD","TBXT","NOTO"))
```

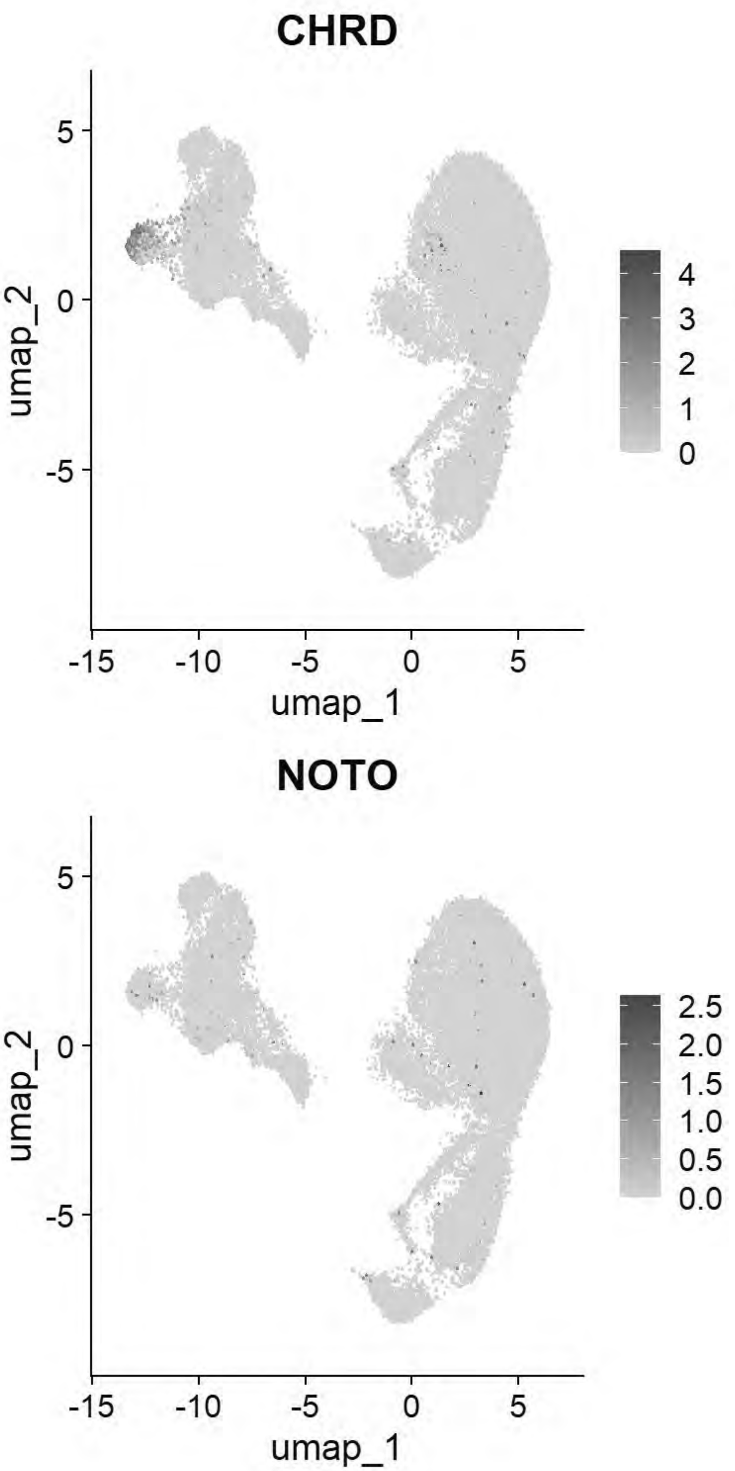


```
# Pajanoja: Ventral Neural Tube
FeaturePlot(chick, features=c("SHH","FOXA1","FOXA2"))
```

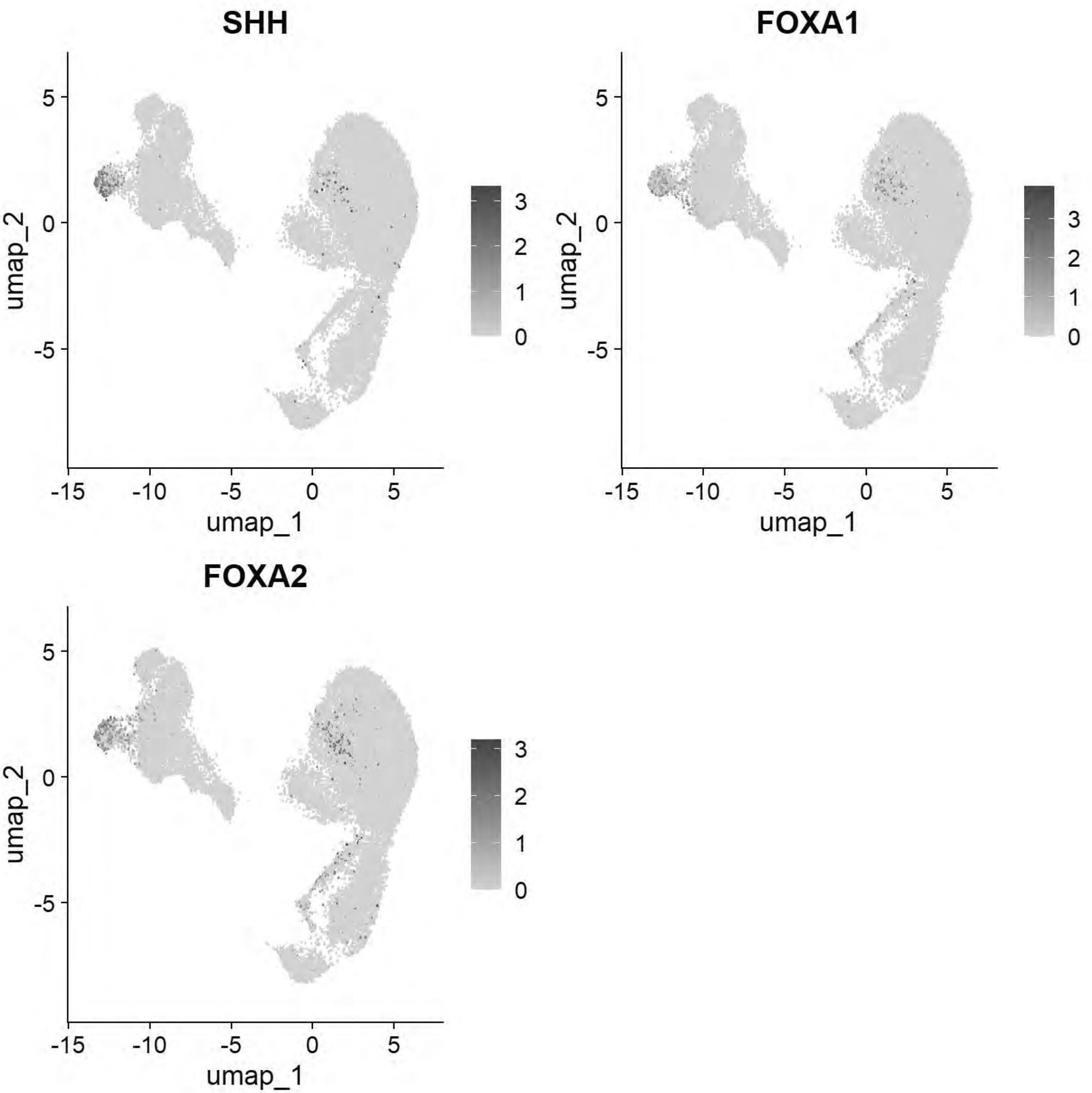


```
# Pajanoja: Neural Tube
FeaturePlot(chick, features=c("CDH2","HES5","MYCN","PAX2","NESTIN"))
```

```
## Warning: The following requested variables were not found: NESTIN
```

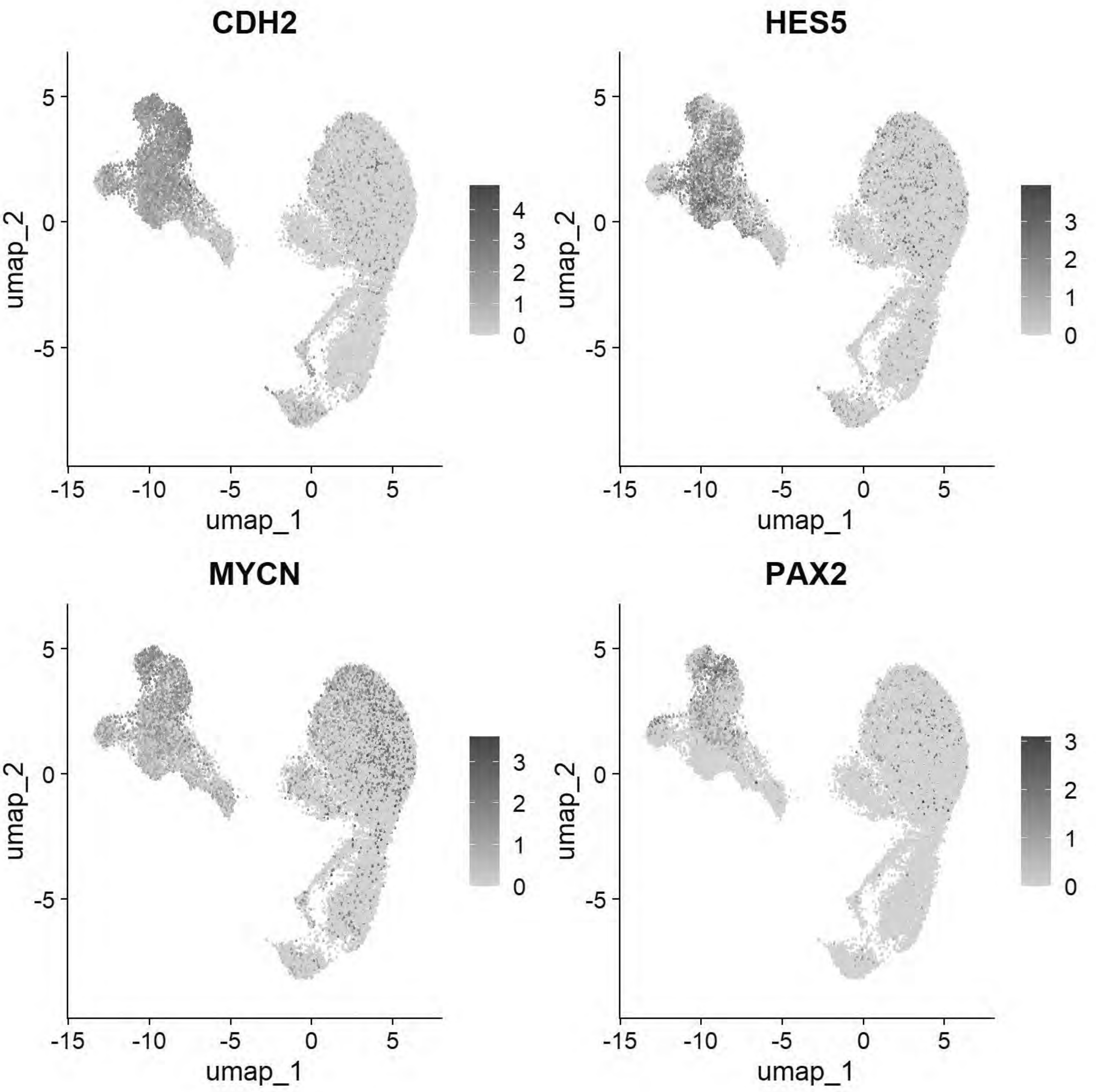


```r
# Pajanoja: Non-Neural Ectoderm/Placode
FeaturePlot(chick, features=c("KRT24","KRT18","DLX5","SIX1","EYA2","TFAP2A"))
```

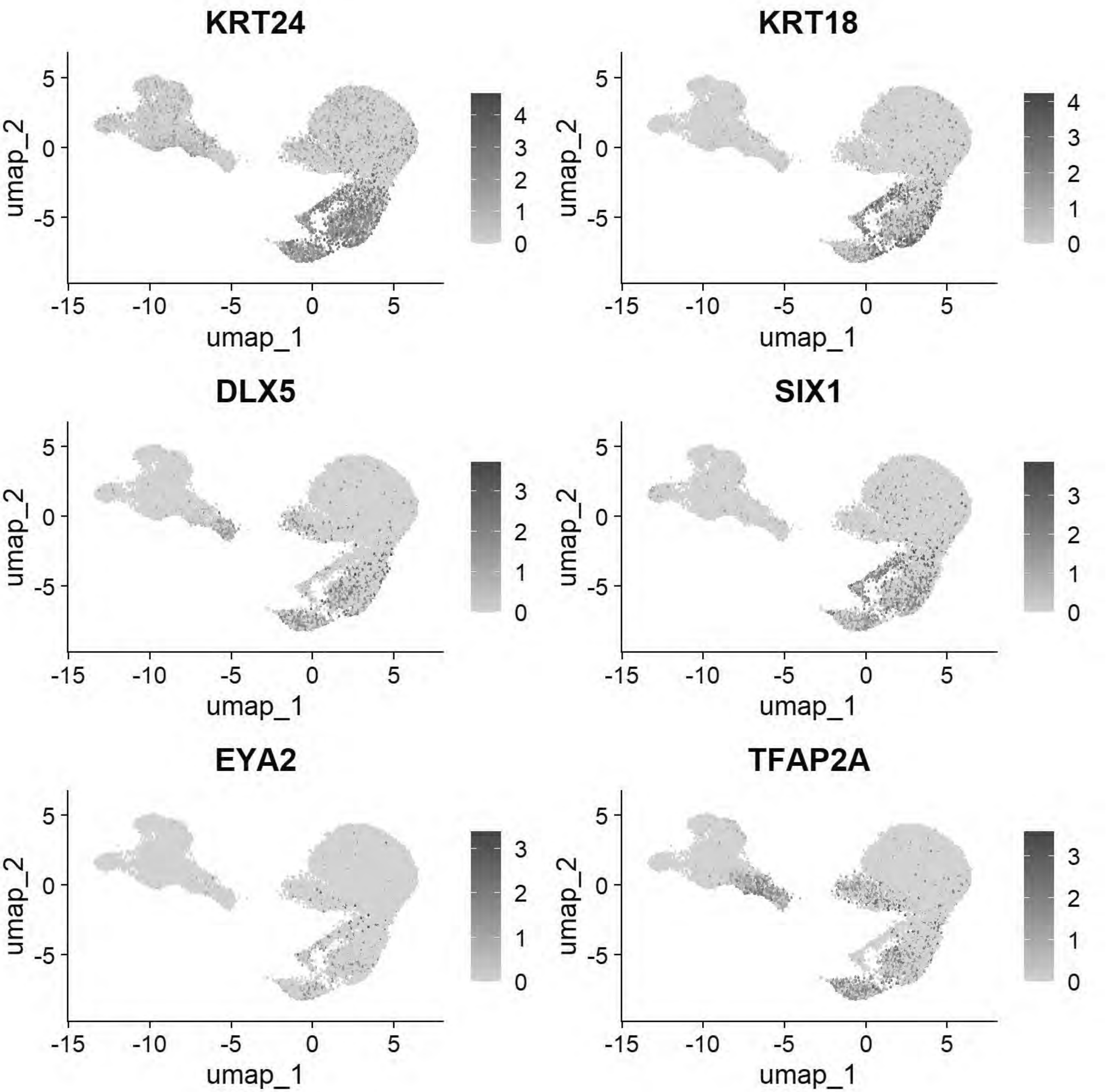


```r
# Pajanoja: Neural crest
FeaturePlot(chick, features=c("FOXD3","TFAP2B","SOX9","ETS1","SNAI2"))
```

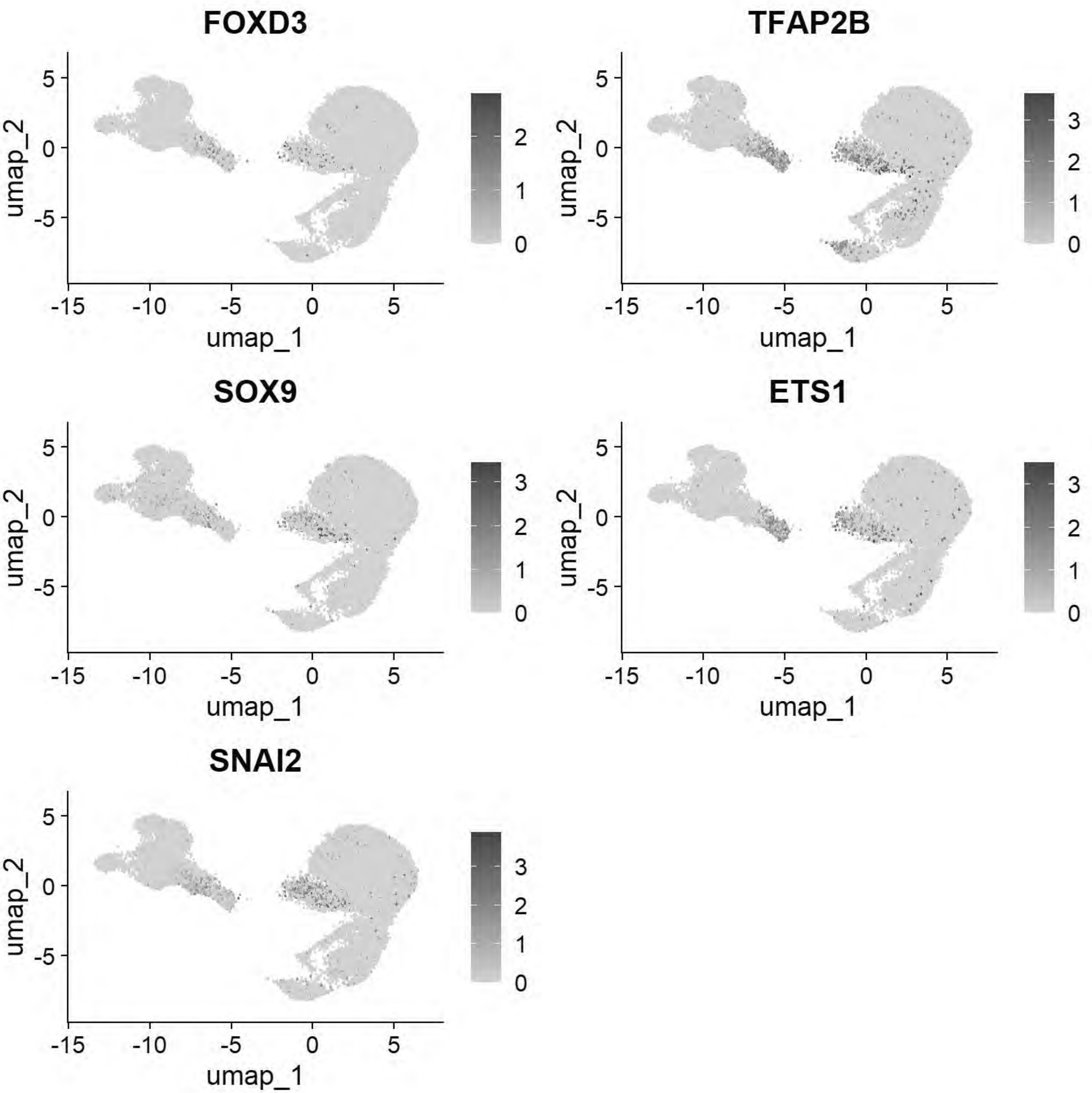


```
gc()
```

```
##             used   (Mb) gc trigger    (Mb)   max used    (Mb)
## Ncells   5072817  271.0   13651909   729.1   21331105  1139.3
## Vcells 994183038 7585.1 2342481578 17871.8 4574774951 34902.8
```

```
chick <- JoinLayers(chick)
chick.markers <- FindAllMarkers(chick, min.pct = 0.5, logfc.threshold = 0.5)
```

```
## Calculating cluster 0
```

```
## Calculating cluster 1
```

```
## Calculating cluster 2
```

```
## Calculating cluster 3
```

```
## Calculating cluster 4
```

```
## Calculating cluster 5
```

```
## Calculating cluster 6
```

```
## Calculating cluster 7
```

```
## Calculating cluster 8
```

```
## Calculating cluster 9
```

```
## Calculating cluster 10
```

```
## Calculating cluster 11
```

```
## Calculating cluster 12
```

```
## Calculating cluster 13
```

```
## Calculating cluster 14
```

```
gc()
```

```
##              used   (Mb) gc trigger    (Mb)   max used    (Mb)
## Ncells    5072184  270.9   13651909   729.1   21331105  1139.3
## Vcells 1191398065 9089.7 2342481578 17871.8 4574774951 34902.8
```

```
chick.markers %>%
  group_by(cluster) %>%
  top_n(n = 5, wt = avg_log2FC) -> top5
DotPlot(chick, features = unique(top5$gene))+coord_flip()
```

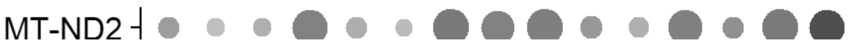

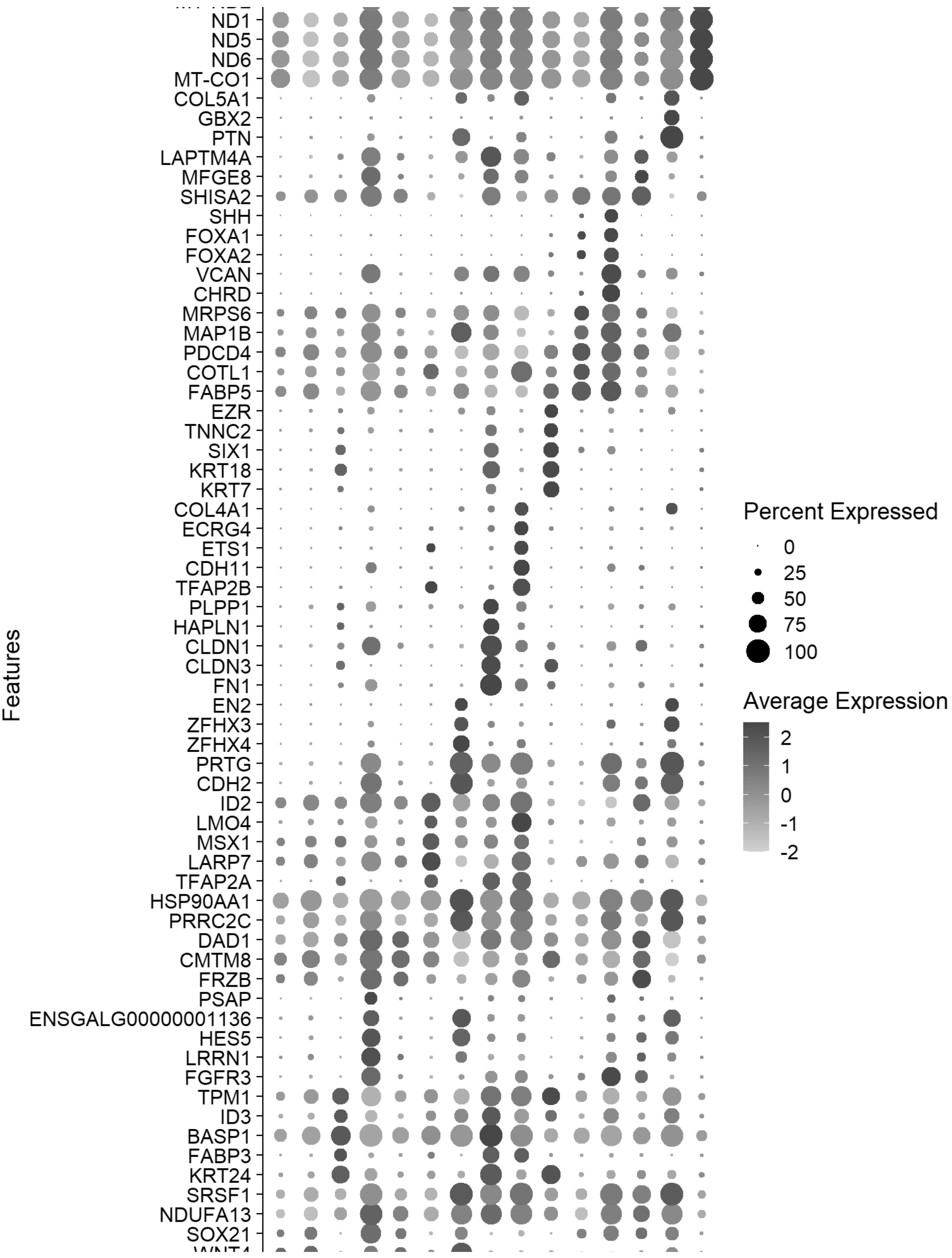
Features
ND1
ND5
ND6
MT-CO1
COL5A1
GBX2
PTN
LAPTM4A
MFGE8
SHISA2
SHH
FOXA1
FOXA2
VCAN
CHRD
MRPS6
MAP1B
PDCD4
COTL1
FABP5
EZR
TNNC2
SIX1
KRT18
KRT7
COL4A1
ECRG4
ETS1
CDH11
TFAP2B
PLPP1
HAPLN1
CLDN1
CLDN3
FN1
EN2
ZFHX3
ZFHX4
PRTG
CDH2
ID2
LMO4
MSX1
LARP7
TFAP2A
HSP90AA1
PRRC2C
DAD1
CMTM8
FRZB
PSAP
ENSGALG00000001136
HES5
LRRN1
FGFR3
TPM1
ID3
BASP1
FABP3
KRT24
SRSF1
NDUFA13
SOX21
Percent Expressed
0
25
50
75
100
Average Expression
2
1
0
-1
-2

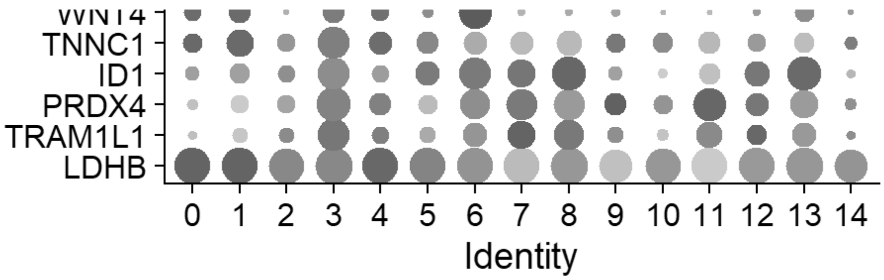


```
gc()
```

```
##               used   (Mb) gc trigger    (Mb)   max used    (Mb)
## Ncells    4983750  266.2   13651909   729.1   21331105  1139.3
## Vcells 1193299665 9104.2 2342481578 17871.8 4574774951 34902.8
```

```
FeaturePlot(chick, features = c("PAX7","SNAI2","SOX9","SOX10"))
```

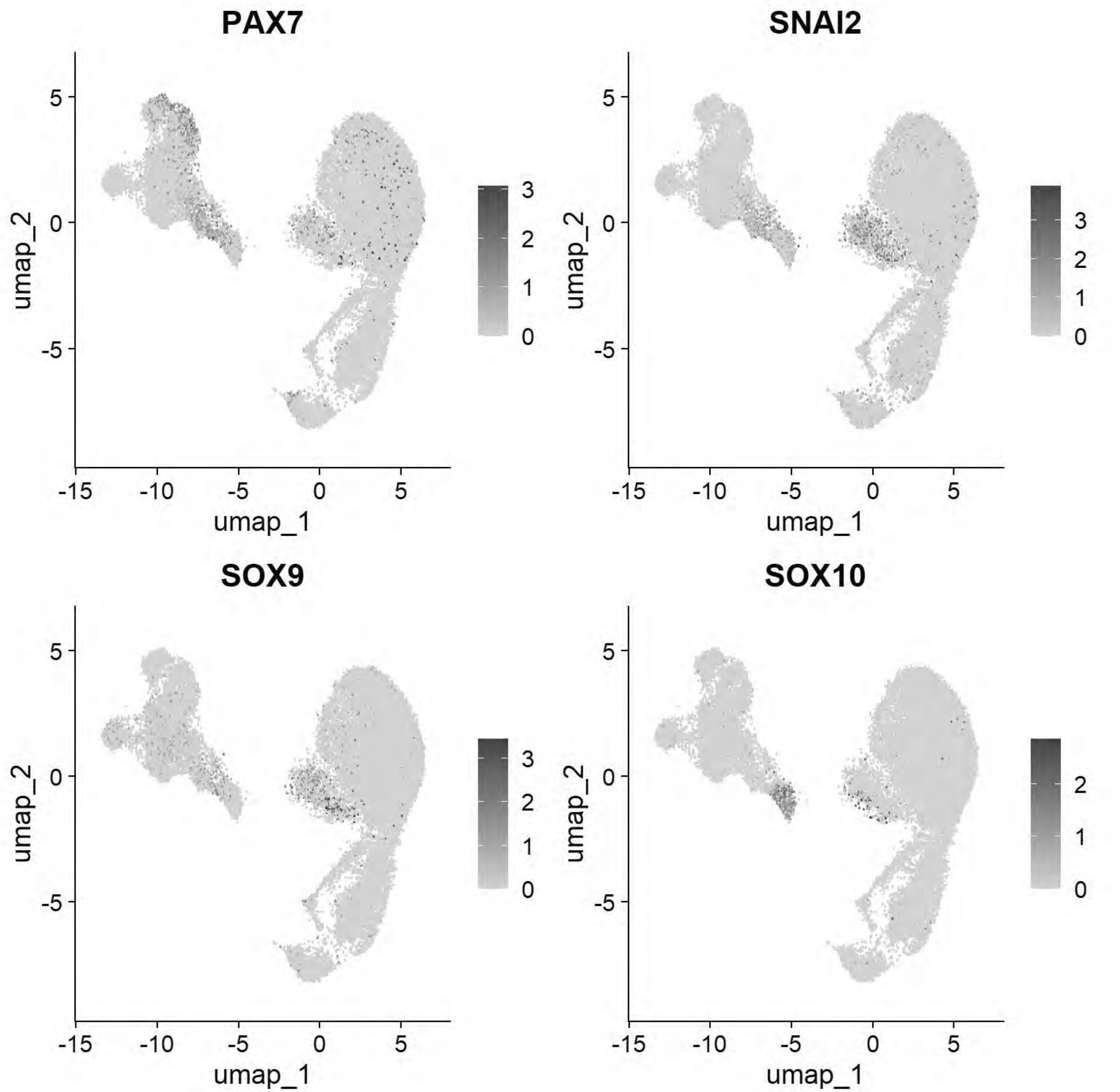


```
FeaturePlot(chick, features = c("SHH","DRAXIN","CDH6","SOX2"))
```

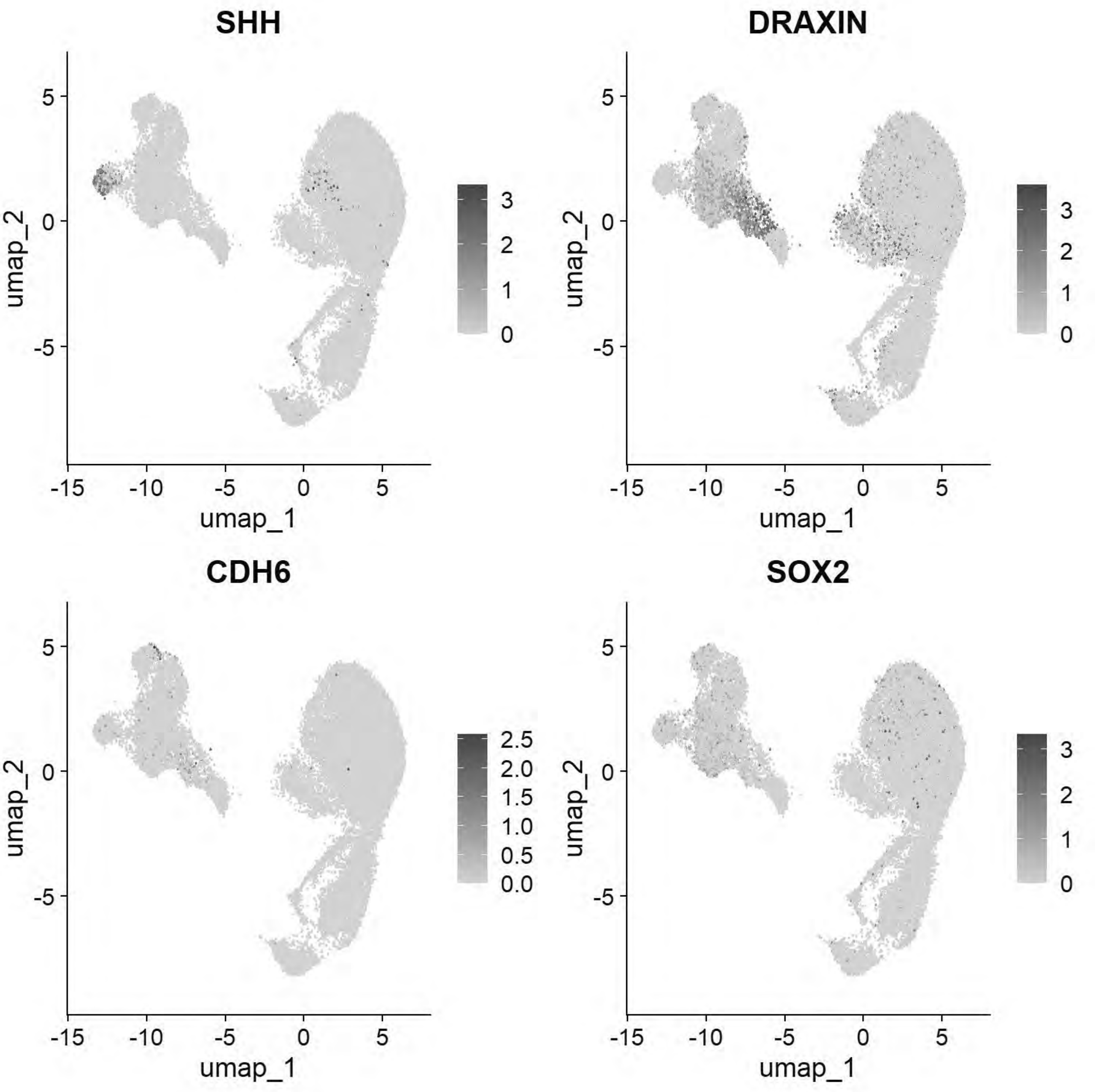


```
FeaturePlot(chick, features = c("PAX7","SHH","PAX6","OLIG2"))
```

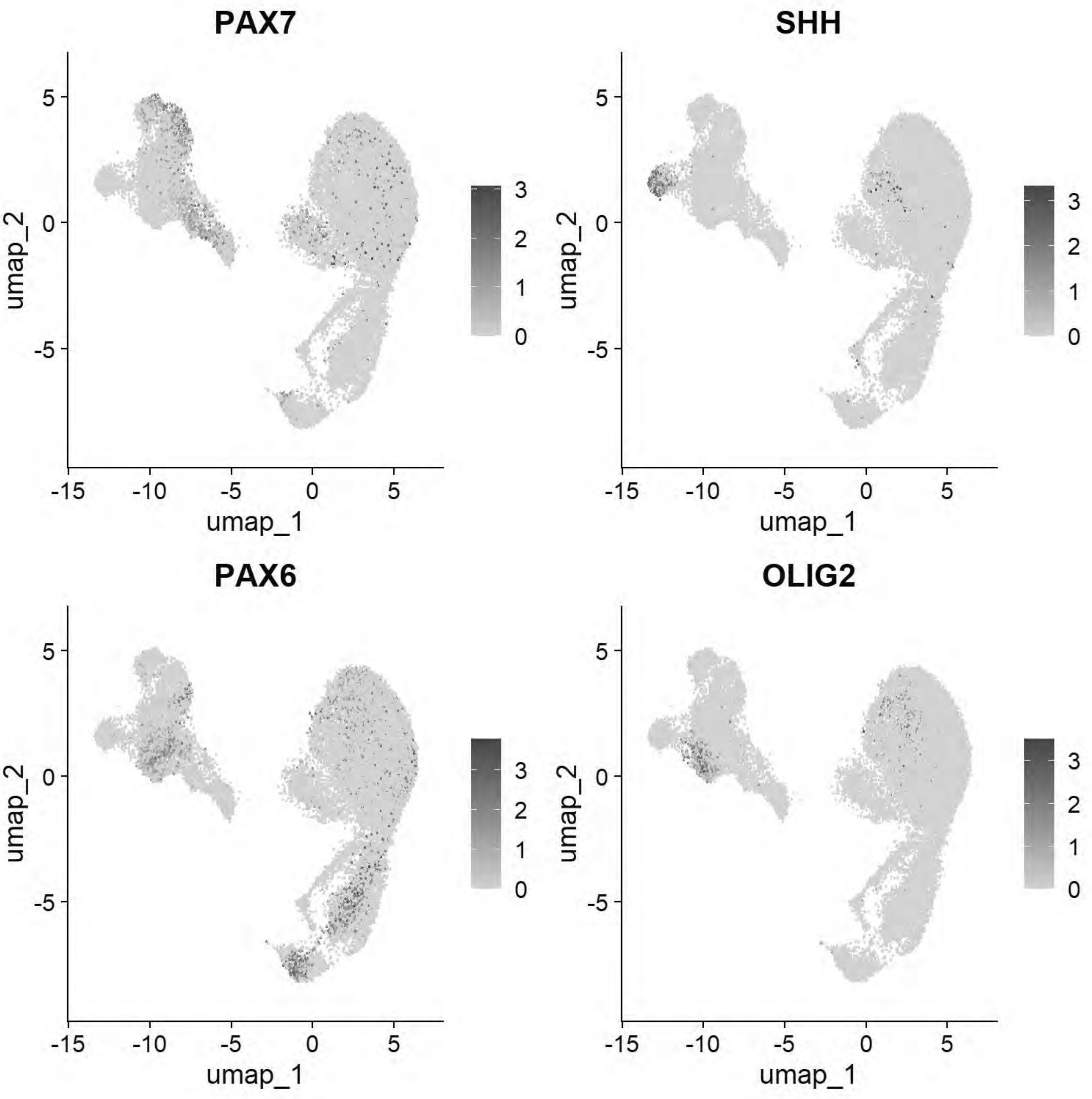


```
FeaturePlot(chick, features = c("PAX6","PAX2","PAX7","OLIG2"))
```

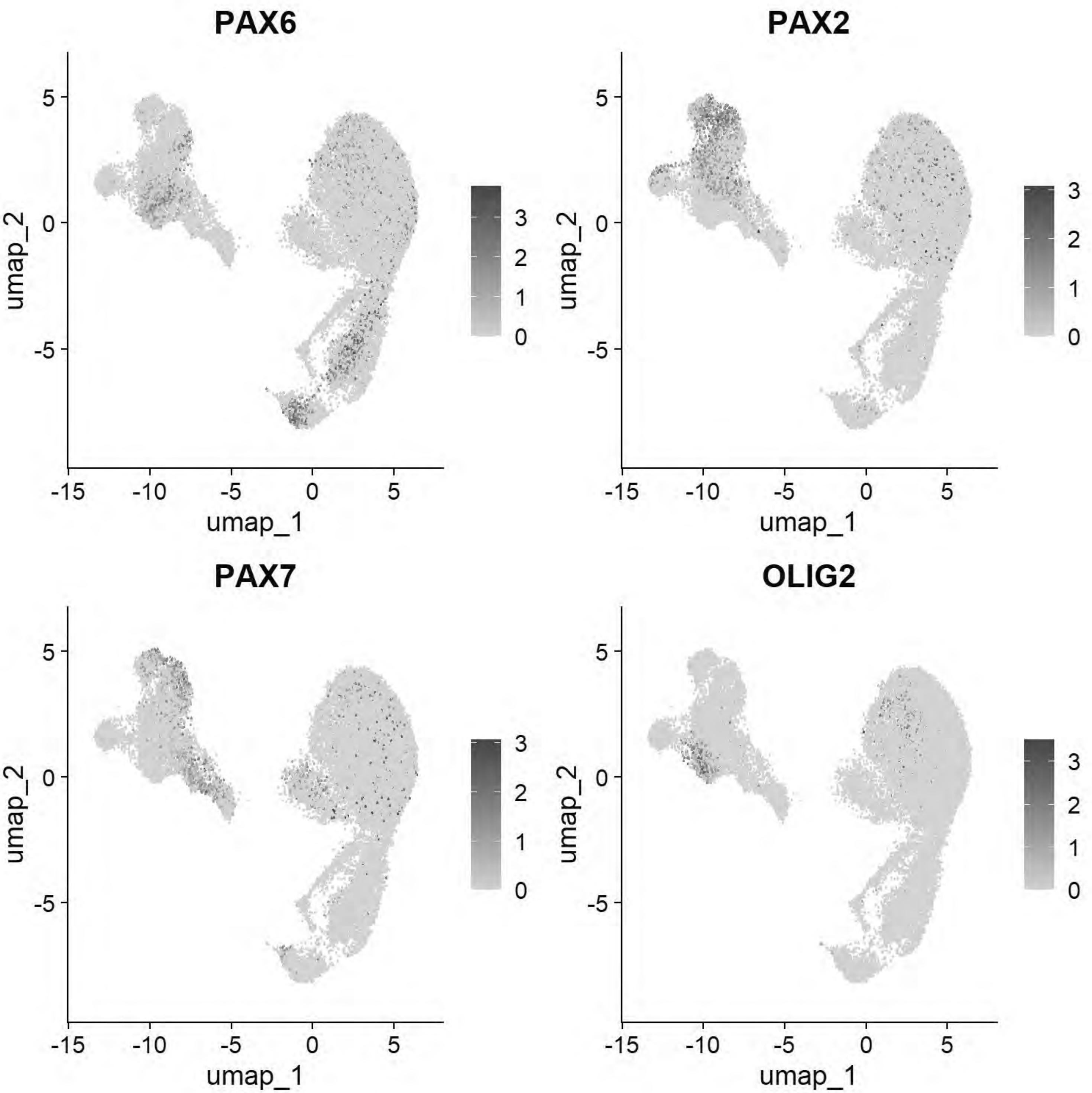


```
DimPlot(chick, group.by = "stage")
```

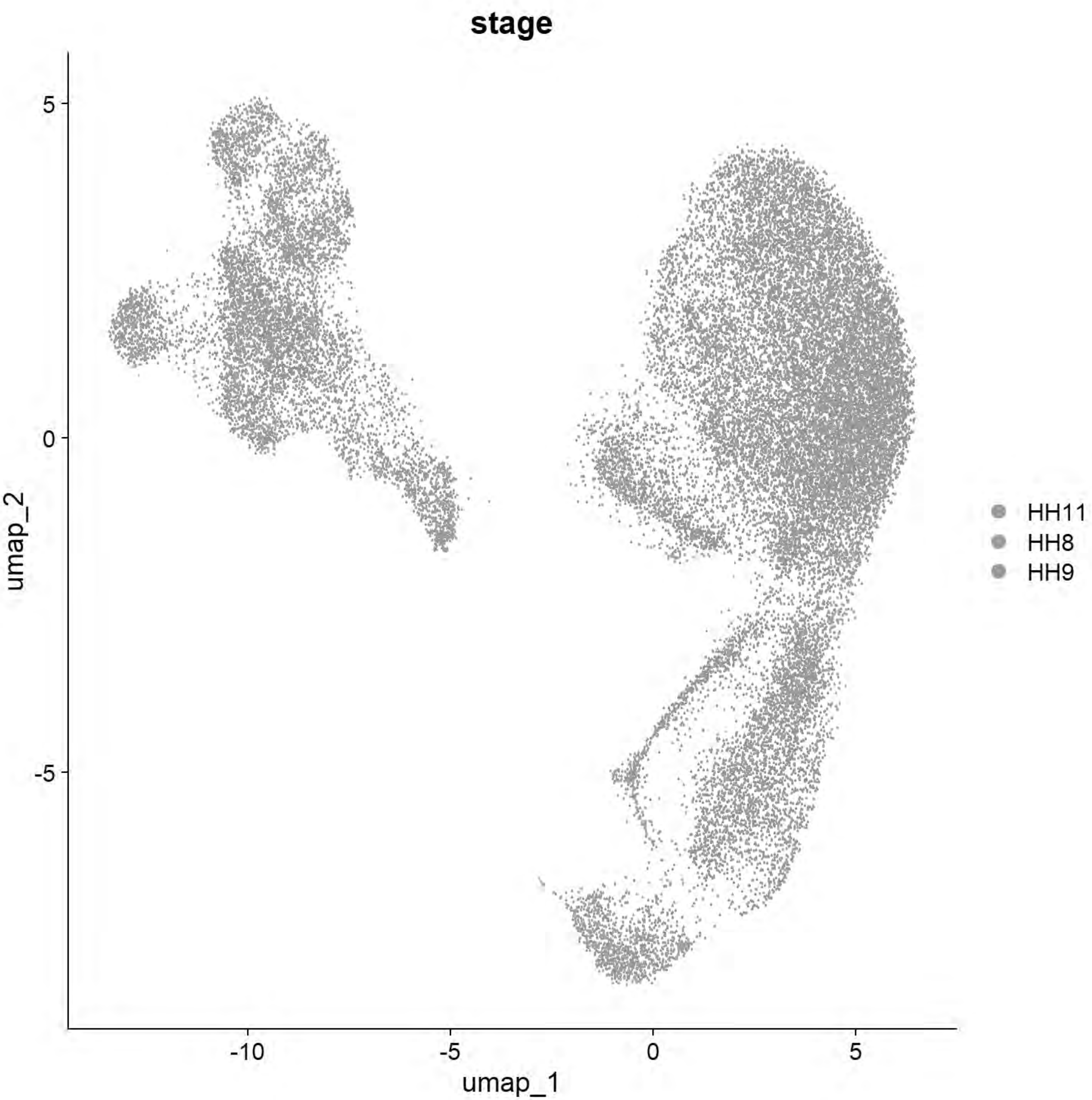


```
#FeaturePlot(chick, features = c("PAX7","TFAP2A","DLX5","BMP4","MSX1","DRAXIN","SOX10"))
```

```
markers=c("DRAXIN","PAX7","CDON","SNAI2",
                            "SOX9","TUBB3","TFAP2A",
                            "SOX10","TWIST1","CDH11",
                            "SOX21","CDH2",
                            "SHH","FOXA2","FOXA1",
                            "PAX2",
                            "WNT4","ZFHX4","NES",
                            "PTN","ZFHX3","GBX2",
```

```
                            "DLX5","SIX1","PAX6",
                            "DCN","WNT2B","KRT24","KRT7")
DotPlot(chick,features=markers)+coord_flip()
```

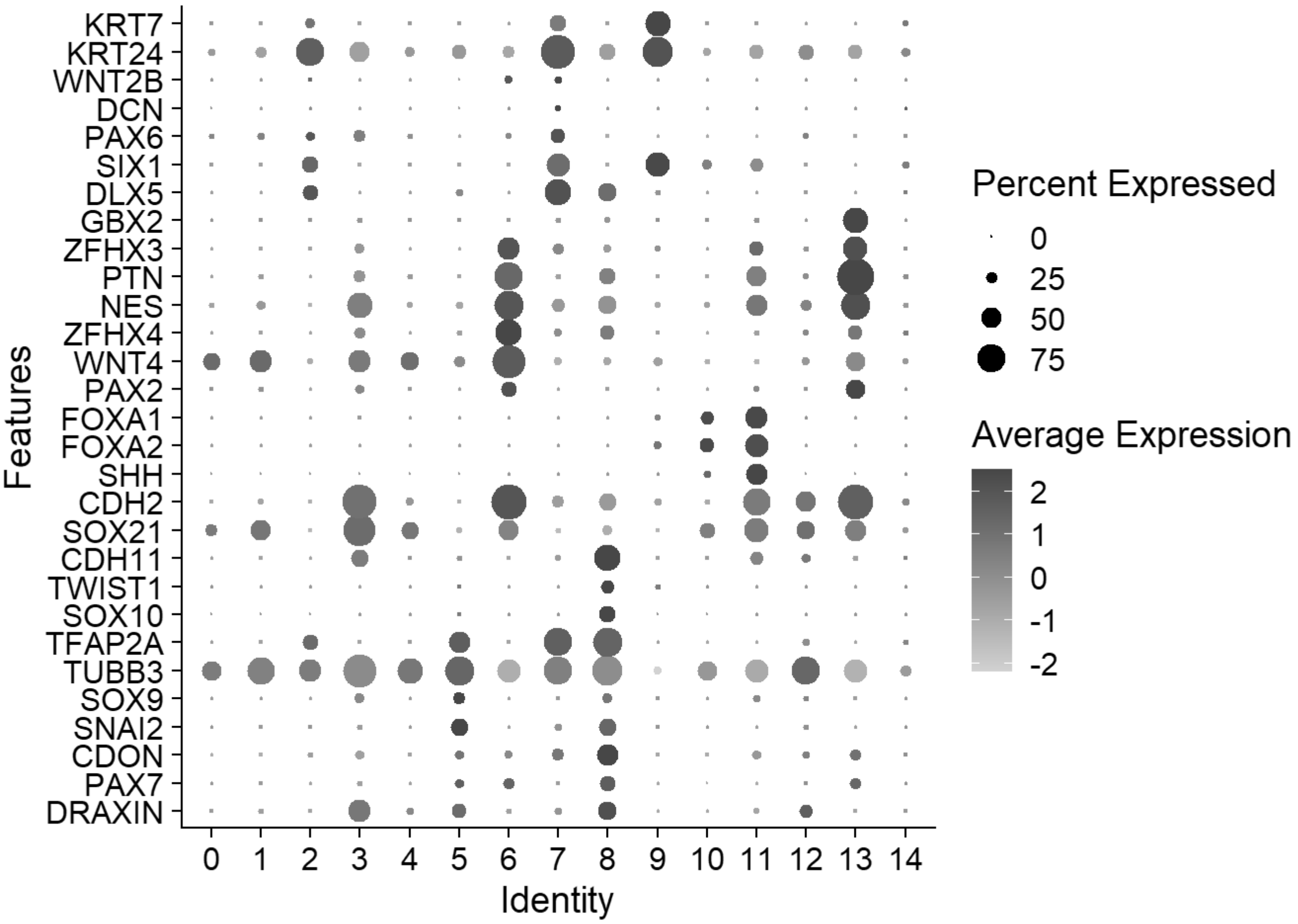


```
#Annotation of Cell Types
cluster_map <- c("0"="Uncommitted 1",
                 "1"="Uncommitted 2",
                 "2"="NNE/PPE",
                 "3"="NP",
                 "4"="Uncommitted 3",
                 "5"="preMig NC",
                 "6"="NT1",
                 "7"="Optic",
                 "8"="EMT/Mig NC",
                 "9"="PPE",
                 "10"="Uncommitted 4",
                 "11"="vNT",
                 "12"="tNP",
                 "13"="NT2",
                 "14"="Mito-High")
chick$celltype1 <- plyr::mapvalues(chick$seurat_clusters, from = names(cluster_map), to = clus
ter_map)

chick$celltype1 <- factor(chick$celltype1,
                          levels = c("preMig NC","EMT/Mig NC",
                                     "Uncommitted 1","Uncommitted 2","Uncommitted 3","tNP","NP
```

```
","NT1","NT2","Uncommitted 4","vNT",
                                     "NNE/PPE","Optic","PPE",
                                     "Mito-High"))
markers=c("SNAI2","SOX9","TUBB3","TFAP2A",
          "SOX10","TWIST1","CDH11",
          "SOX21","CDH2",
          "PAX2","NES",
          "PTN","ZFHX3","WNT4","ZFHX4","GBX2",
          "SHH","FOXA2","FOXA1",
          "DLX5","SIX1","PAX6",
          "DCN","WNT2B","KRT24","KRT7",
          "MT-ND2","MT-CO1")
p_dot=DotPlot(chick, features = markers, group.by = "celltype1") + # cluster.idents dendrogram
s the y-axis
  coord_flip() +
  # Force the dot sizes to scale more dramatically (range from 1 to 10)
  scale_size_continuous(range = c(1, 10)) +
  scale_color_gradientn(colors = c("grey90", "#313695", "#d73027"))+
  theme_light() +
  theme(
    axis.text.x = element_text(angle = 45, hjust = 1, color = "black", size = 12),
    axis.text.y = element_text(face = "italic", color = "black", size = 12),
    axis.title.x = element_blank(),
    axis.title.y = element_blank(),
    panel.grid.major = element_line(color = "grey90")
  )
```

```
## Scale for size is already present.
## Adding another scale for size, which will replace the existing scale.
## Scale for colour is already present.
## Adding another scale for colour, which will replace the existing scale.
```

```
p_dot
```

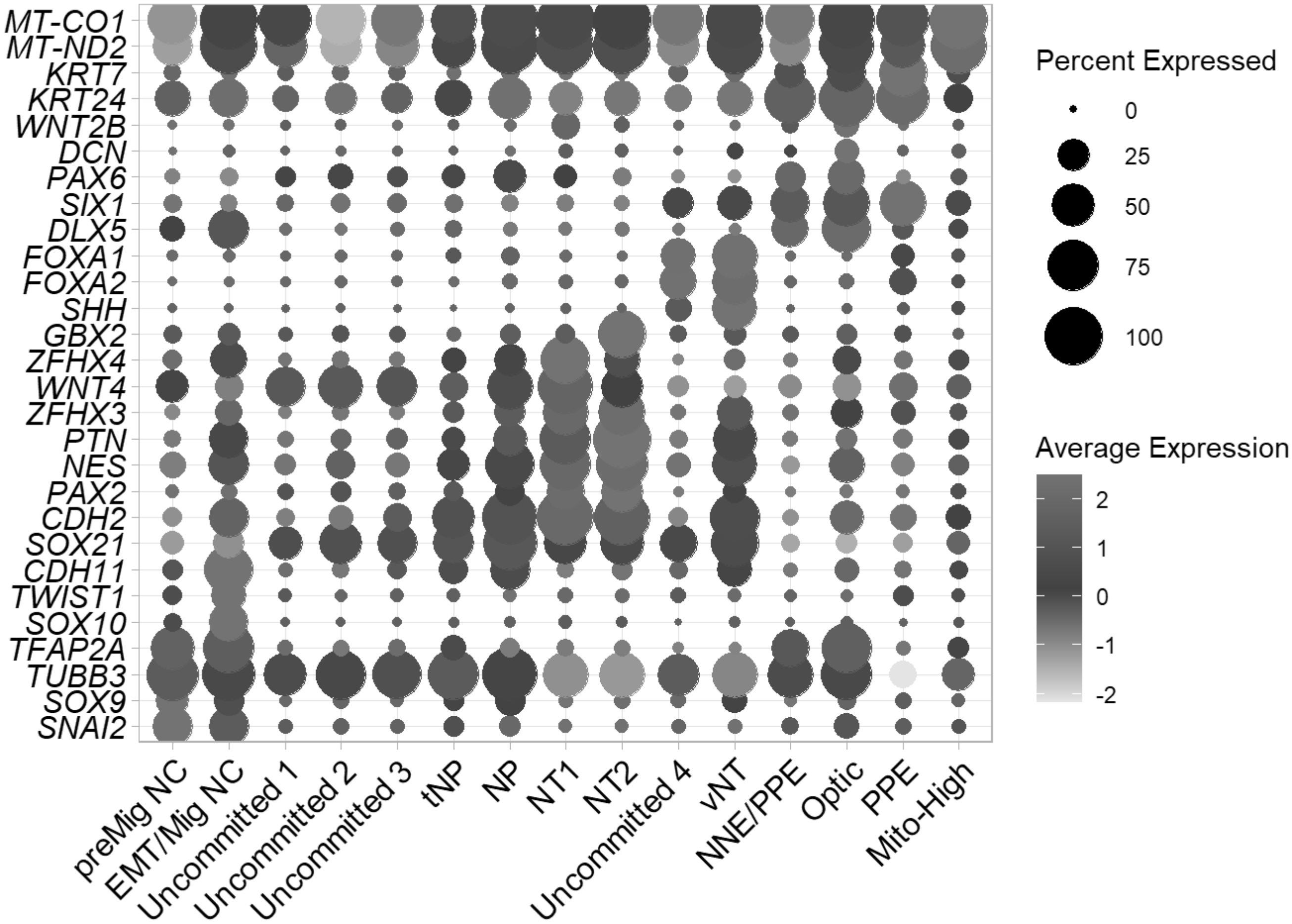


```
ggsave(filename = paste0("E:/Rogers Lab/Rogers NC EMT/Figures/Fig1_dotplot.png"),plot=p_dot, d
evice = 'png', width =6.5,height =7.5)
```

```
# Find Markers & Export directly to CSV
chick <- JoinLayers(chick)
Idents(chick) <- "celltype1" # Setting active identity ensures markers are calculated by cell
type
global_markers <- FindAllMarkers(chick, min.pct = 0.5, logfc.threshold = 0.5)
```

```
## Calculating cluster preMig NC
```

```
## Calculating cluster EMT/Mig NC
```

```
## Calculating cluster Uncommitted 1
```

```
## Calculating cluster Uncommitted 2
```

```
## Calculating cluster Uncommitted 3
```

```
## Calculating cluster tNP
```

```
## Calculating cluster NP
```

```
## Calculating cluster NT1
```

```
## Calculating cluster NT2
```

```
## Calculating cluster Uncommitted 4
```

```
## Calculating cluster vNT
```

```
## Calculating cluster NNE/PPE
```

```
## Calculating cluster Optic
```

```
## Calculating cluster PPE
```

```
## Calculating cluster Mito-High
```

```
write.csv(global_markers, "E:/Rogers Lab/Rogers NC EMT/Supplementary file 7_Ectodermal_Markers
_Annotated.csv", row.names = FALSE)

# Generate and Save Summary Panel
# A) UMAP
celltype_colors <-
c("preMig NC"="#8C96C6",
  "EMT/Mig NC"="#8C6BB1",
  "Uncommitted 1"="#BFD3E6",
  "Uncommitted 2"="#9ECAE1",
  "Uncommitted 3"="#6BAED6",
  "NP"="#41B6C4",
  "tNP"="#7FCDBB",
  "NT1"="#1D91C0",
  "NT2"="#225EA8",
  "Uncommitted 4"="#08519C",
  "vNT"="#08306B",
  "NNE/PPE"="#A1D99B",
  "Optic"="#006D2C",
  "PPE"="#238B45",
  "Mito-High"="#525252")

p_umap <- DimPlot(chick, group.by = "celltype1",cols = celltype_colors, label = FALSE, pt.size
=0.1) + ggtitle(NULL)+NoAxes()+
            NoLegend()
p_umap
```

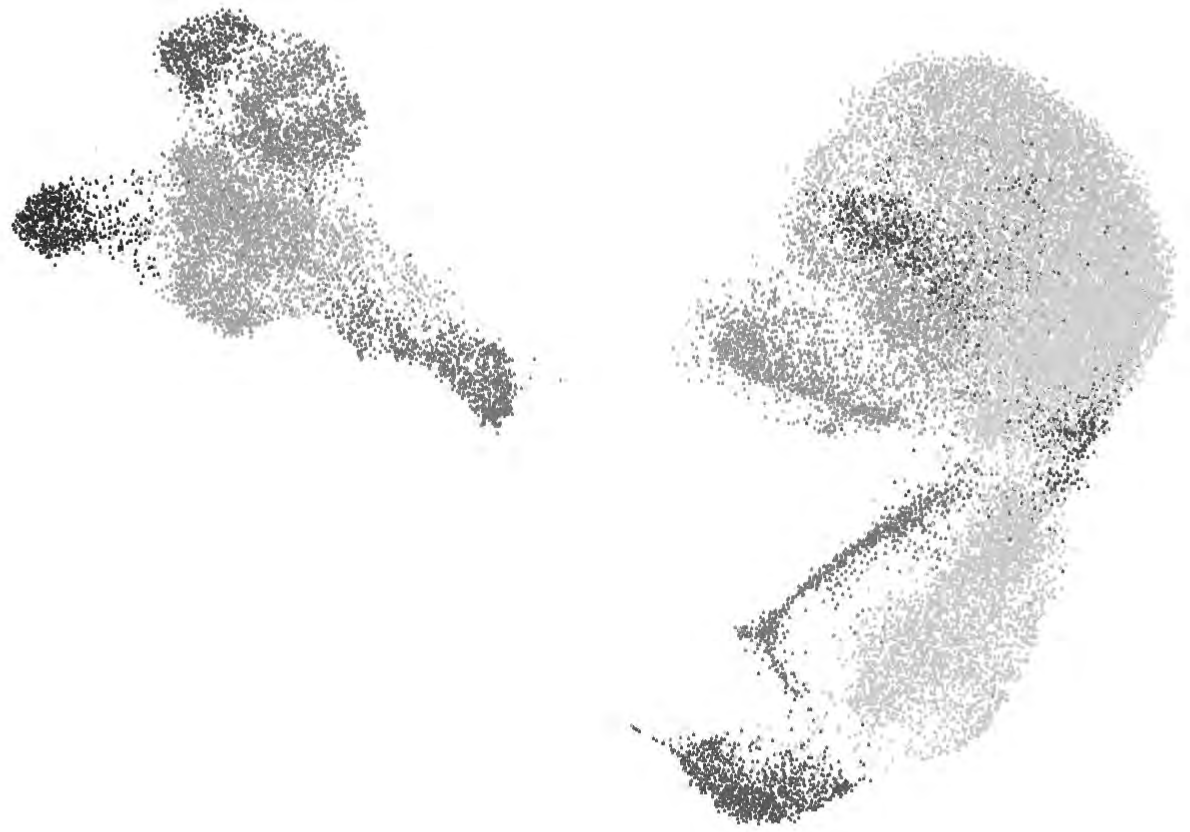

```
ggsave("E:/Rogers Lab/Rogers NC EMT/Figures/Fig1_Ectoderm_UMAP1.png", plot = p_umap, width = 6
, height = 6, dpi = 600)
```

```
chick@meta.data$stage <- factor(x = chick@meta.data$stage, levels = c("HH8","HH9","HH11"))
stage_colors <- c("HH8" = "#F59F2C", "HH9" = "#EF4E42", "HH11" = "#91182B")
p1= DimPlot(chick, group.by = "stage", cols=stage_colors, pt.size = 0.1)+NoAxes()
p1
```

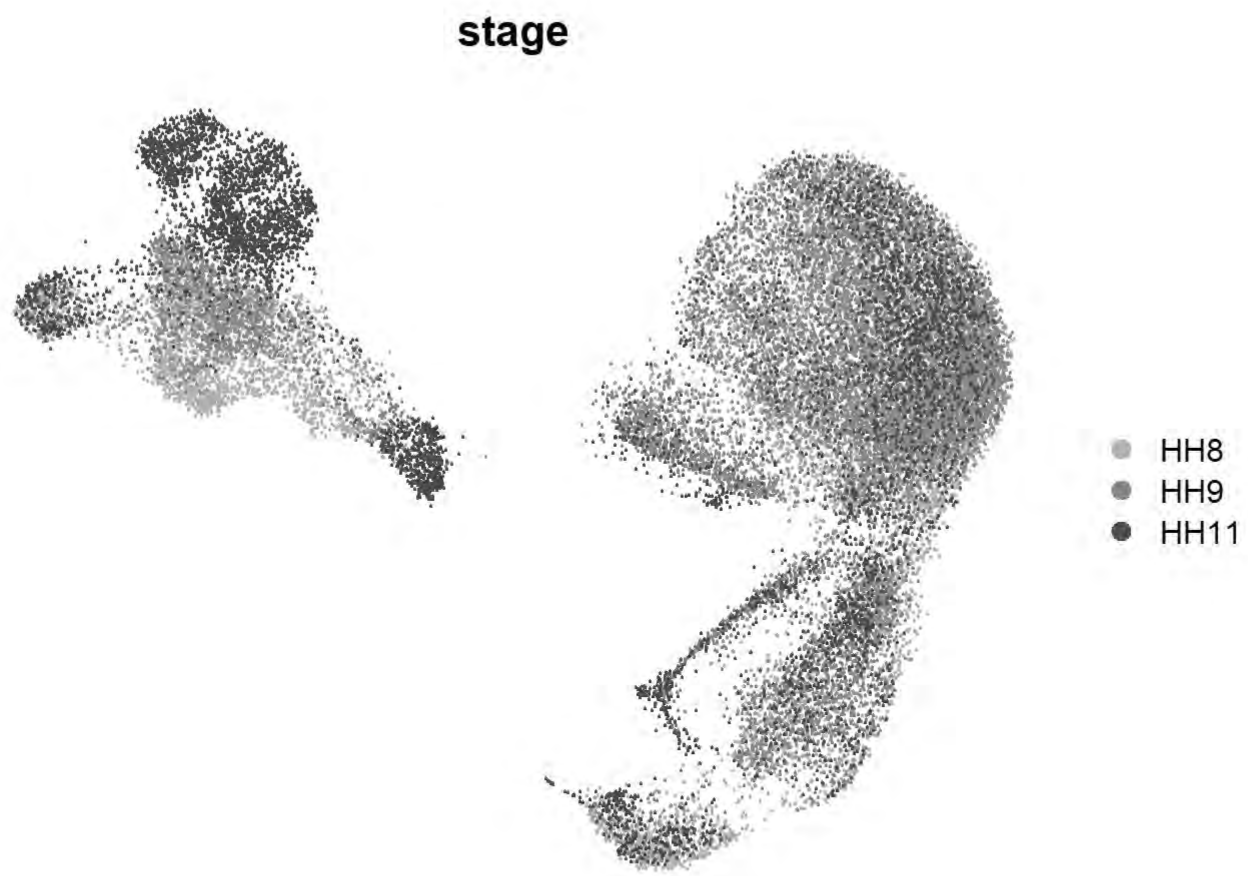


```
ggsave(p1, filename = "E:/Rogers Lab/Rogers NC EMT/Figures/Fig1_umap_stage1.png", width = 5, h
eight = 5, device='png', dpi=600)
p1= DimPlot(chick, group.by = "stage", cols=stage_colors, pt.size = 0.1)+NoAxes()+ggtitle(NULL
)+NoLegend()
ggsave(p1, filename = "E:/Rogers Lab/Rogers NC EMT/Figures/Fig1_umap_stage2.png", width = 5, h
eight = 5, device='png', dpi=600)
```

```
cptbl=table(chick@meta.data$stage,chick@meta.data$celltype1)
rownames(cptbl) <- c("HH8","HH9","HH11")
cptbl  <-  cbind(cptbl, rowSums(cptbl))
colnames(cptbl)[length(cptbl[1,])] <- "TOTAL"

tg = gridExtra::tableGrob(cptbl)
h = grid::convertHeight(sum(tg$heights), "in", TRUE)
w = grid::convertWidth(sum(tg$widths), "in", TRUE)
ggplot2::ggsave("E:/Rogers Lab/Rogers NC EMT/Figures/Fig1_TableCellType1.png", tg, width=w, he
ight=h, device = 'png', dpi =300)

cptbl=table(chick@meta.data$celltype1,chick@meta.data$stage)
rownames(cptbl) <- c("preMig NC","EMT/Mig NC",
                                     "Uncommitted 1","Uncommitted 2","Uncommitted 3","tNP","NP
","NT1","NT2","Uncommitted 4","vNT",
                                     "NNE/PPE","Optic","PPE",
                                     "Mito-High")
cptbl  <-  cbind(cptbl, rowSums(cptbl))
```

```
colnames(cptbl)[length(cptbl[1,])] <- "TOTAL"

tg = gridExtra::tableGrob(cptbl)
h = grid::convertHeight(sum(tg$heights), "in", TRUE)
w = grid::convertWidth(sum(tg$widths), "in", TRUE)
ggplot2::ggsave("E:/Rogers Lab/Rogers NC EMT/Figures/Fig1_TableCellType2.png", tg, width=w, he
ight=h, device = 'png', dpi =300)
```

```
saveRDS(chick,"E:\\Rogers Lab\\Rogers NC EMT\\HH8_9_11_Ectoderm_chick_v2.rds")
```

### Ectodermal Tubulin Mapping

```
tub_subunit_map <- c("TUBA1A"="A","TUBA1B"="A","TUBA1C"="A","TUBA3E"="A","TUBA8B"="A","TUBAL3"
="A",
       "TUBB"="B","TUBB1"="B","TUBB2A"="B","TUBB2B"="B","TUBB3"="B","TUBB4B"="B","TUBB6"="B",
       "TUBG1"="G")
family_text_colors <- c(
  "A" = "#6A3D9A",
  "B" = "#008B8B",
  "G" = "#B15928"
)

# Sort
tub_in_data <- c("TUBA1A","TUBA1B","TUBA1C","TUBA3E","TUBA8B","TUBAL3",
       "TUBB","TUBB1","TUBB2A","TUBB2B","TUBB3","TUBB4B","TUBB6",
       "TUBG1")

sorted_tub_df <- data.frame(Gene = tub_in_data) %>%
  mutate(Family = tub_subunit_map[Gene]) %>%
  filter(!is.na(Family)) %>%
  mutate(Family_Factor = factor(Family, levels = c("A", "B", "G"))) %>%
  arrange(Family_Factor, Gene)

organized_tub <- sorted_tub_df$Gene
axis_text_colors <- family_text_colors[sorted_tub_df$Family]

# Plot
Idents(chick) <- "celltype1"
p <- DotPlot(chick, features = rev(organized_tub)) +
  coord_flip() +
  scale_color_gradientn(colors = c("grey90", "#313695", "#d73027")) +

  scale_size_continuous(range = c(1, 10)) +
  theme_light() +
  theme(
    axis.text.x = element_text(angle = 45, hjust = 1, color = "black", size = 12),
    axis.text.y = element_text(face = "italic", color = rev(axis_text_colors), size = 12),
    axis.title.x = element_blank(),
    axis.title.y = element_blank(),
    panel.grid.major = element_line(color = "grey90")
  )
```

```
## Scale for colour is already present.
## Adding another scale for colour, which will replace the existing scale.
## Scale for size is already present.
## Adding another scale for size, which will replace the existing scale.
```

```
p
```

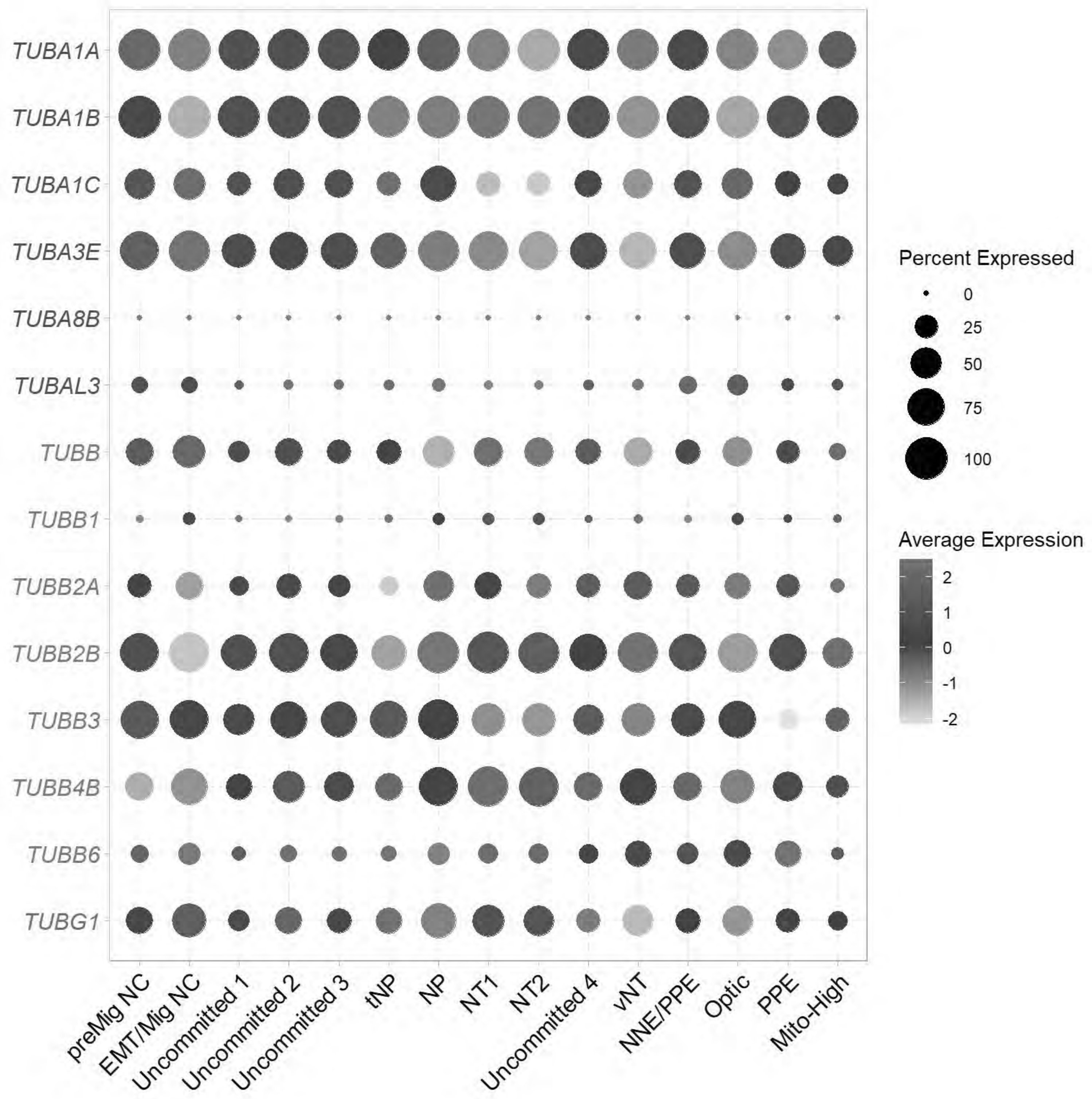


```
ggsave(filename = paste0("E:/Rogers Lab/Rogers NC EMT/Figures/Fig1_tub_categorization.png"), p
lot = p, device = 'png', width = 7.5, height = 5)
```

```
f1=FeaturePlot(chick, features=c("TUBA1A"))+NoAxes()+scale_color_gradientn(colors = c("grey90"
, "#2EACBD", "#08306B"),limits = c(0, 5))
```

```
## Scale for colour is already present.
## Adding another scale for colour, which will replace the existing scale.
```

```
f2=FeaturePlot(chick, features=c("TUBA1B"))+NoAxes()+scale_color_gradientn(colors=c("grey90",
"#2EACBD", "#08306B"),limits = c(0, 5))
```

```
## Scale for colour is already present.
## Adding another scale for colour, which will replace the existing scale.
```

```
f3=FeaturePlot(chick, features=c("TUBB3"))+NoAxes()+scale_color_gradientn(colors=c("grey90", "
#2EACBD", "#08306B"),limits = c(0, 5))
```

```
## Scale for colour is already present.
## Adding another scale for colour, which will replace the existing scale.
```

```
f4=FeaturePlot(chick, features=c("TUBB2A"))+NoAxes()+scale_color_gradientn(colors=c("grey90",
"#2EACBD", "#08306B"),limits = c(0, 5))
```

```
## Scale for colour is already present.
## Adding another scale for colour, which will replace the existing scale.
```

```
f5=FeaturePlot(chick, features=c("TUBB2B"))+NoAxes()+scale_color_gradientn(colors=c("grey90",
"#2EACBD", "#08306B"),limits = c(0, 5))
```

```
## Scale for colour is already present.
## Adding another scale for colour, which will replace the existing scale.
```

```
f6=FeaturePlot(chick, features=c("TUBB4B"))+NoAxes()+scale_color_gradientn(colors=c("grey90",
"#2EACBD", "#08306B"),limits = c(0, 5))
```

```
## Scale for colour is already present.
## Adding another scale for colour, which will replace the existing scale.
```

```
f= ggarrange(plotlist = list(f1,f2,f3,f4,f5,f6), ncol = 3, nrow = 2)
f
```

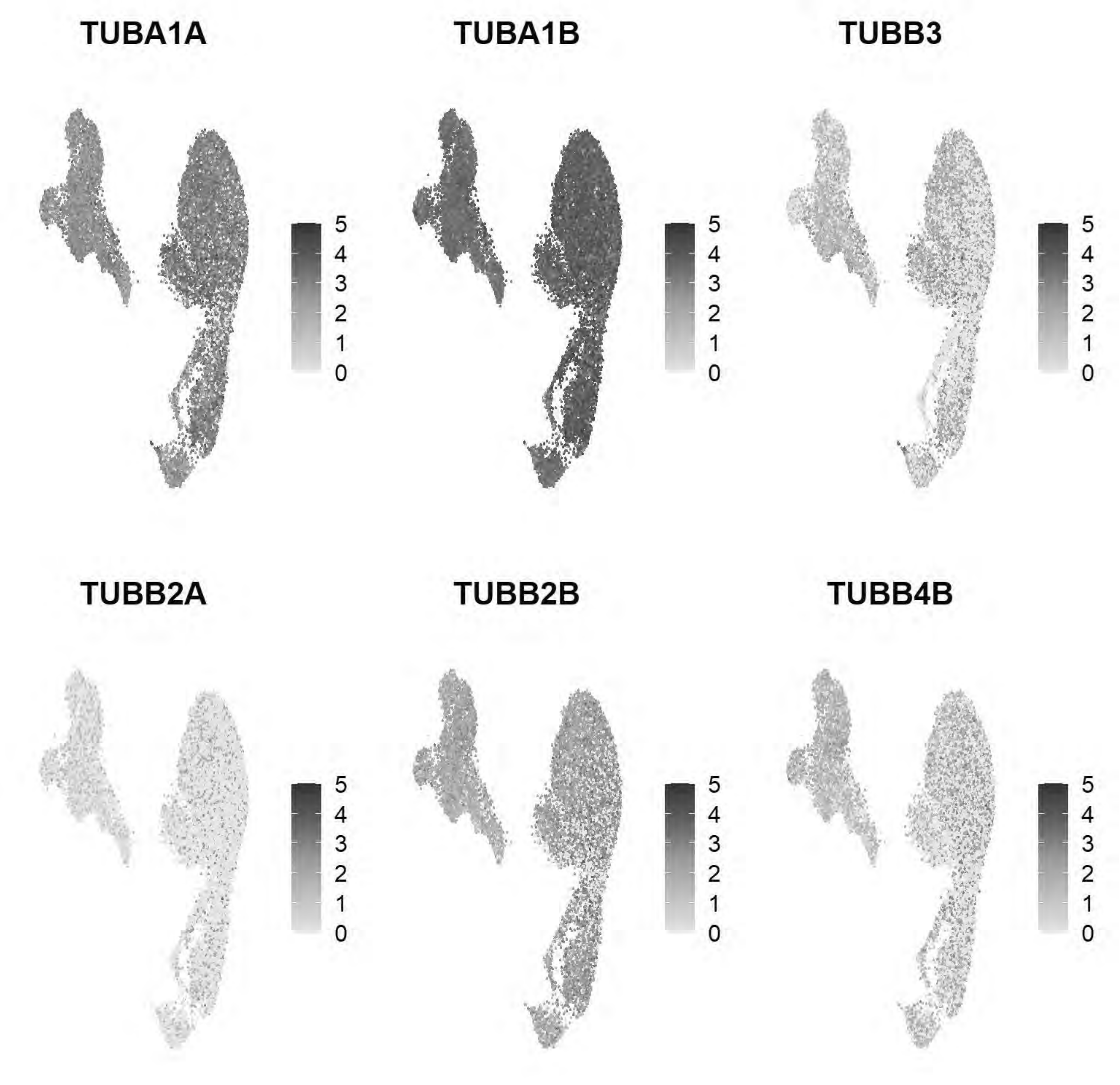


```
ggsave(f, filename = "E:/Rogers Lab/Rogers NC EMT/Figures/Fig1_selecttubs_FeaturePlot1.png", w
idth = 10.5, height = 6, device='png', dpi=600)
```

### Global Dynein Mapping

```
dyn_subunit_map <- c(
  "DYNC1H1"="C1", "DYNC1I1"="C1","DYNC1I2"="C1","DYNC1LI1"="C1","DYNC1LI2"="C1",
  "DYNLL1"="C1+C2","DYNLL2"="C1+C2","DYNLRB1"="C1+C2","DYNLRB2"="C1+C2","DYNLT1"="C1+C2","DYNL
T3"="C1+C2",
```

```
  "DYNLT2"="C2","DYNC2H1"="C2","DYNC2I1"="C2","DYNC2I2"="C2","DYNC2LI1"="C2")

family_text_colors <- c(
  "C1"        = "#008B8B",
  "C1+C2" = "#B15928",
  "C2"        = "#6A3D9A"
)

dyn_in_data <- rownames(chick)[grep("^DYN", toupper(rownames(chick)))]

sorted_dyn_df <- data.frame(Gene = dyn_in_data) %>%
  mutate(Family = dyn_subunit_map[Gene]) %>%
  filter(!is.na(Family)) %>%
  mutate(Family_Factor = factor(Family, levels = c("C1", "C1+C2", "C2"))) %>%
  arrange(Family_Factor, Gene)

organized_dyn <- sorted_dyn_df$Gene
axis_text_colors <- family_text_colors[sorted_dyn_df$Family]

Idents(chick) <- "celltype1"

p <- DotPlot(chick, features = rev(organized_dyn)) +
  coord_flip() +
  scale_color_gradientn(colors = c("grey90", "#313695", "#d73027")) +
  scale_size_continuous(range = c(1, 10)) +
  theme_light() +
  theme(
    axis.text.x = element_text(angle = 45, hjust = 1, color = "black", size = 12),
    axis.text.y = element_text(face = "italic", color = rev(axis_text_colors), size = 12),
    axis.title.x = element_blank(),
    axis.title.y = element_blank(),
    panel.grid.major = element_line(color = "grey90")
  )
```

```
## Scale for colour is already present.
## Adding another scale for colour, which will replace the existing scale.
## Scale for size is already present.
## Adding another scale for size, which will replace the existing scale.
```

```
p
```

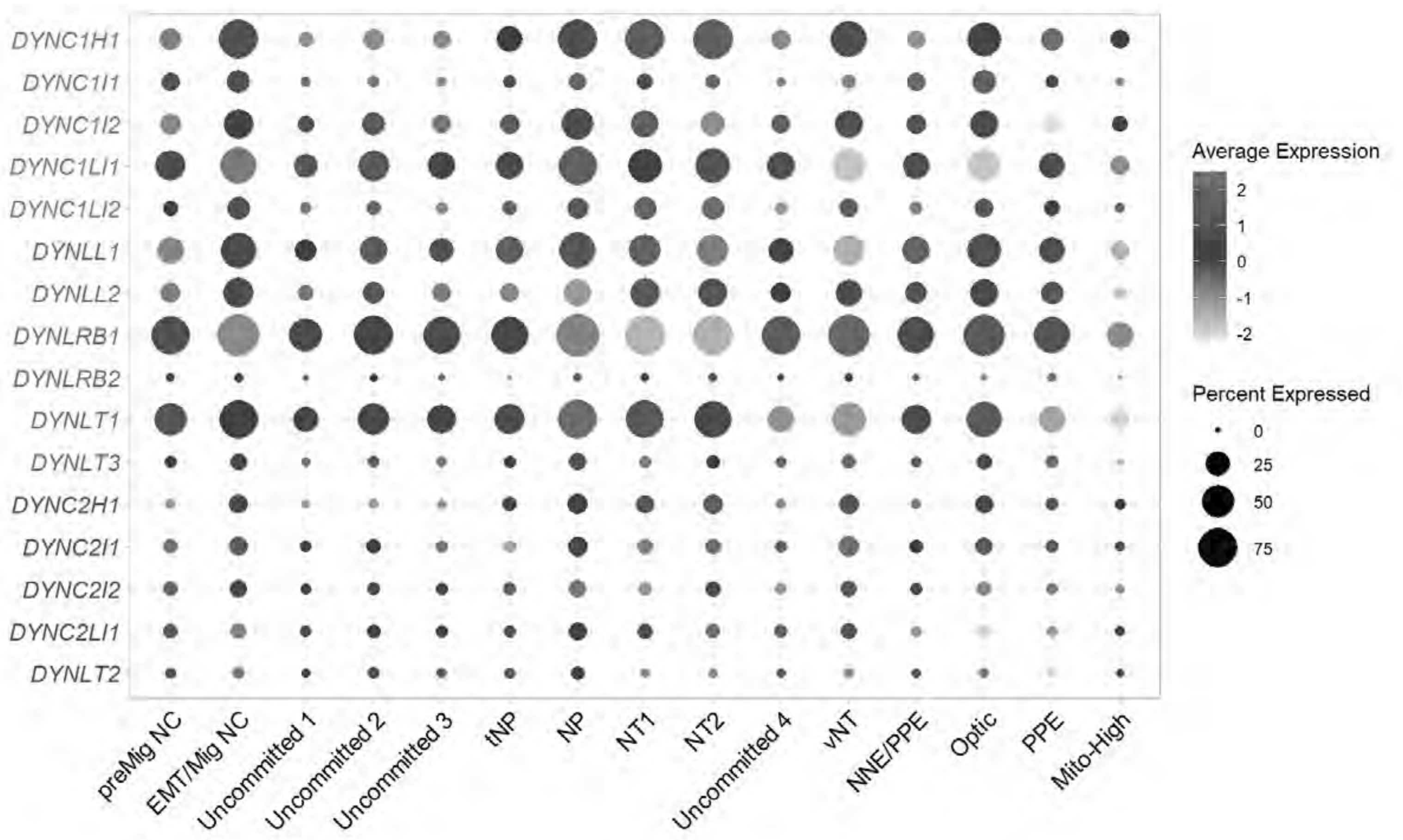


```
ggsave(filename = paste0("E:/Rogers Lab/Rogers NC EMT/Figures/Fig1_dyn_dotplot.png"), plot = p
, device = 'png', width = 7, height = 5)
```

### Global Kinesin Mapping

```
#Gene to superfamily mapping
gene_superfamily_map <- c(
  "KIF2A"="M", "KIF2B"="M", "KIF2C"="M",
  "KIFC1"="C", "KIFC2"="C", "KIFC3"="C",
  "KIF5A"="N-1", "KIF5B"="N-1", "KIF5C"="N-1",
  "KIF6"="N-2", "KIF7"="N-2", "KIF8"="N-2", "KIF9"="N-2", "KIF11"="N-2",
  "KIF1A"="N-3", "KIF1B"="N-3", "KIF1C"="N-3", "KIF13A"="N-3", "KIF13B"="N-3", "KIF14"="N-3",
"KIF16A"="N-3", "KIF16B"="N-3",
  "KIF3A"="N-4", "KIF3B"="N-4", "KIF3C"="N-4", "KIF17"="N-4",
  "KIF4A"="N-5", "KIF4B"="N-5", "KIF21A"="N-5", "KIF21B"="N-5",
  "KIF20A"="N-6", "KIF20B"="N-6", "KIF23"="N-6",
  "KIF10"="N-7",
  "KIF18A"="N-8", "KIF18B"="N-8", "KIF19A"="N-8", "KIF19B"="N-8", "KIF22"="N-8",
  "KIF12"="N-9",
  "KIF15"="N-10",
  "KIF24"="N-11", "KIF25"="N-11", "KIF26A"="N-11", "KIF26B"="N-11"
)

#
family_text_colors <- c(
```

```
  "C"    = "#1A1A1A", # Crisp Black
  "M"    = "#E31A1C", # Bright Red
  "N-1"  = "#33A02C", # Strong Green
  "N-2"  = "#1F78B4", # Strong Blue
  "N-3"  = "#FF7F00", # Vibrant Orange
  "N-4"  = "#6A3D9A", # Vibrant Purple
  "N-5"  = "#008B8B", # Dark Cyan
  "N-6"  = "#B15928", # Burnt Orange
  "N-7"  = "#E7298A", # Hot Pink
  "N-8"  = "#00BA38", # Bright Lime/Green
  "N-9"  = "#619CFF", # Bright Sky Blue
  "N-10" = "#B8860B", # Goldenrod
  "N-11" = "#8B4513"  # Saddle Brown
)

kifs_in_data <- rownames(chick)[grep("^KIF", toupper(rownames(chick)))]
sorted_kif_df <- data.frame(Gene = kifs_in_data) %>%
  mutate(Family = gene_superfamily_map[Gene]) %>%
  filter(!is.na(Family)) %>%
  mutate(Family_Factor = factor(Family, levels = c("C", "M", "N-1", "N-2", "N-3", "N-4", "N-5"
, "N-6", "N-7", "N-8", "N-9", "N-10", "N-11"))) %>%
  arrange(Family_Factor, Gene)

organized_kifs <- sorted_kif_df$Gene
axis_text_colors <- family_text_colors[sorted_kif_df$Family]

Idents(chick) <- "celltype1"
p <- DotPlot(chick, features = rev(organized_kifs)) +
  coord_flip() +
  scale_color_gradientn(colors = c("grey90", "#313695", "#d73027")) +
  scale_size_continuous(range = c(1, 10)) +
  theme_light() +
  theme(
    axis.text.x = element_text(angle = 45, hjust = 1, color = "black", size = 12),
    axis.text.y = element_text(face = "italic", color = rev(axis_text_colors), size = 12),
    axis.title.x = element_blank(),
    axis.title.y = element_blank(),
    panel.grid.major = element_line(color = "grey90")
  )
```

```
## Scale for colour is already present.
## Adding another scale for colour, which will replace the existing scale.
## Scale for size is already present.
## Adding another scale for size, which will replace the existing scale.
```

```
p
```

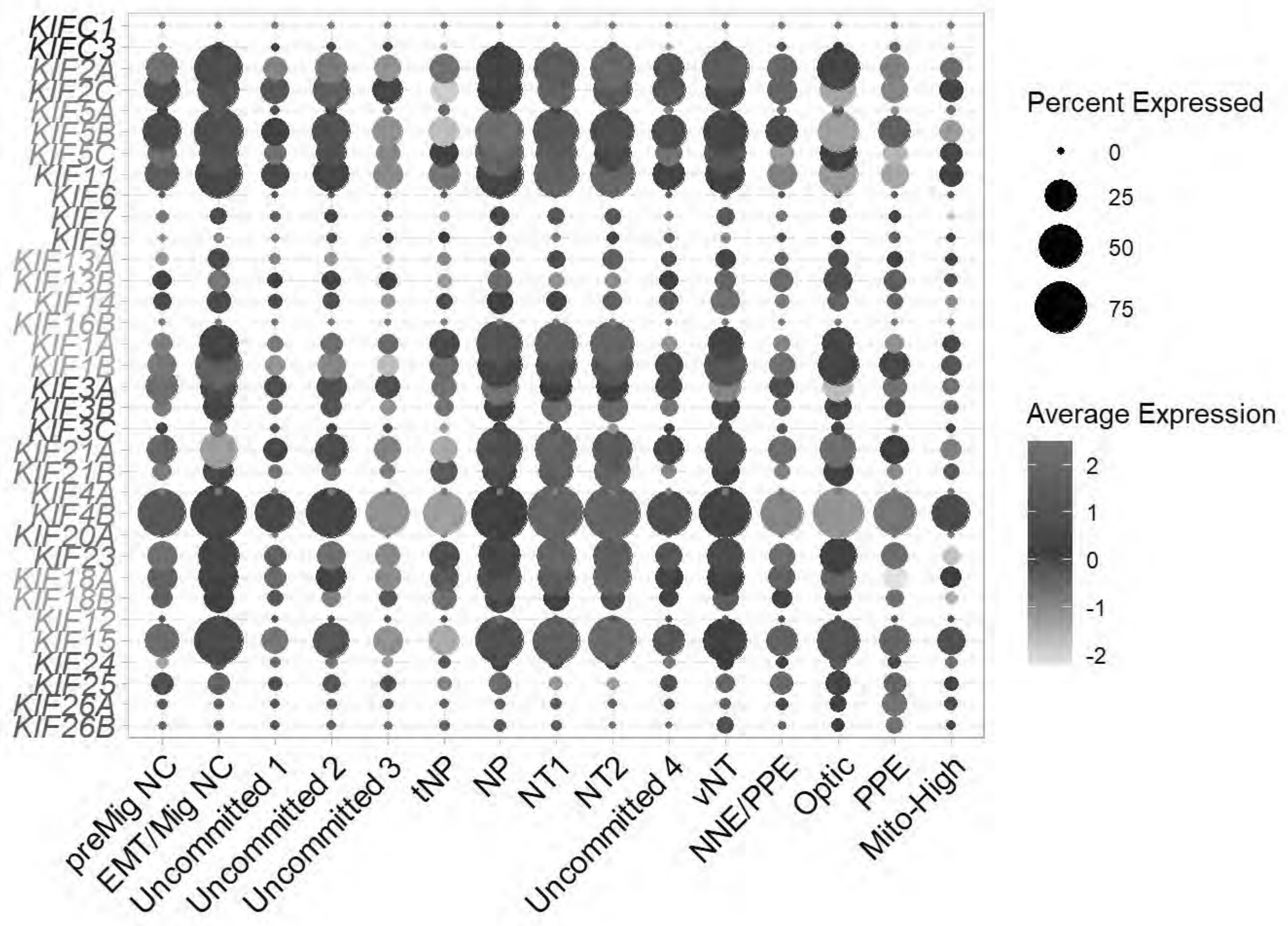


```
ggsave(filename = paste0("E:/Rogers Lab/Rogers NC EMT/Figures/Fig1_kif_dotplot.png"), plot = p
, device = 'png', width = 7, height = 7)
```

```
f1=FeaturePlot(chick, features=c("DYNC1LI1"))+NoAxes()+scale_color_gradientn(colors = c("grey9
0", "#2EACBD", "#08306B"), limit=c(0,4))
```

```
## Scale for colour is already present.
## Adding another scale for colour, which will replace the existing scale.
```

```
f1
```

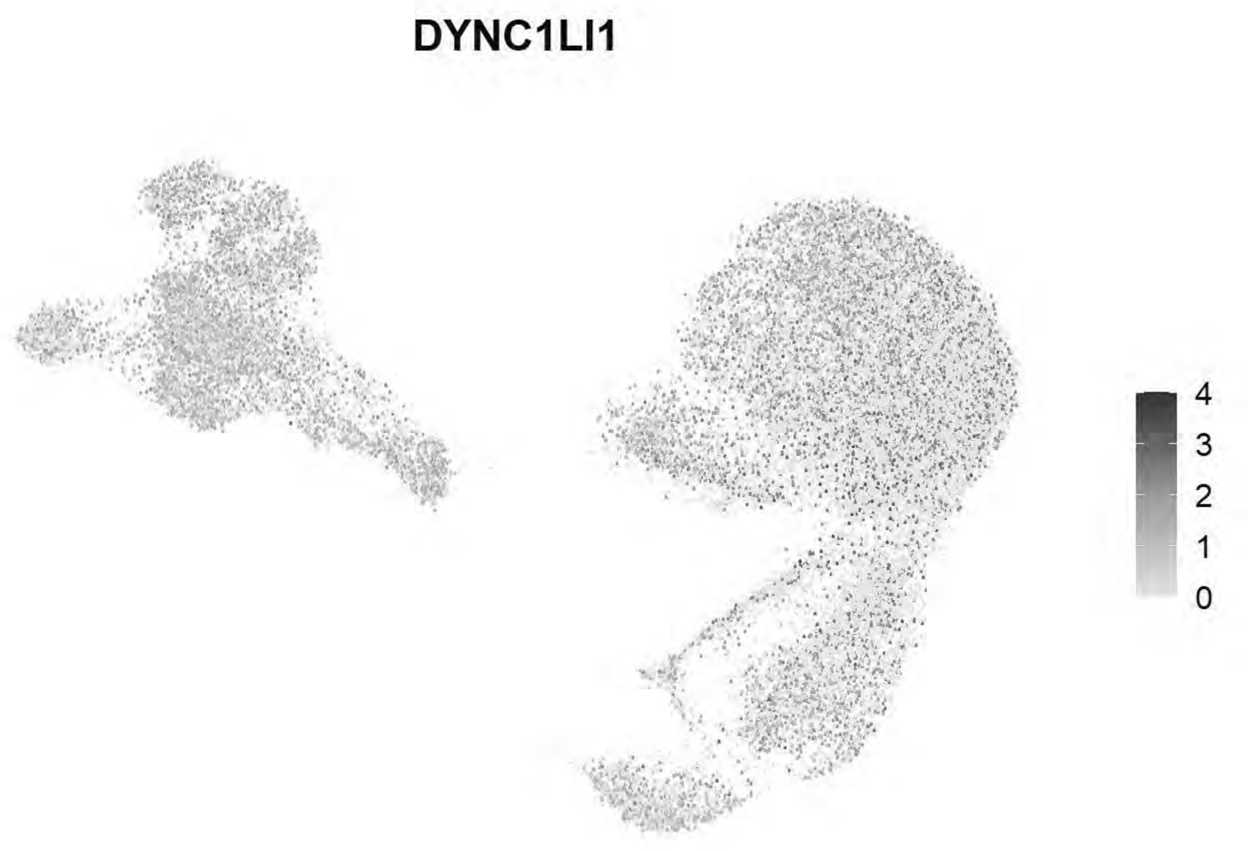


```
f2=FeaturePlot(chick, features=c("KIF11"))+NoAxes()+scale_color_gradientn(colors=c("grey90", "
#2EACBD", "#08306B"), limit=c(0,4))
```

```
## Scale for colour is already present.
## Adding another scale for colour, which will replace the existing scale.
```

```
f2
```

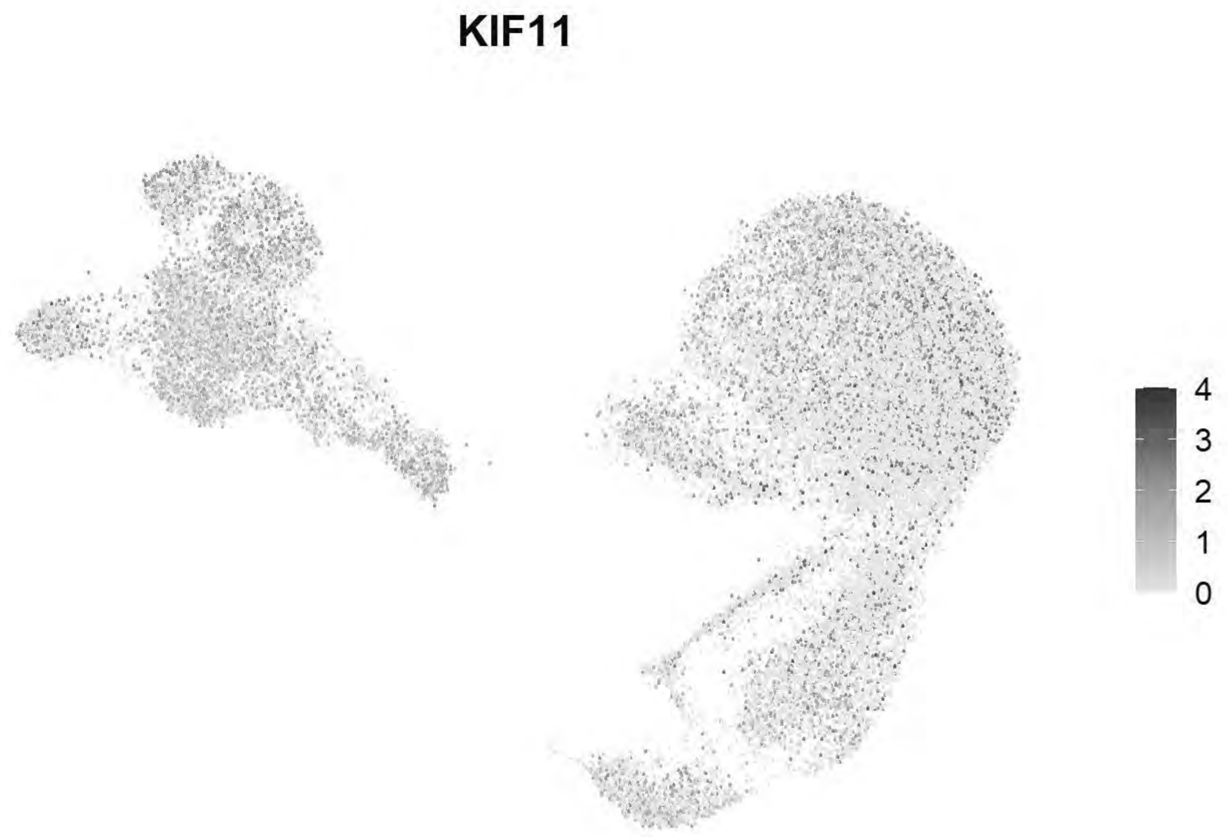


```
ggsave(f1, filename = "E:/Rogers Lab/Rogers NC EMT/Figures/FigX_dync1li1.png", width = 3, height = 3, device='png', dpi=600)
ggsave(f2, filename = "E:/Rogers Lab/Rogers NC EMT/Figures/FigX_Kif11.png", width = 3, height = 3, device='png', dpi=600)
```

### END OF SCRIPT
