## Supplementary File 2 for "A spatial and temporal atlas of tubulin isotype gene expression during vertebrate embryonic development"

A

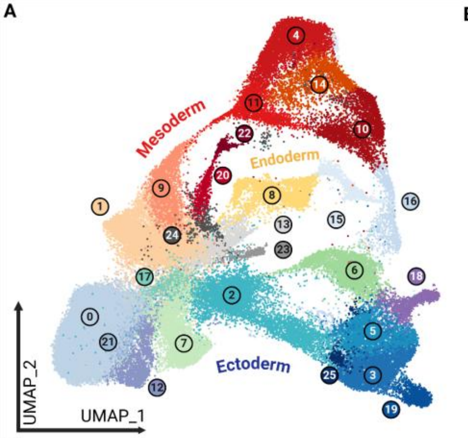


B

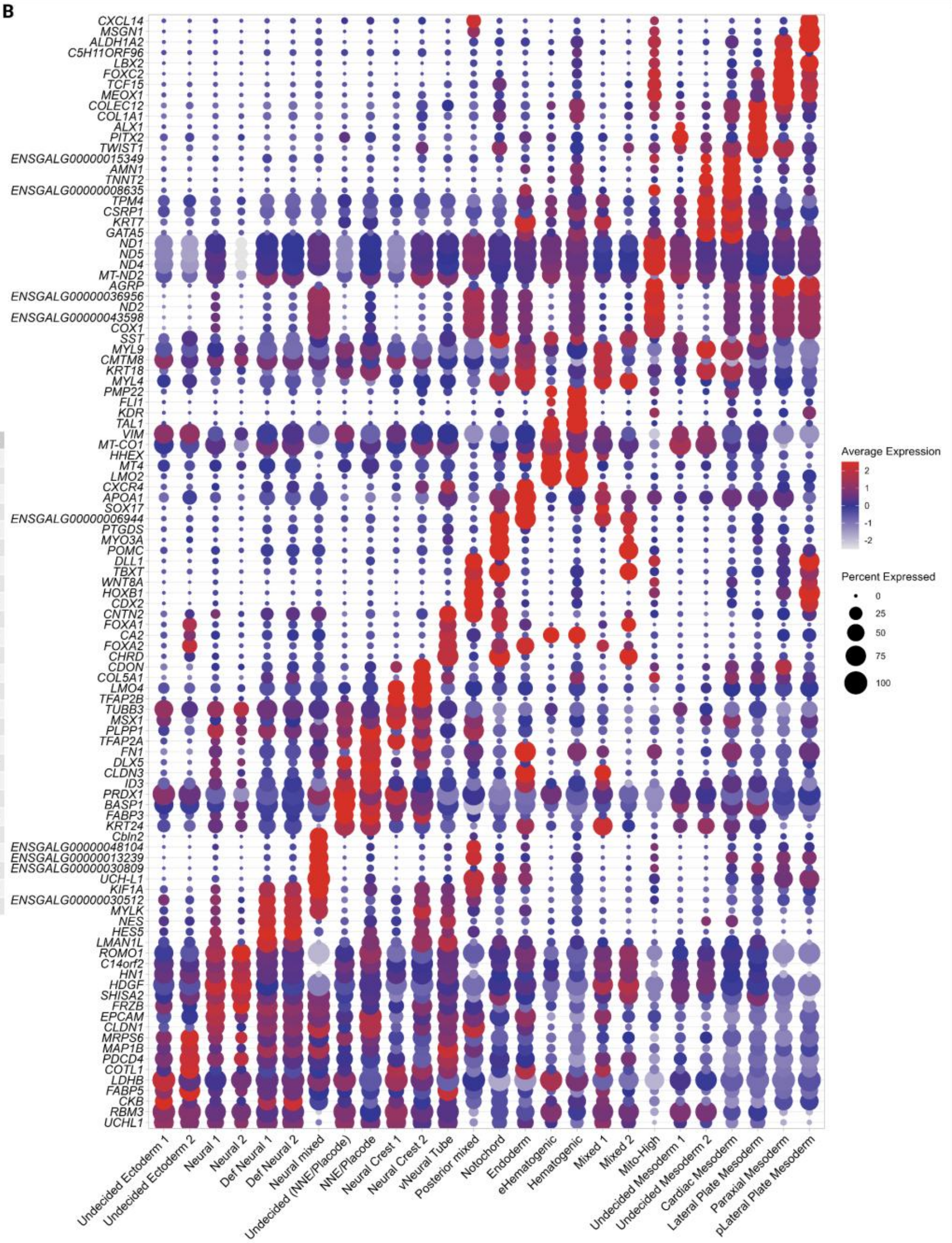


C

| | HH4 | HH5 | HH6 | HH7 | HH8 | HH9 | HH11 | TOTAL |
|---|---|---|---|---|---|---|---|---|
| 0 *Undecided Ectoderm 1* | 8 | 1642 | 5 | 1433 | 4710 | 7351 | 1573 | 16722 |
| 21 *Undecided Ectoderm 2* | 0 | 38 | 3 | 36 | 130 | 337 | 39 | 583 |
| 2 *Neural 1* | 286 | 2291 | 101 | 1777 | 1350 | 266 | 287 | 6358 |
| 17 *Neural 2* | 3 | 169 | 0 | 530 | 466 | 167 | 42 | 1377 |
| 3 *Def Neural 1* | 155 | 1931 | 16 | 1095 | 1013 | 761 | 1038 | 6009 |
| 5 *Def Neural 2* | 113 | 1109 | 13 | 972 | 707 | 443 | 1159 | 4516 |
| 25 *Neural mixed* | 1 | 31 | 64 | 175 | 0 | 0 | 5 | 276 |
| 7 *Undecided (NNE/Placode)* | 12 | 507 | 4 | 291 | 1023 | 1479 | 480 | 3796 |
| 6 *NNE/Placode* | 290 | 1730 | 102 | 762 | 661 | 134 | 297 | 3976 |
| 12 *Neural Crest 1* | 0 | 6 | 0 | 38 | 391 | 1142 | 200 | 1777 |
| 18 *Neural Crest 2* | 3 | 90 | 3 | 136 | 273 | 246 | 569 | 1320 |
| 19 *vNeural Tube* | 9 | 277 | 17 | 312 | 218 | 165 | 248 | 1246 |
| 16 *Posterior mixed* | 93 | 264 | 388 | 605 | 38 | 2 | 25 | 1415 |
| 15 *Notochord* | 32 | 430 | 183 | 591 | 131 | 55 | 48 | 1470 |
| 8 *Endoderm* | 254 | 1416 | 292 | 692 | 490 | 53 | 167 | 3364 |
| 20 *eHematogenic* | 4 | 10 | 9 | 12 | 267 | 456 | 63 | 821 |
| 22 *Hematogenic* | 35 | 21 | 116 | 69 | 121 | 85 | 70 | 517 |
| 13 *Mixed 1* | 10 | 555 | 15 | 207 | 401 | 431 | 115 | 1734 |
| 23 *Mixed 2* | 1 | 146 | 10 | 119 | 132 | 104 | 1 | 513 |
| 24 *Mito-High* | 65 | 120 | 120 | 112 | 23 | 4 | 2 | 446 |
| 1 *Undecided Mesoderm 1* | 32 | 927 | 23 | 803 | 3542 | 2594 | 247 | 8168 |
| 9 *Undecided Mesoderm 2* | 26 | 240 | 11 | 415 | 1651 | 734 | 41 | 3118 |
| 11 *Cardiac Mesoderm* | 148 | 349 | 452 | 651 | 516 | 132 | 23 | 2271 |
| 4 *Lateral Plate Mesoderm* | 78 | 1203 | 510 | 1759 | 563 | 880 | 652 | 5645 |
| 14 *Paraxial Mesoderm* | 44 | 333 | 487 | 835 | 4 | 5 | 11 | 1719 |
| 10 *pLateral Plate Mesoderm* | 234 | 480 | 1128 | 967 | 54 | 4 | 68 | 2935 |

D

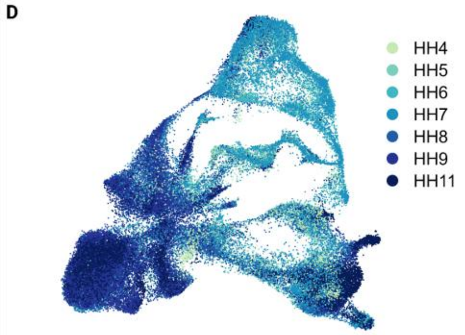


**Fig. S1. Combined integration and clustering of scRNA-sequencing datasets to identify cell types.** Unsupervised clustering of publicly available scRNA-seq data of chick embryos between HH4 and HH11 reveals 26 clusters. (A) UMAP demonstrating the unsupervised clustering results of chick whole embryos. (B) Dot plot demonstrating expression of top 5 differentially enriched genes by cluster. (C) Table of cell count by stage and cluster. (D) UMAP plot colored by stage. Definitive, Def; Non-neural ectoderm, NNE; vNeural, ventral neural; pLateral, posterior lateral; eHematogenic, early hematogenic.

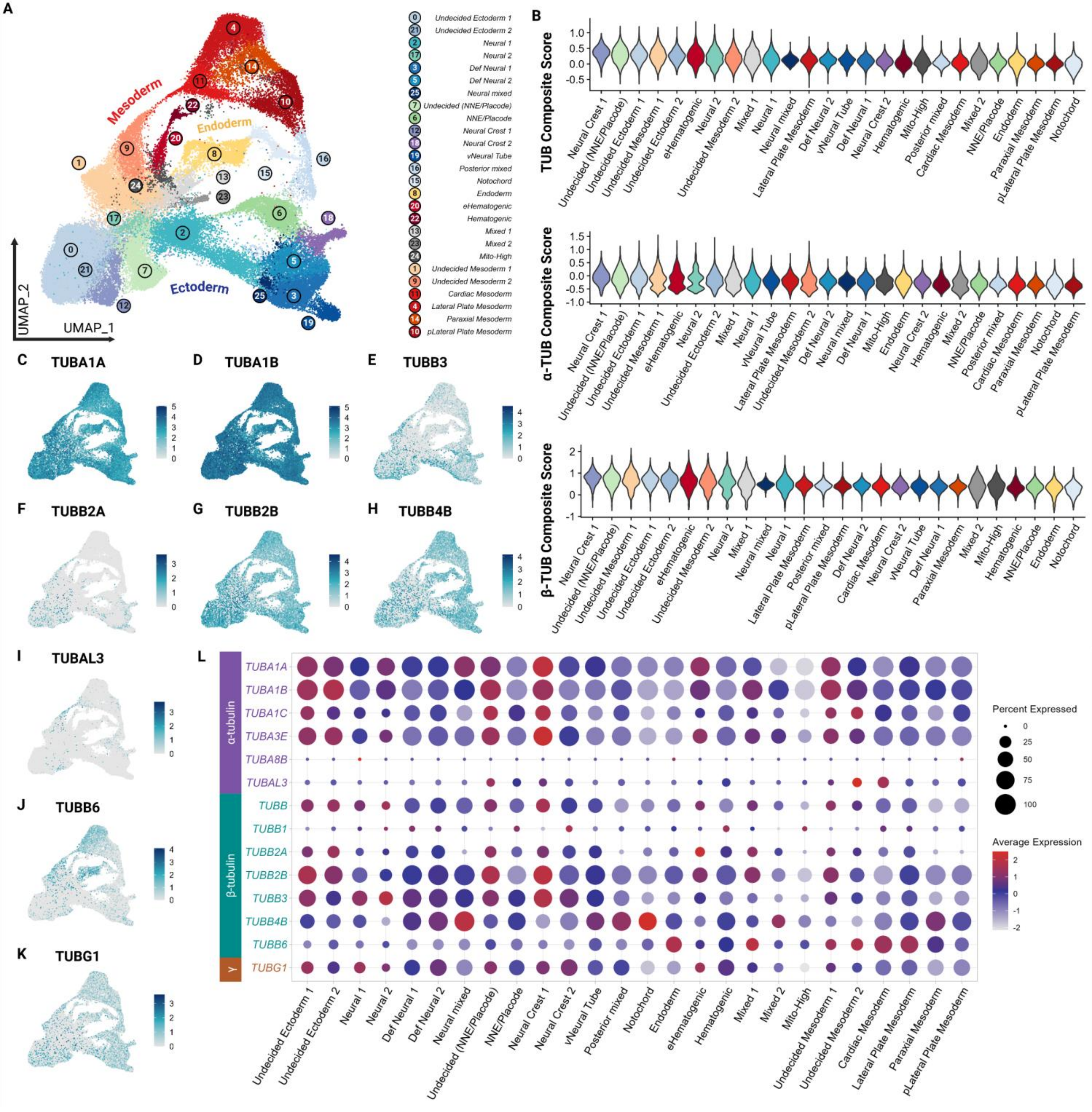


**Fig. S2. Expression of tubulin isotypes are enriched in neural crest and affiliated clusters of the developing chick.** Unsupervised clustering of publicly available scRNA-seq data of chick embryos between HH4 and HH11 reveals 26 clusters. (A) UMAP demonstrating the unsupervised clustering results of chick whole embryos. (B) Violin plots showing expression of composite tubulin scores by cluster. Tubulin scores calculated using *Seurat::AddModuleScore.* TUBB includes all tubulins in dataset, α-TUB includes all α-tubulin isotypes and β-TUBB includes all β-tubulin isotypes. Clusters are ordered in descending values of average score. (C-K) UMAP colored by select tubulin isotypes showing diversity in spatiotemporal expression patterns. (L) Dot plot showing expression of all tubulin genes across clusters. Definitive, Def; Non-neural ectoderm, NNE; vNeural, ventral neural; pLateral, posterior lateral; eHematogenic, early hematogenic.

A

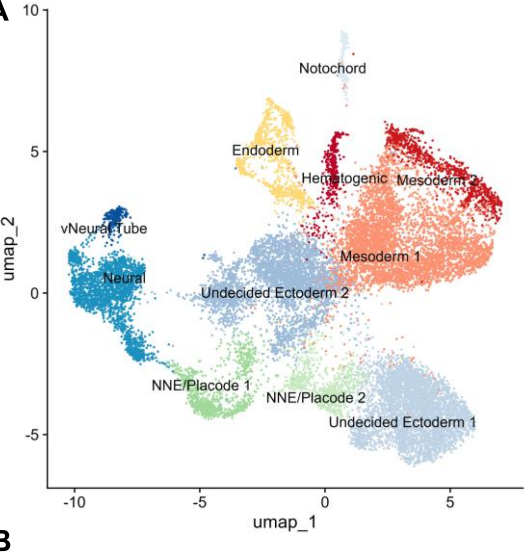


D

| | HH8 | TOTAL |
|---|---|---|
| *Mesoderm 1* | 4620 | 4620 |
| *Undecided Ectoderm 1* | 4059 | 4059 |
| *Undecided Ectoderm 2* | 3807 | 3807 |
| *Neural* | 2157 | 2157 |
| *Endoderm* | 895 | 895 |
| *NNE/Placode* | 879 | 879 |
| *Mesoderm 2* | 863 | 863 |
| *Undecided NNE/Placode* | 745 | 745 |
| *Hematogenic* | 379 | 379 |
| *Notochord* | 249 | 249 |
| *vNeural Tube* | 222 | 222 |

G

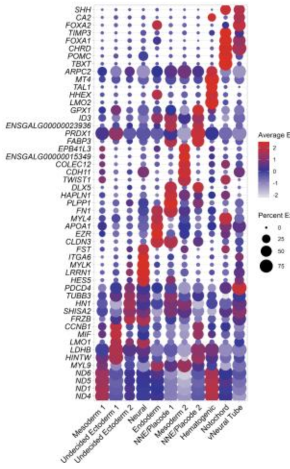

B

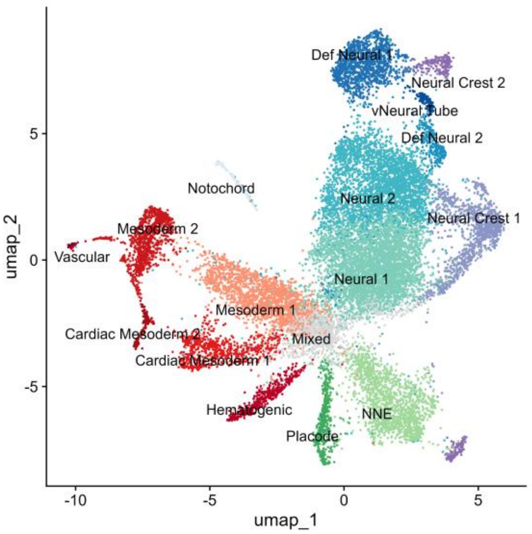


E

| | HH9 | TOTAL |
|---|---|---|
| *Neural 1* | 4422 | 4422 |
| *Neural 2* | 3234 | 3234 |
| *Mesoderm 1* | 1825 | 1825 |
| *NNE* | 1530 | 1530 |
| *Def Neural 1* | 1204 | 1204 |
| *Neural Crest 1* | 1122 | 1122 |
| *Mixed* | 1085 | 1085 |
| *Mesoderm 2* | 899 | 899 |
| *Cardiac Mesoderm 1* | 671 | 671 |
| *Placode* | 469 | 469 |
| *Hematogenic* | 466 | 466 |
| *Neural Crest 2* | 329 | 329 |
| *Def Neural 2* | 310 | 310 |
| *Notochord* | 152 | 152 |
| *vNeural Tube* | 149 | 149 |
| *Cardiac Mesoderm 2* | 130 | 130 |
| *Vascular* | 33 | 33 |

H

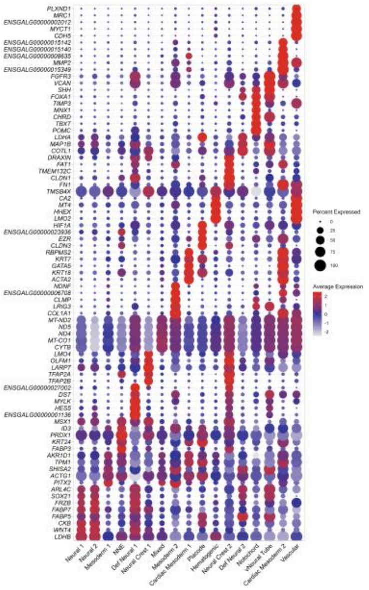

C

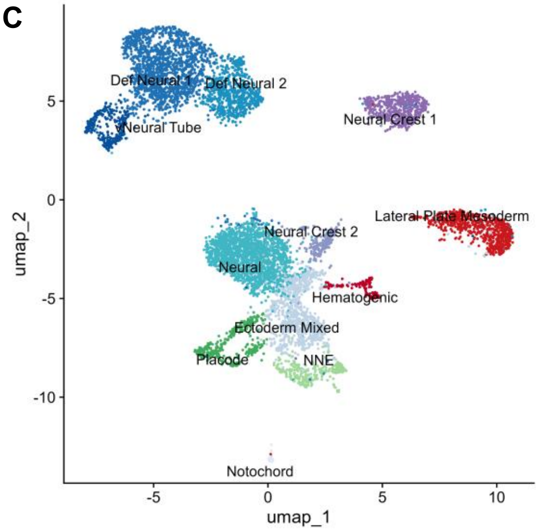


F

| | HH11 | TOTAL |
|---|---|---|
| *Neural* | 1814 | 1814 |
| *Def Neural 1* | 1588 | 1588 |
| *Ectoderm Mixed* | 878 | 878 |
| *Def Neural 2* | 706 | 706 |
| *Lateral Plate Mesoderm* | 671 | 671 |
| *Neural Crest 1* | 540 | 540 |
| *Placode* | 323 | 323 |
| *vNeural Tube* | 292 | 292 |
| *NNE* | 290 | 290 |
| *Neural Crest 2* | 206 | 206 |
| *Hematogenic* | 122 | 122 |
| *Notochord* | 40 | 40 |

I

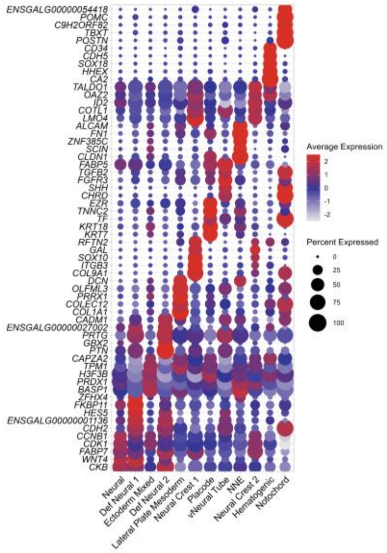

**Fig. S3. Stage specific clustering of scRNA-sequencing datasets.** (A-C) UMAP demonstrating the unsupervised clustering results. (D-F) Table of cell counts by stage and cluster. (G-I) Dot plot demonstrating expression of top 5 differentially enriched genes by cluster. Definitive, Def; Non-neural ectoderm, NNE; vNeural, ventral neural; pLateral, posterior lateral; eHematogenic, early hematogenic.

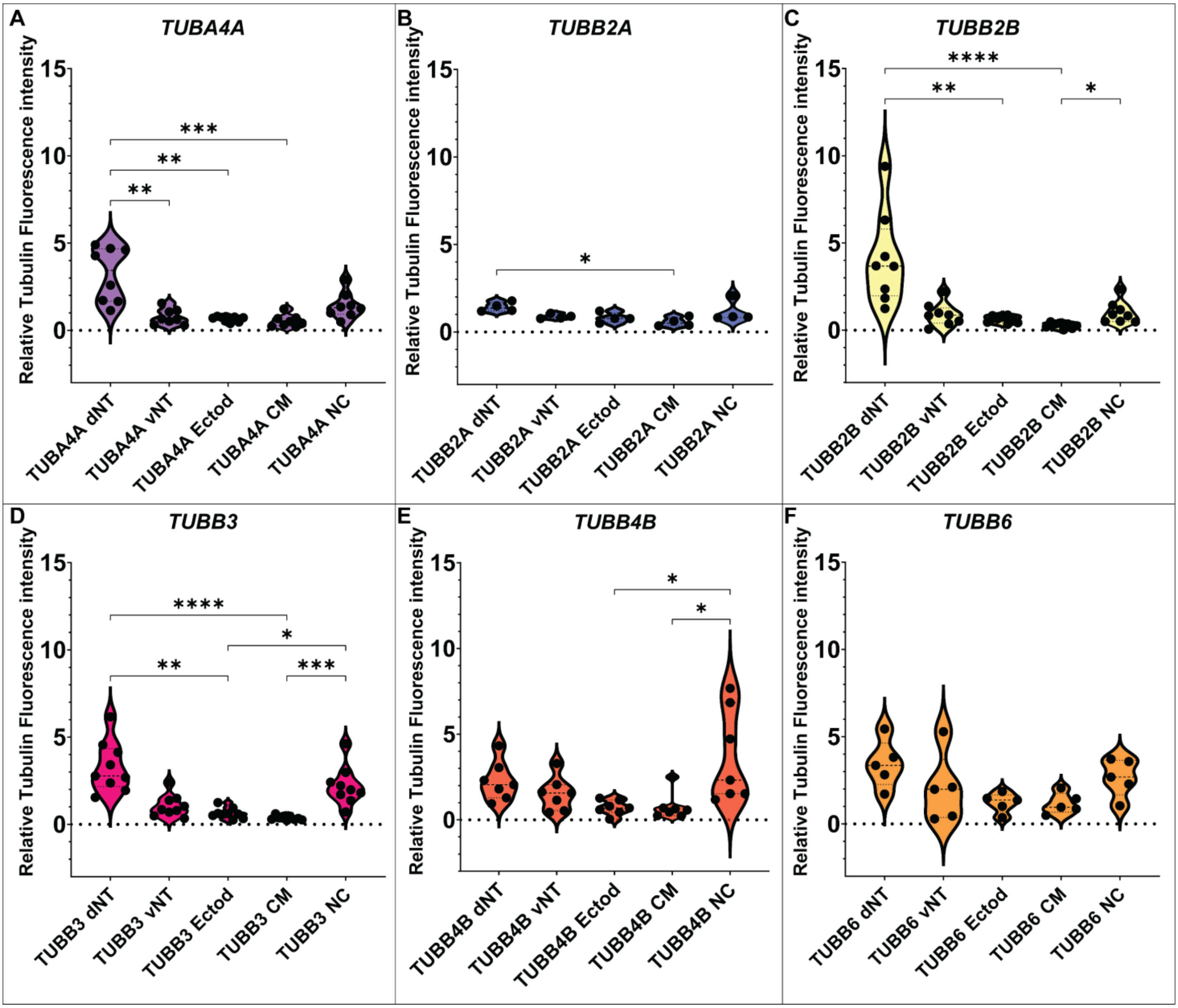


**Fig. S4. Relative fluorescence intensity of genes encoding tubulin isotypes across embryonic tissue regions.** Violin plots showing the distribution of corrected relative tubulin fluorescence intensity for six tubulin isoforms: (A) *TUBA4A* (n= 8), (B) *TUBB2A* (n= 4), (C) *TUBB2B* (n= 8), (D) *TUBB3* (n= 9), (E) *TUBB4B* (n= 7), and (F) *TUBB6* (n= 5) in transverse sections of chicken embryos. Fluorescence intensity was quantified in the dorsal neural tube (dNT), ventral neural tube (vNT), ectoderm (Ectod), cranial mesenchyme (CM), and neural crest (NC) regions using ImageJ/Fiji. Black dots represent measurements from 1-3 sections from unique individuals, while violin plots depict the distribution and density of fluorescence values within each tissue type. The dotted horizontal line indicates background-normalized fluorescence intensity at 0. Statistical significance between indicated tissue comparisons was determined using Kruskal-Wallis test (one-way ANOVA by ranks), with significance denoted as $p < 0.05$ (*), $p < 0.01$ (**), $p < 0.001$ (***), and $p < 0.0001$ (****).

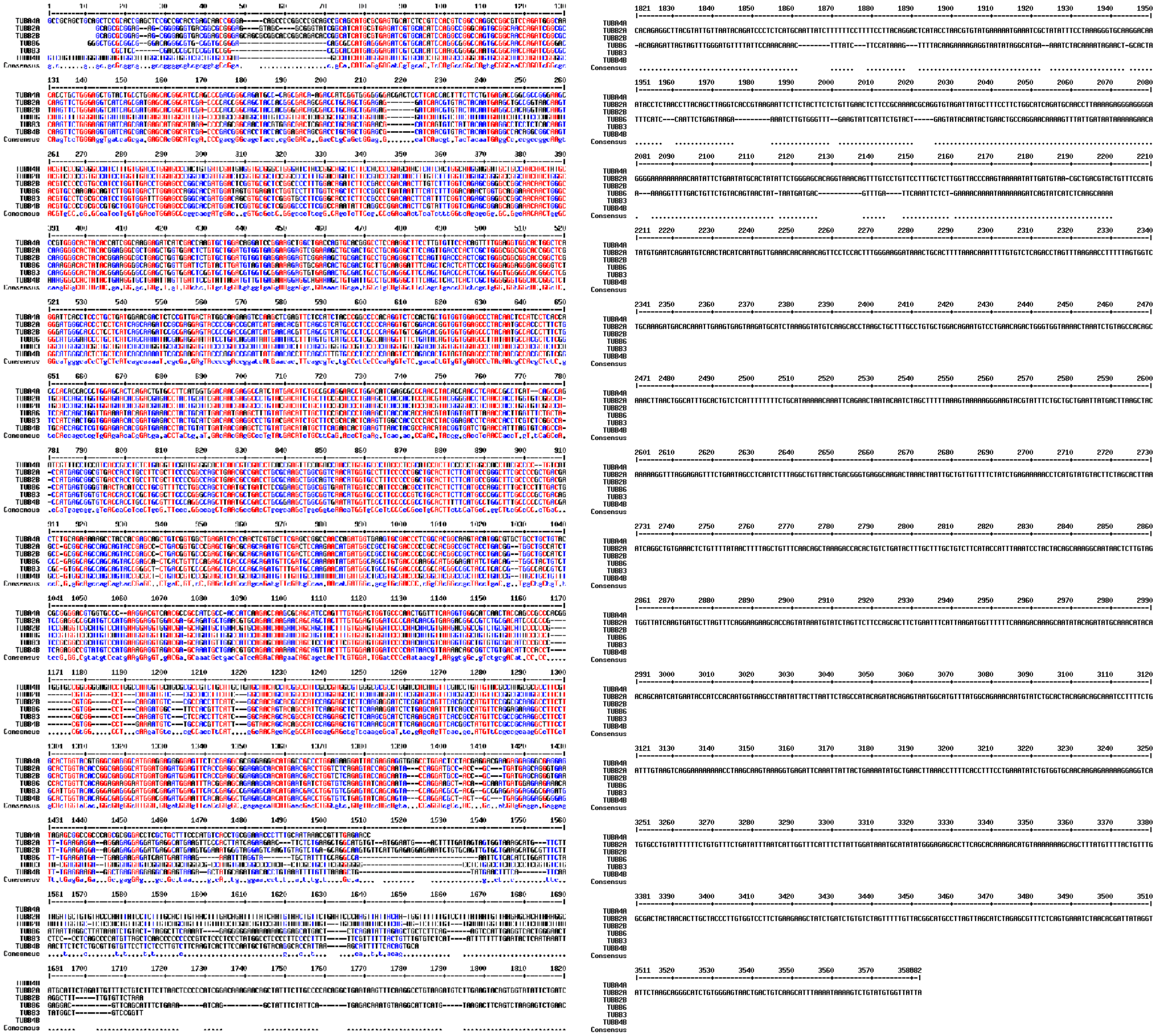

**Fig. S5. Multiple sequence alignments of tubulin mRNA transcripts analyzed during neural crest EMT.** (A) ClustalW multiple sequence alignment of *Gallus gallus TUBA4A*, *TUBB2A*, *TUBB2B*, *TUBB3*, *TUBB4B*, and *TUBB6* complete transcript sequences sourced from NCBI (Table 1). Conserved nucleotides are indicated by consensus alignment formatting (capital letters), while regions of divergence are visible throughout the aligned sequences. Alignment demonstrates broad conservation among tubulin family members with substantial divergence between the α-tubulin isotype *TUBA4A* and β-tubulin transcripts with significant sequence deviation at the 3’ end. Red= high nucleotide consensus, blue low consensus, black= neutral.

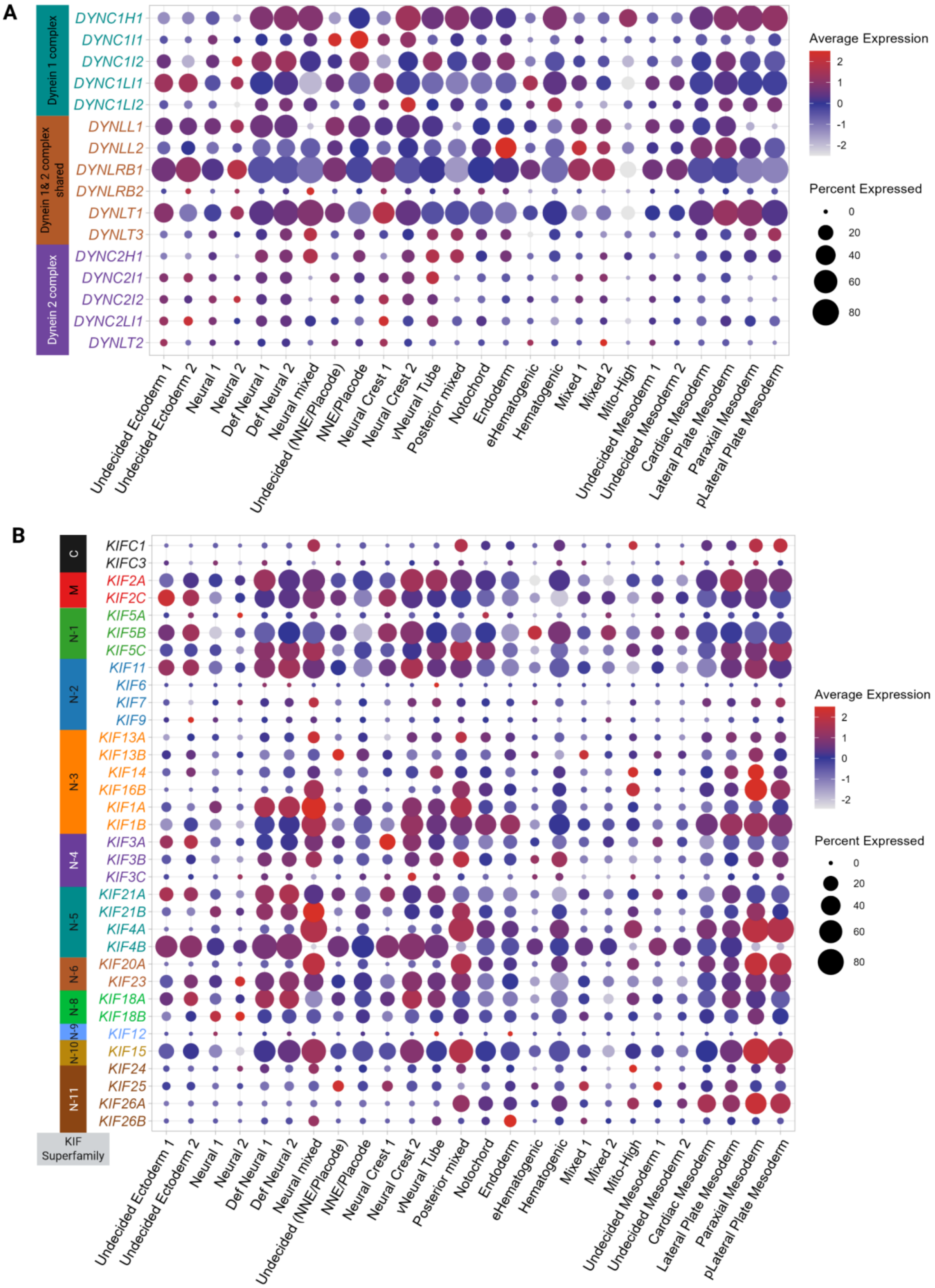


**Fig. S6. Expression map of motor proteins across cell types in the developing chick embryo.** (A) Dot plot showing expression of cytoplasmic dynein subunit genes across clusters, colored by contribution. (B) Dot plot showing expression of kinesin genes across clusters, colored by kinesin superfamilies. Definitive, Def; Non-neural ectoderm, NNE; vNeural, ventral neural; pLateral, posterior lateral; eHematogenic, early hematogenic.

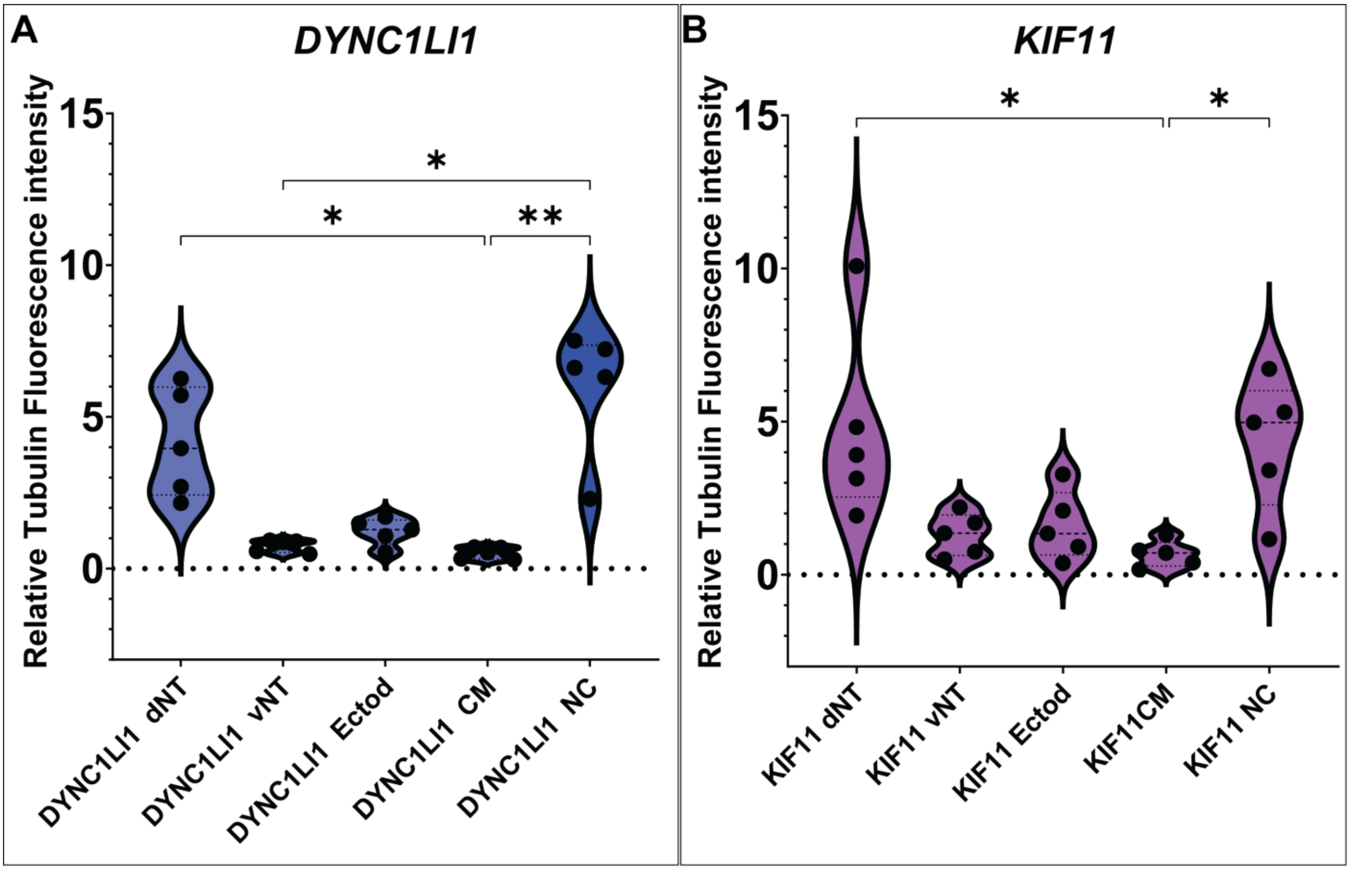


**Fig. S7. Relative fluorescence intensity of microtubule motors across embryonic tissue regions.** Violin plots showing the distribution of corrected relative tubulin fluorescence intensity for six tubulin isoforms: (A) *DYNC1LI1* (n= 5) and (B) *KIF11* (n= 5) in transverse sections of chicken embryos. Fluorescence intensity was quantified in the dorsal neural tube (dNT), ventral neural tube (vNT), ectoderm (Ectod), cranial mesenchyme (CM), and neural crest (NC) regions using ImageJ/Fiji. Black dots represent measurements from 1-3 sections from unique individuals, while violin plots depict the distribution and density of fluorescence values within each tissue type. The dotted horizontal line indicates background-normalized fluorescence intensity at 0. Statistical significance between indicated tissue comparisons was determined using [insert statistical test], with significance denoted as $p < 0.05$ (*), $p < 0.01$ (**), $p < 0.001$ (***), and $p < 0.0001$ (****).
